# Design, Synthesis and Evaluation of WD-repeat containing protein 5 (WDR5) degraders

**DOI:** 10.1101/2021.04.12.439490

**Authors:** Anja Dölle, Bikash Adhikari, Andreas Krämer, Janik Weckesser, Nicola Berner, Lena-Marie Berger, Mathias Diebold, Magdalena M. Szewczyk, Dalia Barsyte-Lovejoy, Cheryl H. Arrowsmith, Jakob Gebel, Frank Löhr, Volker Dötsch, Martin Eilers, Stephanie Heinzlmeir, Bernhard Küster, Christoph Sotriffer, Elmar Wolf, Stefan Knapp

## Abstract

Histone H3K4 methylation serves as post-translational hallmark of actively transcribed genes and is introduced by histone methyltransferases (HMT) and its regulatory scaffolding proteins. One of these is the WD-repeat containing protein 5 (WDR5) that has also been associated with controlling long non-coding RNAs and transcription factors including MYC. The wide influence of dysfunctional HMTs complexes and the typically upregulated MYC levels in diverse tumor types suggested WDR5 as an attractive drug target. Indeed, protein-protein interface inhibitors for two protein interaction interfaces on WDR5 have been developed. While such compounds only inhibit a subset of WDR5 interactions, chemically induced proteasomal degradation of WDR5 might represent an elegant way to target all oncogenic function. This study presents the design, synthesis and evaluation of two diverse WDR5 degrader series based on two WIN site binding scaffolds and shows that linker nature and length strongly influence degradation efficacy.

## Introduction

Eukaryotic transcription by all three nuclear RNA polymerases is controlled by chromatin organization. The accessibility of chromatin is influenced by the distribution of nucleosomes in the genome, the composition of histone variants within nucleosomes, and post-transcriptional modifications of histone proteins. Histone modifications present a crucial layer of epigenetic transcription control and have become a wide research field as their influence on disease development and progression is outstanding.^1^ Tight control of target gene transcription is specifically important for multicellular organisms as deregulated transcription is often associated with aberrant cellular growth and proliferation and might ultimately induce the development of cancer.^2, 3, 4, 5, 6, 7^

Among all possible histone modifications, mono-(me1), di-(me2), or tri-(me3) methylation of lysine residues in histone tails are considered as key hallmark of epigenetic regulation. One important example is the methylation of histone H3 lysine 4 residues (H3K4), which defines regulatory elements such as promoters of RNA polymerase II and enhancer elements, and therefore plays a critical role in transcriptional regulation of most protein coding genes.^8, 9, 10^ Cellular H3K4 methylation levels are determined by the balance between H3K4 demethylases and methyltransferases.^11, 12^ The highly conserved class 2 lysine methyltransferases (KTM2) comprises the mixed lineage leukaemia family (MLL1, MLL2, MLL3, MLL4, MLL5) and SET1A/SET1B enzymes, and are responsible for deposition of most of the H3K4 methylation marks associated with transcription.^13^ With the exception of MLL5, the catalytic activity of the KMT2s is dependent on the assembly of further adaptor proteins. The so called WRAD complex consists of WD repeat-containing protein 5 (WDR5), DPY30, absent, small or homeotic-2 like (ASH2L) and retinoblastoma binding protein 5 (RBBP5).^14, 15^

WDR5 is of particular importance: its propeller shaped WD interaction domain interacts with a large diversity of proteins as well as some long non-coding RNAs. Both surfaces of the doughnut shaped WD domain protein present docking sites, which are called WIN (<u>W</u>DR5-<u>in</u>teracting site) and WBM (<u>W</u>DR5- <u>b</u>inding <u>m</u>otif) sites and both protein interaction sites have been targeted successfully by small molecules.^16, 17, 18, 19, 20, 21^

Interestingly, WDR5 is not only an integral part of the WRAD complex but it also directly binds to the MYC oncoprotein family (c-, L- and N-Myc).^22, 23^ MYC proteins are essential transcription factors and their expression is frequently enhanced and deregulated in human tumours.^24^ Partial genetic inhibition of MYC is well tolerated in adult mice and it induces tumour regression and long-term survival in several tumor models such as lung adenocarcinoma,^25^ but no clinical inhibitor of MYC function is available, so far.^26^ MYC binds to the WBM site of WDR5 with an evolutionary conserved N-terminal region which is called Myc-box IIIb.^27^ Interestingly, the interaction of MYC with WDR5 supports chromatin association of MYC and is required for the MYC-mediated induction of components of the ribosome and protein production.^28^ Recently, first MYC/WDR5 protein-protein interface inhibitors have been developed.^15, 29^

The WDR5 WIN site, located opposite the MYC binding site, is required for WDR5 chromatin recruitment and interaction with KMT2 enzymes, and MLL1 is a particularly dependent on this interaction. The small molecule antagonist OICR-9429 for instance binds to the WIN site of WDR5, and efficiently disrupts its interaction with MLL1.^16, 17^ Its potential as an anti-cancer agent has been demonstrated in leukaemia and some solid tumors.^17, 30, 31^ Interestingly, targeting the WIN site with small molecules evicts WDR5 as well as its interaction partners from chromatin, resulting in changes of MLL1 dependent histone methylation.^27, 30^ Beside WDR5s’ well-established scaffolding function for KMT2 enzymes, long non-coding RNAs are also able to bind to WDR5, adding a further key role in MLL regulation of gene transcription and tumorigenesis.^32, 33^

Both main binding sites of WDR5 are important for key oncogenic functions. We hypothesized therefore, that a WDR5 degrader molecule may be an efficient therapeutic agent for cancer treatment and designed a series of PROTACs (<u>PRO</u>teolysis <u>T</u>argeting <u>C</u>himeras) based on two known WIN ligands at two diverse attachment points for linkers and E3 binding moieties. Herein, we present the development of WDR5 degraders that are based on the scaffold of the antagonist OICR-9429 as well as a modified pyrroloimidazole scaffold published by Wang and coworkers.^16, 17, 18^ The most optimal degraders of both series result in rapid, selective and robust degradation of WDR5, providing chemical tools for further studies on this interesting cancer target.

## Results and discussion

### Synthesis design of WDR5 degraders

In this study, two established WDR5 antagonists were used as basic anchoring scaffolds for PROTAC design: the (trifluoromethyl)-pyridine-2-one OICR-9429 as well as the pyrroloimidazole-based inhibitor that was published by Wang and coworkers.^16, 17, 18^ An overlay of the available crystal structures of the inhibitor complexes revealed (see Figure 1), that the two ligands enter the WDR5 binding pocket at different angles. The attachment points to the E3 ligase binding moiety were chosen at carbonyl groups that were solvent accessible and allowed the exit of linker moieties at different positions of the WIN pocket. We hypothesized that the diversity of linker attachment points might increase the possibility of efficient ternary complex formation and thus successful degradation of WDR5 by the developed degraders.

**Figure 1:**
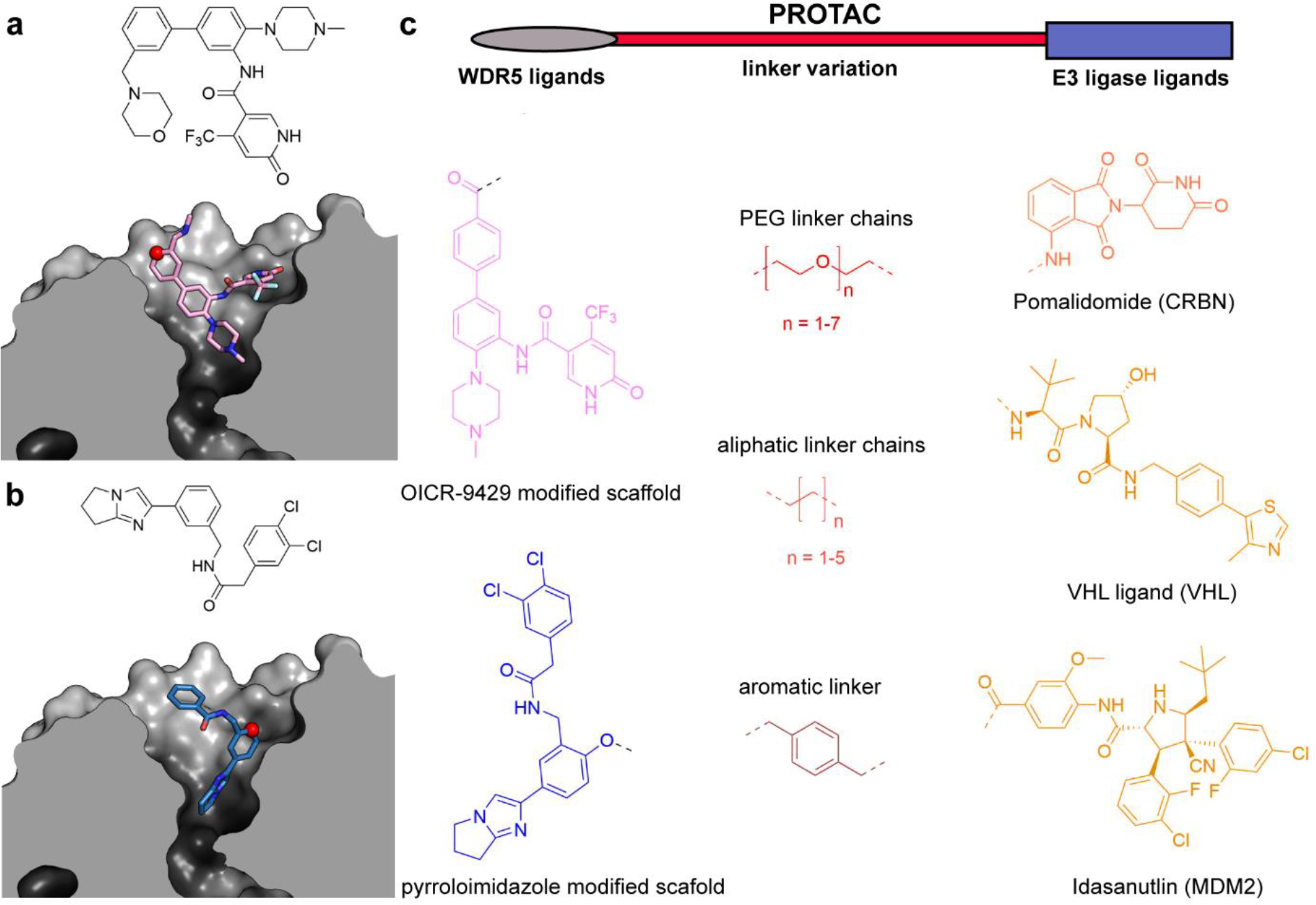
Synthesis design of WDR5 degraders based on available crystal structures. **(a)** Crystal structures of WDR5 in complex with small molecule antagonist OICR-9429 (pink; the chemical structure of OICR-9429 is displayed above) and **(b)** a small molecule published by Wang and coworkers (blue; chemical structure of the chemical probe is displayed above). The red colored spheres indicate the solvent exposed site and represent the attachment point for linkers. pdb entry: 4ql1, 6dak; **(c) (upper panel)** Schematic illustration of a heterobifunctional degrader molecule (also called PROTAC) consisting of a ligand for the target protein (WDR5, herein colored in pink/ blue) and a ligand binding an E3 ligase (orange). **(lower panel)** Chemical structures of two developed degrader series targeting WDR5. The molecules contain either the OICR-9429 derivate (pink) or the pyrroloimidazole derivate (blue) and are connected via different linkers (red) to different E3 ligase ligands (orange).

### Synthesis of OICR-9429-based degraders 7a-e, 8a-j and 9a-c

In the first series the WDR5 antagonist OICR-9429 scaffold was used.^16, 17^ Educts **1** and **2** formed intermediate **3** in a nucleophilic aromatic substitution. The reduction reaction of the aromatic nitro group lead to intermediate **4** which was coupled via a Suzuki-Miyaura cross coupling to obtain biaryl **5**. To maintain affinity for WDR5, an amide coupling of the primary amine **5** with the nicotinic acid was carried out to form amide **6**. The *in situ* deprotection of the carboxylic acid on the attached biaryl system pointed towards the solvent and served as elongation point of several E3 ligase linker **L1** – **L15** (synthesis described in **Supplementary Information**). The E3 ligase linker molecules were synthesized by either attaching the Boc-protected linker to the VHL ligand in an amide formation reaction or by attaching the Boc-protected linker to pomalidomide in a nucleophilic aromatic substitution. Deprotection of intermediate **6** and amide coupling reactions to the corresponding E3 ligase linker resulted in degrader molecules **7a-e** and **8a-j**. The synthetic steps carried out for the synthesis of the OICR-9429 based degrader molecules are shown in Scheme 1 and 2.

**Scheme 1:**
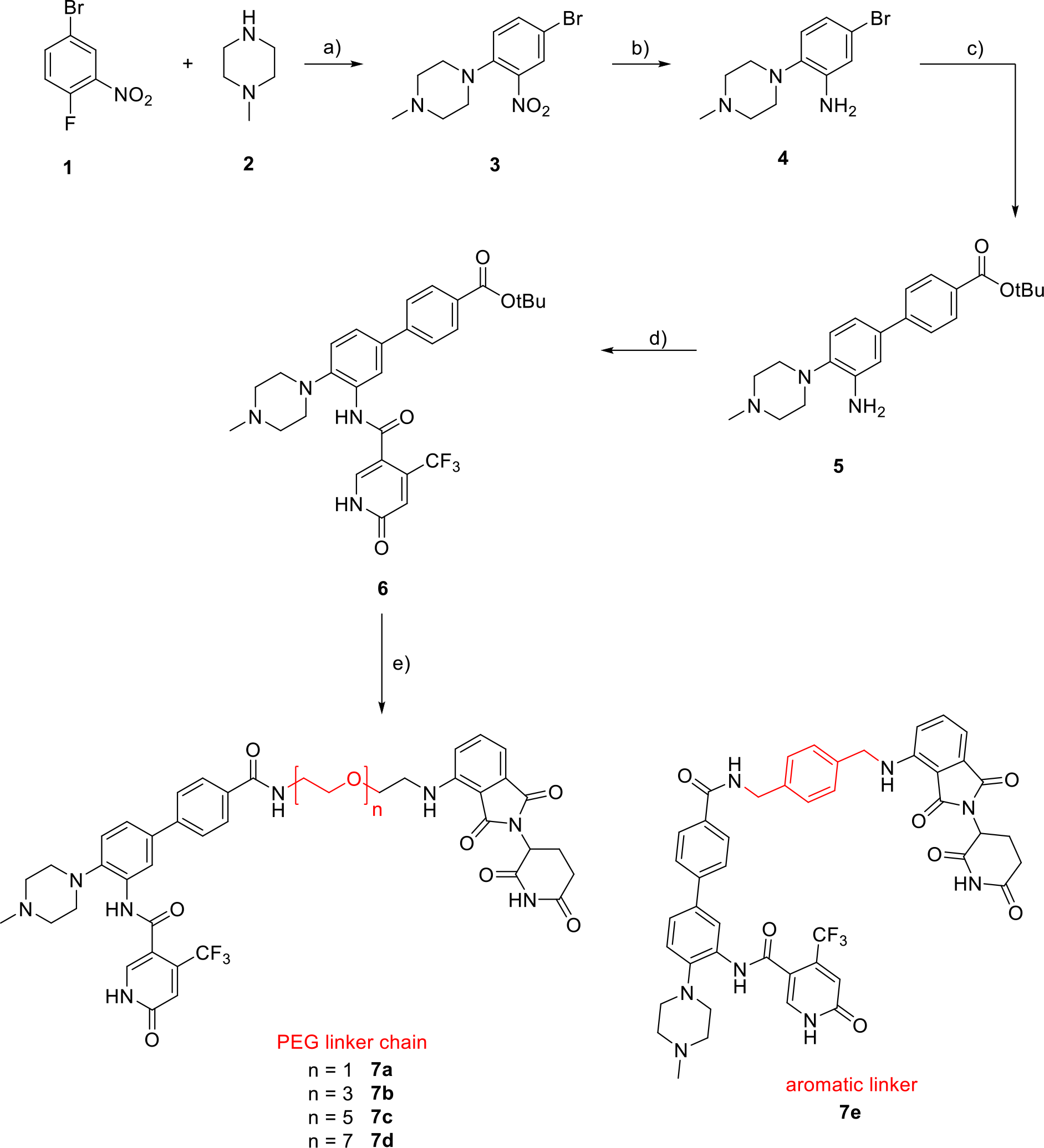
Synthesis route of WDR5 degraders **7a-e** addressing the E3 ligase Cereblon (CRBN): a) DIEA, EtOH, 80 °C, 16 h; b) Zn, NH4Cl, Dioxane/ water (3/1), rt, 30 min; c) (4-(tert-butoxycarbonyl)phenyl)boronic acid, XPhos Pd G3, NaHCO3, Dioxane/ water (3/1), 85 °C, 16 h; d) 1. Carboxylic acid, SOCl2, CH2Cl2/ ACN (1/1), 50 °C, 3 h; 2. pyridine, CH2Cl2/ ACN (1/1), 50 °C, 16 h; e) 1. TFA/ CH2Cl2 (1/1), rt, 1 h; 2. HATU, DIEA, DMF, CRBN ligase ligand linker **L1 – L5**, rt, 3-5 h. The type and nature of the linker is indicated in red.

**Scheme 2:**
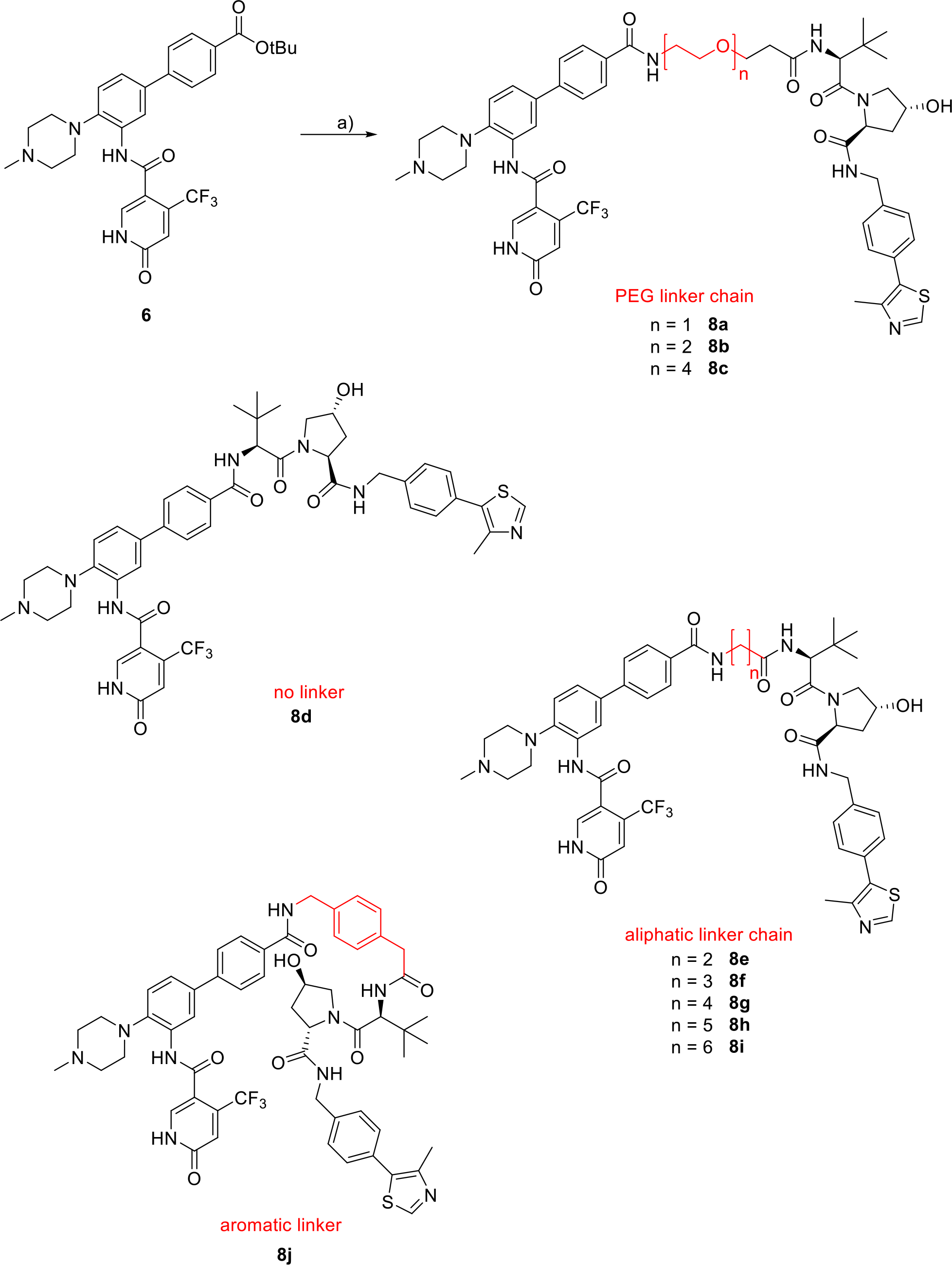
Chemical structures of WDR5 degraders **8a-j** addressing the E3 ligase Von-Hippel-Lindau (VHL). The synthesis steps starting from intermediate **6** are shown: a) 1. TFA/ CH2Cl2 (1/1), rt, 1 h; 2. HATU, DIEA, DMF, linker **L6** – **L14**, rt, 3-5 h. The type and nature of the linker is indicated in red. For the synthesis of the E3 ligase MDM2 targeting degraders, the linker was directly attached to the modified OICR-9429 scaffold **6** to form intermediates **6a-c** and this molecule was subsequently coupled using an amide formation reaction to the MDM2 inhibitor idasanutlin, yielding degrader molecules **9a-c**, as shown in Scheme 3.

**Scheme 3:**
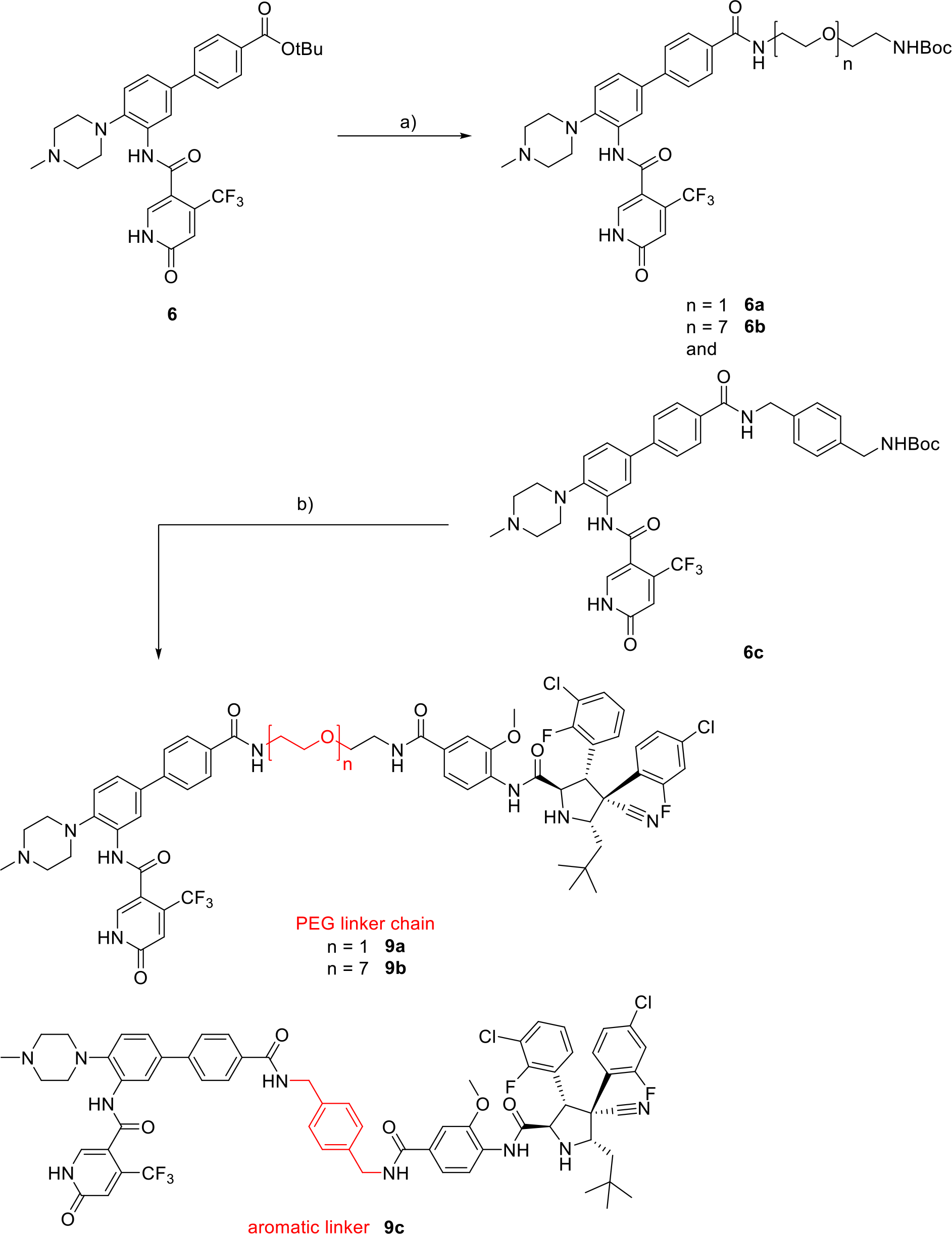
Synthesis route of WDR5 degraders **9a-c** addressing the E3 ligase MDM2. The synthesis steps starting from intermediate **6** are shown: a) 1. TFA/ CH2Cl2 (1/1), rt, 1 h; 2. HATU, DIEA, DMF, linker, rt, 3-5 h; b) 1. TFA/ CH2Cl2 (1/1), rt, 1 h; 2. HATU, DIEA, DMF, idasanutlin, rt, 12 h. The type and nature of the linker is indicated in red.

### Synthesis of pyrroloimidazole-based degraders 17a-g

The modified scaffold of the second degrader series derived from the antagonist published by Wang and coworkers.^18^ The synthetic scheme for the pyrroloimidazole based molecules are shown in Scheme 4. Educt **10** underwent an intramolecular cyclisation to form aryl bromide **11**. Intermediate **11** reacted by a Suzuki-Miyaura cross coupling with (3-cyano-4-methoxyphenyl)boronic acid to form the biaryl system **12**. Reduction of the nitrile group lead to amine **13** which was then coupled with 2-(3,4-dichlorophenyl)acetic acid to yield amide **14**. Ether deprotection of intermediate **14** lead to the free phenol **15**. Phenol **15** was reacted by a nucleophilic substitution reaction with the tosylated linker derivatives to yield intermediates **16a-g**. Deprotection of **16a-g** and followed amide bond formation with the VHL E3 ligase ligand^34^ resulting in the degrader molecules **17a-g**.

**Scheme 4:**
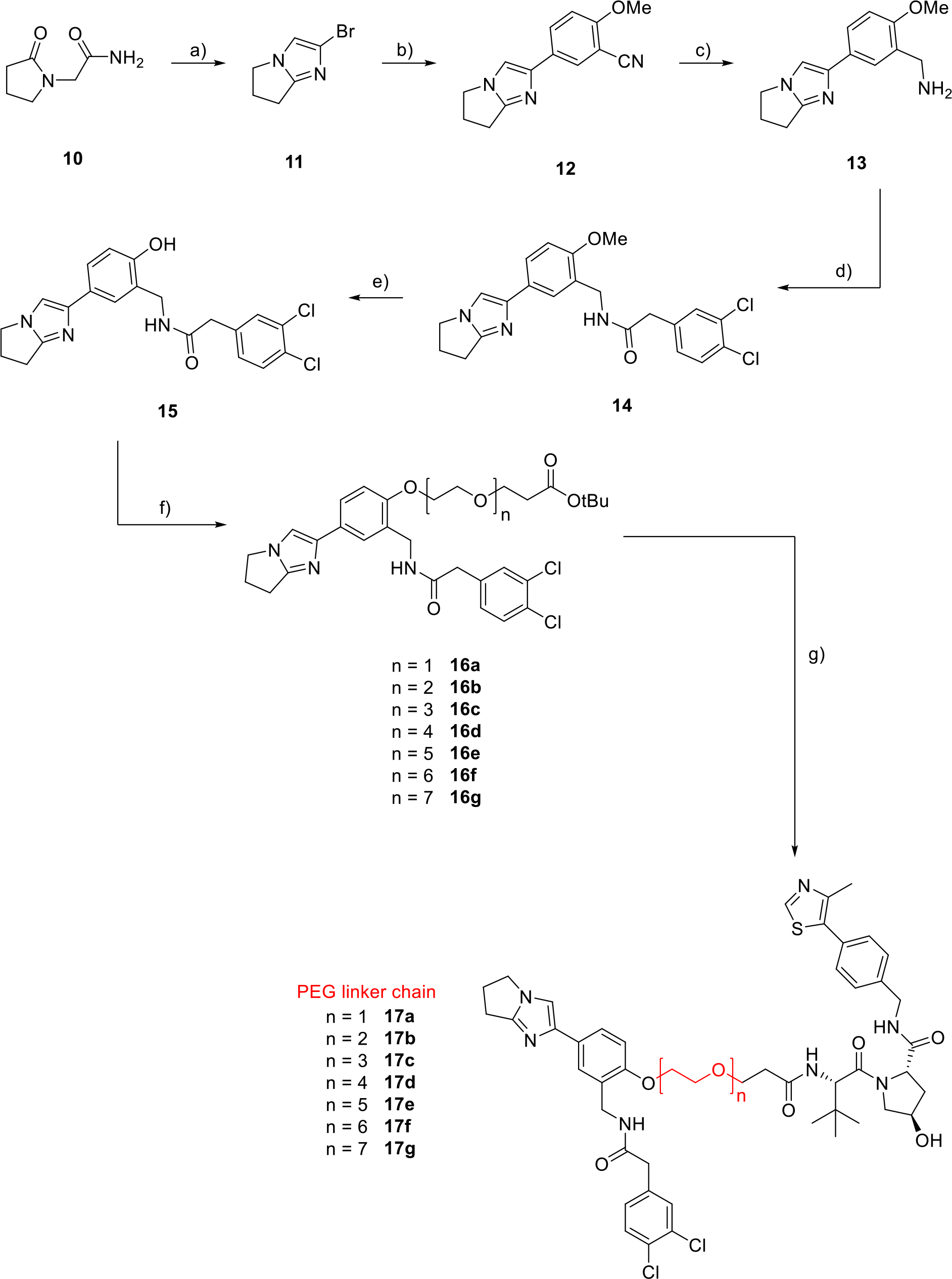
Synthesis of WDR5 degraders **17a-g** adressing the E3 ligase VHL: a) POBr3, MeCN, MW, 70 °C, 2 h; b) (3-cyano-4-methoxyphenyl)boronic acid, XPhos PdG3, NaOH, THF/H2O, 80 °C, 21 h; c) LiAlH4, THF, 60 °C - rt, 21 h; d) 2-(3,4-dichlorophenyl)acetic acid, EDC, HOBt, DIEA, DMF, rt, 16 h; e) 1. BBr3, CH2Cl2, −78 °C – rt, 21 h; 2. NaOH/H2O; f) Linker-OTs, K2CO3, DMF, 70 °C, 16-22.5 h; g) 1. TFA/ CH2Cl2 (1/1), rt, 1.5 h; 2. HATU, DIEA, DMF, VHL hydrochloride linker, rt, 3-18 h. The type and nature of the linker is indicated in red.

### Evaluation of degrader binding to WDR5

To gain a first indications of the binding affinity of the degraders to WDR5, DSF (differential scanning fluorimetry)/temperature shift measurements were carried out with recombinantly expressed WDR5, using the WDR5 antagonist OICR-9429 as a reference compound. The obtained temperature shifts *ΔT_m_* are listed in **Table 1**. Unexpectedly, the inhibitor modification **6** increased thermal stability significantly when compared to the reference OICR-9429 suggesting improved potency. Regarding the degrader molecules, temperature shift data correlated with the nature of the introduced linker: mostly aliphatic linker **8e-i** showed weaker thermal stabilization compared to more polar liker in **8a** and **8c**. However, this might be due to limiting solubility of compounds containing an aliphatic linker. The heterobifunctional molecules **7a**, **8a** and **9a** contained the shortest [PEG]-linker ([PEG]_1_) and they showed higher melting temperature shifts than degraders containing longer [PEG]-linkers, regardless of the used E3 ligase ligand. Beside the heterobifunctional molecules, the inhibitors idasanutlin, VHL ligand 1 and a modified thalidomide were included in the experiments to investigate their binding affinity towards WDR5 (see **Supplementary Information**.) All E3 ligase ligands alone used did not result in significant temperature shifts of WDR5. MDM2 ligand containing degraders **9a-c** lost their affinity towards WDR5 possibly due to their size and associated less favorable physiochemical properties or steric constrains. Performing DSF measurements of MDM2 targeting degrader intermediates **6a-c**, a decrease in temperature shifts was observed.

**Table 1.**
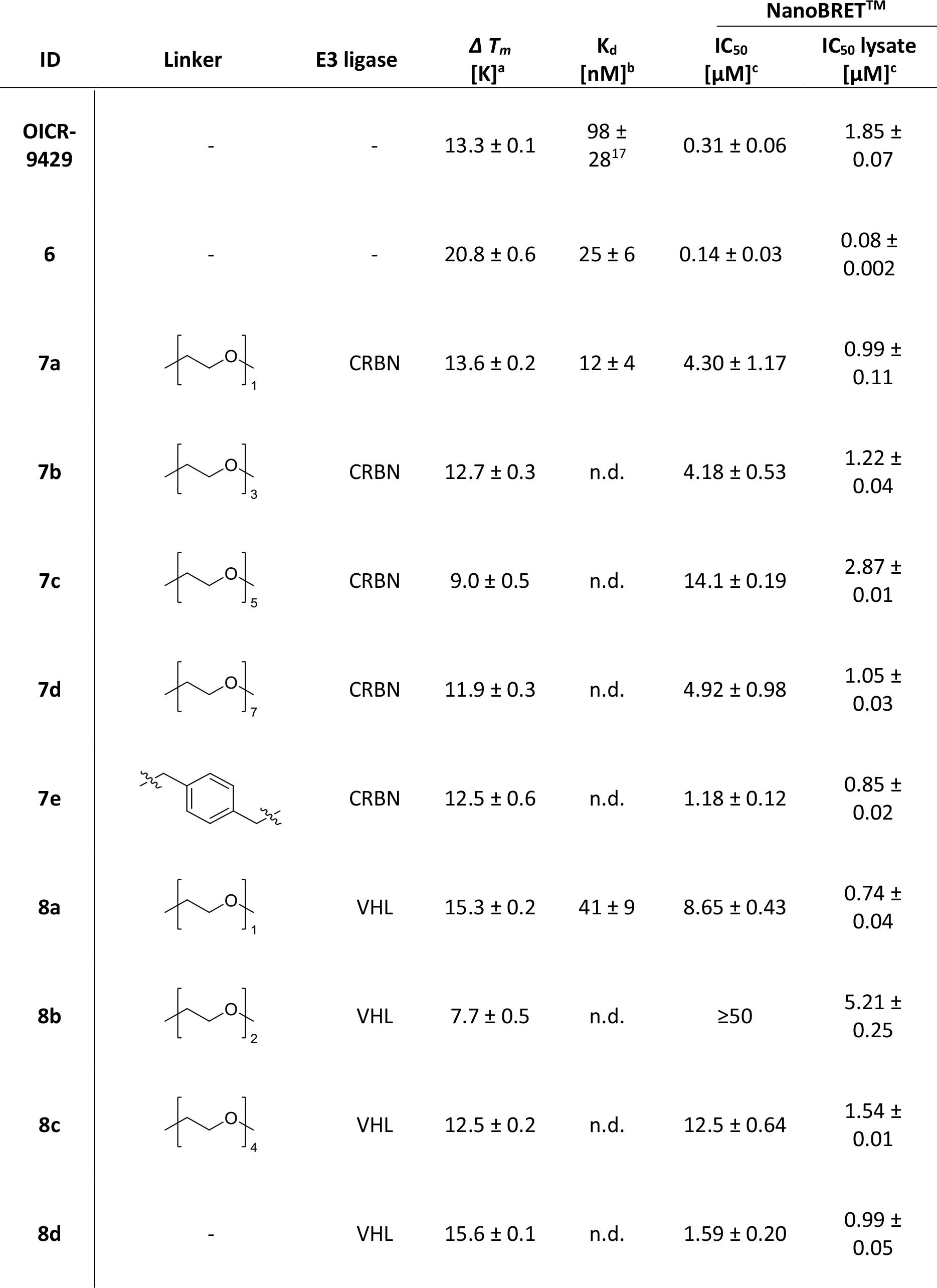
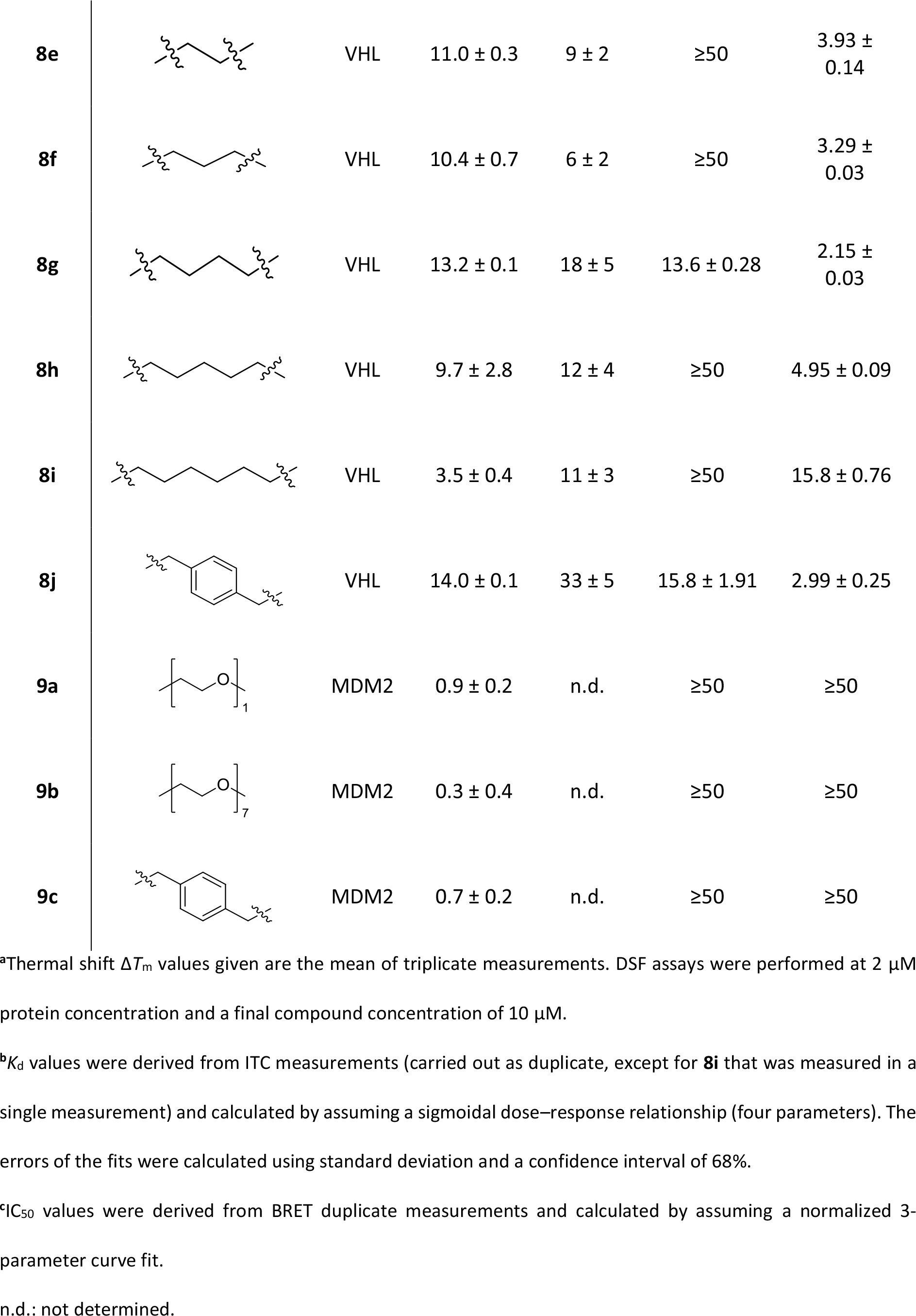
*In vitro* and *in cellulo* data of WDR5 antagonist OICR-9429, the modified inhibitor **6** and degraders **7a-e**, **8a-j** and **9a-c**.

| ID | Linker | E3 ligase | $\Delta T_m$<br>[K]<sup>a</sup> | $K_d$<br>[nM]<sup>b</sup> | NanoBRET<sup>TM</sup> | |
| --- | --- | --- | --- | --- | --- | --- |
|  |  |  |  |  | IC<sub>50</sub><br>[μM]<sup>c</sup> | IC<sub>50</sub> lysate<br>[μM]<sup>c</sup> |
| OICR-9429 | - | - | 13.3 ± 0.1 | 98 ± 28<sup>17</sup> | 0.31 ± 0.06 | 1.85 ± 0.07 |
| **6** | - | - | 20.8 ± 0.6 | 25 ± 6 | 0.14 ± 0.03 | 0.08 ± 0.002 |
| **7a** |  | CRBN | 13.6 ± 0.2 | 12 ± 4 | 4.30 ± 1.17 | 0.99 ± 0.11 |
| **7b** |  | CRBN | 12.7 ± 0.3 | n.d. | 4.18 ± 0.53 | 1.22 ± 0.04 |
| **7c** |  | CRBN | 9.0 ± 0.5 | n.d. | 14.1 ± 0.19 | 2.87 ± 0.01 |
| **7d** |  | CRBN | 11.9 ± 0.3 | n.d. | 4.92 ± 0.98 | 1.05 ± 0.03 |
| **7e** |  | CRBN | 12.5 ± 0.6 | n.d. | 1.18 ± 0.12 | 0.85 ± 0.02 |
| **8a** |  | VHL | 15.3 ± 0.2 | 41 ± 9 | 8.65 ± 0.43 | 0.74 ± 0.04 |
| **8b** |  | VHL | 7.7 ± 0.5 | n.d. | ≥50 | 5.21 ± 0.25 |
| **8c** |  | VHL | 12.5 ± 0.2 | n.d. | 12.5 ± 0.64 | 1.54 ± 0.01 |
| **8d** | - | VHL | 15.6 ± 0.1 | n.d. | 1.59 ± 0.20 | 0.99 ± 0.05 |
| **8e** | | VHL | $11.0 \pm 0.3$ | $9 \pm 2$ | $\geq 50$ | $3.93 \pm 0.14$ |
| **8f** | | VHL | $10.4 \pm 0.7$ | $6 \pm 2$ | $\geq 50$ | $3.29 \pm 0.03$ |
| **8g** | | VHL | $13.2 \pm 0.1$ | $18 \pm 5$ | $13.6 \pm 0.28$ | $2.15 \pm 0.03$ |
| **8h** | | VHL | $9.7 \pm 2.8$ | $12 \pm 4$ | $\geq 50$ | $4.95 \pm 0.09$ |
| **8i** | | VHL | $3.5 \pm 0.4$ | $11 \pm 3$ | $\geq 50$ | $15.8 \pm 0.76$ |
| **8j** | | VHL | $14.0 \pm 0.1$ | $33 \pm 5$ | $15.8 \pm 1.91$ | $2.99 \pm 0.25$ |
| **9a** | | MDM2 | $0.9 \pm 0.2$ | n.d. | $\geq 50$ | $\geq 50$ |
| **9b** | | MDM2 | $0.3 \pm 0.4$ | n.d. | $\geq 50$ | $\geq 50$ |
| **9c** | | MDM2 | $0.7 \pm 0.2$ | n.d. | $\geq 50$ | $\geq 50$ |

<sup>a</sup>Thermal shift $\Delta T_m$ values given are the mean of triplicate measurements. DSF assays were performed at 2 μM protein concentration and a final compound concentration of 10 μM.
<sup>b</sup> $K_d$ values were derived from ITC measurements (carried out as duplicate, except for **8i** that was measured in a single measurement) and calculated by assuming a sigmoidal dose–response relationship (four parameters). The errors of the fits were calculated using standard deviation and a confidence interval of 68%.
<sup>c</sup>IC<sub>50</sub> values were derived from BRET duplicate measurements and calculated by assuming a normalized 3-parameter curve fit.
n.d.: not determined.

In order to determine the binding affinity for WDR5 in solution, ITC (isothermal titration calorimetry) measurements for selected degrader molecules **7a**, **8a** and **8e-j** were performed (see Table 1; for binding curves see **Supplementary Information**.) We chose CRBN-based degrader **7a** and VHL-based degrader **8a** for ITC measurements due to their comparably small molecular weight, their identical linker structure and the high *ΔT_m_* value. Their comparison showed that the binding affinity did not correlate well with *ΔT_m_* shifts, **7a** which showed less thermal stabilizing had a three times higher affinity for WDR5 than **8a** which showed a higher *ΔT_m_* value. Also, DSF measurements revealed that **8e-i** showed a surprising diversity of thermal stabilization (ranging from 3.5 K to 13 K shift) and these ligands were therefore chosen for further characterization by ITC. Interestingly, only a small difference in binding affinity was observed for degraders **8e-I** which decrease with increasing linker length (ethyl- to hexyl-linker.) Similar to **7a** and **8a**, we concluded that high thermal stabilization did not always correlate well binding affinity for these large ligands. For instance, **8g** and **8j** displayed the highest *ΔT_m_* shifts, but showed in ITC measurements the lowest binding affinities in this series. Furthermore, a decrease in solubility was observed for degraders with long aliphatic linker (especially **8h** and **8i**) that made data acquisition using ITC challenging. The limited solubility of some degraders made also comparison of thermodynamic parameters problematic and particularly thermodynamic data measured on degraders harbouring aliphatic and aromatic linkers such as **8h**, **8i** and **8j** need to be treated with caution. Despite all these technical challenges we concluded that all heterobifunctional degrader molecules showed excellent affinity to WDR5 in the one- or two-digit nanomolar K_D_ range. For all PROTACs that we selected for ITC studies, large negative binding enthalpies were observed ranging from −10 to −4.9 kcal/mol (**Supplementary Information Table 3**). Entropy changes (TΔS) were also favourable, except form **8a** (−0.2 kcal/mol), ranging from +0.7 to +4.5 kcal/mol but enthalpy-entropy compensation was observed for degraders with large favourable ΔH values. For instance, compounds **8a**, **8f** and **8i** all showed large negative binding enthalpies of −10 kcal/mol with close to zero entropy changes (−0.2 to 1.1 kcal/mol) whereas compounds with large positive entropy changes showed small binding enthalpy (e.g. 8e: ΔH= −6.3 kcal/mol, TΔS= + 4.5 kcal/mol. These compensation mechanisms could be due to water displacement as the polar [PEG]_1_ linker in **8a** showed an unfavourable binding entropy change while degraders with aliphatic linker feature an increased binding entropy term.

Overall, the ITC experiments highlighted the value of determining binding affinity in an orthogonal assay system as similar *ΔT_m_* shifts resulted in some cases in very different binding affinities and degraders with high *ΔT_m_* shift were not necessarily more affine than a degrader with a moderate *ΔT_m_* shift.

In analogy to degrader molecules based on the modified OICR-9429 scaffold, the highest *ΔT_m_* shift of the pyrroloimidazole-based degraders **17a-g** were observed with the shortest linker moieties **17a** (see Table 2). The subsequent decrease in temperature shift values correlated also for this scaffold with the linker length. The *ΔT_m_* shift did increase slightly for degraders with longer linker moieties, namely **17e-g**, thus, solubility, solvatization in the large Win pocket and interaction with the hydrophobic surface of WDR5 might be a possible explanation for the observed effect. **17b** affinity for WDR5 was examined by ITC showing that **17b** as well as the parent compound **14** had comparable dissociation constants (K_d_s) of 97 nM and 125 nM, respectively. In accordance with the OICR-9429 derived degraders, the binding event showed to be enthalpically favored and also a gain in entropy was observed in the binding measurements. Once more, we assume this observable effect might be related to the characteristic properties of the WDR5 system that have already been discussed in context of the OICR-9429 derived degraders.

**Table 2.**
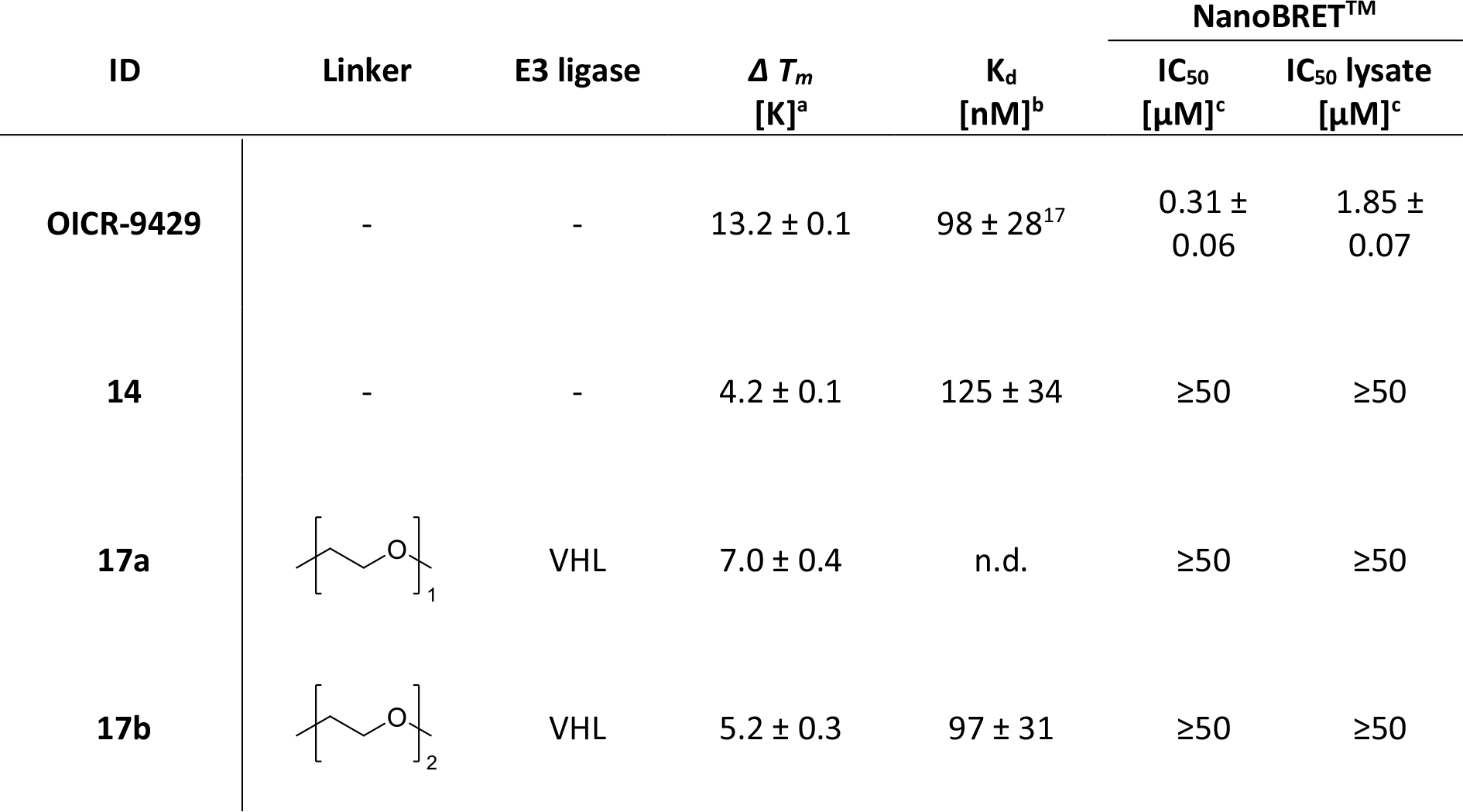
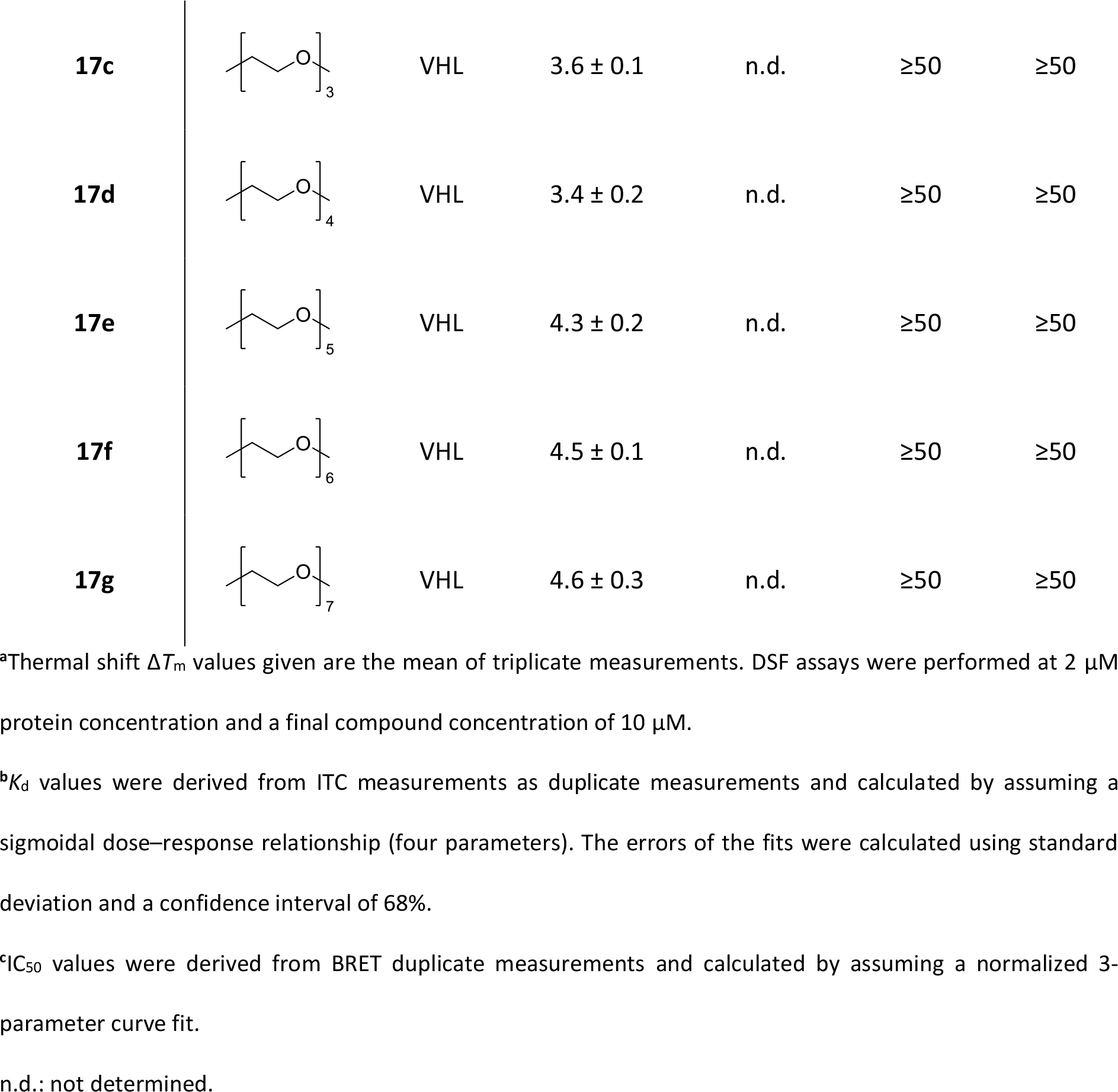
*In vitro* and *in cellulo* data of WDR5 antagonist OICR-9429, the modified molecule **14** and degraders **17a-g**.

| ID | Linker | E3 ligase | $\Delta T_m$<br>[K]<sup>a</sup> | $K_d$<br>[nM]<sup>b</sup> | NanoBRET<sup>TM</sup> | |
| --- | --- | --- | --- | --- | --- | --- |
|  |  |  |  |  | IC<sub>50</sub><br>[μM]<sup>c</sup> | IC<sub>50</sub> lysate<br>[μM]<sup>c</sup> |
| OICR-9429 | - | - | 13.2 ± 0.1 | 98 ± 28<sup>17</sup> | 0.31 ± 0.06 | 1.85 ± 0.07 |
| **14** | - | - | 4.2 ± 0.1 | 125 ± 34 | ≥50 | ≥50 |
| **17a** |  | VHL | 7.0 ± 0.4 | n.d. | ≥50 | ≥50 |
| **17b** |  | VHL | 5.2 ± 0.3 | 97 ± 31 | ≥50 | ≥50 |
| **17c** | | VHL | $3.6 \pm 0.1$ | n.d. | $\geq 50$ | $\geq 50$ |
| **17d** | | VHL | $3.4 \pm 0.2$ | n.d. | $\geq 50$ | $\geq 50$ |
| **17e** | | VHL | $4.3 \pm 0.2$ | n.d. | $\geq 50$ | $\geq 50$ |
| **17f** | | VHL | $4.5 \pm 0.1$ | n.d. | $\geq 50$ | $\geq 50$ |
| **17g** | | VHL | $4.6 \pm 0.3$ | n.d. | $\geq 50$ | $\geq 50$ |

<sup>a</sup>Thermal shift $\Delta T_m$ values given are the mean of triplicate measurements. DSF assays were performed at 2 μM protein concentration and a final compound concentration of 10 μM.
<sup>b</sup> $K_d$ values were derived from ITC measurements as duplicate measurements and calculated by assuming a sigmoidal dose–response relationship (four parameters). The errors of the fits were calculated using standard deviation and a confidence interval of 68%.
<sup>c</sup> $IC_{50}$ values were derived from BRET duplicate measurements and calculated by assuming a normalized 3-parameter curve fit.
n.d.: not determined.

### Cellular permeability and target engagement of degrader molecules

Due to the rather large molecular weight inherent of heterobifunctional molecules, cellular permeability can be limiting factor. In this study, BRET (Bioluminescence Resonance Energy Transfer) experiments were used to determine cell permeability and potency in the cellular context and for comparison also in lysed cells. To establish this assay, three tracer molecules **19a-c** based on molecule **6** were synthesized using BODIPY fluorescent conjugates. The synthetic procedures of the tracer molecules **19a-c** via its intermediates **6a**, **S18a** and **S18b** are summarized in the **Supplementary Information**. The obtained tracer molecules were titrated into cells transfected with N-terminally and C-terminally tagged WDR5 NanoLuc fusion constructs in order to determine the assay performance (see Figure 2a). These experiments revealed **19c** (see Figure 2b) and the C-terminally tagged WDR5 construct as the most suitable Tracer-NanoLuc combination for cellular BRET assays.

**Figure 2:**
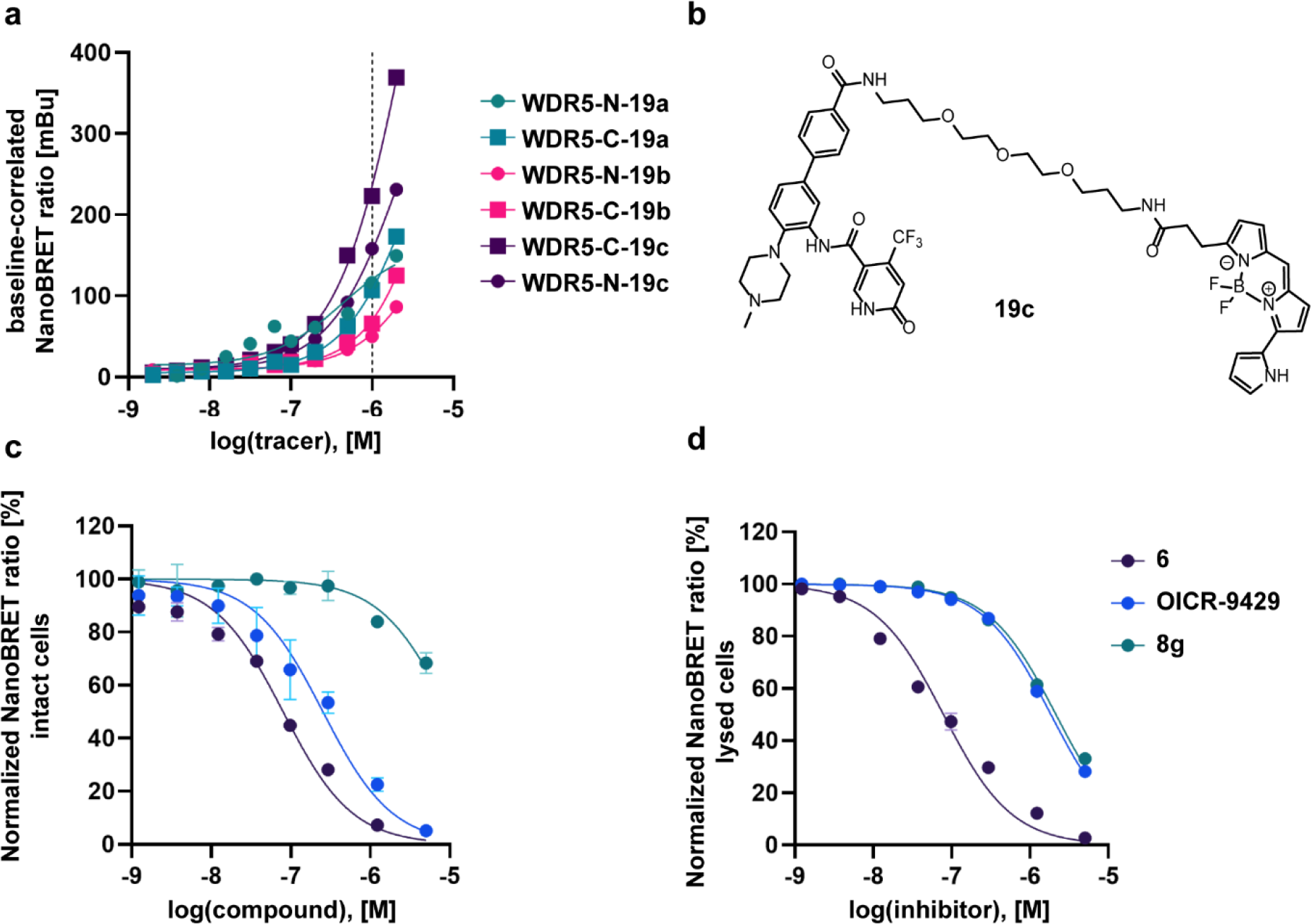
Cellular permeability and target engagement studies were performed with the BRET assay. (a) Tracer titration of all three synthesized tracer molecules and either C-terminal or N-terminal Nanoluc-tagged WDR5 (WDR5-C or WDR5-N). (b) Chemical structure of the most suitable tracer molecule 19c. (c) and (d) NanoBRETTM dose response curves of degrader 8g, ligand 6 and WDR5 antagonist OICR-9429 in live cells (c) as well as in lysed cells (d).

BRET assays were performed in lysed (permeabilized) and living cells (see Figure 2c and d) and both assay formats were used to assess cell penetration as well as cellular affinity for full-length WDR5 (see **Table 1** and **Table 2**). The parent compound of the degrader series **7a-e**, **8a-j** and **9a-c** showed a cellular potency of 139 nM. We observed a significant drop in affinity when parent compound **6** was elongated to the final degrader molecules. Comparing the attached E3 ligase ligands as well as the nature of the linker, central changes could be observed. Most Cereblon (CRBN) targeting degraders **7a-e** had similar cellular potencies. Furthermore, both formats of the assay system (intact and lysed mode) indicated a low µM affinity for WDR5. Interestingly, VHL targeting degraders showed weaker cellular activity potentially due to the peptide like nature of this ligand. While we were able to determine affinity values in lysed cells for all VHL degraders, no matter of the linker nature, the only aliphatic linker containing degrader **8g** showed a two-digit µM affinity in the intact cellular experiment. Comparing **8e-i** to the PEG containing degraders **8a-c**, the directly linked degrader **8d** and the aromatic linked molecule **8j**, weak solubility might be the limiting factor for this observation (as already shown in the ITC experiments of **8e-i**). As MDM2 targeting degraders **9a-c** already showed weak *in vitro* affinity, the BRET measurements confirmed the weak *in cellulo* activity of these molecules to WDR5. Taken together with the DSF results, the MDM2 targeting degraders were excluded from further experiments.

The generated tracer **19c** could not be displaced by the pyrroloimidazole based degraders **17a-g**. A possible hypothesis for this observation might be the different binding modes of the used inhibitor. All in all, the herein established BRET assay for WDR5 indicates that several degrader molecules are cell permeable and bind to WDR5 in cells and lysates, demonstrating their potential as putative degraders *in vivo*.

### PROTAC-mediated degradation of cellular WDR5

Targeted degradation of cellular proteins requires productive complex formation of the PROTAC-bound protein with the respective E3 ligase and proteasomal degradation of the ubiquitylated target protein. We therefore analyzed PROTAC-mediated degradation of WDR5 in cells. The open reading frame of WDR5 was fused with a luciferase peptide (called HiBiT) and stably transduced into the AML cell line MV4-11 (MV4-11^WDR5-HiBiT^). Immunoblots demonstrated that expression levels of WDR5-HiBiT are comparable to endogenously expressed WDR5 (see Figure 3a). We treated MV4-11^WDR5-HiBiT^ cells with various concentrations of the two different degrader series for 24 hours and estimated depletion of WDR5-HiBiT by measuring luciferase activity.

**Figure 3:**
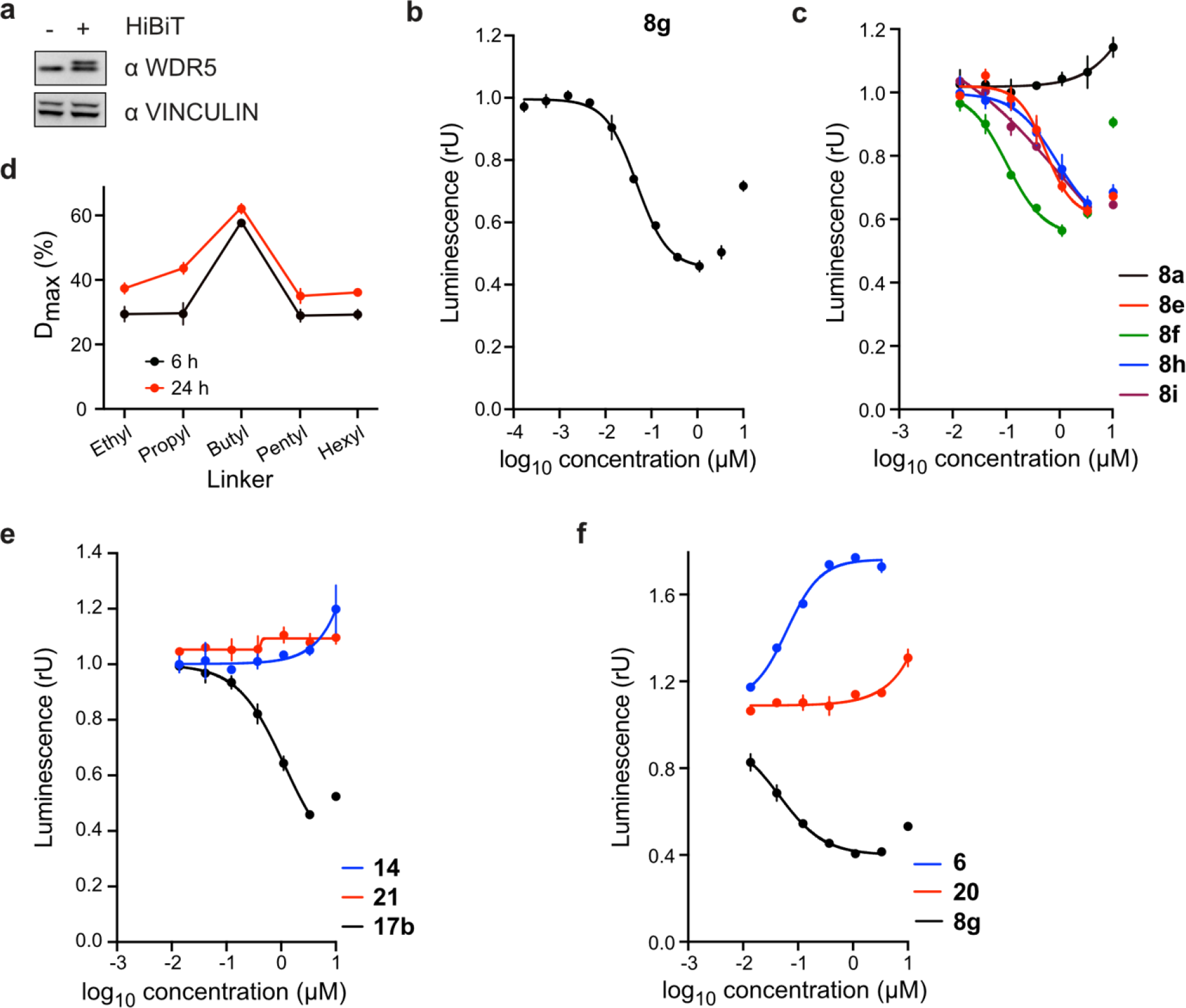
Cellular degradation studies on WDR5. (a) Immunoblot of WDR5. WDR5 was fused to fragment of luciferase (HiBiT) and stably expressed in MV4-11 cells. Naive (-) MV4-11 cells and HiBiT-WDR5 (+) MV4-11 cells. Vinculin was used as a loading control. (b) WDR5 levels based on luciferase measurements. MV4-11WDR5-HiBiT cells were treated with different concentrations of 8g for 24 h, lysed, complemented with the second luciferase fragment (largeBiT) and measured for luciferase activity. (c) WDR5 levels based on luciferase measurements. MV4-11WDR5-HiBiT cells were treated with different concentrations of degraders 8a, 8e, 8f, 8h and 8i for 24 h, lysed, complemented with the second luciferase fragment (largeBiT) and measured for luciferase activity. (d) Quantification of WDR5 Dmax (maximal degradation) from HiBiT assay for degraders with different aliphatic linkers. MV4-11WDR5-HiBiT cells were treated with different concentrations of degraders for 6h or 24 h, lysed, complemented with the second luciferase fragment (largeBiT) and measured for luciferase activity of linkers comprising ethyl: 8e, propyl: 8f, butyl: 8g, pentyl: 8h and hexyl: 8i. (e) and (f) WDR5 levels based on luciferase measurements. MV4-11WDR5-HiBiT cells were treated with different concentrations of 14, 21 and 17b (e), 6, 20, and 8g (f) for 24 h. Cells were then lysed, complemented with the second luciferase fragment (largeBiT) and measured for luciferase activity.

Intriguingly, depletion of WDR5-HiBiT varied substantially between the different degrader classes (see **Table 3** and **Table 4**, additionally data can be found in the **Supplementary Information** Figure 3). While none of the Cereblon based PROTACs showed significant depletion, many VHL ligand-containing degraders demonstrate *cellular* degrader efficacy. The most effective degrader of the OICR-9429 derived series **8g** that linked both functional binding moieties by a butyl chain, showed a maximum depletion of 58 ± 3 % and DC_50_-values of 53 ± 10 nM (see Figure 3b and Table 3). A shortening of the linker by implementing ethyl (**8e**) and propyl (**8f**) chains as well as an elongation with pentyl (**8h**) and hexyl (**8i**) chains significantly reduced degradation efficiency (see Figure 3c and 3d). Strikingly, degrader **8a**, a degrader resembling **8h**, but containing a [PEG]_1_ moiety instead of an aliphatic chain, did not induce degradation of WDR5 (see Figure 3c). The most-efficient pyrroloimidazole-based degrader was **17b** that contained a [PEG]_2_ linker (see Figure 3e). Both, **8g** and **17b** did not induce maximal depletion of WDR5 at high concentrations, a phenomenon called the Hook-effect, resulting from less efficient ternary complex formation at excess degrader levels due to binding site competition (see Figure 3b **and 3e**).^35^ Two negative controls, **20** resembling molecule **8g**, and **21** resembling molecule **17b**, with the inactive variant of the VHL ligand were synthesised to verify the effect of targeted protein degradation. Their biophysical properties can be found in the **Supplementary Information**. Notably, neither negative control **20** and **21**, nor the ligands **6** and **14** alone, induced degradation of WDR5 in cells (see Figure 3e and 3f). The increase on WDR5 levels by **6** is most likely a direct effect on WDR5 stability induced by ligand binding.

**Table 3.**
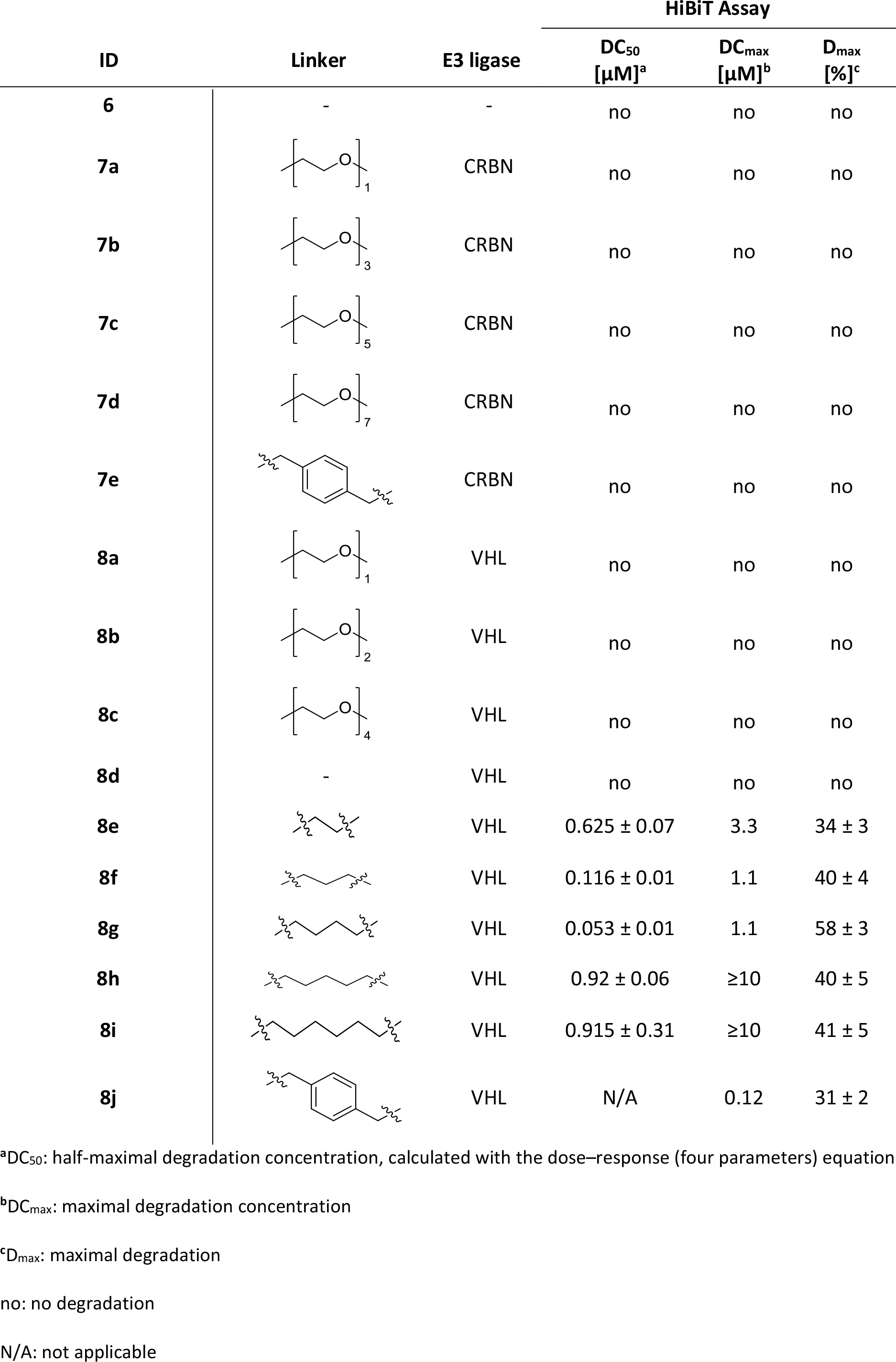
HiBiT data of WDR5 ligand **6** and degraders **7a-e** and **8a-j**.

| ID | Linker | E3 ligase | HiBiT Assay |  |  |
| --- | --- | --- | --- | --- | --- |
|  |  |  | DC<sub>50</sub><br>[μM]<sup>a</sup> | DC<sub>max</sub><br>[μM]<sup>b</sup> | D<sub>max</sub><br>[%]<sup>c</sup> |
| **6** | - | - | no | no | no |
| **7a** |  | CRBN | no | no | no |
| **7b** |  | CRBN | no | no | no |
| **7c** |  | CRBN | no | no | no |
| **7d** |  | CRBN | no | no | no |
| **7e** |  | CRBN | no | no | no |
| **8a** |  | VHL | no | no | no |
| **8b** |  | VHL | no | no | no |
| **8c** |  | VHL | no | no | no |
| **8d** | - | VHL | no | no | no |
| **8e** |  | VHL | 0.625 ± 0.07 | 3.3 | 34 ± 3 |
| **8f** |  | VHL | 0.116 ± 0.01 | 1.1 | 40 ± 4 |
| **8g** |  | VHL | 0.053 ± 0.01 | 1.1 | 58 ± 3 |
| **8h** |  | VHL | 0.92 ± 0.06 | ≥10 | 40 ± 5 |
| **8i** |  | VHL | 0.915 ± 0.31 | ≥10 | 41 ± 5 |
| **8j** |  | VHL | N/A | 0.12 | 31 ± 2 |

<sup>a</sup>DC<sub>50</sub>: half-maximal degradation concentration, calculated with the dose–response (four parameters) equation
<sup>b</sup>DC<sub>max</sub>: maximal degradation concentration
<sup>c</sup>D<sub>max</sub>: maximal degradation
no: no degradation
N/A: not applicable

**Table 4.**
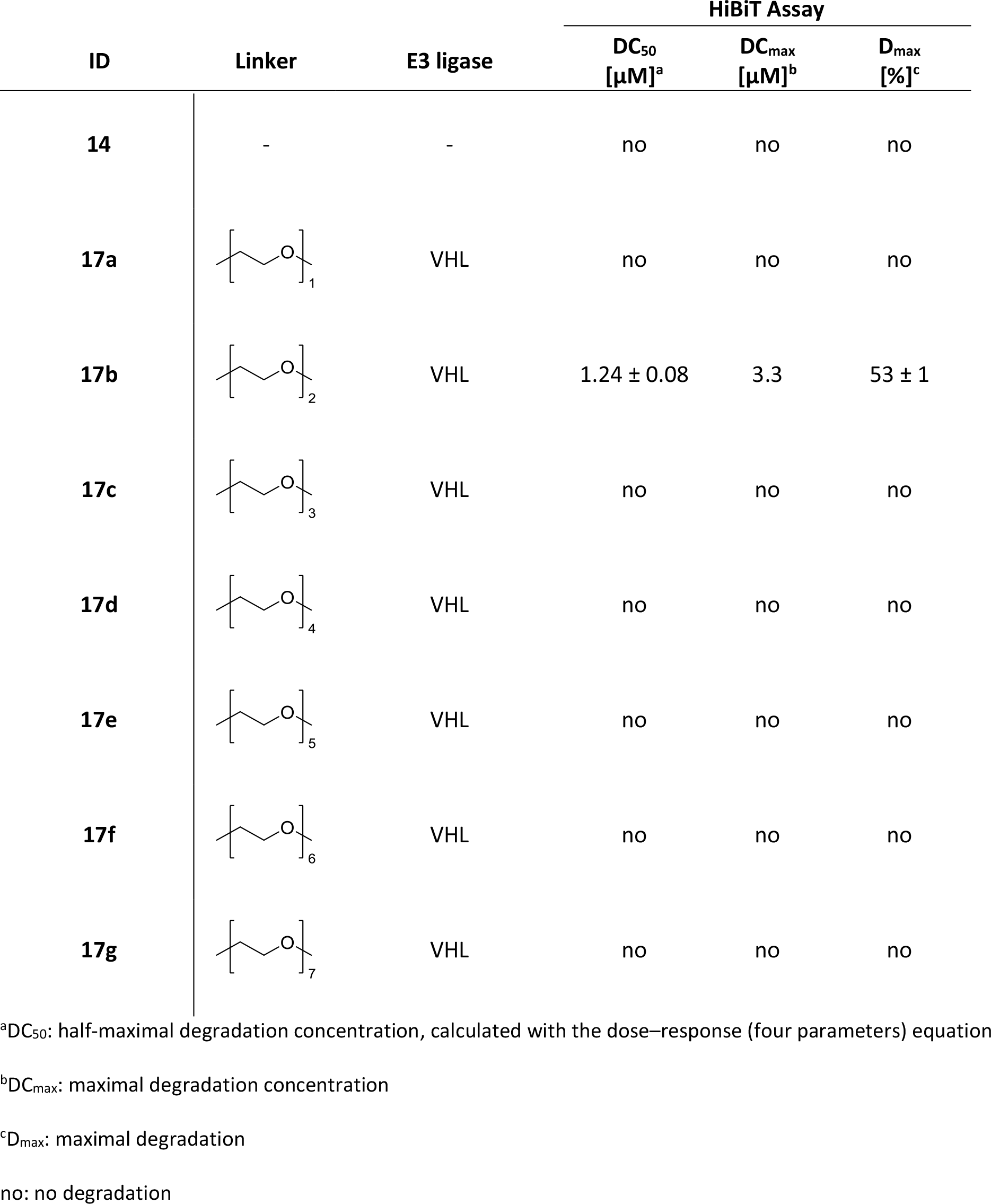
HiBiT data of WDR5 ligand **14** and degraders **17a-g**.

| ID | Linker | E3 ligase | HiBiT Assay |  |  |
| --- | --- | --- | --- | --- | --- |
|  |  |  | DC<sub>50</sub><br>[μM]<sup>a</sup> | DC<sub>max</sub><br>[μM]<sup>b</sup> | D<sub>max</sub><br>[%]<sup>c</sup> |
| **14** | - | - | no | no | no |
| **17a** |  | VHL | no | no | no |
| **17b** |  | VHL | 1.24 ± 0.08 | 3.3 | 53 ± 1 |
| **17c** |  | VHL | no | no | no |
| **17d** |  | VHL | no | no | no |
| **17e** |  | VHL | no | no | no |
| **17f** |  | VHL | no | no | no |
| **17g** |  | VHL | no | no | no |

<sup>a</sup>DC<sub>50</sub>: half-maximal degradation concentration, calculated with the dose–response (four parameters) equation<sup>b</sup>DC<sub>max</sub>: maximal degradation concentration<sup>c</sup>D<sub>max</sub>: maximal degradation
no: no degradation

Next, we analyzed if the degrader molecules **8g** and **17b** also induced degradation of untagged and endogenously expressed WDR5. MV4-11^WDR5-HiBiT^ cells were treated for 24 hours with **8g** or **17b** and were analyzed by immunoblots stained with an anti-WDR5 antibody. Both degraders induced efficient depletion of endogenous WDR5, and the dose-dependency and gratifyingly degradation efficacy of the endogenous protein resembled the depletion of the HiBiT-tagged protein (see Figure 4a and **Supplementary Information** Figure 4). Degraders **8g** and **17b** showed similar level of WDR5 depletion in naive MV4-11 cells even 72 hours post treatment (see Figure 4b and **Supplementary Information** Figure 5a). Immunoblotting also confirmed the depletion efficiency of various other degraders of WDR5 in naive MV4-11 cells as seen in HiBiT data (see **Supplementary Information Figure 5b-5f**). Degraders induce degradation of their targets by inducing ubiquitylation and subsequently proteasomal degradation. We therefore tested, if **8g** and **17b** decreased protein stability of WDR5. We blocked protein translation by incubating cells with cycloheximide in addition to the degraders (or vehicle treated cells), and estimated WDR5 levels by immunoblotting at several time points. Both degraders reduced the stability of WDR5 protein substantially in comparison to the vehicle treated cells (Figure 4c and **Supplementary Information** Figure 5g).

**Figure 4:**
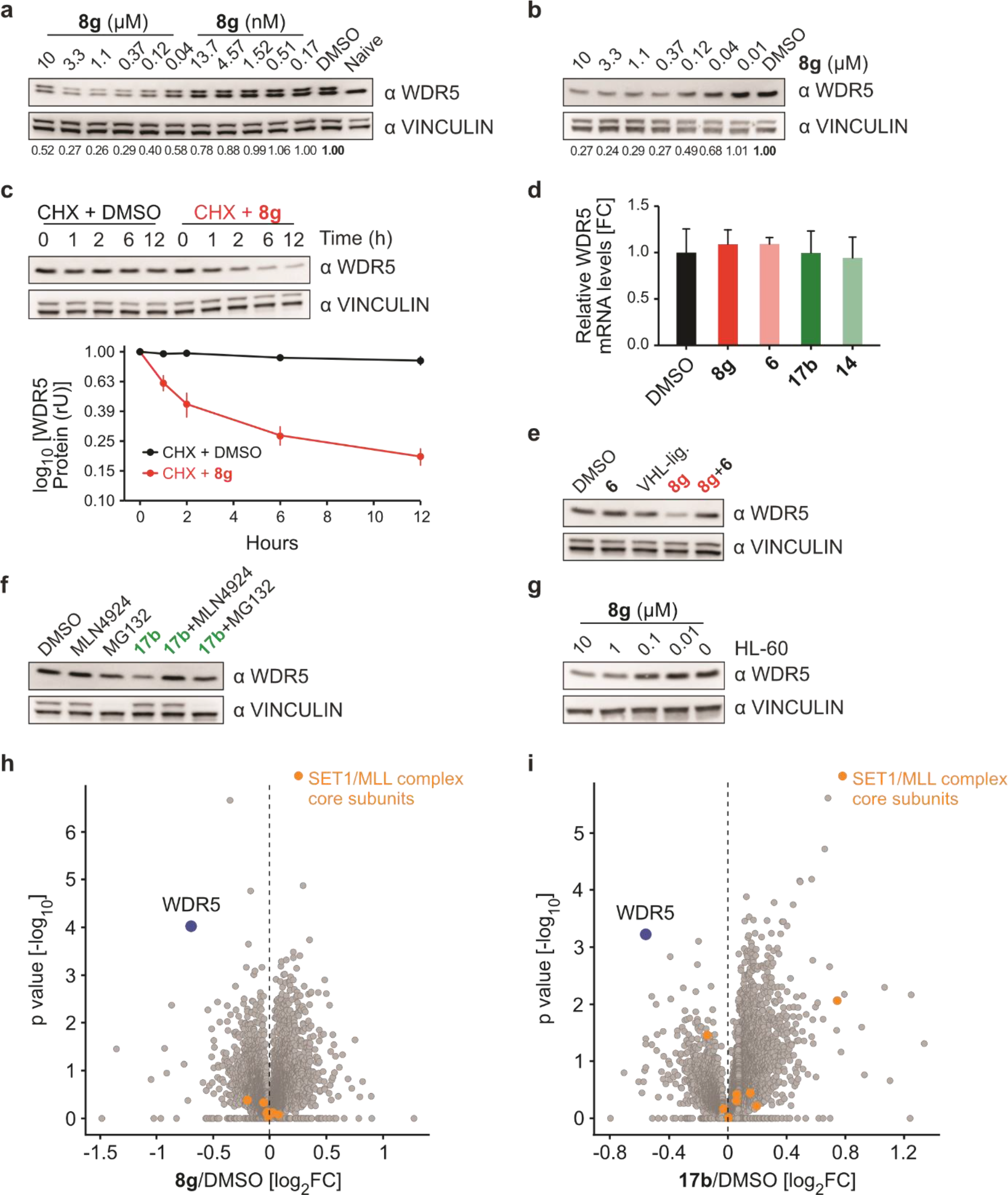
Degrader-induced depletion of WDR5 depends on the ubiquitin system. (a) Immunoblot of WDR5. MV4-11WDR5-HiBiT cells were treated with different concentration of 8g for 24 h and compared with DMSO treated or naive MV4-11 cells. Vinculin was used as loading control (as in all further immunoblot experiments). Quantification is based on both protein bands (endogenous /HiBiT tagged WDR5). (b) Immunoblot of WDR5. Naive MV4-11 cells were treated with different concentration of 8g for 72 h and compared with DMSO treated cells. (c) Immunoblot and quantification of WDR5 levels. WDR5 protein stability was evaluated by treating 1 µM 8g or DMSO incubated MV4-11 cells for 0, 1, 2, 6 and 12 h with cycloheximide (CHX). The data is mean ± s.d from n=2 biological replicates. (d) Quantitative RT-PCR analysis of WDR5 mRNA levels. RNA was extracted from MV4-11 cells incubated with 1 µM 8g, 1 µM 6, 3 µM 17b and 3 µM 14 for 24h. WDR5 expression levels were normalized with a reference gene (B2M). Bars represents mean ± s.d. of n= 3 technical replicates. (e) Immunoblot of WDR5. MV4-11 cells were treated for 6 h with 1 µM 8g or 5 µM 6 or 10 µM VHL-ligand or combination of them. (f) Immunoblot of WDR5. MV4-11 cells were treated for 6h with 3 µM 17b or 5 µM MLN4924 or 10 µM MG132 or combination of them. (g) Immunoblot of WDR5. HL-60 cells were treated with different concentrations of 8g for 24 h. (h) and (i) Volcano plot exhibiting global proteomics change. MV4-11 cells were treated with 1 µM 8g (h) or 5 µM 17b (i) for 9 hours and lysates were analyzed by quantitative proteomics. WDR5 (blue) and other SET1/MLL complex core subunits: KMT2A, KTM2B, KTM2C, KTM2D, SETD1A, RBBP5, ASH2L and DPY30 (orange).

**Figure 5:**
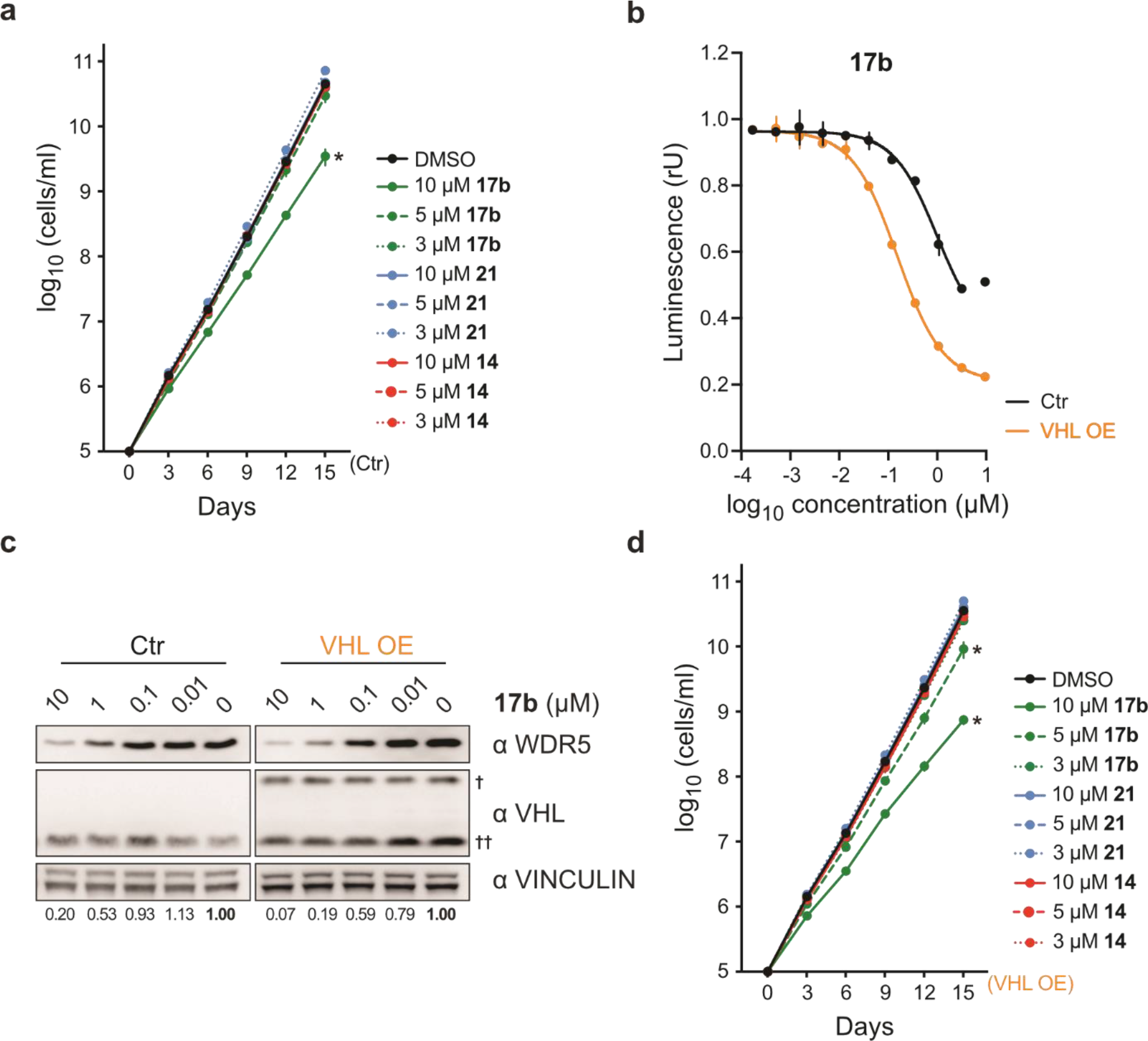
VHL overexpression increases PROTACs/degrader efficiency and cellular responses. (a) Cumulative growth curve in MV4-11 cells. MV4-11 cells were treated with 10 µM, 5 µM and 3 µM of 17b, 21 and 14 and counted at indicated time point. To prevent the overgrowth, cells were reseeded to original density every third day in fresh media with compounds and treatment continued for 15 days. Data represent mean ± s.d. of n=2 biological replicates. Asterisks indicate P-value calculated from 15th day cumulative cell number (two-tailed unpaired t-test assuming equal variance against DMSO treatment). * P ≤ 0.05 (b) WDR5 levels based on luciferase measurements. MV4-11WDR5-HiBiT cells (Ctr) and MV4-11WDR5-HiBiT/VHL cells (VHL OE) were treated with different concentrations of 17b for 24 h, lysed, complemented with the second luciferase fragment (largeBiT) and measured for luciferase activity. (c) Immunoblot of WDR5 and VHL. MV4-11 cells (Ctr) and MV4-11VHL (VHL OE) were treated for 24h with various concentrations of 17b. † overexpressed VHL; †† endogenous VHL (d) Cumulative growth curve in MV4-11VHL cells. MV4-11VHL cells were treated with 10 µM, 5 µM and 3 µM of 17b, 21 and 14 and counted at indicated time points. To prevent the overgrowth, cells were reseeded to original density every third day in fresh media with compounds and treatment continued for 15 days. Data represent mean ± s.d. of n=2 biological replicates. Asterisks indicate P-values calculated from 15th day cumulative cell number (two-tailed unpaired t-test assuming equal variance against DMSO treatment). * P ≤ 0.05

To exclude effects of our compounds on WDR5 transcription, we treated MV4-11 cells with the most effective degraders **8g** and **17b** and their corresponding WDR5 ligands **6** and **14** and quantified mRNA by quantitative PCR (qPCR). As expected, even though both degraders reduced the WDR5 protein level, WDR5 mRNA levels were not decreased (see Figure 4d and **Supplementary Information** Figure 5h). Similarly, **8g** mediated WDR5 degradation was completely abolished by co-incubation of MV4-11 cells with WDR5 ligand **6** showing WDR5 depletion requires binding of **8g** to WDR5 (see Figure 4e). We also rescued WDR5 depletion by **17b** through proteasomal inhibition with MG132 and neddylation inhibition with MLN4924 (see Figure 4f). Finally, we also tested if the degradation of WDR5 was limited to MV4-11 cells or if the degrader compounds were also functional in other cell lines. We, thus treated the human leukemia cell line HL-60 with **8g** and observed WDR5 depletion as seen in MV4-11 cells (see Figure 4g). We concluded that **8g** and **17b** mediate depletion of endogenous WDR5 in various cancer cell lines by inducing protein ubiquitylation and degradation.

In order to determine whether **8g-** and **17b-**mediated degradation was specific to WDR5, we analyzed cellular protein levels by quantitative proteomics. To this end, we treated MV4-11 cells with **8g** and **17b** or with their corresponding ligands and compared their protein content to untreated cells by mass spectrometry. Remarkably, among 5805 proteins detected, only WDR5 was significantly and substantially depleted (-log_10_p > 3, log_2_FC < −0.5) after 9 hours of treatment (see Figure 4h-4i). WDR5 levels were not significantly altered by treatment with the ligands, **6** and **14** (see **Supplementary Information** Figure 6).

**Figure 6:**
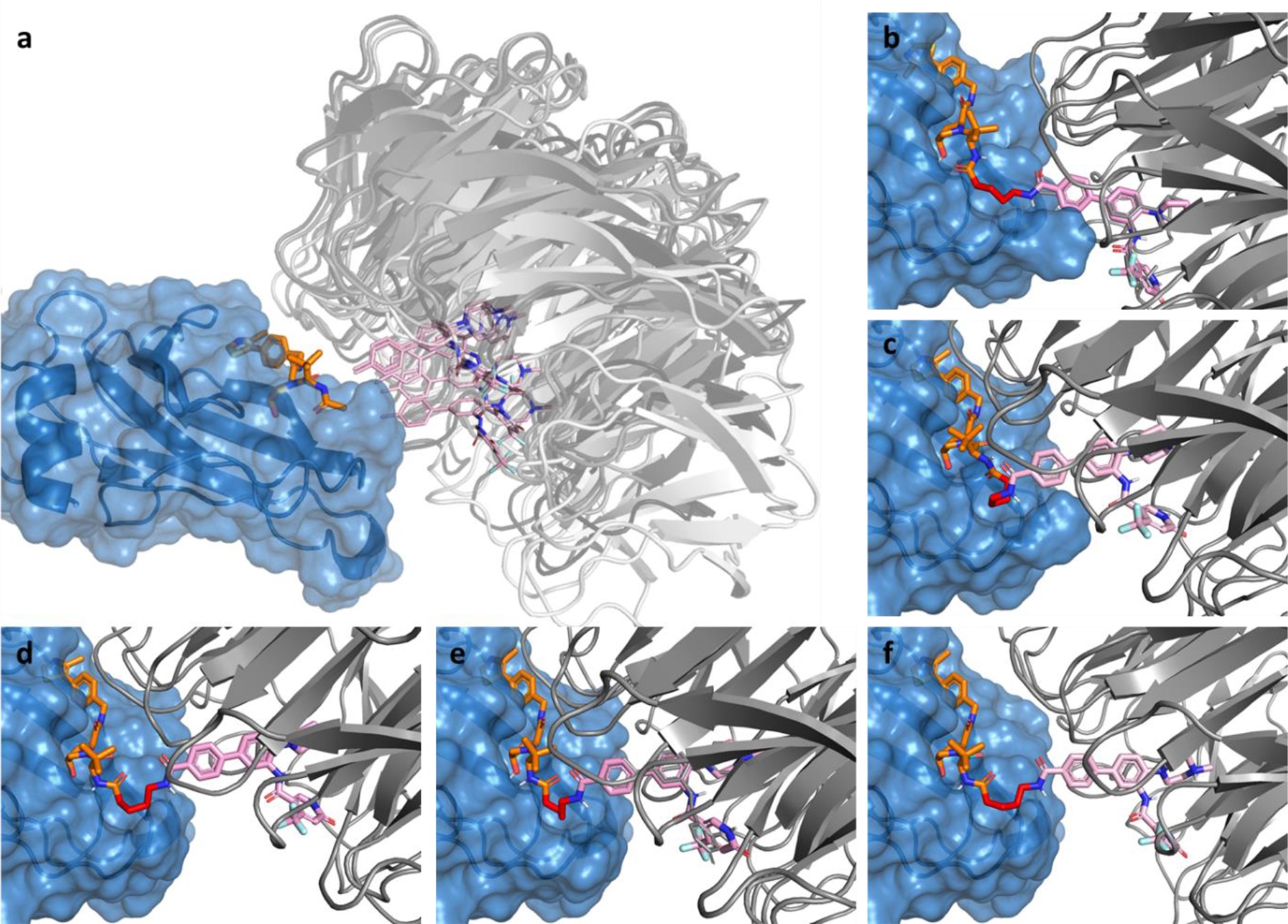
Computational docking studies on WDR5 and VHL. (a) Structures of the selected protein/protein docking solutions compatible with a short linker distance. VHL (blue) and VH032 (orange) form a complex with WDR5 (grey) and a modified OICR-9429 ligand (pink), with the attachment points for linkers in close proximity (<4 Å) (b-f) DSX top-ranked docking pose of degrader 8g docked to each of the selected protein/protein complexes. The functional parts of molecule 8g are coloured in orange for the interaction with VHL, in pink for the interaction with WDR5 and in red for the linker that connects both moieties.

To determine the cellular consequences of WDR5 depletion, we analyzed MV4-11 cell proliferation. We treated MV4-11 cells with different concentration of **17b**, its negative control (**21**) and its ligand (**14**) for 15 days. Only higher concentration (∼10 µM) of **17b** induced a proliferation defect over the course of time whereas neither lower concentration of **17b** nor any incubation with **21** and **14** showed significant growth defect (see Figure 5a). We hypothesized that only high concentrations mediate a long-standing and sufficient depletion of WDR5, which is required to affect cell growth. In fact, all degraders studied here induce only partial degradation of WDR5 and we speculate that more efficient depletion is required to see stronger attenuation of cancer cell growth. As effective degradation via degraders requires a stoichiometric relation between the target protein, the degrader and the E3 ligase, we wondered, whether the expression of VHL might be the limiting factor for degrader efficacy.

To test this hypothesis, we stably expressed exogenous VHL in MV4-11^WDR5-HiBiT^ cells (MV4-11^WDR5-HiBiT/VHL^) (see **Supplementary Information** Figure 5i). Strikingly, HiBiT assay demonstrated superior degradation of WDR5 protein by both **8g** and **17b** in MV4-11^WDR5-HiBiT/VHL^ cells when compared to control cells (see Figure 5b and **Supplementary Information** Figure 3). For **17b**, the D_max_ increased from 52.5 % to 77.8 % whereas DC_50_ decreased from 1.01 µM to 0.155 µM. Immunoblotting confirmed enhanced degradation of endogenous WDR5 after **17b** treatment in MV4-11^WDR5-HiBiT/VHL^ cells (see **Supplementary Information** Figure 5j). To test if enhanced degradation of WDR5 in VHL-overexpressing cells induced a more pronounced cell cycle phenotype, we also generated naive MV4-11 cells with ectopic VHL expression (MV4-11^VHL^) (see **Supplementary Information** Figure 5i). We verified the degradation of endogenous WDR5 in these cells in comparison to control cells (see Figure 5c). Finally, we repeated the cumulative growth analysis in MV4-11^VHL^ cells by incubating the cells with different concentration of **17b**, **21** and **14** for 15 days. In line with stronger WDR5 depletion, high concentration (10 µM) of **17b** showed stronger growth inhibition than that in the control cells (see Figure 5a **and 5d**). Importantly, MV4-11^VHL^ cells also showed growth defects even at lower concentrations of **17b** (5 µM) whereas none of the controls (**21** and **14)** showed notable effects (see Figure 5d). We concluded that degrader-induced partial depletion of WDR5 show moderate but statistically significant cell growth defects in MV4-11 cells.

### Computational studies on WDR5 degraders

Computational docking studies were performed to obtain a structural model that was able to explain the binding of the active degraders **8e-j** to WDR5 and VHL. Therefore, crystal structures of WDR5 in complex with WDR5 antagonist OICR-9429 (PDB: 4QL1)^17^ and VHL in complex with ligand VH032 (PDB: 4W9H)^36^ were prepared for protein/protein docking, as described in the methods section and **Supplementary Information**. In the second step, Protein/protein docking was performed in the Molecular Operating Environment (MOE) to obtain 405 possible protein/protein complexes. For evaluation of the generated protein complexes, the distance between the two linkage sites served as the primary selection criterion. Since the shortest linker used in **8e** showed effective degradation, docking solutions capable of binding both OCIR-9429 and VH032 in their known binding modes while maintaining a linkable distance for the short ethyl moiety were examined. Ten of the obtained protein/protein complexes showed a distance of less than 4 Å between the two critical carbon atoms, with five of these being ranked in the top 20% of the protein/protein docking solutions. None of the complexes stood out as clearly preferred, suggesting rather an ensemble of possible configurations.

In the next step, degraders **8e-j** were docked with GOLD to the five best-ranked of the compatible protein/protein complexes (ranks 52, 53, 59, 78 and 79) and evaluated based on a rescoring with DrugscoreX and RMSD-values with respect to the crystalized binding modes of VHL ligand VH032 and WDR5 antagonist OICR-9429. The docking results show that all active degraders can be placed in these complexes while still occupying their native binding sites in the individual proteins with RMSD values around or below 1 Å. Only degrader **8j** required rearrangements of the scaffolds in most top-ranked docking solutions (**see Supplementary Information**). The structural models obtained by a successive protein/protein- and small-molecule-docking suggest that WDR5 and VHL do not form a dominant ternary complex. Rather, it appears likely that an ensemble of multiple different configurations of the ternary complex may exist in solution, similar (but probably even more diverse) as illustrated in Figure 6.

## Conclusion and Outlook

The MLL/SET HMT complexes as well as the transcription factor family MYC are attractive drug targets. In the strategy presented here, we aimed to modulate both oncogenic functions by degrading the scaffolding protein WDR5 using a PROTAC approach. We used two diverse inhibitor scaffolds allowing for diverse exit vectors from the WDR5 WIN binding site comprising OICR-9429 and a modified pyrroloimidazole based inhibitor in combination with E3 ligase ligands targeting Cereblon, VHL and MDM2. Thus, the study provides interesting SAR developing degraders for this attractive cancer target. In BRET studies, VHL and Cereblon based degraders showed good cell permeability and on target activity in cells. Surprisingly, degraders based on both inhibitor scaffolds led to successful degradation of the target protein indicating good degradability of WDR5. However, only VHL-based degraders led to functional degradation and small changes in linker length and linker type resulted in significant changes in degradation efficacy. Large MDM2 based PROTACs did not show significant binding affinity and were discarded early on in the study. Best degrader molecules showed low nM potency degrading WDR5 in a proteasome and ubiquitin dependent way. However, under the cell lines and condition tested only partial degradation was observed. This could be a kinetic effect depending on the speed of re-synthesis and degradation or potentially due to location of WDR5 on chromatin, which might protect a fraction of the protein from degradation. Nonetheless, the cells could be sensitized to better degradation of protein via overexpression of one of the components of ternary complex (here VHL) which significantly increased the degradation efficiency of the PROTACs. These data suggest that using the PROTAC in cell lines that show higher VHL levels will also increase WDR5 degradation determining sensitivity to the degrader. We established an array of assay systems such as a BRET based target engagement assay and a stable HiBiT cell line, that will allow further testing of future second generation WDR5 degrader molecules with improved potency. However, the molecules presented here are versatile tool molecules that will allow comprehensive evaluation of WDR5 degraders in diverse cancer types and the potential of this strategy for drug discovery.

## Experimental section

### Compound Synthesis

The structures of the synthesized compounds were verified by ^1^H- and ^13^C NMR, and mass spectrometry (ESI/ MALDI). Purity of the final compounds (254, 260 and 280 nm >95%) was determined by analytical HPLC. All commercial chemicals and solvents were used without further purification. Commercially available VHL ligand 1 Hydrochloride was used for the degrader molecules **8a-j** and **17a-g**, while the VHL ligand for the negative control **20** and **21** was obtained analog to Buckley and van Molle^34^. ^1^H NMR and ^13^C NMR spectra were measured in DMSO-d6, MeOD, CD_2_Cl_2_ or CDCl_3_ on a Bruker DPX250, AV300, AV400, AV500, DPX600, AV700 or AV800 spectrometer. Chemical shifts δ are reported in parts per million (ppm). Mass spectra were recorded by the mass spectrometry service team of the Goethe University using a Thermo Fisher Surveyor MSQ system (including TLC-MS for reaction control). High-resolution mass spectra were recorded on a MALDI LTQ ORBITRAP XL device from Thermo Fisher Scientific. Product purification was performed on a PuriFlash Flash Column Chromatography System from Interchim using prepacked silica or RP C18 columns. Product purification was also performed on an Agilent 1260 Infinity II LC System [Eclipse XDB-C18 column (7 μM, 21.2 × 250 mm)] using a gradient of water/MeCN + 0.1% TFA (98:2−5:95) over 40 min with a flow rate of 21 mL/min. Compound purity was analyzed on an Agilent 1260 Infinity II LC System [Eclipse XDB-C18 column (4.6 × 250 mm, 5 μm)] coupled to an Agilent InifinityLab LC/MSD using a gradient of water/MeCN + 0.1% TFA (98:2−5:95) over 25 min at a flow rate of 1 mL/min (see the SI data for HPLC−MS traces of lead compounds). Final compounds **7a-e**, **8a-j** and **9a-c** were synthesized from 5- Bromo-2-Fluoronitrobenzole **1** and 4-Methylpiperazine **2** via intermediates **3−6**, **6a-c** as outlined in Scheme 1. Final compounds **17a-g** were synthesized from Piracetam **10** via intermediates **11-16a-g** as outlined in Scheme 2. More information can be found in the **Supplementary Information**.

### General procedure A (amide coupling)

1.0 eq of the Boc-protected carboxylic acid were dissolved in 1 mL CH_2_Cl_2_ and 1 mL TFA was added. The solution was stirred for 1 h at rt. Excess solvent was removed under reduced pressure. 1.0 eq of the Boc-protected amine were dissolved in 1 mL CH_2_Cl_2_ and 1 mL TFA was added. The solution was stirred for 1 h at rt. Excess solvent was removed under reduced pressure. The crude species was dissolved in 0.5 mL DMF and DIEA was added until the pH of the solution was basic. 1.2 eq HATU was added to the deprotected carboxylic acid. The reaction mixture was stirred for 20 min at rt. The crude species of the deproteced amine rt for 3-5 h. The reaction was stopped with 1 mL water. Saturated NaHCO_3_ solution and saturated NaCl solution were added and the reaction mixture was extracted 4x with EA. The combined organic phases were washed with saturated NaHCO_3_ solution, dried over MgSO_4_ and filtered. The solvent of the organic phase was evapored under reduced pressure. The purification of the crude product was carried out on a preparative HPLC system. **General procedure B** (toslyation and nucleophilic substitution): 1.4 eq Tosylchloride was added portion wise (2x) to 1.0 eq of commercially available alcohol, 0.3 eq DMAP and 1.3 eq triethylamine in 5 mL dichloromethane at −10 °C. The reaction mixture was stirred for 20 min, allowed to warm to room temperature and stirred for another 22 h. The reaction was quenched by adding 3 ml of a saturated solution of NH_4_Cl in water. The organic layer was separated and the remaining aqueous layer was extracted with dichloromethane (3x). The combined organic layers were washed with brine, dried with Na_2_SO_4_ and the solvent was removed under reduced pressure. Purification was achieved by column chromatography (10% MeOH/CH_2_Cl_2_).

2.4 eq of the tosylate linker species in 1.5 mL DMF was added to a solution of the alcohol **15** and 3.2 eq K_2_CO_3_ in 2 mL DMF. The reaction mixture was stirred at 70 °C for 21 h. The reaction was quenched by adding 6 ml water and ethylacetate. The organic layer was separated and the remaining aqueous layer was extracted with ethylacetate (3x). The combined organic layers were dried with MgSO_4_ and the solvent was removed under reduced pressure. Purification was carried out on a preparative HPLC system.

### General procedure C (amide coupling)

This reaction was not performed in an inert atmosphere. A solution of 1.0 eq ester in 4 mL dichloromethane/TFA (1/1) was stirred for 1.5 h. All volatiles were removed under reduced pressure. 1.2 eq HATU was added to a solution of the carboxylic acid and 10.0 eq DIPEA in 2 mL DMF. The reaction mixture was stirred for 15 min, turned orange and 1.1 eq VHL amine hydrochloride was added. The mixture was stirred for another 3.5 h and quenched with water and ethylacetate. The organic layer was separated and the aqueous layer was extracted with ethylacetate (3x). The combined organic layers were dried with MgSO_4_ and the solvent was removed under reduced pressure. Purification was carried out on a preparative HPLC system. The gained product was then dissolved in ethyl acetate and a solution of saturated NaHCO_3_ and saturated NaCl solution and extracted with ethyl acetate. The combined organic phases were dried over MgSO_4_ and the solvent was removed under reduced pressure.

### Synthesis of 1-(4-bromo-2-nitrophenyl)-4-methylpiperazine (3)

5 mL (39.7 mmol, 1.00 eq) 5-Bromo- 2-Fluoro-Nitrobenzene was dissolved in 20 mL EtOH. 4 mL (39.7 mmol, 1.00 eq) N-Methylpiperazine and 13.5 mL (79.4 mmol, 2.00 eq) DIEA were added and the reaction mixture was stirred for 5 h at 80 °C. The reaction was cooled to rt and the solvent was removed under reduced pressure. The reaction mixture was diluted with water and extracted 8x with DCM. The combined organic phases were washed with 1 M HCl, saturated NaCl solution, dried over MgSO_4_ and filtered. The crude product was purified via CC (gradient: 0 % to 10 % MeOH in DCM) to give 11.1 g (37 mmol, 95 %) of an orange powder. R_f_ (5% MeOH/ CH_2_Cl_2_): 0.54. ESI: (calculated): [M+H^+^] 300.03 g/mol, (found): [M+H^+^] 299.98 g/mol. ^1^H NMR (250 MHz, CDCl_3_) δ = 7.94 (d, *^4^J* = 2.3 Hz, 1H), 7.65 (dd, *^3^J* = 8.7 Hz, *^4^J* = 2.4 Hz, 1H), 7.17 (d, *^3^J* = 8.7 Hz, 1H), 3.43 (m, 4H), 3.25 (m, 4H), 2.80 (s, 3H) ppm.

### Synthesis of 5-bromo-2-(4-methylpiperazin-1-yl)aniline (4)

2.00 g (7.40 mmol, 1.00 eq) 1-(4-bromo- 2-nitrophenyl)-4-methylpiperazine was suspended in a mixture of 1,4-Dioxane and water (3:1). 3.20 g (37 mmol, 7.50 eq) Ammonium chloride was added, followed by a slow addition of 2.60 g (37 mmol, 7.50 eq) Zinc dust. The reaction mixture was stirred until a colour change from orange to light pink was observed. The solvent was removed under reduced pressure. The reaction mixture was diluted with saturated NaHCO_3_ solution and extracted 6x with DCM. The combined organic phases were washed with saturated NaCl solution, dried over MgSO_4_, filtered and the solvent was removed under reduced pressure to give 1.57 mg (5.81 mmol, 79%) of a light pink solid. R_f_ (5% MeOH/ CH_2_Cl_2_): 0.3. ESI: (calculated): [M+H^+^] 270.06 g/mol, (found): [M+H^+^] 270.03 g/mol. HPLC: RT = 10.9 min (254 nm, 99%). ^1^H NMR (250 MHz, CDCl_3_) δ = 6.96 – 6.70 (m, 3H), 4.01 (s, 2H), 2.92 (t, *^3^J* = 4.5 Hz, 4H), 2.60 (s, 4H), 2.39 (s, 3H) ppm.

### Synthesis of *tert*-butyl 3’-amino-4’-(4-methylpiperazin-1-yl)-[1,1’-biphenyl]-4-carboxylate (5)

395 mg (1.78 mmol, 1.2 eq) (4-(*tert*-butoxycarbonyl)phenyl)boronic acid was dissolved in an Argon-purged solvent solution (1,4-Dioxane/ Water (3:1)) and 288 mg (7.40 mmol, 5.0 eq) Sodium hydroxide was added. The reaction mixture was stirred for 5 min at rt under Argon atmosphere, then 400 mg (1.48 mmol, 1.0 eq) 5-bromo-2-(4-methylpiperazin-1-yl)aniline and 171 mg (148 µmol, 0.1 eq) Pd(PPh_3_)_4_ were added. The reaction was stirred at 90 °C for 18 h under Argon atmosphere. The reaction mixture was cooled to rt, filtered over celite and washed with MeOH. The solvent was removed under reduced pressure. The reaction mixture was diluted with water and extracted 3x with DCM. The crude product was purified via FC (0 % to 10 % MeOH/ DCM) to give 395 mg (1.08 mmol, 73%) of a white solid. R_f_ (5% MeOH/ CH_2_Cl_2_): 0.28. ESI: (calculated): [M+H^+^] 368.23 g/mol, (found): [M+H^+^] 368.13 g/mol. HPLC: RT = 11.9 min (254 nm, 94%) ^1^H NMR (500 MHz, CD_2_Cl_2_) δ = 7.98 (d, *^3^J* = 8.5 Hz, 2H), 7.59 (d, *^3^J* = 8.5 Hz, 2H), 7.06 (d, *^3^J* = 8.7 Hz, 1H), 6.99 (m, 2H), 3.97 (s, 1H), 2.97 (s, 4H), 2.61 (s, 4H), 2.35 (s, 3H), 1.58 (s, 9H) ppm. ^13^C NMR (126 MHz, CD_2_Cl_2_) δ = 165.9, 145.5, 142.3, 139.8, 136.5, 130.8, 130.1, 126.8, 120.4, 117.6, 113.8, 81.1, 55.0, 51.0, 46.1, 28.3 ppm.

### Synthesis of *tert*-butyl 3’-(6-hydroxy-4-(trifluoromethyl)nicotinamido)-4’-(4-methylpiperazin-1-yl)- [1,1’-biphenyl]-4-carboxylate (6)

56 mg, (272 µmol, 1.00 eq) of 6-Hydroxy-4- (trifluoromethyl)nicotinic acid was dissolved in 1 mL DCM and 228 µL (2.72 mmol, 10 eq) thionyl chloride. The reaction mixture was stirred for 3 h at 50 °C until a colour change from clear to yellow was observed. Excess thionyl chloride was removed under reduced pressure and the acyl chloride was evapored on a high vacuum line for 5 min. The acyl chloride was diluted with 2 mL of DCM and a solution of 3 mL containing 100 mg (272 µmol, 1.00 eq) 3’-amino-4’-(4-methylpiperazin-1-yl)-[1,1’- biphenyl]-4-carboxylate and 44 µL (544 µmol, 2.00 eq) pyridine was added. The reaction mixture was stirred at 50 °C for 18 h. The reaction mixture was cooled to rt, diluted with water and extracted 3x with DCM. The crude product was purified via FC (0 % to 10 % MeOH/ DCM) to give 109 mg (139 µmol, 51%) of a yellow TFA salt with a stoichiometry of 1:2 (product: TFA). R_f_ (5% MeOH/ CH_2_Cl_2_): 0.21. ESI: (calculated): [M+H^+^] 557.23 g/mol, (found): [M+H^+^] 557.08 g/mol. HPLC: RT = 12.9 min (254 nm, 93%). ^1^H NMR (500 MHz, CDCl_3_) δ = 8.97 (s, 1H), 8.69 (s, 1H), 8.04 (d, *^3^J* = 8.3 Hz, 2H), 7.89 (s, 1H), 7.66 (d, *^3^J* = 8.3 Hz, 2H), 7.48 – 7.30 (m, 2H), 6.95 (s, 1H), 3.05 (s, 4H), 2.78 (s, 4H), 2.51 (s, 3H), 1.61 (s, 9H) ppm. ^13^C NMR (75 MHz, DMSO) δ = 164.8, 163.1, 161.1, 143.7, 143.2, 139.5 (m), 138.6 (q, *^2^J* = 32 Hz), 135.0, 134.1, 132.7, 130.1, 129.8, 126.4, 124.0, 122.1 (q, *^1^J* = 273 Hz), 121.0, 118.9 (m), 111.6 (m), 80.7, 52.8, 48.1, 42.3, 27.8 ppm.

### Synthesis of tert-butyl (2-(2-(3’-(6-hydroxy-4-(trifluoromethyl)nicotinamido)-4’-(4-methylpiperazin- 1-yl)-[1,1’-biphenyl]-4-carboxamido)ethoxy)ethyl)carbamate (6a)

20 mg (36 µmol, 1.0eq) tert-butyl 3’-(6-hydroxy-4-(trifluoromethyl)nicotinamido)-4’-(4-methylpiperazin-1-yl)-[1,1’-biphenyl]-4- carboxylate were dissolved in 0.5 mL DCM and 0.5 mL TFA and stirred at rt for 1 h. Excess solvent was evapored. The solid was dissolved in 0.5 mL DMF, then 125 µL (720 µmol, 20 eq) DIEA and 16.4 mg (43 µmol, 1.2 eq) HATU were added. After 15 min, a solution of 7.7 mg (38 µmol, 1.05 eq) tert-butyl (2-(2-aminoethoxy)ethyl)carbamate in 0.5 mL DMF was added. The solution was stirred for 3 h at rt. The reaction mixture was quenched with 2 mL water and 2 mL saturated NaHCO_3_, then the reaction was extracted 3x with EA. The organic phase was dried over MgSO_4_, filtered and the solvent was removed under reduced pressure. The crude product was purified using by HPLC to obtain 18 mg of a TFA salt with unknown stoichiometry as a white solid. ESI: (calculated) [M+H^+^] 687.31 g/mol, (found) [M+H^+^] 687.52 g/mol. HPLC: RT = 11.6 min (254 nm, 96%). ^1^H NMR (400 MHz, DMSO) δ = 9.47 (s, 1H), 8.52 (t, *^3^J* = 5.4 Hz, 1H), 8.13 (d, *^3^J* = 2.1 Hz), 7.97 (s, 1H), 7.93 (d, *^3^J* = 8.4 Hz, 2H), 7.68 (d, *^3^J* = 8.5 Hz, 2H), 7.52 (dd, *^3^J* = 8.4 Hz, *^4^J* = 2.2 Hz), 7.27 (d, *^3^J* = 8.4 Hz, 1H), 6.82 (s, 1H), 6.76 (s, 1H), 3.55 – 3.52 (m, 2H), 3.47 – 3.38 (m, 4H), 3.11 – 3.06 (m, 2H), 2.99 – 2.85 (m, 4H), 2.50 (s, 4H) 2.23 (s, 3H), 1.36 (s, 9H) ppm. ^13^C NMR (101 MHz, DMSO) δ = 168.0, 166.0, 162.8, 161.2, 144.9, 142.2, 140.0, 139.5 (q, ^2^J = 35 Hz), 134.2, 133.00 132.3, 132.2, 128.3, 127.9, 126.0, 123.9, 122.6 (q, ^1^J = 240 Hz), 122.3, 121.0, 120.4, 119.1 (m), 110.3 (m), 77.59, 68.99, 68.7, 54.7, 51.0, 45.7, 39.1, 38.9, 28.2 ppm.

### Synthesis of tert-butyl (1-(3’-(6-hydroxy-4-(trifluoromethyl)nicotinamido)-4’-(4-methylpiperazin-1- yl)-[1,1’-biphenyl]-4-yl)-1-oxo-5,8,11,14,17,20,23-heptaoxa-2-azapentacosan-25-yl)carbamate (6b)

20 mg (36 µmol, 1.0 eq) tert-butyl 3’-(6-hydroxy-4-(trifluoromethyl)nicotinamido)-4’-(4- methylpiperazin-1-yl)-[1,1’-biphenyl]-4-carboxylate were dissolved in 0.5 mL DCM and 0.5 mL TFA and stirred at rt for 1 h. Excess solvent was evapored. The solid was dissolved in 0.5 mL DMF, then 125 µL (720 µmol, 20 eq) DIEA and 16.4 mg (43 µmol, 1.2 eq) HATU were added. After 15 min, a solution of 17.7 mg (38 µmol, 1.05 eq) tert-butyl (23-amino-3,6,9,12,15,18,21-heptaoxatricosyl)carbamate in 0.5 mL DMF was added. The solution was stirred for 3 h at rt. The reaction mixture was quenched with 2 mL water and 2 mL saturated NaHCO_3_, then the reaction was extracted 3x with EA. The organic phase was dried over MgSO_4_, filtered and the solvent was removed under reduced pressure. The crude product was purified using by HPLC to obtain 10 mg of a TFA salt with unknown stoichiometry as white oil. ESI: (calculated) [M+H^+^] 951.47 g/mol, (found) [M+H^+^] 951.95 g/mol. HPLC: RT = 11.8 min (254 nm, 94%). ^1^H NMR (400 MHz, DMSO) δ = 9.46 (s, 1H), 8.53 (t, *^3^J* = 5.7 Hz, 1H), 8.12 (d, *^4^J* = 2.2 Hz, 1H), 7.97 (s, 1H), 7.94 (d, *^3^J* = 8.5 Hz, 2H), 7.68 (d, *^3^J* = 8.5 Hz, 2H), 7.53 (dd, *^3^J* = 8.4 Hz, *^4^J* = 2.2 Hz, 1H), 7.27 (d, *^3^J* = 8.4 Hz, 1H), 6.82 (s, 1H), 6.72 (t, *^3^J* = 5.3 Hz, 1H), 3.61 – 3.47 (m, 28H), 3.44 (dd, *^2^J* = 11.6 Hz, *^3^J* = 5.8 Hz, 2H), 3.36 (t, *^3^J* = 6.1 Hz, 4H), 3.05 (q, *^3^J* = 6.0 Hz, 2H), 2.96 – 2.86 (m, 4H), 2.24 (s, 3H), 1.36 (s, 9H) ppm.

### Synthesis of tert-butyl (4-((3’-(6-hydroxy-4-(trifluoromethyl)nicotinamido)-4’-(4-methylpiperazin-1- yl)-[1,1’-biphenyl]-4-carboxamido)methyl)benzyl)carbamate (6c)

20 mg (36 µmol, 1.0eq) tert-butyl 3’-(6-hydroxy-4-(trifluoromethyl)nicotinamido)-4’-(4-methylpiperazin-1-yl)-[1,1’-biphenyl]-4- carboxylate were dissolved in 0.5 mL DCM and 0.5 mL TFA and stirred at rt for 1 h. Excess solvent was evapored. The solid was dissolved in 0.5 mL DMF, then 125 µL (720 µmol, 20 eq) DIEA and 16.4 mg (43 µmol, 1.2 eq) HATU were added. After 15 min, a solution of 8.90 mg (38 µmol, 1.05 eq) tert-butyl (4-(aminomethyl)benzyl)carbamate in 0.5 mL DMF was added. The solution was stirred for 3 h at rt. The reaction mixture was quenched with 2 mL water and 2 mL saturated NaHCO_3_, then the reaction was extracted 3x with EA. The organic phase was dried over MgSO_4_, filtered and the solvent was removed under reduced pressure. The crude product was purified using by HPLC to obtain 14 mg of TFA salt with unknown stochimetry as a white solid. ESI: (calculated) [M+H^+^] 719.32 g/mol, (found) [M+H^+^] 719.54 g/mol. HPLC: RT = 12.2 min (254 nm, 91%). ^1^H NMR (400 MHz, MeOD) δ 8.29 (d, *^4^J* = 1.8 Hz, 1H), 7.99 (s, 1H), 7.92 (d, *^3^J* = 8.4 Hz, 2H), 7.72 (d, *^3^J* = 8.4 Hz, 2H), 7.52 (dd, *^3^J* = 8.3 Hz, *^4^J* = 2.1 Hz, 1H), 7.35 - 7.33 (m, 3H), 7.25 (d, *^3^J* = 8.1 Hz, 2H), 6.92 (s, 1H), 4.57 (s, 2H), 4.21 (s, 2H), 3.01 (s, 4H), 2.69 (s, 4H), 2.39 (s, 3H), 1.44 (s, 9H).

### Synthesis of *N*-(4’-((2-(2-((2-(2,6-dioxopiperidin-3-yl)-1,3-dioxoisoindolin-4- yl)amino)ethoxy)ethyl)carbamoyl)-4-(4-methylpiperazin-1-yl)-[1,1’-biphenyl]-3-yl)-6-hydroxy-4- (trifluoromethyl)nicotinamide (7a)

The reaction was carried out as described in General Procedure A. The gained product was then dissolved in ethyl acetate and a solution of saturated NaHCO_3_ and saturated NaCl solution and extracted with ethyl acetate. The combined organic phases were dried over MgSO_4_ and the solvent was removed under reduced pressure to give 17.2 mg (20.4 µmol, 71 %) of a yellow solid. MALDI: (calculated) [M+H^+^] 843.31 g/mol, (found) [M+H^+^] 843.41 g/mol. HPLC: RT = 11.6 min (254 nm, 100%). HRMS: (calculated) [M+H^+^] 843.3072 g/mol, (found) [M+H^+^] 843.3067 g/mol, ^1^H NMR (500 MHz, DMSO) δ = 12.54 (s, 1H), 11.08 (s, 1H), 9.52 (s, 1H), 8.53 (t, *^3^J* = 5.6 Hz, 1H), 8.14 (d, *^4^J* = 1.8 Hz, 1H), 8.00 (s, 1H), 7.91 (d, *^3^J* = 8.4 Hz, 2H), 7.66 (d, *^3^J* = 8.4 Hz, 2H), 7.60 – 7.48 (m, 2H), 7.28 (d, *^3^J* = 8.4 Hz, 1H), 7.15 (d, *^3^J* = 8.6 Hz, 1H), 7.02 (d, *^3^J* = 7.0 Hz, 1H), 6.83 (s, 1H), 6.63 (t, *^3^J* = 5.7 Hz, 1H), 5.03 (dd, *^3^J* = 12.8 Hz, *^4^*J = 5.4 Hz, 1H), 3.66 (t, *^3^J* = 5.4 Hz, 2H), 3.61 (t, *^3^J* = 6.0 Hz, 2H), 3.48 (dd, *^3^J* = 13.3 Hz, *^4^J* = 4.9 Hz, 4H), 2.99 (s, 4H), 2.91 – 2.80 (m, 1H), 2.75 (s, 4H), 2.58 - 2.57 (m, 1H), 2.52 (s, 1H), 2.42 (s, 3H), 2.01 - 1.98 (m, 1H) ppm. ^13^C NMR (126 MHz, DMSO) δ = 172.7, 170.1, 168.9, 167.3, 166.0, 162.9, 161.1, 146.4, 144.3, 142.1, 139.3, 138.5 (q, *^2^J* = 33 Hz), 136.2, 134.5, 133.0, 132.3, 132.1, 127.9, 126.0, 123.9, 122.3, 122.0 (q, *^1^J* = 275 Hz), 120.6, 119.0, 117.5, 111.7, 110.7, 109.2, 68.8, 68.8, 54.1, 50.1, 48.5, 44.8, 41.7, 31.0, 25.5, 22.1 ppm.

### Synthesis of *N*-(4’-((2-(2-(2-(2-((2-(2,6-dioxopiperidin-3-yl)-1,3-dioxoisoindolin-4- yl)amino)ethoxy)ethoxy)ethoxy)ethyl)carbamoyl)-4-(4-methylpiperazin-1-yl)-[1,1’-biphenyl]-3-yl)- 6-hydroxy-4-(trifluoromethyl)nicotinamide (7b)

The reaction was carried out as described in General Procedure A. The gained product was then dissolved in ethyl acetate and a solution of saturated NaHCO_3_ and saturated NaCl solution and extracted with ethyl acetate. The combined organic phases were dried over MgSO_4_ and the solvent was removed under reduced pressure to give 8.58 mg (8.95 µmol, 33 %) of a light yellow solid. MALDI: (calculated) [M+H^+^] 959.39 g/mol, (found) [M+H^+^] 959.47 g/mol. HPLC: RT = 11.8 min (254 nm, 98%). HRMS: (calculated) [M+H^+^] 959.3909 g/mol, (found) [M+H^+^] 959.3905 g/mol. ^1^H NMR (500 MHz, DMSO) δ = 11.08 (s, 1H), 9.45 (s, 1H), 8.44 (t, *^3^J* = 5.5 Hz, 1H), 8.12 (d, *^4^J* = 1.7 Hz, 1H), 7.98 (s, 1H), 7.91 (d, *^3^J* = 8.4 Hz, 2H), 7.67 (d, *^3^J* = 8.4 Hz, 2H), 7.61 – 7.54 (m, 1H), 7.51 (dd, *^3^J* = 8.3 Hz, *^4^*J = 2.2 Hz, 1H), 7.26 (d, *^3^J* = 8.4 Hz, 1H), 7.08 (d, *^3^J* = 8.6 Hz, 1H), 7.01 (d, *^3^J* = 7.0 Hz, 1H), 6.81 (s, 1H), 6.65 (t, *^3^J* = 5.9 Hz, 1H), 5.04 (dd, *^3^J* = 12.7 Hz, *^4^*J = 5.4 Hz, 1H), 3.61 – 3.35 (m, 21H), 2.95 – 2.89 (m, 3H), 2.91 - 2.84 (m, 1H), 2.62 – 2.55 (m, 1H), 2.23 (s, 3H), 2.07 – 1.95 (m, 1H), 1.85 – 1.72 (m, 5H) ppm. ^13^C NMR (126 MHz, DMSO) δ = 172.8, 170.1, 168.8, 167.3, 165.8, 162.8, 162.3, 146.4, 144.8, 142.1, 139.0, 138.5 (q, *^2^J* = 32 Hz), 136.2, 134.2, 133.2, 132.2, 132.2, 127.8, 126.0, 123.9, 122.2, 122.0 (q, *^1^J* = 275 Hz), 120.4, 118.8, 117.1, 111.7, 110.3, 109.0, 69.8, 69.7, 69.7, 69.6, 68.3, 68.2, 54.7, 51.0, 48.5, 45.8, 40.4, 36.7, 35.8, 31.0, 29.4, 28.9, 22.1 ppm.

### Synthesis of *N*-(4’-((17-((2-(2,6-dioxopiperidin-3-yl)-1,3-dioxoisoindolin-4-yl)amino)-3,6,9,12,15- pentaoxaheptadecyl)carbamoyl)-4-(4-methylpiperazin-1-yl)-[1,1’-biphenyl]-3-yl)-6-hydroxy-4- (trifluoromethyl)nicotinamide (7c)

The reaction was carried out as described in General Procedure A. The gained product was then dissolved in ethyl acetate and a solution of saturated NaHCO_3_ and saturated NaCl solution and extracted with ethyl acetate. The combined organic phases were dried over MgSO_4_ and the solvent was removed under reduced pressure to give 18 mg (14.4 µmol, 41%) of a yellow oil as TFA salt (1:2/ product: TFA.) MALDI: (calculated) [M+H^+^] 1019.41 g/mol, (found) [M+H^+^] 1019.52 g/mol. HPLC: RT = 11.6 min (254 nm, 100%). HRMS: (calculated) [M+H^+^] 1019.4121 g/mol, (found) [M+H^+^] 1019.4117 g/mol. ^1^H NMR (500 MHz, DMSO) δ = 11.10 (s, 1H), 9.20 (s, 1H), 8.59 (d, *^3^J* = 5.2 Hz, 1H), 8.32 (d, *^4^J* = 1.7 Hz, 1H), 8.14 (s, 1H), 7.94 (d, *^3^J* = 8.3 Hz, 6H), 7.68 (d, *^3^J* = 8.4 Hz, 6H), 7.57 (dd, *^3^J* = 8.4 Hz, *^3^J* = 7.3 Hz, 1H), 7.46 (dd, *^3^J* = 8.3 Hz, *^4^J* = 2.0 Hz, 1H), 7.29 (d, *^3^J* = 8.4 Hz, 1H), 7.13 (t, *^3^J* = 8.4 Hz, 1H), 7.06 – 6.94 (m, 1H), 6.58 (dd, *^2^J* = 12.2 Hz, *^3^J* = 6.4 Hz, 1H), 6.46 (s, 1H), 5.05 (dd, *^3^J* = 12.7 Hz, *^4^J* = 5.4 Hz, 1H), 3.67 – 3.32 (m, 28H), 2.89 (t, *^3^J* = 4.4 Hz, 4H), 2.60 – 2.56 (m, 1H), 2.48 (s, 1H), 2.28 – 2.25 (m, 1H), 2.22 (s, 3H), 2.08 – 1.95 (m, 1H) ppm. ^13^C NMR (126 MHz, DMSO) δ = 172.8, 170.1, 168.9, 167.3, 165.9, 163.1, 161.1, 146.4, 142.9, 142.0, 139.5, 138.6 (q, *^2^J* = 32 Hz), 136.2, 135.3, 133.2, 132.6, 132.1, 128.0, 126.1, 123.9, 122.3, 122.1 (q, *^1^J* = 275 Hz), 121.0, 119.0, 117.4, 111.7, 110.7, 109.2, 69.8, 69.8, 69.7, 69.6, 68.9, 68.9, 52.8, 48.6, 48.1, 42.4, 41.7, 38.9, 31.0, 22.1 ppm.

### Synthesis of *N*-(4’-((23-((2-(2,6-dioxopiperidin-3-yl)-1,3-dioxoisoindolin-4-yl)amino)- 3,6,9,12,15,18,21-heptaoxatricosyl)carbamoyl)-4-(4-methylpiperazin-1-yl)-[1,1’-biphenyl]-3-yl)-6- hydroxy-4-(trifluoromethyl)nicotinamide (7d)

The reaction was carried out as described in General Procedure A. The gained product was then dissolved in ethyl acetate and a solution of saturated NaHCO_3_ and saturated NaCl solution and extracted with ethyl acetate. The combined organic phases were dried over MgSO_4_ and the solvent was removed under reduced pressure to give 20.5 mg (18.5 µmol, 69 %) of a yellow oil. MALDI: (calculated) [M+H^+^] 1107.46 g/mol, (found) [M+H^+^] 1107.52 g/mol. HRMS: (calculated) [M+H^+^] 1107.4645 g/mol, (found) [M+H^+^] 1107.4638 g/mol. HPLC: RT = 11.7 min (254 nm, 97%). ^1^H NMR (500 MHz, DMSO) δ = 12.51 (s, 1H), 11.08 (s, 1H), 9.47 (s, 1H), 8.54 (t, *^3^J* = 5.6 Hz, 1H), 8.12 (d, *^4^J* = 2.0 Hz, 1H), 7.98 (s, 1H), 7.94 (d, *^3^J* = 8.4 Hz, 2H), 7.68 (d, *^3^J* = 8.4 Hz, 2H), 7.60 - 7.55 (m, 1H), 7.52 (dd, *^3^J* = 8.3 Hz, *^4^J* = 2.2 Hz, 1H), 7.26 (d, *^3^J* = 8.4 Hz, 1H), 7.13 (d, *^3^J* = 8.6 Hz, 1H), 7.03 (d, *^3^J* = 7.0 Hz, 1H), 6.82 (s, 1H), 6.59 (t, *^3^J* = 5.7 Hz, 1H), 5.05 (dd, *^3^J* = 12.7 Hz, *^4^J* = 5.4 Hz, 1H), 3.61 (t, *^3^J* = 5.4 Hz, 2H), 3.58 – 3.41 (m, 32H), 3.33 – 3.25 (m, 2H), 2.95 – 2.89 (m, 4H), 2.89 – 2.83 (m, 1H), 2.61 - 2.51 (m, 2H), 2.23 (s, 3H), 2.06 – 1.96 (m, 1H) ppm. ^13^C NMR (126 MHz, DMSO) δ = 172.7, 170.0, 168.9, 167.3, 165.9, 162.8, 161.1, 146.4, 144.9, 142.2, 139.2, 138.5 (q, *^2^J* = 32 Hz), 136.2, 134.2, 132.9, 132.2, 132.1, 127.9, 126.0, 123.9, 122.3, 122.0 (q, *^1^J* = 275 Hz), 120.4, 119.0, 117.4, 111.7, 110.64 109.2, 69.8, 69.8, 69.7, 69.6, 68.9, 68.9, 54.7, 51.0, 48.6, 45.7, 41.7, 31.0, 22.1 ppm.

### Synthesis of N-(4’-((4-(((2-(2,6-dioxopiperidin-3-yl)-1,3-dioxoisoindolin-4- yl)amino)methyl)benzyl)carbamoyl)-4-(4-methylpiperazin-1-yl)-[1,1’-biphenyl]-3-yl)-6-hydroxy-4- (trifluoromethyl)nicotinamide (7e)

The reaction was carried out as described in General Procedure A. The gained product was then dissolved in ethyl acetate and a solution of saturated NaHCO_3_ and saturated NaCl solution and extracted with ethyl acetate. The combined organic phases were dried over MgSO_4_ and the solvent was removed under reduced pressure to give 7.6 mg (6.9 µmol, 30%) of a yellow solid as TFA salt (1:2/ product: TFA). MALDI: (calculated) [M+Na^+^] 897.29 g/mol, [M+H^+^] 875.31 g/mol, (found) [M+Na^+^] 897.2018 g/mol (100), [M+H^+^] 875.2220 g/mol (70). HRMS: (calculated) [M+H^+^] 875.3123 g/mol, (found) [M+H^+^] 875.3120 g/mol. HPLC: RT = 12.1 min (254 nm, 95%). ^1^H NMR (500 MHz, DMSO) δ = 11.09 (s, 1H), 10.11 (s, 1H), 9.61 (s, 1H), 9.08 (t, *^3^J* = 5.9 Hz, 1H), 8.25 - 8. 22 (m, 1H), 8.04 (s, 1H), 7.95 (d, *^3^J* = 8.4 Hz, 2H), 7.69 (d, *^3^J* = 8.4 Hz, 2H), 7.56 (dd, *^3^J* = 8.3 Hz, *^4^J* = 2.1 Hz, 1H), 7.450 - 7.47 (m, 1H), 7.38 – 7.28 (m, 3H), 7.28 – 7.15 (m, 3H), 7.00 (d, *^3^J* = 7.0 Hz, 1H), 6.95 (d, *^3^J* = 8.6 Hz, 1H), 6.84 (s, 1H), 5.08 - 5.04 (m, 1H), 4.56 (d, *^3^J* = 4.4 Hz, 2H), 4.49 (d, *^3^J* = 4.8 Hz, 2H), 3.55 - 3.52 (m, 2H), 3.33 – 3.19 (m, 4H), 3.08 - 3.04 (m, 2H), 2.92 – 2.82 (m, 4H), 2.59 - 2.54 (m, 2H), 2.04 −2.00 (m, 1H) ppm. ^13^C NMR (126 MHz, DMSO) δ = 173.3, 170.5, 169.2, 167.7, 166.3, 163.6, 161.6, 146.6, 143.4, 142.5, 140.6, 139.5, 139.0, 138.6, 136.6, 135.8, 133.6, 133.1, 132.6, 129.0, 128.5, 126.6, 126.4, 126.1, 125.9, 124.4, 122.9, 122.6 (d, *^1^J* = 275 Hz), 121.5, 119.0, 118.1, 111.3, 111.2, 110.0, 53.3, 49.0, 48.6, 45.9, 43.0, 42.9, 31.4, 22.6 ppm.

### Synthesis of 6-hydroxy-N-(4’-((2-(3-(((S)-1-((2S,4R)-4-hydroxy-2-((4-(4-methylthiazol-5- yl)benzyl)carbamoyl)pyrrolidin-1-yl)-3,3-dimethyl-1-oxobutan-2-yl)amino)-3- oxopropoxy)ethyl)carbamoyl)-4-(4-methylpiperazin-1-yl)-[1,1’-biphenyl]-3-yl)-4- (trifluoromethyl)nicotinamide (8a)

The reaction was carried out as described in General Procedure A. The gained product was then dissolved in ethyl acetate and a solution of saturated NaHCO_3_ and saturated NaCl solution and extracted with ethyl acetate. The combined organic phases were dried over MgSO_4_ and the solvent was removed under reduced pressure to give 27.4 mg, 26.6 µmol, 59% of a white solid. MALDI: (calculated): [M+Na^+^] 1050.41 g/mol, (found): [M+Na^+^] 1050.20 g/mol. HRMS: (calculated) [M+Na^+^] 1050.4129 g/mol, (found) [M+Na^+^] 1050.4121 g/mol. HPLC: RT = 11.3 min (254 nm, 97%). ^1^H NMR (500 MHz, DMSO) δ = 9.47 (s, 1H), 8.97 (s, 1H), 8.56 (t, *^3^J* = 6.1 Hz, 1H), 8.50 (t, *^3^J* = 5.5 Hz, 1H), 8.12 (d, *^3^J* = 2.1 Hz, 1H), 8.02 – 7.91 (m, 4H), 7.68 (d, *^3^J* = 8.5 Hz, 2H), 7.52 (dd, *^3^J* = 8.4 Hz, *^4^J*= 2.2 Hz, 1H), 7.45 – 7.33 (m, 4H), 7.26 (d, *^3^J* = 8.4 Hz, 1H), 6.82 (s, 1H), 5.13 (s, 1H), 4.56 (d, *^3^J* = 9.4 Hz, 1H), 4.46 – 4.38 (m, 2H), 4.35 (s, 1H), 4.24 - 4.20 (m, 1H), 3.71 – 3.58 (m, 4H), 3.58 – 3.48 (m, 2H), 3.44 - 3.41 (m, 4H), 2.98 – 2.85 (m, 4H), 2.61 – 2.52 (m, 1H), 2.44 – 2.34 (m, 2H), 2.23 (s, 3H), 2.07 – 1.99 (m, 1H), 1.96 – 1.84 (m, 1H), 0.92 (s, 9H) ppm. ^13^C NMR (126 MHz, DMSO) δ = 171.9, 170.0, 169.6, 165.9, 162.8, 161.2, 151.4, 147.7, 144.9, 142.2, 139.5, 139.2, 138.5 (q, *^2^J* = 32 Hz), 134.2, 132.9, 132.2, 131.2, 129.6, 128.6, 127.9, 127.4, 126.0, 123.9, 122.3, 122.0 (d, *^1^J* = 275 Hz), 120.4, 119.0, 111.7, 68.9, 68.6, 66.6, 58.7, 56.4, 56.3, 54.7, 51.0, 45.7, 41.7, 39.0, 37.9, 35.6, 35.4, 26.3, 15.9 ppm.

### Synthesis of 6-hydroxy-*N*-(4’-((2-(2-(3-(((*S*)-1-((2*S*,4*R*)-4-hydroxy-2-((4-(4-methylthiazol-5- yl)benzyl)carbamoyl)pyrrolidin-1-yl)-3,3-dimethyl-1-oxobutan-2-yl)amino)-3- oxopropoxy)ethoxy)ethyl)carbamoyl)-4-(4-methylpiperazin-1-yl)-[1,1’-biphenyl]-3-yl)-4- (trifluoromethyl)nicotinamide (8b)

The reaction was carried out as described in General Procedure A to give 14.2 mg (10 µmol, 23 %) of a white solid as TFA salt (1:3/ product: TFA.) MALDI: (calculated) [M+Na^+^] 1094.44 g/mol, (found) [M+Na^+^] 1094.44 g/mol. HPLC: RT = 11.3 min (254 nm, 100%). HRMS: (calculated) [M+Na^+^] 1094.4392 g/mol, (found) [M+Na^+^] 1094.4384 g/mol. ^1^H NMR (600 MHz, DMSO) δ = 9.92 (s, 1H), 9.59 (s, 1H), 8.97 (s, 1H), 8.56 (t, *^3^J* = 5.6 Hz, 2H), 8.22 (s, 1H), 8.04 (s, 1H), 7.95 - 7.90 (m, 3H), 7.69 (d, *^3^J* = 8.2 Hz, 2H), 7.55 (dd, *^3^J* = 8.3 Hz, *^4^J* = 1.4 Hz, 1H), 7.42 - 7.37 (m, 4H), 7.31 (d, *^3^J* = 8.3 Hz, 1H), 6.83 (s, 1H), 4.55 (d, *^3^J* = 9.4 Hz, 1H), 4.49 – 4.38 (m, 2H), 4.35 (s, 1H), 4.22 (dd, *^3^J* = 15.8 Hz, *^4^J* = 5.5 Hz, 2H), 3.68 - 3.65 (m, 1H), 3.64 – 3.56 (m, 3H), 3.56 – 3.47 (m, 8H), 3.46 - 4.42 (m, 2H), 3.28 - 3.25 (m, 2H), 3.24 - 3.19 (m, 2H), 3.06 - 3.02 (m, 2H), 2.87 (s, 3H), 2.53 (s, 1H), 2.44 (s, 3H), 2.40 – 2.29 (m, 1H), 2.08 – 2.00 (m, 1H), 1.96 – 1.86 (m, 1H), 0.93 (s, 9H) ppm. ^13^C NMR (126 MHz, DMSO) δ = 171.9, 170.0, 169.6, 165.9, 163.1, 161.1, 151.5, 147.7, 142.9, 142.0, 139.5, 139.1, 138.5 (q, *^2^J* = 32 Hz), 135.3, 133.2, 132.64, 131.2, 129.6, 128.6, 128.0, 127.4, 126.1, 123.9, 122.32, 122.1 (q, *^1^J* = 275 Hz), 121.0, 119.0, 118.8, 111.6, 69.6, 69.5, 69.0, 68.9, 66.9, 58.7, 56.4, 56.3, 52.85, 48.1, 42.4, 41.7, 38.0, 35.7, 35.4, 26.3, 15.9 ppm.

### Synthesis of 6-hydroxy-*N*-(4’-(((*S*)-17-((2*S*,4*R*)-4-hydroxy-2-((4-(4-methylthiazol-5- yl)benzyl)carbamoyl)pyrrolidine-1-carbonyl)-18,18-dimethyl-15-oxo-3,6,9,12-tetraoxa-16- azanonadecyl)carbamoyl)-4-(4-methylpiperazin-1-yl)-[1,1’-biphenyl]-3-yl)-4- (trifluoromethyl)nicotinamide (8c)

The reaction was carried out as described in General Procedure A to give 11.5 mg (7.11 µmol, 16 %) of a white solid as TFA salt (1:4/ product: TFA.) MALDI: (calculated) [M+H^+^] 1160.51 g/mol, (found) [M+H^+^] 1160.58 g/mol. HPLC: RT = 11.4 min (254 nm, 99%). HRMS: (calculated) [M+Na^+^] 1182.4916 g/mol, (found) [M+Na^+^] 1182.4913 g/mol. ^1^H NMR (500 MHz, DMSO) δ = 9.76 (s, 1H), 9.57 (s, 1H), 8.97 (s, 1H), 8.56 (t, *^3^J* = 5.4 Hz, 2H), 8.21 (d, *^4^J* = 1.8 Hz, 1H), 8.03 (s, 1H), 7.95 (d, *^3^J* = 8.4 Hz, 2H), 7.90 (d, *^3^J* = 9.4 Hz, 1H), 7.69 (d, *^3^J* = 8.4 Hz, 2H), 7.56 (dd, *^3^J* = 8.3 Hz, *^4^J* = 2.1 Hz, 1H), 7.40 (q, *^3^J* = 8.3 Hz, 5H), 7.31 (d, *^3^J* = 8.4 Hz, 1H), 6.84 (s, 1H), 5.12 (s, 1H), 4.54 (d, *^3^J* = 9.4 Hz, 1H), 4.48 – 4.39 (m, 2H), 4.35 (s, 1H), 4.21 (dd, *^3^J* = 15.9 Hz, *^4^J* =5.5 Hz, 1H), 3.68 - 3.64 (m, 1H), 3.64 – 3.57 (m, 4H), 3.56 – 3.41 (m, 20H), 3.25 – 3.15 (m, 2H), 3.05 - 3.00 (m, 2H), 2.86 (s, 3H), 2.44 (s, 3H), 2.37 - 2.30 (m, 1H), 2.07 – 1.99 (m, 1H), 1.95 – 1.85 (m, 1H), 0.93 (s, 9H) ppm. ^13^C NMR (126 MHz, DMSO) δ = 171.9, 170.0, 169.5, 165.9, 163.1, 161.1, 151.5, 147.6, 142.9, 142.0, 139.5, 139.1, 138.5 (q, *^2^J* = 33 Hz), 135.4, 133.2, 132.7, 131.2, 129.6, 128.6, 128.0, 127.4, 126.1, 123.9, 122.3, 122.1 (q, *^1^J* = 277 Hz), 121.0, 118.9, 111.7, 69.8, 69.7, 69.7, 69.6, 69.5, 68.9, 68.9, 66.9, 58.7, 56.4, 56.3, 52.8, 48.1, 42.4, 41.7, 38.0, 35.7, 35.4, 26.3, 15.9 ppm.

### Synthesis of 6-hydroxy-N-(4’-(((S)-1-((2S,4R)-4-hydroxy-2-((4-(4-methylthiazol-5- yl)benzyl)carbamoyl)pyrrolidin-1-yl)-3,3-dimethyl-1-oxobutan-2-yl)carbamoyl)-4-(4- methylpiperazin-1-yl)-[1,1’-biphenyl]-3-yl)-4-(trifluoromethyl)nicotinamide (8d)

The reaction was carried out as described in General Procedure A to give 22.2 mg, 24.3 µmol, 67% of a white solid. ESI: (calculated) [M+H^+^] 913.37 g/mol, (found) [M+H^+^] 913.63 g/mol. MADLI: (calculated) [M+Na^+^] 935.3496 g/mol, (found) [M+Na^+^] 935.0969 g/mol. HRMS: (calculated) [M+Na^+^] 935.3496 g/mol, (found) [M+Na^+^] 935.3486 g/mol. HPLC: RT = 11.8 min (254 nm, 96%). ^1^H NMR (600 MHz, DMSO) δ = 9.94 (s, 1H), 9.59 (s, 1H), 8.99 (s, 1H), 8.59 (t, *^3^J* = 5.7 Hz, 1H), 8.23 (s, 1H), 8.03 (d, *^3^J* = 10.0 Hz, 2H), 7.98 (d, *^3^J* = 7.8 Hz, 2H), 7.69 (d, *^3^J* = 7.8 Hz, 2H), 7.56 (d, *^3^J* = 8.3 Hz, 1H), 7.41 (q, *^3^J* = 8.1 Hz, 4H), 7.33 (d, *^3^J* = 8.3 Hz, 1H), 6.84 (s, 1H), 4.80 (d, *^3^J* = 9.0 Hz, 1H), 4.47 (t, *^3^J* = 8.1 Hz, 1H), 4.47 – 4.41 (m, 2H), 4.39 (s, 1H), 4.25 (dd, *^3^J* = 15.7 Hz, *^4^J* = 5.3 Hz, 1H), 3.75 (s, 2H), 3.54 – 3.52 (m, 2H), 3.29 – 3.17 (m, 2H), 3.23 – 3.19 (m, 2H), 3.06 – 3.02 (m, 2H), 2.87 (s, 3H), 2.45 (s, 3H), 2.10 – 2.03 (m, 1H), 1.96 – 1.90 (m, 1H), 1.05 (s, 9H) ppm. ^13^C NMR (151 MHz, DMSO) δ = 171.9, 169.5, 166.2, 162.9, 161.1, 151.5, 147.8, 142.9, 142.4, 139.5, 139.4, 138.8 (q, *^2^J* = 33 Hz), 134.5, 132.7, 132.4, 131.2, 129.7, 128.7, 128.5, 127.5, 126.0, 124.0, 122.3, 121.1 (q, *^1^J* = 275 Hz), 120.6, 119.1, 111.6, 68.9, 58.8, 57.3, 56.5, 52.8, 48.1, 41.7, 40.4, 37.9, 35.6, 26.5, 15.9 ppm.

### Synthesis of 6-hydroxy-N-(4’-((3-(((S)-1-((2S,4R)-4-hydroxy-2-((4-(4-methylthiazol-5- yl)benzyl)carbamoyl)pyrrolidin-1-yl)-3,3-dimethyl-1-oxobutan-2-yl)amino)-3- oxopropyl)carbamoyl)-4-(4-methylpiperazin-1-yl)-[1,1’-biphenyl]-3-yl)-4- (trifluoromethyl)nicotinamide (8e)

The reaction was carried out as described in General Procedure A to give 26.8 mg, 18.6 µmol, 43% of a white solid as TFA salt (1:4/ product: TFA.) MALDI: (calculated) [M+Na^+^] 1006.39 g/mol, (found) [M+Na^+^] 1006.42 g/mol. HRMS: (calculated) [M+Na^+^] 1006.3867 g/mol, (found): [M+Na^+^] 1006.3870 g/mol. HPLC: RT = 11.2 min (254 nm, 100%). ^1^H NMR (600 MHz, DMSO) δ = 10.24 (s, 1H), 9.92 (s, 1H), 8.97 (s, 1H), 8.56 (t, *^3^J* = 5.5 Hz, 1H), 8.50 - 8.48 (m, 1H), 8.34 (s, 1H), 8.16 (s, 1H), 8.01 (d, *^3^J* = 9.2 Hz, 1H), 7.94 (d, *^3^J* = 8.3 Hz, 2H), 7.67 (d, *^3^J* = 7.8 Hz, 2H), 7.59 (d, *^3^J* = 8.5 Hz, 1H), 7.40 (dd, *^2^J* = 22.6 Hz, *^3^*J = 7.6 Hz, 4H), 7.34 (d, *^3^J* = 8.3 Hz, 1H), 7.31 (s, 1H), 4.57 (d, *^3^J* = 9.2 Hz, 1H), 4.47 – 4.39 (m, 2H), 4.36 (s, 1H), 4.22 (dd, *^3^J* = 15.9 Hz, *^4^J* = 5.3 Hz, 1H), 3.72 – 3.64 (m, 2H), 3.59 – 3.52 (m, 2H), 3.49 (dd, *^3^J* = 15.9 Hz, *^4^J* = 7.6 Hz, 2H), 3.31 (d, *^3^J* = 11.4 Hz, 2H), 3.06 (s,4H), 2.86 (s, 3H), 2.62 – 2.54 (m, 2H), 2.44 (s, 3H), 2.09 – 1.99 (m, 1H), 1.95 – 1.87 (m, 1H), 0.93 (s, 9H) ppm. ^13^C NMR (151 MHz, DMSO) δ = 171.9, 170.4, 169.6, 165.8, 163.1, 161.1, 151.5, 147.6, 142.9, 142.0, 139.5, 139.0, 138.5 (q, *^2^J* = 32 Hz), 135.3, 133.3, 132.6, 131.2, 129.6, 128.6, 127.9, 127.4, 126.1, 124.0, 122.4, 122.1 (q, *^1^J* = 279 Hz), 121.0, 118.9, 111.7, 68.9, 58.7, 56.5, 56.4, 52.9, 48.1, 42.4, 41.7, 38.0, 36.3, 35.3, 34.9, 26.4, 15.9 ppm.

### Synthesis of 6-hydroxy-N-(4’-((4-(((S)-1-((2S,4R)-4-hydroxy-2-((4-(4-methylthiazol-5- yl)benzyl)carbamoyl)pyrrolidin-1-yl)-3,3-dimethyl-1-oxobutan-2-yl)amino)-4-oxobutyl)carbamoyl)- 4-(4-methylpiperazin-1-yl)-[1,1’-biphenyl]-3-yl)-4-(trifluoromethyl)nicotinamide (8f)

The reaction was carried out as described in General Procedure A to give 14.7 mg, 14.7 µmol, 41% of a white solid. ESI: (calculated) [M+H^+^] 998.42 g/mol, (found) [M+H^+^] 998.35 g/mol (70). MALDI: (calculated): [M+Na^+^] 1020.40 g/mol, (found) [M+Na^+^] 1020.34 g/mol. HRMS: (calculated) [M+Na^+^] 1020.4024 g/mol, (found) [M+Na^+^] 1020.4010 g/mol. HPLC: RT = 11.3 min (254 nm, 100%). ^1^H NMR (600 MHz, DMSO) δ = 9.82 (s, 1H), 9.58 (s, 1H), 8.97 (s, 1H), 8.55 (t, *^3^J* = 6.0 Hz, 1H), 8.52 (t, *^3^J* = 5.5 Hz, 1H), 8.21 (s, 1H), 8.03 (s, 1H), 7.95 (d, *^3^J* = 8.2 Hz, 3H), 7.69 (d, *^3^J* = 8.3 Hz, 2H), 7.56 (dd, *^3^J* = 8.3 Hz, *^4^J* = 1.9 Hz, 1H), 7.40 (dd, *^2^J* = 22.3 Hz, *^3^J* = 8.1 Hz, 4H), 7.32 (d, *^3^J* = 8.4 Hz, 1H), 6.84 (s, 1H), 4.56 (d, *^3^J* = 9.3 Hz, 1H), 4.43 (dd*^, 3^J* = 14.8 Hz, *^4^J* = 6.9 Hz, 2H), 4.36 (s, 1H), 4.22 (dd, *^3^J* = 15.8 Hz, *^4^J* = 5.4 Hz, 1H), 3.71 - 3.63 (m, 2H), 3.52 (d, *^3^J* = 11.2 Hz, 2H), 3.28 (d, *^3^J* = 7.4 Hz, 4H), 3.22 - 3.18 (m, 2H), 3.05 - 3.01 (m, 2H), 2.87 (s, 3H), 2.44 (s, 3H), 2.37 – 2.30 (m, 1H), 2.25 - 2.20 (m, 1H), 2.05 - 2.00 (m, 1H), 1.95 – 1.87 (m, 1H), 1.81 – 1.72 (m, 2H), 0.95 (s, 9H) ppm. ^13^C NMR (201 MHz, DMSO) δ = 171.9, 171.8, 169.6, 165.8, 162.8, 161.0, 151.4, 147.7, 144.8, 142.0, 139.5, 139.1, 138.7 (q, *^2^J* = 33 Hz), 134.3, 133.2, 132.2, 131.1, 129.6, 128.6, 127.9, 127.4, 126.0, 124.0, 122.4, 122.0 (q, *^1^J* = 276 Hz), 120.4, 119.0, 111.6, 68.9, 58.5, 56.2, 56.1, 52.6, 48.0, 42.1, 41.4, 38.7, 37.7, 35.1, 32.4, 26.2, 25.4, 15.6 ppm.

### Synthesis of 6-hydroxy-*N*-(4’-((5-(((*S*)-1-((2*S*,4*R*)-4-hydroxy-2-((4-(4-methylthiazol-5- yl)benzyl)carbamoyl)pyrrolidin-1-yl)-3,3-dimethyl-1-oxobutan-2-yl)amino)-5- oxopentyl)carbamoyl)-4-(4-methylpiperazin-1-yl)-[1,1’-biphenyl]-3-yl)-4- (trifluoromethyl)nicotinamide (8g)

The reaction was carried out as described in General Procedure A to give 7.42 mg (7.33 µmol, 23 %) of a white solid. ESI: (calculated) [M+H^+^] 1012.43 g/mol, (found) [M+H^+^] 1012.62 g/mol. HPLC: RT = 11.4 min (254 nm, 100%). HRMS: (calculated) [M+Na^+^] 1034.4180 g/mol, (found) [M+Na^+^] 1034.4171 g/mol. ^1^H NMR (500 MHz, DMSO) δ = 9.77 (s, 1H), 9.57 (s, 1H), 8.97 (s, 1H), 8.55 (t, *^3^J* = 6.1 Hz, 1H), 8.50 (t, *^3^J* = 5.6 Hz, 1H), 8.21 (d, *^4^J* = 1.8 Hz, 1H), 8.03 (s, 1H), 7.94 (d, *^3^J* = 8.4 Hz, 2H), 7.86 (d, *^3^J* = 9.3 Hz, 1H), 7.68 (d, *^3^J* = 8.4 Hz, 2H), 7.55 (dd, *^3^J* = 8.3 Hz, *^4^J* = 2.1 Hz, 1H), 7.42 −7.39 (m, 4H), 7.32 (d, *^3^J* = 8.4 Hz, 1H), 6.84 (s, 1H), 5.12 (s, 1H), 4.54 (d, *^3^J* = 9.4 Hz, 1H), 4.48 – 4.38 (m, 2H), 4.35 (s, 1H), 4.21 (dd, *^3^J* = 15.9 Hz, *^4^J* = 5.5 Hz, 1H), 3.70 – 3.63 (m, 2H), 3.55 - 3.46 (m, 2H), 3.29 – 3.23 (m, 4H), 3.07 - 3.02 (m, 2H), 2.86 (s, 3H), 2.53 – 2.50 (m, 2H), 2.44 (s, 3H), 2.35 – 2.26 (m, 1H), 2.19 - 2.14 (m, 1H), 2.09 – 1.99 (m, 1H), 1.93 - 1.88 (m, 1H), 1.60 – 1.47 (m, 4H), 0.94 (s, 9H) ppm. ^13^C NMR (126 MHz, DMSO) δ = 172.0, 172.0, 169.7, 165.8, 163.2, 151.4, 147.7, 144.6, 142.1, 139.5, 139.1, 138.1 (d, *^2^J* = 32 Hz), 134.4, 133.2, 132.5, 131.2, 129.6, 128.6, 127.9, 127.4, 126.0, 123.7, 121.8, 122.3 (q, *^1^J* = 276 Hz), 120.5, 117.7, 111.7, 68.9, 58.7, 56.3, 56.3, 54.8, 51.1, 45.8, 41.6, 38.7, 37.9, 35.2, 34.7, 28.9, 26.4, 23.1, 15.9 ppm.

### Synthesis of 6-hydroxy-*N*-(4’-((6-(((*S*)-1-((2*S*,4*R*)-4-hydroxy-2-((4-(4-methylthiazol-5- yl)benzyl)carbamoyl)pyrrolidin-1-yl)-3,3-dimethyl-1-oxobutan-2-yl)amino)-6-oxohexyl)carbamoyl)- 4-(4-methylpiperazin-1-yl)-[1,1’-biphenyl]-3-yl)-4-(trifluoromethyl)nicotinamide (8h)

The reaction was carried out as described in General Procedure A to give 15.5 mg (11.3 µmol, 26 %) of a white solid as TFA salt (1:3/ product: TFA.) MALDI: (calculated) [M+Na^+^] 1048.43 g/mol, [M+H^+^] 1026.45 g/mol, (found) [M+Na^+^] 1048.43 g/mol, [M+H^+^] 1026.44 g/mol. HPLC: RT = 11.4 min (254 nm, 100%). HRMS: (calculated) [M+Na^+^] 1048.4337 g/mol, (found) [M+Na^+^] 1048.4338 g/mol. ^1^H NMR (500 MHz, DMSO) δ = 9.96 (s, 1H), 9.59 (s, 1H), 8.98 (s, 1H), 8.56 (t, *J* = 6.0 Hz, 1H), 8.48 (t, *J* = 5.6 Hz, 1H), 8.22 (d, *J* = 2.0 Hz, 1H), 8.04 (s, 1H), 7.94 (d, *J* = 8.5 Hz, 2H), 7.84 (d, *J* = 9.3 Hz, 1H), 7.68 (d, *J* = 8.4 Hz, 2H), 7.55 (dd, *J* = 8.3, 2.2 Hz, 1H), 7.45 – 7.36 (m, 4H), 7.32 (d, *J* = 8.4 Hz, 1H), 6.84 (s, 1H), 4.54 (d, *J* = 9.4 Hz, 1H), 4.47 – 4.38 (m, 2H), 4.35 (s, 1H), 4.22 (dd, *J* = 15.9, 5.4 Hz, 1H), 3.68 - 3.65 (m, 2H), 3.53 (d, *J* = 11.2 Hz, 2H), 3.28 - 3.20 (m, 6H), 3.04 (t, *J* = 11.3 Hz, 2H), 2.87 (s, 3H), 2.44 (s, 3H), 2.31 – 2.22 (m, 1H), 2.17 - 2.10 (m, 1H), 2.05 - 2.01 (m, 1H), 1.93 - 1.87 (m, 1H), 1.56 - 1.50 (m, 4H), 1.35 – 1.25 (m, 2H), 0.93 (s, 9H) ppm. ^13^C NMR (126 MHz, DMSO) δ = 172.1, 172.0, 169.7, 165.7, 163.1, 161.1, 151.5, 147.7, 142.9, 141.9, 139.5, 139.0, 138.5 (q, *^2^J* = 33 Hz), 135.4, 133.5, 132.6, 131.2, 129.6, 128.6, 127.9, 127.4, 126.1, 123.9, 122.3, 122.1 (q, *^1^J* = 276 Hz) 121.0, 118.9, 118.8, 111.6, 68.9, 58.7, 56.4, 56.3, 52.8, 48.1, 42.4, 41.7, 38.0, 35.2, 34.9, 29.0, 26.4, 26.2, 25.3, 15.9 ppm.

### Synthesis of 6-hydroxy-*N*-(4’-((7-(((*S*)-1-((2*S*,4*R*)-4-hydroxy-2-((4-(4-methylthiazol-5- yl)benzyl)carbamoyl)pyrrolidin-1-yl)-3,3-dimethyl-1-oxobutan-2-yl)amino)-7- oxoheptyl)carbamoyl)-4-(4-methylpiperazin-1-yl)-[1,1’-biphenyl]-3-yl)-4- (trifluoromethyl)nicotinamide (8i)

The reaction was carried out as described in General Procedure A to give 4.67 mg, 4.49 µmol, 14 % of a yellow solid (TFA salt with a stoichiometry of 1:4/ product: TFA). MALDI: (calculated) [M+H^+^] 1040.21 g/mol, (found) [M+H^+^] 1040.25 g/mol. HPLC: RT = 11.6 min (254 nm, 98%). HRMS: (calculated) [M+Na^+^] 1062.4494 g/mol, (found) [M+Na^+^] 1062.4483 g/mol. ^1^H NMR (500 MHz, DMSO) δ = 12.55 (s, 1H), 9.52 (s, 1H), 8.97 (s, 1H), 8.55 (t, *^3^J* = 6.0 Hz, 1H), 8.46 (t, *^3^J* = 5.6 Hz, 1H), 8.17 (s, 1H), 8.01 (s, 1H), 7.93 (d, *^3^J* = 8.4 Hz, 2H), 7.84 (d, *^3^J* = 9.4 Hz, 1H), 7.68 (d, *^3^J* = 8.4 Hz, 2H), 7.54 (dd, *^3^J* = 8.3 Hz, *^4^*J = 2.1 Hz, 1H), 7.40 (q, *^3^J* = 8.3 Hz, 4H), 7.30 (d, *^3^J* = 8.4 Hz, 1H), 6.83 (s, 1H), 5.11 (d, *^3^J* = 3.6 Hz, 1H), 4.54 (d, ^3^*J* = 9.4 Hz, 1H), 4.49 – 4.38 (m, 2H), 4.35 (s, 1H), 4.21 (dd, *^2^J* = 16.0 Hz, *^3^*J = 5.4 Hz, 1H), 3.72 – 3.58 (m, 2H), 3.26 (dd, *^3^J* = 13.3 Hz, *^4^*J = 6.7 Hz, 4H), 3.05 (s, 4H), 2.44 (s, 3H), 2.32 – 2.28 (m, 1H), 2.19 – 2.07 (m, 1H), 2.07 – 1.98 (m, 1H), 1.93 - 1.88 (m, 1H), 1.52 - 1.47 (m, 4H), 1.31 - 1.23 (m, 3H), 0.93 (s, 9H) ppm. ^13^C NMR (126 MHz, DMSO) δ = 172.1, 171.9, 169.7, 165.7, 163.1, 161.1, 151.4, 147.7, 142.9, 141.8, 139.5, 139.1, 138.5 (q, *^2^J* = 32 Hz), 135.4, 133.5, 132.6, 131.2, 129.6, 128.6, 127.9, 127.4, 126.1, 123.9, 123.1, 122.3, 122.1 (q, *^1^J* = 273 Hz), 121.0, 119.0, 111.6, 68.9, 58.7, 56.3, 56.3, 52.8, 48.1, 42.3, 41.6, 38.4, 38.0, 35.2, 34.8, 29.1, 26.4, 25.4, 15.9 ppm.

### Synthesis of 6-hydroxy-N-(4’-((4-(2-(((S)-1-((2S,4R)-4-hydroxy-2-((4-(4-methylthiazol-5- yl)benzyl)carbamoyl)pyrrolidin-1-yl)-3,3-dimethyl-1-oxobutan-2-yl)amino)-2- oxoethyl)benzyl)carbamoyl)-4-(4-methylpiperazin-1-yl)-[1,1’-biphenyl]-3-yl)-4- (trifluoromethyl)nicotinamide (8j)

The reaction was carried out as described in General Procedure A. The gained product was then dissolved in ethyl acetate and a solution of saturated NaHCO_3_ and saturated NaCl solution and extracted with ethyl acetate. The combined organic phases were dried over MgSO_4_ and the solvent was removed under reduced pressure to give 14.4 mg, 13.6 µmol, 30% of a white solid. MALDI: (calculated) [M+H^+^] 1060.44 g/mol, (found) [M+H^+^] 1060.17 g/mol. HRMS: (calculated) [M+Na^+^] 1082.4180 g/mol, (found) [M+Na^+^] 1082.4171 g/mol. HPLC: RT = 11.6 min (254 nm, 100%). ^1^H NMR (500 MHz, DMSO) δ = 9.47 (s, 1H), 9.04 (t, *^3^J* = 6.0 Hz, 1H), 8.98 (s, 1H), 8.56 (t, *^3^J* = 6.1 Hz, 1H), 8.12 (d, *^4^J* = 2.1 Hz, 1H), 8.09 (d, *^3^J* = 9.3 Hz, 1H), 8.01 – 7.93 (m, 3H), 7.69 (d, *^3^J* = 8.5 Hz, 2H), 7.53 (dd, *^3^J* = 8.4 Hz, *^4^J* = 2.2 Hz, 1H), 7.40 (q, *^3^J* = 8.4 Hz, 4H), 7.27 (d, *^3^J* = 8.4 Hz, 1H), 7.24 (s, 4H), 6.82 (s, 1H), 5.11 (d, *^3^J* = 3.1 Hz, 1H), 4.51 (d, *^3^J* = 9.4 Hz, 1H), 4.47 (d, *^3^J* = 5.8 Hz, 2H), 4.45 – 4.39 (m, 2H), 4.33 (s, 1H), 4.22 (dd, *^3^J* = 15.9 Hz, *^4^J* = 5.5 Hz, 1H), 3.71 – 3.58 (m, 3H), 3.44 (d, *^3^J* = 13.9 Hz, 2H), 2.91 (t, *^3^J* = 4.3 Hz, 4H), 2.50 (s, 4H), 2.44 (s, 3H), 2.24 (s, 3H), 2.07 – 1.97 (m, 1H), 1.92 - 1.86 (m, 1H), 0.92 (s, 9H) ppm. ^13^C NMR (126 MHz, DMSO) δ = 171.9, 170.0, 169.5, 165.8, 162.8, 161.1, 151.4, 147.72, 145.0, 142.3, 139.5, 139.2, 138.5 (q, *^2^J* = 32 Hz), 137.6, 135.1, 134.2, 132.9, 132.2, 131.2, 129.6, 129.0, 128.6, 128.0, 127.4, 127.0, 126.1, 124.0, 122.4, 122.0 (d, *^1^J* = 275 Hz), 120.4, 119.0, 111.6, 68.9, 58.7, 56.5, 56.4, 54.7, 51.0, 45.7, 42.4, 41.7, 41.5, 37.9, 35.4, 26.3, 15.9 ppm.

### Synthesis of N-(4’-((2-(2-(4-((2R,3S,4R,5S)-3-(3-chloro-2-fluorophenyl)-4-(4-chloro-2-fluorophenyl)- 4-cyano-5-neopentylpyrrolidine-2-carboxamido)-3-methoxybenzamido)ethoxy)ethyl)carbamoyl)-4- (4-methylpiperazin-1-yl)-[1,1’-biphenyl]-3-yl)-6-hydroxy-4-(trifluoromethyl)nicotinamide (9a)

15.2 mg (21 µmol, 1.05 eq) Intermediate **6a** was dissolved in 2 mL TFA/ CH_2_Cl_2_ and stirred for 1h at rt. Excess solvent was removed under reduced pressure. 13 mg (21 µmol, 1.0 eq) Idasanutlin, 9.6 mg (25µmol, 1.2 eq) HATU and 7 µL (42 µmol, 2.0 eq) DIEA were dissolved in 1 mL DMF and stirred for 15 min at rt. Then, a solution of deprotected **6a** and 73µL (420 µmol, 10 eq) DIEA in 1 mL DMF were added to the solution and stirred for 12 h at rt. The reaction mixture was quenched with 2 mL water and 2 mL saturated NaHCO_3_, then the reaction was extracted 3x with EA. The organic phase was dried over MgSO_4_, filtered and the solvent was removed under reduced pressure. The crude product was purified by HPLC. The gained product was then dissolved in ethyl acetate and a solution of saturated NaHCO_3_ and saturated NaCl solution and extracted with ethyl acetate. The combined organic phases were dried over MgSO_4_ and the solvent was removed under reduced pressure to give 3.0 mg, 2.5 µmol, 12% of a clear oil. MALDI: (calculated) [M+H^+^] 1184.49 g/mol, (found) [M+H^+^] 1184.40 g/mol. HRMS: (calculated) [M+Na^+^] 1206.3805 g/mol, (found) [M+Na^+^] 1206.3821 g/mol. HPLC: RT = 14.4 min (254 nm, 98%). ^1^H NMR (400 MHz, DMSO) δ = 10.45 (s, 1H), 10.40 (s, 1H), 9.45 (s, 1H), 8.53 (t, ^3^*J* = 5.5 Hz, 1H), 8.48 (t, ^3^*J* = 5.5 Hz, 1H), 8.35 (s, 1H), 8.32 (d, ^3^*J* = 8.5 Hz, 2H), 8.13 (d, ^4^*J* = 1.9 Hz, 1H), 7.92 (d, ^3^*J* = 8.4 Hz, 2H), 7.73 (t, ^3^*J* = 7.3 Hz, 2H), 7.67 (d, ^3^*J* = 8.4 Hz, 2H), 7.60 - 7.56 (m, 4H), 7.53 - 7.50 (m, 3H), 7.44 – 7.38 (m, 1H), 7.38 – 7.31 (m, 4H), 7.25 (d, ^3^*J* = 8.4 Hz, 1H), 6.81 (s, 1H), 4.65 – 4.54 (m, 4H), 4.40 - 4.33 (m, 2H), 3.99 - 3.95 (m, 2H), 3.92 (s, 3H), 3.90 (s, 3H), 3.60 - 3.57 (m, 5H), 3.48 - 3.43 (m, 8H), 2.92 – 2.88 (m, 4H), 1.69 - 1.60 (m, 2H), 1.39 - 1.36 (m, 1H), 1.28 - 1.24 (m, 2H), 0.96 (s, 9H) ppm. Contains rotameres.

### Synthesis of N-(4’-((1-(4-((2R,3S,4R,5S)-3-(3-chloro-2-fluorophenyl)-4-(4-chloro-2-fluorophenyl)-4- cyano-5-neopentylpyrrolidine-2-carboxamido)-3-methoxyphenyl)-1-oxo-5,8,11,14,17,20,23- heptaoxa-2-azapentacosan-25-yl)carbamoyl)-4-(4-methylpiperazin-1-yl)-[1,1’-biphenyl]-3-yl)-6- hydroxy-4-(trifluoromethyl)nicotinamide (9b)

8.9 mg (9 µmol, 1.0 eq) Intermediate **6b** was dissolved in 2 mL TFA/ CH_2_Cl_2_ and stirred for 1h at rt. Excess solvent was removed under reduced pressure. 5.5 mg (9 µmol, 1.0 eq) Idasanutlin, 4 mg (11 µmol, 1.2 eq) HATU and 3 µL (18 µmol, 2.0 eq) DIEA were dissolved in 1 mL DMF and stirred for 15 min at rt. Then, a solution of deprotected **6b** and 31 µL (180 µmol, 10 eq) DIEA in 1 mL DMF were added to the solution and stirred for 12 h at rt. The reaction mixture was quenched with 2 mL water and 2 mL saturated NaHCO_3_, then the reaction was extracted 3x with EA. The organic phase was dried over MgSO_4_, filtered and the solvent was removed under reduced pressure. The crude product was purified by HPLC. The gained product was then dissolved in ethyl acetate and a solution of saturated NaHCO_3_ and saturated NaCl solution and extracted with ethyl acetate. The combined organic phases were dried over MgSO_4_ and the solvent was removed under reduced pressure to give 3.3 mg, 2.2 µmol, 25% of a clear oil. MALDI: (calculated) [M+H^+^] 1448.56 g/mol, (found) [M+H^+^] 1448.53 g/mol. HRMS: (calculated) [M+Na^+^] 1470.5378 g/mol, (found) [M+Na^+^] 1470.5392 g/mol. HPLC: RT = 14.6 min (254 nm, 96%). ^1^H NMR (400 MHz, DMSO) δ = 10.47 (s, 1H), 10.40 (s, 1H), 9.46 (s, 1H), 8.53 (t, *^3^J* = 5.5 Hz, 1H), 8.48 (t, *^3^J* = 5.4 Hz, 1H), 8.36 (d, *^3^J* = 8.8 Hz, 1H), 8.32 (d, *^3^J* = 8.4 Hz, 2H), 8.12 (d, *^4^J* = 2.1 Hz, 1H), 7.92 (d, *^3^J* = 8.5 Hz, 2H), 7.73 (t, *^3^J* = 7.2 Hz, 2H), 7.67 (d, *^3^J* = 8.5 Hz, 2H), 7.60 - 7.56 (m, 5H), 7.54 – 7.51 (m, 2H), 7.50 - 7.48 (m, 1H), 7.40 (dd, *^3^J* = 8.4 Hz, *^4^J* = 2.4 Hz, 1H), 7.38 – 7.34 (m, 3H), 7.34 – 7.32 (m, 1H), 7.25 (d, *^3^J* = 8.4 Hz, 1H), 6.82 (s, 1H), 4.65 – 4.54 (m, 4H), 4.41 - 4.33 (m, 2H), 3.99 - 3.95 (m, 2H), 3.92 (s, 3H), 3.90 (s, 3H), 3.60 - 3.57 (m, 5H), 3.49 – 3.41 (m, 8H), 2.91 - 2.90 (m, 4H), 1.68 - 1.61 (m, 2H), 1.44 – 1.33 (m, 2H), 0.96 (s, 9H) ppm. Contains rotameres.

### Synthesis of N-(4’-((4-((4-((2R,3S,4R,5S)-3-(3-chloro-2-fluorophenyl)-4-(4-chloro-2-fluorophenyl)-4- cyano-5-neopentylpyrrolidine-2-carboxamido)-3-methoxybenzamido)methyl)benzyl)carbamoyl)-4- (4-methylpiperazin-1-yl)-[1,1’-biphenyl]-3-yl)-6-hydroxy-4-(trifluoromethyl)nicotinamide (9c)

13.5 mg (24 µmol, 1.0 eq) Intermediate **6c** was dissolved in 2 mL TFA/ CH_2_Cl_2_ and stirred for 1h at rt. Excess solvent was removed under reduced pressure. 15.7 mg (25 µmol, 1.1 eq) Idasanutlin, 11 mg (29 µmol, 1.2 eq) HATU and 8 µL (49 µmol, 2.0 eq) DIEA were dissolved in 1 mL DMF and stirred for 15 min at rt. Then, a solution of deprotected **6c** and 84 µL (490 µmol, 10 eq) DIEA in 1 mL DMF were added to the solution and stirred for 12 h at rt. The reaction mixture was quenched with 2 mL water and 2 mL saturated NaHCO_3_, then the reaction was extracted 3x with EA. The organic phase was dried over MgSO_4_, filtered and the solvent was removed under reduced pressure. The crude product was purified by HPLC. The gained product was then dissolved in ethyl acetate and a solution of saturated NaHCO_3_ and saturated NaCl solution and extracted with ethyl acetate. The combined organic phases were dried over MgSO_4_ and the solvent was removed under reduced pressure to give 3.7 mg, 3.0 µmol, 13% of a clear oil. MALDI: (calculated) [M+K^+^] 1254.36 g/mol, (found) [M+K^+^] 1254.32 g/mol. HRMS: (calculated) [M+Na^+^] 1238.3856 g/mol, (found) [M+Na^+^] 1238.3894 g/mol. HPLC: RT = 14.7 min (254 nm, 98%). ^1^H NMR (250 MHz, DMSO) δ = 10.40 (s, 1H), 9.13 (s, 1H), 9.04 (s, 1H), 8.96 (s, 1H), 8.49 (s, 3H), 8.44 (s, 1H), 8.31 (d, *^3^J* = 9.1 Hz, 2H), 8.20 (d, *^3^J* = 11.9 Hz, 1H), 7.98 (d, *^3^J* = 7.9 Hz, 2H), 7.69 (d, *^3^J* = 7.4 Hz, 2H), 7.61 (s, 2H), 7.57 – 7.52 (m, 2H), 7.52 – 7.47 (m, 1H), 7.39 (s, 1H), 7.36 (s, 2H), 7.29 (s, 4H), 6.52 (s, 1H), 6.13 (s, 1H), 4.59 - 4.58 (m, 2H), 4.51 – 4.41 (m, 4H), 3.91 (s, 4H), 3.57 - 3.50 (m, 6H), 3.47 (d, *^3^J* = 4.8 Hz, 2H), 3.11 (d, *^3^J* = 5.1 Hz, 2H), 3.03 - 2.99 (m, 2H), 2.44 – 2.37 (m, 1H), 0.97 (s, 9H) ppm. Contains rotameres. Synthesis of 6-hydroxy-N-(4’-((5-(((S)-1-((2S,4S)-4-hydroxy-2-((4-(4-methylthiazol-5- yl)benzyl)carbamoyl)pyrrolidin-1-yl)-3,3-dimethyl-1-oxobutan-2-yl)amino)-5- oxopentyl)carbamoyl)-4-(4-methylpiperazin-1-yl)-[1,1’-biphenyl]-3-yl)-4- (trifluoromethyl)nicotinamide (20). The reaction was carried out as described in General Procedure A with intermediate 6 and tert-butyl (5-(((S)-1-((2S,4S)-4-hydroxy-2-((4-(4-methylthiazol-5- yl)benzyl)carbamoyl)pyrrolidin-1-yl)-3,3-dimethyl-1-oxobutan-2-yl)amino)-5-oxopentyl)carbamate (L15). The gained product was then dissolved in ethyl acetate and a solution of saturated NaHCO_3_ and saturated NaCl solution and extracted with ethyl acetate. The combined organic phases were dried over MgSO_4_ and the solvent was removed under reduced pressure to give 9 mg, 8.9 µmol, 18% of a white oil. MALDI: (calculated) [M-CH_3_-tBu+2H^+^] 941.35 g/mol, (found) [M-CH_3_-tBu+2H^+^] 941.45 g/mol. HRMS: (calculated) [M+Na^+^] 1034.4180 g/mol, (found) [M+Na^+^] 1034.4169 g/mol. HPLC: RT = 11.4 min (254 nm, 100%). ^1^H NMR (250 MHz, MeOD) δ = 9.03 (s, 1H), 8.26 (d, *^4^J* = 1.9 Hz, 1H), 8.03 (s, 1H), 7.91 (d, *^3^J* = 8.3 Hz, 2H), 7.71 (d, *^3^J* = 8.3 Hz, 2H), 7.56 (dd, *^3^J* = 8.5 Hz, *^4^J* = 2.0 Hz, 1H), 7.44 (dd, *^2^J* = 15.1 Hz, *^3^J* = 7.8Hz, 4H), 7.37 (d, *^4^J* = 2.1 Hz, 1H), 6.94 (s, 1H), 4.63 (s, 1H), 4.56 (s, 1H), 4.50 (s, 2H), 4.35 (d, *^3^J* = 15.7 Hz, 1H), 3.91 (d, *^3^J* = 10.7 Hz, 1H), 3.80 (dd, *^3^J* = 10.9 Hz, *^4^J* = 3.6 Hz, 1H), 3.64 - 3.60 (m, 2H), 3.44 - 3.39 (m, 2H), 3.36 – 3.24 (m, 4H), 3.20 - 3.15 (m, 2H), 2.97 (s, 3H), 2.47 (s, 3H), 2.39 - 2.33 (m, 2H), 2.28 – 2.15 (m, 1H), 2.13 - 2.08 (m, 1H), 1.79 – 1.59 (m, 4H), 1.04 (s, 9H) ppm.

### Synthesis of 2-Bromo-6,7-dihydro-5H-pyrrolo[1,2-a]imidazole (11)

This reaction was performed in two equal batches. 1.00 g (7.03 mmol, 1.0 eq) Piracetam **10** and 4.00 g (14.0 mmol, 2.0 eq) POBr_3_ were dissolved in 20 ml acetonitrile and heated in a microwave under stirring for 80 min at 5 W to 70 °C. Solid POBr_3_ which had formed at the top of the vial was brought back into the reaction mixture with a spatula and the reaction mixture was heated in the microwave for another 45 min. After cooling to rt, the batches were combined and quenched with 35 ml of water. K_2_CO_3_ was added until no gas formation was observed and the solvent was removed under reduced pressure. The aqueous mixture was extracted with dichloromethane (5×35 ml), washed with brine and dried with MgSO_4_. The solvent was removed under reduced pressure. Purification by flash column chromatography (CH_2_Cl_2_/MeOH) to give 2.16 g, 5.76 mmol, 82% of a colorless solid. R_f_ (10% MeOH/ CH_2_Cl_2_) 0.70. ESI: (calculated) [M+H^+^] 189.99 g/mol, (found) [M+H^+^] 189.09 g/mol; m.p. 95.7°C. ^1^H NMR (600 MHz, CD_3_OD) δ = 7.05 (s, 1H), 4.02 (t, *^3^J* = 7.20 Hz, 2H), 2.82 (t, *^3^J* = 7.40 Hz, 2H), 2.60-2.54 (m, 2H) ppm. ^13^C NMR (125.8 MHz, CD_3_OD) δ = 155.4, 116.6, 116.0, 46.5, 26.4, 24.1 ppm.

### Synthesis of 5-(6,7-Dihydro-5H-pyrrolo[1,2-a]imidazol-2-yl)-2-methoxybenzonitrile (12)

A solution of 538 mg (13.5 mmol, 5.2 eq) NaOH and 693 mg (3.92 mmol, 1.5 eq) (3-cyano-4- methoxyphenyl)boronic acid in 24 ml tetrahydrofuran/water (3/1) was stirred at 50 °C for 5 min. Bromide **11** (480 mg, 2.57 mmol, 1.0 eq) and 113 mg (0.13 mmol. 0.1 eq) XPhos PdG3 catalyst were added and the reaction mixture was stirred for 21 h at 80 °C. The two batches were combined and the organic solvent was removed under reduced pressure. The remaining aqueous layer was extracted with dichloromethane (3x). The combined organic layers were washed with brine, dried with MgSO_4_ and the solvent was removed under reduced pressure. Purification by flash column chromatography (EA) to give 386 mg, 1.62 mmol, 63% of a colorless solid. R_f_ (EA) 0.43. ESI: (calculated) [M+H^+^] 240.12 g/mol, (found) [M+H^+^] 240.05 g/mol. M.p. 160.9 °C. ^1^H NMR (600 MHz, CDCl_3_) δ = 7.99 (dd, *^3^J* = 8.82 Hz, *^4^J* = 1.90 Hz, 1H), 7.87 (d, *^4^J* = 1.90 Hz, 1H), 7.13 (s, 1H), 6.98 (d, *^3^J* = 8.82 Hz, 1H), 4.05 (t, *^3^J* = 6.98 Hz, 2H), 3.94 (s, 3H), 2.97 (t, *^3^J* = 7.48 Hz, 2H), 2.68-2.62 (m, 2H) ppm. ^13^C NMR (125.8 MHz, CDCl_3_) δ = 160.1, 155.1, 143.6, 130.8, 129.8, 127.8, 116.6, 111.7, 110.3, 102.0, 56.3, 45.3, 26.2, 23.2 ppm.

### Synthesis of (5-(6,7-dihydro-5H-pyrrolo[1,2-a]imidazol-2-yl)-2-methoxyphenyl)methanamine (13)

16.1 mL (16.1 mmol, 5.0 eq) LiAlH_4_ (1 M in THF) was added dropwise to a colorless solution of 772 mg (3.23 mmol, 1.0 eq) nitrile **12** in 32 ml tetrahydrofuran via a syringe. The reaction mixture was stirred at 60 °C for 2.5 h, got orange and was stirred over night at room temperature. The reaction was quenched with an aqueous solution of 1 M NaOH (35 ml), filtered and the solvent was removed under reduced pressure. The remaining aqueous phase was extracted with dichloromethane (2x). The combined organic layers were washed with brine and dried with MgSO_4_. The solvent was removed under reduced pressure. 691 mg, 2.84 mmol, 88% of the title compound **12** was isolated as sticky yellow solid and used without further purification. ESI (calculated) [M+H^+^] 244.15 g/mol, (found) [M+H^+^] 244.22 g/mol. ^1^H NMR (600 MHz, CDCl_3_) δ = 7.59 (t, *^4^J* = 1.28 Hz, 1H), 7.57 (dd, *^3^J* = 8.26 Hz, *^4^J* = 1.28 Hz, 1H), 7.07 (s, 1H), 6.84 (d, *^3^J* = 8.26 Hz, 1H), 3-97 (t, *^3^J* = 6.67 Hz, 2H), 3.84 (s, 3H), 3.81 (s, 2H), 2.89 (t, *^3^J* = 7.56 Hz, 2H), 2.58 (quin, *^3^J* = 7.23 Hz, 2H), 1.92 (br, s) ppm. The spectrum contains minor impurities. ^13^C NMR (125.8 MHz, DMSO) δ = 155.8, 154.5, 145.2, 128.3, 127.9, 125.4, 124.5, 111.0, 110.5, 55.9, 44.7, 39.6, 26.1, 22.3 ppm.

### Synthesis of 2-(3,4-Dichlorophenyl)-N-(5-(6,7-dihydro-5H-pyrrolo[1,2-a]imidazol-2-yl)-2- methoxybenzyl)acetamide (14)

118 mg EDC hydrochloride (0.647 mmol, 1.5 eq) was added to a solution of the amine **13** (100 mg, 0.411 mmol, 1.0 eq), 126 mg (0.617 mmol, 1.5 eq) 2-(3,4- Dichlorophenyl)acetic acid, 94 mg (0.65 μmol, 1.5 eq) HOBT monohydrate and 0.11 mL (0.63 mmol, 1.5 eq) DIPEA in 1 ml dimethylformamide at 0 °C. The reaction mixture was stirred at room temperature overnight. The reaction was quenched with water and extracted with ethylacetate (4x). The combined organic layers were dried with MgSO_4_ and the solvent was removed under reduced pressure. Purification by flash column chromatography (CH_2_Cl_2_/MeOH) yielded the title compound **14** 110 mg, 0.259 mmol, 62% of a colorless solid. R_f_ (10% MeOH/CH_2_Cl_2_) 0.59. ESI: (calculated) [M+H^+^] 430.11 g/mol, (found) [M+H^+^] 430.09 g/mol. HPLC: RT = 12.4 min (254 nm, 100%). ^1^H NMR (600 MHz, CDCl_3_) δ = 7.64 (dd, *^3^J* = 8.44 Hz, *^4^J* = 1.79 Hz, 1H), 7.53 (s, 1H), 7.37 (s, 1H), 7.36 (d, *^3^J* = 5.57 Hz, 1H), 7.11 (dd, *^3^J* = 8.20 Hz, *^4^J* = 1.51 Hz, 1H), 7.01 (s, 1H), 6.23 (t, *^3^J* = 4.60 Hz, 1H), 4.42 (d, *^3^J* = 5.60 Hz, 2H), 3.99 (t, *^3^J* = 7.14 Hz, 2H), 3.76 (s, 3H), 7.49 (s, 2H), 2.90 (t, *^3^J* = 7.46 Hz, 2H), 2.60 (quin, *^3^J* = 7.46, 2H). ^13^C NMR (125.8 MHz, CDCl_3_) δ = 169.6, 156.6, 154.4, 145.0, 135.7, 132.6, 131.4, 131.2, 130.6, 129.0, 126.7, 126.1, 126.0, 125.2, 110.6, 109.7, 55.5, 45.3, 42.7, 39.7, 26.1, 23.2 ppm.

### Synthesis of 2-(3,4-dichlorophenyl)-N-(5-(6,7-dihydro-5H-pyrrolo[1,2-a]imidazol-2-yl)-2- hydroxybenzyl) acetamide (15)

1.22 mL (1.22 mmol, 5.0 eq) of a solution 1M BBr_3_ in dichloromethane was added to a solution of 105 mg (0.244 mmol, 1.0 eq) the protected phenol **14** in 15 mL dichloromethane at −78 °C via syringe. After 30 min, the reaction mixture was allowed to warm to rt and turned orange while stirred for another 21 h. The reaction was quenched by adding 6.5 ml of an aqueous solution of 1 M NaOH and stirred for 4 h. The organic layer was separated and the remaining aqueous layer was extracted with dichloromethane (2x). The combined organic layers were dried with MgSO_4_ and the solvent was removed under reduced pressure to give 101 mg, 0.242 mmol, 99% of a colorless solid without further purification. R_f_ (10% MeOH/CH_2_Cl_2_) 0.41. ESI: (calculated) [M+H^+^] 416.10 g/mol, (found) [M+H^+^] 416.09 g/mol. ^1^H NMR (500 MHz, DMSO) δ = 9.45 (s, 1H), 8.50 (t, *^3^J* = 5.70 Hz, 1H), 7.58-7.55 (m, 2H), 7.42-7.39 (m, 2H), 7.31 (dd, *^3^J* = 8.31 Hz, *^4^J* = 1.85 Hz, 1H), 7.15 (s, 1H), 6.77-6.74 (m, 1H), 4.23-4.20 (m, 2H), 3.94 (t, *^3^J* = 7.01 Hz, 2H), 3.54 (s, 2H), 2.73 (t, *^3^J* = 7.13 Hz), 2.54-2.47 (m, 2H) ppm. ^13^C NMR (125.8 MHz, DMSO) δ = 169.6, 153.8, 153.4, 145.1, 137.6, 131.0, 130.7, 130.3, 129.5, 129.1, 126.4, 124.7, 124.4, 123.8, 115.0, 109.2, 44.3, 41.0, 37.8, 25.6, 22.4 ppm.

### Synthesis of tert-butyl 3-(2-(2-((2-(3,4-dichlorophenyl)acetamido)methyl)-4-(6,7-dihydro-5H- pyrrolo[1,2-a]imidazol-2-yl)phenoxy)ethoxy)propanoate (16a)

The reaction was carried out as described in General Procedure B using tert-butyl 3-(2-hydroxyethoxy)propanoate as commercial available starting material, obtaining the product as TFA salt with unknown stoichiometry as a colorless oil (31 mg). MALDI: (calculated) [M+H^±^] 588.21 g/mol, (found) [M+H^±^] 588.25 g/mol. ^1^H NMR (500 MHz, CDCl_3_) δ = 7.72 (t, *^3^J* = 6.00 Hz, 1H), 7.50 (d, *^4^J* = 2.10 Hz, 1H), 7.45 (d, *^4^J* = 1.98 Hz, 1H), 7.38 (dd, *^3^J* = 8.44 Hz, *^4^J* = 2.10 Hz, 1H), 7.31 (d, *^3^J* = 8.22 Hz, 1H), 7.16 (dd, *^3^J* = 8.22 Hz, *^4^J* = 1.98 Hz, 1H), 7.14 (s, 1H) 6.78 (d, *^3^J* = 8.57 Hz, 1H), 4.41 (d, *^3^J* = 6.00 Hz, 2H), 4.16 (t, *^3^J* = 7.25 Hz, 2H), 4.10-4.07 (m, 2H), 3.80-3.76 (m, 4H), 3.59 (s, 2H), 3.20 (t, *^3^J* = 7.50 Hz, 2H), 2.77-2.70 (m, 2H), 2.50 (t, *^3^J* = 6.12 Hz, 2H), 1.42 (s, 9H) ppm. ^13^C NMR (125.8 MHz, CDCl_3_) δ = 171.2, 171.0, 157.4, 152.0, 139.2, 136.0, 132.3, 131.3, 130.9, 130.44, 129.1, 128.2, 125.9, 119.6, 112.2, 110.9, 81.0, 69.3, 68.1, 67.3, 47.6, 42.1, 38.1, 36.4, 28.2, 25.7, 23.5 ppm.

### Synthesis of tert-butyl 3-(2-(2-(2-((2-(3,4-dichlorophenyl)acetamido)methyl)-4-(6,7-dihydro-5H- pyrrolo[1,2-a]imidazol-2-yl)phenoxy)ethoxy)ethoxy)propanoate (16b)

The reaction was carried out as described in General Procedure B using tert-butyl 3-(2-(2-hydroxyethoxy)ethoxy)propanoate as commercial available starting material, as TFA salt with unknown stoichiometry as a colorless oil (74 mg). MALDI: (calculated) [M+H]^±^ 632.23 g/mol, (found) [M+H]^±^ 632.19 g/mol. ^1^H NMR (500 MHz, CDCl_3_) δ = 7.63 (t, *^3^J* = 6.00 Hz, 1H), 7.56 (d, *^4^J* = 1.91 Hz, 1H), 7.47 (dd, *^3^J* = 8.50 Hz, *^4^J* = 1.91 Hz, 1H), 7.45 (d, *^4^J* = 1.70 Hz, 1H), 7.32 (d, *^3^J* = 8.17 Hz, 1H), 7.17 (dd, *^3^J* = 8.17 Hz, *^4^J* = 1.70 Hz, 1H), 7.12 (s, 1H), 6.86 (d, *^3^J* = 8.50 Hz, 1H), 4.45 (d, *^3^J* = 6.00 Hz, 2H), 4.21 (t, *^3^J* = 7.24 Hz, 2H), 4.16-4.12 (m, 2H), 3.86- 3.82 (m, 2H), 3.79-3.60 (m, 6H), 3.57 (s, 2H), 3.30 (t, *^3^J* = 7.48 Hz, 2H), 2.83-2.75 (m, 2H), 2.48 (t, *^3^J* = 6.42 Hz, 2H), 1.42 (s, 9H) ppm. Spectrum contains CH_2_Cl_2_. ^13^C NMR (125.8 MHz, CDCl_3_): δ = 171.1, 170.9, 157.5, 152.3, 139.7, 136.1, 132.3, 131.3, 130.9, 130.4, 129.1, 128.4, 126.1, 126.01, 119.7, 112.5, 110.6, 80.9, 70.8, 70.4, 69.6, 68.1, 67.0, 53.6 (CH_2_Cl_2_), 47.6, 42.2, 38.8, 36.36, 28.2, 25.8, 23.6 ppm.

### Synthesis of tert-butyl 3-(2-(2-(2-(2-((2-(3,4-dichlorophenyl)acetamido)methyl)-4-(6,7-dihydro-5H- pyrrolo[1,2-a]imidazol-2-yl)phenoxy)ethoxy)ethoxy)ethoxy)propanoate (16c)

The reaction was carried out as described in General Procedure B using tert-butyl 3-(2-(2-(2- hydroxyethoxy)ethoxy)ethoxy)propanoate as commercial available starting material, as TFA salt with unknown stoichiometry as a colorless oil (43 mg). MALDI: (calculated) [M+H^±^] 676.26 g/mol, (found) [M+H^±^] 676.25 g/mol. ^1^H NMR (500 MHz, CDCl_3_) δ = 7.89 (t, *^3^J* = 6.08 Hz, 1H), 7.51 (d, *^4^J* = 2.10 Hz, 1H), 7.45 (d, *^4^J* = 1.99 Hz, 1H), 7.38 (dd, *^3^J* = 8.51 Hz, *^4^J* = 2.10 Hz, 1H), 7.30 (d, *^3^J* = 8.19 Hz, 1H), 7.17 (dd, *^3^J* = 8.19 Hz, *^4^J* = 1.99 Hz, 1H), 7.14 (s, 1H), 6.78 (d, *^3^J* = 8.51 Hz, 1H), 4.41 (d, *^3^J* = 6.08 Hz, 2H), 4.14 (t, *^3^J* = 7.32 Hz, 2H), 4.12-4.09 (m, 2H), 3.84-3.81 (m, 2H), 3.72-3.53 (m, 12H), 3.18 (t, *^3^J* = 7.66 Hz, 2H), 2.75-2.67 (m, 2H), 2.45 (t, *^3^J* = 6.46 Hz, 2H), 1.41 (s, 9H) ppm. ^13^C NMR (125.8 MHz, CDCl_3_) δ = 171.0, 171.0, 157.3, 152.0, 139.1, 136.2, 132.2, 131.2, 130.7, 130.4, 129.1, 128.3, 125.8, 119.7, 112.2, 111.0, 80.8, 70.8, 70.5, 70.5, 70.4, 69.6, 68.1, 67.0, 47.6, 42.1, 38.7, 36.3, 28.2, 25.7, 23.4 ppm.

### Synthesis of tert-butyl 1-(2-((2-(3,4-dichlorophenyl)acetamido)methyl)-4-(6,7-dihydro-5H- pyrrolo[1,2-a]imidazol-2-yl)phenoxy)-3,6,9,12-tetraoxapentadecan-15-oate (16d)

The reaction was carried out as described in General Procedure B using tert-butyl 1-hydroxy-3,6,9,12- tetraoxapentadecan-15-oate as commercial available starting material, as TFA salt with unknown stoichiometry as a colorless oil (44 mg). MALDI: (calculated) [M+H]^±^ 720.28 g/mol, (found) [M+H]^±^ 720.33 g/mol; ^1^H NMR (500 MHz, CDCl_3_) δ = 7.93 (t, *^3^J* = 6.03 Hz, 1H), 7.47 (d, *^4^J* = 2.19 Hz, 1H), 7.41 (d, *^4^J* = 2.02 Hz, 1H), 7.39 (dd, *^3^J* = 8.52 Hz, *^4^J* = 2.19 Hz, 1H), 7.30 (d, *^3^J* = 8.27 Hz, 1H), 7.19 (s, 1H), 7.13 (dd, *^3^J* = 8.27 Hz, *^4^J* = 2.02 Hz, 1H), 6.79 (d, *^3^J* = 8.52 Hz, 1H), 4.41 (d, *^3^J* = 6.03 Hz, 2H), 4.18 (t, *^3^J* = 7.31 Hz, 2H), 4.13 - 4.08 (m, 2H), 3.85-3.81 (m, 2H), 3.76-3.52 (m, 16H), 3.20 (t, *^3^J* = 7.53 Hz, 2H), 2.79-2.71 (m, 2H), 2.46 (t, *^3^J* = 6.45 Hz, 2H), 1.41 (s, 9H) ppm. ^13^C NMR (125.8 MHz, CDCl_3_) δ = 171.6, 171.1, 157.5, 152.0, 139.0, 135.8, 132.3, 131.2, 131.0, 130.5, 129.1, 127.9, 126.2, 126.0, 119.5, 112.4, 111.1, 80.8, 70.7, 70.6, 70.5, 70.4, 70.4, 70.3, 69.5, 68.1, 66.9, 47.7, 41.9, 39.1, 36.3, 28.1, 25.7, 23.4 ppm.

### Synthesis of tert-butyl 1-(2-((2-(3,4-dichlorophenyl)acetamido)methyl)-4-(6,7-dihydro-5H- pyrrolo[1,2-a]imidazol-2-yl)phenoxy)-3,6,9,12,15-pentaoxaoctadecan-18-oate (16e)

The reaction was carried out as described in General Procedure B tert-butyl 1-hydroxy-3,6,9,12,15- pentaoxaoctadecan-18-oate as commercial available starting material, as TFA salt with unknown stoichiometry as a colorless oil (39 mg). MALDI: (calculated) [M+H^±^] 764.31 g/mol, (found) [M+H^±^] 764.36 g/mol. ^1^H NMR (500 MHz, CDCl_3_) δ = 7.92 (t, *^3^J* = 6.09 Hz, 1H), 7.53-7.49 (m, 1H), 7.44 (d, *^4^J* = 1.95 Hz, 1H), 7.42-7.37 (m, 1H), 7.29 (d, *^3^J* = 8.24 Hz, 1H), 7.18-7.15 (m, 2H), 6.81-6.77 (m, 1H), 4.40(d, *^3^J* = 6.09 Hz, 2H), 4.14 (t, *^3^J* = 7.72 Hz, 2H), 4.12-4.08 (m, 2H), 3.84-3.81 (m, 2H), 3.75-3.53 (m, 20H), 3.21-3.15 (m, 2H), 2.74 −2.67 (m, 2H), 2.46 (t, *^3^J* = 6.46 Hz, 2H), 1.43-1.41 (m, 9H) ppm. ^13^C NMR (125.8 MHz, CDCl_3_) δ = 171.1, 171.0, 157.3, 152.0, 139.1, 136.2, 132.1, 131.2, 130.7, 130.4, 129.1, 128.3, 125.8, 125.8, 119.7, 112.2, 110.9, 80.7, 70.8, 70.7, 70.7, 70.6, 70.5, 70.5, 70.5, 70.3, 69.5, 68.1, 66.9, 47.6, 42.0, 38.7, 36.3, 28.2, 25.7, 23.4 ppm.

### Synthesis of tert-butyl 1-(2-((2-(3,4-dichlorophenyl)acetamido)methyl)-4-(6,7-dihydro-5H- pyrrolo[1,2-a]imidazol-2-yl)phenoxy)-3,6,9,12,15,18-hexaoxahenicosan-21-oate (16f)

The reaction was carried out as described in General Procedure B using tert-butyl 1-hydroxy-3,6,9,12,15,18- hexaoxahenicosan-21-oate as commercial available starting material, as TFA salt with unknown stoichiometry as a colorless oil (36 mg). MALDI: (calculated) [M+H^±^] 808.34 g/mol, (found) [M+H^±^] 808.39 g/mol. ^1^H NMR (500 MHz, CDCl_3_) δ = 7.91 (t, *^3^J* = 5.71 Hz, 1H), 7.51-7.48 (m, 1H), 7.43 (d, *^4^J* = 1.86 Hz, 1H), 7.42-7.38 (m, 1H), 7.32-7.28 (m, 1H), 7.17 (s, 1H), 7.15 (dd, (m, *^3^J* = 8.23 Hz, *^4^J* = 1.86 Hz, 1H), 6.82-6.77 (m, 1H), 4.41 (d, *^3^J* = 5.71 Hz, 2H), 4.16 (t, *^3^J* = 6.88 Hz, 2H), 4.13-4.08 (m, 2H), 3.84-3.81 (m, 2H), 3.75 - 3.53 (m, 24H), 3.23-3.15 (m, 2H), 2.7-2.69 (m, 2H), 2.47 (t, *^3^J* = 6.70 Hz, 2H), 1.44-1.38 (m, 9H) ppm. ^13^C NMR (125.8 MHz, CDCl_3_) δ = 171.2, 171.1, 157.4, 152.0, 139.1, 136.1, 132.2, 131.2, 130.8, 130.4, 129.1, 128.2, 126.0, 125.9, 119.6, 112.3, 111.0, 80.7, 70.7, 70.7, 70.7, 70.7, 70.6, 70.5, 70.5, 70.4, 70.4, 70.3, 69.5, 68.1, 66.9, 47.6, 42.0, 38.8, 36.3, 28.2, 25.7, 23.5 ppm.

### Synthesis of tert-butyl 1-(2-((2-(3,4-dichlorophenyl)acetamido)methyl)-4-(6,7-dihydro-5H- pyrrolo[1,2-a]imidazol-2-yl)phenoxy)-3,6,9,12,15,18,21-heptaoxatetracosan-24-oate (16g)

The reaction was carried out as described in General Procedure B using tert-butyl 1-hydroxy- 3,6,9,12,15,18,21-heptaoxatetracosan-24-oate as commercial available starting material, as TFA salt with unknown stoichiometry as a colorless oil (59 mg). MALDI: (calculated) [M+H^±^] 852.36 g/mol, (found) [M+H^±^] 852.42 g/mol. ^1^H NMR (500 MHz, CDCl_3_) δ = 7.98-7.89 (m, 1H), 7.50-7.46 (m, 1H), 7.43- 7.41 (m, 1H), 7.40-7.36 (m, 1H), 7.30-7.27 (m, 1H), 7.17 (s, 1H), 7.16-7.12 (m, 1H), 6.80-6.76 (m, 1H), 4.40 (d, *^3^J* = 5.38 Hz, 2H), 4.17-4.12 (m, 2H), 4.11-4.07 (m, 2H), 3.83-3.79 (m, 2H), 3.74-3.51 (m, 28H), 3.21-3.14 (m, 2H), 2.75-2.67 (m, 2H), 2.48-2.44 (m 2H), 1.42-1.40 (m, 9H) ppm. ^13^C NMR (125.8 MHz, CDCl_3_): δ = 171.1, 171.0, 157.4, 152.0, 139.0, 136.1, 132.1, 131.2, 130.7, 130.4, 129.1, 128.2, 125.9, 125.8, 119.6, 112.2, 111.0, 80.7, 70.7, 70.7, 70.7, 70.6, 70.5, 70.5, 70.5, 70.5, 70.5, 70.4, 70.4, 70.3, 69.5, 68.0, 66.9, 47.6, 42.0, 38.8, 36.3, 28.1, 25.6, 23.4 ppm.

### Synthesis of (2S,4R)-1-((S)-2-(3-(2-(2-((2-(3,4-dichlorophenyl)acetamido)methyl)-4-(6,7-dihydro-5H- pyrrolo[1,2-a]imidazol-2-yl)phenoxy)ethoxy)propanamido)-3,3-dimethylbutanoyl)-4-hydroxy-N-(4- (4-methylthiazol-5-yl)benzyl)pyrrolidine-2-carboxamide (17a)

The reaction was carried out as described in General Procedure B using intermediate **16a**, yielding in 17 mg, 18 µmol, 40% of a white solid. MALDI: (calculated) [M+2H^±^] 945.34 g/mol, (found) [M+2H^±^] 945.67 g/mol. HRMS: (calculated) [M+Na^±^] 966.3153 g/mol, (found) [M+Na^±^] 966.3156 g/mol. HPLC: R_t_ = 12.3 min (254 nm, 100%). ^1^H NMR (500 MHz, DMSO-*d6*) δ = 8.97 (s, 1H), 8.56 (t, *^3^J* = 5.70 Hz, 1H), 8.42 (t, *^3^J* = 5.85 Hz, 1H), 7.95 (d, *^3^J* = 9.40 Hz, 1H), 7.58 (d, *^2^J* = 2.11 Hz, 1H), 7.56 (d, *^3^J* = 8.33 Hz, 1H), 7.53 (dd, *^3^J* = 8.54 Hz, *^4^J* = 2.10 Hz, 1H), 7.43 (d, *^4^J* = 2.10 Hz, 1H), 7.42 (d, *^3^J* = 8.40 Hz, 2H), 7.38 (d, *^3^J* = 8.40 Hz, 2H), 7.32 (dd, *^3^J* = 8.33 Hz, *^4^J* = 2.11 Hz, 1H), 7.25 (s, 1H, H-3e), 6.94 (d, *^3^J* = 8.54 Hz, 1H), 5.14 (bs, 1H), 4.56 (d, *^3^J* = 9.40 Hz, 1H), 4.46-4.39 (m, 2H), 4.35 (br, s, 1H), 4.26 (d, *^3^J* = 5.85 Hz, 2H), 4.22 (dd, *^2^J* = 15.86 Hz, *^3^J* = 5.70 Hz, 1H), 4.09 Hz, (t, *^3^J* = 4.64 Hz, 2H), 3.98 (t, *^3^J* = 7.16 Hz, 2H), 3.75-3.61 (m, 6H), 3.56 (s, 2H), 2.78 (t, *^3^J* = 7.04 Hz, 2H), 2.62-2.55 (m, 1H), 2.50 (m, 2H), 2.44 (s, 3H), 2.42-2.36 (m, 1H), 2.07-2.01 (m, 1H), 1.94-1.87 (m, 1H), 0.92 (s, 9H) ppm. ^13^C NMR (126 MHz, DMSO) δ = 171.9, 169.9, 169.5, 169.3, 154.4, 154.0, 151.4, 147.7, 144.6, 139.5, 137.7, 131.1, 131.0, 130.7, 130.3, 129.6, 129.5, 129.0, 128.8, 128.6, 127.8, 127.4, 126.8, 123.6, 111.9, 109.7, 68.9, 68.6, 67.7, 67.2, 58.7, 56.3, 56.3, 44.3, 41.6, 41.2, 37.9, 37.5, 35.7, 35.3, 26.3, 25.6, 22.5, 15.9 ppm.

### Synthesis of (2S,4R)-1-((S)-2-(3-(2-(2-(2-((2-(3,4-dichlorophenyl)acetamido)methyl)-4-(6,7-dihydro- 5H-pyrrolo[1,2-a]imidazol-2-yl)phenoxy)ethoxy)ethoxy)propanamido)-3,3-dimethylbutanoyl)-4- hydroxy-N-(4-(4-methylthiazol-5-yl)benzyl)pyrrolidine-2-carboxamide (17b)

The reaction was carried out as described in General Procedure B using intermediate **16b**, yielding in 30 mg, 30 µmol, 48% of a white solid. MALDI: (calculated) [M+2H^±^] 989.36 g/mol, (found) [M+2H^±^] 989.89 g/mol. HRMS: (calculated) [M+Na^±^] 1010.3415 g/mol, (found) [M+Na^±^] 1010.3442 g/mol. HPLC: R_t_ = 12.3 min (254 nm, 100%). ^1^H NMR (500 MHz, DMSO-*d6*) δ = 8.97 (s, 1H), 8.56 (t, *^3^J* = 5.82 Hz, 1H), 8.41 (t, *^3^J* = 5.64 Hz, 1H), 7.91 (d, *^3^J* = 9.38 Hz, 1H), 7.58 (d, *^4^J* = 2.04 Hz, 1H), 7.57 (d, *^3^J* = 8.31 Hz, 1H), 7.54 (dd, *^3^J* = 8.48 Hz, *^4^J* = 2.25 Hz, 1H), 7.43 (d, *^4^J* = 2.25 Hz, 1H), 7.42 (d, *^3^J* = 8.37 Hz, 2H), 7.38 (d, *^3^J* = 8.37 Hz, 2H), 7.32 (dd, *^3^J* = 8.31 Hz, *^4^J* = 2.04 Hz, 1H), 7.23 (s, 1H), 6.94 (d, *^3^J* = 8.48 Hz, 1H), 5.14 (br, s, 1H), 4.55 (d, *^3^J* = 9.38 Hz, 1H), 4.46-4.40 (m, 2H), 4.35 (bs, 1H), 4.26 (d, *^3^J* = 5.82 Hz, 2H), 4.22 (dd, *^2^J* = 15.89 Hz, *^3^J* = 5.64 Hz, 1H), 4.09 Hz, (t, *^3^J* = 4.56 Hz, 2H), 3.97 (t, *^3^J* = 7.02 Hz, 2H), 3.72 (t, *^3^J* = 4.56 Hz, 2H), 3.69-3.46 (m, 10H), 2.78 (t, *^3^J* = 6.62 Hz, 2H), 2.62-2.55 (m, 1H), 2.50 (m, 2H), 2.44 (s, 3H), 2.40-2.32 (m, 1H), 2.06-2.00 (m, 1H), 1.94-1.87 (m, 1H), 0.93 (s, 9H) ppm. ^13^C NMR (126 MHz, DMSO) δ = 171.9, 169.9, 169.5, 169.3, 154.4, 154.0, 151.4, 147.7, 144.7, 139.5, 137.7, 131.14, 131.0, 130.7, 130.3, 129.6, 129.5, 129.0, 128.6, 127.9, 127.4, 126.8, 123.6, 111.9, 109.7, 69.9, 69.5, 69.0, 68.8, 67.7, 66.9, 59.7, 58.7, 56.3, 56.3, 44.3, 41.6, 41.2, 37.9, 37.5, 35.6, 35.3, 26.3, 25.6, 22.4, 15.9 ppm.

### Synthesis of (2S,4R)-1-((S)-14-(tert-butyl)-1-(2-((2-(3,4-dichlorophenyl)acetamido)methyl)-4-(6,7- dihydro-5H-pyrrolo[1,2-a]imidazol-2-yl)phenoxy)-12-oxo-3,6,9-trioxa-13-azapentadecan-15-oyl)-4- hydroxy-N-(4-(4-methylthiazol-5-yl)benzyl)pyrrolidine-2-carboxamide (17c)

The reaction was carried out as described in General Procedure B using intermediate **16c**, yielding in 24 mg, 23.4 µmol, 52% of a white solid. MALDI: (calculated) [M+2H^±^] 1033.40 g/mol, (found) [M+2H^±^] 1033.98 g/mol. HRMS: (calculated) [M+Na^±^] 1054.3677 g/mol, (found) [M+Na^±^] 1036.3689 g/mol. HPLC: R_t_ = 12.4 min (254 nm, 100%). ^1^H NMR (500 MHz, DMSO-*d6*) δ = 8.97 (s, 1H), 8.57 (t, *^3^J* = 5.60 Hz, 1H), 8.42 (t, *^3^J* = 5.80 Hz, 1H), 7.90 (d, *^3^J* = 9.42 Hz, 1H), 7.58 (d, *^4^J* = 2.08, 1H), 7.57 (d, *^3^J* = 8.33 Hz, 1H), 7.54 (dd, *^3^J* = 8.48 Hz, *^4^J* = 2.20 Hz, 1H), 7.43 (d, *^4^J* = 2.20 Hz, 1H), 7.41 (d, *^3^J* = 8.39 Hz, 2H), 7.38 (d, *^3^J* = 8.39 Hz, 2H), 7.32 (dd, *^3^J* = 8.33 Hz, *^4^J* = 2.08 Hz, 1H), 7.24 (s, 1H), 6.95 (d, *^3^J* = 8.48 Hz, 1H), 5.15 (br, s, 1H), 4.55 (d, *^3^J* = 9.42 Hz, 1H), 4.46-4.40 (m, 2H), 4.35 (br, s, 1H), 4.26 (d, *^3^J* = 5.60 Hz, 2H), 4.22 (dd, *^2^J* = 15.99 Hz, *^3^J* = 5.80 Hz, 1H), 4.11-4.08 Hz, (m, 2H), 3.97 (t, *^3^J* = 6.93 Hz, 2H), 3.74-3.70 (m, 2H), 3.69-3.42 (m, 14H), 2.77 (t, *^3^J* = 6.93 Hz, 2H), 2.57-2.53 (m, 1H), 2.50 (m, 2H), 2.44 (s, 3H), 2.38-2.30 (m, 1H), 2.07-2.01 (m, 1H), 1.94-1.87 (m, 1H), 0.93 (s, 9H) ppm. ^13^C NMR (126 MHz, DMSO) δ = 171.9, 169.9, 169.5, 169.3, 154.5, 154.0, 151.4, 147.7, 144.6, 139.5, 137.7, 131.1, 131.0, 130.7, 130.3, 129.6, 129.5, 129.1, 128.8, 128.6, 127.8, 127.4, 126.8, 123.6, 112.0, 109.8, 70.0, 69.8, 69.72, 69.46, 69.0, 68.9, 67.8, 66.9, 58.7, 56.3, 56.3, 44.3, 41.6, 41.2, 37.9, 37.6, 35.7, 35.3, 26.3, 25.6, 22.5, 15.9 ppm.

### Synthesis of (2S,4R)-1-((S)-17-(tert-butyl)-1-(2-((2-(3,4-dichlorophenyl)acetamido)methyl)-4-(6,7- dihydro-5H-pyrrolo[1,2-a]imidazol-2-yl)phenoxy)-15-oxo-3,6,9,12-tetraoxa-16-azaoctadecan-18- oyl)-4-hydroxy-N-(4-(4-methylthiazol-5-yl)benzyl)pyrrolidine-2-carboxamide (17d)

The reaction was carried out as described in General Procedure B using intermediate **16d**, yielding in 6.9 mg, 6.4 µmol, 12% of a yellow oil. MALDI: (calculated) [M+2H^±^] 1076.41 g/mol, (found) [M+2H^±^] 1076.10 g/mol. HRMS: (calculated) [M+Na^±^] 1098.3939 g/mol, (found) [M+Na^±^] 1098.3954 g/mol. HPLC: R_t_ = 12.4 min (254 nm, 100%). ^1^H NMR (500 MHz, DMSO-*d6*) δ = 8.97 (s, 1H), 8.56 (t, *^3^J* = 5.55 Hz, 1H), 8.40 (t, *^3^J* = 5.76 Hz, 1H), 7.90 (d, *^3^J* = 9.38 Hz, 1H), 7.58-7.55 (m, 2H), 7.54 (dd, *^3^J* = 8.48 Hz, *^4^J* = 2.07 Hz, 1H), 7.43 (d, *^4^J* = 2.07 Hz, 1H), 7.42 (d, *^3^J* = 8.31 Hz, 2H), 7.38 (d, *^3^J* = 8.31 Hz, 2H), 7.32 (dd, *^3^J* = 8.17 Hz, *^4^J* = 1.93 Hz, 1H), 7.24 (s, 1H), 6.95 (d, *^3^J* = 8.48 Hz, 1H), 5.13 (br, s, 1H), 4.55 (d, *^3^J* = 9.38 Hz, 1H), 4.46- 4.39 (m, 2H), 4.35 (br, s, 1H), 4.26 (d, *^3^J* = 5.55 Hz, 2H), 4.22 (dd, *^2^J* = 15.87 Hz, *^3^J* = 5.76 Hz, 1H), 4.12- 4.08 Hz, (m, 2H), 3.97 (t, *^3^J* = 7.02 Hz, 2H), 3.75-3.71 (m, 2H), 3.69-3.42 (m, 18H), 2.76 (t, *^3^J* = 7.29 Hz, 2H), 2.56-2.50 (m, 1H), 2.50 (m, 2H), 2.44 (s, 3H), 2.37-2.30 (m, 1H), 2.07-2.00 (m, 1H), 1.94-1.87 (m, 1H), 0.93 (s, 9H) ppm. ^13^C NMR (126 MHz, DMSO) δ = 171.9, 169.9, 169.5, 169.3, 154.5, 154.0, 151.4, 147.7, 144.6, 139.5, 137.7, 131.1, 131.0, 130.7, 130.3, 129.6, 129.5, 129.0, 128.8, 128.6, 127.8, 127.4, 126.8, 123.6, 112.0, 109.8, 70.0, 69.8, 69.7, 6978, 69.45, 69.0, 68.8, 67.8, 66.9, 58.7, 56.3, 56.3, 44.3, 41.6, 41.2, 37.9, 37.6, 35.6, 35.3, 26.3, 25.6, 22.5, 15.9 ppm.

### Synthesis of (2S,4R)-1-((S)-20-(tert-butyl)-1-(2-((2-(3,4-dichlorophenyl)acetamido)methyl)-4-(6,7- dihydro-5H-pyrrolo[1,2-a]imidazol-2-yl)phenoxy)-18-oxo-3,6,9,12,15-pentaoxa-19-azahenicosan- 21-oyl)-4-hydroxy-N-(4-(4-methylthiazol-5-yl)benzyl)pyrrolidine-2-carboxamide (17e)

The reaction was carried out as described in General Procedure B using intermediate **16e**, yielding in 17 mg, 15.2 µmol, 36% of a yellow oil. MALDI: (calculated) [M+3H^±^] 1122.45 g/mol, (found) [M+3H^±^] 1122.14 g/mol. HRMS: (calculated) [M+Na^±^] 1142.4202 g/mol, (found) [M+Na^±^] 1142.4229 g/mol. HPLC: R_t_ = 12.4 min (254 nm, 100%). ^1^H NMR (500 MHz, DMSO-*d6*) δ = 8.98 (s, 1H), 8.56 (t, *^3^J* = 5.98 Hz, 1H), 8.40 (t, *^3^J* = 5.68 Hz, 1H), 7.90 (d, *^3^J* = 9.41 Hz, 1H), 7.82 (s, 1H), 7.59 (dd, 1H, *^3^J* = 8.55, *^4^J* = 2.05, 1H), 7.56 (d, *^3^J* = 8.00 Hz, 1H), 7.55 (s, 1H), 7.45 (d, *^4^J* = 2.05 Hz, 1H), 7.42 (d, *^3^J* = 8.32 Hz, 2H), 7.38 (d, *^3^J* = 8.32 Hz, 2H), 7.28 (dd, *^3^J* = 8.00 Hz, *^4^J* = 1.95 Hz, 1H), 7.15 (d, *^3^J* = 8.55 Hz, 1H), 4.55 (d, *^3^J* = 9.41 Hz, 1H), 4.46-4.40 (m, 2H), 4.37-4.33 (m, 1H), 4.28 (d, *^3^J* = 5.68 Hz, 2H), 4.25-4.19 (m, 3H), 4.18 (t, *^3^J* = 4.42 Hz, 2H), 3.78-3.74 (m, 2H), 3.70-3.41 (m, 22H), 3.18 (t, *^3^J* = 7.53 Hz, 2H), 2.73-2.65 (m, 2H), 2.57-2.51 (m, 1H), 2.44 (s, 3H), 2.38-2.31 (m, 1H), 2.07-2.01 (m, 1H), 1.94-1.88 (m, 1H), 0.93 (s, 9H) ppm. ^13^C NMR (126 MHz, DMSO) δ = 171.9, 169.9, 169.5, 169.2, 154.5, 154.0, 151.4, 147.70, 139.5, 137.68, 131.1, 131.0, 130.7, 130.3, 129.6, 129.5, 129.0, 128.8, 128.6, 128.0, 127.41, 126.81, 123.7, 123.6, 112.0, 109.8, 70.0, 69.8, 69.7, 69.7, 69.4, 69.0, 68.8, 67.8, 66.9, 58.7, 56.3, 44.4, 41.6, 41.2, 37.9, 37.6, 35.6, 35.3, 26.3, 22.5, 15.9 ppm.

### Synthesis of (2S,4R)-1-((S)-23-(tert-butyl)-1-(2-((2-(3,4-dichlorophenyl)acetamido)methyl)-4-(6,7- dihydro-5H-pyrrolo[1,2-a]imidazol-2-yl)phenoxy)-21-oxo-3,6,9,12,15,18-hexaoxa-22-azatetracosan- 24-oyl)-4-hydroxy-N-(4-(4-methylthiazol-5-yl)benzyl)pyrrolidine-2-carboxamide (17f)

The reaction was carried out as described in General Procedure B using intermediate **16f**, yielding in 6.2 mg, 5.3 µmol, 27% of a yellow oil. MALDI: (calculated) [M+3H^±^] 1166.48 g/mol, (found) [M+3H^±^] 1166.20 g/mol. HRMS: (calculated) [M+Na^±^] 1186.4464 g/mol, (found) [M+Na^±^] 1186.4497 g/mol. HPLC: R_t_ = 12.4 min (254 nm, 100%). ^1^H NMR (400 MHz, DMSO-*d6*, water supp.) δ = 8.98 (s, 1H), 8.55 (t, *^3^J* = 5.69 Hz, 1H), 8.38 (t, *^3^J* = 5.52 Hz, 1H), 7.90 (d, *^3^J* = 9.22 Hz, 1H), 7.82 (s, 1H), 7.61-7.53 (m, 3H), 7.44 (d, *^4^J* = 1.64 Hz, 1H), 7.42 (d, *^3^J* = 8.27 Hz, 2H), 7.38 (d, *^3^J* = 8.27 Hz, 2H), 7.28 (dd, *^3^J* = 8.18 Hz, *^4^J* = 1.64 Hz, 1H), 7.15 (d, *^3^J* = 8.62 Hz, 1H), 4.55 (d, *^3^J* = 9.22 Hz, 1H), 4.47-4.39 (m, 2H, H-10a), 4.35 (br, s, 1H), 4.27 (d, *^3^J* = 5.52 Hz, 2H), 4.26-4.20 (m, 3H), 4.18 (t, *^3^J* = 4.19 Hz, 2H), 3.76 (t, *^3^J* = 4.19 Hz, 2H), 3.70-3.42 (m, 26H), 3.18 (t, *^3^J* = 7.58 Hz, 2H), 2.74-2.64 (m, 2H), 2.58-2.50 (m, 1H), 2.44 (s, 3H), 2.39- 2.30 (m, 1H), 2.06-1.99 (m, 1H), 1.95-1.86 (m, 1H), 0.93 (s, 9H) ppm. ^13^C NMR (126 MHz, DMSO) δ = 171.9, 169.9, 169.5, 169.2, 154.5, 154.0, 151.4, 147.7, 139.5, 137.7, 131.1, 131.0, 130.7, 130.3, 129.6, 129.5, 129.0, 128.8, 128.6, 128.0, 127.4, 126.8, 123.7, 123.6, 112.0, 109.8, 70.0, 69.8, 69.7, 69.7, 69.4, 69.0, 68.8, 67.8, 66.9, 58.7, 56.3, 44.4, 41.6, 41.2, 37.9, 37.5, 35.6, 35.3, 26.3, 25.6, 22.5, 15.9 ppm.

### Synthesis of (2S,4R)-1-((S)-26-(tert-butyl)-1-(2-((2-(3,4-dichlorophenyl)acetamido)methyl)-4-(6,7- dihydro-5H-pyrrolo[1,2-a]imidazol-2-yl)phenoxy)-24-oxo-3,6,9,12,15,18,21-heptaoxa-25- azaheptacosan-27-oyl)-4-hydroxy-N-(4-(4-methylthiazol-5-yl)benzyl)pyrrolidine-2- carboxamide (17g)

The reaction was carried out as described in General Procedure B using intermediate **16g**, yielding in 18 mg, 15 µmol, 36% of a yellow oil. MALDI: (calculated) [M+3H^±^] 1210.50 g/mol, (found) [M+3H^±^] 1210.16 g/mol. HRMS: (calculated) [M+Na^±^] 1230.4726 g/mol, (found) [M+Na^±^] 1230.4747 g/mol. HPLC: R_t_ = 12.4 min (254 nm, 100%). ^1^H NMR (400 MHz, DMSO-*d6*, water supp.) δ = 8.98 (s, 1H), 8.56 (t, *^3^J* = 6.03 Hz, 1H), 8.39 (t, *^3^J* = 5.59 Hz, 1H), 7.90 (d, *^3^J* = 9.34 Hz, 1H), 7.82 (s, 1H), 7.59 (dd, *^3^J* = 8.50 Hz, *^4^J* = 2.15 Hz, 1H), 7.57-7.54 (m, 2H), 7.45 (d, *^4^J* = 1.83 Hz, 1H), 7.42 (d,*^3^J* = 8.15 Hz, 2H), 7.38 (d, *^3^J* = 8.29 Hz, 2H), 7.28 (dd, *^3^J* = 8.18 Hz, *^4^J* = 1.89 Hz, 1H), 7.15 (d, *^3^J* = 8.63 Hz, 1H), 4.55 (d, *^3^J* = 9.15 Hz, 1H), 4.47-4.39 (m, 2H), 4.38-4.33 (m, 1H), 4.28 (d, *^3^J* = 5.38 Hz, 2H), 4.26-4.20 (m, 3H), 4.18 (t, *^3^J* = 4.71 Hz, 2H), 3.76 (t, *^3^J* = 4.46 Hz, 2H), 3.73-3.43 (m, 30H), 3.18 (t, *^3^J* = 7.78 Hz, 2H), 2.73-2.64 (m, 2H), 2.58-2.50 (m, 1H), 2.44 (s, 3H), 2.39-2.30 (m, 1H), 2.08-2.00 (m, 1H), 1.95-1.86 (m, 1H), 0.93 (s, 9H) ppm. ^13^C NMR (126 MHz, DMSO) δ = 171.9, 169.9, 169.5, 169.3, 154.5, 154.02, 151.4, 147.7, 144.6, 139.5, 137.7, 131.1, 131.0, 130.7, 130.3, 129.6, 129.5, 129.0, 128.6, 127.9, 127.4, 126.8, 123.6, 123.6, 112.0, 109.8, 70.0, 69.8, 69.7, 69.7, 69.5, 69.0, 68.9, 67.8, 66.9, 58.7, 56.3, 56.3, 44.3, 41.6, 41.2, 37.9, 37.6, 35.7, 35.3, 26.3, 25.6, 22.5, 15.9 ppm.

### Synthesis of (2S,4S)-1-((S)-2-(3-(2-(2-(2-((2-(3,4-dichlorophenyl)acetamido)methyl)-4-(6,7-dihydro- 5H-pyrrolo[1,2-a]imidazol-2-yl)phenoxy)ethoxy)ethoxy)propanamido)-3,3-dimethylbutanoyl)-4- hydroxy-N-(4-(4-methylthiazol-5-yl)benzyl)pyrrolidine-2-carboxamide (21)

The reaction was carried out as described in General Procedure B, yielding in 17 mg, 17.2 µmol, 39% of a yellow solid. MALDI: (calculated) [M+H^+^] 988.36 g/mol, (found) [M+H^+^] 988.31 g/mol. HRMS: (calculated) [M+H^+^] 988.3596 g/mol, (found) [M+H^+^] 988.3242 g/mol. HPLC: R_t_ = 11.8 min (254 nm, 100%). ^1^H NMR (500 MHz, DMSO) δ = 8.98 (s, 1H), 8.44 – 8.37 (m, 2H), 7.81 (s, 1H), 7.59 (dd, *^3^J* = 8.5 Hz, *^4^J* = 2.3 Hz, 1H), 7.58 - 7.54 (m, 2H), 7.46 – 7.39 (m, 3H), 7.38 - 7.34 (m, 2H), 7.28 (dd, *^3^J* = 8.2 Hz, *^4^J* = 1.9 Hz, 1H), 7.16 (d, *^3^J* = 8.7 Hz, 1H), 4.54 (t, *^3^J* = 7.5 Hz, 1H), 4.51 – 4.42 (m, 1H), 4.39 – 4.30 (m, 4H), 4.28 - 4.26 (m 3H), 4.25 – 4.21 (m, 3H), 4.20 – 4.14 (m, 3H), 3.77 – 3.73 (m, 2H), 3.64 – 3.52 (m, 8H), 3.51 – 3.48 (m, 2H), 3.38 - 3.36 (m, 1H), 3.18 (t, *^3^J* = 7.5 Hz, 2H), 2.74 – 2.65 (m, 2H), 2.44 (s, 3H), 2.31 - 2.28 (m, 1H), 2.24 - 2.20 (m, 1H), 2.07 – 2.01 (m, 1H), 1.97 – 1.87 (m, 1H), 1.81 - 1.77 (m, 1H) ppm. ^13^C NMR (75 MHz, DMSO) δ = 172.2, 170.6, 169.3, 168.7, 154.4, 154.0, 151.4, 147.8, 144.7, 139.2, 137.7, 131.1, 131.0, 130.7, 130.4, 129.8, 129.5, 129.1, 128.7, 128.6, 127.9, 127.4, 126.8, 123.6, 112.0, 109.8, 82.0, 69.9, 69.7, 69.3, 69.0, 68.7, 67.8, 66.2, 58.8, 56.5, 54.7, 44.3, 41.7, 41.2, 37.5, 36.9, 34.3, 25.6, 22.5, 15.9 ppm.

### Protein Purification

WDR5A was expressed and purified as described elsewhere.^17^ Plasmids of WDR5A (aa 1-334 and aa 33-334) were a kind gift of M. Vedadi from the SGC Toronto. Briefly, WDR5 was overexpressed in *E. coli* BL21 using TB media. Protein expression was induced by addition of 0.5 mM IPTG. Cell were grown overnight at 18 °C. Next morning, the cells were harvested and resuspended in Lysis buffer (50 mM HEPES buffer, pH 7.5, 500 mM NaCl, 20 mM imidazole, 0.5 mM TCEP and 5% glycerol). For purification, the cells were lysed by sonication. After centrifugation, the supernatant was loaded onto a Nickel-Sepharose column equilibrated with 30 mL lysis buffer. The column was washed with 100 mL lysis buffer. WDR5 was eluted by an imidazole step gradient (50, 100, 200, 300 mM). Fractions containing WDR5 were pooled together, concentrated, and loaded onto a Superdex 200 16/60 HiLoad gel filtration column equilibrated with final buffer (25 mM HEPES pH 7.5, 300 mM NaCl and 0.5 mM TCEP. The protein was concentrated to approx. 400 µM. The buffer was kept and used for ITC experiments.

### Differential Scanning Fluorimetry

Ligand binding to protein was detected using DSF on an MX3005P qPCR system from Agilent Technologies as described elsewhere.^37^ Briefly, protein was buffered in 25 mM HEPES (pH 7.5), 500 mM NaCl, 0.5 mM TCEP and diluted to a final concentration of 2 μM, and the fluorescent dye SYPRO Orange was added at a dilution of 1:1000. Compounds were dissolved in DMSO (10 mM) and added at a final concentration of 10 μM to 20 μL of protein-dye mix in a 96-well plate. Real-time melting curves were then recorded by heating the samples from 25 to 96 °C in 71 cycles (heating rate of 270 K/h, excitation/emission filters = 492/610 nm), and the melting point, T_m_, was calculated using the Boltzmann equation.

### Isothermal Titration Calorimetry

Binding constant (K_d_), stoichiometry (n) and thermodynamic binding parameters (ΔH, ΔS and ΔG) of ligand−protein interactions were determined on a Nano-ITC from TA Instruments as described elsewhere.^38^ Briefly, compounds were diluted to a final concentration of 10−25 μM in buffer and placed into the sample cell. Proteins (80−120 μM) were added using an initial injection of 3 or 4 μL, followed by 12-30 injections of 6 or 8 μL at 22 °C. Collected data were corrected by subtraction of pure DMSO injection heats. Data were analyzed by assuming a sigmoidal dose−response relationship (four parameters). Errors of fits were calculated using standard deviation and a confidence interval of 68% in GraphPad Prism.

### NanoBRET^TM^

The NanoBRET^TM^ assay was performed as described previously.^39, 40^ Briefly, Full-length WDR5 was cloned in frame as N- or C-terminal NanoLuc-fusion pNLF1 vector using ligation independent in-fusion cloning (Takara Bio) and sequence verified. Plasmids were transfected into HEK293T cells using FuGENE HD (Promega, E2312) and proteins were allowed to express for 20 h. Serially diluted inhibitor and Tracer molecule **19c** at a concentration of 1 µM determined previously as the Tracer **19c** K_D,app_ were pipetted into white 384-well plates (Greiner 781 207) using an Echo acoustic dispenser (Labcyte). The corresponding protein-transfected cells were added and reseeded at a density of 2 x 10^5^ cells/ml after trypsinization and resuspending in Opti-MEM without phenol red (Life Technologies). The system was allowed to equilibrate for 2 hours at 37 °C/5% CO_2_ prior to BRET measurements. To measure BRET, NanoBRET^TM^ NanoGlo Substrate and Extracellular NanoLuc Inhibitor (Promega, N2540) was added as per the manufacturer’s protocol, and filtered luminescence was measured on a PHERAstar plate reader (BMG Labtech) equipped with a luminescence filter pair (450 nm BP filter (donor) and 610 nm LP filter (acceptor). Competitive displacement data were then graphed using GraphPad Prism 8 software using a normalized 3-parameter curve fit.

### Cell culture

Human MV4-11 (male) and human HL-60 (female) cells were cultured in RPMI-1640 medium whereas human HEK293 (female) cells were cultured in DMEM medium at 37 °C in 5% CO_2_. Both medium were supplemented with 10% FBS and 1% penicillin/streptomycin solution.

### Cloning

WDR5-HiBiT was cloned by PCR amplification of vector containing full length WDR5 using forward primer: CGCACCGGTATGGCGACGGAGGAGAAGAAGC and reverse primer: CGCGACGCGTTTAGCTAATCTTCTTGAACAGCCGCCAGCCGCTCACACCGGAGCTCCCGCAGTCACTCTTCCAC AGT. The PCR product was inserted into pRRL-PGK vector using AgeI/MluI restriction sites. HA-tagged VHL was cloned by PCR amplification of cDNA from MV4-11 cells as template (forward primer: CGCACCGGTATGTACCCTTACGACGTGCCCGACTACGCCGGGAGCTCCGGTCCCCGGAGGGCGGAGAAC and reverse primer: GGACTAGTTCAATCTCCCATCCGTTGATGTG) and inserted into pRRL-SFFV vector using AgeI/SpeI sites. The sequence of the cloned VHL was identified as isoform 1 by sanger sequencing.

### Cell line generation

Lentiviral infection was used to generate stable MV4-11^WDR5-HiBiT^, MV4-11^VHL^ and MV4-11^WDR5-HiBiT/VHL^ cells. Lentivirus was produced using plasmids psPAX2, pMD2.G and HiBiT-WDR5 or HA-VHL plasmid in HEK293 cells. MV4-11 cells were infected with filtered virus supernatant and selected after 72 h of infection for generation of MV4-11^WDR5-HiBiT^ and MV4-11^VHL^ cells. MV4-11^WDR5-HiBiT^ cells were used to prepare MV4-11^WDR5-HiBiT/VHL^ cells.

### HiBiT assay

HiBiT assay was performed as described previously.^35^ MV4-11^WDR5-HiBiT^ were seeded and treated with serial dilutions of compounds for 6 h or 24h. Nano-Glo HiBiT Lytic Detection System (Promega) was used for the assay. Luminescence was measured on a GloMax 96 Microplate Luminometer (Promega). DC_50_ was calculated using lower concentrations (showing sigmoidal behaviour) with the dose–response (four parameters) equation.

### Immunoblotting

After the treatment, cells were lysed in RIPA lysis buffer (50 mM HEPES pH 7.9, 140 mM NaCl, 1 mM EDTA, 1% Triton X-100, 0.1% SDS, 0.1% sodium deoxycholate) supplemented with protease and phosphatase inhibitors (Sigma) for 20 min at 4 °C head-over-tail. Supernatant were collected after centrifugation. Bicinchoninic acid (BCA) assay was used for protein quantification. Per sample equal amounts of protein were separated by Bis-Tris-PAGE and transferred to PVDF membranes (Millipore). The membranes were incubated with 5% (w/v) non-fat dry milk in TBS-T (20 mM Tris-HCl, pH 7.5, 150 mM NaCl, 0.1% (v/v) Tween-20 for 1 h at room temperature for blocking and then incubated with the primary antibody overnight at 4 °C. For visualisation, horseradish peroxidase (HRP)-labelled secondary antibodies were used and detected using chemiluminescent HRP substrate (Millipore) in LAS4000 Mini (Fuji). The signal was quantified using ImageJ (version 1.53g) or Image Studio Lite (LI-COR Biosciences, version 5.2.5). Vinculin was used as a loading control. Antibodies used in this study were: WDR5 (Santa Cruz Biotechnology; sc-393080), HA (Santa Cruz Biotechnology; sc-805), VHL (Santa Cruz Biotechnology; sc-135657) and vinculin (Sigma; V9131).

### Cycloheximide chase assay

Cycloheximide chase assay was performed as described previously.^35^ MV4-11 cells were treated with 50 µg/ml CHX with or without PROTACs for different time points. The cells were harvested in RIPA buffer and probed for immunoblotting. The intensity of WDR5 band at 0 h was set as 1. The mean ± SD from n=2 biological experiments were plotted as log_10_ values. **RT-qPCR.** RT-qPCR was performed as described previously.^35^ Total RNA was extracted using peqGOLD TriFast (Peqlab). cDNA synthesis was carried out and cDNA were analysed by qPCR on a StepOnePlus Real-Time PCR System (Thermo Fisher Scientific) using the SYBR Green Master Mix (Thermo Fisher Scientific). Equal amounts of cDNA and SYBR Green Master Mix were added along with WDR5 primers (forward: CCAGTCTCGGCCGTTCATTT, reverse: CGTTCGGGGAGAACTTCACA). For analysis, expression was normalized to β2-microglobulin expression. qPCR was done in technical triplicates.

### Cell proliferation assay

MV4-11 or MV4-11^VHL^ cells were seeded at density of 1x 10^5^ cells per ml and treated with compounds. Cells were counted every third day and reseeded to original density of 1x 10^5^ cells per ml in fresh media with compounds. The mean cumulative cell number ± SEM (n=2 biological experiments) were plotted as log_10_ values over the course of time. P-value were calculated from cumulative cell number from end point using two-tailed unpaired t-test assuming equal variance against DMSO treated cells.

### Quantitative proteomics

4 million MV4-11 cells (in 10 ml) were seeded at least in triplicates for each treatment evening before the treatment. Cells were treated with either 1 µM **6**, 1 µM **8g**, 5 µM **17b**, 5 µM 1**4** or DMSO as control for 9 hours. After treatment, cells were washed twice with ice cold PBS supplemented with protease and phosphatase inhibitor and lysed in SDS lysis buffer (2% SDS in 40 mM Tris-HCl, pH 7.6). In order to reduce viscosity, the sample was sonicated using a sonication water bath (10 cycles, 15 sec sonication, 15 sec pause on ice), boiled at 95 °C for 10 min and trifluoroacetic acid was added to a final concentration of 1 %. To neutralize the sample (final pH 7.6-8.0), 300 mM N-methylmorpholin was added to a final concentration of 2 %. The protein concentration in cell lysate was determined using the Pierce^TM^ BCA Protein Assay Kit (ThermoScientific) according to the protocol of the manufacturer. The beads suspension for sp3 sample workup was prepared by mixing magnetic SeraMag-A and SeraMag-B beads (10 µl per sample of each type; Cytiva) in a ratio of 1:1, washing them three times with ddH_2_O and resuspending them in 10 µl ddH_2_O per sample. A total of 200 µg per sample was mixed with 10 µl beads suspension. Acetonitrile (ACN) was added to a final concentration of 70 % and incubated at room temperature, 18 min, 800 rpm. After discarding the supernatant, beads were washed twice using 200 µl 80% ethanol. For reduction and alkylation, beads were resuspended in 70 µl of 2 mM CaCl_2_ in 40 mM Tris pH 7.6. Proteins were reduced with 10 mM dithiothreitol (DTT) for 45 min at 37 °C and 800 rpm, and alkylated with 55 mM chloroacetamide (CAA) at room temperature in the dark for 30 min. Proteins were digested (1:50 trypsin/substrate weight) overnight at 37 °C and 1000 rpm. Samples were centrifuged (5 min, 20000 rcf), sonicated 3 times for 30 sec and supernatant was collected. Beads were washed once with 100 µl ddH_2_O, sonicated 3 times for 30 sec, and supernatants were combined with previous supernatants. Samples were acidified with formic acid (FA) to a final concentration of 1 %. Peptides were desalted using tC18 RP solid-phase extraction cartridges (Waters Corp.; wash solvent: 0.1% FA; elution solvent: 0.1% FA in 50% acetonitrile (ACN). Samples were frozen at −80 °C freezer, dried in a SpeedVac, reconstituted in 0.1 % FA and peptide concentration was determined using a NanoDrop and stored at −20 °C until LC-MS^2^ analysis. A micro-flow LC-MSMS setup with a Q Exactive HF-X mass spectrometer (Thermo Fisher Scientific) was used as described in detail in previous publications^41, 42^. 50 µg peptides dissolved in 0.1 % FA were directly injected onto the microflow LC system. Online chromatography was performed using a commercially available Thermo Fisher Scientific Acclaim PepMap 100 C18 LC column (2 µm particle size, 1 mm ID × 150 mm; catalog number 164711). Column temperature was maintained at 55 °C using the integrated column oven. Peptides were delivered at a flow rate of 50 µl/min and separated using a two-step linear gradient (120 min) ranging from 1-24 % (105 min) and 24-35 % (15 min) of LC solvent B (0.1 % FA, 3 % DMSO in ACN) in LC solvent A (0.1 % FA, 3 % DMSO^43^. The Q Exactive HF-X was operated as follows: positive polarity; spray voltage 4 kV, capillary temperature 320 °C; vaporizer temperature 200 °C. The flow rates of sheath gas, aux gas and sweep gas were set to 40, 3, and 0, respectively. TopN was set to 50. Full MS was readout in the orbitrap, resolution was set to 120,000 and the mass range was set to 360–1300. Full MS AGC target value was 3E6 with a maximum IT of 100 ms and RF lens value was set to 40. Peptide match was set to preferred and default charge state was set to 2. The dynamic exclusion duration was set to 40 s and exclude isotopes was switched on. For readout of MS2 spectra, orbitrap resolution was set to 15,000 and the mass range was set to 200– 2000. The isolation width was set to 1.3 m/z, the first mass was fixed at 100 m/z, NCE was 28. The AGC target value was set to 1E5 at a maximum IT of 22 ms.

Protein and peptide identification and quantification was performed using MaxQuant^44^ (version 1.6.12.0) by searching the MS^2^ data against all canonical protein sequences as annotated in the UniProt reference database (human proteins only, downloaded 24.08.2020) using the search engine Andromeda^45^. Carbamidomethylated cysteine was set as fixed modification; oxidation of methionine and N-terminal protein acetylation were set as variable modification. Trypsin/P was specified as proteolytic enzyme and up to two missed cleavage sites were allowed. The minimum peptide length was set to seven and all data were adjusted to 1 % peptide-spectrum-match (PSM) and 1 % protein false discovery rate (FDR). LFQ based quantification was enabled including the match between runs option and without normalization.

Data analysis was performed using the Perseus software suite^46^ (version 1.6.14.0) and Microsoft Excel on identified and quantified protein groups as provided in the proteinGroups.txt file. Proteingroups.txt was filtered for contaminants and reverse hits, and median centric normalization and log2 transformation were performed. The **14** treated replicate 1 showed high differences to the other conditions and was not considered for further analysis. Entries were filtered for at least three valid values in one condition. Two-sample t-test were performed (S0:0.1, permutation-based FDR: 5%, number of randomizations: 250). For principal component analysis (PCA) remaining missing values were replaced from normal distribution (width 0.3, down shift: 1.8).

The mass spectrometry proteomics data and complete MaxQuant search results have been deposited to the ProteomeXchange Consortium (http://www.proteomexchange.org/) via the PRIDE^47^ partner repository with the data set identifier PXD025257 (Reviewer account details: Username, Password: gZzaqRKs).

### Computational studies on WDR5 degraders. Structure Preparation

Preparation of protein structures started from the published crystal structures of WDR5 in complex with OICR-9429 (PDB: 4QL1, resolution: 1.50 Å)^17^ and of VHL in complex with VH032 (PDB: 4W9H, resolution: 2.10 Å).^36^ The structures were curated within the Molecular Operating Environment (MOE) 2019.01023^48^ using its structure preparation utility. Chain F of structure 4W9H, containing VHL and VH032 was used, and the termini capped. The dimethylarsenic-cystein in position 77 was mutated to cystein, side chains of unresolved amino acids were added automatically and the system was protonated (Protonate3D4)^49^ at the pH of the crystallization buffer (6.3). Chain A of the WDR5 structure 4QL1 was similarly prepared; its termini were capped, and missing side chains were added automatically. Alternate conformations with occupancies of 0.5 were resolved based on visual inspection (conformation A was selected for residues 81, 84, 91, 154, 209, 256) and conformers with highest occupancies were chosen where possible (conformation C for residue 325). The structure file contained an unknown atom which was removed. Also, the partially resolved methylene-morpholino substituent of OICR-9429 was deleted and a para-positioned methyl group added to the terminal aromatic ring, where the amide bond to the linker in the studied PROTACs is formed. The structure was protonated at pH 6.5. All water molecules, ions and other small molecules were removed from the structures.

### Protein-Protein Docking

The prepared structures of WDR5 with the modified OICR-9429 ligand and VHL with VH032 were docked onto each other using MOE’s Protein/Protein Docking (PPD) utility. VHL and VH032 were defined as “receptor”, whereas WDR5 and the modified OICR-9429 were used as “ligand”. Receptor- and ligand-sites were defined around the small molecules. Hydrophobic patch potentials were enabled, and antibody-specific options turned off. Termination criteria were set to 800 iterations and a gradient of 0.001 kcal/(mol*Å) during the final minimization. The maximum number of returned poses after pre-placement, placement and refinement was set to 100,000, 10,000 and 1000, respectively. The obtained protein-protein complexes were evaluated based on the score assigned by MOE and the distance between the atoms in the modified OICR-9429 and VH032 ligands which are linked in the final PROTACs. Ten protein/protein complexes showed a distance under 4 Å between the two critical carbon atoms, with five being ranked in the top 20% (ranks 52, 53, 59, 78 and 79). These five complexes were used as receptors for a subsequent small-molecule docking.

### Small-Molecule Docking

A protocol for small-molecule docking was verified by redocking the modified OCIR-9429 and VH032 ligands to their respective receptors. Small-molecule docking was performed with GOLD V 5.8.15^50^ using the implemented scoring function ChemPLP. For each ligand, 100 docking solutions were generated without allowing for early termination. Default values for the genetic algorithm were kept but with number of operations fixed to 100,000. Docking solutions were evaluated based on a rescoring with DrugscoreX6^51^ (V0.90, CSD potentials) and RMSD-values were calculated with fconv V 1.247.^52^ The DSX top-ranked pose of VH032 showed an RMSD-value of 0.27 Å, the modified OICR-9429 ligand achieved 0.36 Å, indicating perfect reproduction of the crystallographically observed binding modes. Degraders **8e-j** were built and minimized (MMFF94x, gradient: 0.0001 kcal/(mol·Å)) using MOE and docked to the protein-protein complexes (obtained by PPD as described above) as receptor structures. 50 docking solutions were created for each ligand-receptor pair. The non-hydrogen atoms of VH032 and the p-tolyl-phenyl-piperazinyl moiety of OICR-9429 were used as mild scaffold constraints during docking (constraint weight: 1.0) with the binding site defined as region within 6 Å of these scaffolds.

The small-molecule docking solutions were again evaluated based on a rescoring with DrugscoreX as well as the RMSD-values with respect to teh same scaffolds used as constraints.

Figures of structures were created using PyMOL Molecular Graphics System, Version 2.2.3. Schrödinger, LLC.

## Supporting information

Biochemistry supplement

Chemistry supplement

formula strings

proteomic data

## Supporting Information Availability

The Supporting Information is available free of charge on the ACS Publication website

- Supporting Information contains thermal shift data; ITC data; BRET assay data; HiBiT data; Immunoblotting data; Proteomics data; Computational data; experimental details for synthesis of all degraders, intermediates and negative controls.
- Appendix with spectra for all degraders, intermediates and negative controls.
- Molecular formula strings.
- Quantitative proteomics table.

## Abbreviations

ASH2L: absent, small or homeotic-2 like
BRET: bioluminescence Resonance Energy Transfer
DSF: differential scanning fluorimetry
HMT: Histone methyltransferase
ITC: isothermal titration calorimetry
KMT: Lysine methyltransferase
MbIIIb: Myc box IIIb
MLL: mixed lineage leukemia
PROTAC: Proteolysis targeting chimera
RBBP5: retinoblastoma binding protein 5
SET: Su(var)3-9, Enhancer-of-zeste and Trithorax
WBM: WDR5-binding
WDR5: WD-repreat containing protein 5
WIN: WDR5-interacting

## Author Contributions

A.D. synthesized all OICR-9429 based compounds, all E3 ligase linkers and negative controls. A.D. evaluated all compounds in biophysical studies (DSF and calorimetry assays) with help of J.W. and J.G. B.A. performed all cellular studies. J.W. synthesized all pyrroloimidazole based compounds. J.G., F. L. and V.D. contributed to the chemical analysis of degrader. A.K. expressed and purified recombinant protein. L.-M.B. performed the BRET assay for all compounds. M.D. performed all computational modelling. C.S. supervised computational modelling studies. M. M. S. cloned the plasmids for BRET assay. D. B.-L. supervised cloning procedure for BRET assay clones. C.H.A. and M. E. gave critical input on the project and edited the manuscript. S.H., N.B. performed the proteomic study and data analysis together with B. A., B.K. supervised proteomic study. S.K. supervised the synthesis and biophysical experiments, and E.W. supervised the cell-based assays. E.W. and S.K. designed the study. A.D., B.A., E.W. and S.K. wrote the manuscript that was approved by all authors.

## Notes

B.K. is cofounder and shareholder of msAId GmbH and OmicScouts GmbH. B.K. has no operational role in either company. The authors declare no additional competing financial interest.

## Acknowledgments.

We thank Nina Krause and Lennart Laube from the University of Frankfurt for their synthesis of the pomalidomide linkers as well as Achim Glaesmann and Dominic Löw from the University of Frankfurt for their synthetic support in the lab. We thank Masoud Vedadi for donation of WDR5 plasmids.

