## Supplementary material for "Design, Synthesis and Evaluation of WD-repeat containing protein 5 (WDR5) degraders": Biochemistry supplement

**Table of content**


1. Differential scanning fluorimetrie data
2. Isothermal titration calorimetry data
3. HiBiT data
4. Immunoblotting data
5. Proteomics data
6. Computational data
7. Synthesis of E3 ligase linker L0-L15
8. Synthesis of NanoBRET Tracer molecules 19a-c

### 1. Differential scanning fluorimetrie data

**SI Table 1:** Thermal shift experiments of pyrroloimidazole-based molecule **14**, degraders **17a-g** and negative control **21** binding to WDR5. $\Delta T_m$: thermal shift change; SD: standard deviation; R1-R3: replicate 1-3. As negative controls, DMSO and VHL ligand 1 were used. As positive control OICR-9429 was used. The concentrations for the experiments were 2 µM WDR5 and 10 µM compound.

| ID | $\Delta T_m$ [K] | SD [K] | R1 $\Delta T_m$ [K] | R2 $\Delta T_m$ [K] | R3 $\Delta T_m$ [K] |
|---|---|---|---|---|---|
| **14** | 4,2 | 0,1 | 4,4 | 4,2 | 4,1 |
| **17b** | 5,2 | 0,3 | 4,8 | 5,4 | 5,3 |
| **17a** | 7,0 | 0,4 | 7,1 | 6,5 | 7,4 |
| **17c** | 3,6 | 0,1 | 3,5 | 3,7 | 3,5 |
| **17d** | 3,4 | 0,2 | 3,4 | 3,1 | 3,6 |
| **17g** | 4,6 | 0,3 | 4,3 | 4,6 | 5,0 |
| **17f** | 4,5 | 0,1 | 4,6 | 4,3 | 4,5 |
| **17e** | 4,3 | 0,2 | 4,1 | 4,3 | 4,6 |
| **VHL** | -0,3 | 0,2 | -0,4 | -0,1 | -0,5 |
| **DMSO** | -0,1 | 0,0 | -0,1 | -0,1 | 0,0 |
| **OICR-9429** | 13,2 | 0,1 | 13,2 | 13,2 | 13,3 |
| **21** | 7,7 | 0,2 | 8,0 | 7,5 | 7,7 |

**SI Table 2:** Thermal shift experiments of OICR-9429 derived molecule **6**, intermediates **6a-c,** degraders **7a-e**, **8a-j**, **9a-c** and negative control **20**. $\Delta T_m$: thermal shift change; SD: standard deviation; R1-R3: replicate 1-3; n.d.: not determined. As negative controls, DMSO and Thalidomide, idasanutlin and VHL ligand 1 were used. As positive control OICR-9429 was used. The concentrations for the experiments were 2 µM WDR5 and 10 µM compound.

| ID | $\Delta T_m$ [K] | SD [K] | R1 $\Delta T_m$ [K] | R2 $\Delta T_m$ [K] | R3 $\Delta T_m$ [K] |
|---|---|---|---|---|---|
| **6** | 20.8 | 0.6 | 20.3 | 21.5 | 20.7 |
| **7a** | 13.6 | 0.2 | 13.4 | 13.7 | 13.6 |
| **7b** | 12.7 | 0.3 | 13.0 | 12.6 | 12.3 |
| **7c** | 9.0 | 0.5 | 8.9 | 9.5 | 8.6 |
| **7d** | 11.9 | 0.3 | 11.8 | 12.3 | 11.7 |
| **7e** | 12.5 | 0.6 | 12.8 | 11.8 | 12.8 |
| **8a** | 15.3 | 0.2 | 15.2 | 15.3 | 15.6 |
| **8b** | 7.7 | 0.5 | 7.6 | 7.4 | 8.3 |
| **8c** | 12.5 | 0.2 | 12.4 | 12.3 | 12.7 |
| **8d** | 15.6 | 0.0 | 15.6 | 15.6 | 15.5 |
| **8e** | 11.0 | 0.3 | 10.9 | 11.5 | 10.7 |
| **8f** | 10.4 | 0.7 | 10.8 | 11.0 | 9.4 |
| **8g** | 13.2 | 0.1 | 13.2 | 13.1 | 13.1 |
| **8h** | 9.7 | 2.8 | 8.2 | 7.9 | 12.9 |
| **8i** | 3.5 | 0.4 | 3.8 | 3.0 | 3.7 |
| **8j** | 14.0 | 0.0 | 14.0 | 14.0 | n.d. |
| **9a** | 0.9 | 0.2 | 1.2 | 0.8 | 0.8 |
| **9b** | 0.3 | 0.4 | 0.7 | -0.3 | 0.5 |
| **9c** | 0.7 | 0.2 | 0.6 | 0.6 | 1.0 |
| **OICR-9429** | 13.3 | 0.1 | 13.2 | 13.1 | 13.6 |
| **DMSO** | 0.0 | 0.3 | -0.1 | -0.4 | 0.3 |
| **20** | 12.2 | 0.1 | 12.2 | 12.4 | 12.1 |
| **VHL** | -0.4 | 0.1 | -0.3 | -0.5 | -0.5 |
| **Thalidomide** | 0.0 | -0.1 | 0.3 | 0.0 | -0.5 |
| **Idasanutlin** | -0.2 | 0.1 | -0.1 | -0.1 | -0.4 |
| **6a** | 5.7 | 0.4 | 6.1 | 5.2 | 5.7 |
| **6b** | 4.1 | 0.6 | 5.0 | 3.4 | 4.0 |
| **6c** | 4.2 | 0.7 | 3.4 | 4.2 | 5.1 |

### 2. Isothermal titration calorimetry data

**SI Table 3:** Thermodynamic properties of OICR-9429 derived molecule **6**, degraders **7a**, **8a**, **8e-j** and pyrroloimidazole-based inhibitor **14** and degrader **17b**. $K_d$: dissociation constant; SD: standard deviation; n: stoichiometry; ΔH: enthalpy change; T: temperature (292.15 K); ΔS: entropy change.

| ID | $K_d$ [nM] | SD [nM] | n | ΔH [kcal/mol] | TΔS [kcal/mol] |
|---|---|---|---|---|---|
| **6** | 25 | 6 | 1.0 | -9.4 | 0.8 |
| **7a** | 12 | 4 | 1.0 | -8.1 | 2.6 |
| **8a** | 41 | 9 | 1.0 | -10 | -0.2 |
| **8e** | 9 | 2 | 1.1 | -6.3 | 4.5 |
| **8f** | 6 | 2 | 1.0 | -10 | 1.1 |
| **8g** | 18 | 5 | 1.1 | -7.4 | 3.2 |
| **8h** | 12 | 4 | 1.0 | -9.3 | 1.4 |
| **8i** | 11 | 3 | 1.1 | -10 | 0.7 |
| **8j** | 33 | 5 | 1.0 | -7.9 | 2.2 |
| **14** | 125 | 34 | 1.0 | -4.9 | 4.4 |
| **17b** | 97 | 31 | 1.0 | -6.6 | 2.8 |

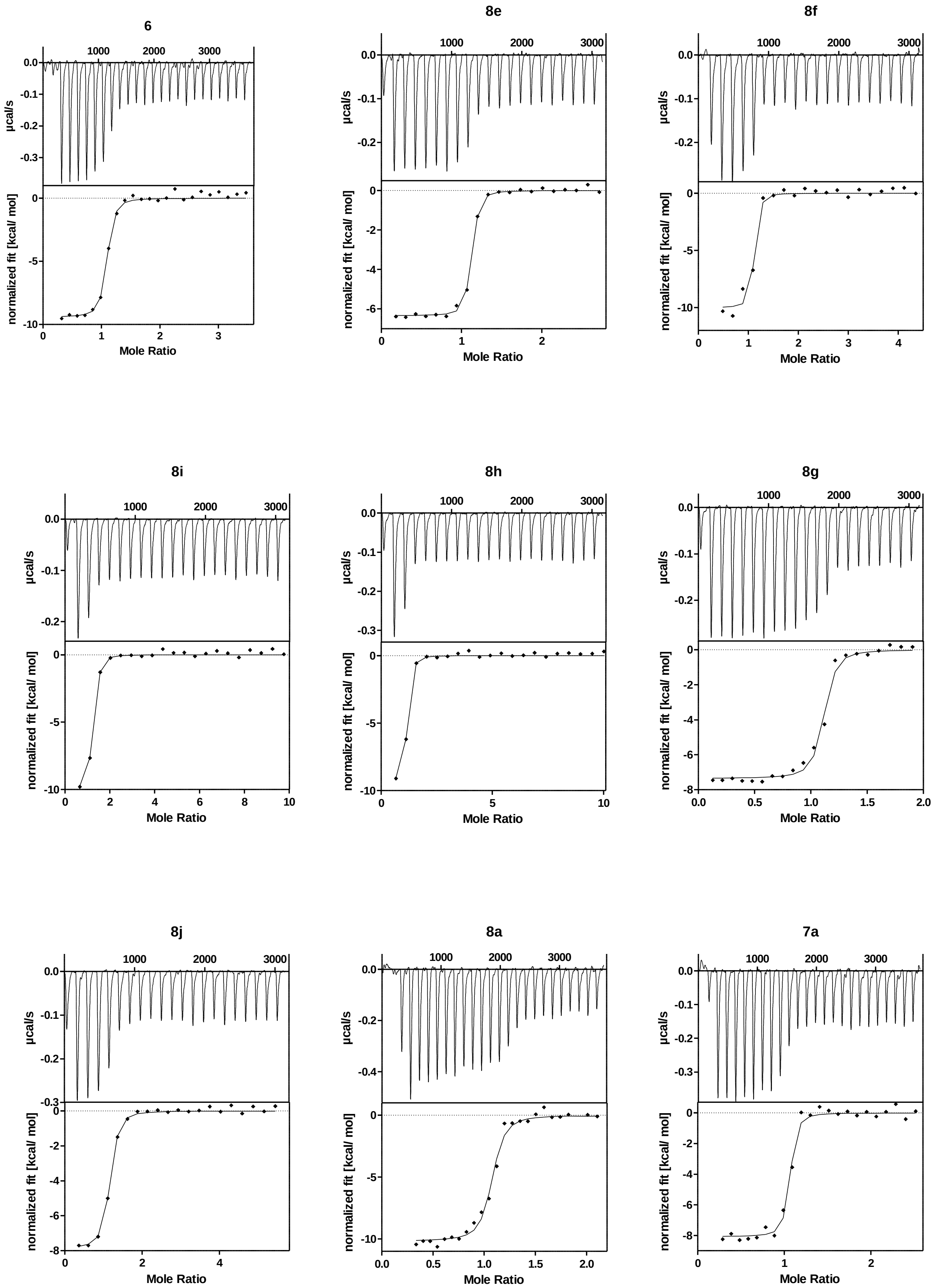


**SI Figure 1: ITC curves of OICR-9429 derived molecule** 6 **and degraders** 7a**,** 8a**,** 8e-j**.**

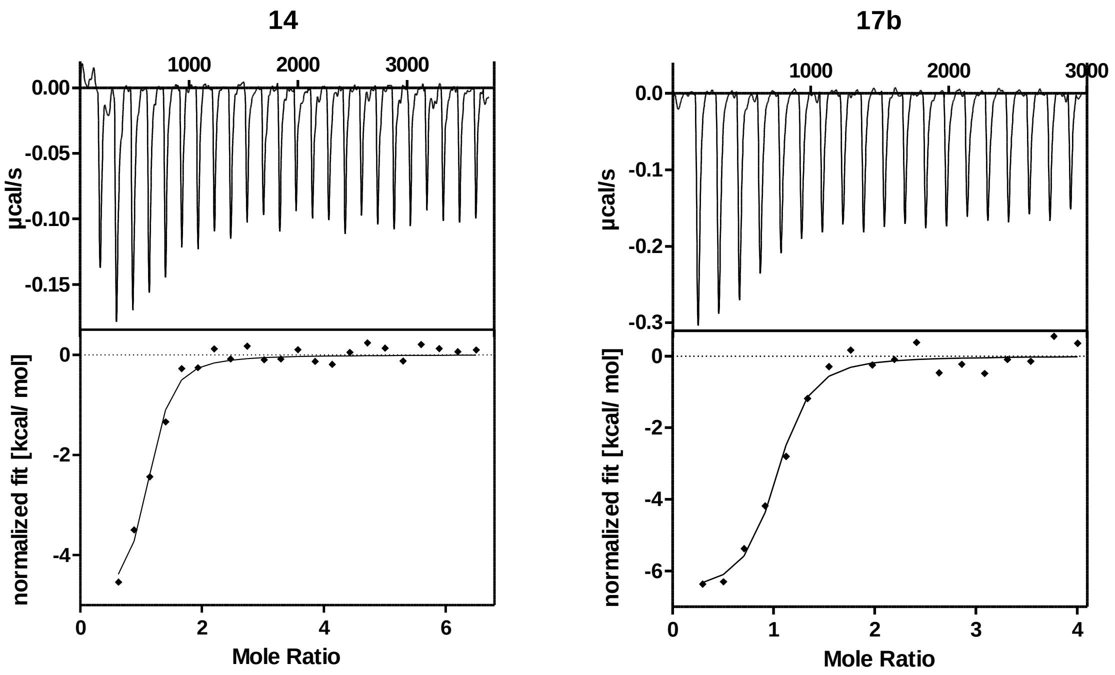


**SI Figure 2: ITC curves of pyrroloimidazole-based inhibitor** 14 **and degrader 17b.**

**3. HiBiT assay data**

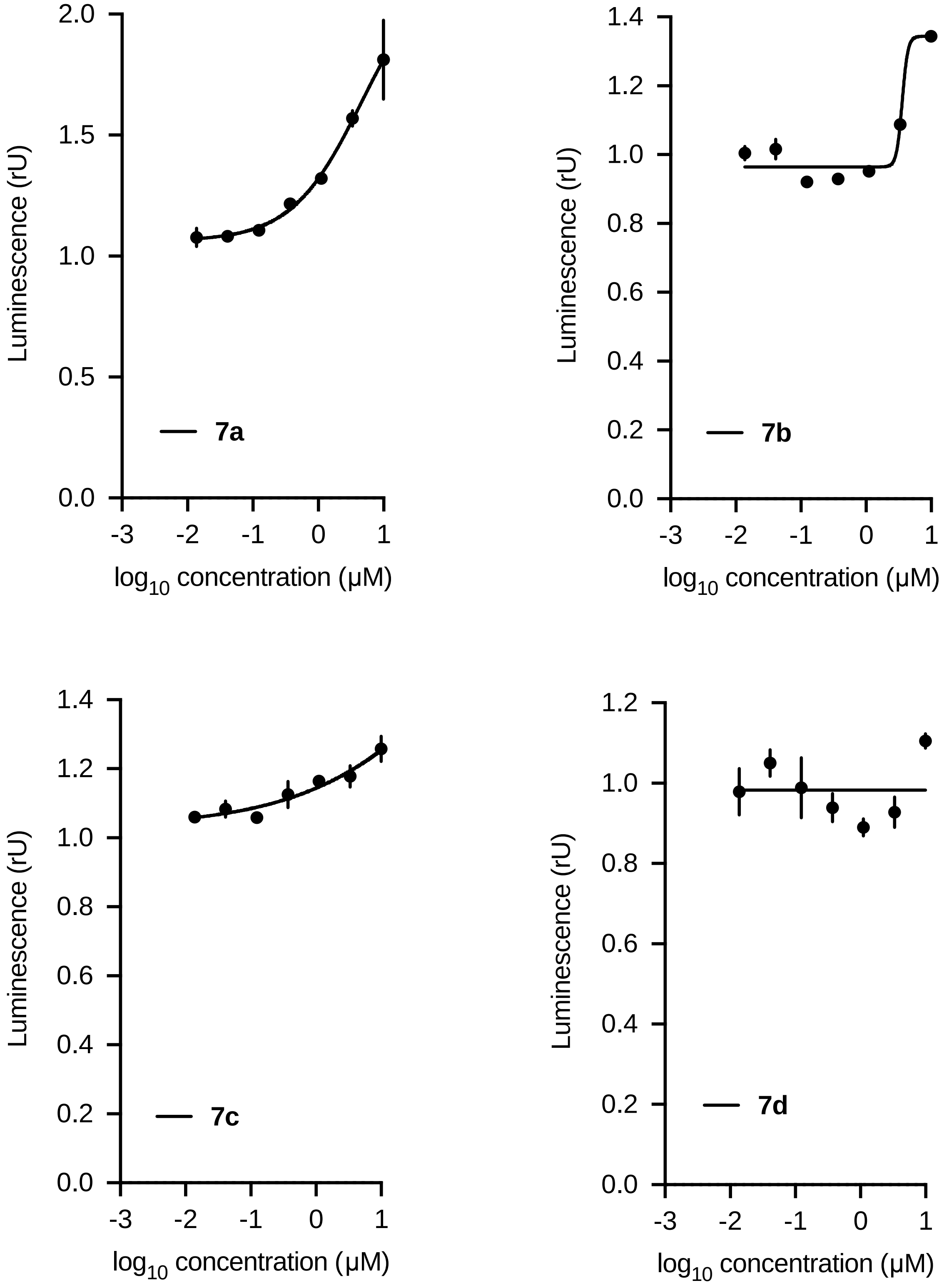

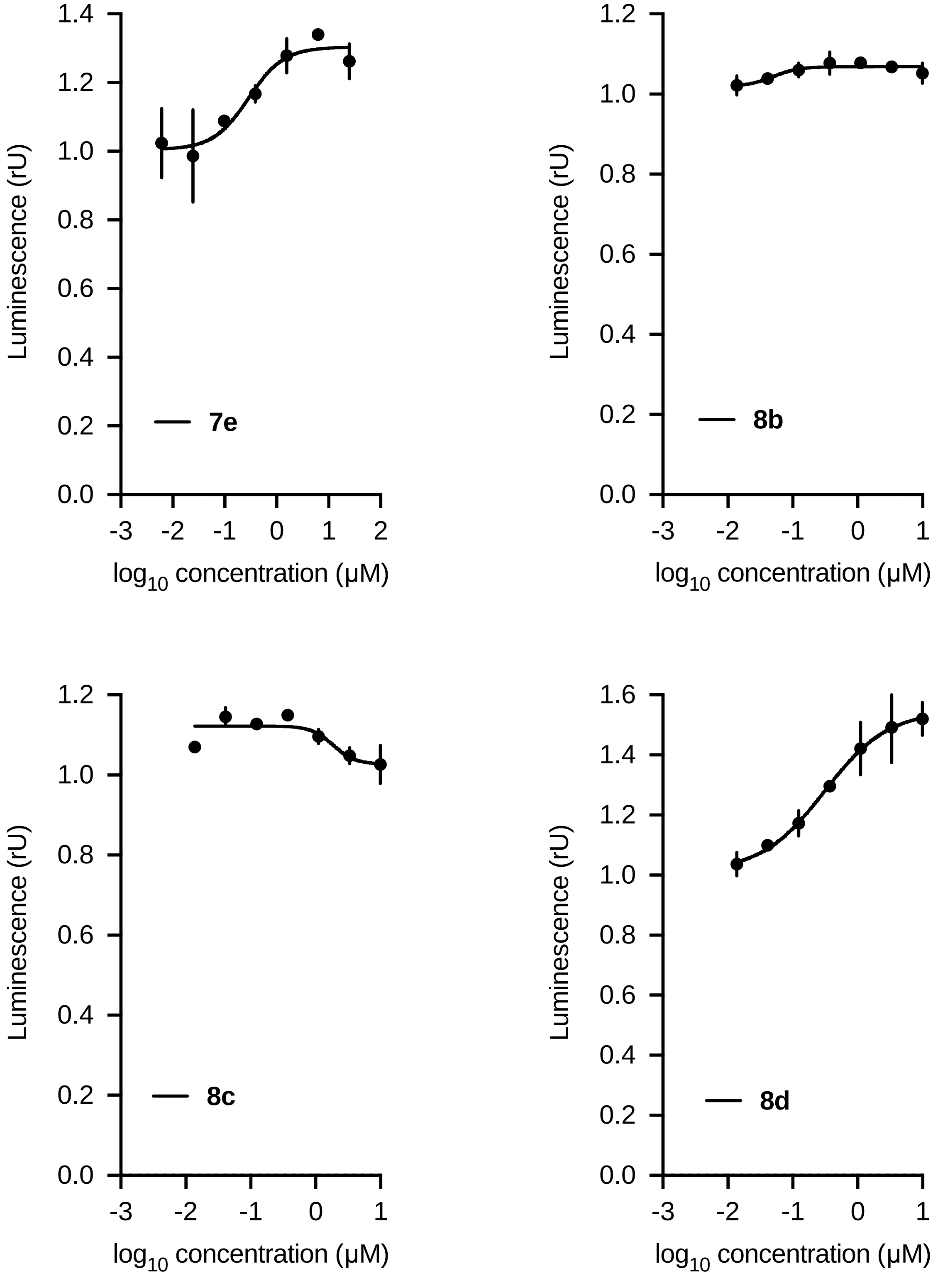

Luminescence (rU)
log10 concentration (μM)
7e
8b
8c
8d

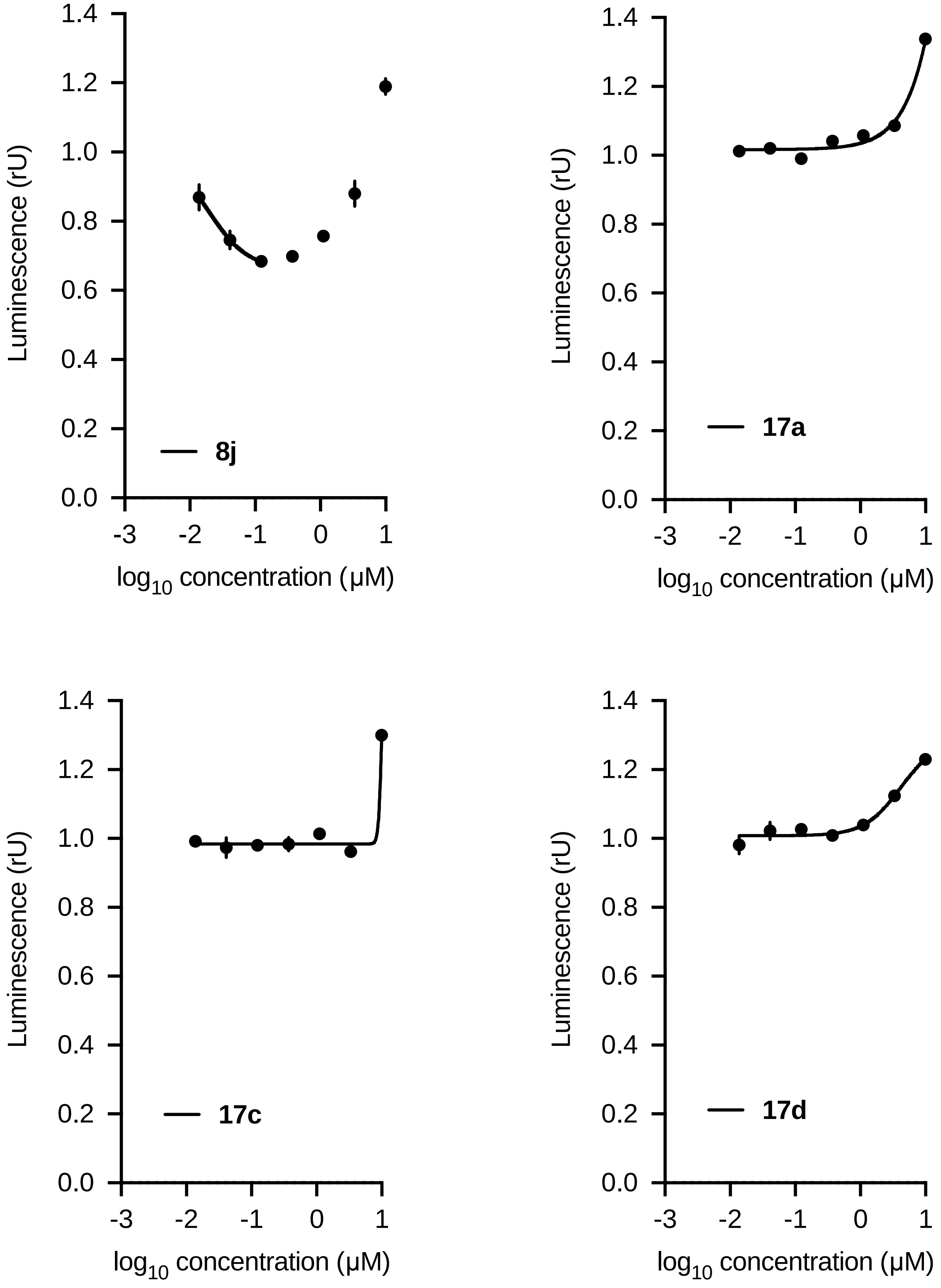
Luminescence (rU)
log10 concentration (μM)
8j
17a
17c
17d

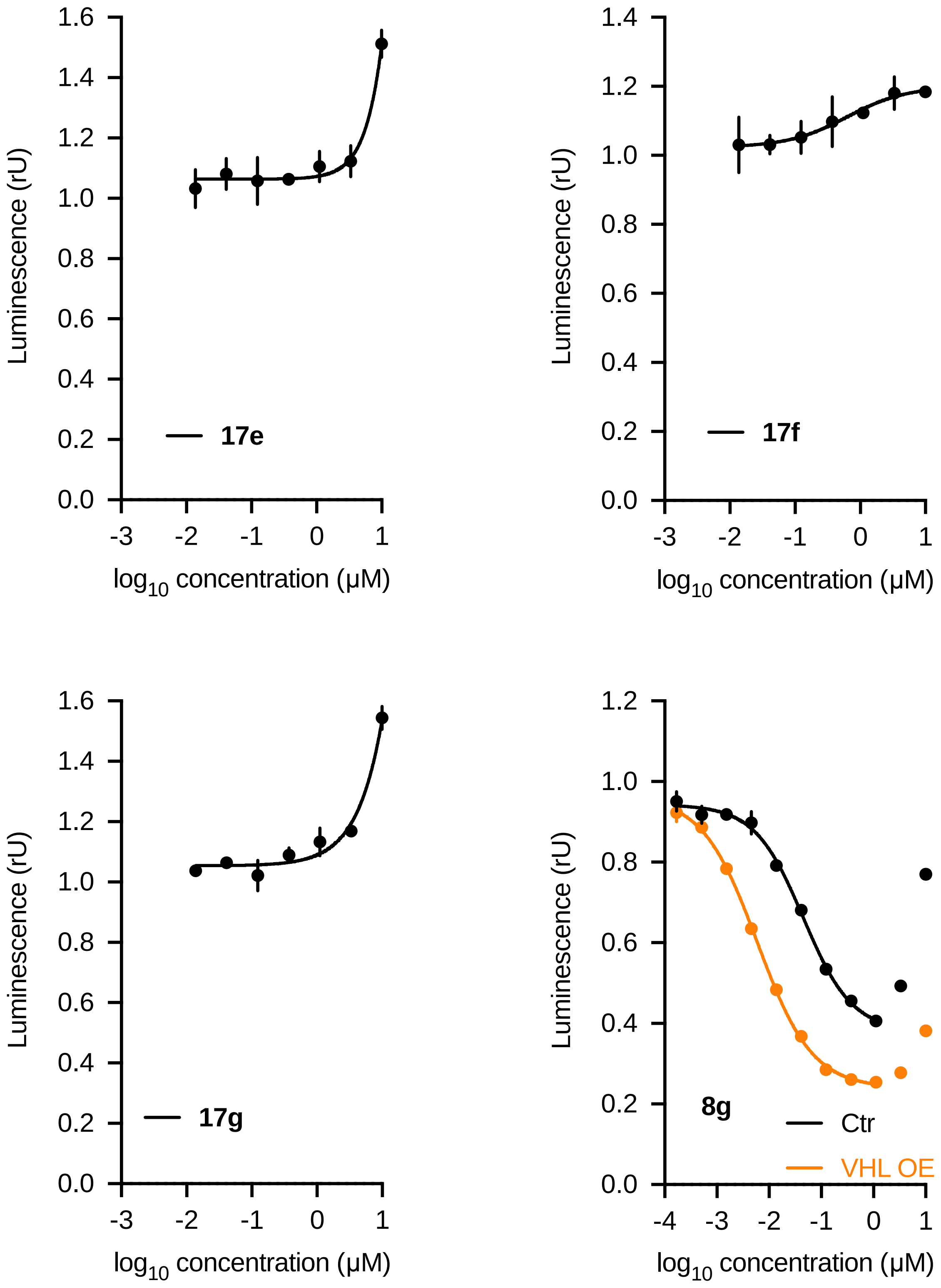


**SI Figure 3: HiBiT data:** WDR5 levels based on luciferase measurements. MV4-11$^{WDR5\text{-}HiBiT}$ cells were treated with different concentrations of **degraders** for 24 h (9h for **7e**), lysed, complemented with the second luciferase fragment (largeBiT) and measured for luciferase activity. The fit was nonlinear fit obtained from the dose-response (four parameters) equation.

### 4. Immunoblotting data

#### Immunoblot data I:

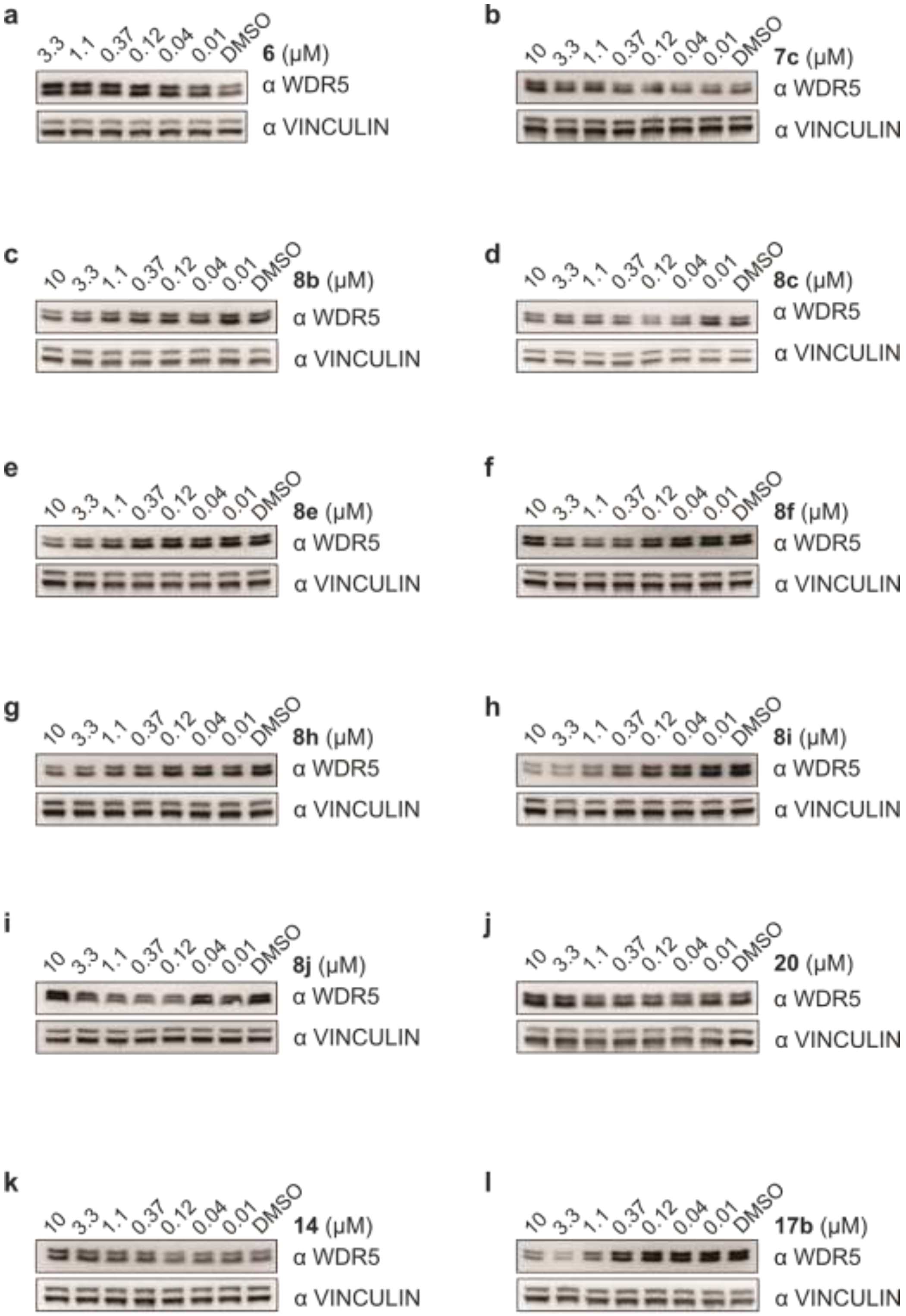


**SI Figure 4: Immunoblot data I:** Immunoblot of WDR5. MV4-11$^{WDR5\text{-}HiBiT}$ cells were treated with different concentration of PROTACs for 24 h and compared with DMSO treated cells.

**Immunoblot data II:**

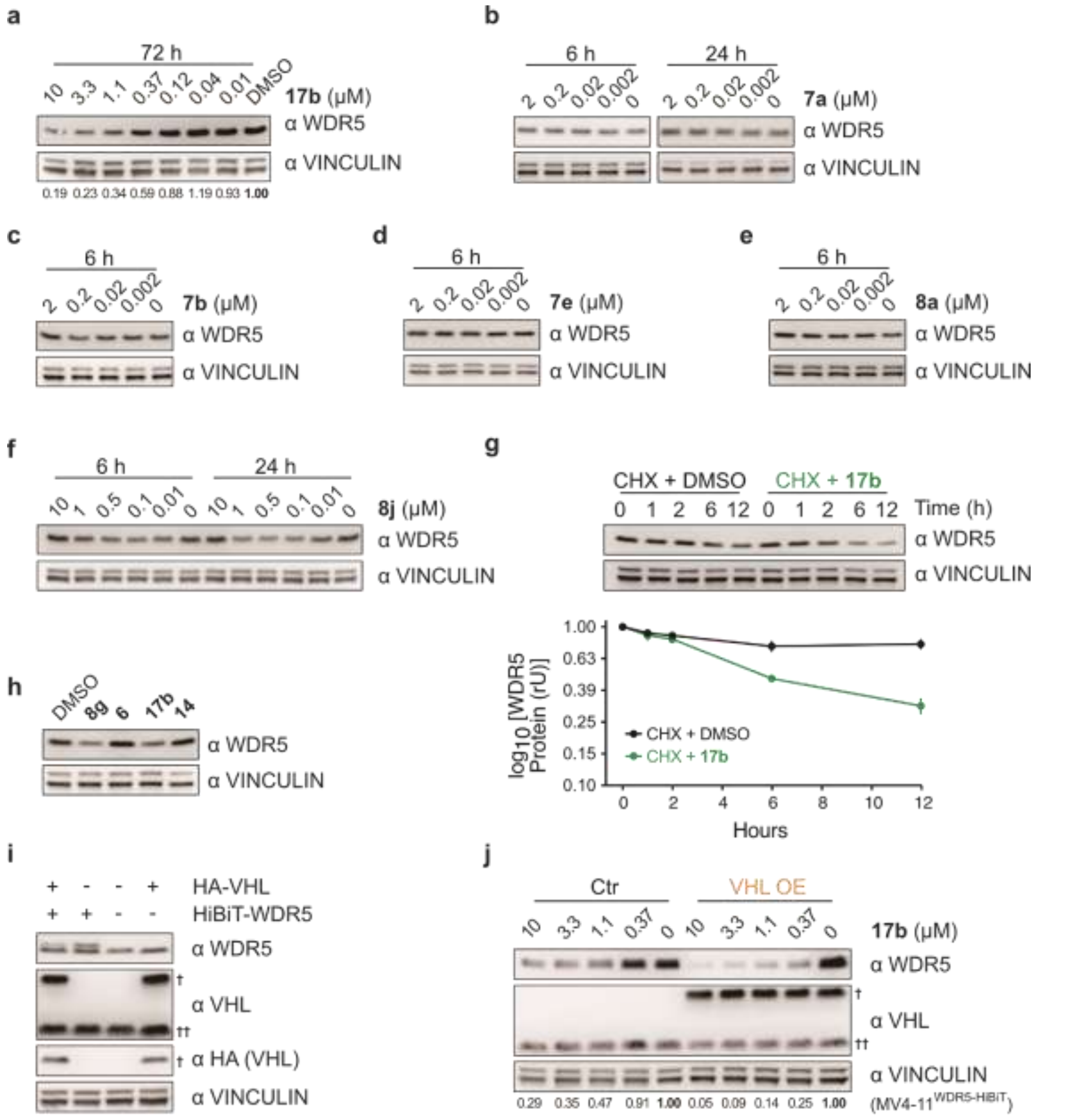


**SI Figure 5: Immunoblot data II: (a)** Immunoblot of WDR5. MV4-11 cells were treated with different concentration of **17b** for 72 h and compared with DMSO treated cells. **(b)-(f)** Immunoblot of WDR5. MV4-11 cells were treated for indicated time with **(b) 7a**, **(c) 7b**, **(d) 7e**, **(e) 8a** and **(f) 8j. (g)** Immunoblot and quantification of WDR5 levels. The WDR5 protein stability was evaluated by treating 3 µM **17b** or DMSO incubated MV4-11 cells for 0, 1, 2, 6 and 12 h with cycloheximide (CHX). The quantification is from n=2 biological replicates. **(h)** Immunoblot of WDR5. MV-11 cells were incubated with 1 µM **8g**, 1 µM **6**, 3 µM **17b** and 3 µM **14** for 24h. This immunoblot corresponds to the RT-PCR from figure 4d. **(i)** Immunoblot of WDR5 and VHL. HA tagged VHL was stably expressed in MV4-11$^{\text{WDR5-HiBiT}}$ cells and MV4-11 cells. † overexpressed VHL; †† endogenous VHL. **(j)** Immunoblot of WDR5 and VHL. MV4-11$^{\text{WDR5-HiBiT}}$ (Ctr) and MV4-11$^{\text{WDR5-HiBiT/VHL}}$ (VHL OE) were treated for 24h with various concentrations of **17b**. † overexpressed VHL; †† endogenous VHL

### 5. Proteomics data

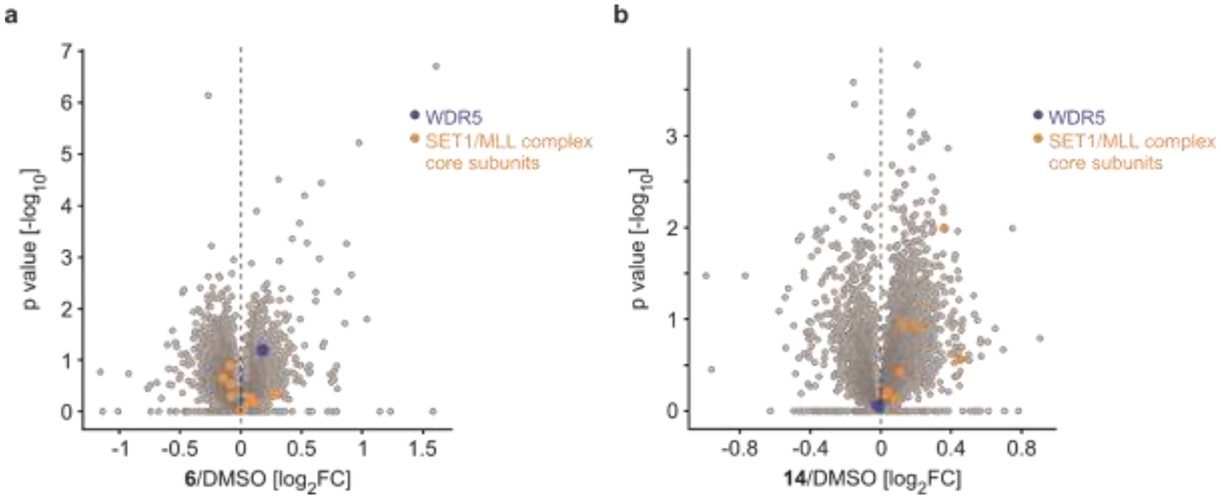


**SI Figure 6: Proteomics data: (a) and (b)** Volcano plot showing global proteomics change. MV4-11 cells were treated with 1 µM **6 (a)** or 5 µM **14 (b)** for 9 hours and lysates were analyzed by quantitative proteomics. WDR5 (blue) and other SET1/MLL complex core subunits: KMT2A, KTM2B, KTM2C, KTM2D, SETD1A, RBBP5, ASH2L and DPY30 (orange) were labelled.

### 6. Computational data

Computational docking studies were conducted to obtain a structural model compatible with the binding of the identified active degraders (consisting of the VHL-binding moiety VH032 linked to a modified OICR-9429, a known WDR5 inhibitor) to the known binding sites of the WDR5 and VHL recognition units. Compounds with linkers ranging from two to six methylene groups (i.e. ethyl- to hexyl–linkers) proved to be effective as degraders. Accordingly, a structural model had to be identified which can explain the binding of the ethyl-linked ligand as well as that of the hexyl-linked compound. Crystal structures of WDR5 in complex with OICR-9429 (PDB: 4ql1, resolution: 1.50 Å) and VHL in complex with VH032 (PDB: 4w9h, resolution: 2.10 Å) were prepared for protein/protein docking, as described in the methods section. OICR-9429 was modified by replacing its partially resolved methylene-morpholino moiety with a methyl-group, which served as linkage site towards VH032 (see **SI Fig. 7**).

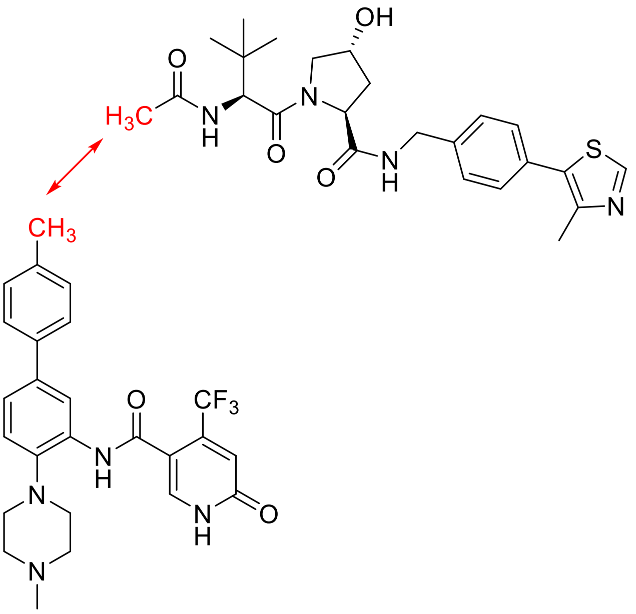


**SI Figure 7:** Structures of OICR-9429 (bottom) and VH032 (top) used during protein/protein docking. The distance between the carbon atoms of the highlighted methyl groups was evaluated.

Structural overlays of published WDR5 and VHL crystal structures are shown in **SI Figure 8**. They illustrate that the VHL and WDR5 structures show only little fluctuations and align very well with their counterparts in the docking solution. The secondary structure elements are perfectly superimposed, and minor differences are seen only in some WDR5 loop regions distant from the binding site and in the conformations of a few long and polar side chains of VHL and WDR5. In summary, the surfaces of VHL and WDR5 are sufficiently well conserved to be amenable to the applied protein/protein docking and refinement procedure.

Protein/protein docking was performed in the Molecular Operating Environment (MOE) with VHL and VH032 defined as the “receptor” and WDR5 with modified OICR-9429 acting as “ligand”. The small molecules were chosen to define the binding sites. The 405 obtained protein/protein complexes were ranked by their score assigned by MOE, as well as by the distance between the atoms serving as linkage sites (as indicated in **SI Fig. 7**).

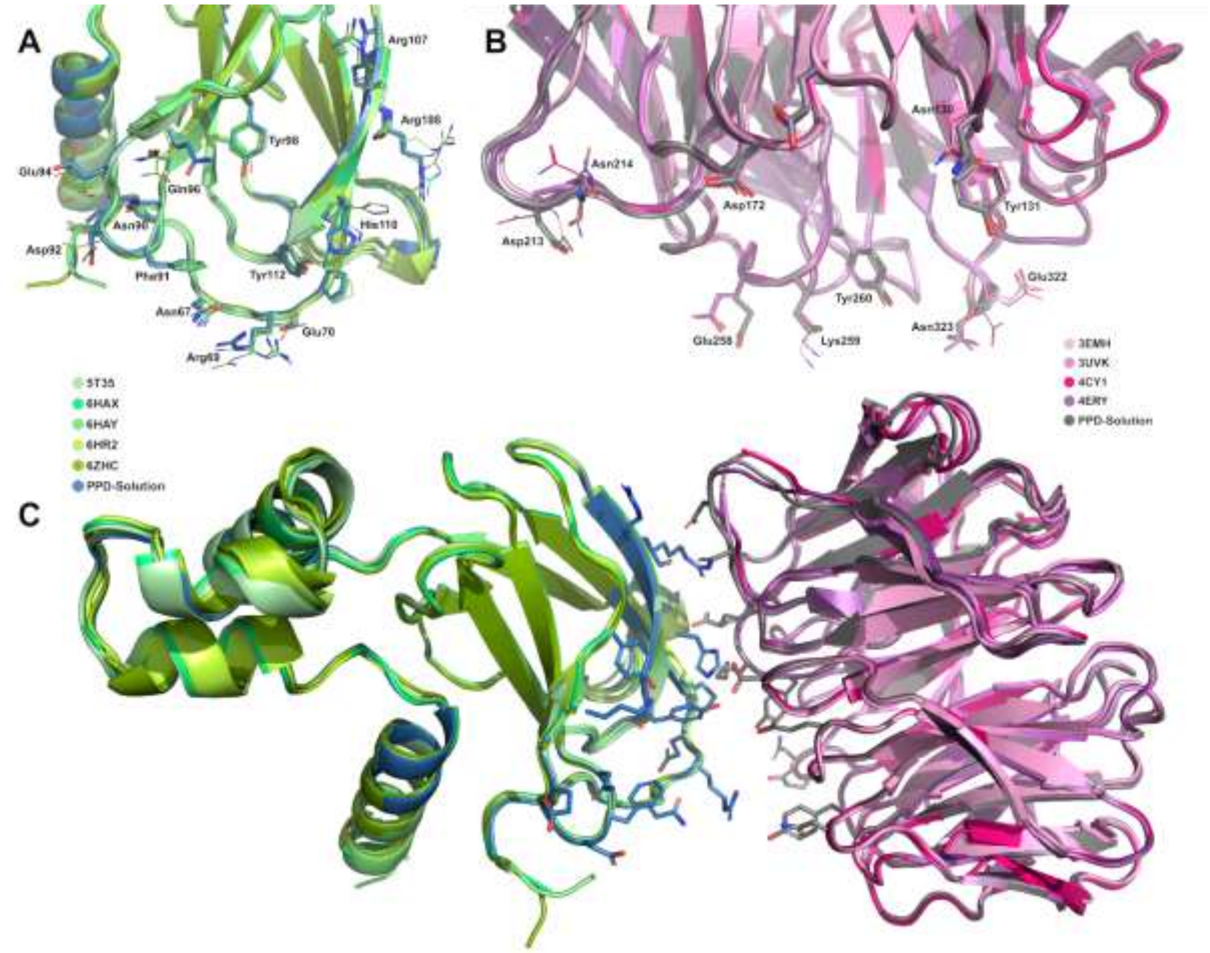


**SI Figure 8:** Comparison of VHL and WDR5 crystal structures with protein/protein docking solution. **A:** Structural overlay of VHL from the protein/protein docking solution (blue) with all VHL crystal structures complexed with a PROTAC ligand and a target protein available in the PDB at the time of writing (shown in green colors). The five structures (PDB codes: 5T35, 6HAX, 6HAY, 6HR2, 6ZHC) were aligned on the VHL structure from the best-ranked of the compatible protein/protein docking solutions. The structures are illustrated in cartoon representation; side chains of residues involved in protein/protein contacts in any of the crystal structures and the compatible protein/protein docking solutions are shown explicitly. **B:** Structural overlay of WDR5 from the protein/protein docking solution (gray) with the four best-resolved crystal structures of WDR5 with peptide ligands from the PDB (3EMH, 3UVK, 4CY1, 4ERY; shown in reddish colors). The four structures were aligned on the WDR5 structure from the best-ranked of the compatible protein/protein docking solution. The structures are illustrated in cartoon representation; side chains of residues involved in protein/protein contacts in the compatible protein/protein docking solutions are shown explicitly. **C:** Overview of the aligned VHL and WDR5 structures from panels A and B in the orientation of the protein/protein docking solution. Only the side chains of the docking solution are shown as sticks.

Since the compound with the shortest linker still showed effective degradation, the resulting protein/protein complex must be capable of fitting both recognition units in their known binding modes while maintaining a linkable distance for the ethyl-linker. Accordingly, the distance between the two linkage sites was used as the primary selection criterion. Ten protein/protein complexes showed a distance under 4 Å between the two critical carbon atoms (**SI Fig. 9a**), with five being ranked in the top 20% (ranks 52, 53, 59, 78 and 79). These five complexes were used as receptors for a subsequent small-molecule docking.

To check for suitable binding modes of all degraders with carbon chain linkers, small-molecule docking was performed with GOLD, using the selected protein/protein complex as receptor. Parts of the OCIR-9429 and VH032 scaffolds in this structure were used to define the binding site and to provide mild scaffold constraints to guide the recognition units into their known binding sites (**SI Fig. 9b**).

The degraders **8e-j** were docked to each of the five selected protein/protein complexes and evaluated based on a rescoring with DrugscoreX and RMSD-values with respect to the crystallized binding modes of VH032 and OICR-9429 (**SI Fig. 9b**).

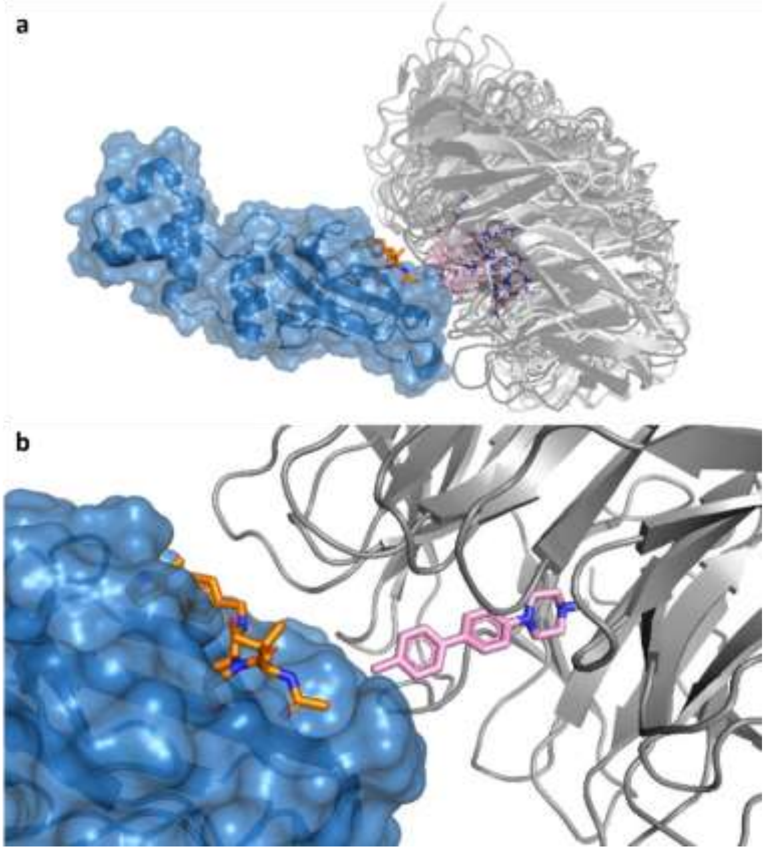


**SI Figure 9:** Structure of the protein/protein docking solutions. **a:** The ten protein/protein complexes with distances below 4 Å between the two critical carbon atoms. VHL (blue) and VH032 (orange) form a complex with WDR5 (grey) and a modified OICR-9429 ligand (pink). The structures were all superimposed on VHL, of which only on conformation is shown (rank 52). **b:** Scaffolds of VH032 and OICR-9429 used as constraints during docking and as reference structures for RMSD calculations of small-molecule docking solutions are shown in ball-and-stick representation in pink and orange for OICR-9429 and VH032, respectively. Only one protein/protein docking solution (rank 52) is shown.

Results of the DSX top-ranked docking poses for each protein/protein complex of interest are shown in **SI Table 4** as well as in **SI Figure 10-17**.

**SI Table 4:** RMSD-values and scores of the DSX top-ranked docking solutions.

**Protein/Protein Rank 52**

| Ligand | DSX-Score | RMSD [Å] OICR-9429 | RMSD [Å] VH032 |
|---|---|---|---|
| **8e** | -330.3 | 0.60 | 0.59 |
| **8f** | -331.8 | 0.62 | 0.37 |
| **8g** | -337.4 | 0.51 | 0.75 |
| **8h** | -331.0 | 0.55 | 0.83 |
| **8i** | -328.1 | 0.56 | 1.33 |
| **8j** | -327.1 | 1.63 | 3.90 |

**Protein/Protein Rank 53**

| Ligand | DSX-Score | RMSD [Å] OICR-9429 | RMSD [Å] VH032 |
|---|---|---|---|
| **8e** | -351.3 | 1.03 | 0.70 |
| **8f** | -355.1 | 0.62 | 0.85 |
| **8g** | -350.0 | 0.72 | 0.71 |
| **8h** | -340.6 | 0.73 | 0.68 |
| **8i** | -349.6 | 0.64 | 1.32 |
| **8j** | -300.6 | 1.09 | 1.70 |

**Protein/Protein Rank 59**

| Ligand | DSX-Score | RMSD [Å] OICR-9429 | RMSD [Å] VH032 |
|---|---|---|---|

| | | | |
|---|---|---|---|
| **8e** | -346.3 | 0.53 | 0.55 |
| **8f** | -352.3 | 0.58 | 0.80 |
| **8g** | -341.5 | 0.44 | 0.81 |
| **8h** | -340.8 | 0.62 | 0.39 |
| **8i** | -339.9 | 0.81 | 0.86 |
| **8j** | -306.8 | 1.03 | 1.61 |

**Protein/Protein Rank 78**

| Ligand | DSX-Score | RMSD [Å] OICR-9429 | RMSD [Å] VH032 |
|---|---|---|---|
| **8e** | -352.6 | 0.73 | 0.38 |
| **8f** | -353.1 | 0.49 | 0.47 |
| **8g** | -347.3 | 0.65 | 0.76 |
| **8h** | -333.3 | 0.74 | 0.71 |
| **8i** | -332.9 | 0.85 | 1.01 |
| **8j** | -330.6 | 1.59 | 3.64 |

**Protein/Protein Rank 79**

| Ligand | DSX-Score | RMSD [Å] OICR-9429 | RMSD [Å] VH032 |
|---|---|---|---|
| **8e** | -318.3 | 0.85 | 0.42 |
| **8f** | -337.9 | 0.67 | 0.75 |
| **8g** | -335.6 | 0.47 | 1.56 |
| **8h** | -317.5 | 0.49 | 1.41 |
| **8i** | -308.0 | 0.62 | 1.81 |
| **8j** | -297.4 | 0.48 | 1.39 |

a

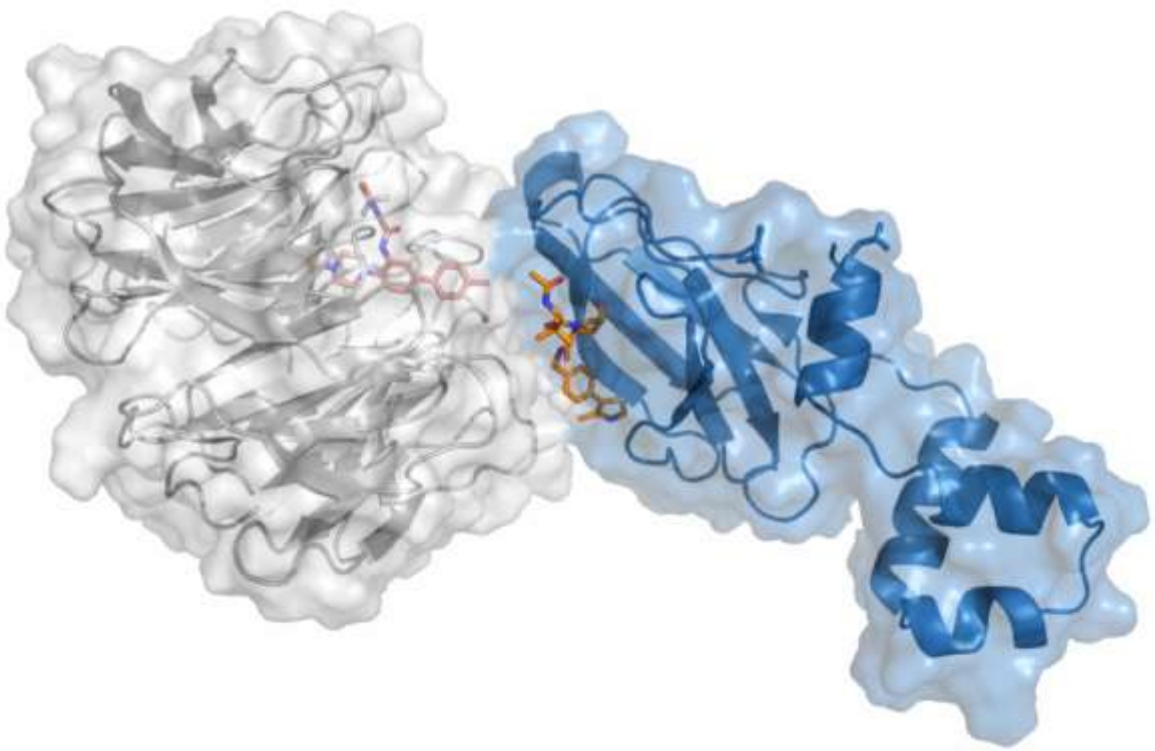

b

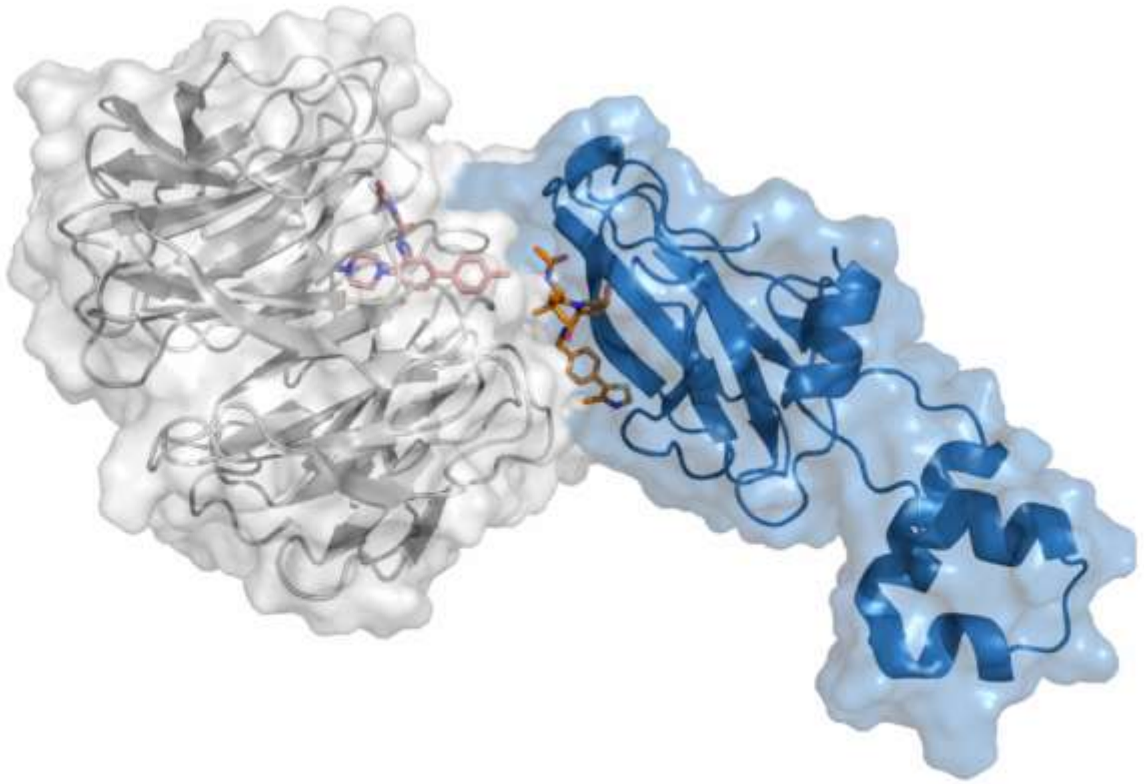

**SI Figure 10:** Structure of the protein/protein docking solutions of **a:** rank 52 and **b:** rank 53. VHL (blue) and VH032 (orange) form a complex with WDR5 (grey) and a modified OICR-9429 ligand (pink).

a

b

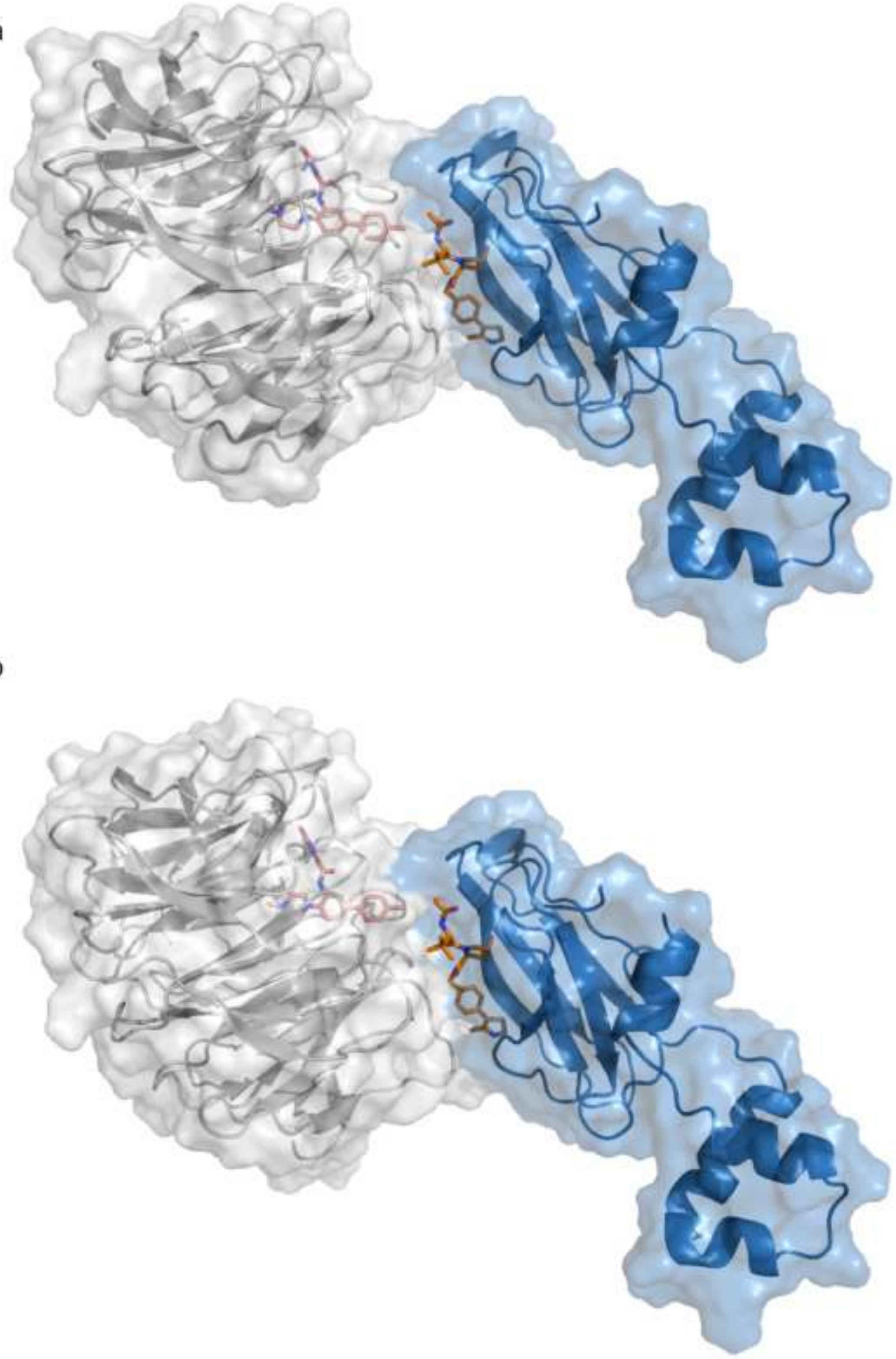

**SI Figure 11:** Structure of the protein/protein docking solutions of **a:** rank 59 and **b:** rank 78. VHL (blue) and VH032 (orange) form a complex with WDR5 (grey) and a modified OICR-9429 ligand (pink).

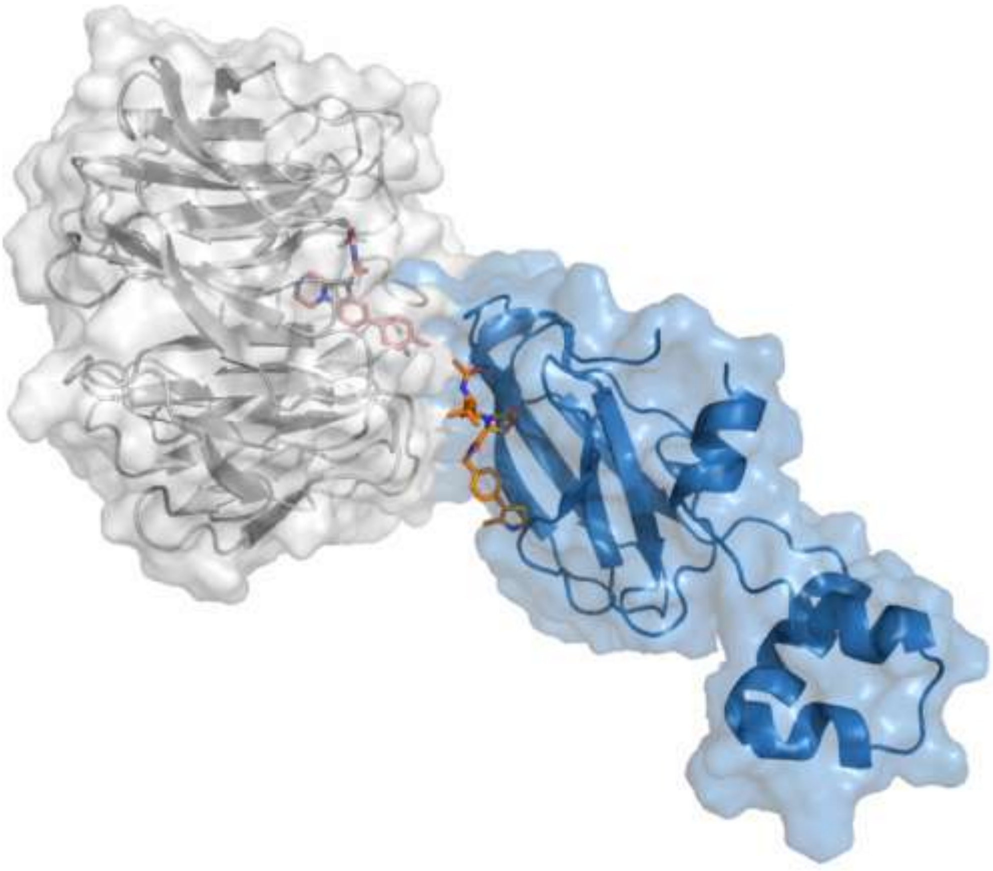

**SI Figure 12:** Structure of the protein/protein docking solutions of rank 79. VHL (blue) and VH032 (orange) form a complex with WDR5 (grey) and a modified OICR-9429 ligand (pink).

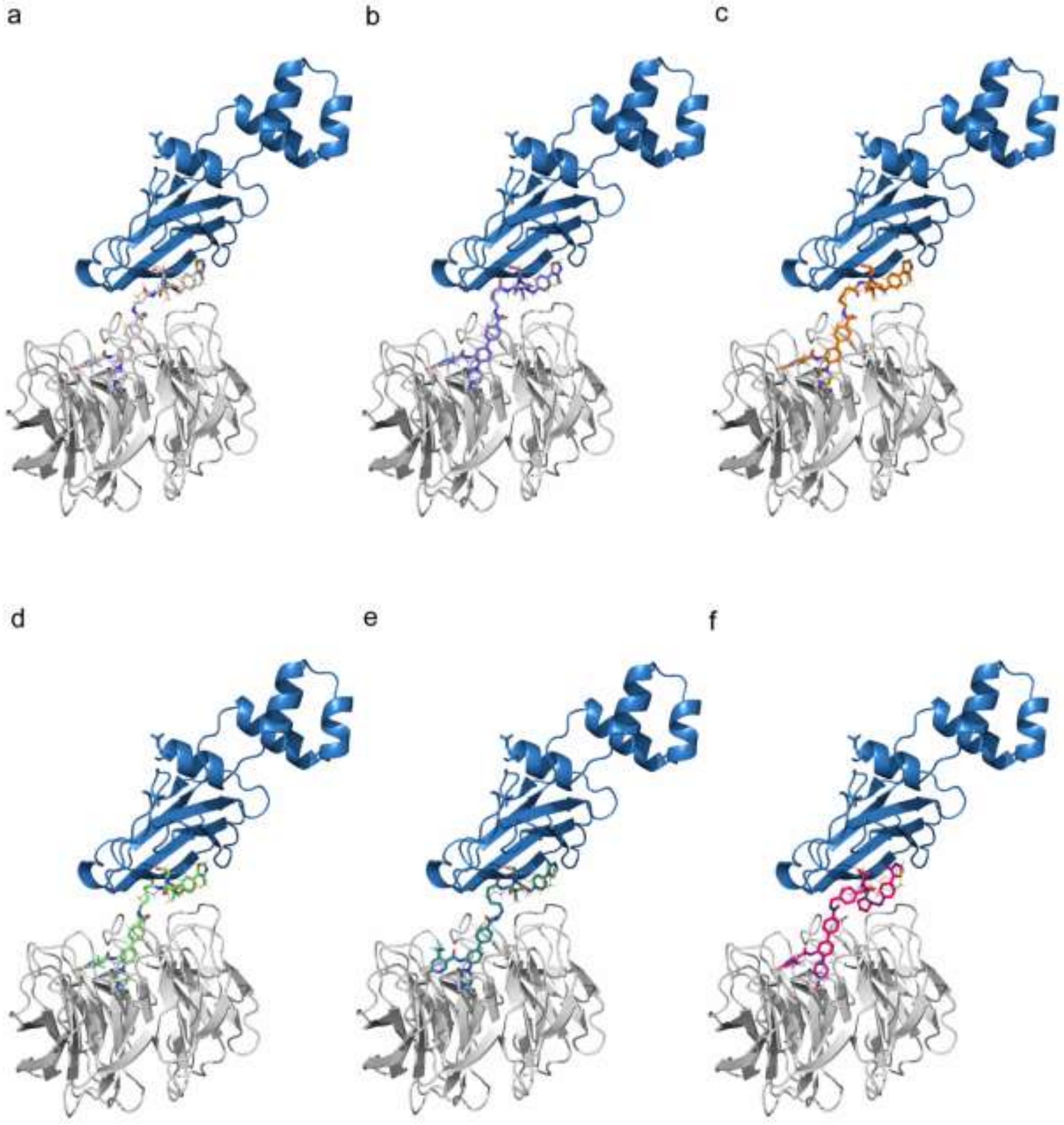


**SI Figure 13:** Protein-protein docking complex rank 52 with docking solutions for molecules **a: 8e**, **b: 8f**, **c: 8g**, **d: 8h**, **e: 8i** and **f: 8j**. VHL (blue) forms a complex with WDR5 (grey) and the coloured heterobifunctional degrader molecules. A modified OICR-9429 ligand (shown in lines in pink) and VH032 (shown in lines in orange) indicate the ligand positions of the protein-protein docking.

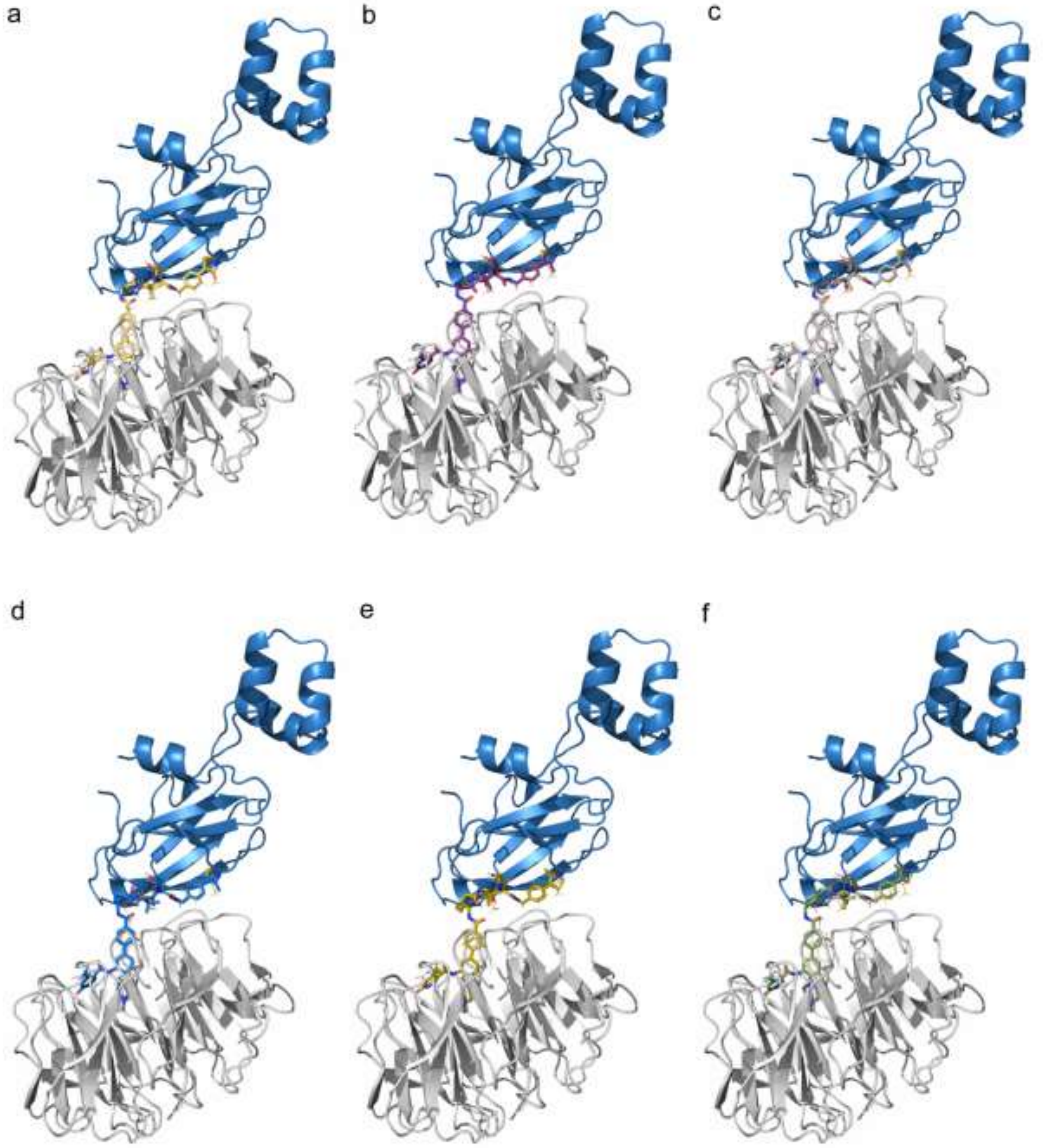


**SI Figure 14:** Protein-protein docking complexes of rank 53 for molecules **a: 8e**, **b: 8f**, **c: 8g**, **d: 8h**, **e: 8i** and **f: 8j**. VHL (blue) forms a complex with WDR5 (grey) and the coloured heterobifunctional degrader molecules. A modified OICR-9429 ligand (shown in lines in pink) and VH032 (shown in lines in orange) indicate the ligand positions of the protein-protein docking.

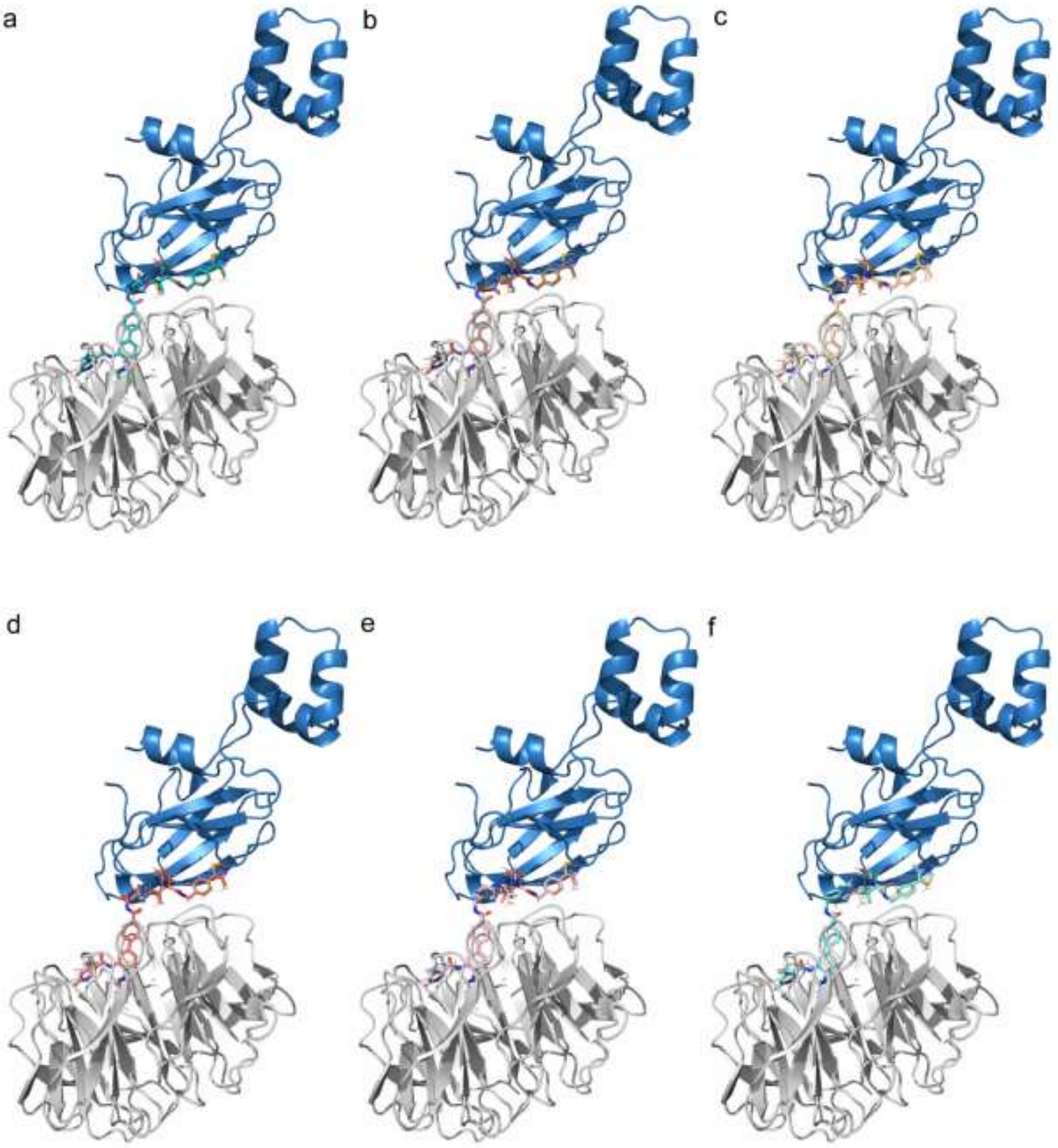


**SI Figure 15:** Protein-protein docking complexes of rank 59 for molecules **a: 8e**, **b: 8f**, **c: 8g**, **d: 8h**, **e: 8i** and **f: 8j**. VHL (blue) forms a complex with WDR5 (grey) and the coloured heterobifunctional degrader molecules. A modified OICR-9429 ligand (shown in lines in pink) and VH032 (shown in lines in orange) indicate the ligand positions of the protein-protein docking.

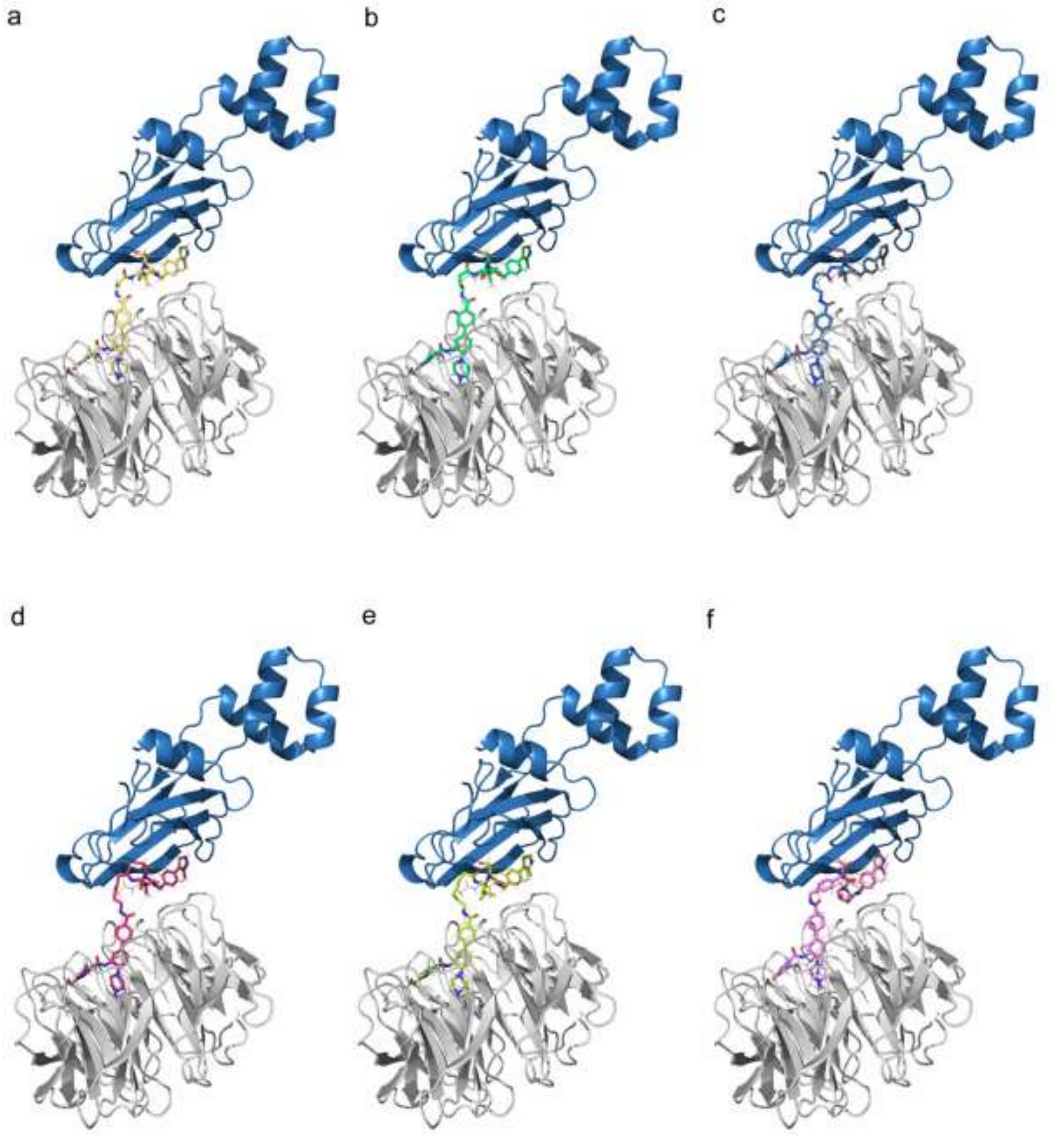


**SI Figure 16:** Protein-protein docking complexes of rank 78 for molecules **a: 8e**, **b: 8f**, **c: 8g**, **d: 8h**, **e: 8i** and **f: 8j**. VHL (blue) forms a complex with WDR5 (grey) and the coloured heterobifunctional degrader molecules. A modified OICR-9429 ligand (shown in lines in pink) and VH032 (shown in lines in orange) indicate the ligand positions of the protein-protein docking.

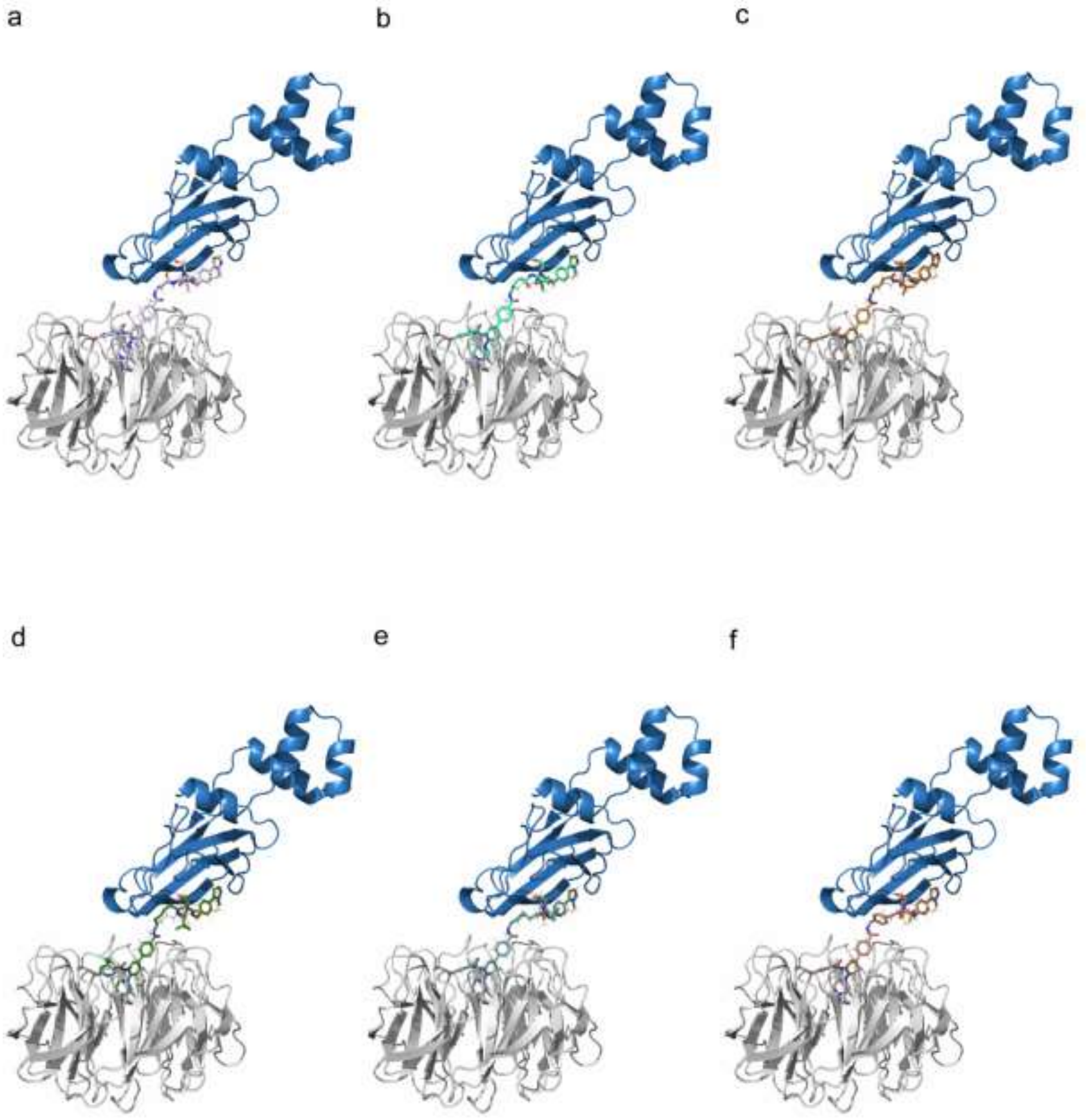


**SI Figure 17:** Protein-protein docking complexes of rank 79 for molecules **a: 8e**, **b: 8f**, **c: 8g**, **d: 8h**, **e: 8i** and **f: 8j**. VHL (blue) forms a complex with WDR5 (grey) and the coloured heterobifunctional degrader molecules. A modified OICR-9429 ligand (shown in lines in pink) and VH032 (shown in lines in orange) indicate the ligand positions of the protein-protein docking.

### 7. Synthesis of E3 ligase linker

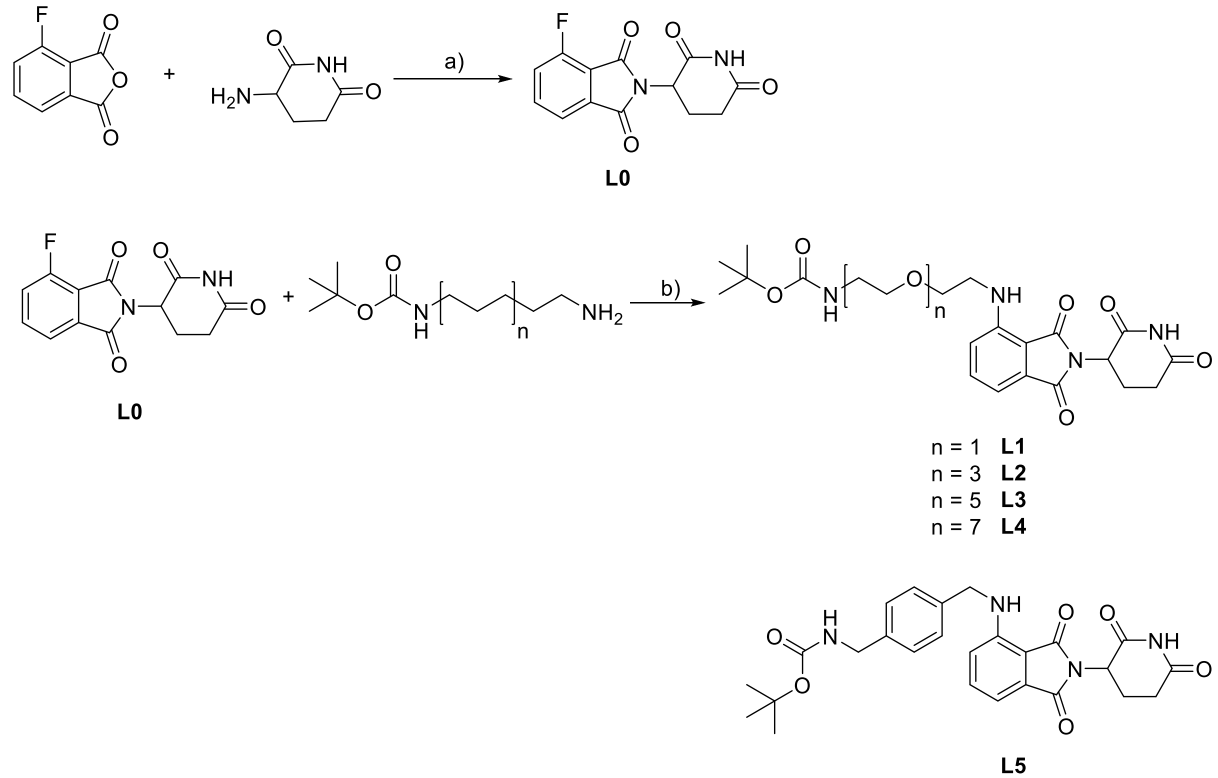


**SI Scheme 1**: Synthesis route of E3 ligase ligand linker **L1-L5**: a) pyridine, 90 °C, 18 h; b) DIEA, DMF, 80°C, 16 h.

#### Synthesis of 2-(2,6-dioxopiperidin-3-yl)-4-fluoroisoindoline-1,3-dione (L0)

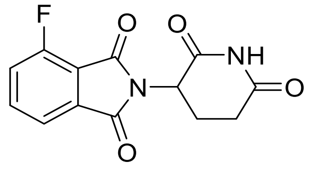

2.00 g (12.0 mmol, 1.00 eq) 4-Fluoroisobenzofuran-1,3-dione and 1.98 g (12.0 mmol, 1.00 eq) 3-Aminopiperidine-2,6-dione were dissolved in 15 mL pyridine and stirred for 18 h at 110 °C. The solution was cooled to rt and the pyridine was removed under reduced pressure. The crude solution was dissolved in water and DCM. The aqueous phase was separated and extracted 6x with DCM. The combined organic phases were washed with 1 M HCl and saturated NaCl solution. The combined organic phases were dried over $MgSO_4$, filtered and excess solvent was removed under reduced pressure. The crude product was purified by column chromatography to obtain 1.13 g, 4.09 mmol, 33% of a yellow solid.

Yield: 1.13 g, 4.09 mmol, 33% of a yellow solid.

$R_f$ (5% MeOH/ $CH_2Cl_2$): 0.53.

ESI: (calculated): [$M+2H^+$] 278.06 g/mol

(found): [$M+2H^+$] 278.96 g/mol.

$^1$H NMR (250 MHz, DMSO) δ = 11.13 (s, 1H, 17-H), 7.95 (ddd, $^3J$ = 8.3 Hz, $^3J$ = 7.5 Hz, $^4J$ = 4.6 Hz, 1H), 7.80 – 7.70 (m, 2H), 5.18 – 5.10 (m, 1H), 2.96 – 2.81 (m, 1H), 2.65 – 2.43 (m, 2H), 2.06 (m, 1H) ppm.

**Synthesis of tert-butyl (2-(2-((2-(2,6-dioxopiperidin-3-yl)-1,3-dioxoisoindolin-4-yl)amino)ethoxy)ethyl)carbamate (L1)**

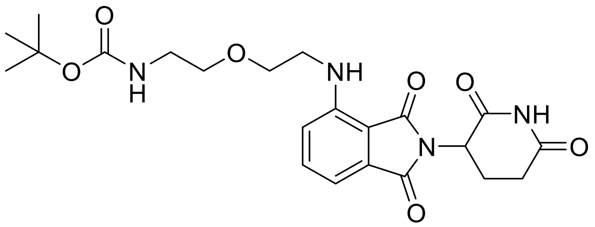

150 mg (543 µmol, 1.00 eq) 2-(2,6-dioxopiperidin-3-yl)-4-fluoroisoindoline-1,3-dione, 111 mg (543 µmol, 1.00 eq) tert-butyl (2-(2-aminoethoxy)ethyl)carbamate and 189 µL (1.09 mmol, 2.00 eq) DIEA were dissolved in 5 mL NMP and stirred at 80 °C for 16 h. The solution was cooled to rt and diluted with EA and saturated $NaHCO_3$ and saturated NaCl solution. The mixture was extracted 3x with EA, dried over $MgSO_4$, filtered and excess solvent was removed under reduced pressure. The crude product was purified by Flash Chromatography to obtain 102 mg, 177 µmol 33% of a yellow solid.

Yield: 133 mg, 288 µmol, 53% of a yellow solid.

$R_f$ (30% Cyclohexane/EA): 0.46.

ESI: (calculated): [$M+Na^+$] 483.19 g/mol

(found): [$M+Na^+$] 483.23 g/mol.

HPLC: RT = 12.8 min (254 nm, 100%).

$^{1}H$ NMR (600 MHz, MeOD) δ = 7.52 (t, $^{3}J$ = 7.8 Hz, 1H), 7.06 – 7.02 (m, 2H), 5.05 (dd, $^{3}J$ = 12.4 Hz, $^{4}J$ = 5.3 Hz, 1H), 3.68 (t, $^{3}J$ = 5.2 Hz, 2H), 3.68 – 3.46 (m, 4H), 3.23 (t, $^{3}J$ = 5.5 Hz, 2H), 2.85 (ddd, $^{2}J$ = 19.2 Hz, $^{3}J$ = 14.3 Hz, $^{4}J$ = 5.2 Hz, 1H), 2.78 – 2.66 (m, 2H), 2.16 – 2.06 (m, 1H), 1.41 (s, 9H) ppm.

$^{13}C$ NMR (126 MHz, MeOD) δ = 174.6, 171.5, 170.7, 169.2, 158.4, 148.2, 137.2, 133.8, 118.3, 112.1, 111.2, 80.1, 70.9, 70.4, 50.2, 49.0, 43.2, 41.3, 32.2, 28.7, 23.8 ppm.

**Synthesis of tert-butyl (2-(2-(2-(2-((2-(2,6-dioxopiperidin-3-yl)-1,3-dioxoisoindolin-4-yl)amino)ethoxy)ethoxy)ethoxy)ethyl)carbamate (L2)**

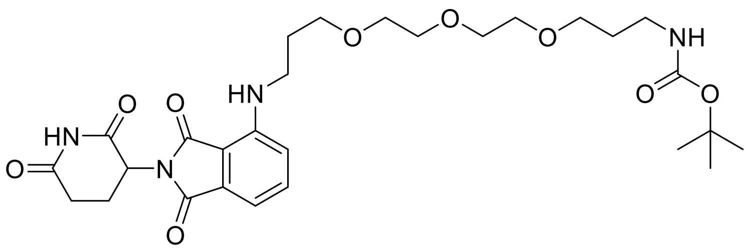


100 mg (362 µmol, 1.00 eq) 2-(2,6-dioxopiperidin-3-yl)-4-fluoroisoindoline-1,3-dione, 84.5 mg (362 µmol, 1.00 eq) tert-butyl (3-(2-(2-(3-aminopropoxy)ethoxy)ethoxy)propyl)carbamate and 126 µL (724 µmol, 2.00 eq) DIEA were dissolved in 5 mL NMP and stirred at 80 °C for 16 h. The solution was cooled to rt and diluted with EA and saturated $NaHCO_3$ and saturated NaCl solution. The mixture was extracted 3x with EA, dried over $MgSO_4$, filtered and excess solvent was removed under reduced pressure. The crude product was purified by Flash Chromatography to obtain 102 mg, 177 µmol 33% of a yellow solid.

Yield: 102 mg, 177 µmol, 33% of a yellow solid.

$R_f$ (30% Cyclohexane/EA): 0.25.

ESI: (calculated): [M+Na$^{+}$] 599.27 g/mol

(found): [M+Na$^{+}$] 599.36 g/mol.

HPLC: RT = 11.1 min (254 nm, 100%).

$^{1}H$ NMR (600 MHz, MeOD) δ = 7.55 (dd, $^{3}J$ = 8.5 Hz, $^{3}J$ = 7.2 Hz, 1H), 7.05 (dd, $^{2}J$ = 13.7, $^{3}J$ = 7.8 Hz, 2H), 5.07 – 5.04 (m, 1H), 3.69 – 3.68 (m, 2H), 3.65 – 3.62 (m, 10H), 3.45 – 3.42 (m, 2H), 3.07 (t, $^{3}J$ = 6.4 Hz, 2H), 2.89 – 2.83 (m, 1H), 2.76 – 2.70 (m, 2H), 2.12 – 2.10 (m, 1H), 1.94 – 1.89 (m, 4H) ppm.

$^{13}C$ NMR (126 MHz, MeOD) δ = 174.6, 171.7, 170.7, 169.3, 148.2, 137.3, 133.9, 118.0, 111.8, 111.0, 71.4, 71.2, 71.1, 70.4, 70.2, 50.2, 41.4, 39.95, 32.2, 30.1, 28.1, 23.8, 18.5 ppm.

**Synthesis of tert-butyl (17-((2-(2,6-dioxopiperidin-3-yl)-1,3-dioxoisoindolin-4-yl)amino)-3,6,9,12,15-pentaoxaheptadecyl)carbamate (L3)**

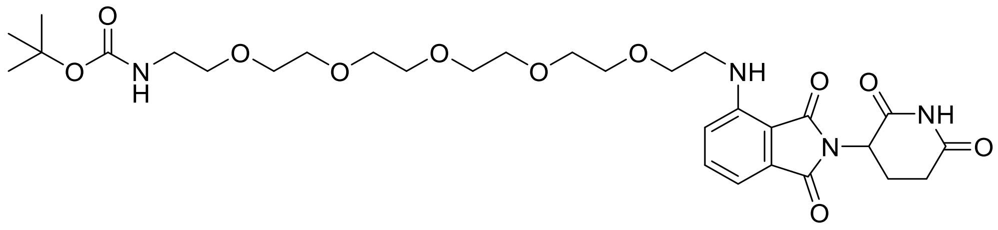

200 mg (720 µmol, 1.0 eq) 2-(2,6-dioxopiperidin-3-yl)-4-fluoroisoindoline-1,3-dione, 84.5 mg (720 µmol, 1.0 eq) tert-butyl (17-amino-3,6,9,12,15-pentaoxaheptadecyl)carbamate and 242 µL (1.45 mmol, 2.0 eq) DIEA were dissolved in 5 mL NMP and stirred at 80 °C for 16 h. The solution was cooled to rt and diluted with EA and saturated $NaHCO_3$ and saturated NaCl solution. The mixture was extracted 3x with EA, dried over $MgSO_4$, filtered and excess solvent was removed under reduced pressure. The crude product was purified by Flash Chromatography to obtain 210 mg, 330 µmol, 46% of a yellow oil.

Yield: 210 mg, 330 µmol, 46% of a yellow oil.

$R_f$ (33% Cyclohexane/ 67% EA): 0.13.

ESI: (calculated): [M+Na$^{+}$] 659.29 g/mol

(found): [M+Na$^{+}$] 659.30 g/mol.

HPLC: RT = 12.8 min (254 nm, 93%).

$^1$H NMR (250 MHz, DMSO) δ = 11.08 (s, 1H), 7.58 (dd, $^3J$ = 8.4 Hz, $^3J$ = 7.2 Hz, 1H), 7.15 (d, $^3J$ = 8.6 Hz, 1H), 7.04 (d, $^3J$ = 7.0 Hz, 1H), 6.73 (t, $^3$J = 4.7 Hz, 1H), 6.60 (t, $^3J$ = 5.6 Hz, 1H), 5.05 (dd, $^3J$ = 12.5 Hz, $^4J$ = 5.4 Hz, 1H), 3.62 (t, $^3J$ = 5.3 Hz, 2H), 3.58 – 3.43 (m, 16H), 3.41 – 3.23 (m, 4H), 3.05 (q, $^3J$ = 6.0 Hz, 2H), 2.99 – 2.77 (m, 1H), 2.61 - 2.55 (m, 2H), 2.04 - 2.02 (m, 1H), 1.36 (s, 9H) ppm.

**Synthesis of tert-butyl (23-((2-(2,6-dioxopiperidin-3-yl)-1,3-dioxoisoindolin-4-yl)amino)-3,6,9,12,15,18,21-heptaoxatricosyl)carbamate (L4)**

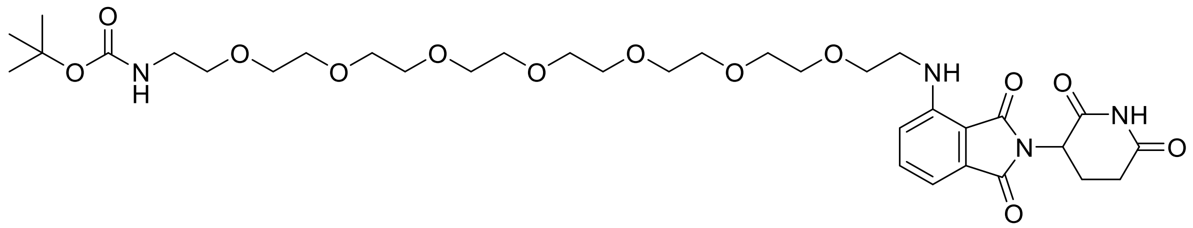

200 mg (720 µmol, 1.0 eq) 2-(2,6-dioxopiperidin-3-yl)-4-fluoroisoindoline-1,3-dione, 337 mg (720 µmol, 1.0 eq) tert-butyl (23-amino-3,6,9,12,15,18,21-heptaoxatricosyl)carbamate and 242 µL (1.45 mmol, 2.0 eq) DIEA were dissolved in 5 mL NMP and stirred at 80 °C for 16 h. The solution was cooled to rt and diluted with EA and saturated $NaHCO_3$ and saturated NaCl solution. The mixture was extracted 3x with EA, dried over $MgSO_4$, filtered and excess solvent was removed under reduced pressure. The crude product was purified by Flash Chromatography to obtain 271 mg, 374 µmol, 52% of a yellow oil.

Yield: 271 mg, 374 µmol, 52% of a yellow oil.

$R_f$ (50% Cyclohexane/ 50% EA): 0.17.

ESI: (calculated): [M+Na$^+$] 747.34 g/mol

(found): [M+Na$^+$] 747.41 g/mol.

HPLC: RT = 12.7 min (254 nm, 100%).

$^1$H NMR (600 MHz, MeOD) δ = 7.52 (dd, $^3J$ = 8.4, 7.2 Hz, 1H), 7.06 - 7.02 (m, 2H), 5.06 - 5.03 (m, 1H), 3.71 (t, $^3J$ = 5.3 Hz, 2H), 3.64 - 3.61 (m, 4H), 3.62 – 3.57 (m, 20H), 3.50 - 3.47 (m, 4H), 3.21 (t, $^3J$ = 5.5 Hz, 2H), 2.93 – 2.81 (m, 1H), 2.76 - 2.70 (m, 2H), 2.13 - 2.11 (m, 1H), 1.43 (s, 9H) ppm.

$^{13}$C NMR (126 MHz, MeOD) δ = 174.5, 171.3, 170.6, 169.1, 158.2, 148.1, 137.2, 133.7, 118.2, 112.0, 111.1, 80.0, 71.6, 71.6, 71.6, 71.5, 71.5, 71.4, 71.2, 71.0, 70.5, 50.2, 43.2, 41.2, 32.2, 28.80, 23.77 ppm.

**Synthesis of tert-butyl (4-(((2-(2,6-dioxopiperidin-3-yl)-1,3-dioxoisoindolin-4-yl)amino)methyl)benzyl)carbamate (L5)**

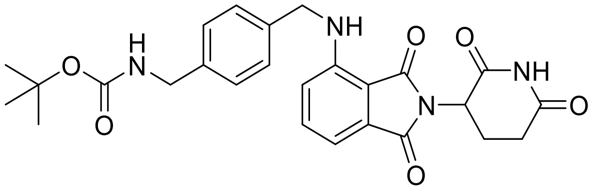

150 mg (543 µmol, 1.0 eq) 2-(2,6-dioxopiperidin-3-yl)-4-fluoroisoindoline-1,3-dione, 128 mg (543 µmol, 1.0 eq) tert-butyl (4-(aminomethyl)benzyl)carbamate and 284 µL (1.63 mmol, 2.0 eq) DIEA were dissolved in 7 mL DMF and stirred at 80 °C for 16 h. The solution was cooled to rt and diluted with EA and saturated $NaHCO_3$ and saturated NaCl solution. The mixture was extracted 3x with EA, dried over $MgSO_4$, filtered and excess solvent was removed under reduced pressure. The crude product was purified by Flash Chromatography to obtain 148 mg, 301 µmol, 55 % of a yellow oil.

Yield: 148 mg, 301 µmol, 55 % of a yellow oil.

$R_f$ (5% MeOH/ $CH_2Cl_2$): 0.31.

ESI: (calculated): [M+$Na^+$] 515.19 g/mol

(found): [M+$Na^+$] 515.27 g/mol.

HPLC: RT = 13.4 min (254 nm, 76%).

$^1$H NMR (250 MHz, MeOD) δ = 7.46 (dd, $^3J$ = 8.3 Hz, $^3J$ = 7.3 Hz, 1H), 7.30 - 7.27 (m, 3H), 7.19 -7.16 (m, 1H), 7.04 (d, $^3J$ = 7.1 Hz, 1H), 6.93 (d, $^3J$ = 8.5 Hz, 1H), 4.79 - 4.73 (m, 1H), 4.55 (s, 2H), 4.21 (s, 2H), 2.89 – 2.61 (m, 1H), 2.49 - 2.47 (m, 1H), 2.39 - 2.33 (m, 1H), 2.12 - 2.01 (m, 1H), 1.42 (s, 9H) ppm.

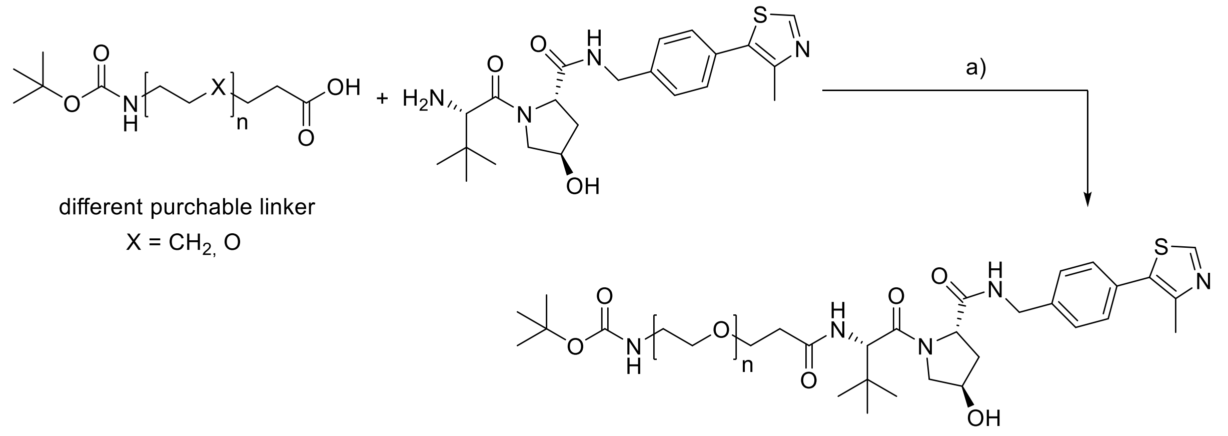


n = 1 **L6**
n = 2 **L7**
n = 4 **L8**

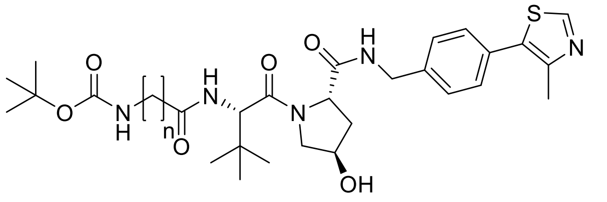

n = 2 **L9**
n = 3 **L10**
n = 4 **L11**
n = 5 **L12**
n = 6 **L13**

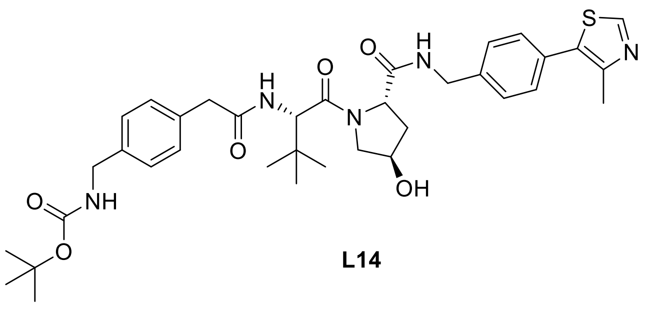

**L14**

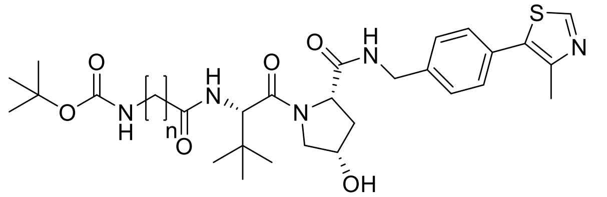

n = 2 **L15**

**SI Scheme 2**: Synthesis route of E3 ligase ligand linker **L6-L15**: a) HATU, DIEA, DMF, 4 h.

**Synthesis of tert-butyl (2-(3-(((S)-1-((2S,4R)-4-hydroxy-2-((4-(4-methylthiazol-5-yl)benzyl)carbamoyl)pyrrolidin-1-yl)-3,3-dimethyl-1-oxobutan-2-yl)amino)-3-oxopropoxy)ethyl)carbamate (L6)**

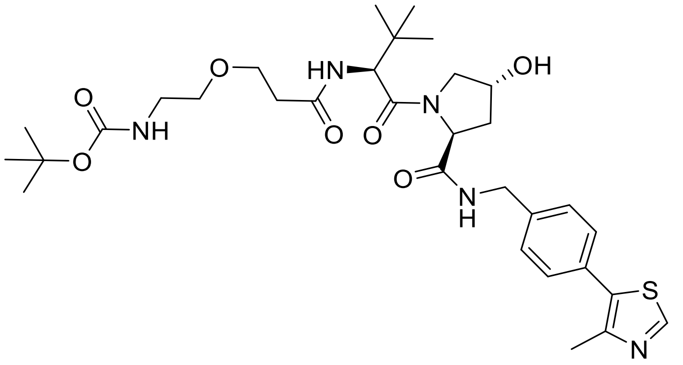

71 mg (307 µmol, 1.0 eq) 3-(2-((tert-butoxycarbonyl)amino)ethoxy)propanoic acid were dissolved in 5 mL dry DMF, 140 mg (368 µmol, 1.2 eq) HATU and 107 µL (613 µmol, 2.0 eq) DIEA were added. The solution was stirred for 5 min at rt, then 150 mg (322 µmol, 1.1 eq) VHL ligand 1 hydrochloride were added to the solution and stirred under Argon atmosphere for 4 h. The reaction mixture was quenched with 2 mL water and 2 mL saturated $NaHCO_3$, then the reaction was extracted 5x with EA. The organic phase was dried over $MgSO_4$, filtered and the solvent was removed under reduced pressure. The crude product was purified using an RP-FC system to obtain 80 mg, 124 µmol, 40% of a white solid.

Yield: 80 mg, 124 µmol, 40% of a white solid.

$R_f$ (10 %MeOH/ $CH_2Cl_2$): 0.15.

ESI: (calculated): [M+H$^+$] 646.31 g/mol

(found): [M+H$^+$] 646.32 g/mol.

HPLC: RT = 12.1 min (254 nm, 99%).

$^1$H NMR (500 MHz, DMSO) δ = 8.98 (s, 1H), 8.55 (t, $^3J$ = 6.0 Hz, 1H), 7.92 (d, $^3J$ = 9.4 Hz, 1H), 7.42 (d, $^3J$ = 8.4 Hz, 2H), 7.38 (d, $^3J$ = 8.4 Hz, 2H), 6.69 (t, $^3J$ = 5.5 Hz, 1H), 5.12 (d, $^3J$ = 3.6 Hz, 1H), 4.56 (d, $^3J$ =

9.4 Hz, 1H), 4.49 – 4.39 (m, 2H), 4.35 (s, 1H), 4.22 (dd, $^3J$ = 15.8 Hz, $^4J$ = 5.4 Hz, 1H), 3.66 (d, $^3J$ = 4.0 Hz, 1H), 3.63 (s, 1H), 3.57 (d, $^3J$ = 11.3 Hz, 2H), 3.40 – 3.33 (m, 2H), 3.05 (q, $^3J$ = 6.0 Hz, 2H), 2.59 – 2.50 (m, 1H), 2.44 (s, 3H), 2.40 – 2.29 (m, 1H), 2.06 – 1.97 (m, 1H), 1.92 (dd, $^3J$ = 8.5 Hz, $^4J$ = 4.5 Hz, 1H), 1.36 (s, 9H), 0.93 (s, 9H) ppm.

$^{13}$C NMR (126 MHz, DMSO) δ = 171.9, 170.0, 169.5, 155.5, 151.4, 147.7, 139.5, 131.2, 129.6, 128.6, 127.4, 77.6, 68.87, 66.6, 58.7, 56.4, 56.3, 41.67, 39.9, 35.4, 28.2, 26.3, 15.9 ppm.

**Synthesis of tert-butyl (2-(2-(3-(((S)-1-((2S,4R)-4-hydroxy-2-((4-(4-methylthiazol-5-yl)benzyl)carbamoyl)pyrrolidin-1-yl)-3,3-dimethyl-1-oxobutan-2-yl)amino)-3-oxopropoxy)ethoxy)ethyl)carbamate (L7)**

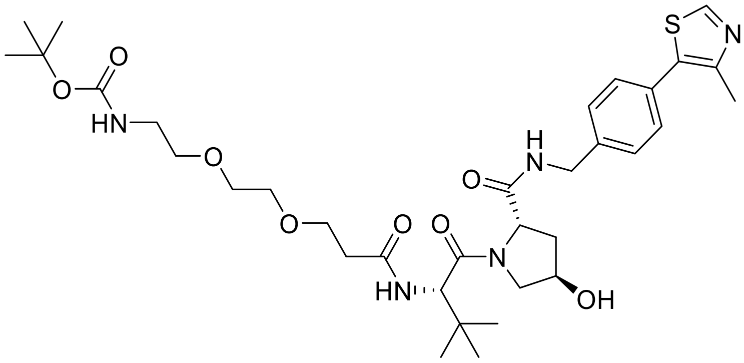


49 mg (178 µmol, 1.0 eq) 2,2-dimethyl-4-oxo-3,8,11-trioxa-5-azatetradecan-14-oic acid were dissolved in 5 mL dry DMF, 82 mg (214 µmol, 1.2 eq) HATU and 62 µL (356 µmol, 2.0 eq) DIEA were added. The solution was stirred for 5 min at rt, then 100 mg (214 µmol, 1.2 eq) VHL ligand 1 hydrochloride were added to the solution and stirred under Argon atmosphere for 4 h. The reaction mixture was quenched with 2 mL water and 2 mL saturated $NaHCO_3$, then the reaction was extracted 5x with EA. The organic phase was dried over $MgSO_4$, filtered and the solvent was removed under reduced pressure. The crude product was purified using an RP-FC system to obtain 97 mg, 141 µmol, 80 % of a white solid.

Yield: 97 mg, 141 µmol, 80 % of a white solid.

$R_f$ (10 % MeOH/ $CH_2Cl_2$): 0.32.

ESI: (calculated): [$M+Na^+$] 712.33 g/mol

(found): [$M+Na^+$] 712.32 g/mol.

HPLC: RT = 12.1 min (254 nm, 100 %).

$^1$H NMR (500 MHz, MeOD) δ = 8.87 (s, 1H), 7.89 (d, $^3J$ = 9.2 Hz, 1H), 7.47 (d, $^3J$ = 8.3 Hz, 2H), 7.44 – 7.39 (m, 2H), 4.66 (s, 1H), 4.56 (dd, $^2J$ = 20.9 Hz, $^3J$ = 12.2 Hz, 2H), 4.50 (s, 1H), 4.36 (d, $^3J$ = 15.5 Hz, 1H), 3.89 (d, $^3J$ = 11.0 Hz, 1H), 3.80 (dd, $^3J$ = 11.0 Hz, $^4J$ = 3.9 Hz, 1H), 3.74 (dt, $^3J$ = 10.9 Hz, $^4J$ = 5.3 Hz, 2H), 3.60 (s, 4H), 3.49 (t, $^3J$ = 5.6 Hz, 2H), 3.20 (m, 2H), 2.58 (m, 1H), 2.52 – 2.48 (m, 1H), 2.48 (s, 3H), 2.21 (m, 1H), 2.09 (m, 1H), 1.42 (s, 9H), 1.04 (s, 9H) ppm.

$^{13}$C NMR (126 MHz, MeOD) δ = 174.4, 173.7, 172.1, 158.4, 152.8, 149.0, 140.3, 133.4, 131.5, 130.4, 129.0, 80.1, 71.4, 71.2, 71.1, 68.3, 60.8, 58.9, 58.0, 43.7, 41.3, 38.9, 37.3, 36.8, 28.8, 27.0, 15.8 ppm.

**Synthesis of tert-butyl ((S)-17-((2S,4R)-4-hydroxy-2-((4-(4-methylthiazol-5-yl)benzyl)carbamoyl)pyrrolidine-1-carbonyl)-18,18-dimethyl-15-oxo-3,6,9,12-tetraoxa-16-azanonadecyl)carbamate (L8)**

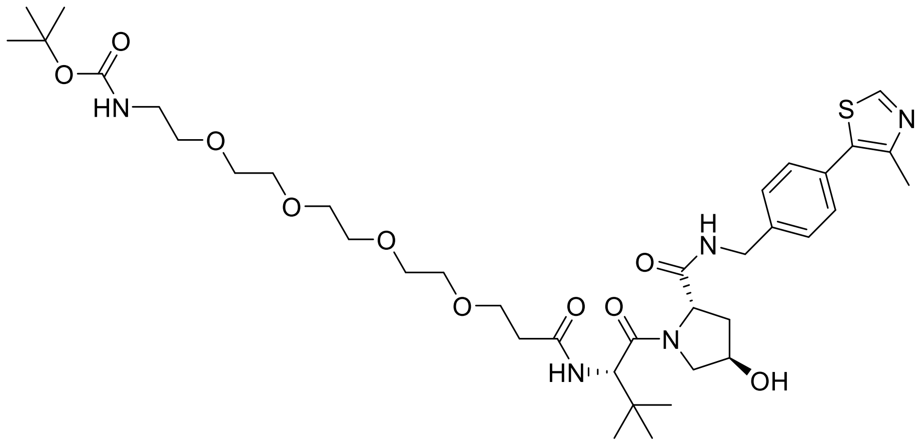

65 mg (178 µmol, 1.0 eq) 2,2-dimethyl-4-oxo-3,8,11,14,17-pentaoxa-5-azaicosan-20-oic acid were dissolved in 5 mL dry DMF, 82 mg (214 µmol, 1.2 eq) HATU and 62 µL (356 µmol, 2.0 eq) DIEA were added. The solution was stirred for 5 min at rt, then 100 mg (214 µmol, 1.2 eq) VHL ligand 1 hydrochloride were added to the solution and stirred under Argon atmosphere for 4 h. The reaction mixture was quenched with 2 mL water and 2 mL saturated $NaHCO_3$, then the reaction was extracted 5x with EA. The organic phase was dried over $MgSO_4$, filtered and the solvent was removed under reduced pressure. The crude product was purified using an RP-FC system to obtain 42 mg, 53.9 µmol, 30% of a colourless oil.

Yield: 42 mg, 53.9 µmol, 30% of a colourless oil.

$R_f$ (10% MeOH/ $CH_2Cl_2$): 0.22.

ESI: (calculated): [M+Na$^+$] 800.39 g/mol

(found): [M+Na$^+$] 800.48 g/mol.

HPLC: RT = 12.1 min (254 nm, 99%).

$^1$H NMR (500 MHz, MeOD) δ = 8.88 (s, 1H), 7.47 (d, $^3J$ = 8.3 Hz, 2H), 7.42 (d, $^3J$ = 8.3 Hz, 2H), 4.65 (s, 1H), 4.60 – 4.52 (m, 2H), 4.50 (s, 1H), 4.36 (d, $^3J$ = 15.5 Hz, 1H), 3.89 (d, $^3J$ = 11.1 Hz, 1H), 3.80 (dd, $^3J$ = 11.0 Hz, $^4J$ = 3.9 Hz, 1H), 3.76 – 3.70 (m, 2H), 3.63 – 3.61 (m, 10H), 3.60 – 3.58 (m, 2H), 3.49 (t, $^3J$ = 5.6 Hz, 2H), 3.21 (t, $^3J$ = 5.6 Hz, 2H), 2.56 (dd, $^3J$ = 7.5 Hz, $^4J$ = 5.3 Hz, 1H), 2.48 (s, 4H), 2.26 – 2.18 (m, 1H), 2.13 – 2.04 (m, 1H), 1.43 (s, 9H), 1.04 (s, 9H) ppm.

$^{13}$C NMR (126 MHz, MeOD) δ = 174.5, 173.7, 172.1, 158.4, 152.8, 149.0, 140.3, 133.4, 131.5, 130.4, 129.0, 80.1, 71.6, 71.5, 71.4, 71.3, 71.1, 68.3, 60.8, 58.9, 58.0, 43.7, 41.3, 38.9, 37.4, 36.8, 28.8, 27.0, 15.8 ppm.

**Synthesis of tert-butyl (3-(((S)-1-((2S,4R)-4-hydroxy-2-((4-(4-methylthiazol-5-yl)benzyl)carbamoyl)pyrrolidin-1-yl)-3,3-dimethyl-1-oxobutan-2-yl)amino)-3-oxopropyl)carbamate (L9)**

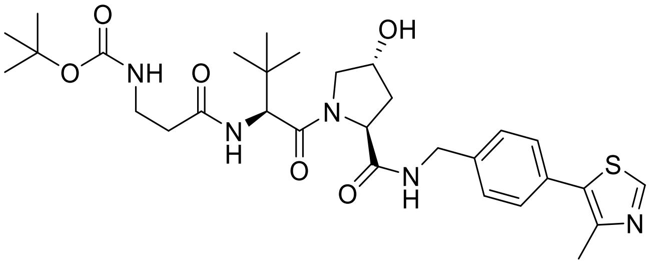


40 mg (214 µmol, 1.0 eq) 3-((tert-butoxycarbonyl)amino)propanoic acid were dissolved in 2 mL dry DMF, 98 mg (257 µmol, 1.2 eq) HATU and 75 µL (428 µmol, 2.0 eq) DIEA were added. The solution was stirred for 5 min at rt, then 100 mg (214 µmol, 1.0 eq) VHL ligand 1 hydrochloride were added to the solution and stirred under Argon atmosphere for 4 h. The reaction mixture was quenched with 2 mL water and 2 mL saturated $NaHCO_3$, then the reaction was extracted 5x with EA. The organic phase was

dried over $MgSO_4$, filtered and the solvent was removed under reduced pressure. The crude product was purified using an RP-FC system to obtain 62 mg, 103 µmol, 48% of a white solid.

Yield: 62 mg, 103 µmol, 48% of a white solid.

$R_f$ (10% MeOH/ $CH_2Cl_2$): 0.42.

ESI: (calculated): [M+2H$^+$] 603.31 g/mol

(found): [M+2H$^+$] 603.26 g/mol.

$^1$H NMR (300 MHz, MeOD) δ = 9.01 (s, 1H), 7.48 (d, $^3J$ = 8.0 Hz, 2H), 7.42 (d, $^3J$ = 7.4 Hz, 2H), 4.59 (d, $^3J$ = 12.9 Hz, 2H), 4.53 – 4.50 (m, 2H), 4.36 (d, $^3J$ = 15.2 Hz, 1H), 3.92 (d, $^3J$ = 11.4 Hz, 1H), 3.80 (d, $^3J$ = 10.2 Hz, 1H), 3.38 – 3.24 (m, 2H), 2.53 – 2.43 (m, 2H), 2.49 (s, 3H), 2.31 – 2.15 (m, 1H), 2.10 (m, 1H), 1.42 (s, 9H), 1.04 (s, 9H) ppm.

$^{13}$C NMR (75 MHz, MeOD) δ = 174.5, 173.7, 172.3, 153.3, 148.2, 140.6, 131.1, 130.5, 130.4, 129.6, 129.0, 80.2, 71.1, 60.8, 59.1, 58.0, 49.0, 43.7, 38.9, 38.6, 38.1, 36.8, 36.5, 28.7, 27.0, 15.4 ppm.

**Synthesis of tert-butyl (4-(((S)-1-((2S,4R)-4-hydroxy-2-((4-(4-methylthiazol-5-yl)benzyl)carbamoyl)pyrrolidin-1-yl)-3,3-dimethyl-1-oxobutan-2-yl)amino)-4-oxobutyl)carbamate (L10)**

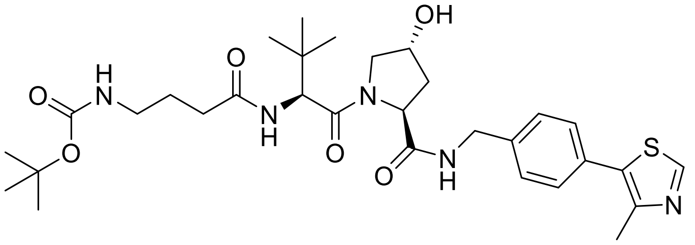


44 mg (214 µmol, 1.0 eq) 4-((tert-butoxycarbonyl)amino)butanoic acid were dissolved in 2 mL dry DMF, 98 mg (257 µmol, 1.2 eq) HATU and 75 µL (428 µmol, 2.0 eq) DIEA were added. The solution was stirred for 5 min at rt, then 100 mg (214 µmol, 1.0 eq) VHL ligand 1 hydrochloride were added to the solution and stirred under Argon atmosphere for 4 h. The reaction mixture was quenched with 2 mL

water and 2 mL saturated $NaHCO_3$, then the reaction was extracted 5x with EA. The organic phase was dried over $MgSO_4$, filtered and the solvent was removed under reduced pressure. The crude product was purified using an RP-FC system to obtain 61 mg, 99.2 µmol, 46% of a white solid.

Yield: 61 mg, 99.2 µmol, 46% of a white solid.

$R_f$ (10% MeOH/ $CH_2Cl_2$): 0.45.

ESI: (calculated): [M+Na$^+$] 639.30 g/mol

(found): [M+Na$^+$] 639.43 g/mol.

$^1$H NMR (300 MHz, MeOD) δ = 8.95 (s, 1H), 7.47 (d, $^3J$ = 8.4 Hz, 2H), 7.44 – 7.38 (m, 2H), 4.55 (m, 4H), 4.36 (d, $^3J$ = 15.5 Hz, 1H), 3.91 (m, 1H), 3.80 (m, 1H), 3.05 (t, $^3J$ = 6.8 Hz, 2H), 2.48 (s, 3H), 2.29 (m, 2H), 2.24 – 2.15 (m, 1H), 2.08 (m, 1H), 1.85 – 1.65 (m, 2H), 1.43 (s, 9H), 1.04 (s, 9H) ppm.

$^{13}$C NMR (75 MHz, MeOD) δ = 175.4, 174.5, 172.4, 153.1, 148.6, 140.4, 133.7, 131.2, 130.4, 129.4, 129.0, 79.9, 71.1, 60.8, 59.2, 58.0, 43.7, 40.7, 38.9, 36.5, 33.9, 28.8, 27.4, 27.0, 15.6 ppm.

**Synthesis of tert-butyl (5-(((S)-1-((2S,4R)-4-hydroxy-2-((4-(4-methylthiazol-5-yl)benzyl)carbamoyl)pyrrolidin-1-yl)-3,3-dimethyl-1-oxobutan-2-yl)amino)-5-oxopentyl)carbamate (L11)**

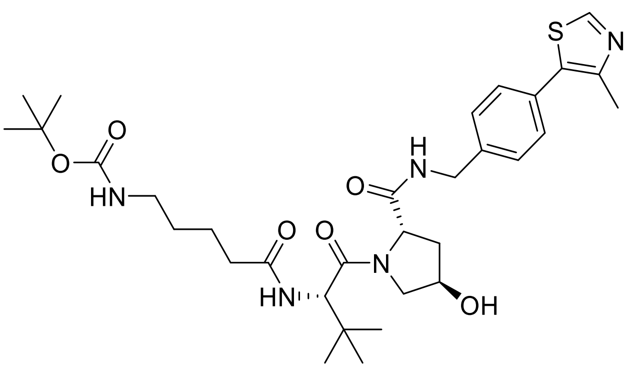


39 mg (178 µmol, 1.0 eq) 5-((tert-butoxycarbonyl)amino)pentanoic acid were dissolved in 5 mL dry DMF, 82 mg (214 µmol, 1.2 eq) HATU and 62 µL (356 µmol, 2.0 eq) DIEA were added. The solution was stirred for 5 min at rt, then 100 mg (214 µmol, 1.2 eq) VHL ligand 1 hydrochloride were added to the solution and stirred under Argon atmosphere for 4 h. The reaction mixture was quenched with 2 mL

water and 2 mL saturated $NaHCO_3$, then the reaction was extracted 5x with EA. The organic phase was dried over $MgSO_4$, filtered and the solvent was removed under reduced pressure. The crude product was purified using an RP-FC system to obtain 59 mg, 93.8 µmol, 53 % of a white solid.

Yield: 59 mg, 93.8 µmol, 53 % of a white solid.

$R_f$ (10 % MeOH/ 90 % $CH_2Cl_2$): 0.37.

ESI: (calculated): [M+Na$^+$] 652.31 g/mol

(found): [M+Na$^+$] 652.27 g/mol.

HPLC: RT = 12.1 min (254 nm, 100 %).

$^1$H NMR (500 MHz, MeOD) δ = 8.87 (s, 1H), 7.46 (d, $^3J$ = 8.3 Hz, 2H), 7.41 (d, $^3J$ = 8.3 Hz, 2H), 4.63 (s, 1H), 4.56 (dd, $^2J$ = 19.3 Hz, $^3J$ = 11.8 Hz, 2H), 4.50 (s, 1H), 4.36 (d, $^2J$ = 15.5 Hz, 1H), 3.90 (d, $^3J$ = 11.1 Hz, 1H), 3.80 (dd, $^3J$ = 10.9 Hz, $^4J$ = 3.9 Hz, 1H), 3.04 (t, $^3J$ = 6.9 Hz, 2H), 2.47 (s, 3H), 2.30 (m, 2H), 2.22 (m, 1H), 2.09 (m, 1H), 1.68 – 1.58 (m, 2H), 1.49 (m, 2H), 1.42 (s, 9H), 1.04 (s, 9H) ppm.

$^{13}$C NMR (126 MHz, MeOD) δ = 166.3, 165.0, 162.8, 149.0, 143.3, 139.5, 130.8, 123.9, 122.0, 120.9, 119.5, 70.3, 61.6, 51.3, 49.5, 48.5, 34.2, 31.4, 29.4, 27.0, 26.7, 21.0, 19.3, 17.6, 14.7, 6.3 ppm.

**Synthesis of tert-butyl (6-(((*S*)-1-((2*S*,4*R*)-4-hydroxy-2-((4-(4-methylthiazol-5-yl)benzyl)carbamoyl)pyrrolidin-1-yl)-3,3-dimethyl-1-oxobutan-2-yl)amino)-6-oxohexyl)carbamate (L12)**

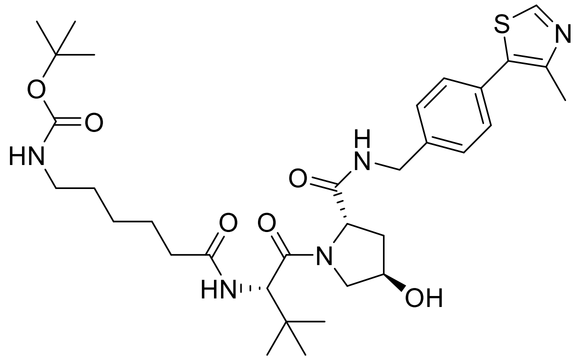

41 mg (178 µmol, 1.0 eq) 6-((tert-butoxycarbonyl)amino)hexanoic acid were dissolved in 5 mL dry DMF, 82 mg (214 µmol, 1.2 eq) HATU and 62 µL (356 µmol, 2.0 eq) DIEA were added. The solution was stirred for 5 min at rt, then 100 mg (214 µmol, 1.2 eq) VHL ligand 1 hydrochloride were added to the solution and stirred under Argon atmosphere for 4 h. The reaction mixture was quenched with 2 mL water and 2 mL saturated $NaHCO_3$, then the reaction was extracted 5x with EA. The organic phase was dried over $MgSO_4$, filtered and the solvent was removed under reduced pressure. The crude product was purified using an RP-FC system to obtain 100 mg, 155 µmol, 87 % of a white solid.

Yield: 100 mg, 155 µmol, 87 % of a white solid.

$R_f$ (5% MeOH/ $CH_2Cl_2$): 0.24.

ESI: (calculated): [M+$Na^+$] 666.32 g/mol

(found): [M+$Na^+$] 666.30 g/mol.

HPLC: RT = 12.3 min (254 nm, 100 %).

$^1$H NMR (500 MHz, MeOD) δ = 8.88 (s, 1H), 7.47 (d, $^3J$ = 8.4 Hz, 2H), 7.41 (d, $^3J$ = 8.3 Hz, 2H), 4.63 (s, 1H), 4.58 (m, 1H), 4.53 (m, 1H) 4.50 (s, 1H), 4.36 (m, 1H), 3.91 (d, $^3J$ = 11.0 Hz, 1H), 3.80 (dd, $^3J$ = 10.9 Hz, $^4J$ = 3.9 Hz, 1H), 3.01 (m, 2H), 2.47 (s, 3H), 2.28 (m, 2H), 2.21 (m, 1H), 2.13 – 2.05 (m, 1H), 1.66 – 1.59 (m, 2H), 1.51 – 1.44 (m, 2H), 1.42 (s, 9H), 1.34 (m, 2H), 1.04 (s, 9H) ppm.

$^{13}$C NMR (126 MHz, MeOD) δ = 175.9, 174.5, 172.3, 158.5, 152.8, 149.0, 140.3, 133.4, 131.5, 130.4, 129.0, 79.8, 71.1, 60.8, 59.0, 58.0, 43.7, 41.2, 38.9, 36.5, 30.6, 28.8, 27.5, 27.0, 26.7, 15.8 ppm.

**Synthesis of tert-butyl (7-(((*S*)-1-((2*S*,4*R*)-4-hydroxy-2-((4-(4-methylthiazol-5-yl)benzyl)carbamoyl)pyrrolidin-1-yl)-3,3-dimethyl-1-oxobutan-2-yl)amino)-7-oxoheptyl)carbamate (L13)**

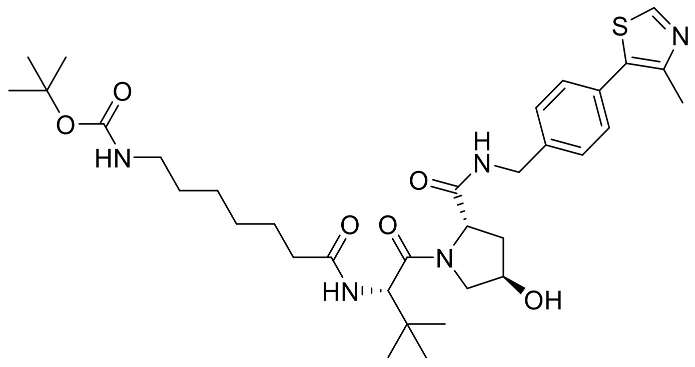


44 mg (178 µmol, 1.0 eq) 7-((tert-butoxycarbonyl)amino)heptanoic acid were dissolved in 5 mL dry DMF, 82 mg (214 µmol, 1.2 eq) HATU and 62 µL (356 µmol, 2.0 eq) DIEA were added. The solution was stirred for 5 min at rt, then 100 mg (214 µmol, 1.2 eq) VHL ligand 1 hydrochloride were added to the solution and stirred under Argon atmosphere for 4 h. The reaction mixture was quenched with 2 mL water and 2 mL saturated $NaHCO_3$, then the reaction was extracted 5x with EA. The organic phase was dried over $MgSO_4$, filtered and the solvent was removed under reduced pressure. The crude product was purified using an RP-FC system to obtain 76 mg, 115 µmol, 65% of a white solid.

Yield: 76 mg, 115 µmol, 65 % of a white solid.

$R_f$ (5% MeOH/ 95 % $CH_2Cl_2$): 0.28.

ESI: (calculated): [M+Na$^+$] 680.34 g/mol

(found): [M+Na$^+$] 680.31 g/mol.

HPLC: RT = 12.5 min (254 nm, 94%).

$^1$H NMR (500 MHz, MeOD) δ = 8.87 (s, 1H), 8.64 (t, $^3J$ = 6.0 Hz, 1H), 7.80 (d, $^3J$ = 9.0 Hz, 1H, NH), 7.46 (d, $^3J$ = 8.3 Hz, 2H), 7.41 (d, $^3J$ = 8.3 Hz, 2H), 4.66 – 4.62 (m, 1H), 4.61 – 4.56 (m, 1H), 4.54 (d, $^3J$ = 15.5 Hz, 1H), 4.50 (s, 1H), 4.36 (d, $^3J$ = 15.5 Hz, 1H), 3.91 (d, $^3J$ = 11.1 Hz, 1H), 3.80 (dd, $^3J$ = 10.9 Hz, $^4J$ = 3.9 Hz,

1H), 3.01 (t, $^{3}J$ = 7.0 Hz, 2H), 2.47 (s, 3H), 2.37 – 2.23 (m, 2H), 2.22 – 2.18 (m, 1H), 2.08 (m, 1H), 1.61 (dt, $^{3}J$ = 14.2 Hz, $^{4}J$ = 7.2 Hz, 2H), 1.49 – 1.43 (m, 2H), 1.42 (s, 9H), 1.35 – 1.31 (m, 4H), 1.04 (s, 9H) ppm. $^{13}$C NMR (126 MHz, MeOD) δ = 176.0, 174.4, 172.3, 158.5, 152.8, 149.0, 140.3, 133.4, 131.5, 130.3, 129.0, 79.7, 71.1, 60.8, 59.0, 58.0, 43.7, 41.3, 38.9, 36.5, 30.8, 29.9, 28.8, 27.5, 27.0, 26.9, 15.8 ppm.

**Synthesis of tert-butyl (4-(2-(((S)-1-((2S,4R)-4-hydroxy-2-((4-(4-methylthiazol-5-yl)benzyl)carbamoyl)pyrrolidin-1-yl)-3,3-dimethyl-1-oxobutan-2-yl)amino)-2-oxoethyl)benzyl)carbamate (L14)**

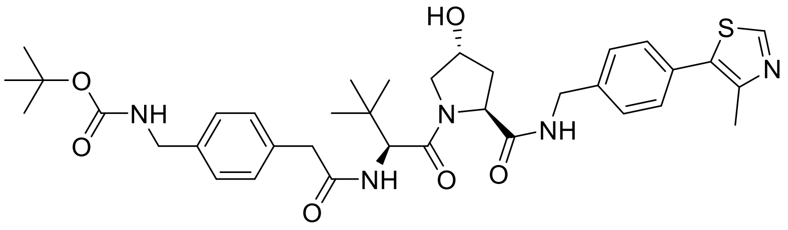


81 mg (307 µmol, 1.0 eq) 2-(4-(((tert-butoxycarbonyl)amino)methyl)phenyl)acetic acid were dissolved in 5 mL dry DMF, 140 mg (368 µmol, 1.2 eq) HATU and 107 µL (613 µmol, 2.0 eq) DIEA were added. The solution was stirred for 5 min at rt, then 150 mg (322 µmol, 1.1 eq) VHL ligand 1 hydrochloride were added to the solution and stirred under Argon atmosphere for 4 h. The reaction mixture was quenched with 2 mL water and 2 mL saturated $NaHCO_3$, then the reaction was extracted 5x with EA. The organic phase was dried over $MgSO_4$, filtered and the solvent was removed under reduced pressure. The crude product was purified using an RP-FC system to obtain 65 mg, 95.9 µmol, 31% of a yellow solid.

Yield: 65 mg, 95.9 µmol, 31% of a yellow solid.

$R_f$ (10% MeOH/ $CH_2Cl_2$): 0.11.

ESI: (calculated): [M+H$^+$] 678.33 g/mol

(found): [M+H$^+$] 678.34 g/mol.

HPLC: RT = 12.5 min (254 nm, 100%).

$^1$H NMR (500 MHz, DMSO) δ = 8.98 (s, 1H), 8.57 (t, $^3J$ = 6.1 Hz, 1H), 8.07 (d, $^3J$ = 9.3 Hz, 1H), 7.42 (d, $^3J$ = 8.4 Hz, 2H), 7.39 (d, $^3J$ = 8.4 Hz, 2H), 7.33 (t, $^3J$ = 6.3 Hz, 1H), 7.20 (d, $^3J$ = 8.0 Hz, 2H), 7.13 (d, $^3J$ = 8.0 Hz, 2H), 5.11 (d, $^3J$ = 3.6 Hz, 1H), 4.51 (d, $^3J$ = 9.4 Hz, 1H), 4.47 – 4.40 (m, 2H), 4.34 (s, 1H), 4.22 (dd, $^3J$ = 15.8 Hz, $^4J$ = 5.5 Hz, 1H), 4.08 (d, $^3J$ = 6.1 Hz, 2H), 3.66 (m, 1H), 3.62 (d, $^3J$ = 13.2 Hz, 2H), 3.41 (d, $^3J$ = 13.8 Hz, 1H), 2.45 (s, 3H), 2.06 – 1.98 (m, 1H), 1.90 (m, 1H), 1.38 (s, 9H), 0.91 (s, 9H) ppm.

$^{13}$C NMR (126 MHz, DMSO) δ = 171.9, 170.0, 169.5, 155.7, 151.4, 147.7, 139.5, 138.1, 134.9, 131.1, 129.6, 128.9, 128.6, 127.4, 126.7, 77.7, 68.8, 58.7, 56.4, 56.4, 43.1, 41.6, 40.0, 35.4, 28.2, 26.3, 15.9 ppm.

**Synthesis of tert-butyl (5-(((S)-1-((2S,4S)-4-hydroxy-2-((4-(4-methylthiazol-5-yl)benzyl)carbamoyl)pyrrolidin-1-yl)-3,3-dimethyl-1-oxobutan-2-yl)amino)-5-oxopentyl)carbamate (L15)**

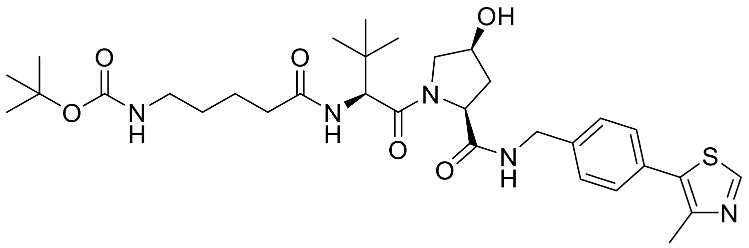

5.63 g (30.9 mmol, 1.00 eq) 4-Bromobenzonitrile and 4.5 mL (30.9 mmol, 1.00 eq) 4-Methylthiazole were dissolved in 25 mL Dimethylacetamide and 2 mL water. 6.00 g (61.8 mmol, 2.00 eq) potassium acetate and 32 mg (3.09 mmol, 0.10 eq) palladium(II)acetate were added. The mixture was stirred for 24 h at 115°C. Excess solvent was removed under reduced pressure. The reaction mixture was dissolved with water and ethyl acetate. The phases were separated and the aqueous phase was extracted four times with ethylacetate. The combined organic phases were dried over MgSO4, filtered and excess solvent was removed under reduced pressure. The purification of the crude product was done by column chromatography to obtain 2.36 g (11.6 mmol, 38%) of 4-(4-methylthiazol-5-yl)benzonitrile.

This product was dissolved in THF and added slowly to 24 mL (23.1 mmol, 2.00 eq) of a cold solution of 1M $LiAlH_4$ in THF. The reaction mixture was allowed to reach rt and then heated at 40 °C for 18 h. The reaction mixture was cooled to rt and water was added slowly. The solution was filtered and washed with 10% MeOH/ $CH_2Cl_2$. The solvent was removed under reduced pressure, then the crude mixture was diluted with $CH_2Cl_2$. The phases were separated, the aqueous phase was extracted four times with $CH_2Cl_2$. The combined organic phases were dried over MgSO4, filtered and excess solvent was removed under reduced pressure to obtain 2.00 g (9.79 mmol, 84 %) of crude (4-(4-methylthiazol-5-yl)phenyl)methanamine.

351 mg (1.52 mmol, 1.05 eq) (2S,4S)-1-(tert-butoxycarbonyl)-4-hydroxypyrrolidine-2-carboxylic acid and 661 mg (1.74 mmol, 1.20 eq) HATU were dissolved in 7 mL DMF. 505 µL (2.89 mmol, 2.00 eq) DIEA were added and the mixture was stirred for 15 min at rt. 296 mg (1.45mmol, 1.00 eq) (4-(4-methylthiazol-5-yl)phenyl)methanamine was added to the mixture and stirred for 3 h at rt. The reaction was stopped with 1 mL water. Saturated $NaHCO_3$ solution and saturated NaCl solution were added and the reaction mixture was extracted 4x with EA. The combined organic phases were washed with saturated $NaHCO_3$ solution, dried over $MgSO_4$ and filtered. The solvent of the organic phase was evapored under reduced pressure. The purification of the crude product was done by Flash Chromatography to obtain 286 mg (685 µmol, 47%) of tert-butyl (2S,4S)-4-hydroxy-2-((4-(4-methylthiazol-5-yl)benzyl)carbamoyl)pyrrolidine-1-carboxylate.

The obtained product was dissolved in 1 mL TFA and 1 mL $CH_2Cl_2$ and stirred for 1h at rt. Excess solvent was removed under reduced pressure. 158 mg (685 µmol, 1.00 eq) (S)-2-((tert-butoxycarbonyl)amino)-3,3-dimethylbutanoic acid and 312 mg (822 µmol, 1.20 eq) HATU were dissolved in 5 mL DMF. 238 µL (1.37 mmol, 2.00 eq) DIEA were added and the mixture was stirred for 15 min at rt. 286 mg (685 µmol, 1.00 eq) of deprotected tert-butyl (2S,4S)-4-hydroxy-2-((4-(4-methylthiazol-5-yl)benzyl)carbamoyl)pyrrolidine-1-carboxylate was dissolved in 1 mL DMF and DIEA was added until the pH was basic. The mixture was added to the active ester species and stirred for 3 h at rt. The reaction was stopped with 1 mL water. Saturated $NaHCO_3$ solution and saturated NaCl solution were added and the reaction mixture was extracted 4x with EA. The combined organic phases were washed with saturated $NaHCO_3$ solution, dried over $MgSO_4$ and filtered. The solvent of the organic phase was evapored under reduced pressure. The purification of the crude product was done by Flash Chromatography to obtain 242 mg (456 µmol, 67%) tert-butyl ((S)-1-((2S,4S)-4-hydroxy-2-((4-(4-methylthiazol-5-yl)benzyl)carbamoyl)pyrrolidin-1-yl)-3,3-dimethyl-1-oxobutan-2-yl)carbamate.

114 mg (216 µmol, 1.00 eq) tert-butyl ((S)-1-((2S,4S)-4-hydroxy-2-((4-(4-methylthiazol-5-yl)benzyl)carbamoyl)pyrrolidin-1-yl)-3,3-dimethyl-1-oxobutan-2-yl)carbamate were dissolved in 1 mL TFA and 1 mL $CH_2Cl_2$ and stirred for 1h at rt. Excess solvent was removed under reduced pressure. 47 mg (216 µmol, 1.00 eq) 5-((tert-butoxycarbonyl)amino)pentanoic acid, 98 mg (258µmol, 1.20 eq) HATU and 75 µL (429 µmol, 2.00 eq) DIPEA were dissolved in 1 mL DMF and stirred for 15 min at rt. The deprotected amine was dissolved in 1 mL DMF and DIEA was added until the pH was basic. The mixture was added to the active ester species and stirred for 3 h at rt. The reaction was stopped with 1 mL water. Saturated $NaHCO_3$ solution and saturated NaCl solution were added and the reaction mixture was extracted 4x with EA. The combined organic phases were washed with saturated $NaHCO_3$ solution, dried over $MgSO_4$ and filtered. The solvent of the organic phase was evapored under reduced pressure. The purification of the crude product was carried out on a preparative HPLC system to obtain 60 mg (95.3 µmol, 44%) tert-butyl (5-(((S)-1-((2S,4S)-4-hydroxy-2-((4-(4-methylthiazol-5-yl)benzyl)carbamoyl)pyrrolidin-1-yl)-3,3-dimethyl-1-oxobutan-2-yl)amino)-5-oxopentyl)carbamate.

ESI: (calculated) $[M+Na^+]$ 652.31 g/mol

(found) $[M+Na^+]$ 652.31 g/mol.

HPLC: RT = 12.0 min (254 nm, 100%).

$^1H$ NMR (500 MHz, MeOD) δ = 8.92 (s, 1H), 7.53 – 7.35 (m, 4H), 4.63 (s, 1H), 4.59 – 4.52 (m, 2H), 4.51 - 4.48 (m, 1H), 4.36 (d, $^3J$ = 15.5 Hz, 1H), 3.91 - 3.89 (m, 1H), 3.80 (dd, $^3J$ = 11.0 Hz, $^4J$ = 3.9 Hz, 1H), 3.04 (t, $^3J$ = 6.9 Hz, 2H), 2.48 (s, 3H), 2.38 – 2.17 (m, 3H), 2.11 - 2.06 (m, 1H), 1.68 – 1.56 (m, 2H), 1.51 - 1.46 (m, 2H), 1.42 (s, 9H), 1.04 (s, 9H) ppm.

$^{13}C$ NMR (126 MHz, MeOD) δ = 175.7, 174.5, 172.3, 158.5, 153.0, 148.7, 140.4, 133.6, 131.3, 130.4, 129.0, 79.8, 71.1, 60.8, 59.0, 58.0, 49.0, 43.7, 40.9, 38.9, 36.5, 36.2, 30.5, 28.8, 27.0, 24.2, 15.7 ppm.

### 8. Synthesis of BRET Tracer molecules

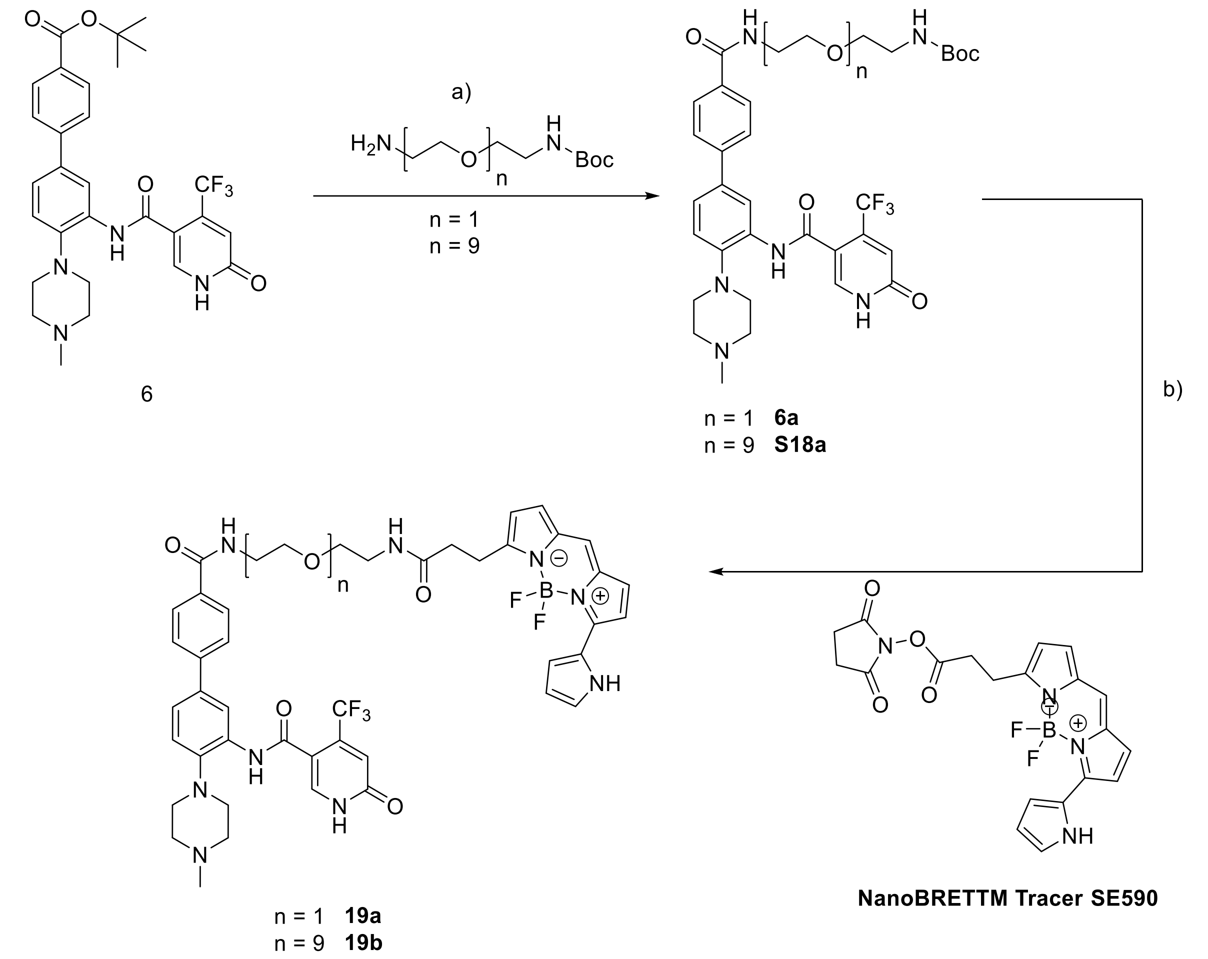


**SI Scheme 3**: Synthesis of NanoBRET™ tracer molecules **19a-b** via intermediates **6a** and **S18a** for WDR5. a) 1) TFA/ CH2Cl2 (1:1), rt, 2h; 2) HATU, DIEA, DMF, rt, 3h; b) DIEA, DMF, rt, 3 h.

**Synthesis of tert-butyl (1-(3'-(6-hydroxy-4-(trifluoromethyl)nicotinamido)-4'-(4-methylpiperazin-1-yl)-[1,1'-biphenyl]-4-yl)-1-oxo-5,8,11,14,17,20,23,26,29-nonaoxa-2-azahentriacontan-31-yl)carbamate (S18a)**

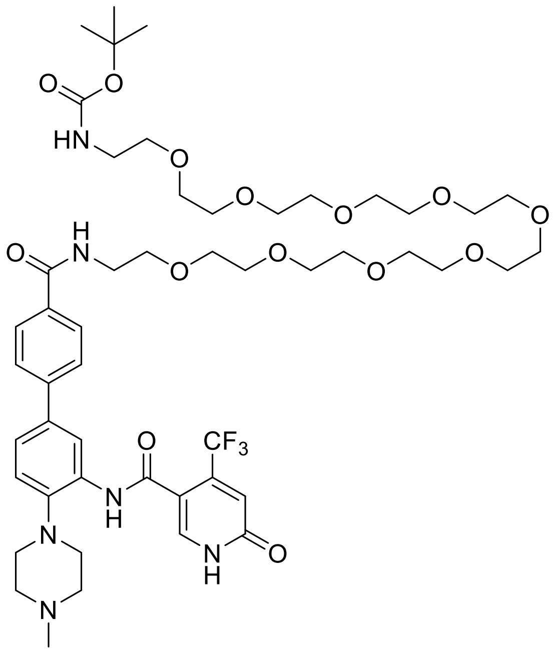


22 mg (40 µmol, 1.0eq) tert-butyl 3'-(6-hydroxy-4-(trifluoromethyl)nicotinamido)-4'-(4-methylpiperazin-1-yl)-[1,1'-biphenyl]-4-carboxylate were dissolved in 0.5 mL DCM and 0.5 mL TFA and stirred at rt for 1 h. Excess solvent was evapored. The solid was dissolved in 0.5 mL DMF, then 137 µL (720 µmol, 20 eq) DIEA and 18 mg (47 µmol, 1.2 eq) HATU were added. After 15 min, a solution of 23.1 mg (42 µmol, 1.05 eq) tert-butyl (29-amino-3,6,9,12,15,18,21,24,27-nonaoxanonacosyl)carbamate in 0.5 mL DMF was added. The solution was stirred for 3 h at rt. The reaction mixture was quenched with 2 mL water and 2 mL saturated $NaHCO_3$, then the reaction was extracted 3x with EA. The organic phase was dried over $MgSO_4$, filtered and the solvent was removed under reduced pressure. The crude product was purified using by HPLC to obtain 41 mg, 35.6 µmol, 89% of a colourless oil.

Yield: 41 mg, 35.6 µmol, 89% of a colourless oil.

MALDI: (calculated) [M-Boc+H$^+$] 939.47 g/mol

(found) [M-Boc+H$^+$] 939.43 g/mol.

$^1$H NMR (400 MHz, MeOD) δ = 8.28 (d, $^4J$ = 2.1 Hz, 1H), 8.04 (s, 1H), 7.94 (d, $^3J$ = 8.6 Hz, 2H), 7.78 – 7.71 (m, 2H), 7.58 (dd, $^3J$ = 8.4 Hz, $^4J$ = 2.2 Hz, 1H), 7.40 (d, $^3J$ = 8.4 Hz, 1H), 6.94 (s, 1H), 3.72 – 3.56 (m, 48H), 2.97 (s, 3H), 1.43 (s, 9H) ppm.

**Synthesis of N-(4'-((2-(2-(3-(5,5-difluoro-7-(1H-pyrrol-2-yl)-5H-5l4,6l4-dipyrrolo[1,2-c:2',1'-f][1,3,2]diazaborinin-3-yl)propanamido)ethoxy)ethyl)carbamoyl)-4-(4-methylpiperazin-1-yl)-[1,1'-biphenyl]-3-yl)-6-hydroxy-4-(trifluoromethyl)nicotinamide (19a)**

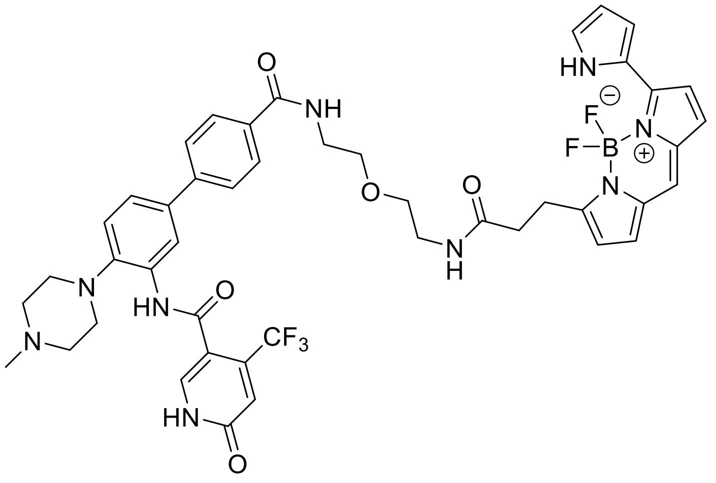


12 mg (14 µmol, 1.0 eq) tert-butyl (2-(2-(3'-(6-hydroxy-4-(trifluoromethyl)nicotinamido)-4'-(4-methylpiperazin-1-yl)-[1,1'-biphenyl]-4-carboxamido)ethoxy)ethyl)carbamate **6a** were dissolved in 0.5 mL DCM and 0.5 mL TFA and stired at rt for 1 h. Excess solvent was evapored. The solid was dissolved in 0.5 mL DMF, then 19 µL (111 µmol, 8.0 eq) DIEA and 6.2 mg (15 µmol, 1.1 eq) 2,5-dioxopyrrolidin-1-yl-3-(5,5-difluoro-7-(1H-pyrrol-2-yl)-5H-5l4,6l4-dipyrrolo[1,2-c:2',1'-f][1,3,2]diazaborinin-3-yl)propanoate in 0.5 mL DMF was added. The solution was stirred for 3 h at rt.

The reaction mixture was quenched with 2 mL water and 2 mL saturated $NaHCO_3$, then the reaction was extracted 3x with EA. The organic phase was dried over $MgSO_4$, filtered and the solvent was removed under reduced pressure. The crude product was purified using by HPLC to obtain 6.35 mg, 7.08 µmol, 77 % of a purple solid.

Yield: 6.35 mg, 7.08 µmol, 77 % of a purple solid.

$R_f$ (20% MeOH/ $CH_2Cl_2$): 0.70.

HPLC: RT = 12.1 min (254 nm, 95%).

HRMS: (calculated) [M+$Na^+$] 920.3449 g/mol

(found) [M+$Na^+$] 920.3459 g/mol.

$^1$H NMR (500 MHz, MeOD) δ = 8.21 (d, $^4J$ = 2.1 Hz, 1H), 8.01 (s, 1H), 7.85 (d, $^3J$ = 8.4 Hz, 2H), 7.64 (d, $^3J$ = 8.4 Hz, 2H), 7.46 (dd, $^3J$ = 8.4 Hz, $^4$J = 2.1 Hz, 1H), 7.27 (d, $^3J$ = 8.4 Hz, 1H), 7.21 – 7.07 (m, 4H), 7.02 – 6.90 (m, 2H), 6.85 (d, $^3J$ = 3.9 Hz, 1H), 6.32 (dd, $^3J$ = 3.8 Hz, $^4$J = 2.6 Hz, 1H), 6.26 (d, $^3J$ = 3.9 Hz, 1H), 3.61 (m, 8H), 3.40 (t, $^3J$ = 5.2 Hz, 2H), 3.35 – 3.20 (m, 6H), 3.13 (t, $^3J$ = 12.8 Hz, 2H), 2.96 (s, 3H), 2.61 (t, $^3J$ = 7.6 Hz, 2H) ppm.

$^{13}$C NMR (126 MHz, MeOD) δ = 174.97, 169.9, 165.3, 163.9, 156.2, 152.1, 144.3, 143.9, 141.4 (q, J = 32.9 Hz), 139.7, 138.9, 138.6, 134.9, 134.4, 133.8, 133.1, 129.0, 127.9, 127.4, 126.0, 124.9, 124.6, 123.8, 123.4 (q, J = 273 Hz), 122.2, 121.0, 120.6 (d, J = 6.5 Hz), 119.0, 117.2, 114.4, 112.3, 70.5, 70.4, 55.0, 50.3, 43.7, 40.9, 40.4, 36.1, 25.6 ppm.

**Synthesis of N-(4'-((33-(5,5-difluoro-7-(1H-pyrrol-2-yl)-5H-5l4,6l4-dipyrrolo[1,2-c:2',1'-f][1,3,2]diazaborinin-3-yl)-31-oxo-3,6,9,12,15,18,21,24,27-nonaoxa-30-azatritriacontyl)carbamoyl)-4-(4-methylpiperazin-1-yl)-[1,1'-biphenyl]-3-yl)-6-hydroxy-4-(trifluoromethyl)nicotinamide (19b)**

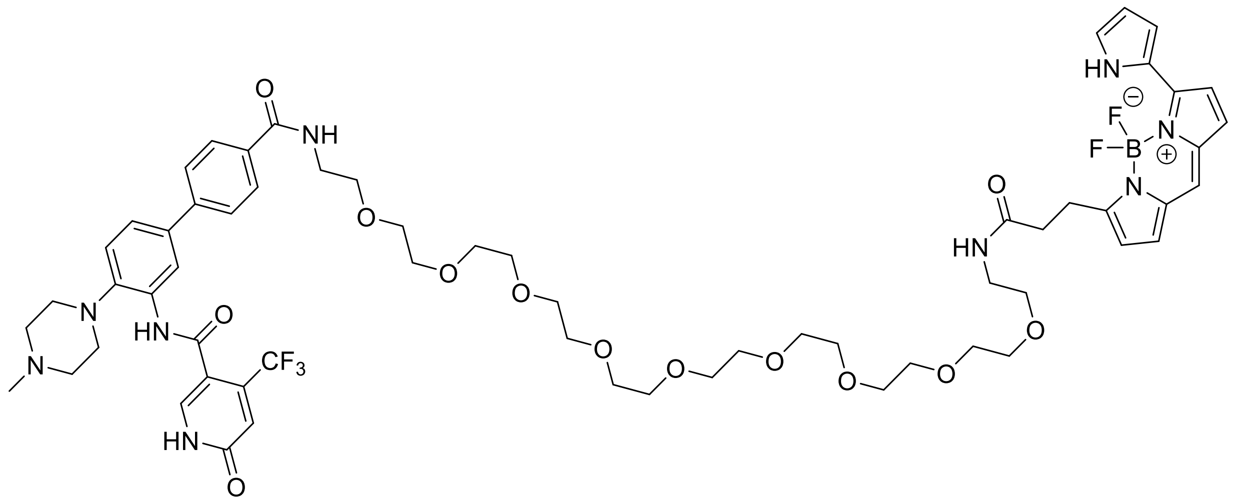

12.5 mg (12 µmol, 1.0 eq) tert-butyl (1-(3'-(6-hydroxy-4-(trifluoromethyl)nicotinamido)-4'-(4-methylpiperazin-1-yl)-[1,1'-biphenyl]-4-yl)-1-oxo-5,8,11,14,17,20,23,26,29-nonaoxa-2-azahentriacontan-31-yl)carbamate were dissolved in 0.5 mL DCM and 0.5 mL TFA and stired at rt for 1 h. Excess solvent was evapored. The solid was dissolved in 0.5 mL DMF, then 17 µL (10 µmol, 2.0 eq) DIEA and 5.8 mg (14 µmol, 1.1 eq) 2,5-dioxopyrrolidin-1-yl-3-(5,5-difluoro-7-(1H-pyrrol-2-yl)-5H-5l4,6l4-dipyrrolo[1,2-c:2',1'-f][1,3,2]diazaborinin-3-yl)propanoate in 0.5 mL DMF was added. The solution was stirred for 3 h at rt. The reaction mixture was quenched with 2 mL water and 2 mL saturated $NaHCO_3$, then the reaction was extracted 3x with EA. The organic phase was dried over $MgSO_4$, filtered and the solvent was removed under reduced pressure. The crude product was purified using by HPLC to obtain 4.88 mg, 3.9 µmol, 33% of a purple solid.

Yield: 4.88 mg, 3.9 µmol, 33% of a purple solid.

HPLC: RT = 12.4 min (254 nm, 100%).

HRMS: (calculated) [M+Na$^+$] 1272.5546 g/mol

(found) [M+Na$^+$] 1272.5583 g/mol.

$^1$H NMR (400 MHz, MeOD) δ = 8.27 (d, $^4J$ = 2.1 Hz, 1H), 8.02 (s, 1H), 7.92 (d, $^3J$ = 8.5 Hz, 2H), 7.72 (d, $^3J$ = 8.5 Hz, 2H), 7.56 (dd, $^3J$ = 8.4 Hz, $^4J$ = 2.2 Hz, 1H), 7.36 (d, $^3J$ = 8.4 Hz, 1H), 7.22 (s, 1H), 7.21 – 7.13 (m, 3H), 7.00 (d, $^4J$ = 4.6 Hz, 1H), 6.94 (s, 1H), 6.91 (d, $^4J$ = 3.9 Hz, 1H), 6.35 - 6.34 (m, 1H), 6.32 (d, $^4J$ = 4.0 Hz, 1H), 3.75 – 3.47 (m, 44H), 3.36 (t, $^3J$ = 5.4 Hz, 2H), 3.27 (t, $^3J$ = 7.7 Hz, 4H), 2.95 (s, 3H), 2.70 – 2.56 (m, 2H) ppm.

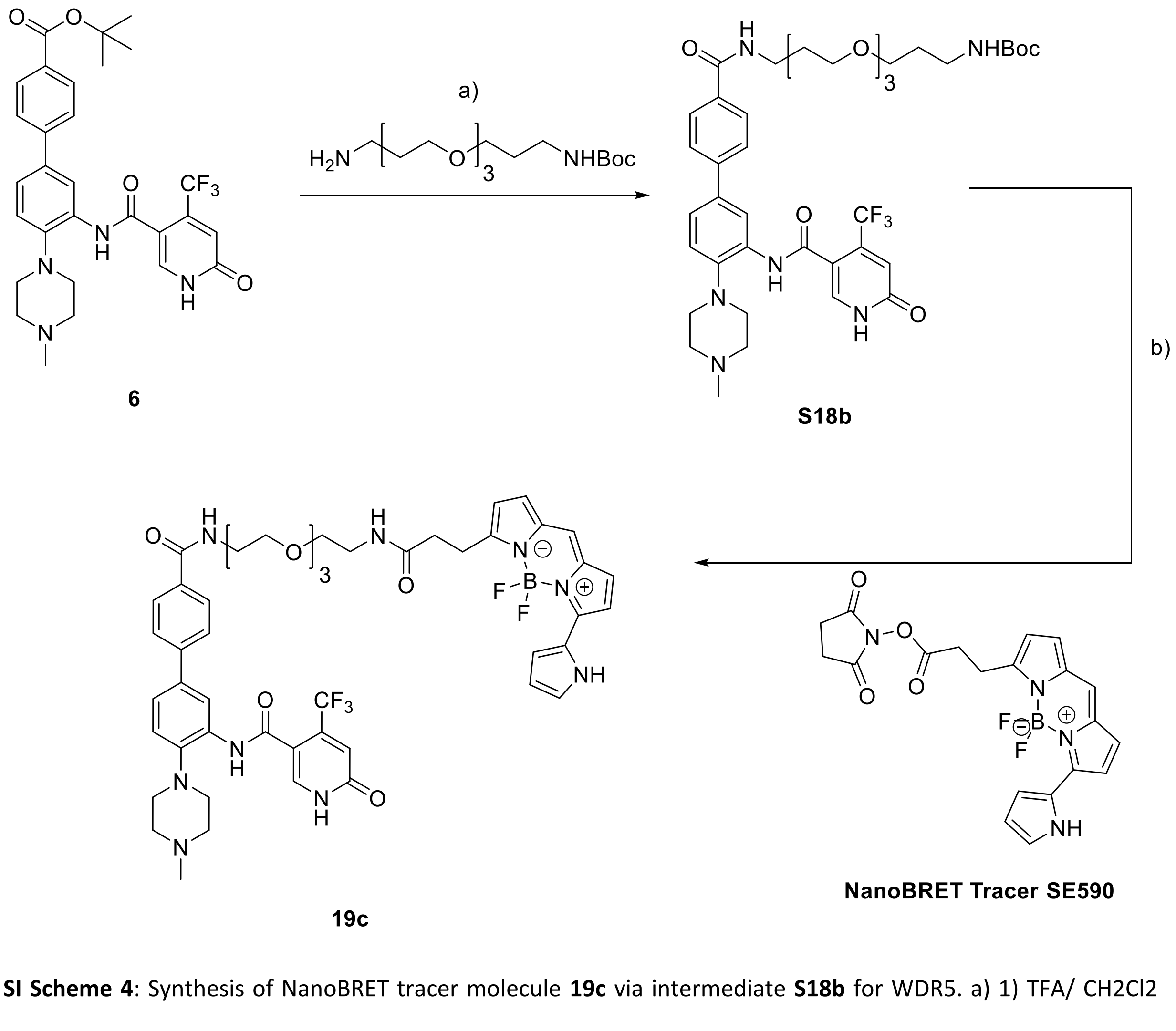


**SI Scheme 4**: Synthesis of NanoBRET tracer molecule **19c** via intermediate **S18b** for WDR5. a) 1) TFA/ CH2Cl2 (1:1), rt, 2h; 2) HATU, DIEA, DMF, rt, 3h; b) DIEA, DMF, rt, 3 h.

**Synthesis of tert-butyl (1-(3'-(6-hydroxy-4-(trifluoromethyl)nicotinamido)-4'-(4-methylpiperazin-1-yl)-[1,1'-biphenyl]-4-yl)-1-oxo-6,9,12-trioxa-2-azapentadecan-15-yl)carbamate (S18b)**

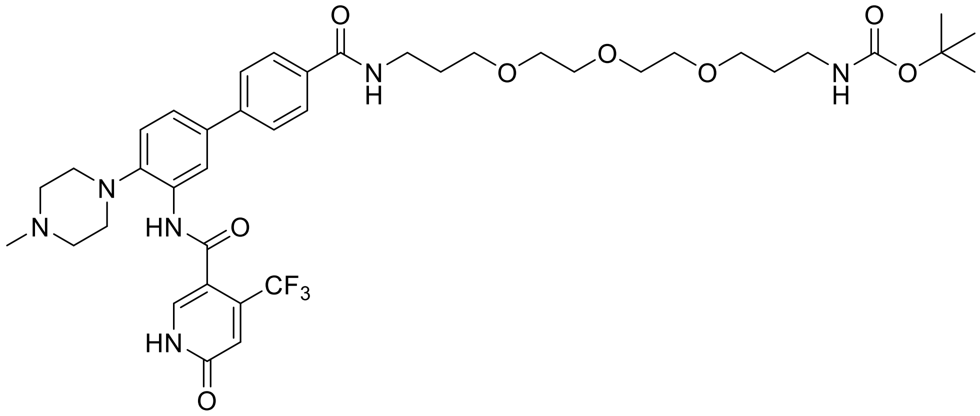


84 mg (151 µmol, 1.0 eq) tert-butyl 3'-(6-hydroxy-4-(trifluoromethyl)nicotinamido)-4'-(4-methylpiperazin-1-yl)-[1,1'-biphenyl]-4-carboxylate were dissolved in 1 mL DCM and 1 mL TFA and stirred at rt for 1 h. Excess solvent was evapored. The solid was dissolved in 2 mL DMF, then 526 µL (3.02 mmol, 20 eq) DIEA and 68.9 mg (181 µmol, 1.2 eq) HATU were added. After 15 min, a solution of 50.8 mg (158 µmol, 1.05 eq) tert-butyl (3-(2-(2-(3-aminopropoxy)ethoxy)ethoxy)propyl)carbamate in 1 mL DMF was added. The solution was stirred for 3 h at rt. The reaction mixture was quenched with 2 mL water and 2 mL saturated $NaHCO_3$, then the reaction was extracted 3x with EA. The organic phase was dried over $MgSO_4$, filtered and the solvent was removed under reduced pressure. The crude product was purified using by HPLC to obtain 94.5 mg, 118 µmol, 78% of a colourless oil.

Yield: 94.5 mg, 118 µmol, 78% of a colourless oil.

ESI: (calculated) $[M+Na^+]$ 825.37 g/mol

(found) $[M+Na^+]$ 825.05 g/mol.

HPLC: RT = 11.9 min (254nm, 100%).

$^1$H NMR (600 MHz, MeOD) δ= 8.29 (s, 1H), 8.00 (s, 1H), 7.89 (d, $^3J$ = 8.4 Hz, 2H, 7.71 (d, $^3J$ = 8.4 Hz, 2H), 7.51 (dd, $^3J$ = 8.3 Hz, $^4J$ = 2.0 Hz), 7.33 (d, $^3J$ = 8.3 Hz, 1H), 6.92 (s, 1H), 3.68 – 3.64 (m, 2H), 3.63 – 3.57 (m, 6H), 3.55 – 3.51 (m, 2H), 3.50 (t, $^3J$ = 6.7 Hz, 2H), 3.46 (t, $^3J$ = 6.2 Hz, 2H), 3.09 (t, $^3J$ = 6.8 Hz, 2H), 3.01 (t, $^3J$ = 4.2 Hz, 4H), 2.69 (s, 4H), 2.39 (s, 3H), 1.96 – 1.82 (m, 2H), 1.76 – 1.62 (m, 2H), 1.41 (s, 9H) ppm.

$^{13}$C NMR (151 MHz, MeOD) δ = 169.7, 165.0, 164.3, 164.3, 145.3, 144.7, 141.41 (d, J = 33.7 Hz), 140.5, 139.3, 137.5, 134.4, 133.9, 129.5, 128.4, 127.3, 125.0, 123.4 (q, J = 275 Hz), 123.2, 122.7, 121.6, 119.8 (m), 114.6 (d, J = 8.2 Hz), 79.8, 72.5, 71.6, 71.3, 71.2, 70.6, 70.4, 56.2, 52.4, 45.5, 38.6, 37.8, 30.4, 30.0, 29.2, 28.3 ppm.

**Synthesis of N-(4'-((17-(5,5-difluoro-7-(1H-pyrrol-2-yl)-5H-5l4,6l4-dipyrrolo[1,2-c:2',1'-f][1,3,2]diazaborinin-3-yl)-15-oxo-4,7,10-trioxa-14-azaheptadecyl)carbamoyl)-4-(4-methylpiperazin-1-yl)-[1,1'-biphenyl]-3-yl)-6-hydroxy-4-(trifluoromethyl)nicotinamide (19c)**

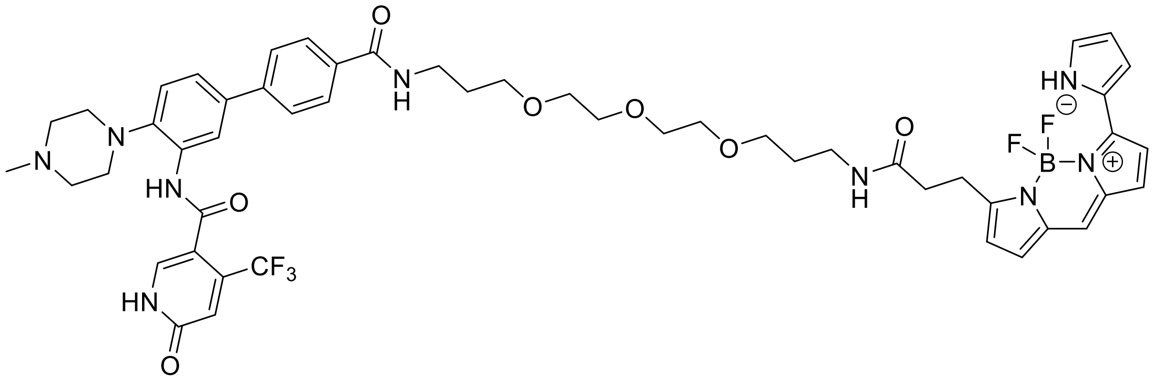

10 mg (12 µmol, 1.0 eq) tert-butyl (1-(3'-(6-hydroxy-4-(trifluoromethyl)nicotinamido)-4'-(4-methylpiperazin-1-yl)-[1,1'-biphenyl]-4-yl)-1-oxo-6,9,12-trioxa-2-azapentadecan-15-yl)carbamate were dissolved in 0.5 mL DCM and 0.5 mL TFA and stired at rt for 1 h. Excess solvent was evapored. The solid was dissolved in 0.5 mL DMF, then 17 µL (10 µmol, 2.0 eq) DIEA and 5.8 mg (14 µmol, 1.1 eq)

2,5-dioxopyrrolidin-1-yl-3-(5,5-difluoro-7-(1H-pyrrol-2-yl)-5H-5l4,6l4-dipyrrolo[1,2-c:2',1'-f][1,3,2]diazaborinin-3-yl)propanoate in 0.5 mL DMF was added. The solution was stirred for 3 h at rt. The reaction mixture was quenched with 2 mL water and 2 mL saturated $NaHCO_3$, then the reaction was extracted 3x with EA. The organic phase was dried over $MgSO_4$, filtered and the solvent was removed under reduced pressure. The crude product was purified using by HPLC to obtain 8.07 mg, 7.96 µmol, 66% of a purple solid.

Yield: 8.1 mg, 7.9 µmol, 66% of a purple solid.

$R_f$ (20% MeOH/ $CH_2Cl_2$): 0.76.

HPLC: RT = 12.4 min (254 nm, 100%).

HRMS: (calculated) $[M+Na^+]$ 1036.4286 g/mol

(found) $[M+Na^+]$ 1036.4303 g/mol.

$^1$H NMR (600 MHz, MeOD) δ = 8.26 (d, $^4J$ = 1.9 Hz, 1H), 8.03 (s, 1H), 7.88 (d, $^3J$ = 8.3 Hz, 2H), 7.70 (d, $^3J$ = 8.4 Hz, 2H), 7.54 (dd, $^3J$ = 8.4 Hz, $^4J$ = 2.0 Hz, 1H), 7.35 (d, $^3J$ = 8.4 Hz, 1H), 7.23 – 7.13 (m, 4H), 6.99 (d, $^4J$ = 4.6 Hz, 1H), 6.94 (s, 1H), 6.89 (d, $^4J$ = 3.9 Hz, 1H), 6.38 – 6.32 (m, 1H), 6.28 (d, 4*J* = 3.9 Hz, 1H), 3.62 - 3.57 (m, 10H), 3.50 - 3.47 (m, 4H), 3.43 (t, $^3J$ = 6.1 Hz, 2H), 3.28 - 3.22 (m, 8H), 3.13 (t, $^3J$ = 11.6 Hz, 2H), 2.94 (s, 3H), 2.59 (t, $^3J$ = 7.7 Hz, 2H), 1.87 (p, $^3J$ = 6.3 Hz, 2H), 1.70 (p, $^3J$ = 6.4 Hz, 2H) ppm.
