## Supplementary material for "Design, Synthesis and Evaluation of WD-repeat containing protein 5 (WDR5) degraders": Chemistry supplement

**Table of content**

**1.** Appendix

1.1 Appendix for E3 Ligase Linker L0-L15

1.2 Appendix for intermediates 6a-c, S18a,b and BRET Tracer molecules 19a-c

1.3 Appendix for intermediates 2-6

1.4 Appendix for heterobifunctional molecules 7a-e, 8a-j, 9a-c

1.5 Appendix for intermediates 11-16a-g

1.6 Appendix for heterobifunctional molecules 17a-g

1.7 Appendix for negative controls 20 and 21

### 1.1 Appendix for E3 Ligase Linker L0-L15

#### ESI, $^1$H-NMR of 2-(2,6-dioxopiperidin-3-yl)-4-fluoroisoindoline-1,3-dione L0

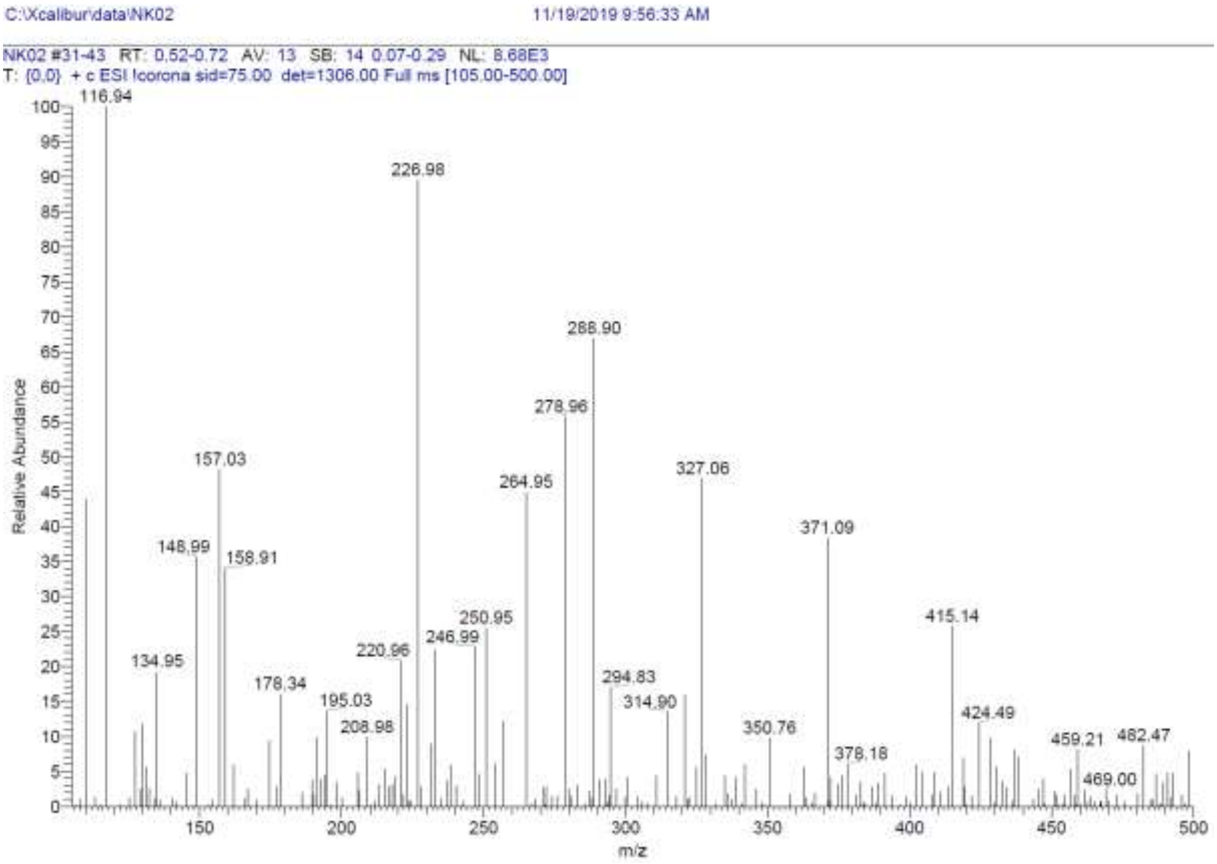

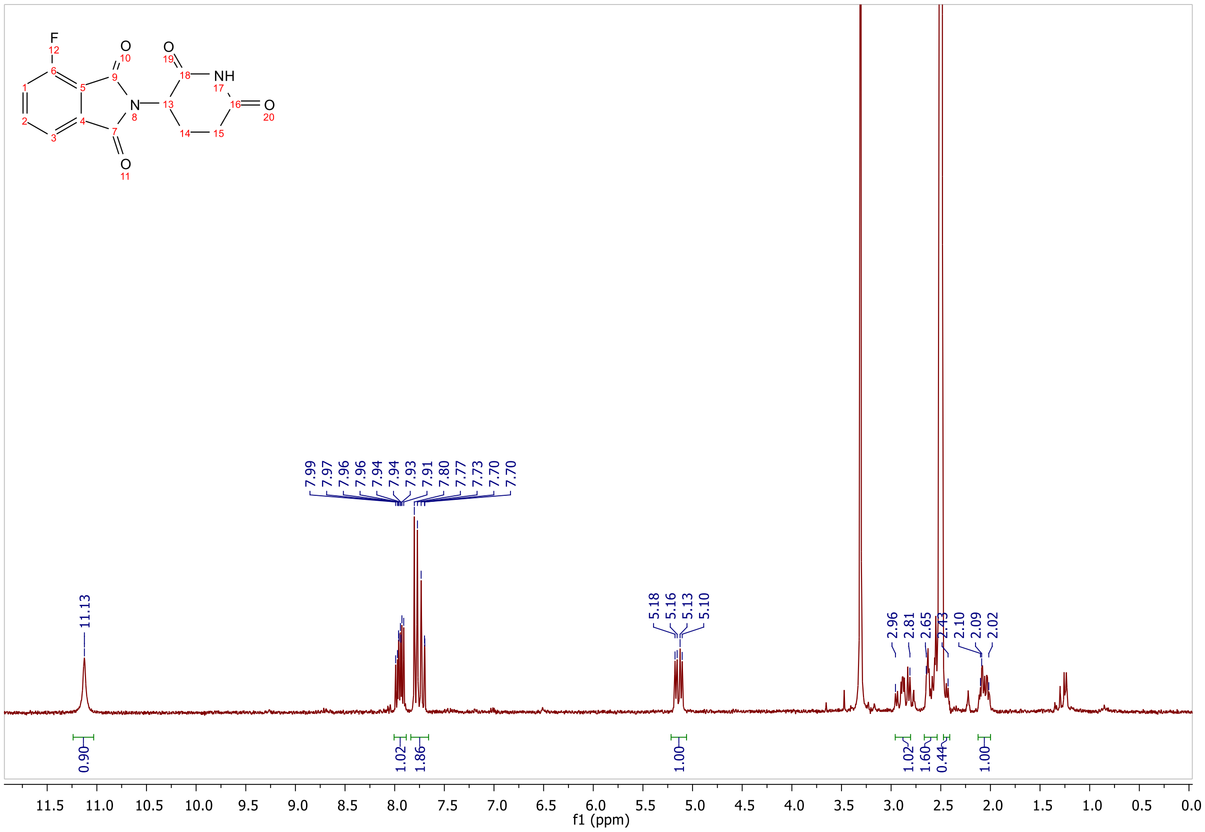

**ESI, HPLC, $^{1}$H-NMR and $^{13}$C-NMR of tert-butyl (2-(2-((2-(2,6-dioxopiperidin-3-yl)-1,3-dioxoisoindolin-4-yl)amino)ethoxy)ethyl)carbamate L1**

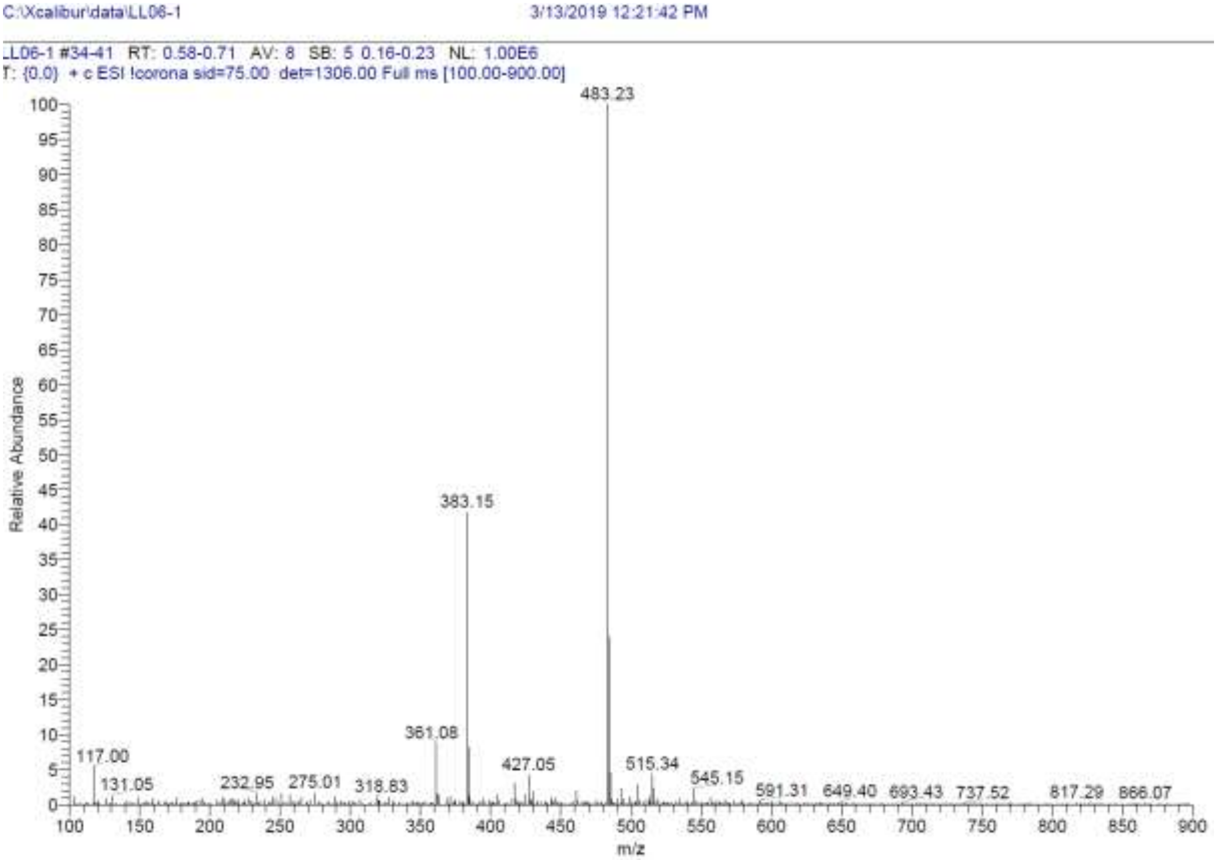

**Signal:** MWD1 A, Sig=254,4 Ref=off

| RT [min] | Type | Width [min] | Area | Height | Area% | Name |
|---|---|---|---|---|---|---|
| 3.313 | BV | 0.0953 | 471.4035 | 74.0216 | 0.8010 | |
| 12.868 | MM | 0.2825 | 58377.2656 | 3443.8289 | 99.1990 | |
| | | Sum | 58848.6691 | | | |

**Signal:** MWD1 B, Sig=280,4 Ref=off

| RT [min] | Type | Width [min] | Area | Height | Area% | Name |
|---|---|---|---|---|---|---|
| 12.805 | MM | 0.2581 | 53369.2109 | 3445.7402 | 100.0000 | |
| | | Sum | 53369.2109 | | | |

**Signal:** MWD1 F, Sig=260,4 Ref=off

| RT [min] | Type | Width [min] | Area | Height | Area% | Name |
|---|---|---|---|---|---|---|
| 3.313 | VV | 0.0926 | 356.4311 | 58.9223 | 0.5828 | |
| 12.774 | MM | 0.2846 | 60802.2852 | 3561.0684 | 99.4172 | |

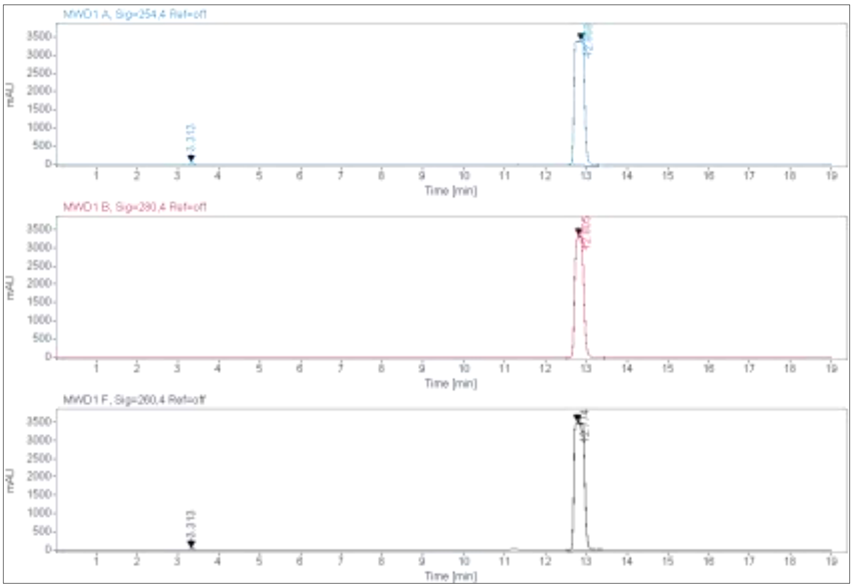
MWD1 A, Sig=254,4 Ref=off
MWD1 B, Sig=280,4 Ref=off
MWD1 F, Sig=260,4 Ref=off
mAU
Time [min]
3.313

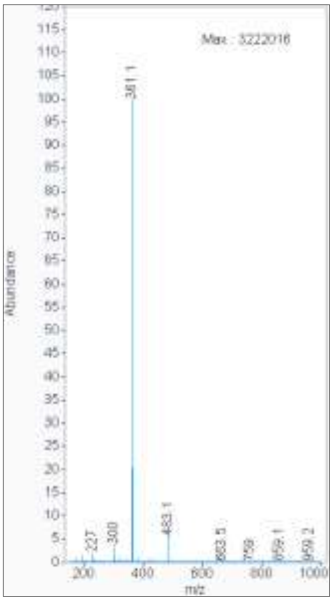
Max: 3222016
361.1
483.1
Abundance
m/z

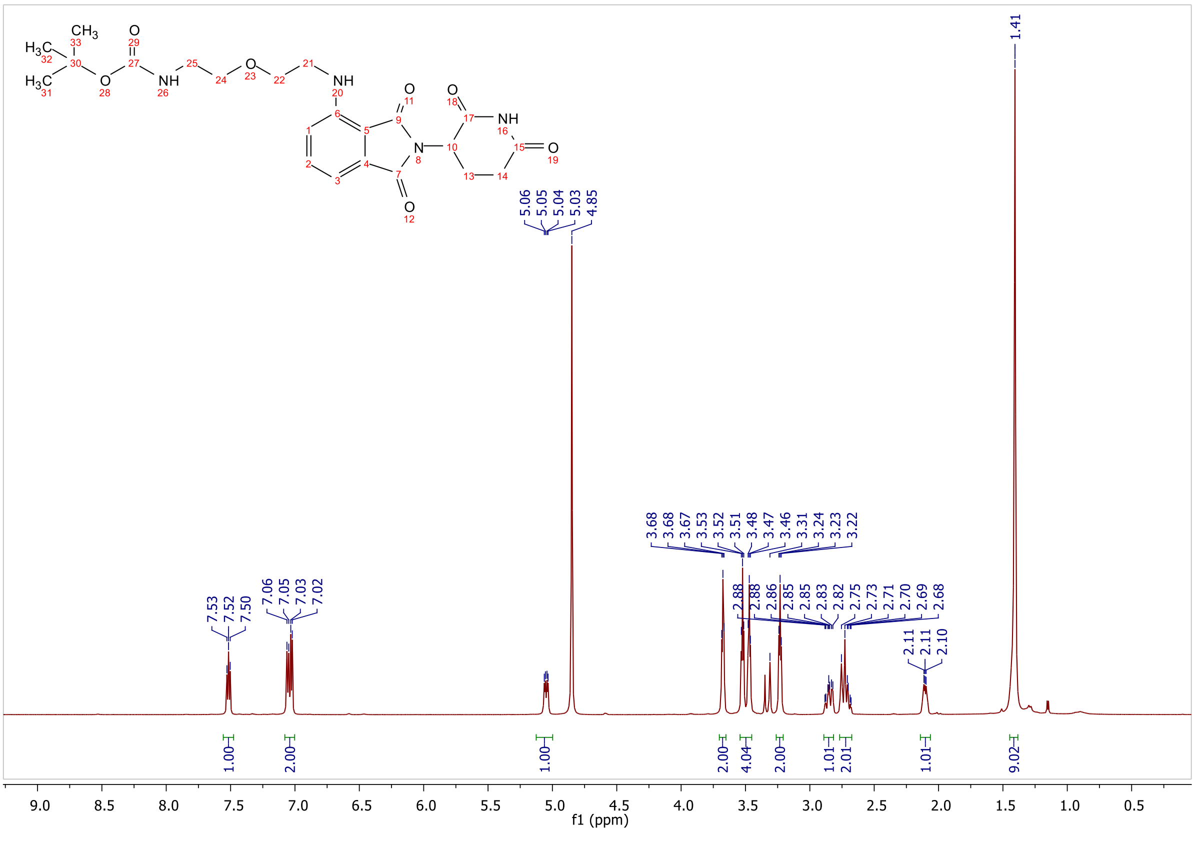
H3C
CH3
O
NH
N
f1 (ppm)
1.41

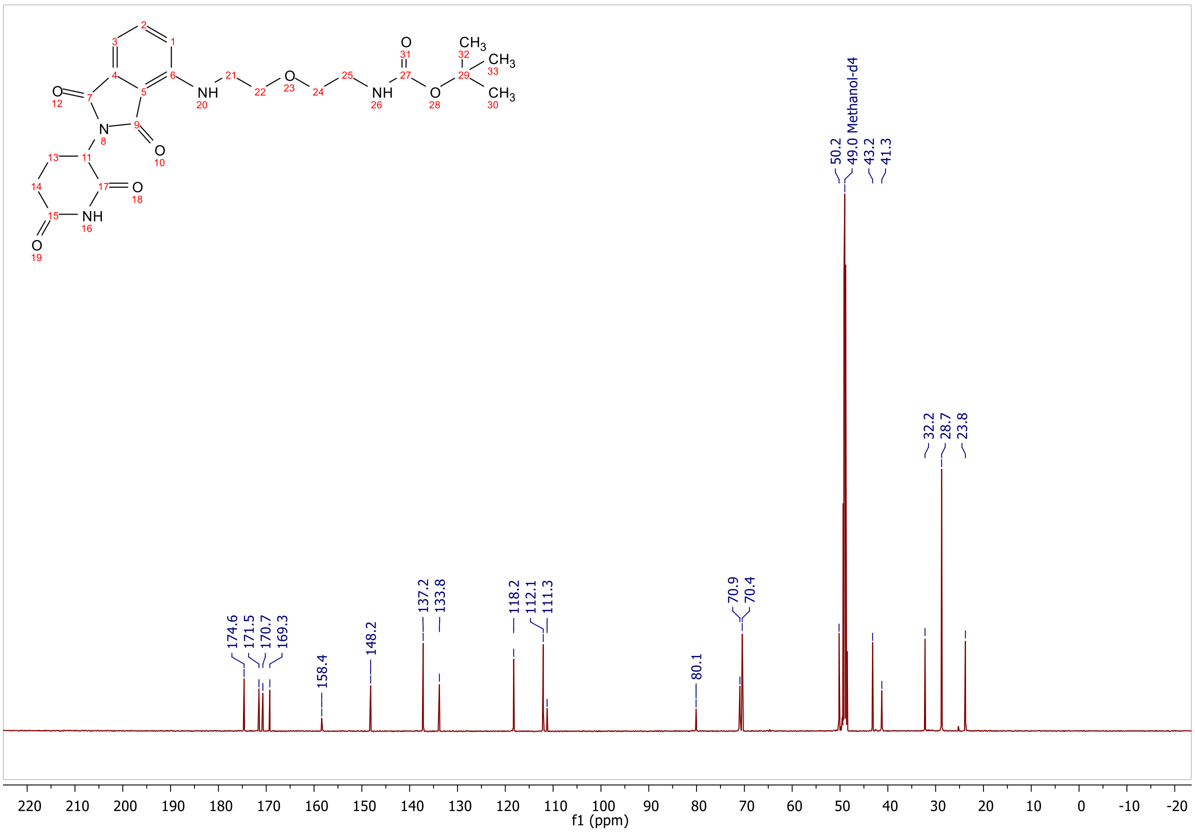

NH
O
NH
O
O
CH3
CH3
CH3
O
N
O
O
NH
O
174.6
171.5
170.7
169.3
158.4
148.2
137.2
133.8
118.2
112.1
111.3
80.1
70.9
70.4
50.2
49.0 Methanol-d4
43.2
41.3
32.2
28.7
23.8
220 210 200 190 180 170 160 150 140 130 120 110 100 90 80 70 60 50 40 30 20 10 0 -10 -20
f1 (ppm)

**ESI, HPLC, [1]H-NMR and [13]C-NMR of tert-butyl (2-(2-(2-(2-((2-(2,6-dioxopiperidin-3-yl)-1,3-dioxoisoindolin-4-yl)amino)ethoxy)ethoxy)ethoxy)ethyl)carbamate L2**

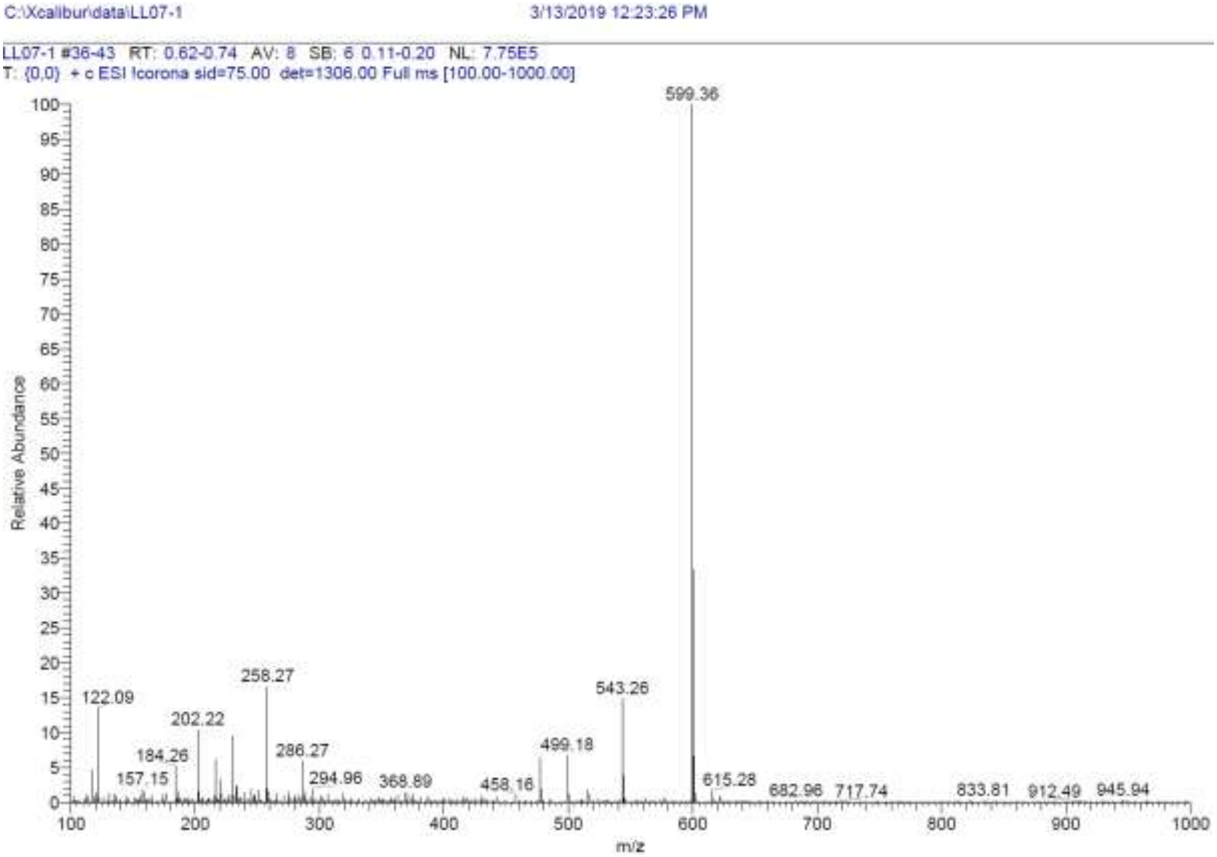

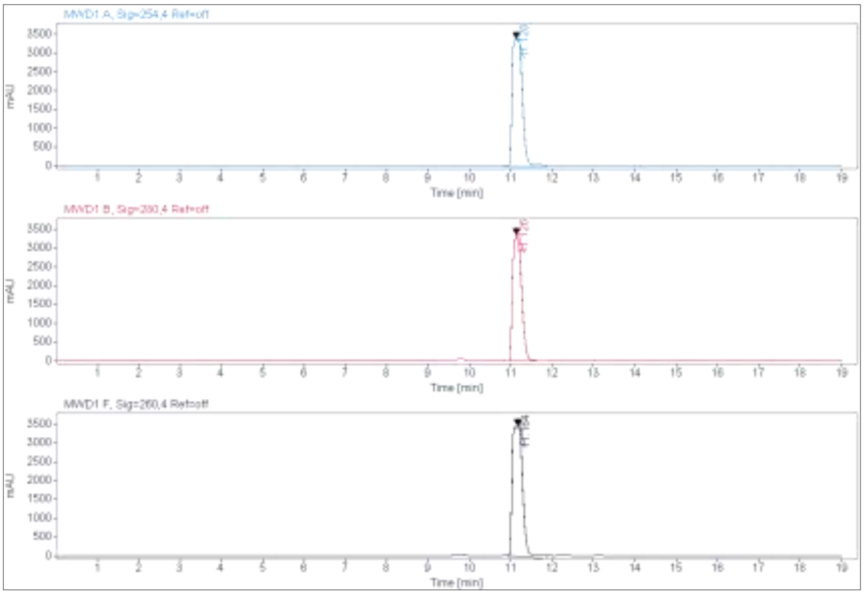

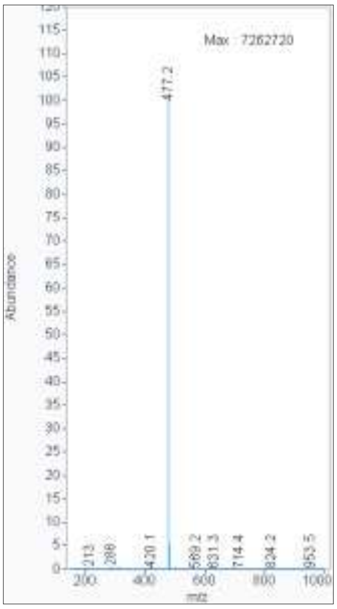

**Signal:** MWD1 A, Sig=254,4 Ref=off

| RT [min] | Type | Width [min] | Area | Height | Area% | Name |
|---|---|---|---|---|---|---|
| 11.128 | MM | 0.2843 | 58034.3398 | 3402.1462 | 100.0000 | |
| | | Sum | 58034.3398 | | | |

**Signal:** MWD1 B, Sig=280,4 Ref=off

| RT [min] | Type | Width [min] | Area | Height | Area% | Name |
|---|---|---|---|---|---|---|

| | | | | | |
|---|---|---|---|---|---|
| 11.126 | VV | 0.1779 | 50324.2734 | 3357.0464 | 100.0000 |
| | | Sum | 50324.2734 | | |

**Signal:** MWD1 F, Sig=260,4 Ref=off

| RT [min] | Type | Width [min] | Area | Height | Area% | Name |
|---|---|---|---|---|---|---|
| 11.164 | MM | 0.2940 | 61209.2969 | 3470.2668 | 100.0000 | |

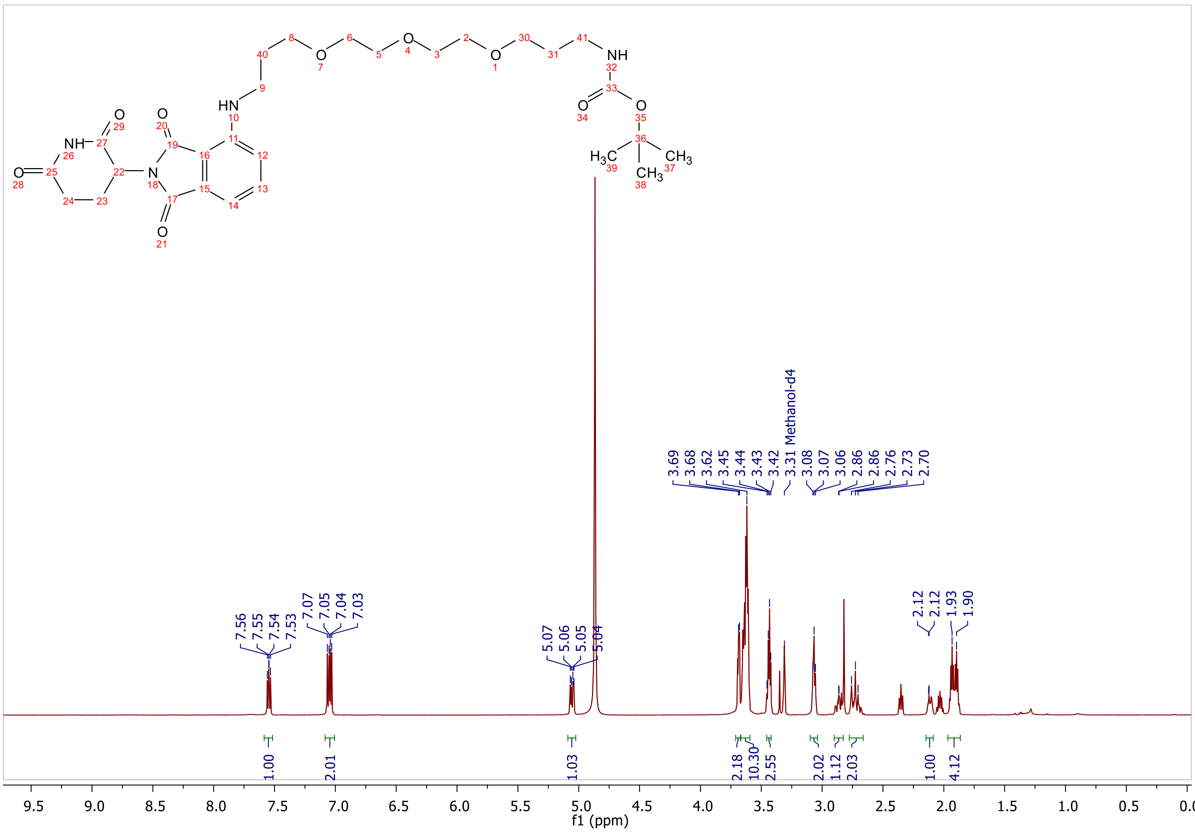

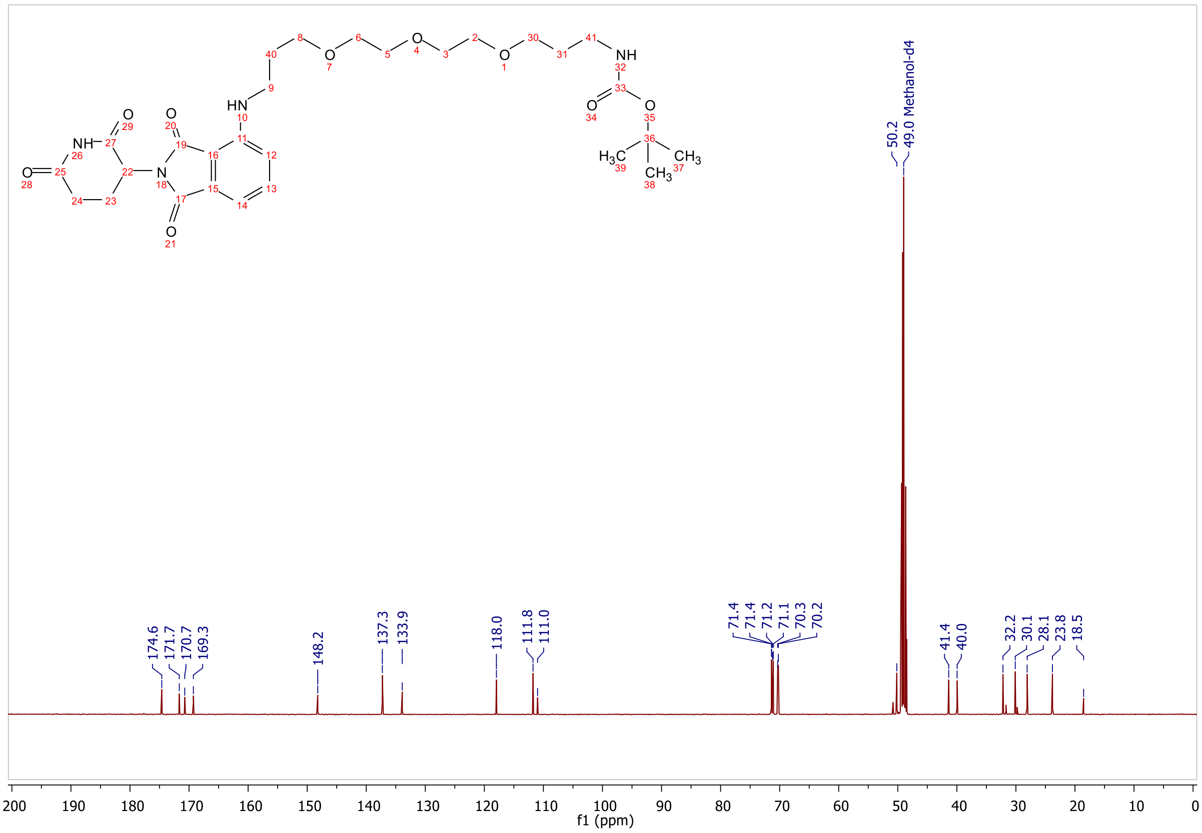
174.6
171.7
170.7
169.3
148.2
137.3
133.9
118.0
111.8
111.0
71.4
71.4
71.2
71.1
70.3
70.2
50.2
49.0 Methanol-d4
41.4
40.0
32.2
30.1
28.1
23.8
18.5
f1 (ppm)

**ESI, HPLC, $^{1}$H-NMR of tert-butyl (17-((2-(2,6-dioxopiperidin-3-yl)-1,3-dioxoisoindolin-4-yl)amino)-3,6,9,12,15-pentaoxaheptadecyl)carbamate L3**

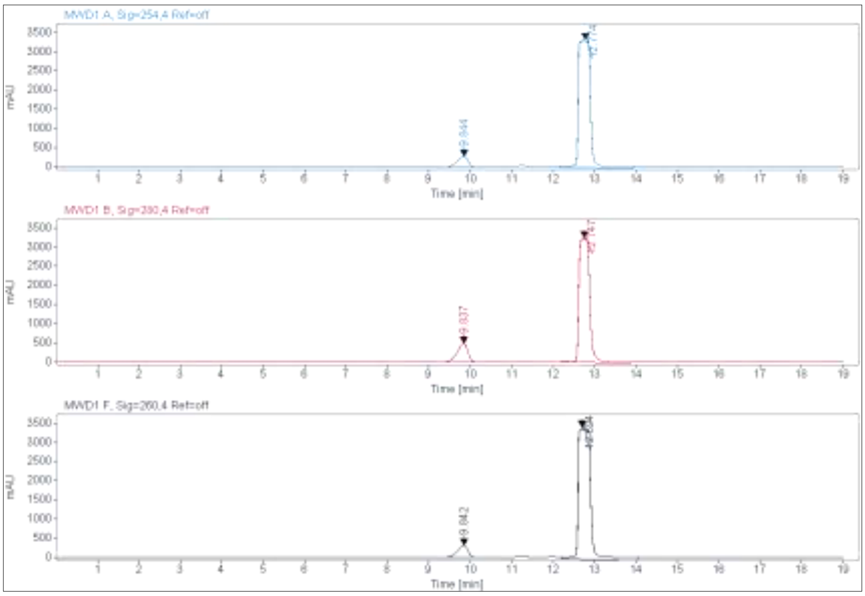

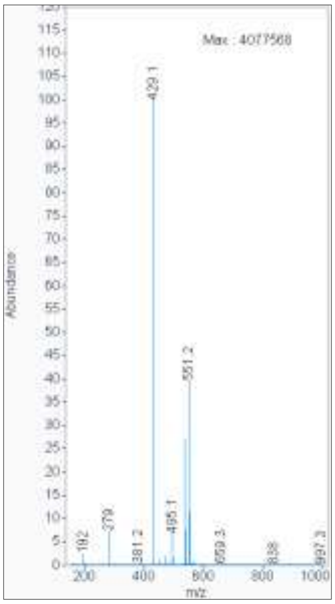

**Signal:** MWD1 A, Sig=254,4 Ref=off

| RT [min] | Type | Width [min] | Area | Height | Area% | Name |
|---|---|---|---|---|---|---|
| 9.844 | VV | 0.2442 | 4875.1143 | 277.2130 | 6.8073 | |
| 12.774 | MM | 0.3324 | 66741.1328 | 3346.6414 | 93.1927 | |
| | | Sum | 71616.2471 | | | |

**Signal:** MWD1 B, Sig=280,4 Ref=off

| RT [min] | Type | Width [min] | Area | Height | Area% | Name |
|---|---|---|---|---|---|---|
| 9.837 | VV | 0.2515 | 8863.7100 | 489.0014 | 13.2680 | |
| 12.747 | MM | 0.2964 | 57941.5820 | 3258.0781 | 86.7320 | |
| | | Sum | 66805.2920 | | | |

**Signal:** MWD1 F, Sig=260,4 Ref=off

| RT [min] | Type | Width [min] | Area | Height | Area% | Name |
|---|---|---|---|---|---|---|
| 9.842 | VV | 0.2456 | 5633.9717 | 318.0999 | 7.2044 | |
| 12.694 | MM | 0.3503 | 72567.7969 | 3452.9812 | 92.7956 | |

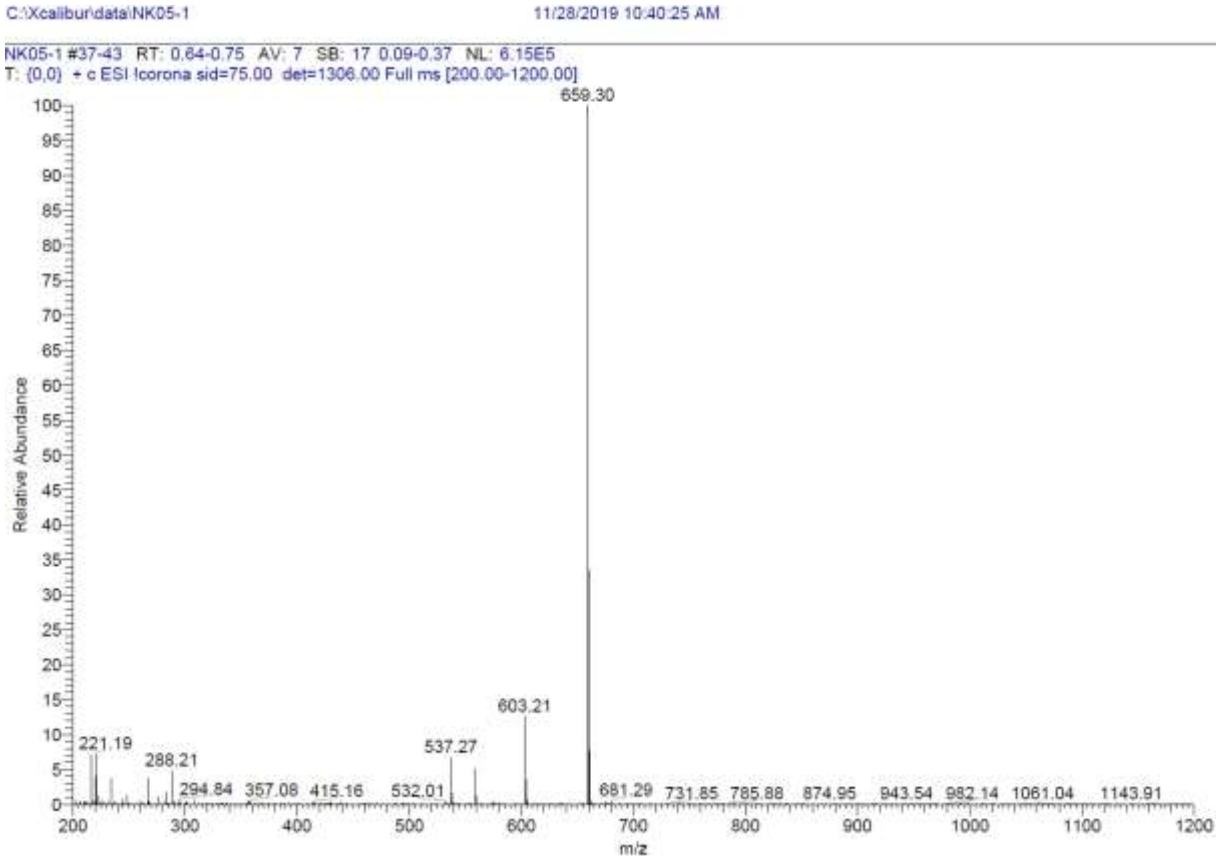
C:\Xcalibur\data\NK05-1
11/28/2019 10:40:25 AM
NK05-1 #37-43 RT: 0.64-0.75 AV: 7 SB: 17 0.09-0.37 NL: 6.15E5
T: {0,0} + c ESI !corona sid=75.00 det=1306.00 Full ms [200.00-1200.00]
659.30
603.21
537.27
221.19
288.21
294.84
357.08
415.16
532.01
681.29
731.85
785.88
874.95
943.54
982.14
1061.04
1143.91
Relative Abundance
m/z

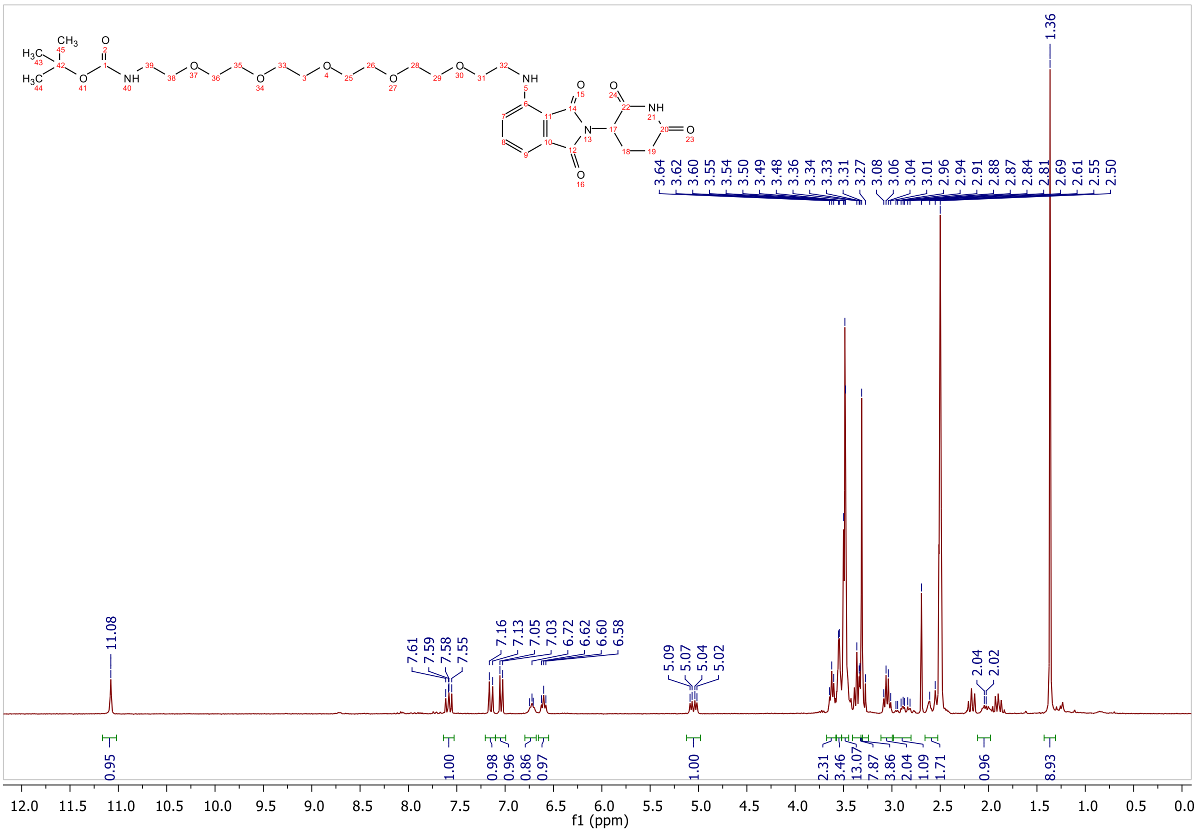
11.08
7.61
7.59
7.58
7.55
7.16
7.13
7.05
7.03
6.72
6.62
6.60
6.58
5.09
5.07
5.04
5.02
3.64
3.62
3.60
3.55
3.54
3.50
3.49
3.48
3.36
3.34
3.33
3.31
3.27
3.08
3.06
3.04
3.01
2.96
2.94
2.91
2.88
2.87
2.84
2.81
2.69
2.61
2.55
2.50
2.04
2.02
1.36
0.95
1.00
0.98
0.96
0.86
0.97
1.00
2.31
3.46
13.07
7.87
3.86
2.04
1.09
1.71
0.96
8.93
f1 (ppm)

### ESI, HPLC, $^{1}H$-NMR and $^{13}C$-NMR of tert-butyl (23-((2-(2,6-dioxopiperidin-3-yl)-1,3-dioxoisoindolin-4-yl)amino)-3,6,9,12,15,18,21-heptaoxatricosyl)carbamate L4

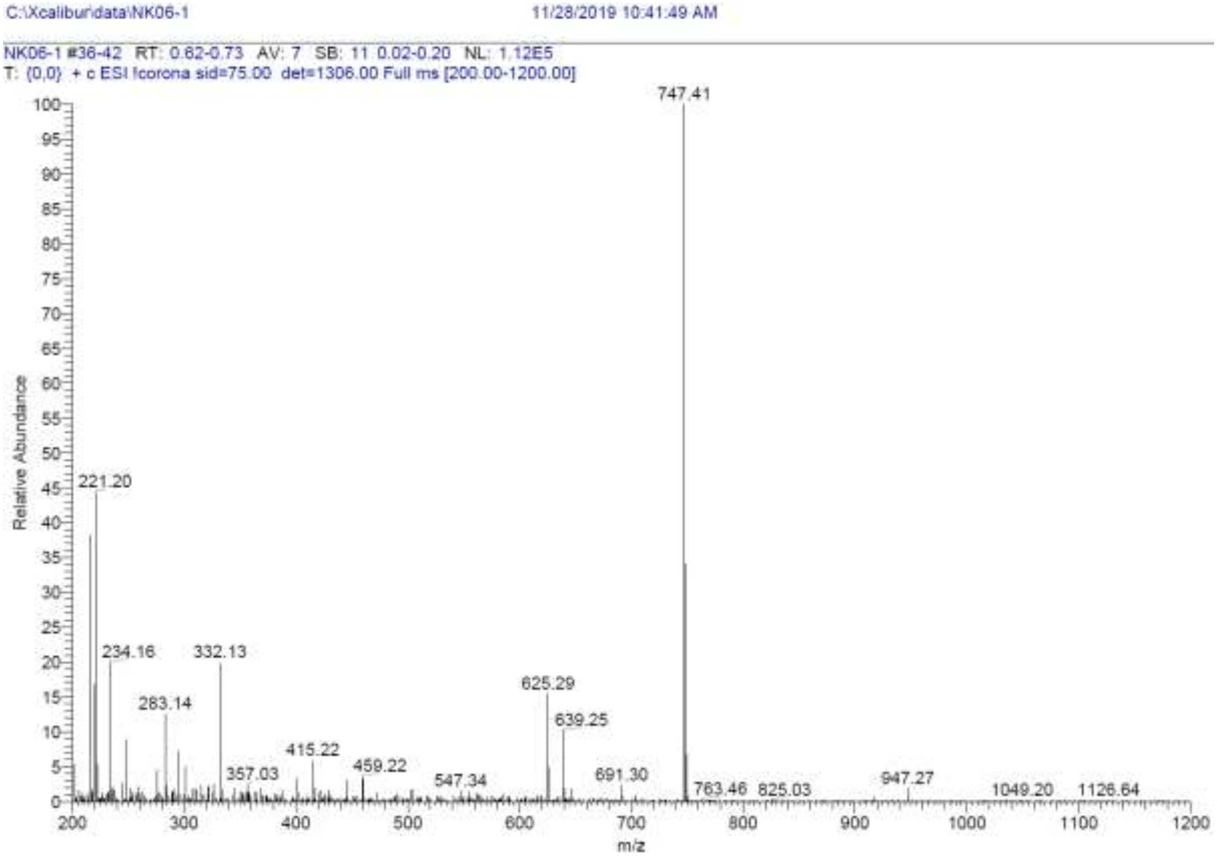

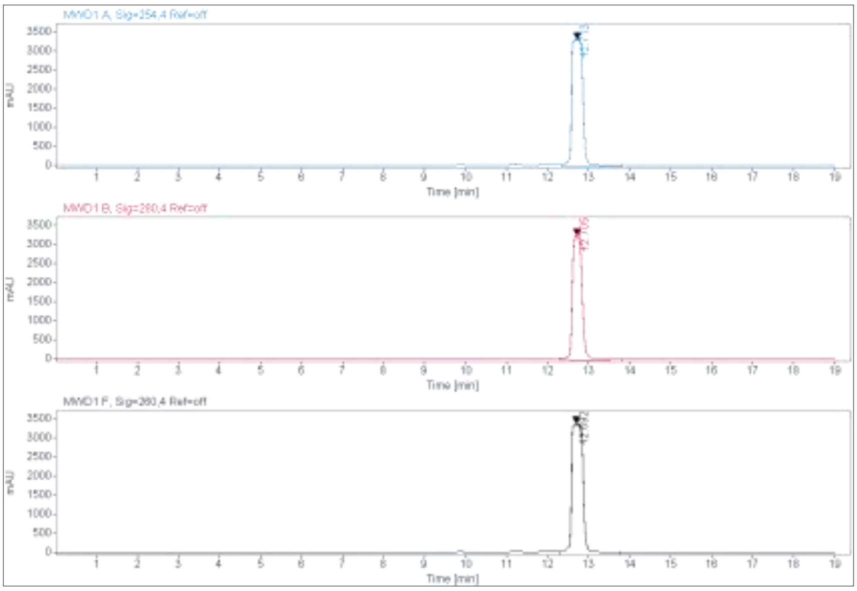

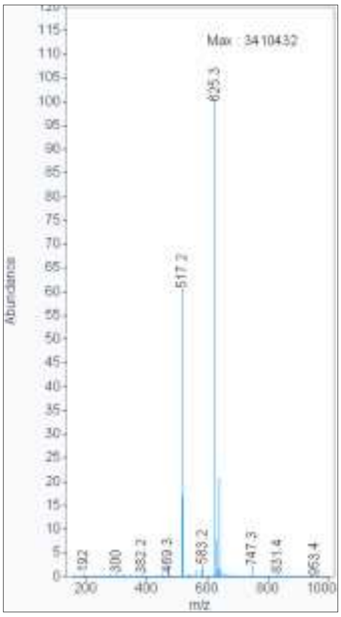

**Signal:** MWD1 A, Sig=254,4 Ref=off

| RT [min] | Type | Width [min] | Area | Height | Area% | Name |
|---|---|---|---|---|---|---|
| 12.713 | MM | 0.3183 | 64034.5742 | 3352.9675 | 100.0000 | |
| | | Sum | 64034.5742 | | | |

**Signal:** MWD1 B, Sig=280,4 Ref=off

| RT [min] | Type | Width [min] | Area | Height | Area% | Name |
|---|---|---|---|---|---|---|

| | | | | | |
|---|---|---|---|---|---|
| 12.705 | MM | 0.2795 | 54819.2344 | 3269.3167 | 100.0000 |
| | | Sum | 54819.2344 | | |

**Signal:** MWD1 F, Sig=260,4 Ref=off

| RT [min] | Type | Width [min] | Area | Height | Area% | Name |
|---|---|---|---|---|---|---|
| 12.692 | MM | 0.3332 | 69116.1953 | 3456.9661 | 100.0000 | |

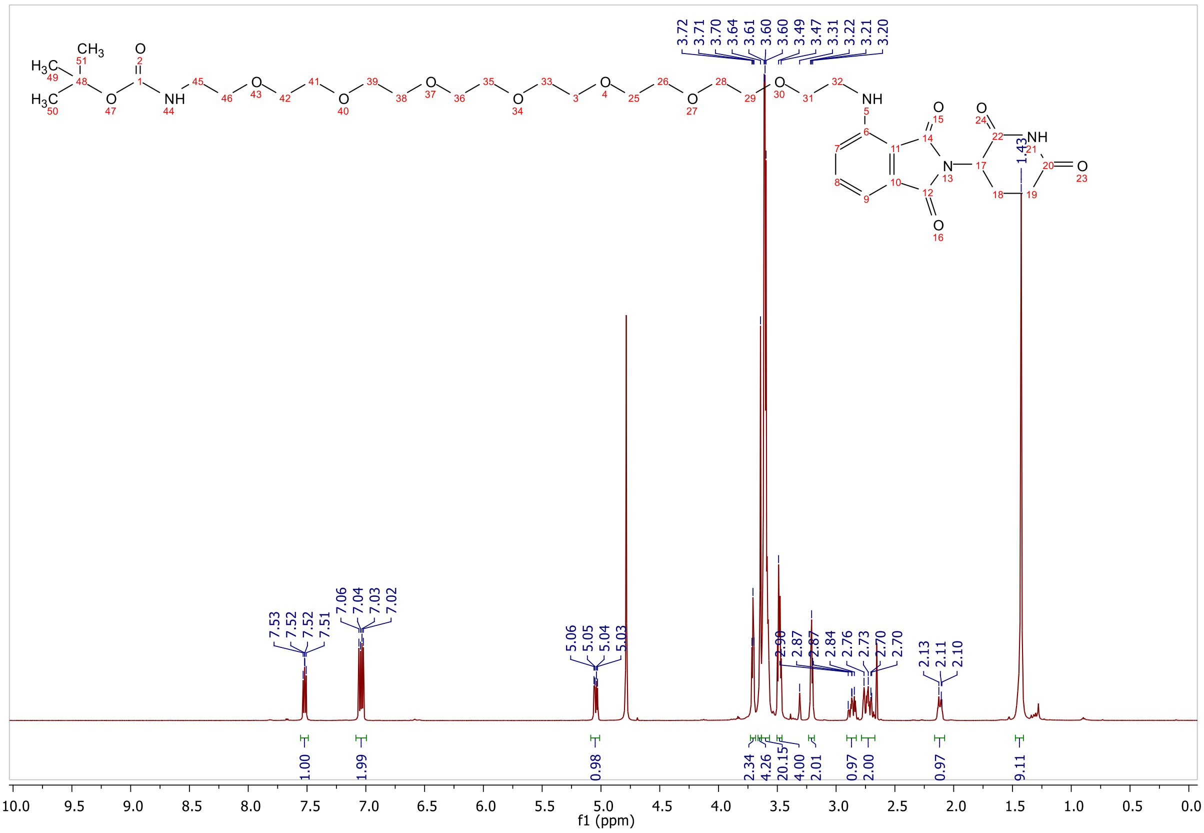

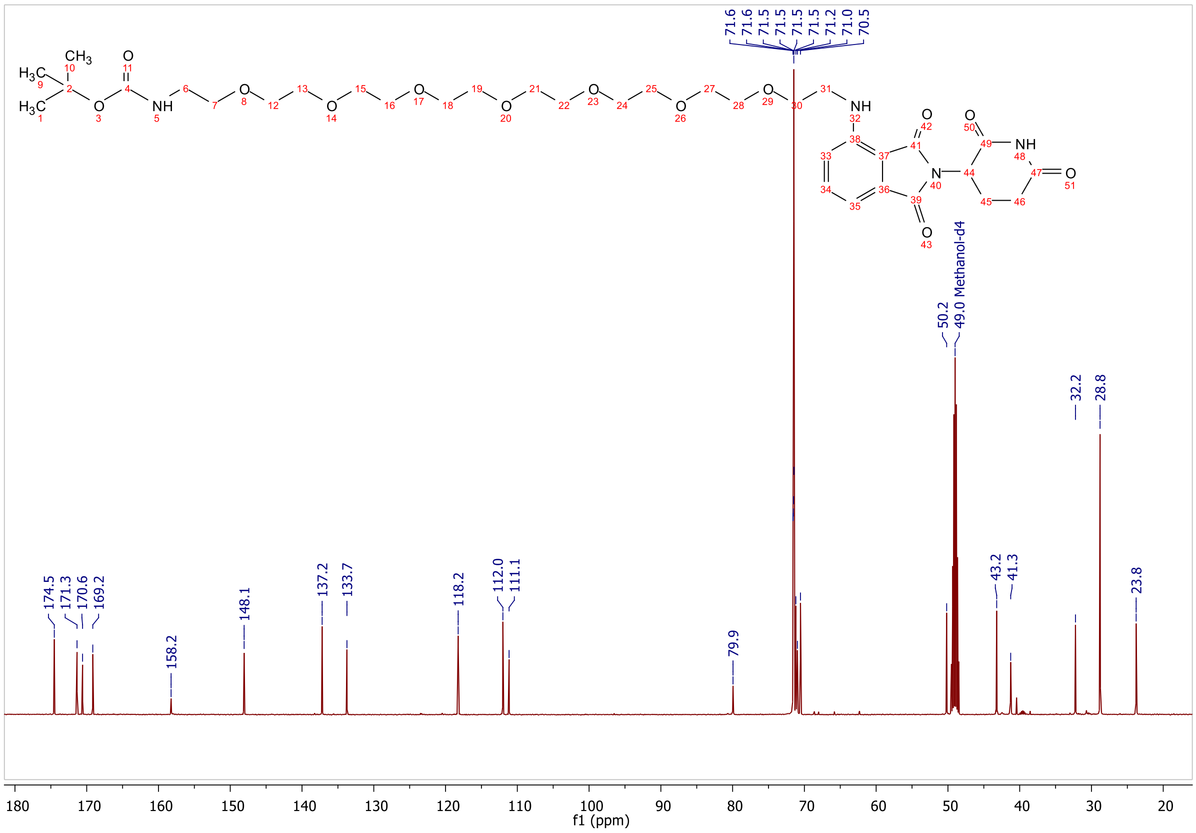
71.6
71.6
71.5
71.5
71.5
71.5
71.2
71.0
70.5
50.2
49.0 Methanol-d4
32.2
28.8
174.5
171.3
170.6
169.2
158.2
148.1
137.2
133.7
118.2
112.0
111.1
79.9
43.2
41.3
23.8
f1 (ppm)

**ESI, HPLC, $^{1}$H-NMR of tert-butyl (4-(((2-(2,6-dioxopiperidin-3-yl)-1,3-dioxoisoindolin-4-yl)amino)methyl)benzyl)carbamate L5**

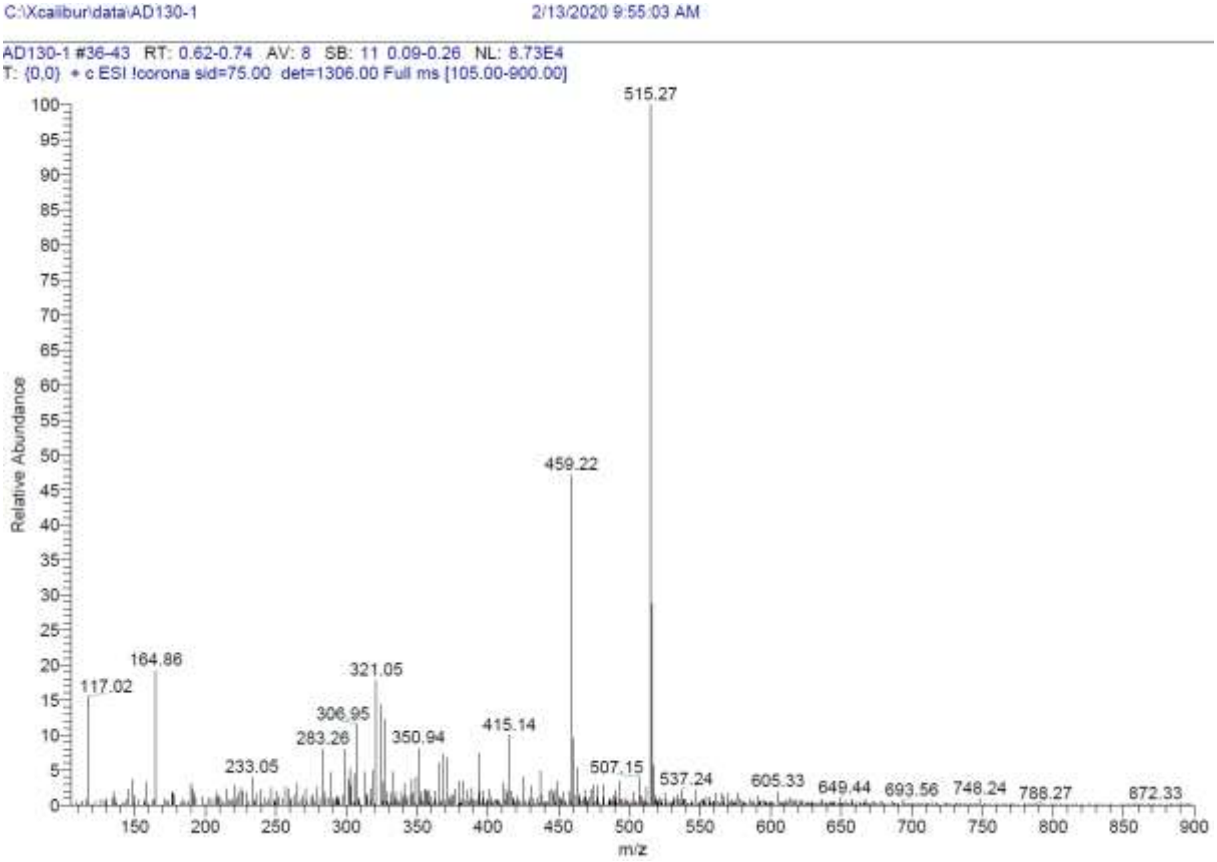

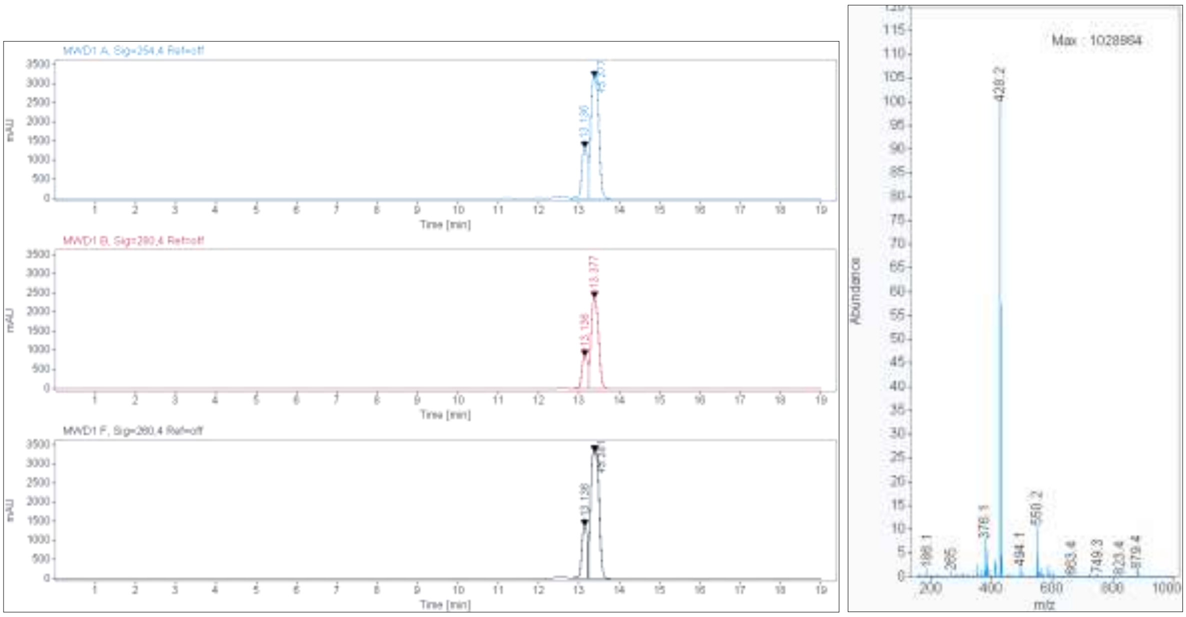

**Signal:** MWD1 A, Sig=254,4 Ref=off

| RT [min] | Type | Width [min] | Area | Height | Area% |
|---|---|---|---|---|---|
| 13.136 | VV | 0.1695 | 14003.7656 | 1322.6830 | 24.3641 |
| 13.377 | VV | 0.1652 | 43473.3477 | 3166.8340 | 75.6359 |
| | | Sum | 57477.1133 | | |

**Signal:** MWD1 B, Sig=280,4 Ref=off

| RT [min] | Type | Width [min] | Area | Height | Area% |
|---|---|---|---|---|---|

| | | | | | |
|---|---|---|---|---|---|
| 13.136 | VV | 0.1658 | 8677.5889 | 844.7153 | 22.1242 |
| 13.377 | VV | 0.1573 | 30544.4863 | 2370.8906 | 77.8758 |
| | | Sum | 39222.0752 | | |

**Signal:** MWD1 F, Sig=260,4 Ref=off

| RT [min] | Type | Width [min] | Area | Height | Area% |
|---|---|---|---|---|---|
| 13.136 | MM | 0.1585 | 12919.3477 | 1358.9250 | 21.3525 |
| 13.381 | VV | 0.1748 | 47585.8594 | 3308.0881 | 78.6475 |

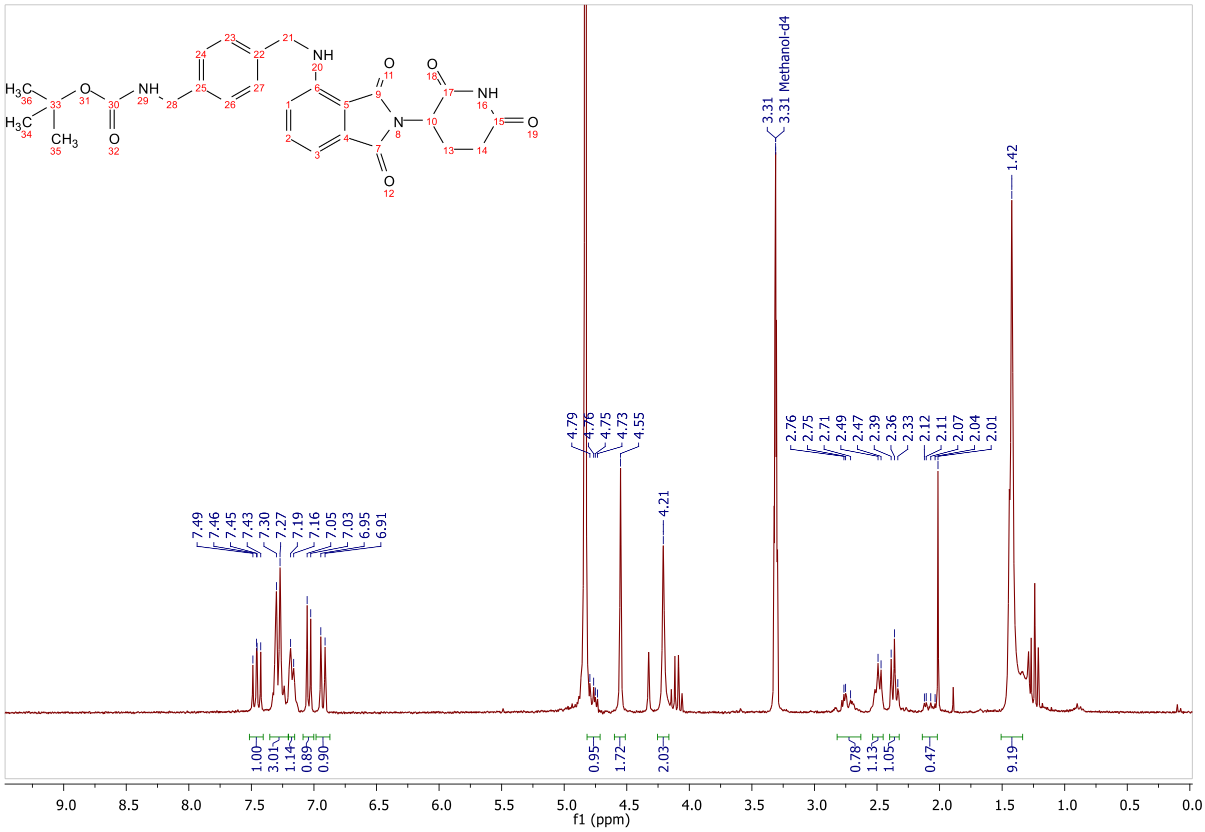

**ESI, HPLC, $^{1}$H-NMR and $^{13}$C-NMR of tert-butyl (2-(3-(((S)-1-((2S,4R)-4-hydroxy-2-((4-(4-methylthiazol-5-yl)benzyl)carbamoyl)pyrrolidin-1-yl)-3,3-dimethyl-1-oxobutan-2-yl)amino)-3-oxopropoxy)ethyl)carbamate L6**

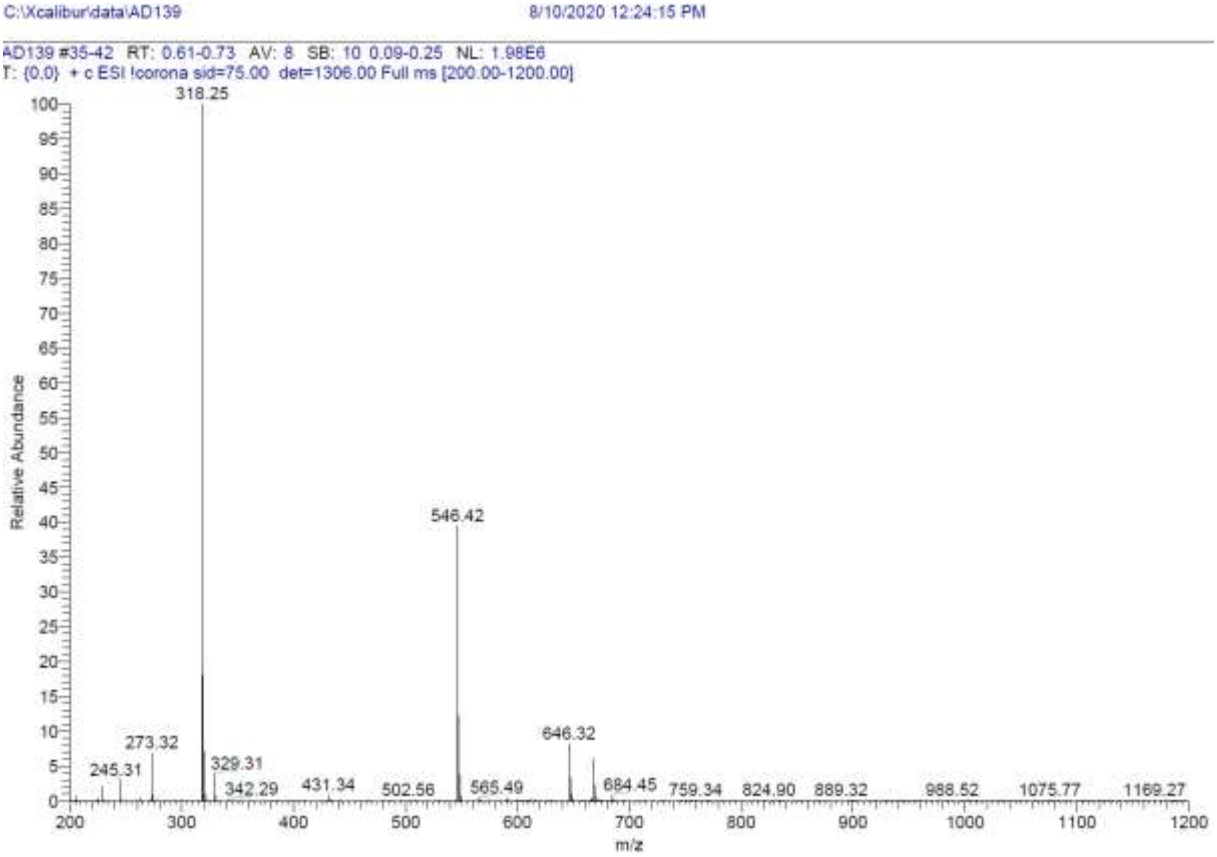

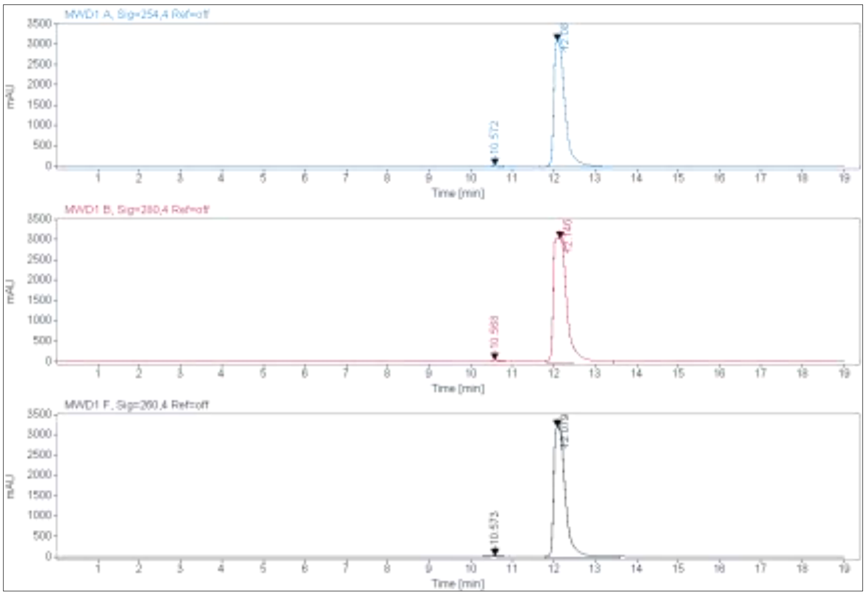

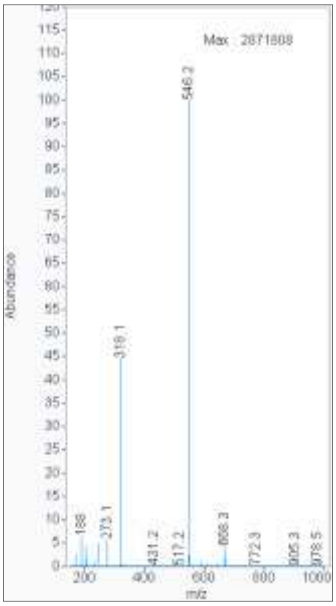

**Signal:** MWD1 A, Sig=254,4 Ref=off

| RT [min] | Type | Width [min] | Area | Height | Area% |
|---|---|---|---|---|---|
| 10.572 | MM | 0.2555 | 672.4956 | 43.8669 | 1.1729 |
| 12.080 | VV | 0.2221 | 56661.4414 | 3088.2632 | 98.8271 |
| | | Sum | 57333.9370 | | |

**Signal:** MWD1 B, Sig=280,4 Ref=off

| RT [min] | Type | Width [min] | Area | Height | Area% |
|---|---|---|---|---|---|
| 10.568 | VV | 0.2175 | 784.3678 | 51.8485 | 1.0468 |
| 12.146 | MM | 0.4020 | 74143.3516 | 3073.8215 | 98.9532 |
| | | Sum | 74927.7194 | | |

**Signal:** MWD1 F, Sig=260,4 Ref=off

| RT [min] | Type | Width [min] | Area | Height | Area% |
|---|---|---|---|---|---|
| 10.573 | MM | 0.2640 | 536.8670 | 33.8943 | 0.8248 |
| 12.079 | MM | 0.3343 | 64556.0977 | 3218.7188 | 99.1752 |

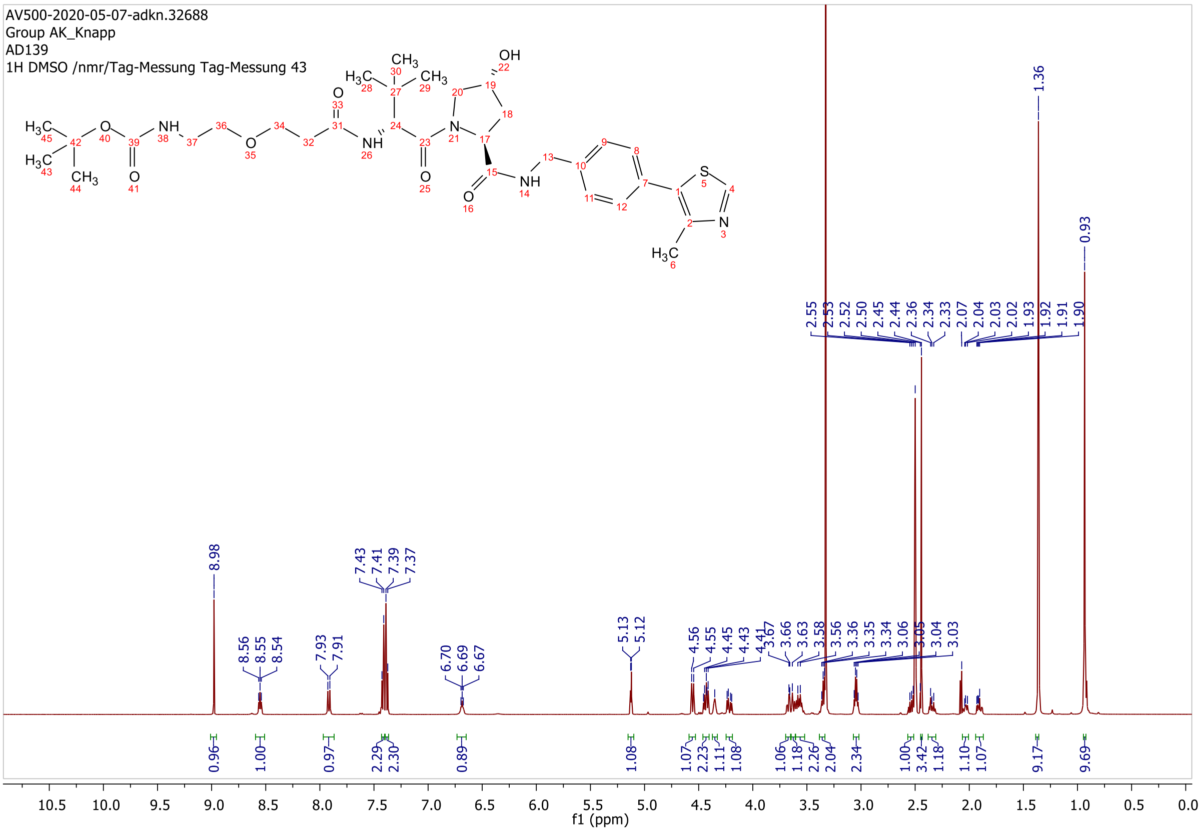

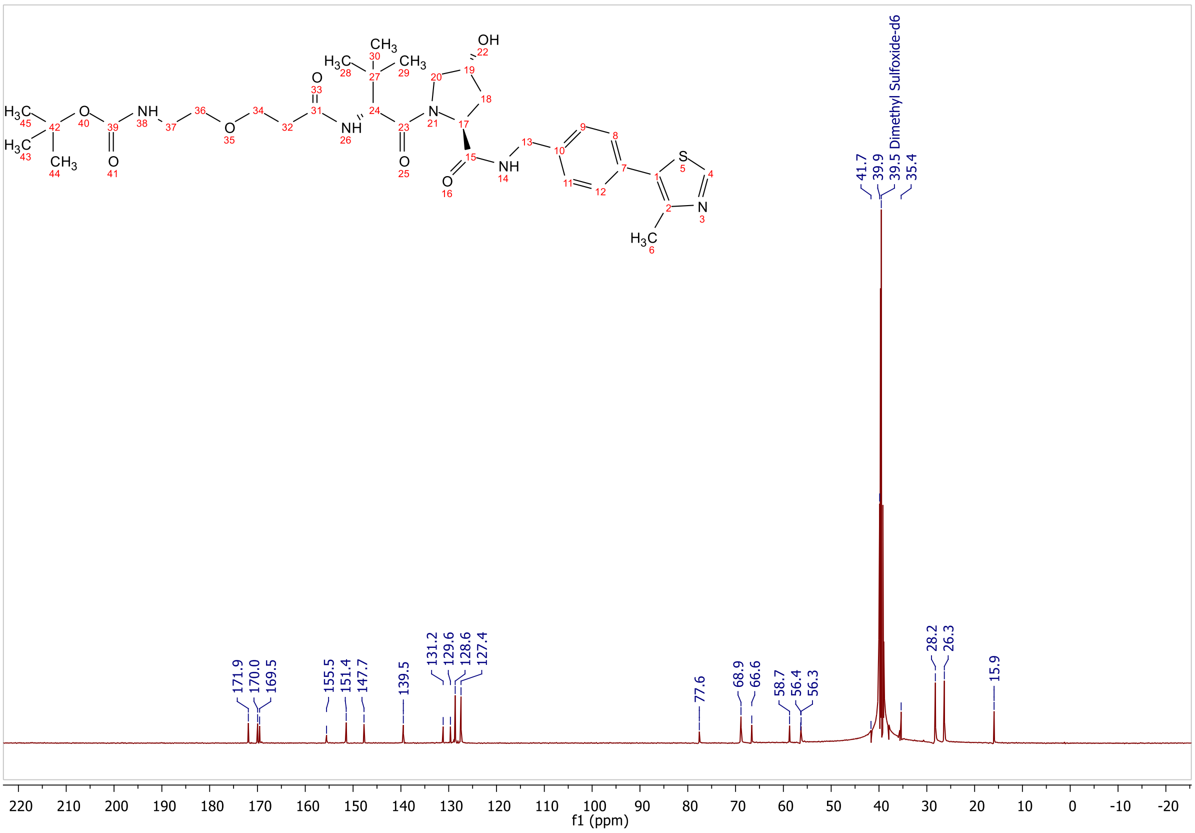

171.9
170.0
169.5
155.5
151.4
147.7
139.5
131.2
129.6
128.6
127.4
77.6
68.9
66.6
58.7
56.4
56.3
41.7
39.9
39.5 Dimethyl Sulfoxide-d6
35.4
28.2
26.3
15.9
f1 (ppm)

**ESI, HPLC, $^{1}$H-NMR and $^{13}$C-NMR of tert-butyl (2-(2-(3-(((S)-1-((2S,4R)-4-hydroxy-2-((4-(4-methylthiazol-5-yl)benzyl)carbamoyl)pyrrolidin-1-yl)-3,3-dimethyl-1-oxobutan-2-yl)amino)-3-oxopropoxy)ethoxy)ethyl)carbamate L7**

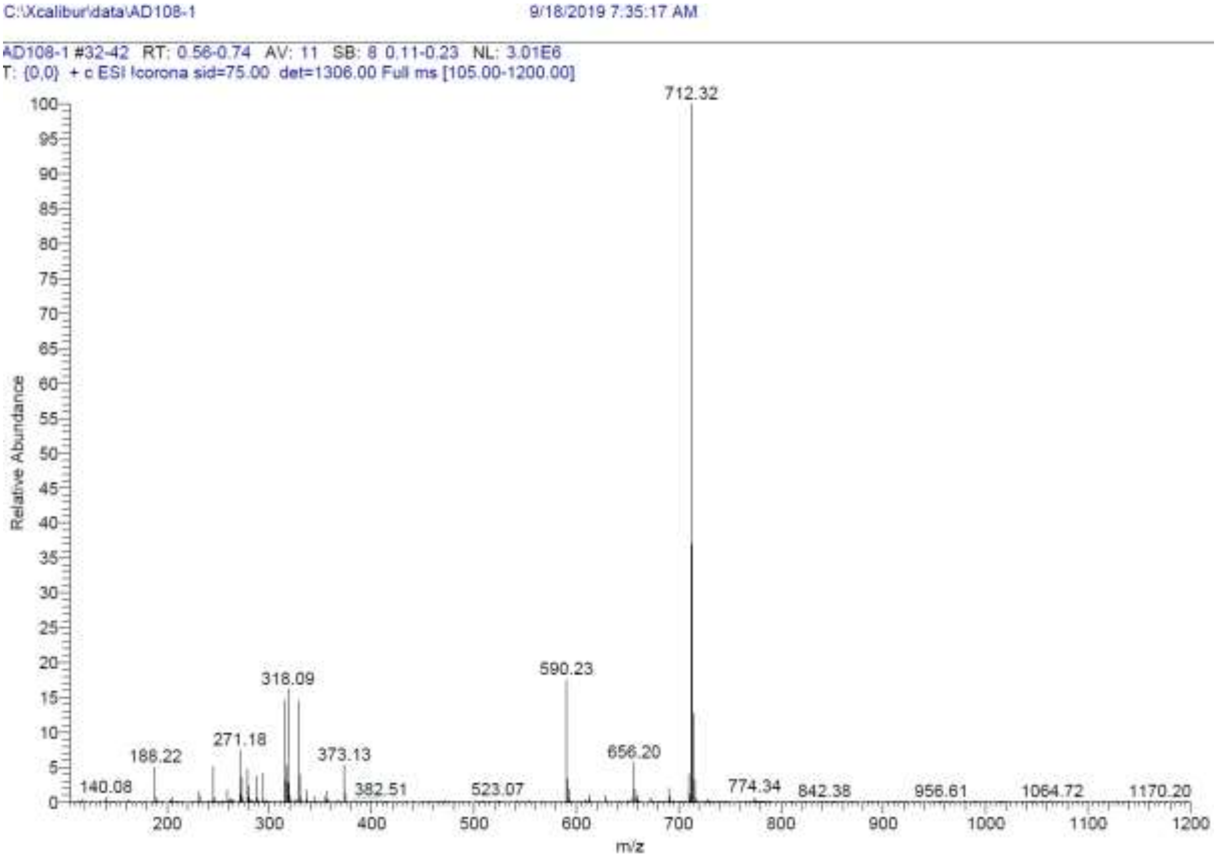

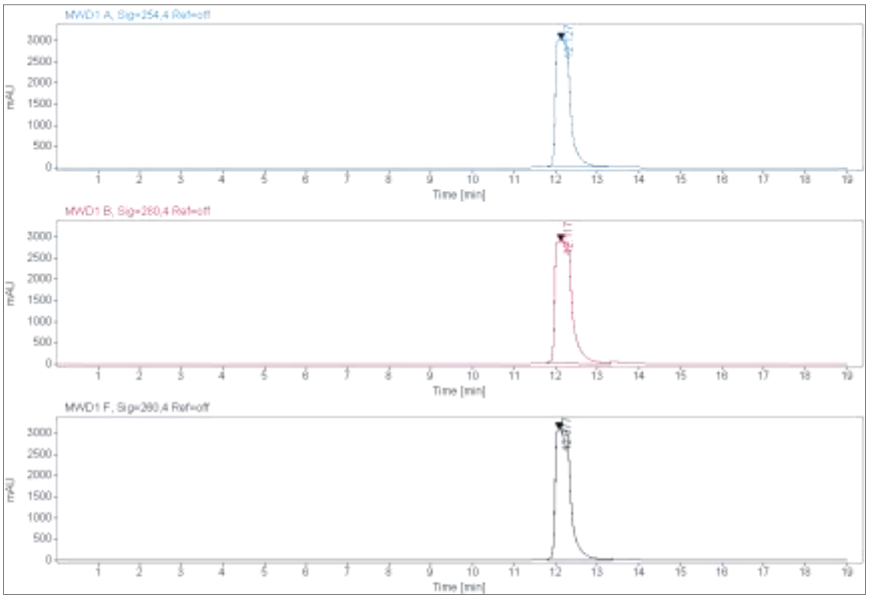

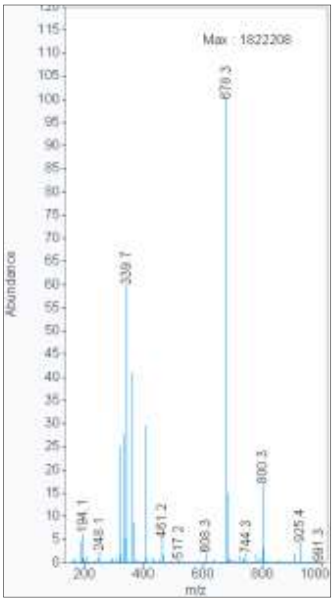

**Signal:** MWD1 A, Sig=254,4 Ref=off

| RT [min] | Type | Width [min] | Area | Height | Area% | Name |
|---|---|---|---|---|---|---|
| 12.127 | MM | 0.4125 | 74880.8047 | 3025.3701 | 100.0000 | |
| | | Sum | 74880.8047 | | | |

**Signal:** MWD1 B, Sig=280,4 Ref=off

| RT [min] | Type | Width [min] | Area | Height | Area% | Name |
|---|---|---|---|---|---|---|
| 12.117 | MM | 0.4749 | 82822.3672 | 2906.9238 | 100.0000 | |
| | | Sum | 82822.3672 | | | |

**Signal:** MWD1 F, Sig=260,4 Ref=off

| RT [min] | Type | Width [min] | Area | Height | Area% | Name |
|---|---|---|---|---|---|---|
| 12.077 | MM | 0.4308 | 80162.5781 | 3100.9529 | 100.0000 | |

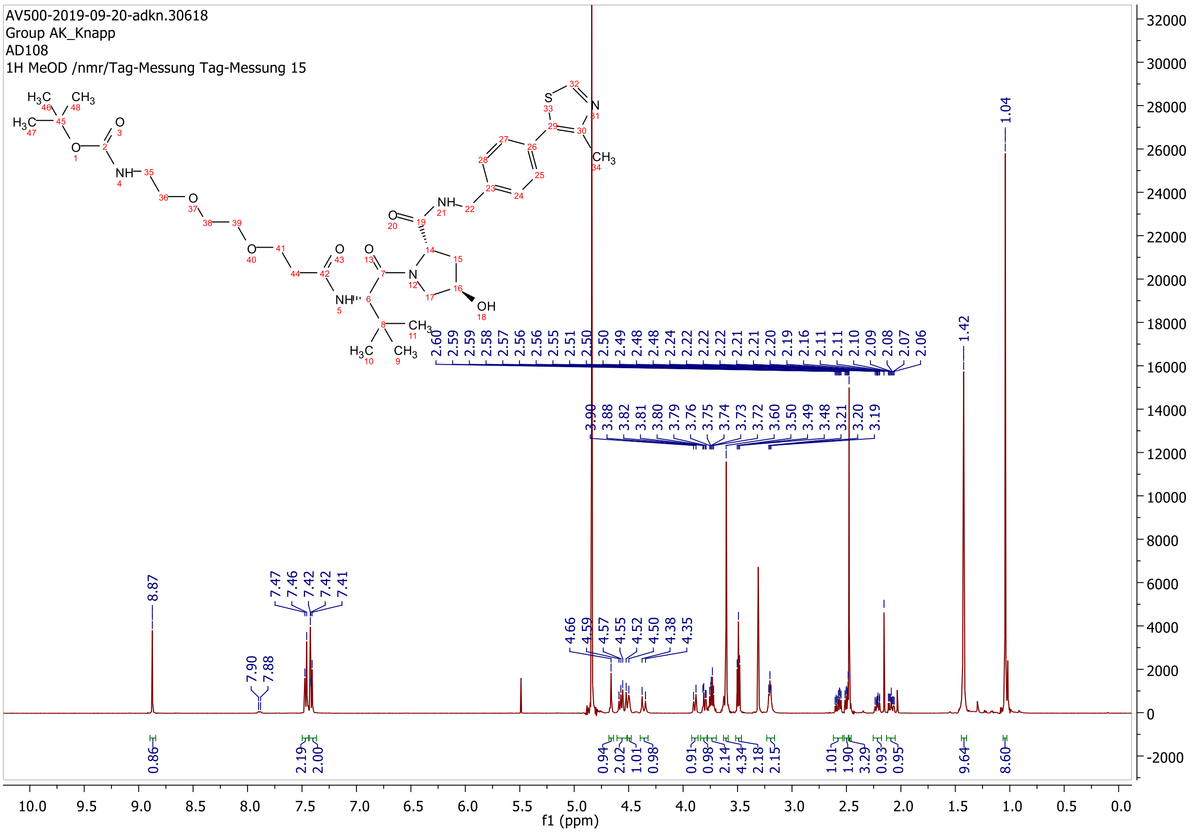

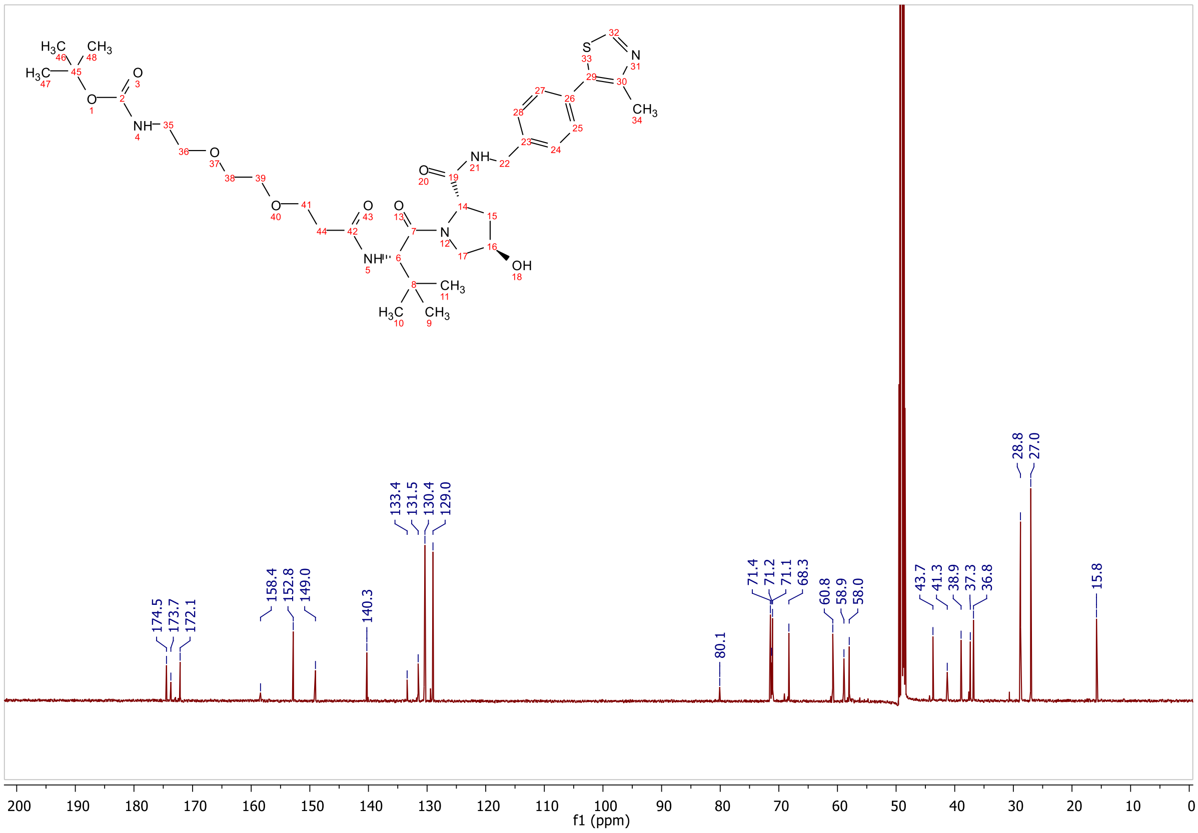

174.5
173.7
172.1
158.4
152.8
149.0
140.3
133.4
131.5
130.4
129.0
80.1
71.4
71.2
71.1
68.3
60.8
58.9
58.0
43.7
41.3
38.9
37.3
36.8
28.8
27.0
15.8
f1 (ppm)

**ESI, HPLC, $^1$H-NMR and $^{13}$C-NMR of tert-butyl ((S)-17-((2S,4R)-4-hydroxy-2-((4-(4-methylthiazol-5-yl)benzyl)carbamoyl)pyrrolidine-1-carbonyl)-18,18-dimethyl-15-oxo-3,6,9,12-tetraoxa-16-azanonadecyl)carbamate L8**

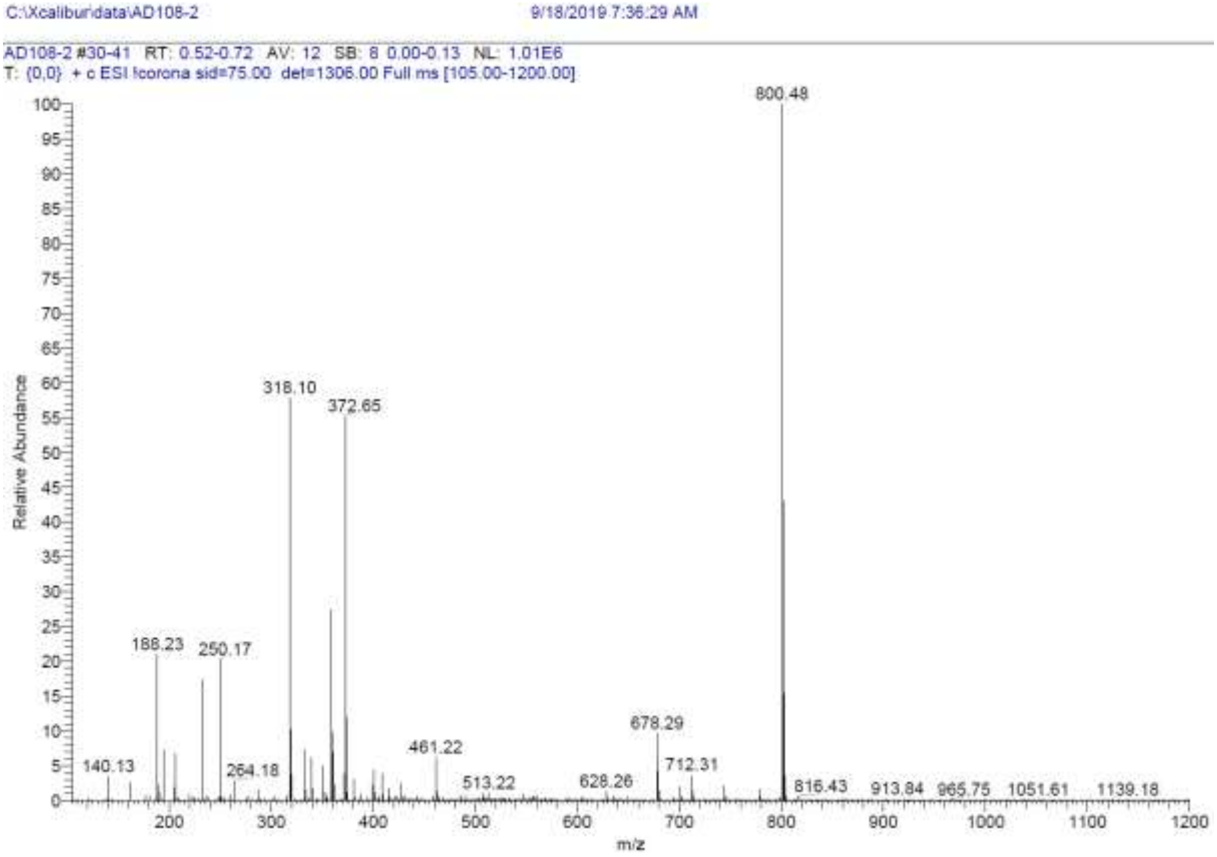

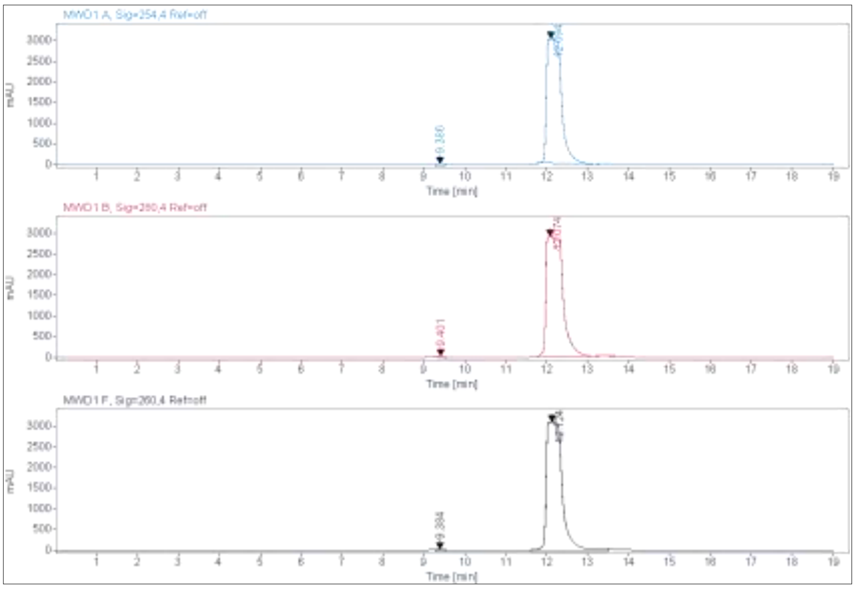

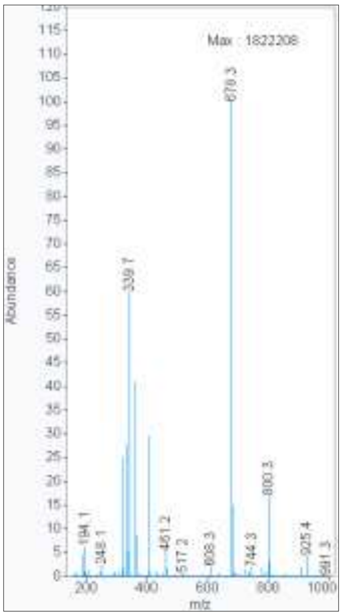

**Signal:** MWD1 A, Sig=254,4 Ref=off

| RT [min] | Type | Width [min] | Area | Height | Area% |
|---|---|---|---|---|---|
| 9.386 | MM | 0.2141 | 578.0328 | 44.9890 | 0.7538 |
| 12.094 | MM | 0.4188 | 76100.4609 | 3028.6030 | 99.2462 |
| | | Sum | 76678.4938 | | |

**Signal:** MWD1 B, Sig=280,4 Ref=off

| RT [min] | Type | Width [min] | Area | Height | Area% |
|---|---|---|---|---|---|
| 9.401 | MM | 0.2216 | 570.3035 | 42.8859 | 0.6772 |
| 12.074 | MM | 0.4768 | 83642.8047 | 2923.6580 | 99.3228 |
| | | Sum | 84213.1082 | | |

**Signal:** MWD1 F, Sig=260,4 Ref=off

| RT [min] | Type | Width [min] | Area | Height | Area% |
|---|---|---|---|---|---|
| 9.384 | MM | 0.2119 | 613.2241 | 48.2244 | 0.6948 |
| 12.124 | MM | 0.4631 | 87650.6875 | 3154.5259 | 99.3052 |
| | | Sum | 88263.9116 | | |

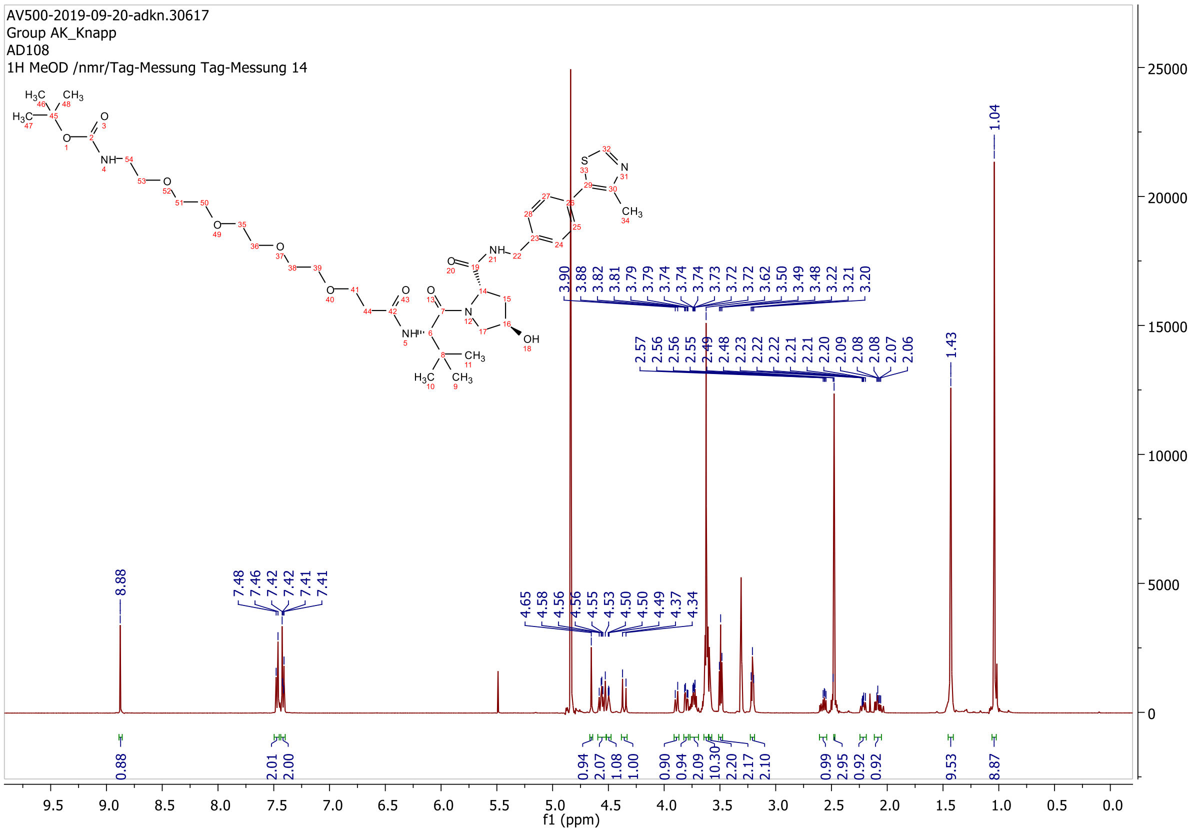

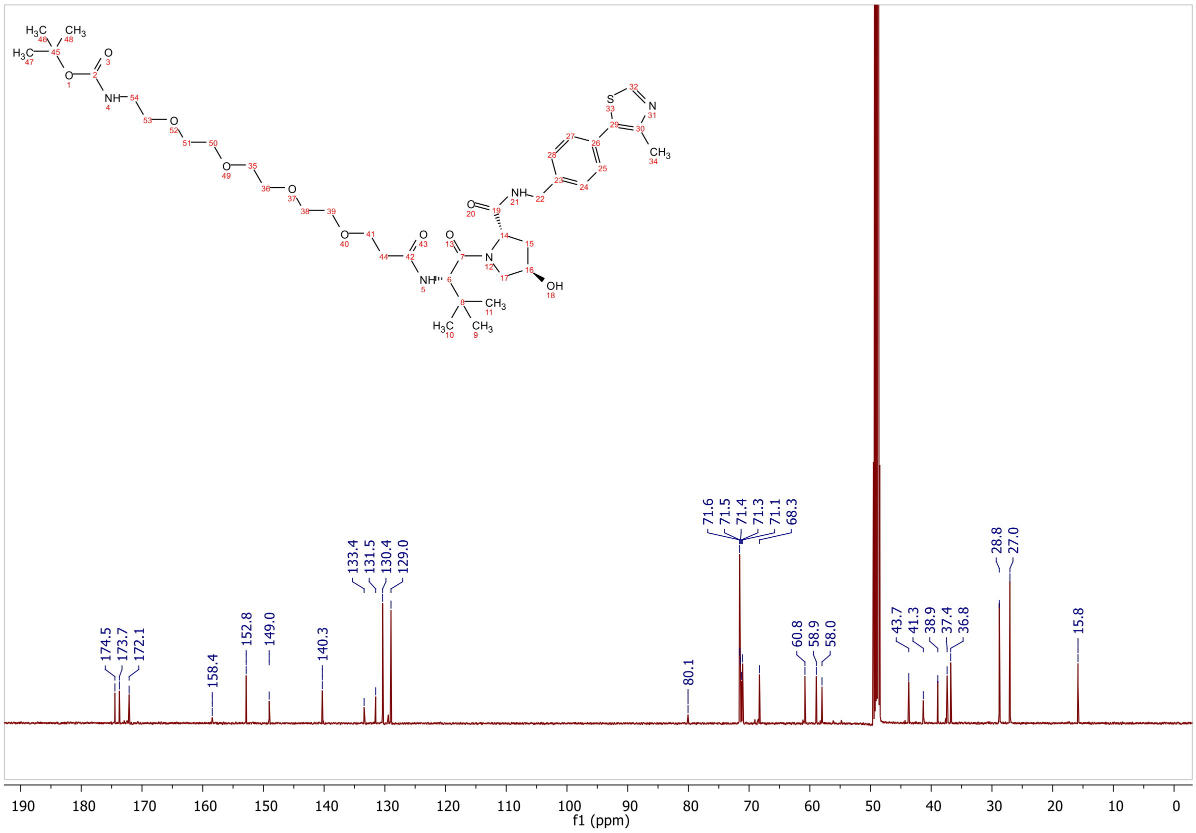

174.5
173.7
172.1
158.4
152.8
149.0
140.3
133.4
131.5
130.4
129.0
80.1
71.6
71.5
71.4
71.3
71.1
68.3
60.8
58.9
58.0
43.7
41.3
38.9
37.4
36.8
28.8
27.0
15.8
f1 (ppm)

**ESI, $^{1}$H-NMR and $^{13}$C-NMR of tert-butyl (3-(((S)-1-((2S,4R)-4-hydroxy-2-((4-(4-methylthiazol-5-yl)benzyl)carbamoyl)pyrrolidin-1-yl)-3,3-dimethyl-1-oxobutan-2-yl)amino)-3-oxopropyl)carbamate L9**

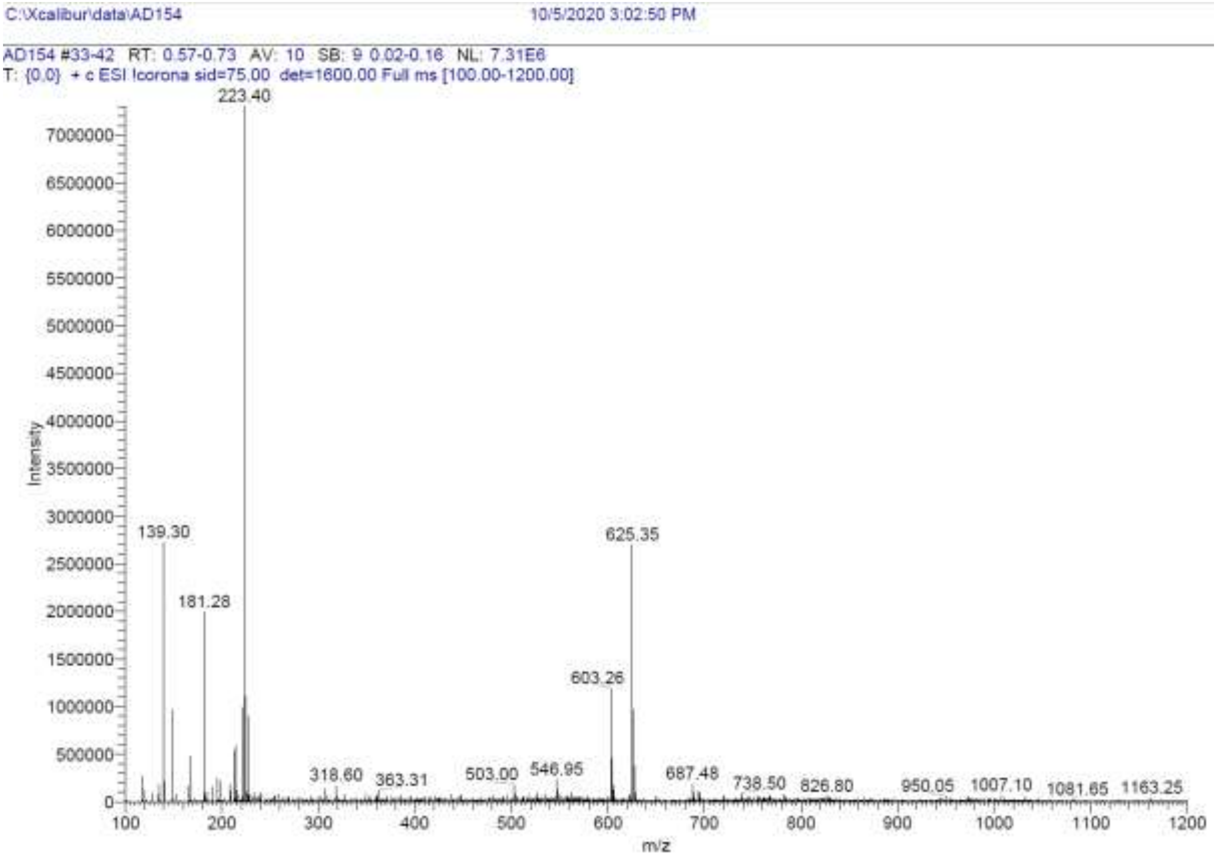

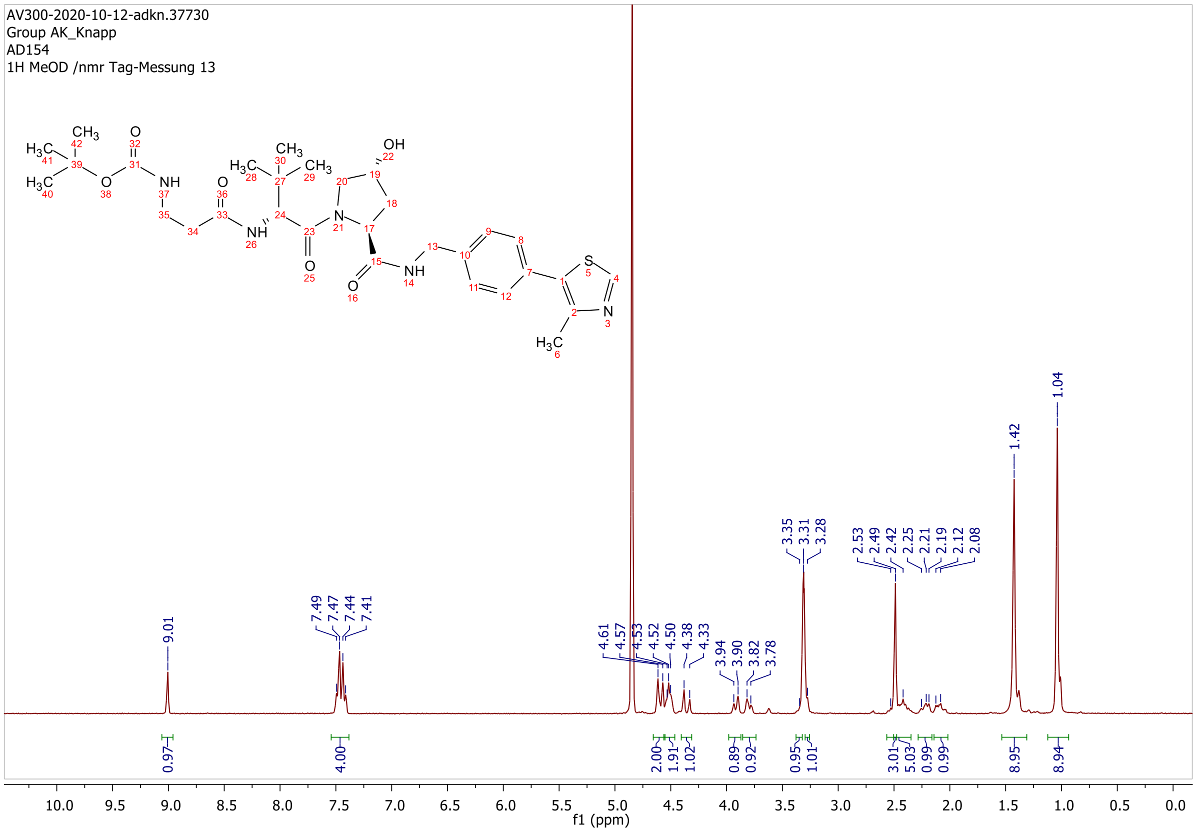

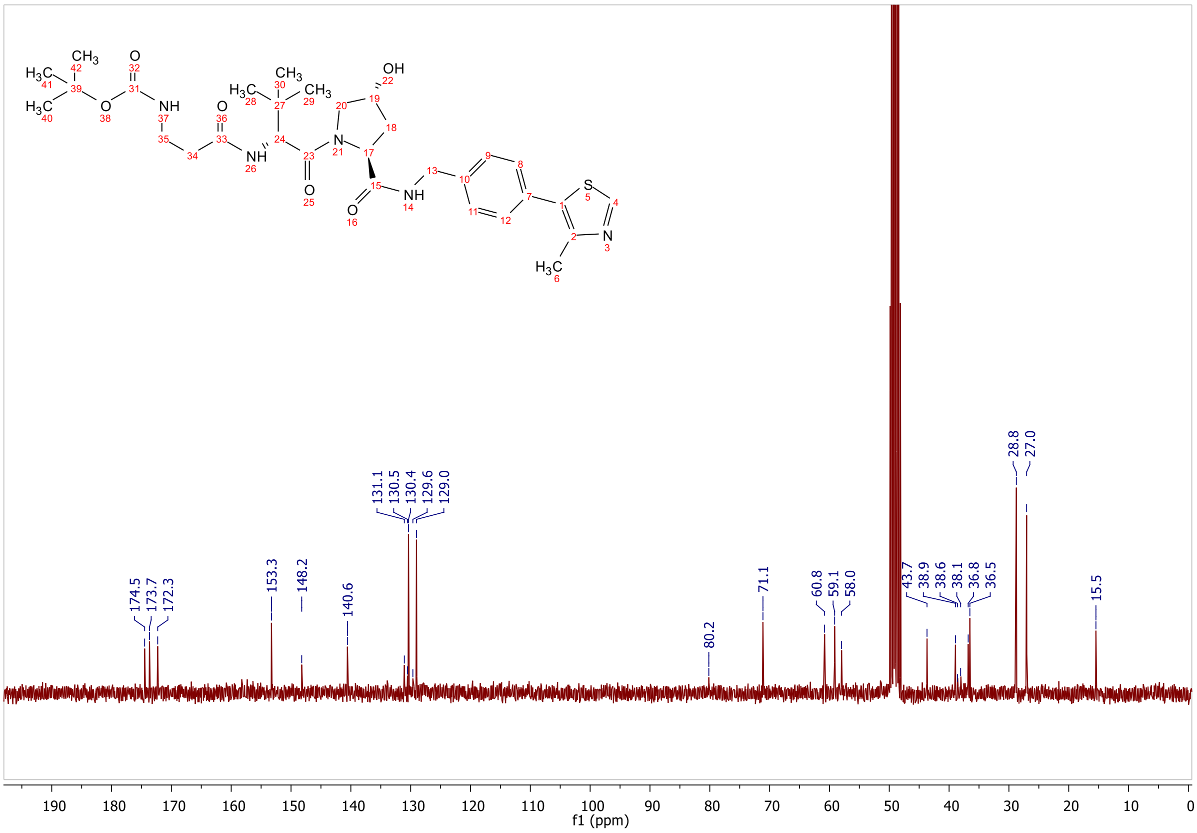
174.5
173.7
172.3
153.3
148.2
140.6
131.1
130.5
130.4
129.6
129.0
80.2
71.1
60.8
59.1
58.0
43.7
38.9
38.6
38.1
36.8
36.5
28.8
27.0
15.5
f1 (ppm)

**ESI, $^{1}$H-NMR and $^{13}$C-NMR of tert-butyl (4-(((S)-1-((2S,4R)-4-hydroxy-2-((4-(4-methylthiazol-5-yl)benzyl)carbamoyl)pyrrolidin-1-yl)-3,3-dimethyl-1-oxobutan-2-yl)amino)-4-oxobutyl)carbamate L10**

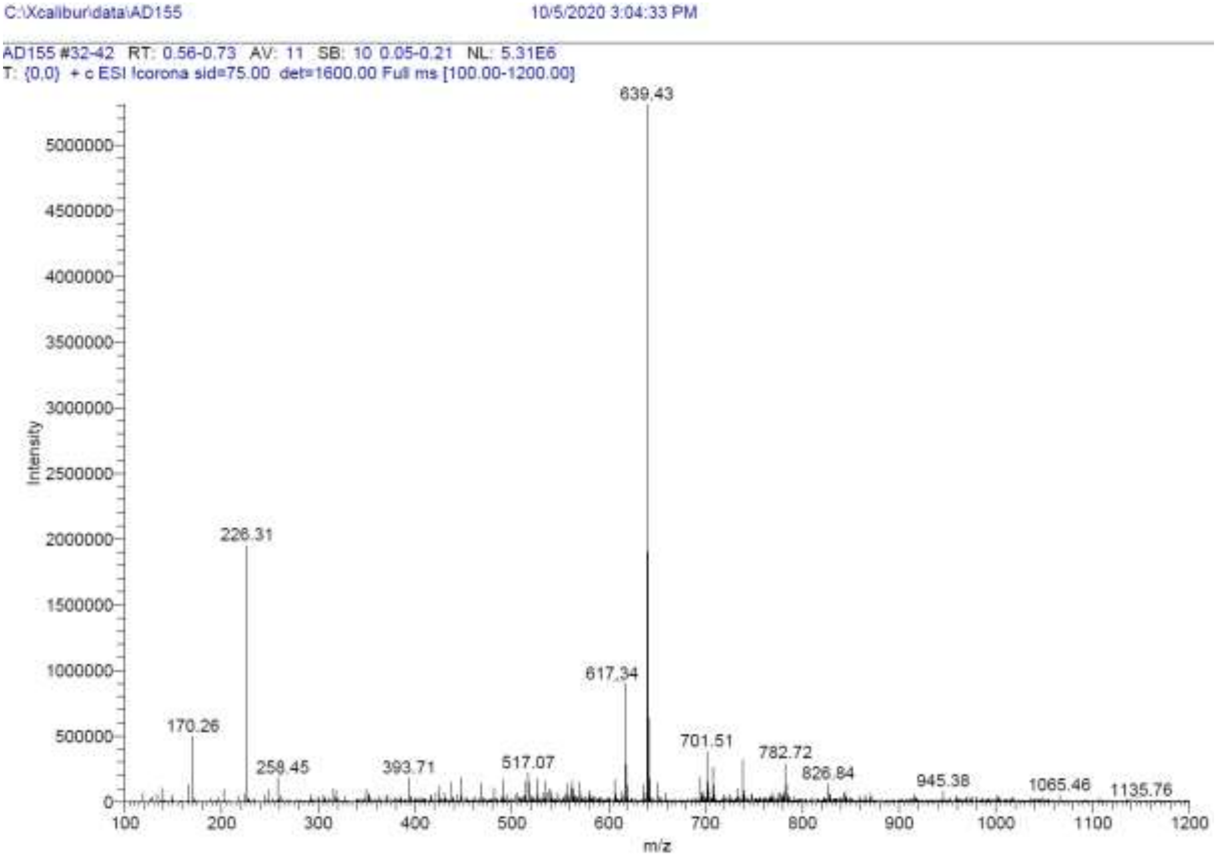

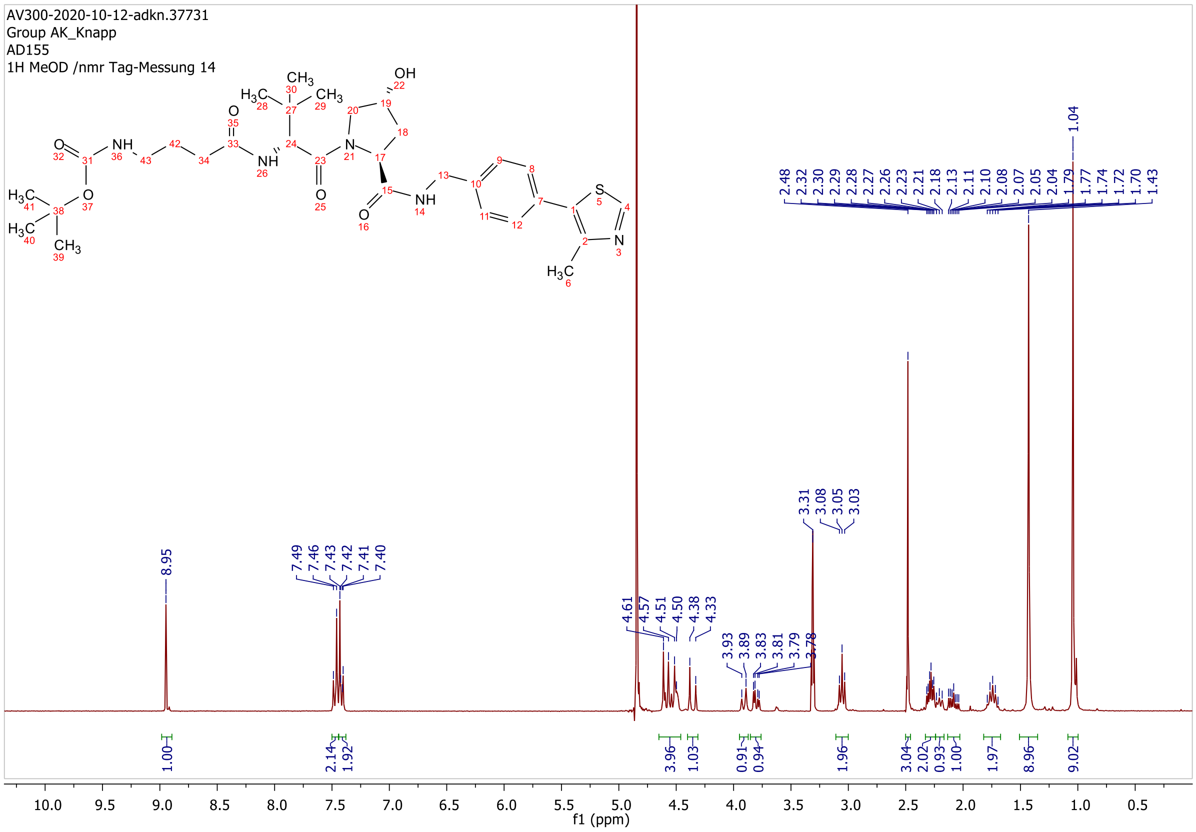

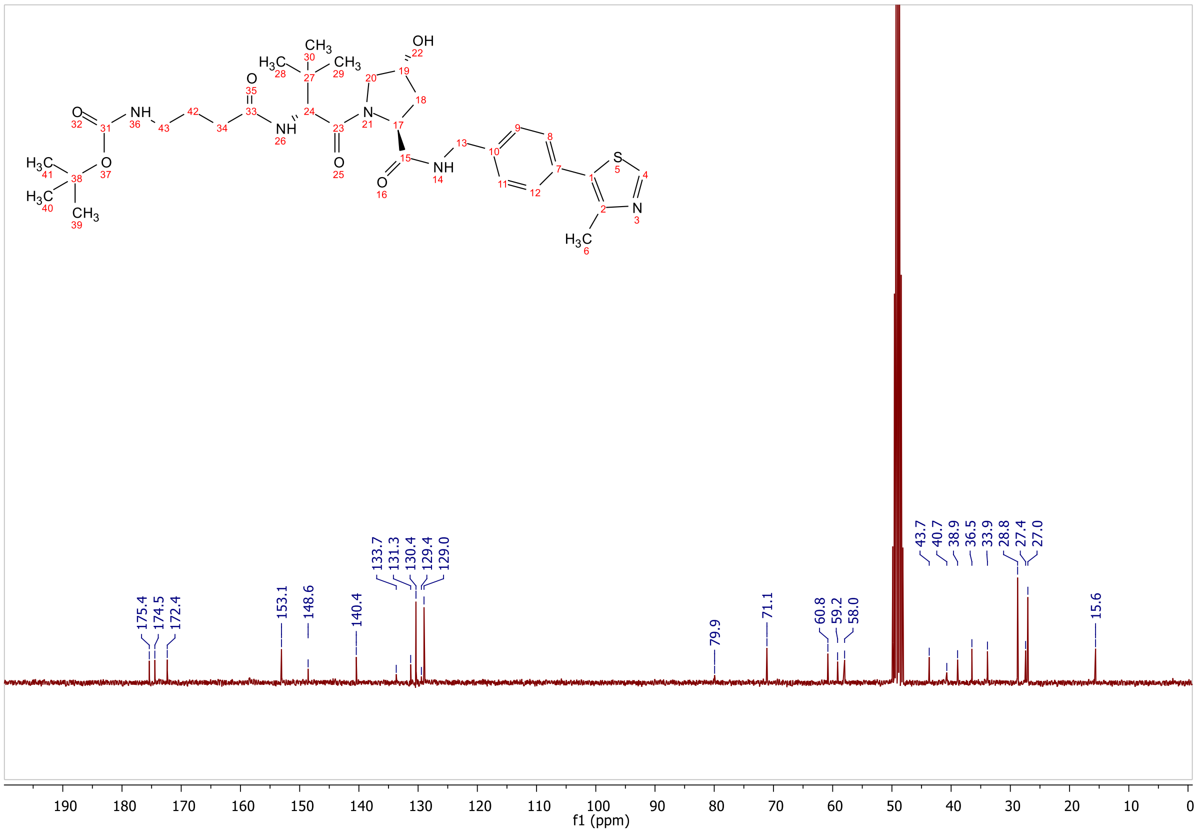

175.4
174.5
172.4
153.1
148.6
140.4
133.7
131.3
130.4
129.4
129.0
79.9
71.1
60.8
59.2
58.0
43.7
40.7
38.9
36.5
33.9
28.8
27.4
27.0
15.6
f1 (ppm)

**ESI, HPLC, ¹H-NMR and ¹³C-NMR of tert-butyl (5-(((S)-1-((2S,4R)-4-hydroxy-2-((4-(4-methylthiazol-5-yl)benzyl)carbamoyl)pyrrolidin-1-yl)-3,3-dimethyl-1-oxobutan-2-yl)amino)-5-oxopentyl)carbamate L11**

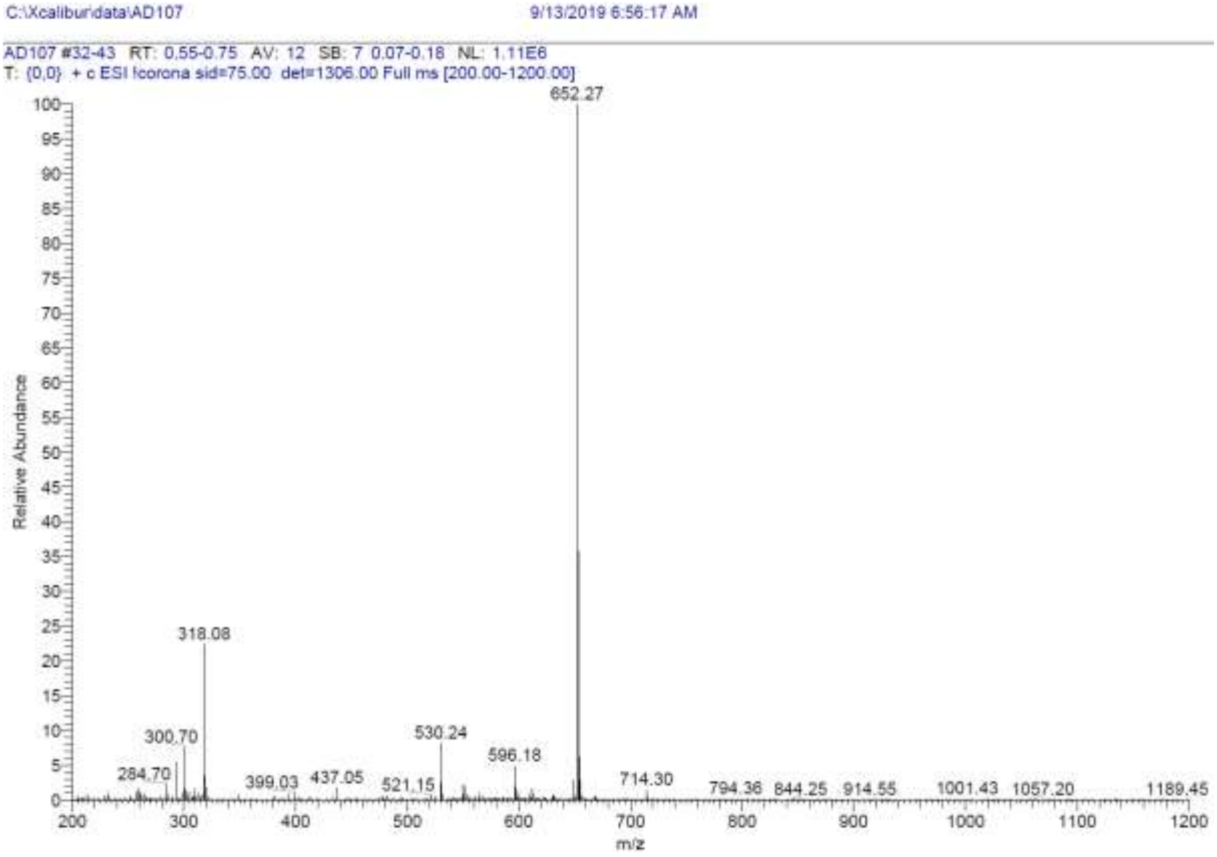

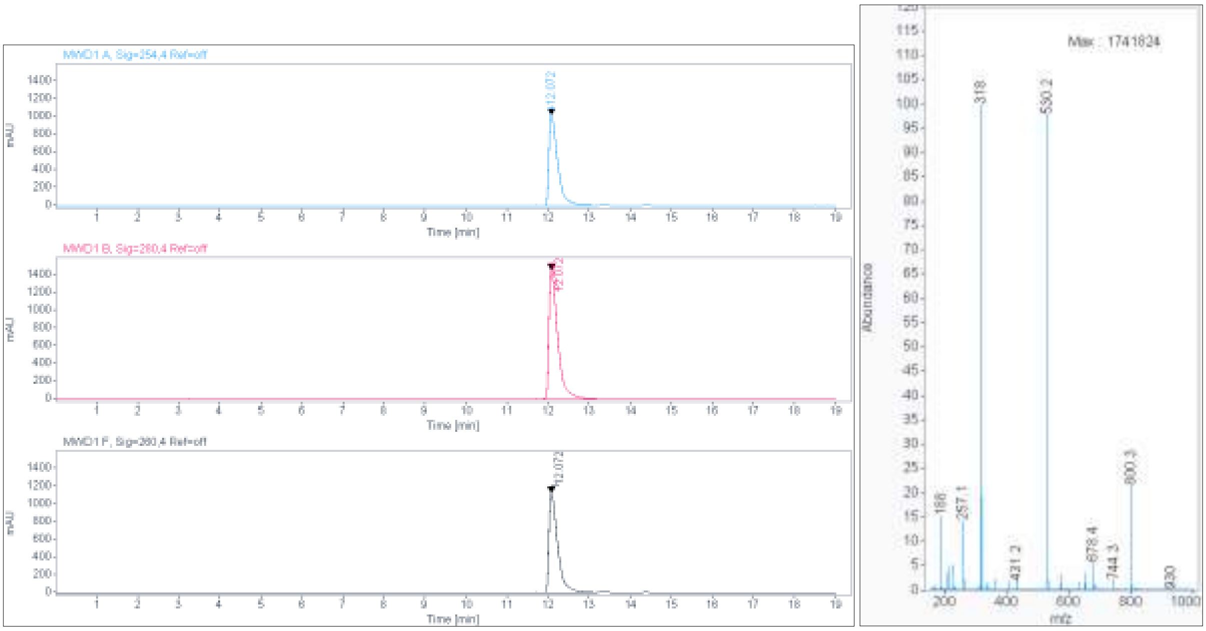

**Signal:** MWD1 A, Sig=254,4 Ref=off

| RT [min] | Type | Width [min] | Area | Height | Area% | Name |
|---|---|---|---|---|---|---|
| 12.072 | VV | 0.2259 | 15183.3105 | 1018.8142 | 100.0000 | |
| | | Sum | 15183.3105 | | | |

**Signal:** MWD1 B, Sig=280,4 Ref=off

| RT [min] | Type | Width [min] | Area | Height | Area% | Name |
|---|---|---|---|---|---|---|
| 12.072 | VV | 0.2262 | 21853.1191 | 1455.2832 | 100.0000 | |
| | | Sum | 21853.1191 | | | |

**Signal:** MWD1 F, Sig=260,4 Ref=off

| RT [min] | Type | Width [min] | Area | Height | Area% | Name |
|---|---|---|---|---|---|---|
| 12.072 | VV | 0.2257 | 16775.8438 | 1126.5007 | 100.0000 | |

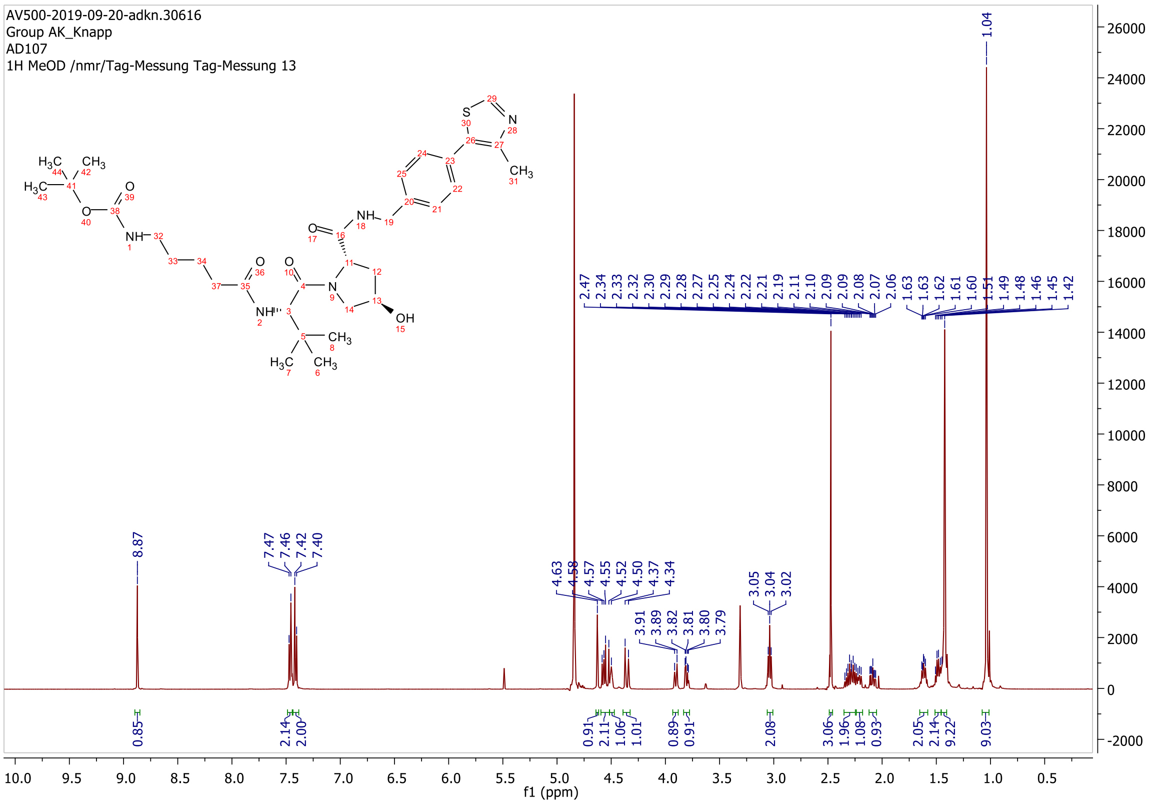

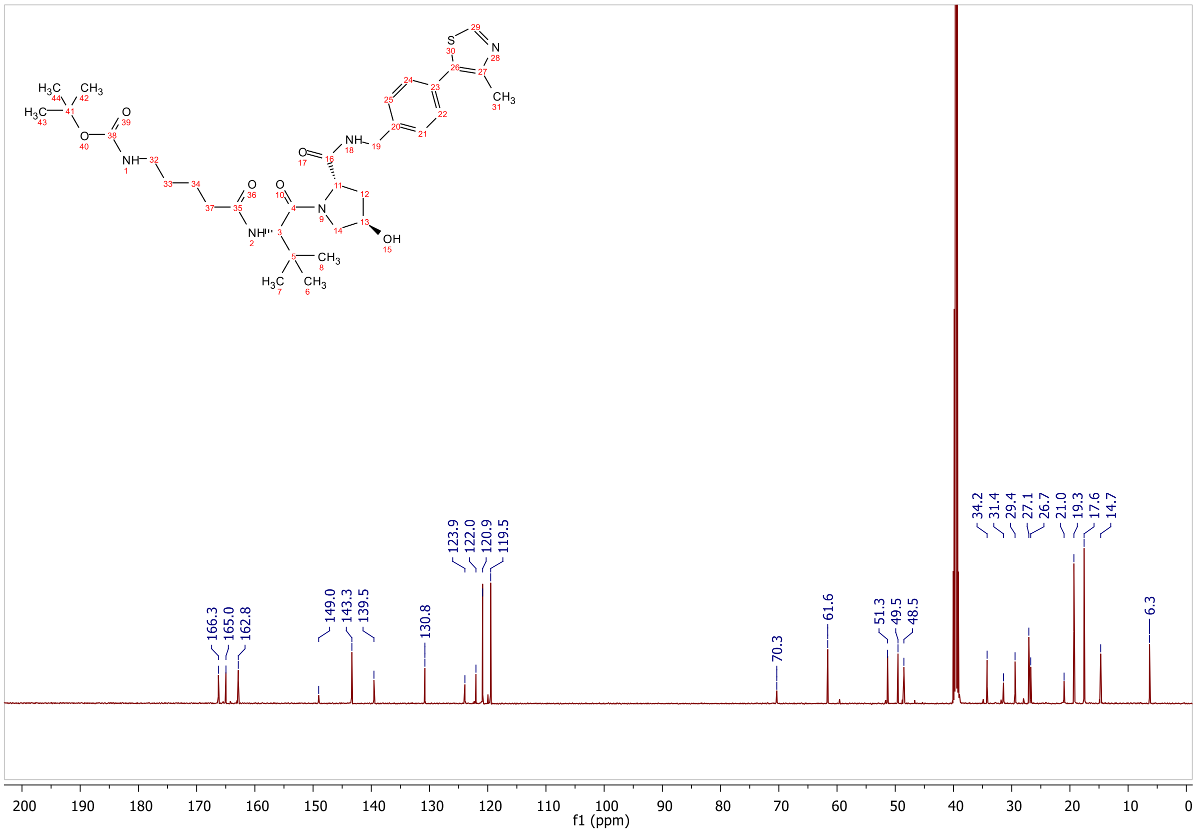
166.3
165.0
162.8
149.0
143.3
139.5
130.8
123.9
122.0
120.9
119.5
70.3
61.6
51.3
49.5
48.5
34.2
31.4
29.4
27.1
26.7
21.0
19.3
17.6
14.7
6.3
f1 (ppm)

**ESI, HPLC, $^{1}$H-NMR and $^{13}$C-NMR of tert-butyl (6-(((*S*)-1-((2*S*,4*R*)-4-hydroxy-2-((4-(4-methylthiazol-5-yl)benzyl)carbamoyl)pyrrolidin-1-yl)-3,3-dimethyl-1-oxobutan-2-yl)amino)-6-oxohexyl)carbamate L12**

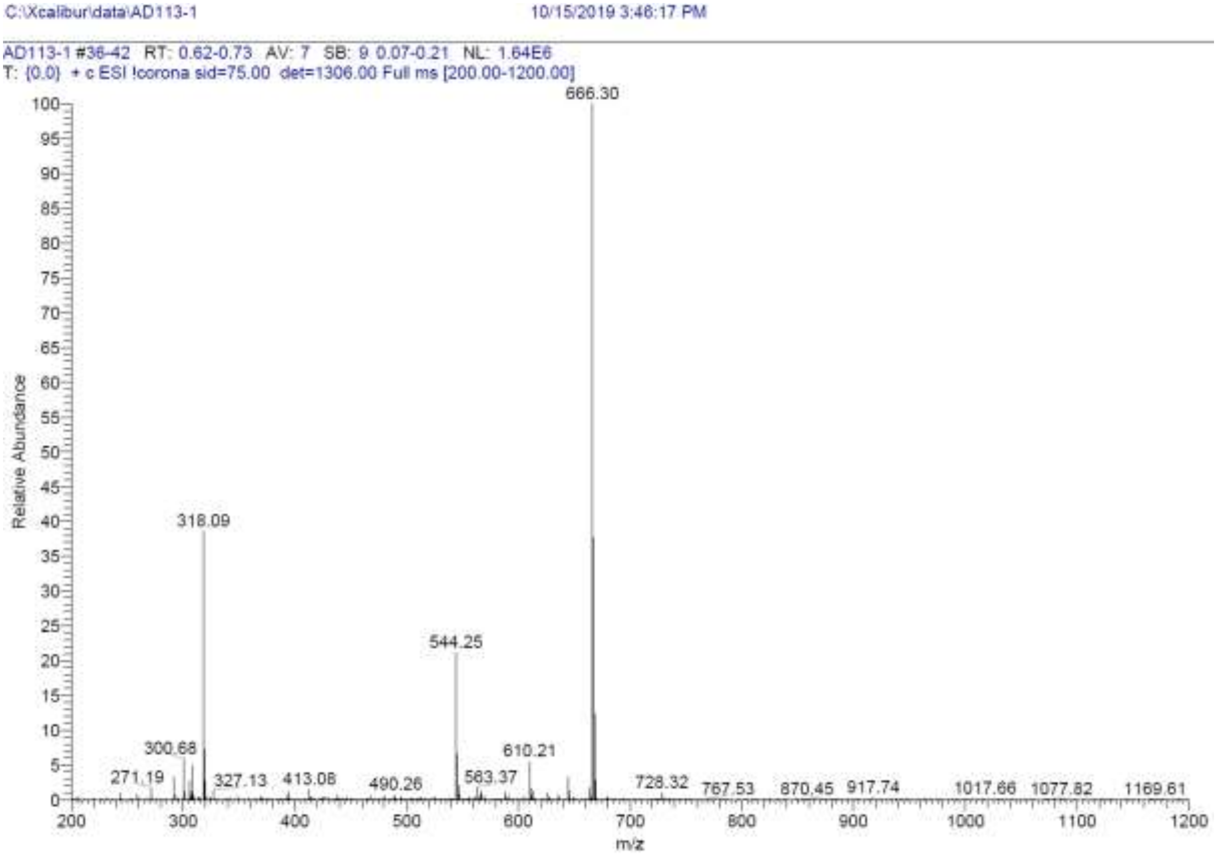

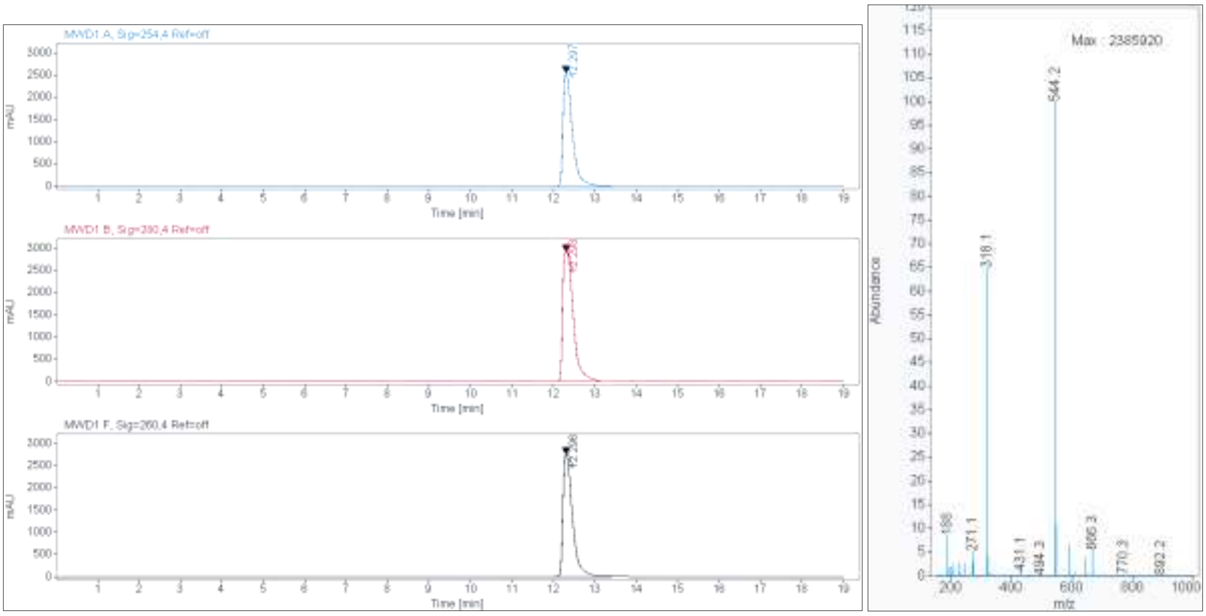

**Signal:** MWD1 A, Sig=254,4 Ref=off

| RT [min] | Type | Width [min] | Area | Height | Area% | Name |
|---|---|---|---|---|---|---|
| 12.297 | VV | 0.2391 | 40065.7813 | 2553.0674 | 100.0000 | |
| | | Sum | 40065.7813 | | | |

**Signal:** MWD1 B, Sig=280,4 Ref=off

| RT [min] | Type | Width [min] | Area | Height | Area% | Name |
|---|---|---|---|---|---|---|
| 12.298 | MM | 0.2941 | 51400.3750 | 2912.6912 | 100.0000 | |
| | | Sum | 51400.3750 | | | |

**Signal:** MWD1 F, Sig=260,4 Ref=off

| RT [min] | Type | Width [min] | Area | Height | Area% | Name |
|---|---|---|---|---|---|---|
| 12.296 | VV | 0.2173 | 44116.5742 | 2763.2627 | 100.0000 | |

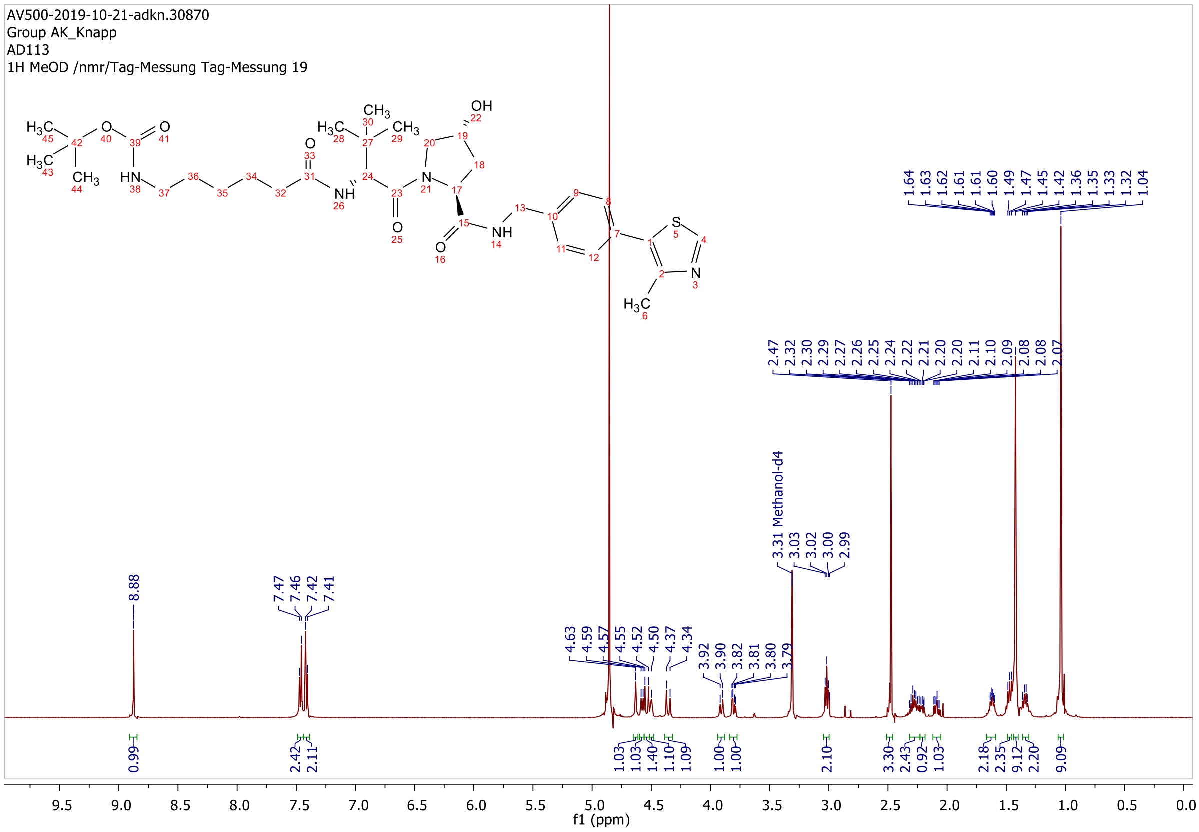

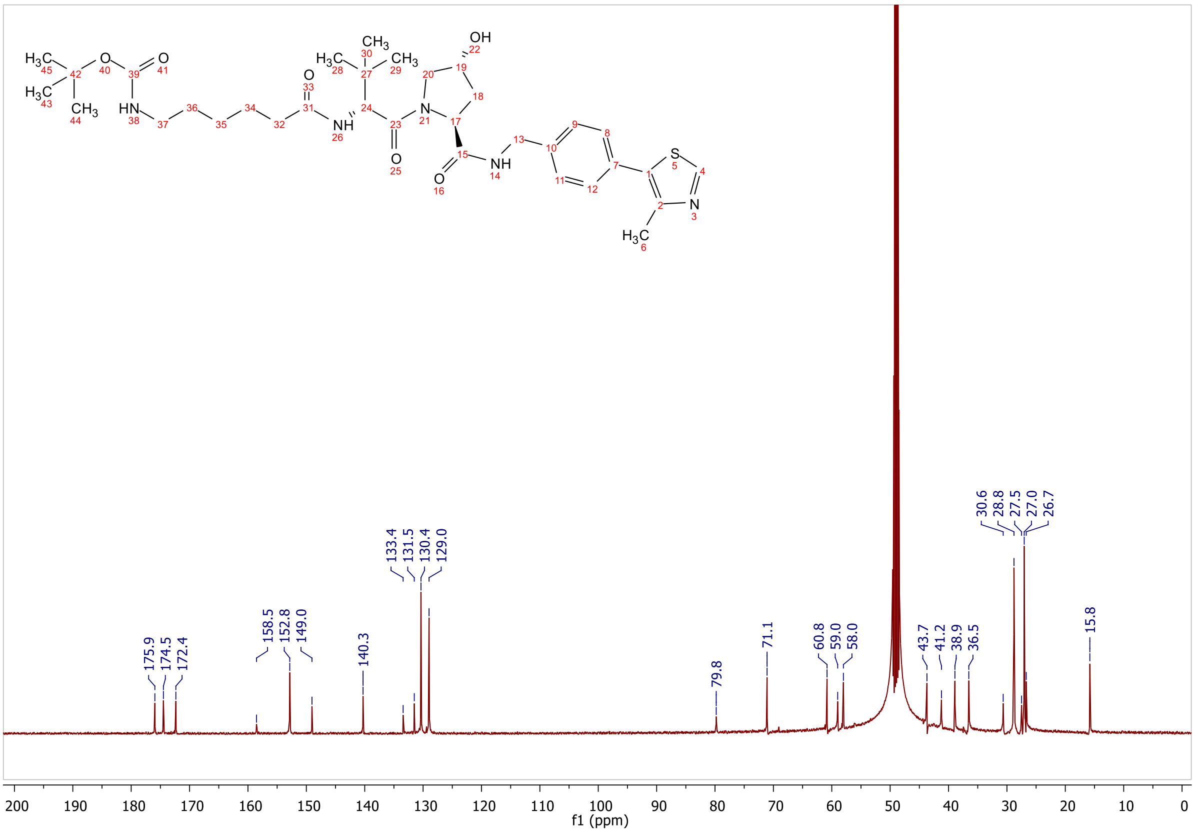
175.9
174.5
172.4
158.5
152.8
149.0
140.3
133.4
131.5
130.4
129.0
79.8
71.1
60.8
59.0
58.0
43.7
41.2
38.9
36.5
30.6
28.8
27.5
27.0
26.7
15.8
f1 (ppm)

**ESI, HPLC, $^{1}$H-NMR and $^{13}$C-NMR of tert-butyl (7-(((*S*)-1-((2*S*,4*R*)-4-hydroxy-2-((4-(4-methylthiazol-5-yl)benzyl)carbamoyl)pyrrolidin-1-yl)-3,3-dimethyl-1-oxobutan-2-yl)amino)-7-oxoheptyl)carbamate L13**

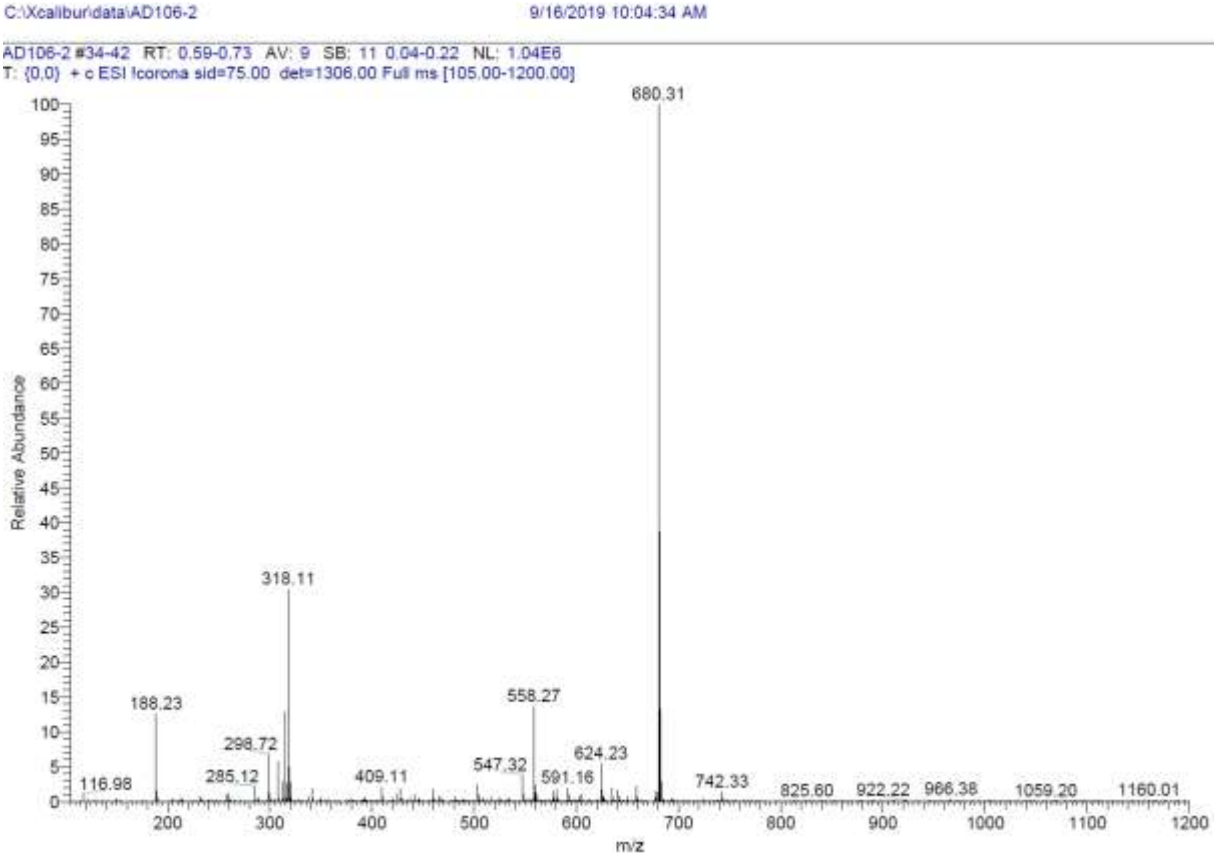

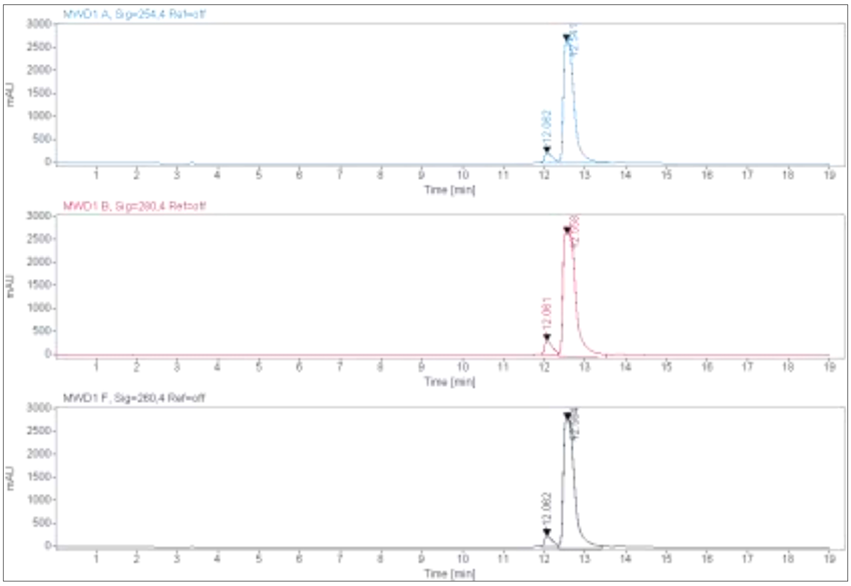

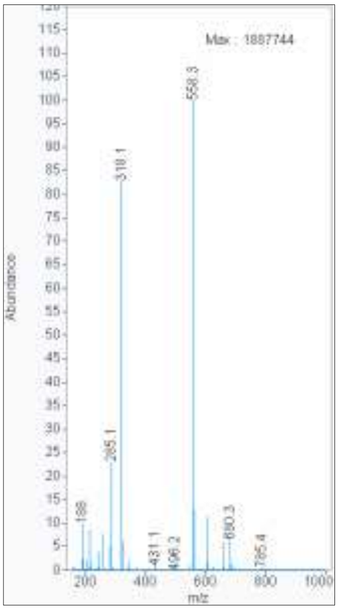

**Signal:** MWD1 A, Sig=254,4 Ref=off

| RT [min] | Type | Width [min] | Area | Height | Area% | Name |
|---|---|---|---|---|---|---|
| 12.062 | VV | 0.2181 | 3252.9497 | 223.1664 | 6.2811 | |
| 12.541 | VV | 0.2368 | 48536.3945 | 2660.7231 | 93.7189 | |
| | | Sum | 51789.3442 | | | |

**Signal:** MWD1 B, Sig=280,4 Ref=off

| RT [min] | Type | Width [min] | Area | Height | Area% | Name |
|---|---|---|---|---|---|---|
| 12.061 | MM | 0.2245 | 4209.9492 | 312.5555 | 6.5658 | |
| 12.558 | MM | 0.3742 | 59909.6172 | 2668.2478 | 93.4342 | |
| | | Sum | 64119.5664 | | | |

**Signal:** MWD1 F, Sig=260,4 Ref=off

| RT [min] | Type | Width [min] | Area | Height | Area% | Name |
|---|---|---|---|---|---|---|
| 12.062 | VV | 0.2166 | 3560.8386 | 246.4372 | 6.1048 | |
| 12.564 | MM | 0.3241 | 54767.2188 | 2816.3435 | 93.8952 | |

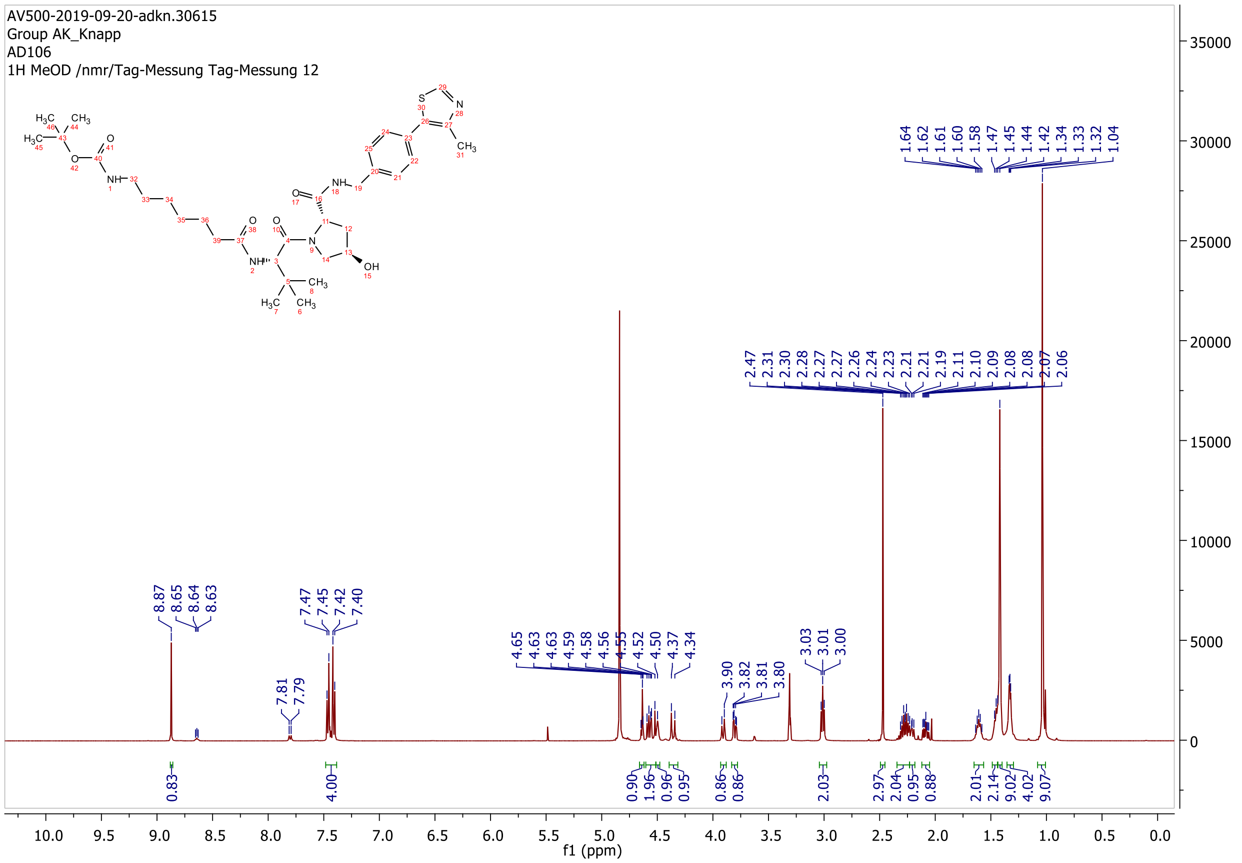

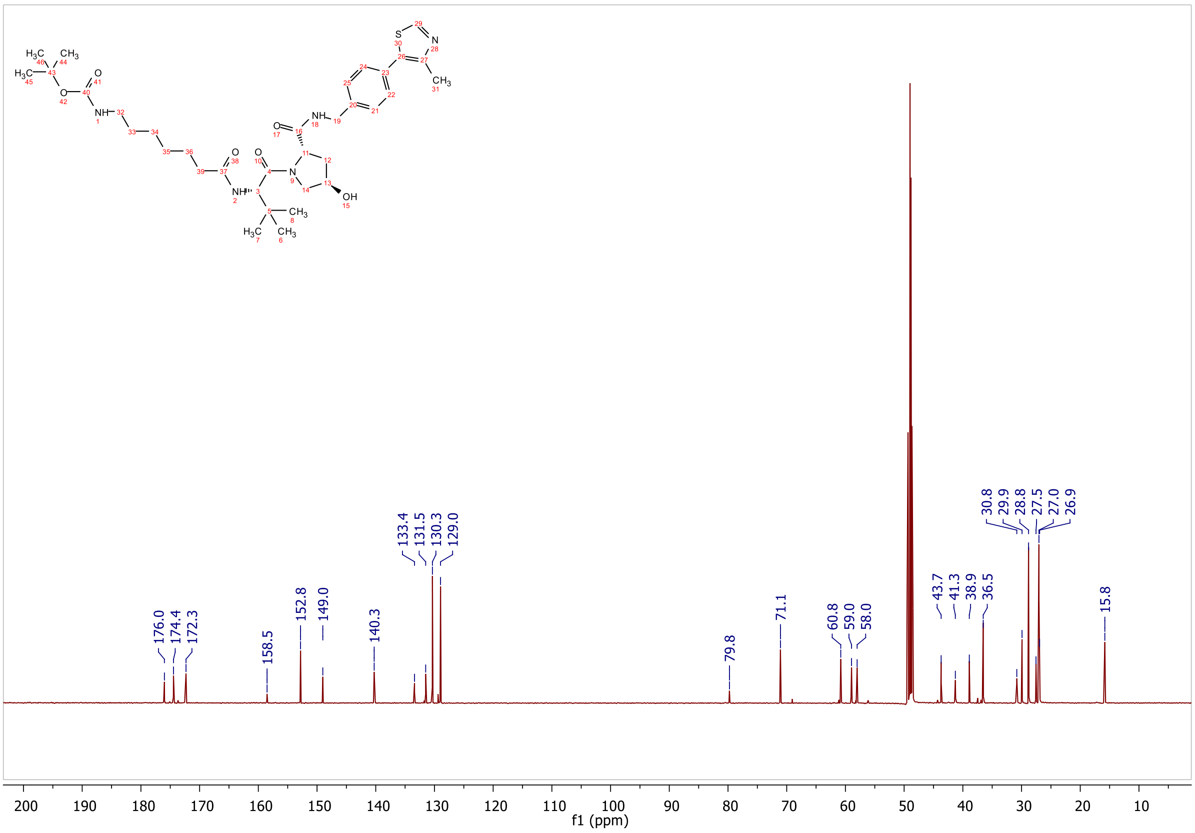

176.0
174.4
172.3
158.5
152.8
149.0
140.3
133.4
131.5
130.3
129.0
79.8
71.1
60.8
59.0
58.0
43.7
41.3
38.9
36.5
30.8
29.9
28.8
27.5
27.0
26.9
15.8
f1 (ppm)

**ESI, HPLC, $^{1}$H-NMR and $^{13}$C-NMR of tert-butyl (4-(2-(((S)-1-((2S,4R)-4-hydroxy-2-((4-(4-methylthiazol-5-yl)benzyl)carbamoyl)pyrrolidin-1-yl)-3,3-dimethyl-1-oxobutan-2-yl)amino)-2-oxoethyl)benzyl)carbamate L14**

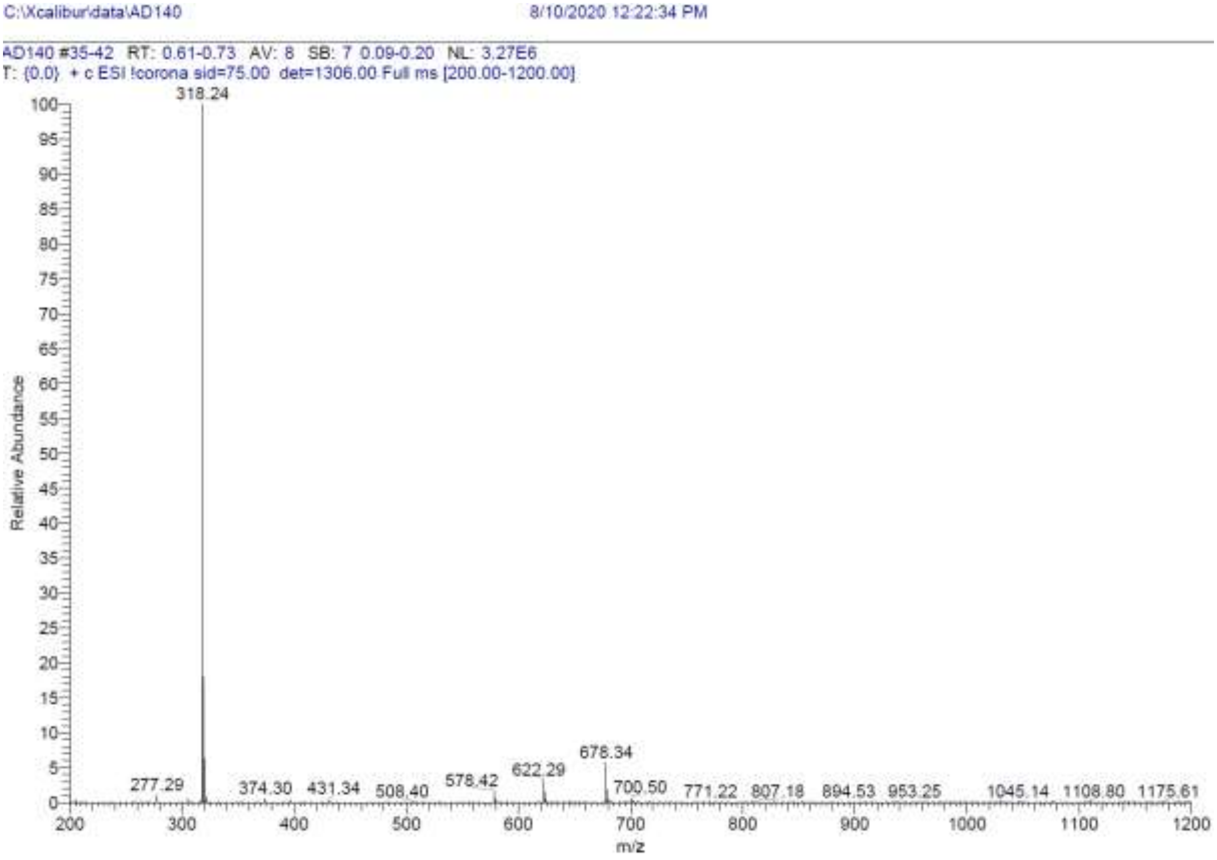

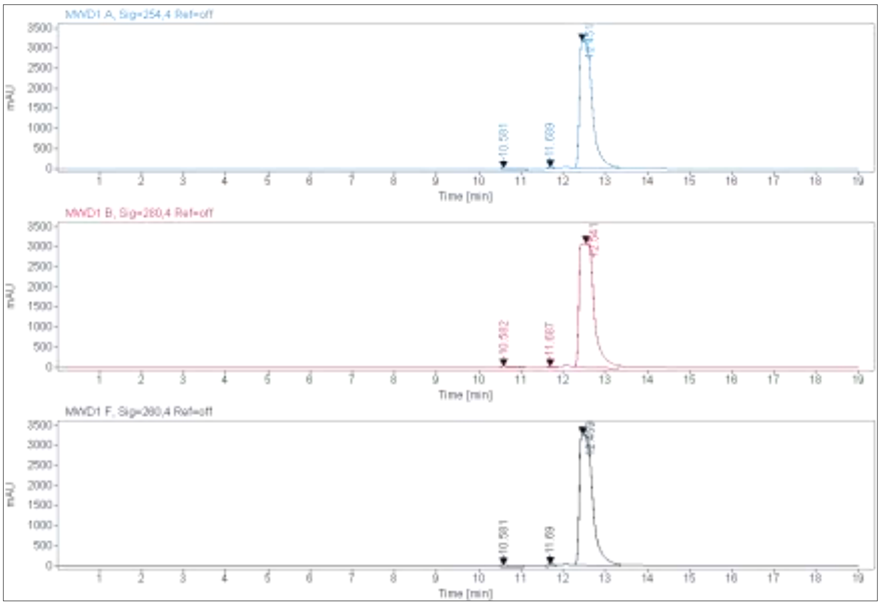

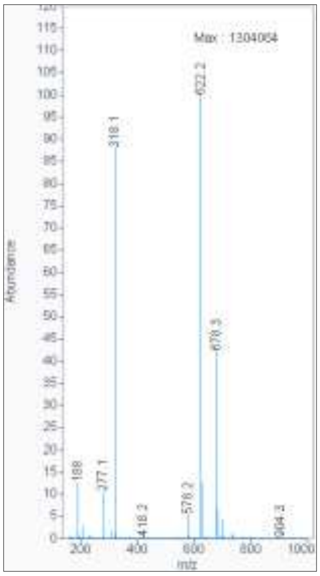

**Signal:** MWD1 A, Sig=254,4 Ref=off

| RT [min] | Type | Width [min] | Area | Height | Area% |
|---|---|---|---|---|---|
| 10.581 | MM | 0.4447 | 1334.5194 | 50.0107 | 1.9565 |
| 11.689 | MM | 0.1568 | 414.5808 | 44.0782 | 0.6078 |
| 12.451 | MM | 0.3501 | 66460.5781 | 3164.2292 | 97.4357 |
| | | Sum | 68209.6783 | | |

**Signal:** MWD1 B, Sig=280,4 Ref=off

| RT [min] | Type | Width [min] | Area | Height | Area% |
|---|---|---|---|---|---|
| 10.582 | MM | 0.3997 | 1567.3302 | 65.3503 | 2.0085 |
| 11.687 | MM | 0.2019 | 685.3602 | 56.5747 | 0.8783 |
| 12.541 | MM | 0.4061 | 75781.6641 | 3110.1145 | 97.1132 |
| | | Sum | 78034.3544 | | |

**Signal:** MWD1 F, Sig=260,4 Ref=off

| RT [min] | Type | Width [min] | Area | Height | Area% |
|---|---|---|---|---|---|
| 10.581 | MM | 0.4067 | 1300.0765 | 53.2741 | 1.7700 |
| 11.690 | MM | 0.1873 | 729.8845 | 64.9543 | 0.9937 |
| 12.459 | MM | 0.3645 | 71421.4219 | 3265.3564 | 97.2363 |

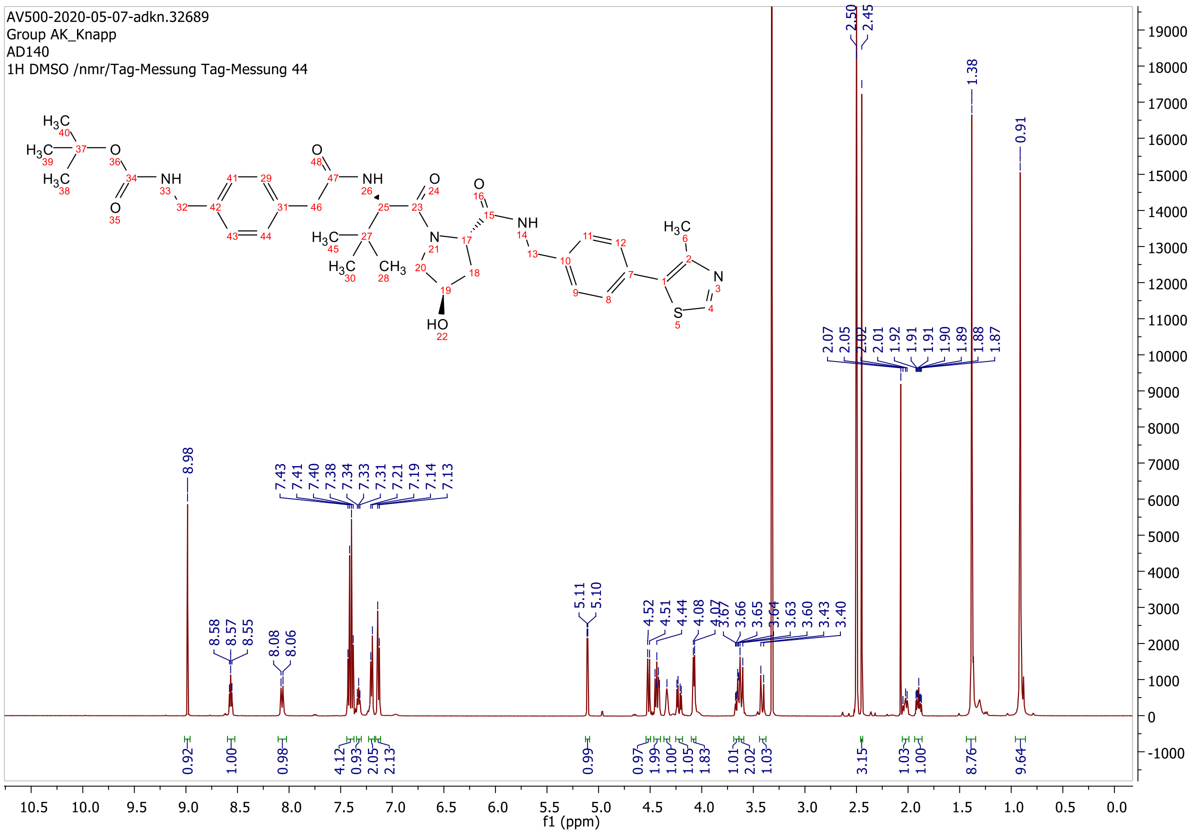

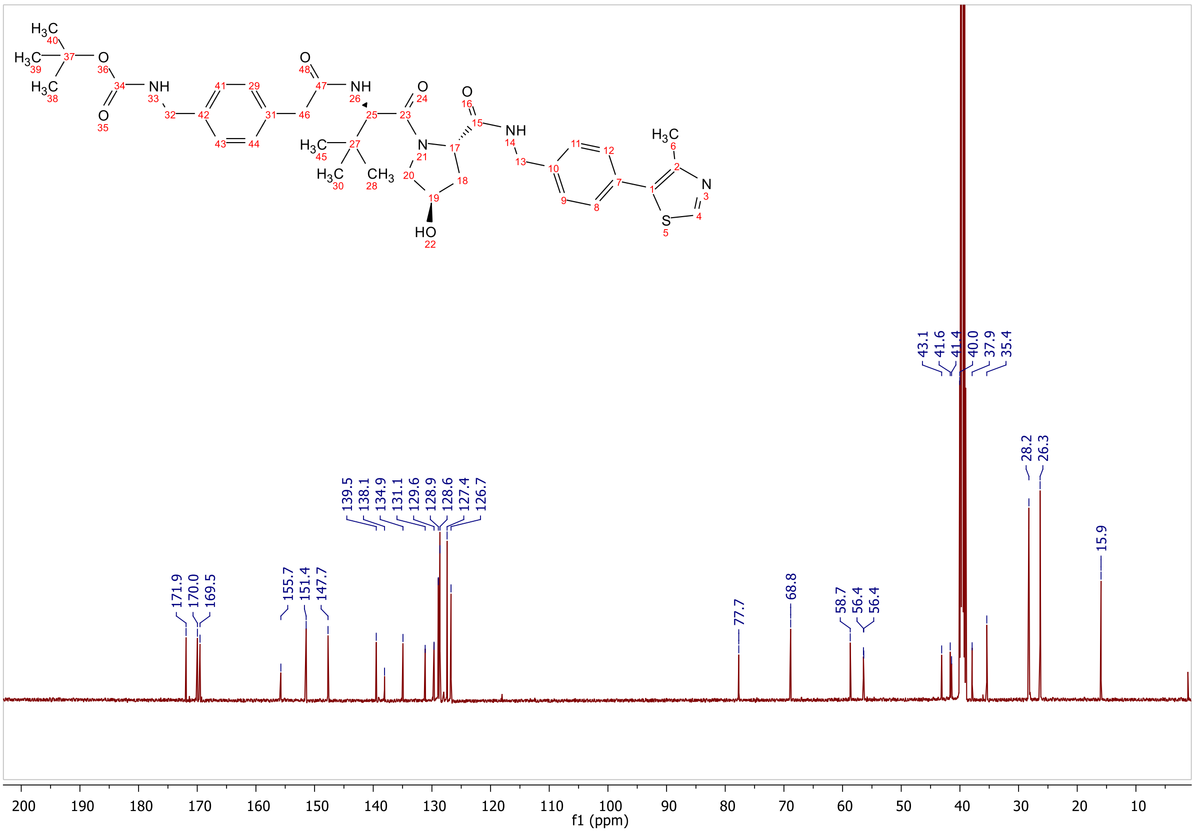
171.9
170.0
169.5
155.7
151.4
147.7
139.5
138.1
134.9
131.1
129.6
128.9
128.6
127.4
126.7
77.7
68.8
58.7
56.4
56.4
43.1
41.6
41.4
40.0
37.9
35.4
28.2
26.3
15.9
f1 (ppm)

**ESI, HPLC, $^{1}$H-NMR and $^{13}$C-NMR of tert-butyl (5-(((S)-1-((2S,4S)-4-hydroxy-2-((4-(4-methylthiazol-5-yl)benzyl)carbamoyl)pyrrolidin-1-yl)-3,3-dimethyl-1-oxobutan-2-yl)amino)-5-oxopentyl)carbamate L15**

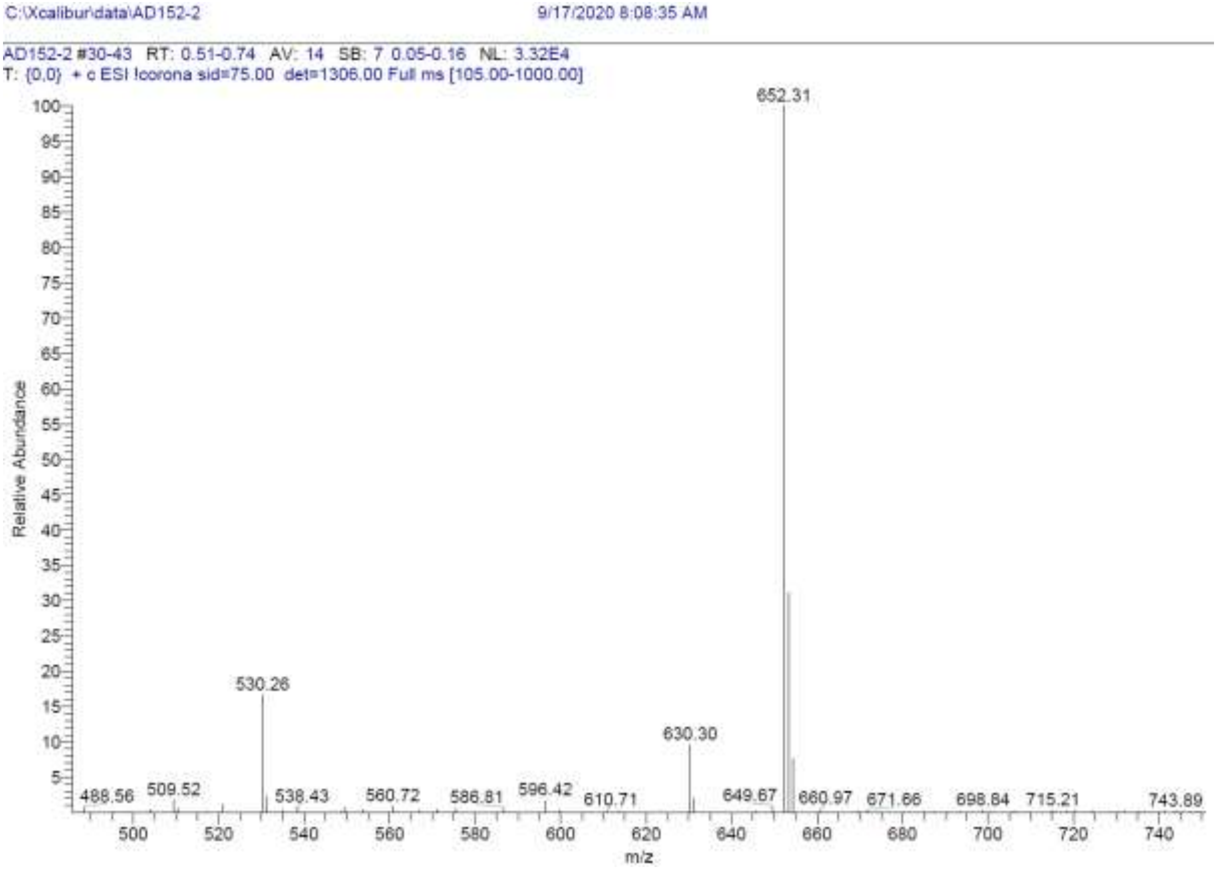

| RT [min] | Type | Width [min] | Area | Height | Area% | Name |
|---|---|---|---|---|---|---|
| 12.025 | VV | 0.2315 | 1145.2445 | 72.0177 | 100.0000 | |
| | | Sum | 1145.2445 | | | |

**Signal:** MWD1 B, Sig=280,4 Ref=off

| RT [min] | Type | Width [min] | Area | Height | Area% | Name |
|---|---|---|---|---|---|---|
| 12.025 | VV | 0.2326 | 1672.4189 | 104.5707 | 100.0000 | |
| | | Sum | 1672.4189 | | | |

**Signal:** MWD1 F, Sig=260,4 Ref=off

| RT [min] | Type | Width [min] | Area | Height | Area% | Name |
|---|---|---|---|---|---|---|
| 12.025 | VV | 0.2292 | 1240.3011 | 78.9803 | 100.0000 | |

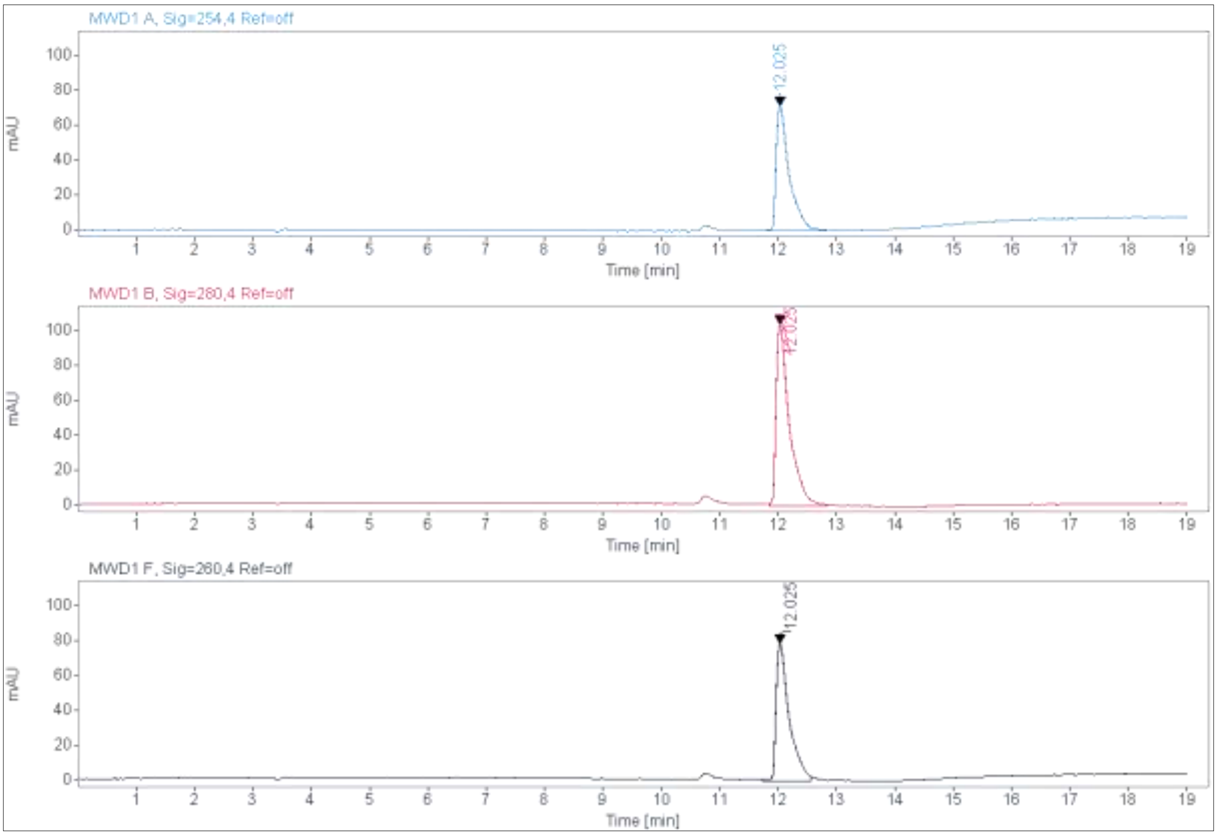
MWD1 A, Sig=254,4 Ref=off
MWD1 B, Sig=280,4 Ref=off
MWD1 F, Sig=260,4 Ref=off
12.025
mAU
Time [min]

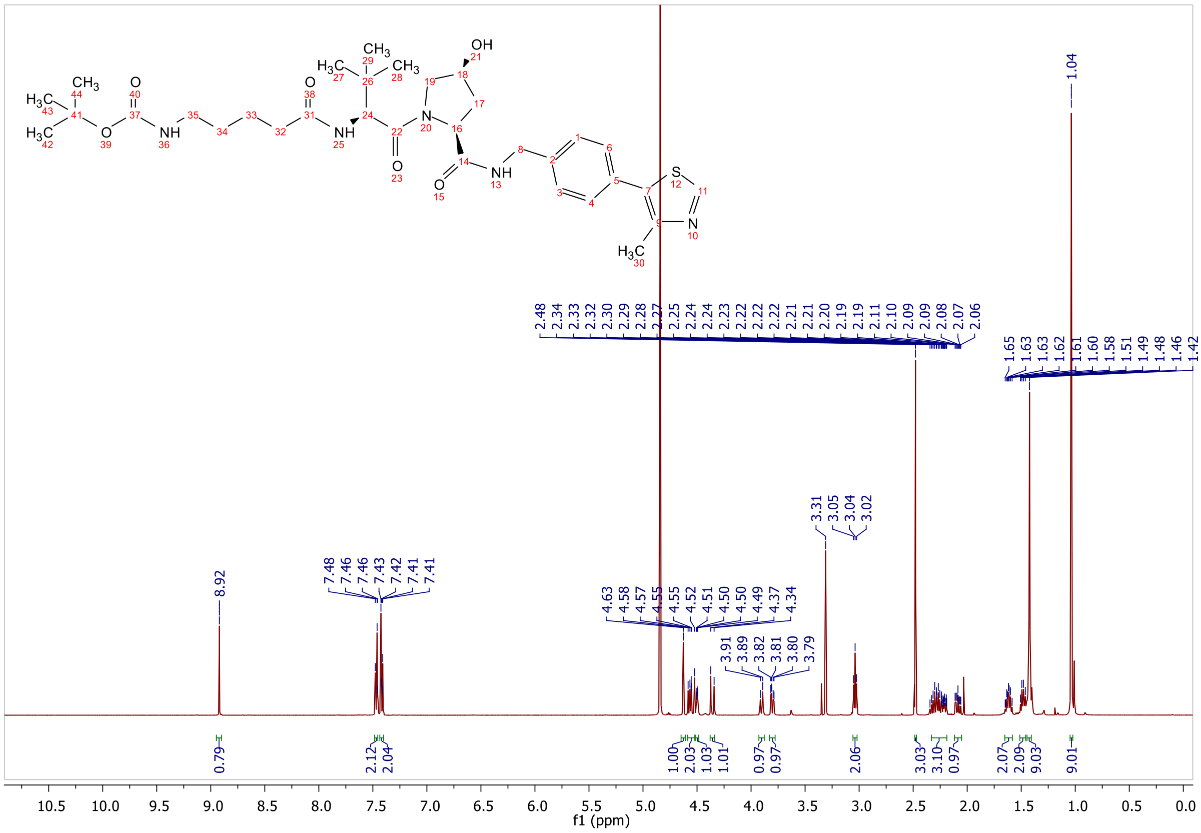
f1 (ppm)

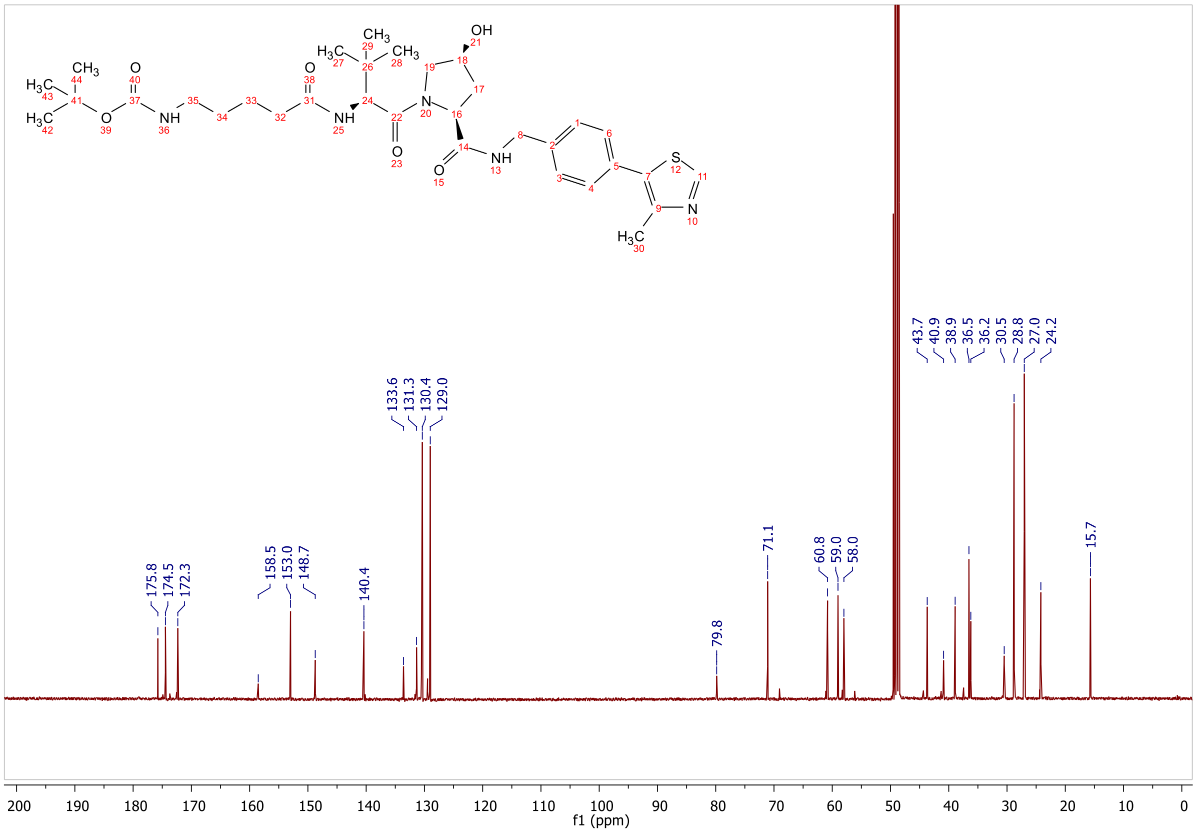
175.8
174.5
172.3
158.5
153.0
148.7
140.4
133.6
131.3
130.4
129.0
79.8
71.1
60.8
59.0
58.0
43.7
40.9
38.9
36.5
36.2
30.5
28.8
27.0
24.2
15.7
f1 (ppm)

#### 1.2 Appendix for intermediates 6a-c, S18a,b and BRET Tracer molecules 19a-c

**ESI, HPLC, $^{1}$H-NMR and $^{13}$C-NMR of tert-butyl (2-(2-(3'-(6-hydroxy-4-(trifluoromethyl)nicotinamido)-4'-(4-methylpiperazin-1-yl)-[1,1'-biphenyl]-4-carboxamido)ethoxy)ethyl)carbamate 6a**

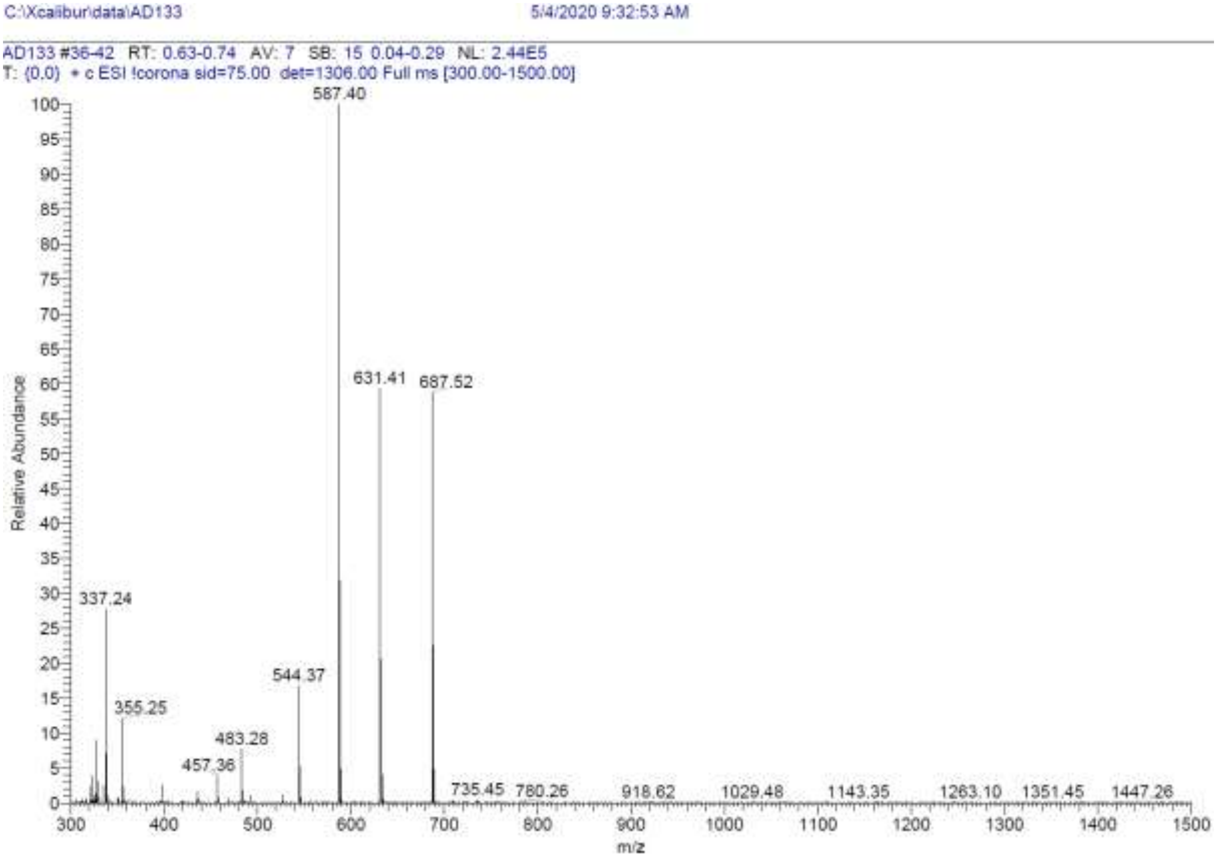

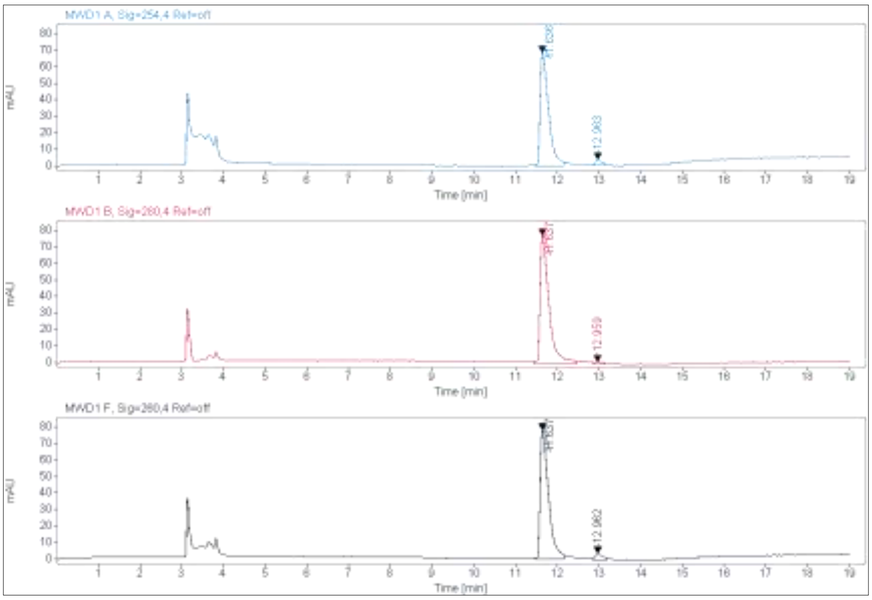

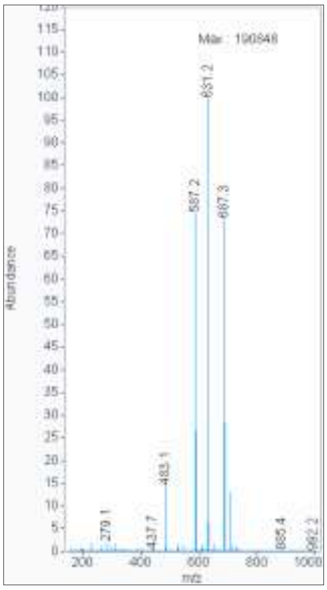

**Signal:** MWD1 A, Sig=254,4 Ref=off

| RT [min] | Type | Width [min] | Area | Height | Area% |
|---|---|---|---|---|---|
| 11.636 | VV | 0.2088 | 960.0562 | 68.8025 | 96.2284 |
| 12.963 | MM | 0.1685 | 37.6289 | 3.7220 | 3.7716 |
| | | Sum | 997.6851 | | |

**Signal:** MWD1 B, Sig=280,4 Ref=off

| RT [min] | Type | Width [min] | Area | Height | Area% |
|---|---|---|---|---|---|
| 11.637 | VV | 0.2215 | 1175.1613 | 78.1245 | 98.9523 |
| 12.959 | MM | 0.2024 | 12.4427 | 1.0247 | 1.0477 |
| | | Sum | 1187.6039 | | |

**Signal:** MWD1 F, Sig=260,4 Ref=off

| RT [min] | Type | Width [min] | Area | Height | Area% |
|---|---|---|---|---|---|
| 11.637 | VV | 0.2084 | 1095.8492 | 78.2365 | 95.5942 |
| 12.962 | MM | 0.2064 | 50.5057 | 4.0791 | 4.4058 |

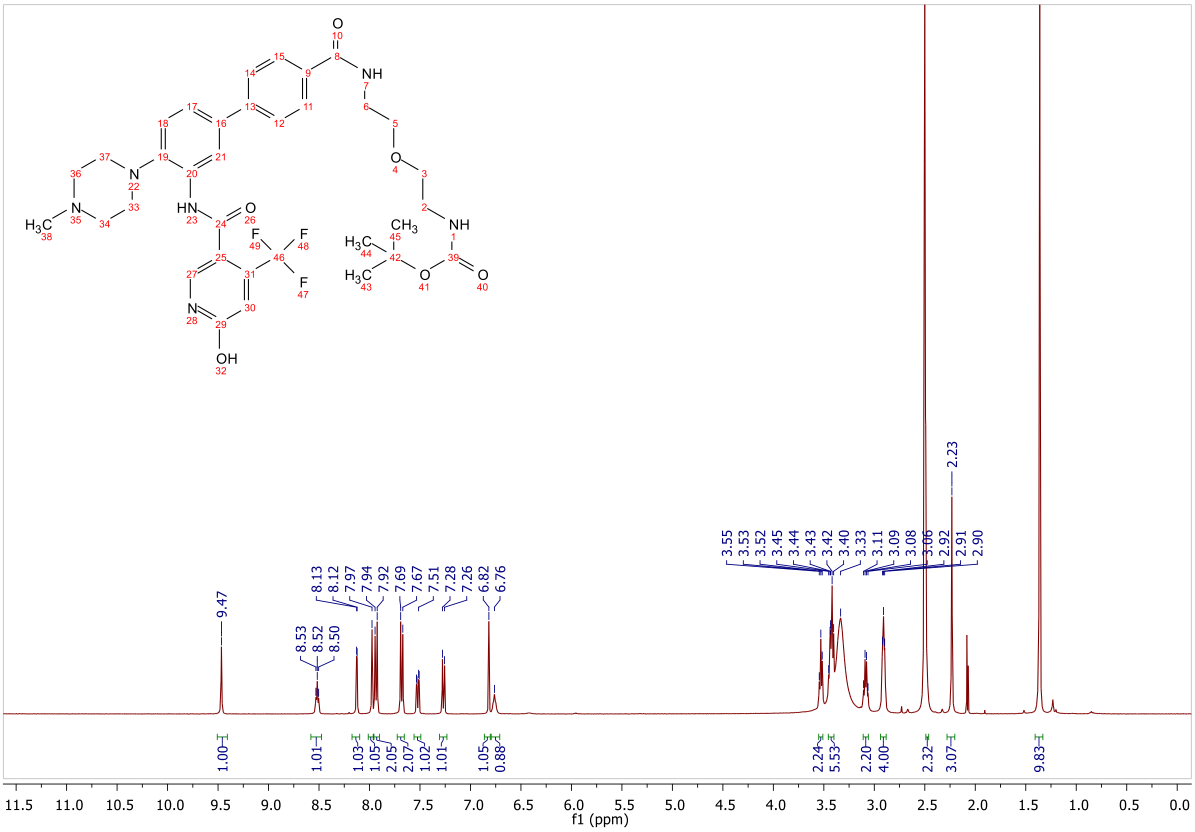

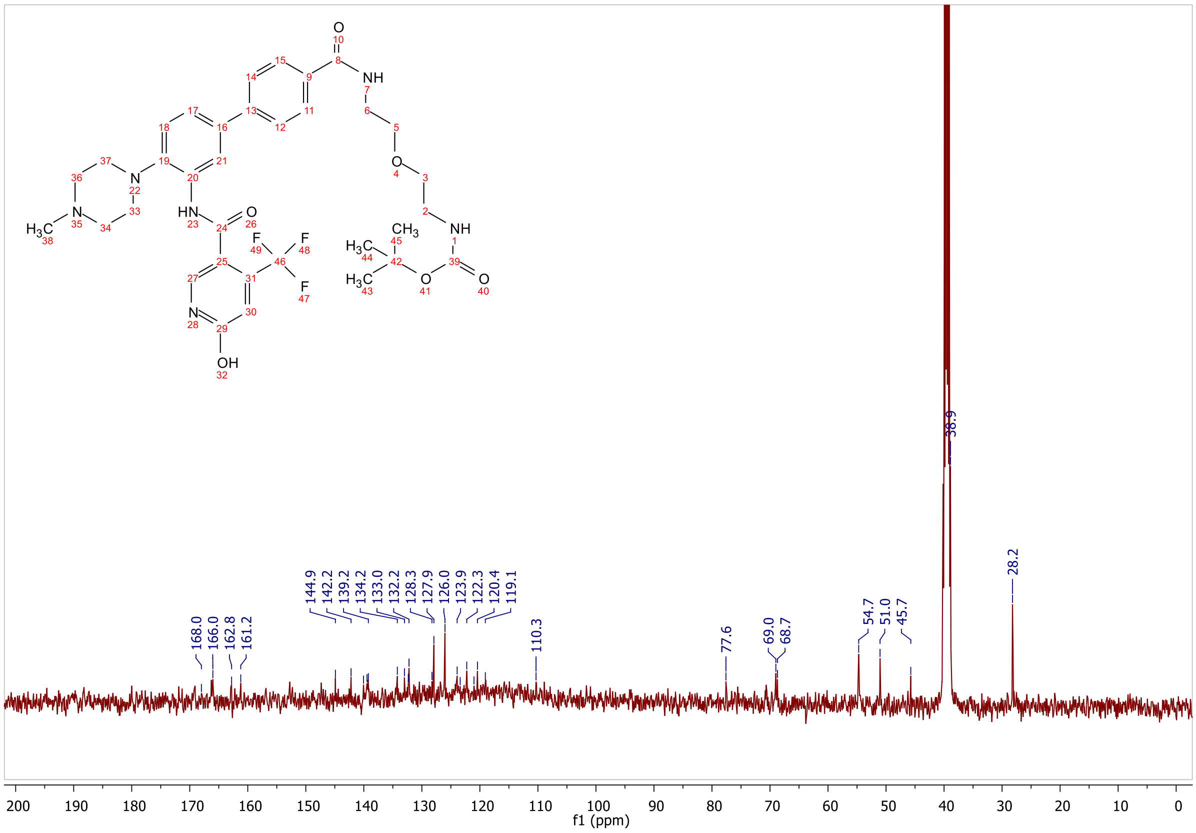
168.0
166.0
162.8
161.2
144.9
142.2
139.2
134.2
133.0
132.2
128.3
127.9
126.0
123.9
122.3
120.4
119.1
110.3
77.6
69.0
68.7
54.7
51.0
45.7
38.9
28.2
f1 (ppm)

**ESI, HPLC, $^1$H-NMR of tert-butyl (1-(3'-(6-hydroxy-4-(trifluoromethyl)nicotinamido)-4'-(4-methylpiperazin-1-yl)-[1,1'-biphenyl]-4-yl)-1-oxo-5,8,11,14,17,20,23-heptaoxa-2-azapentacosan-25-yl)carbamate 6b**

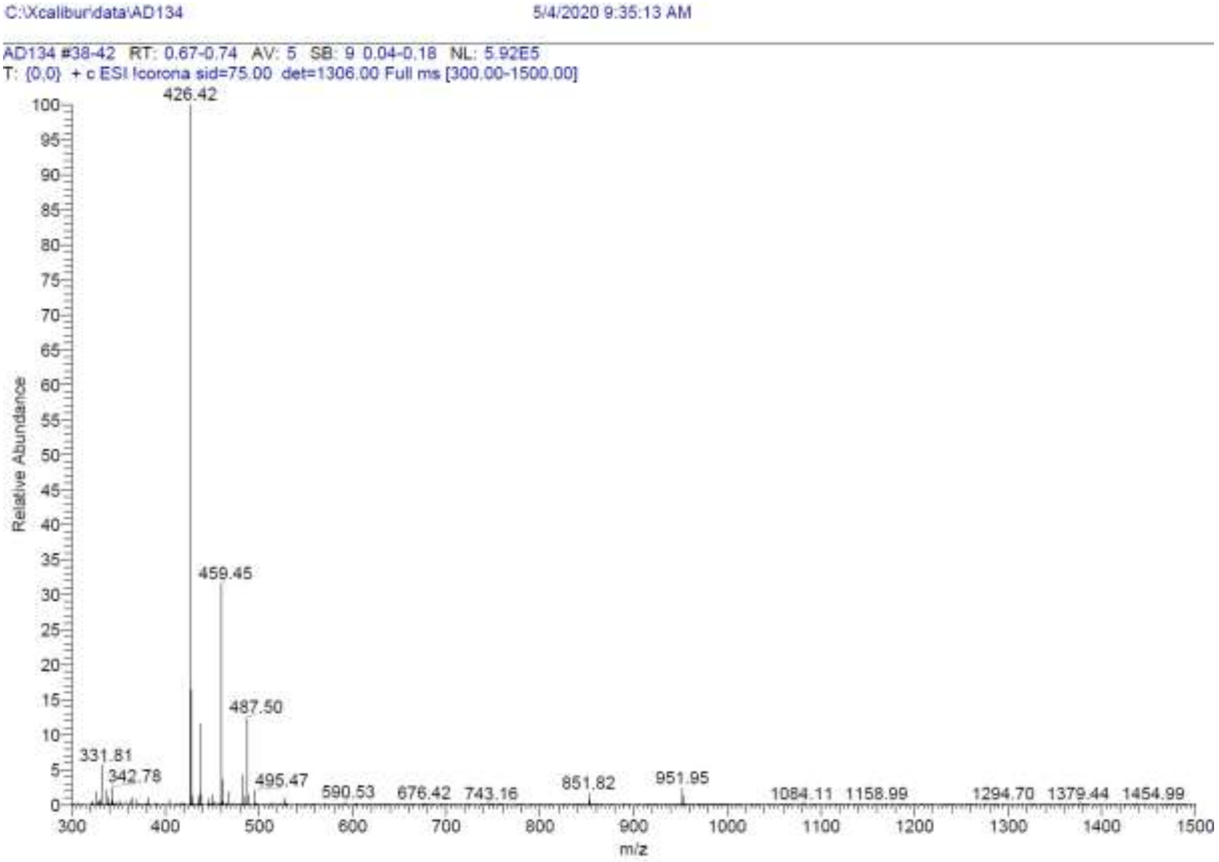

Sum 42114393.0000

**Signal:** MWD1 A, Sig=254,4 Ref=off

| RT [min] | Type | Width [min] | Area | Height | Area% |
|---|---|---|---|---|---|
| 11.751 | VV | 0.2234 | 917.7254 | 61.0386 | 94.3249 |
| 12.958 | MM | 0.2044 | 55.2154 | 4.5021 | 5.6751 |
| | | Sum | 972.9408 | | |

**Signal:** MWD1 B, Sig=280,4 Ref=off

| RT [min] | Type | Width [min] | Area | Height | Area% |
|---|---|---|---|---|---|
| 11.751 | VV | 0.2259 | 1091.4231 | 70.0104 | 98.2189 |
| 12.954 | MM | 0.2499 | 19.7914 | 1.3201 | 1.7811 |
| | | Sum | 1111.2145 | | |

**Signal:** MWD1 F, Sig=260,4 Ref=off

| RT [min] | Type | Width [min] | Area | Height | Area% |
|---|---|---|---|---|---|
| 11.751 | VV | 0.2199 | 1049.4673 | 69.5902 | 94.7728 |
| 12.958 | MM | 0.2140 | 57.8829 | 4.5091 | 5.2272 |

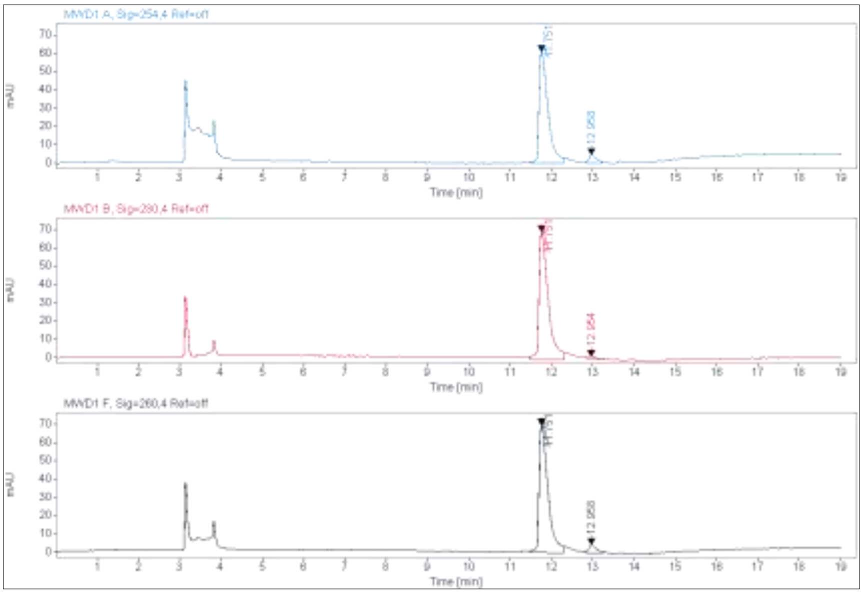
MWD1 A, Sig=254,4 Ref=off
MWD1 B, Sig=280,4 Ref=off
MWD1 F, Sig=260,4 Ref=off
mAU
Time [min]

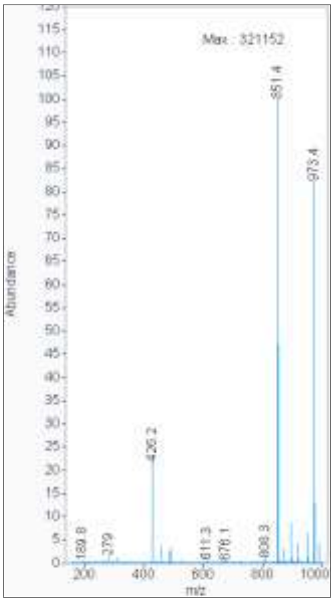
Max : 321152
851.4
973.4
426.2
Abundance
m/z

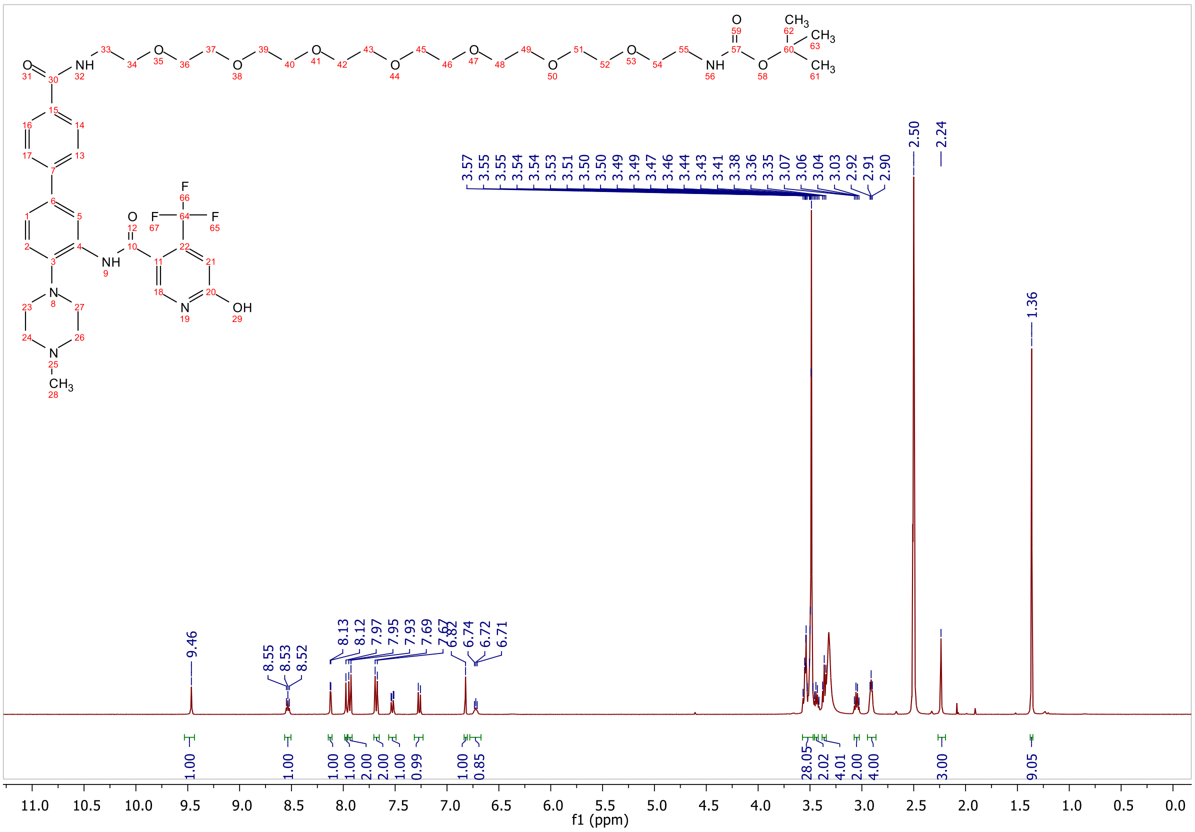
f1 (ppm)

**ESI, HPLC and $^{1}$H-NMR of tert-butyl (4-((3'-(6-hydroxy-4-(trifluoromethyl)nicotinamido)-4'-(4-methylpiperazin-1-yl)-[1,1'-biphenyl]-4-carboxamido)methyl)benzyl)carbamate 6c**

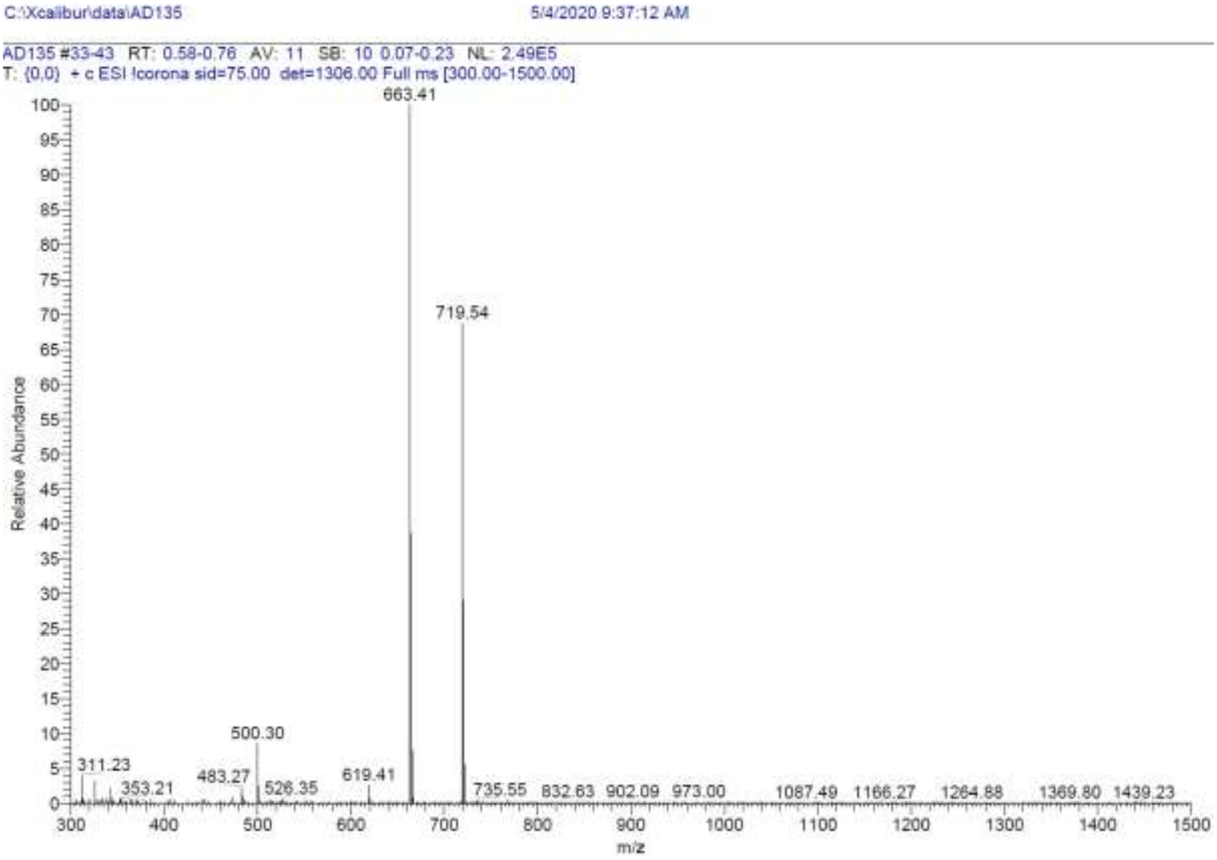

**Signal:** MWD1 A, Sig=254,4 Ref=off

| RT [min] | Type | Width [min] | Area | Height | Area% |
|---|---|---|---|---|---|
| 12.212 | VV | 0.2107 | 791.6523 | 56.4203 | 91.4191 |
| 12.949 | MM | 0.2487 | 74.3068 | 4.9802 | 8.5809 |
| | | Sum | 865.9591 | | |

**Signal:** MWD1 B, Sig=280,4 Ref=off

| RT [min] | Type | Width [min] | Area | Height | Area% |
|---|---|---|---|---|---|
| 12.211 | VV | 0.2149 | 943.0983 | 64.3726 | 96.0057 |
| 12.766 | MM | 0.2462 | 39.2378 | 2.6562 | 3.9943 |
| | | Sum | 982.3361 | | |

**Signal:** MWD1 F, Sig=260,4 Ref=off

| RT [min] | Type | Width [min] | Area | Height | Area% |
|---|---|---|---|---|---|
| 12.212 | VV | 0.2120 | 907.8795 | 64.2070 | 92.2728 |
| 12.951 | MM | 0.2594 | 76.0286 | 4.8847 | 7.7272 |

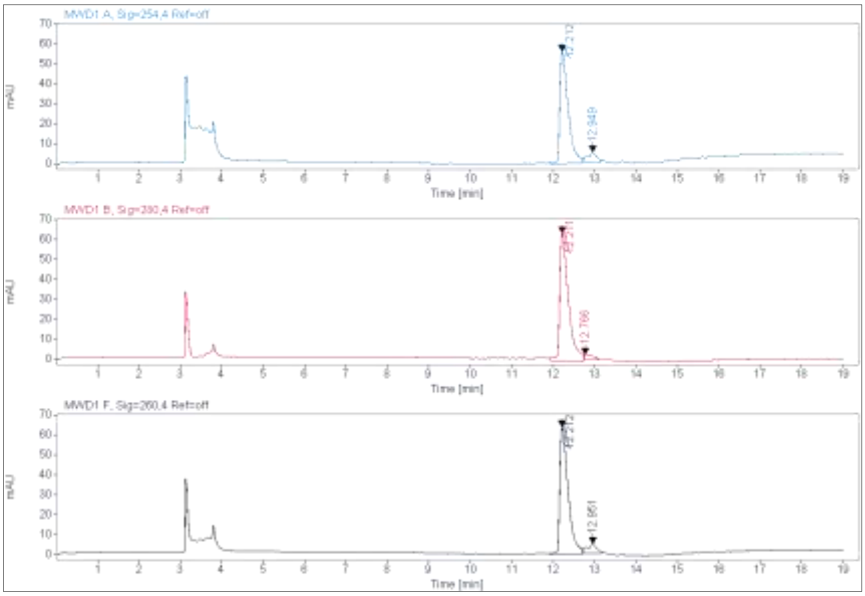

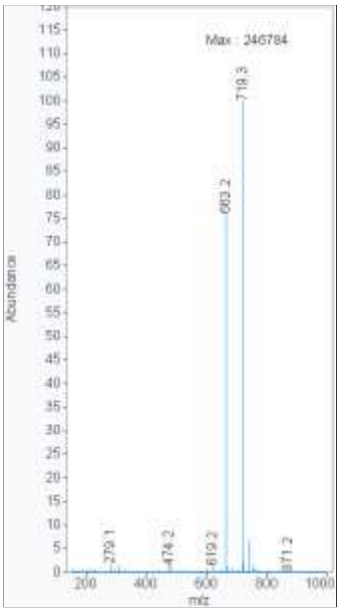

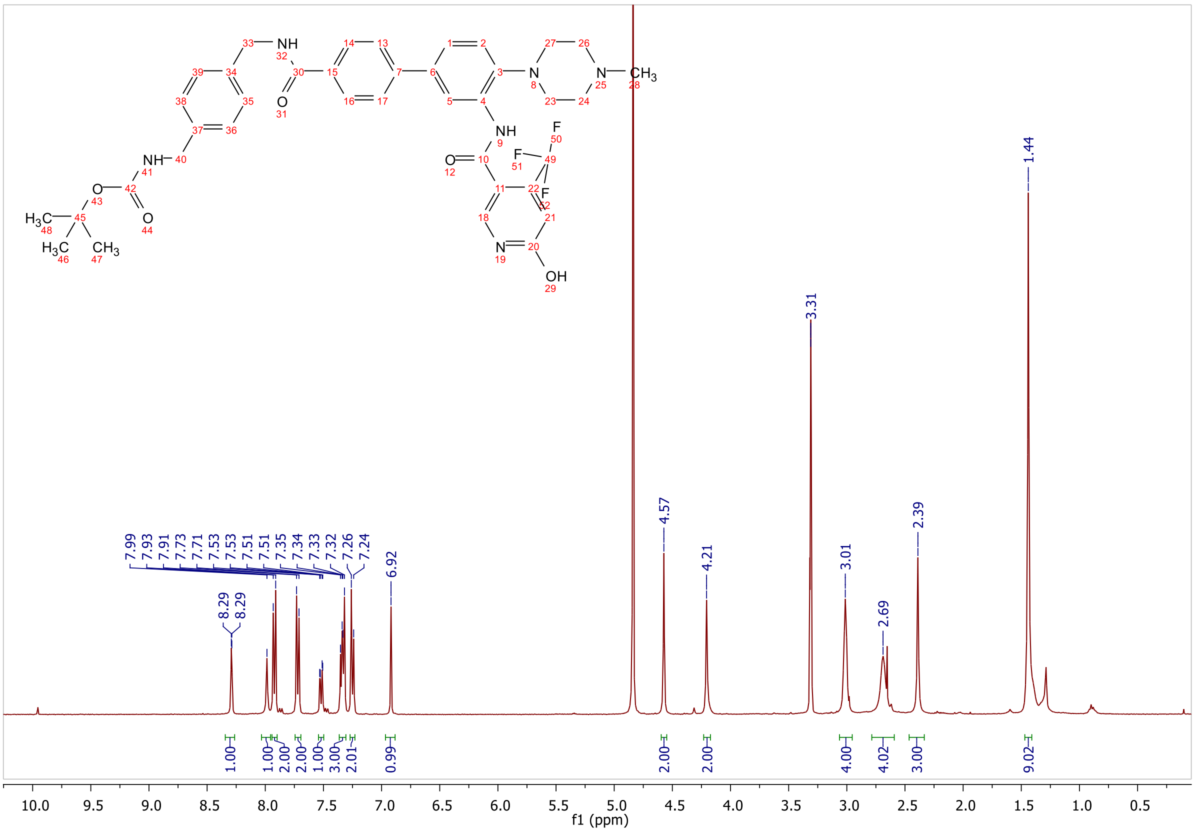

**MALDI, $^{1}$H-NMR of tert-butyl (1-(3'-(6-hydroxy-4-(trifluoromethyl)nicotinamido)-4'-(4-methylpiperazin-1-yl)-[1,1'-biphenyl]-4-yl)-1-oxo-5,8,11,14,17,20,23,26,29-nonaoxa-2-azahentriacontan-31-yl)carbamate S18a**

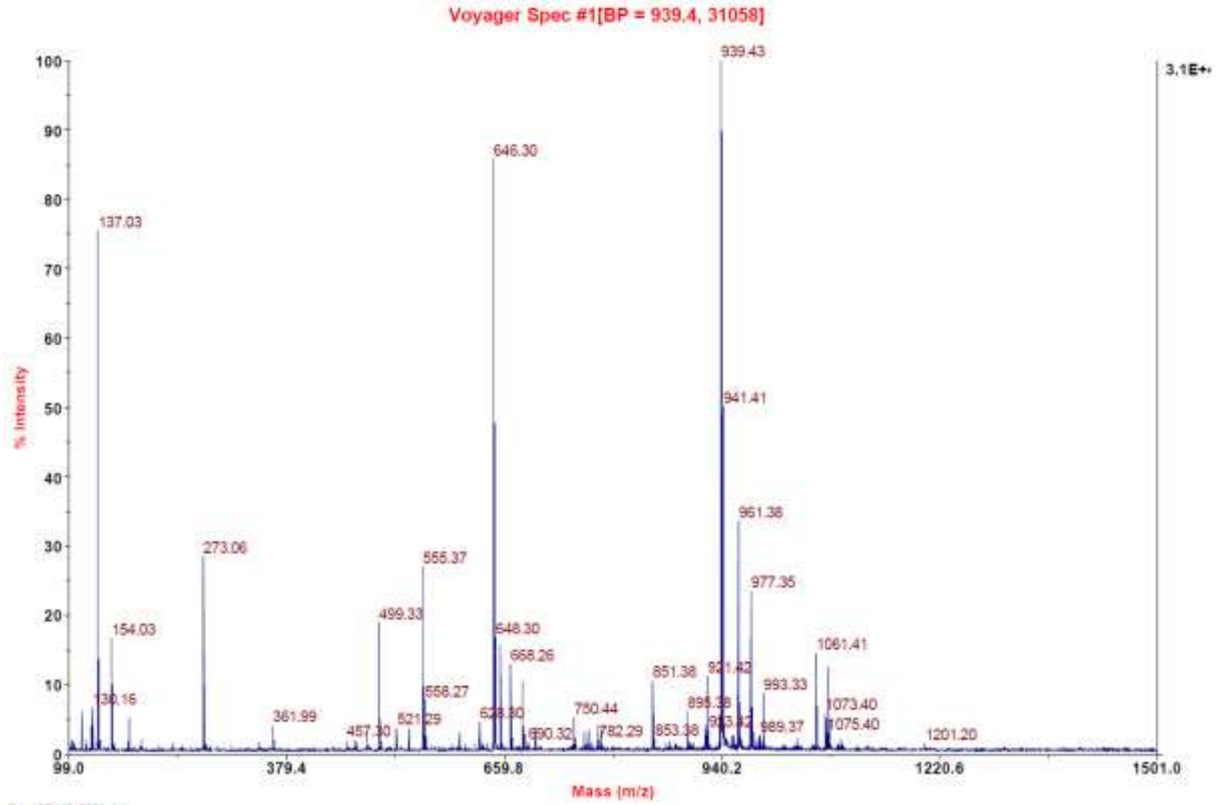

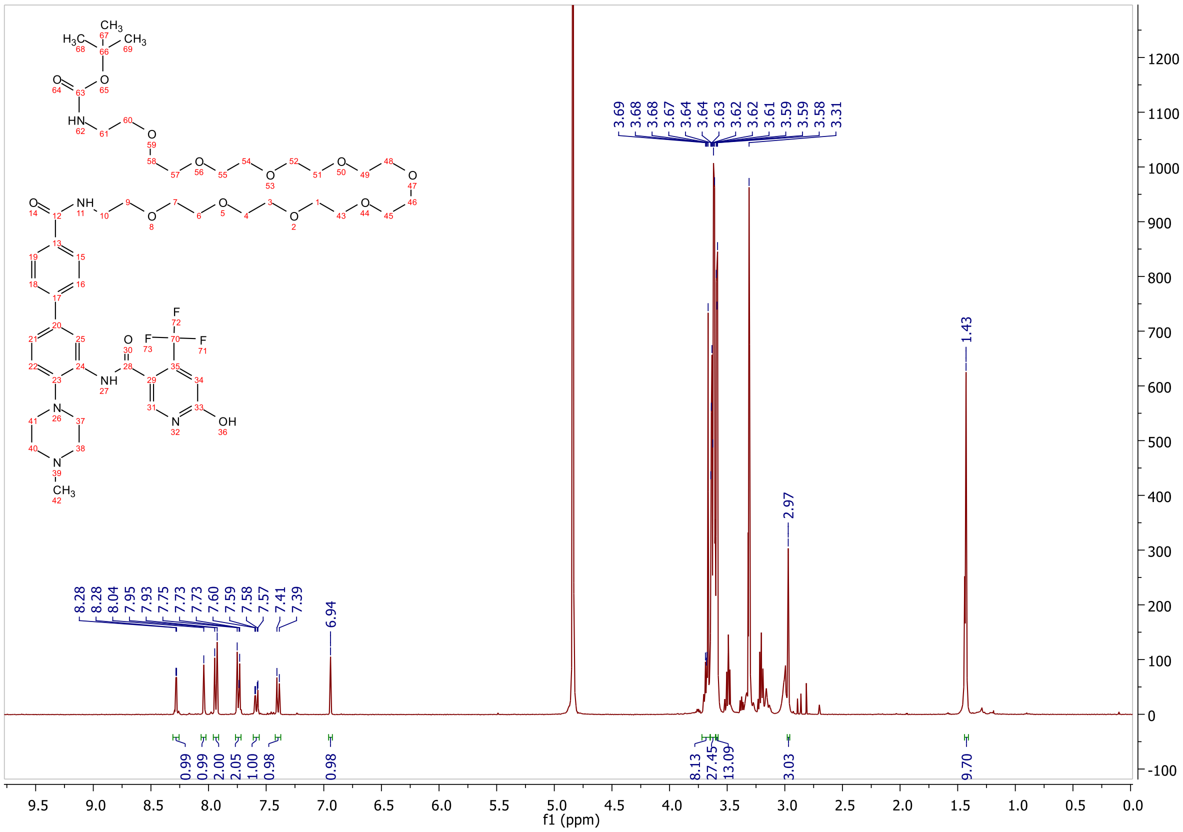

**ESI, HPLC, [1]H-NMR and [13]C-NMR of tert-butyl (1-(3'-(6-hydroxy-4-(trifluoromethyl)nicotinamido)-4'-(4-methylpiperazin-1-yl)-[1,1'-biphenyl]-4-yl)-1-oxo-6,9,12-trioxa-2-azapentadecan-15-yl)carbamate S18b**

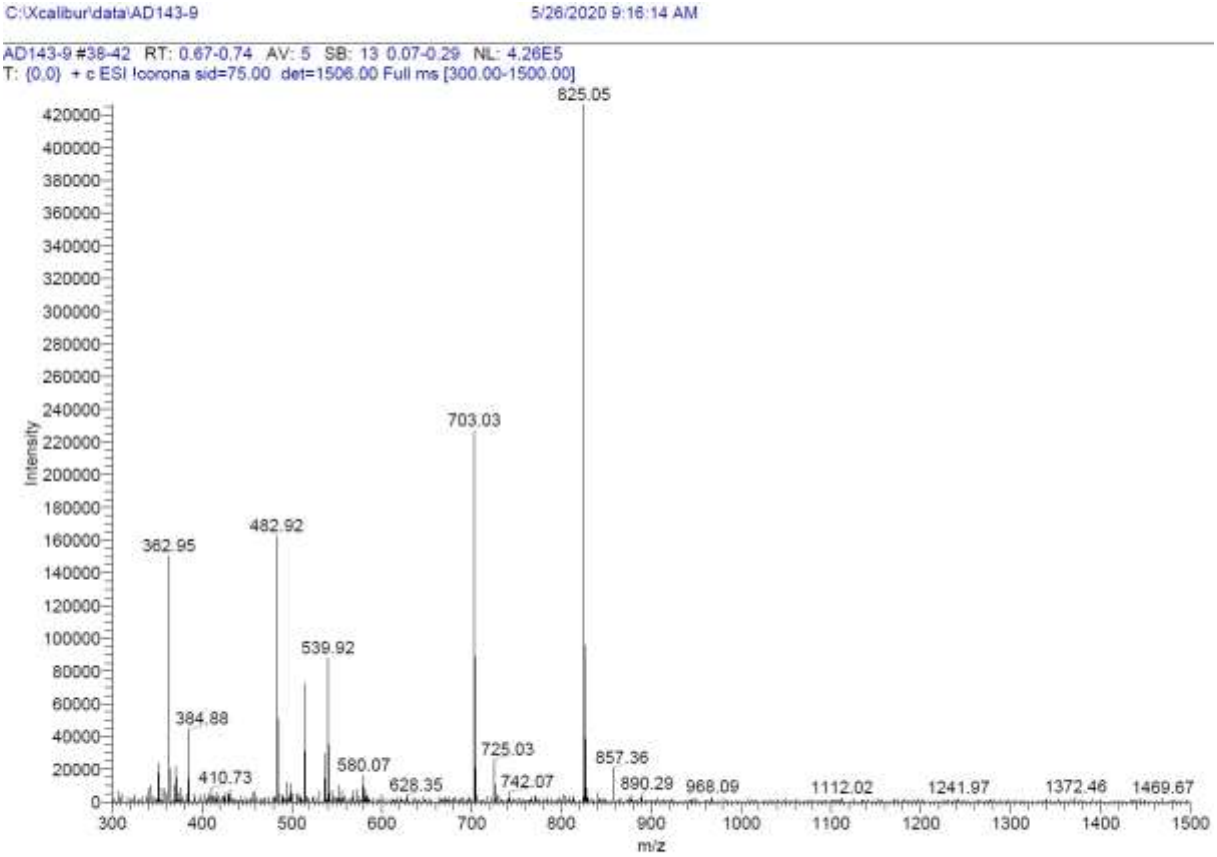

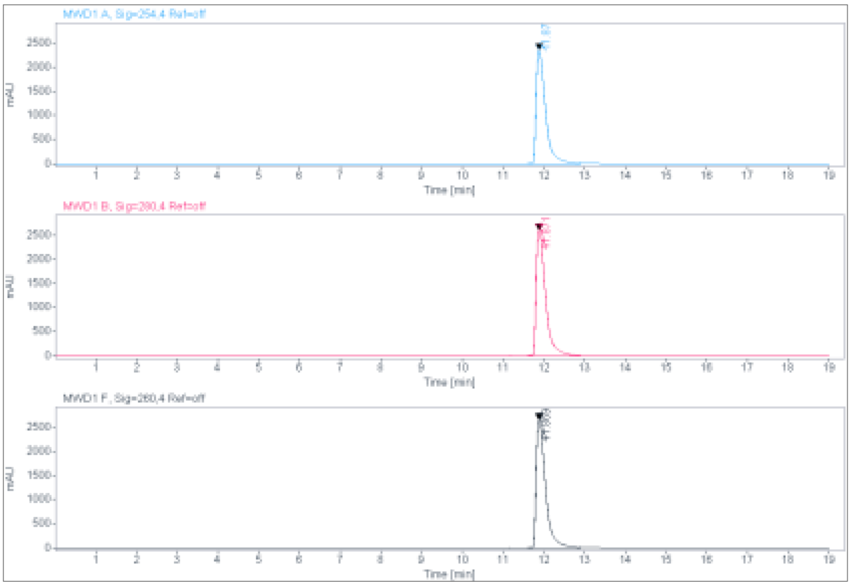

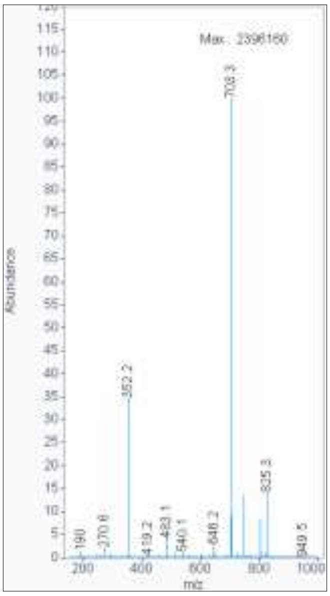

**Signal:** MWD1 A, Sig=254,4 Ref=off

| RT [min] | Type | Width [min] | Area | Height | Area% | Name |
|---|---|---|---|---|---|---|
| 11.870 | BV | 0.2371 | 37841.2813 | 2359.7388 | 100.0000 | |
| | | Sum | 37841.2813 | | | |

**Signal:** MWD1 B, Sig=280,4 Ref=off

| RT [min] | Type | Width [min] | Area | Height | Area% | Name |
|---|---|---|---|---|---|---|

| | | | | | |
|---|---|---|---|---|---|
| 11.871 | VV | 0.2203 | 42077.1367 | 2606.3718 | 100.0000 |
| | | Sum | 42077.1367 | | |

**Signal:** MWD1 F, Sig=260,4 Ref=off

| RT [min] | Type | Width [min] | Area | Height | Area% | Name |
|---|---|---|---|---|---|---|
| 11.869 | VV | 0.2388 | 42766.1367 | 2656.3064 | 100.0000 | |

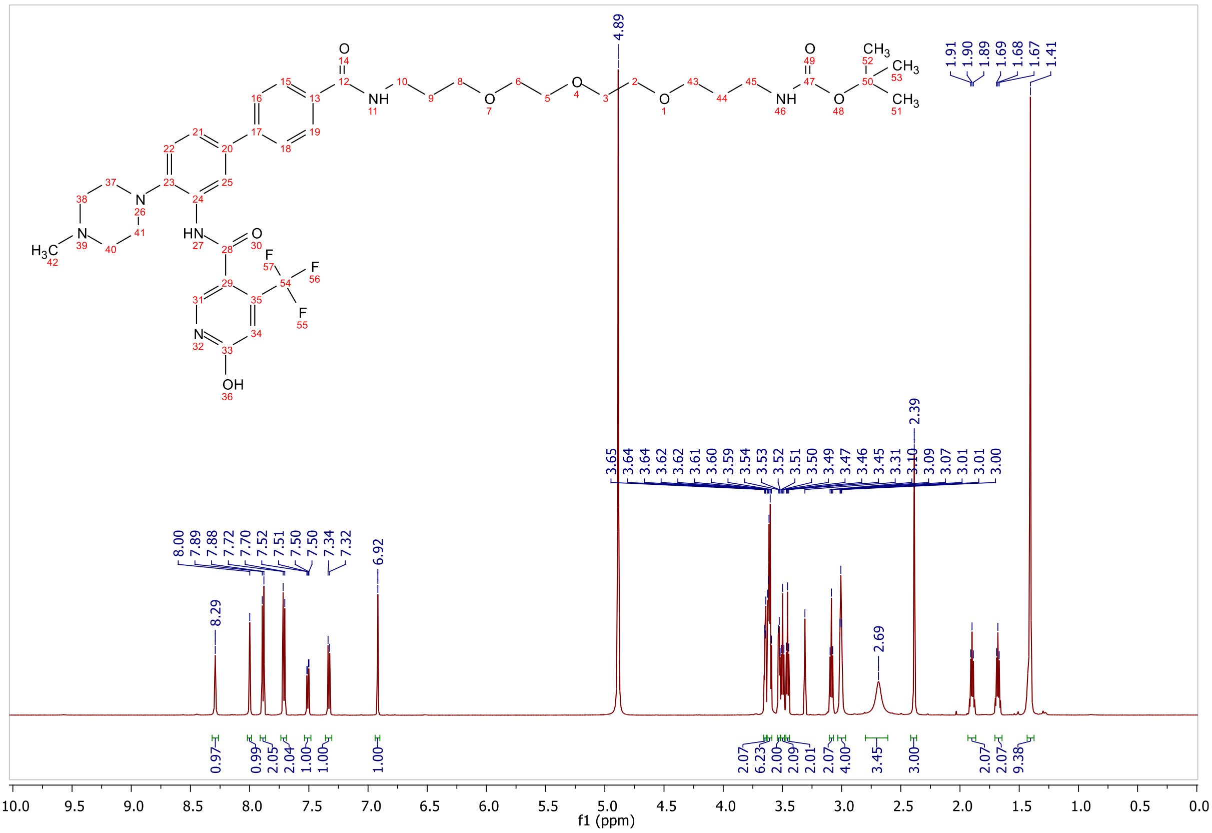

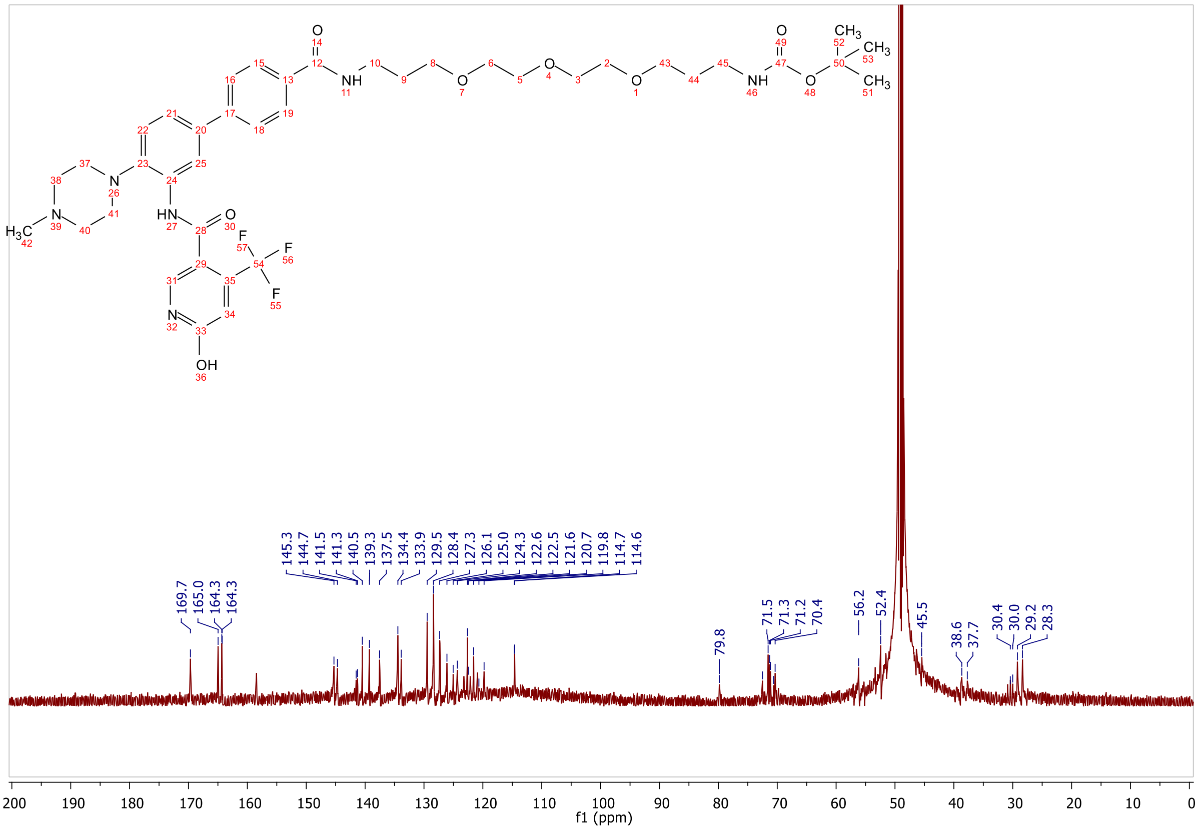
169.7
165.0
164.3
164.3
145.3
144.7
141.5
141.3
140.5
139.3
137.5
134.4
133.9
129.5
128.4
127.3
126.1
125.0
124.3
122.6
122.5
121.6
120.7
119.8
114.7
114.6
79.8
71.5
71.3
71.2
70.4
56.2
52.4
45.5
38.6
37.7
30.4
30.0
29.2
28.3
f1 (ppm)

**HPLC, HRMS, $^{1}H$-NMR and $^{13}C$-NMR of N-(4'-((2-(2-(3-(5,5-difluoro-7-(1H-pyrrol-2-yl)-5H-5l4,6l4-dipyrrolo[1,2-c:2',1'-f][1,3,2]diazaborinin-3-yl)propanamido)ethoxy)ethyl)carbamoyl)-4-(4-methylpiperazin-1-yl)-[1,1'-biphenyl]-3-yl)-6-hydroxy-4-(trifluoromethyl)nicotinamide 19a**

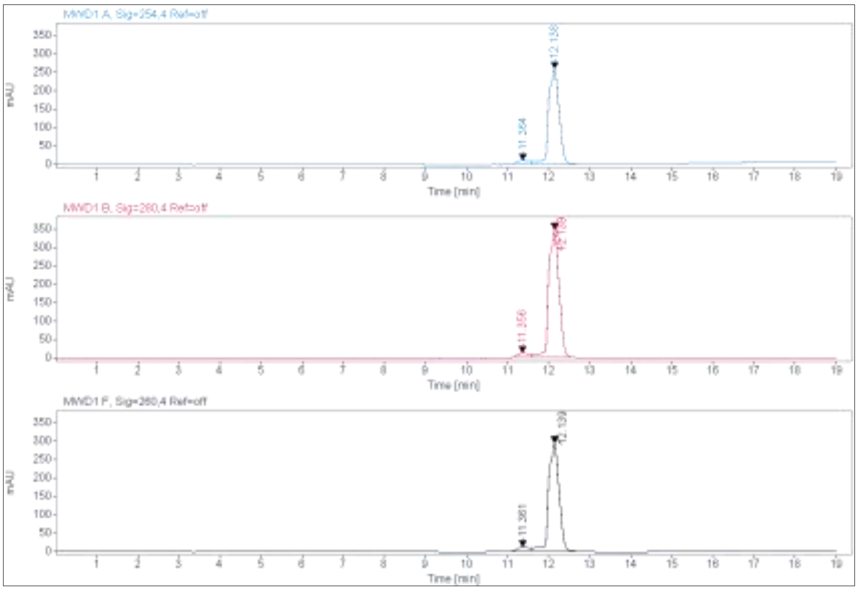

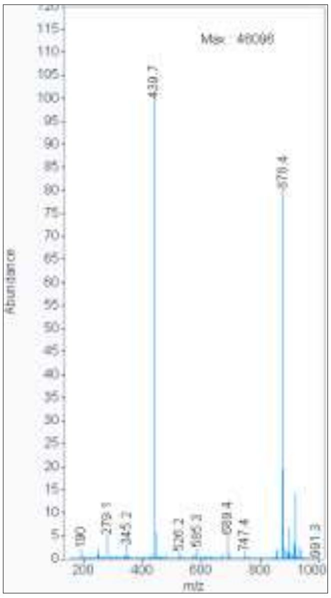

**Signal:** MWD1 A, Sig=254,4 Ref=off

| RT [min] | Type | Width [min] | Area | Height | Area% |
|---|---|---|---|---|---|
| 11.364 | MM | 0.3356 | 228.0416 | 11.3250 | 4.6822 |
| 12.138 | MM | 0.2965 | 4642.3574 | 260.9971 | 95.3178 |
| | | Sum | 4870.3990 | | |

**Signal:** MWD1 B, Sig=280,4 Ref=off

| RT [min] | Type | Width [min] | Area | Height | Area% |
|---|---|---|---|---|---|
| 11.356 | MM | 0.3379 | 265.3242 | 13.0870 | 4.1852 |
| 12.139 | MM | 0.2911 | 6074.2964 | 347.7672 | 95.8148 |
| | | Sum | 6339.6206 | | |

**Signal:** MWD1 F, Sig=260,4 Ref=off

| RT [min] | Type | Width [min] | Area | Height | Area% |
|---|---|---|---|---|---|
| 11.361 | MM | 0.3556 | 252.2446 | 11.8215 | 4.6842 |
| 12.139 | MM | 0.2909 | 5132.7949 | 294.1020 | 95.3158 |

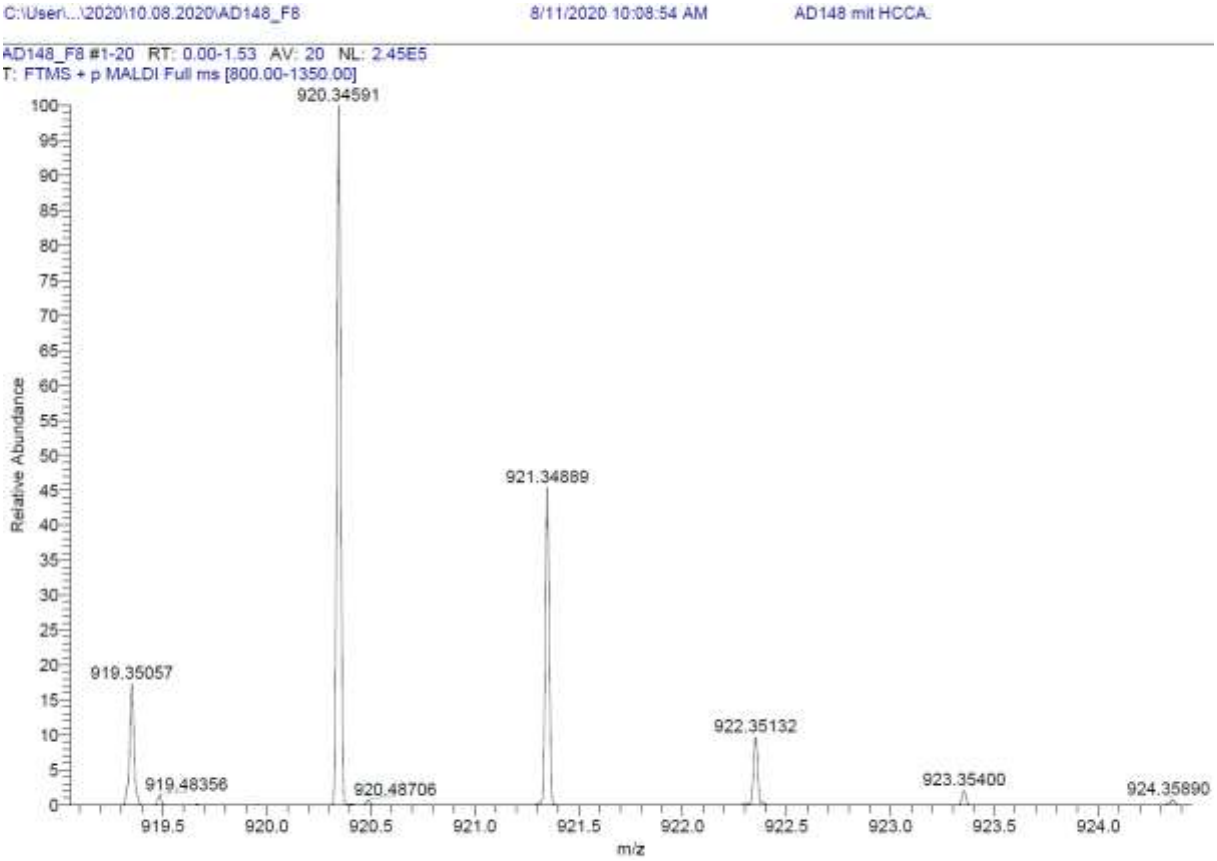
C:\User\...\2020\10.08.2020\AD148_F8
8/11/2020 10:08:54 AM
AD148 mit HCCA
AD148_F8 #1-20 RT: 0.00-1.53 AV: 20 NL: 2.45E5
T: FTMS + p MALDI Full ms [800.00-1350.00]
920.34591
921.34889
919.35057
922.35132
923.35400
924.35890
919.48356
920.48706
Relative Abundance
m/z

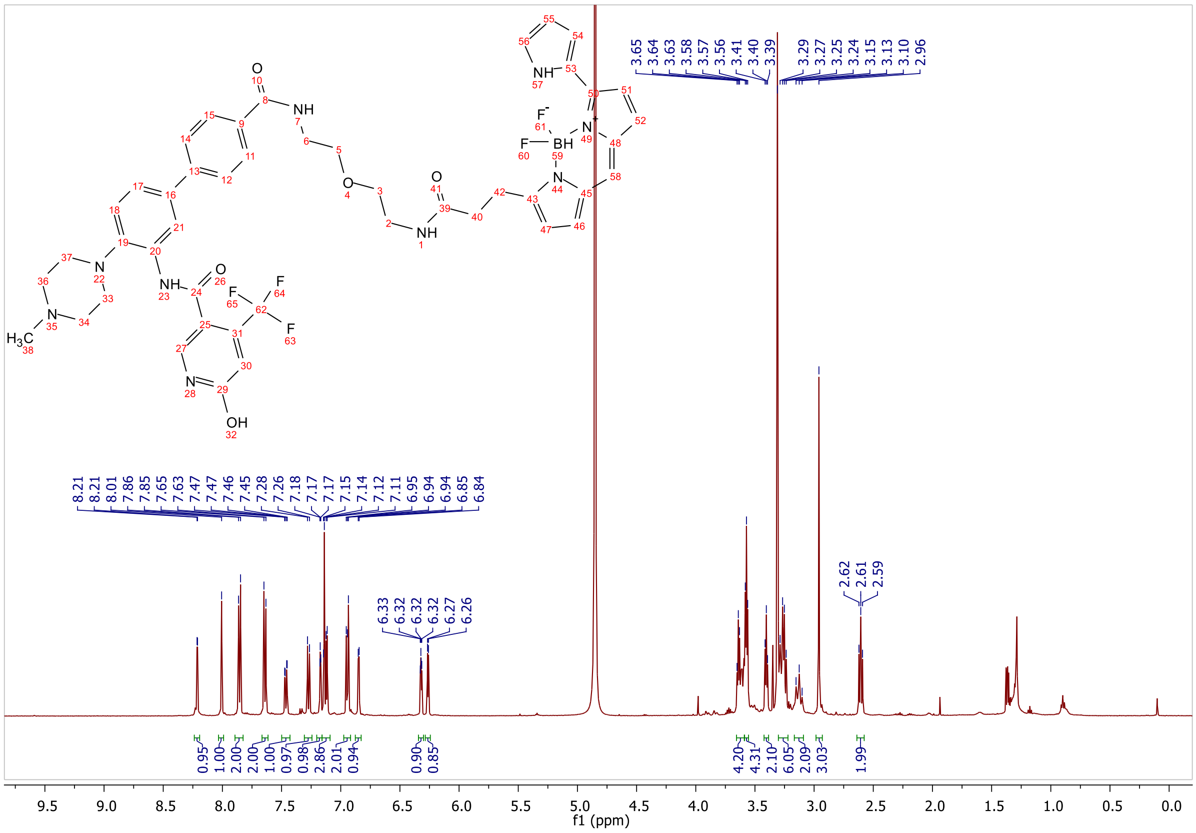
f1 (ppm)

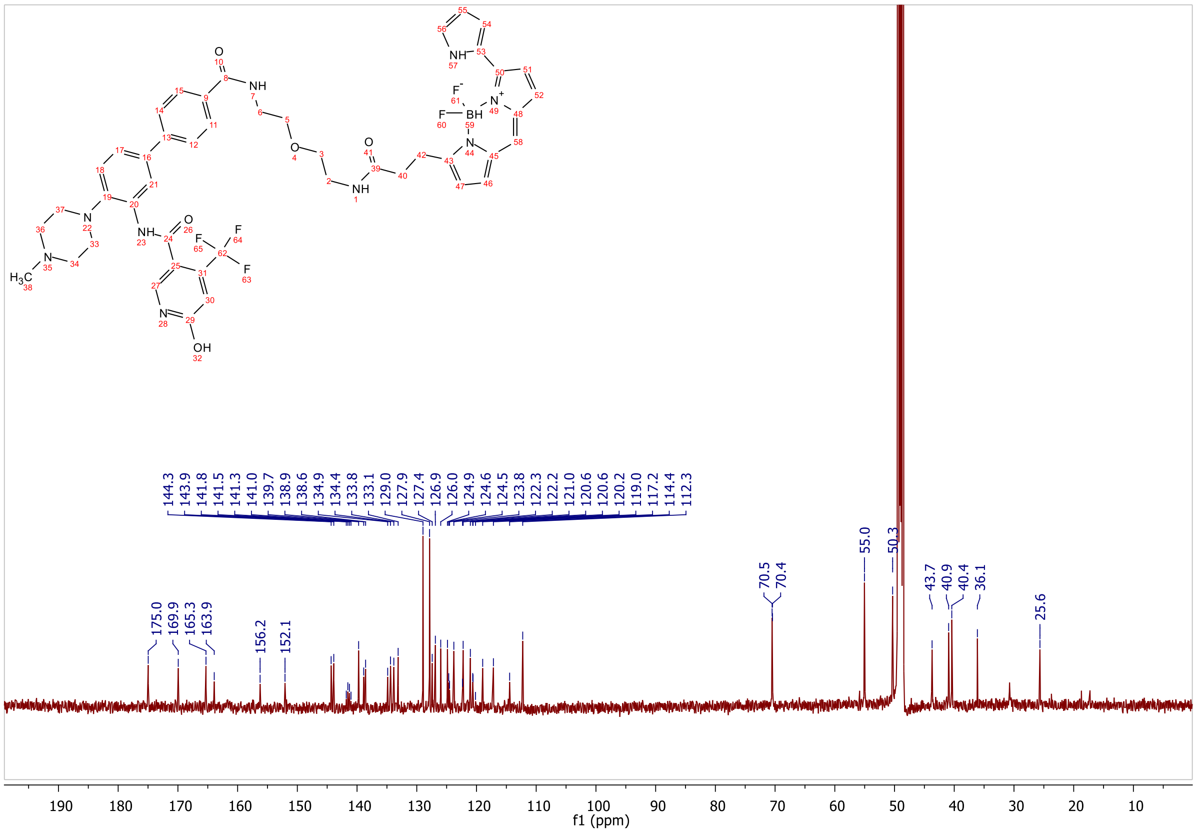

175.0
169.9
165.3
163.9
156.2
152.1
144.3
143.9
141.8
141.5
141.3
141.0
139.7
138.9
138.6
134.9
134.4
133.8
133.1
129.0
127.9
127.4
126.9
126.0
124.9
124.6
124.5
123.8
122.3
122.2
121.0
120.6
120.6
120.2
119.0
117.2
114.4
112.3
70.5
70.4
55.0
50.3
43.7
40.9
40.4
36.1
25.6
f1 (ppm)

**HPLC, HRMS, $^{1}$H-NMR of N-(4'-((33-(5,5-difluoro-7-(1H-pyrrol-2-yl)-5H-5l4,6l4-dipyrrolo[1,2-c:2',1'-f][1,3,2]diazaborinin-3-yl)-31-oxo-3,6,9,12,15,18,21,24,27-nonaoxa-30-azatritriacontyl)carbamoyl)-4-(4-methylpiperazin-1-yl)-[1,1'-biphenyl]-3-yl)-6-hydroxy-4-(trifluoromethyl)nicotinamide 19b**

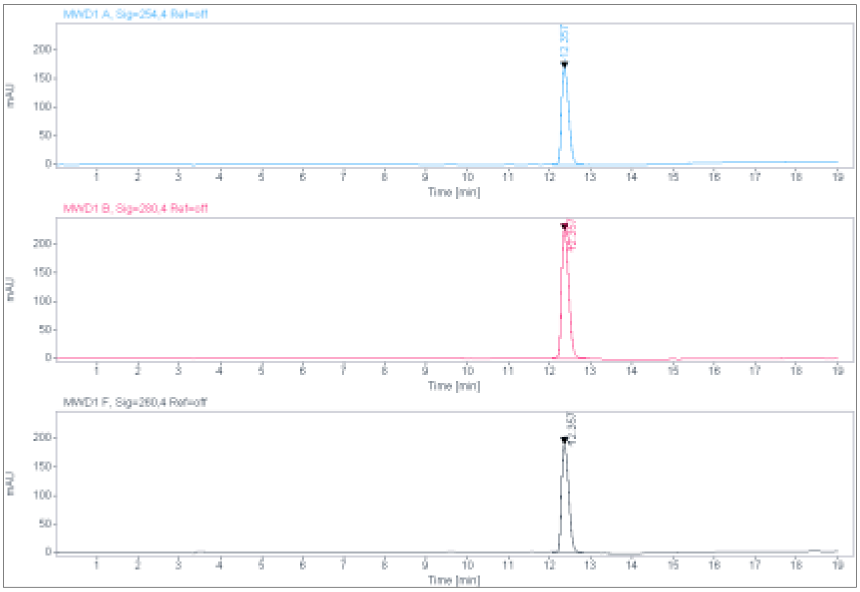

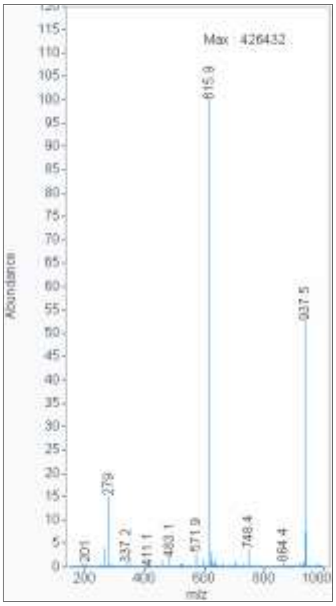

**Signal:** MWD1 A, Sig=254,4 Ref=off

| RT [min] | Type | Width [min] | Area | Height | Area% | Name |
|---|---|---|---|---|---|---|
| 12.357 | VV | 0.1987 | 2072.8545 | 168.3288 | 100.0000 | |
| | | Sum | 2072.8545 | | | |

**Signal:** MWD1 B, Sig=280,4 Ref=off

| RT [min] | Type | Width [min] | Area | Height | Area% | Name |
|---|---|---|---|---|---|---|
| 12.357 | VV | 0.2005 | 2813.1299 | 225.5552 | 100.0000 | |
| | | Sum | 2813.1299 | | | |

**Signal:** MWD1 F, Sig=260,4 Ref=off

| RT [min] | Type | Width [min] | Area | Height | Area% | Name |
|---|---|---|---|---|---|---|
| 12.357 | VV | 0.1988 | 2353.3545 | 190.8938 | 100.0000 | |

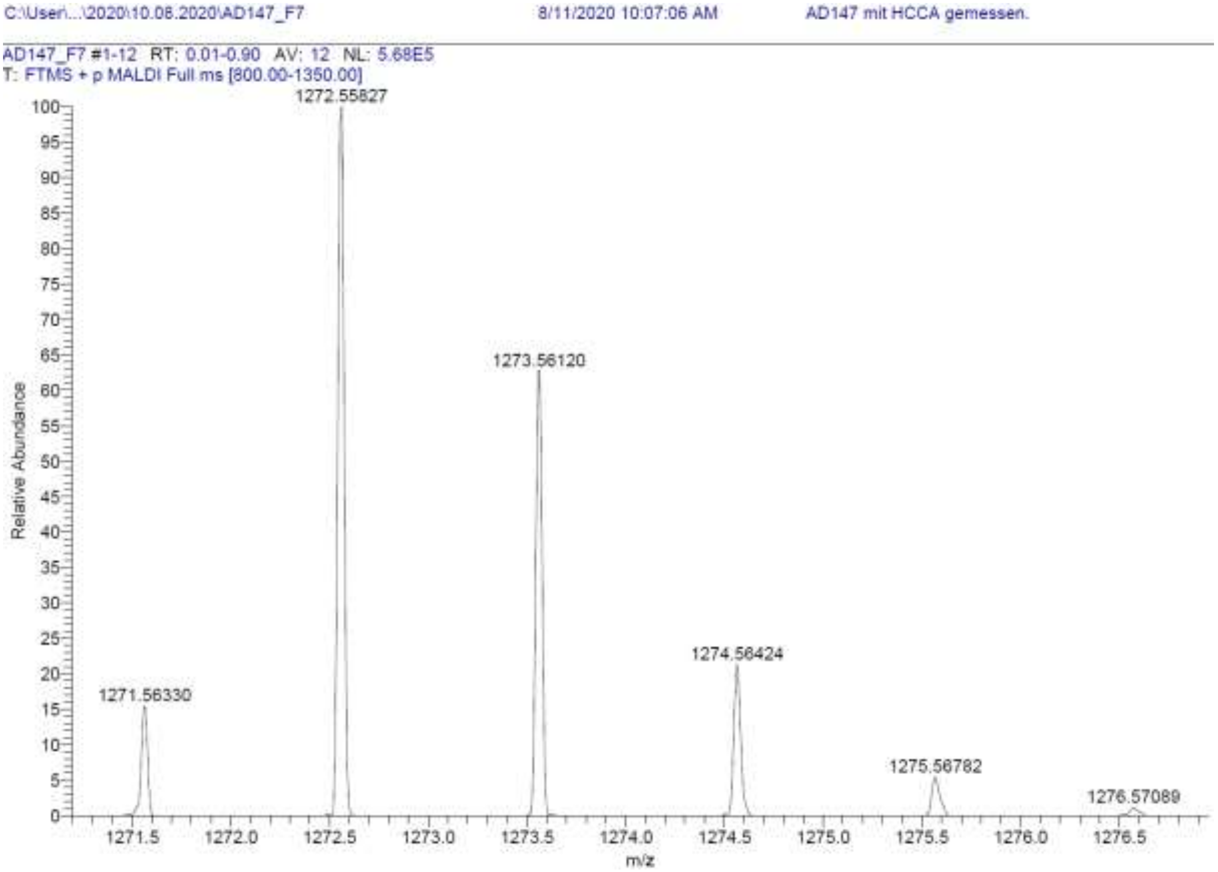
C:\User\...\2020\10.08.2020\AD147_F7
8/11/2020 10:07:06 AM
AD147 mit HCCA gemessen.
AD147_F7 #1-12 RT: 0.01-0.90 AV: 12 NL: 5.68E5
T: FTMS + p MALDI Full ms [800.00-1350.00]
1271.56330
1272.55827
1273.56120
1274.56424
1275.56782
1276.57089
Relative Abundance
m/z

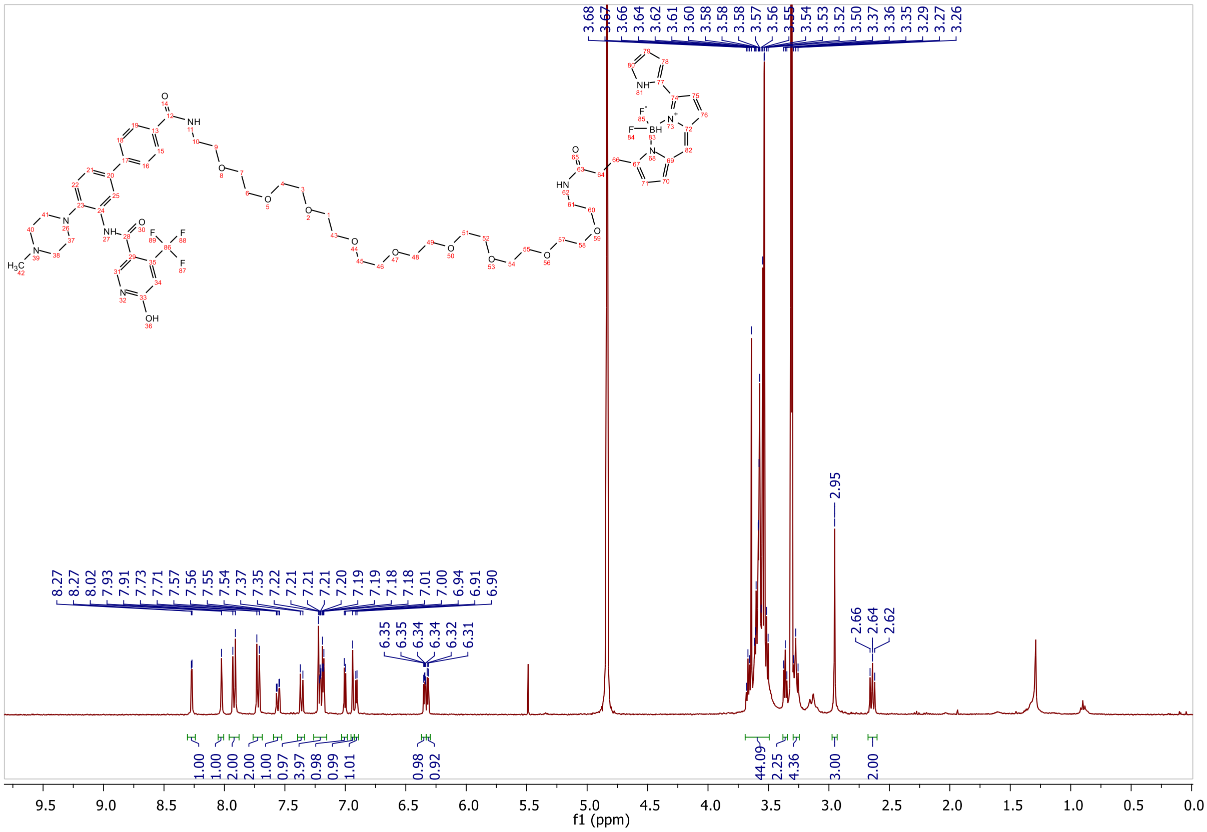
3.68 3.67 3.66 3.64 3.62 3.61 3.60 3.58 3.58 3.57 3.56 3.55 3.54 3.53 3.52 3.50 3.37 3.36 3.35 3.29 3.27 3.26
8.27 8.27 8.02 7.93 7.91 7.73 7.71 7.57 7.56 7.55 7.54 7.37 7.35 7.22 7.21 7.21 7.21 7.20 7.19 7.19 7.18 7.18 7.01 7.00 6.94 6.91 6.90
6.35 6.35 6.34 6.34 6.32 6.31
2.95
2.66 2.64 2.62
1.00 1.00 2.00 2.00 1.00 0.97 3.97 0.98 0.99 1.01 0.98 0.92 44.09 2.25 4.36 3.00 2.00
f1 (ppm)

**HPLC, HRMS, $^{1}$H-NMR and $^{1}$H-$^{13}$C-HMBC of N-(4'-((17-(5,5-difluoro-7-(1H-pyrrol-2-yl)-5H-5l4,6l4-dipyrrolo[1,2-c:2',1'-f][1,3,2]diazaborinin-3-yl)-15-oxo-4,7,10-trioxa-14-azaheptadecyl)carbamoyl)-4-(4-methylpiperazin-1-yl)-[1,1'-biphenyl]-3-yl)-6-hydroxy-4-(trifluoromethyl)nicotinamide 19c**

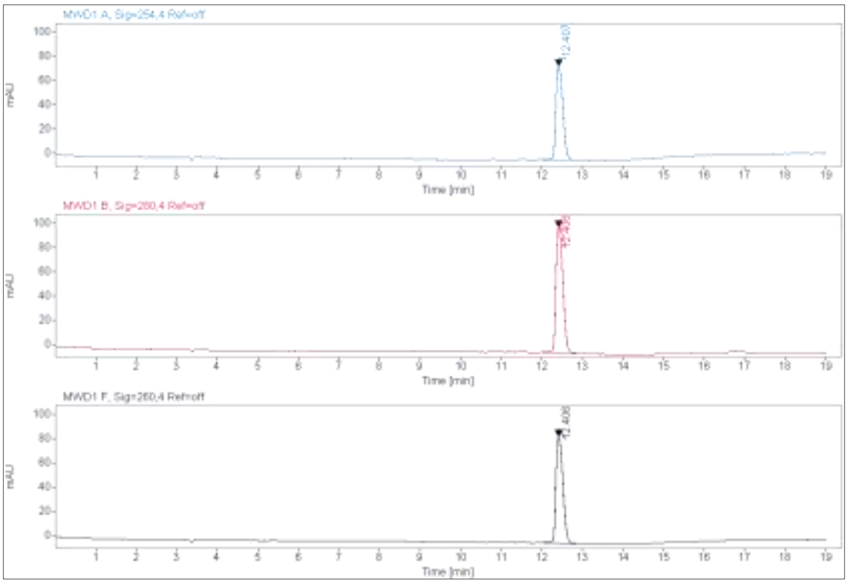

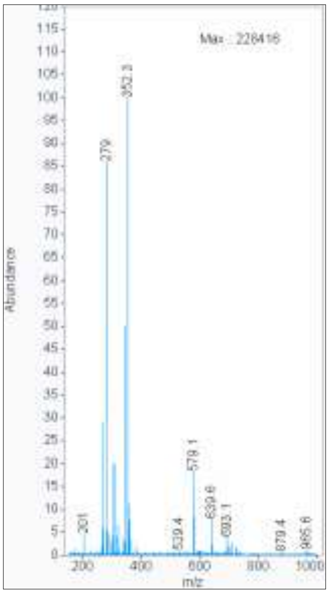

**Signal:** MWD1 A, Sig=254,4 Ref=off

| RT [min] | Type | Width [min] | Area | Height | Area% | Name |
|---|---|---|---|---|---|---|
| 12.407 | VV | 0.2010 | 936.6609 | 77.5274 | 100.0000 | |
| | | Sum | 936.6609 | | | |

**Signal:** MWD1 B, Sig=280,4 Ref=off

| RT [min] | Type | Width [min] | Area | Height | Area% | Name |
|---|---|---|---|---|---|---|
| 12.406 | VV | 0.2034 | 1279.1853 | 104.0625 | 100.0000 | |
| | | Sum | 1279.1853 | | | |

**Signal:** MWD1 F, Sig=260,4 Ref=off

| RT [min] | Type | Width [min] | Area | Height | Area% | Name |
|---|---|---|---|---|---|---|
| 12.406 | VV | 0.2009 | 1061.8375 | 87.9442 | 100.0000 | |

C:\User\...\2020\10.08.2020\AD146_F6 8/11/2020 10:05:47 AM AD146 mit HCCA gemessen.

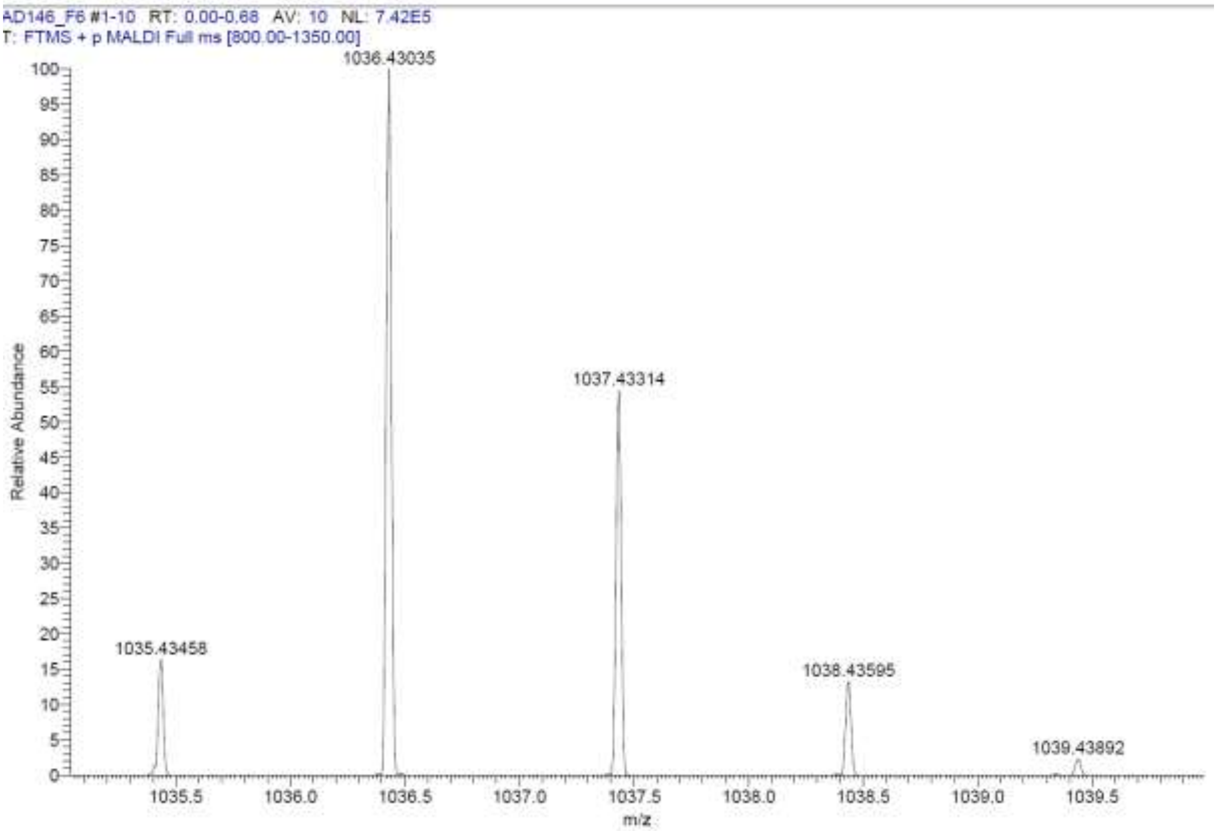

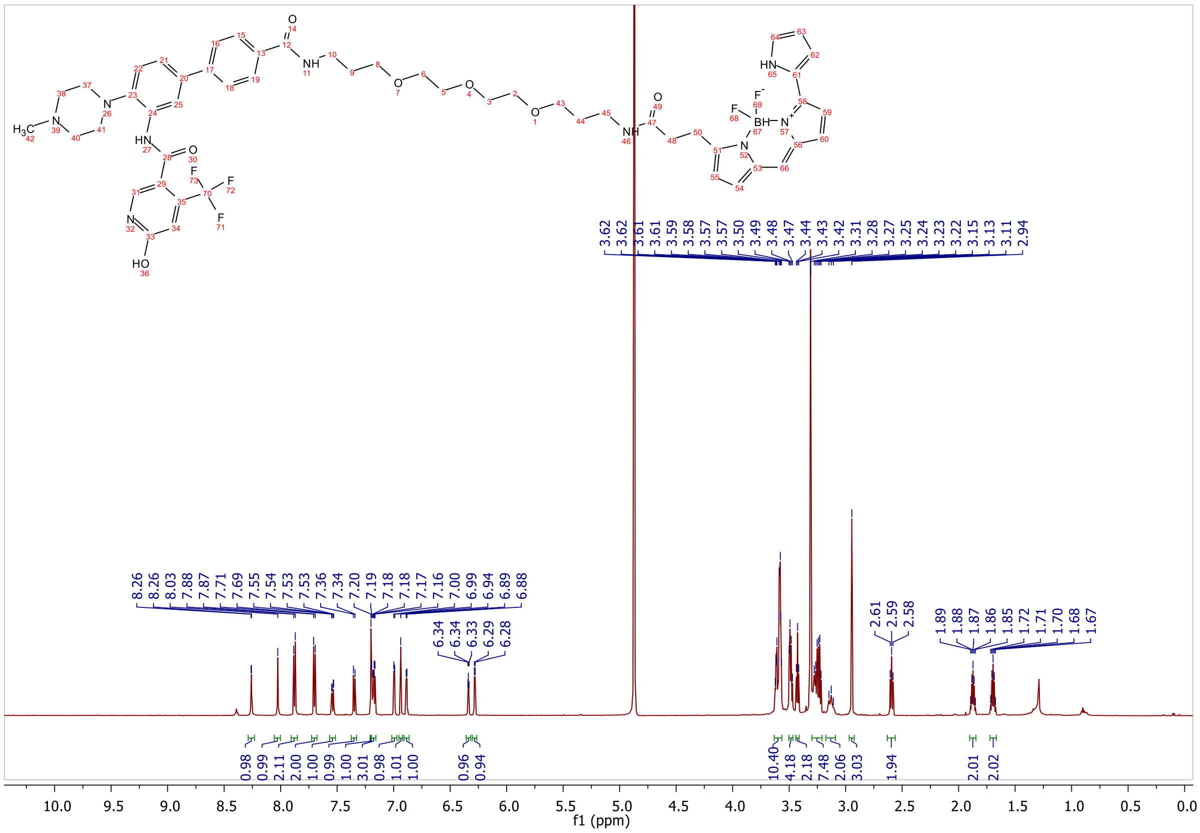

### 1.3 Appendix of intermediates 3-6

**ESI, [1]H-NMR of 1-(4-bromo-2-nitrophenyl)-4-methylpiperazine 3**

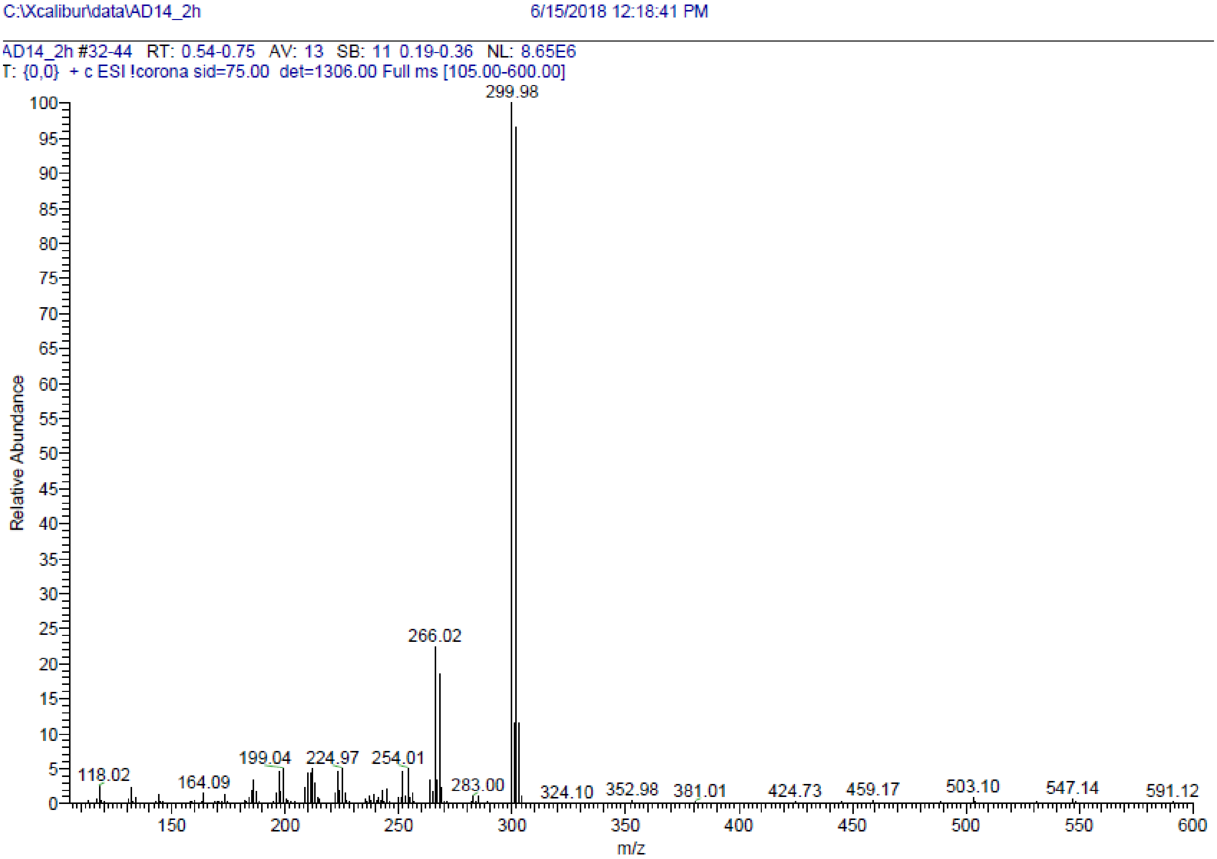

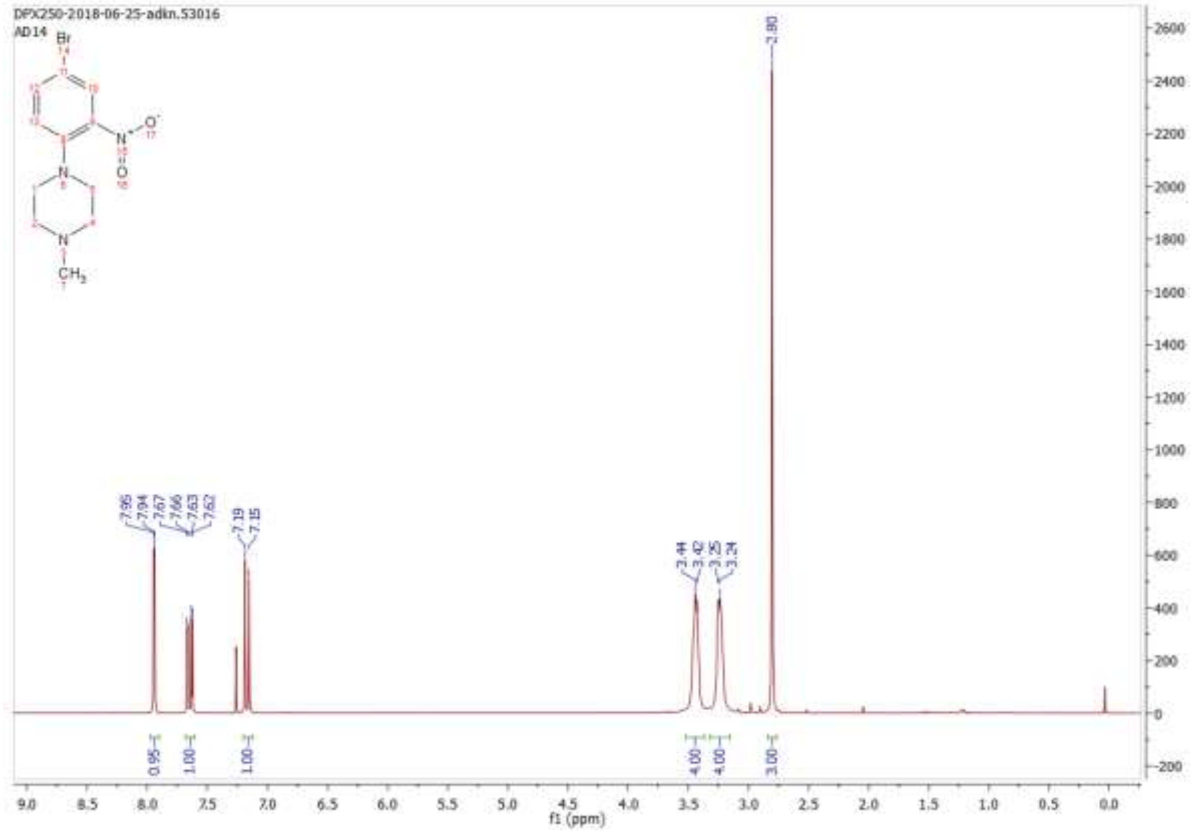

ESI, [1]H-NMR, HPLC of 5-bromo-2-(4-methylpiperazin-1-yl)aniline 4

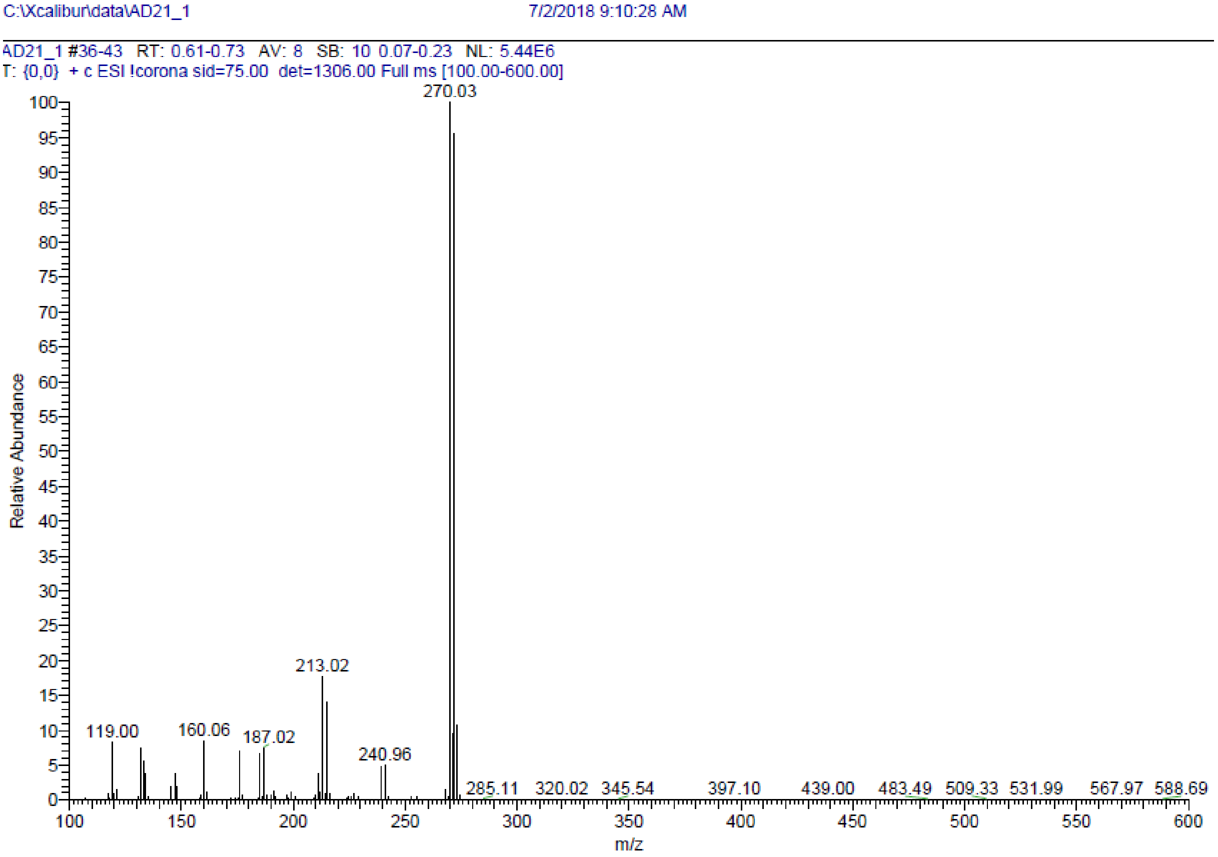

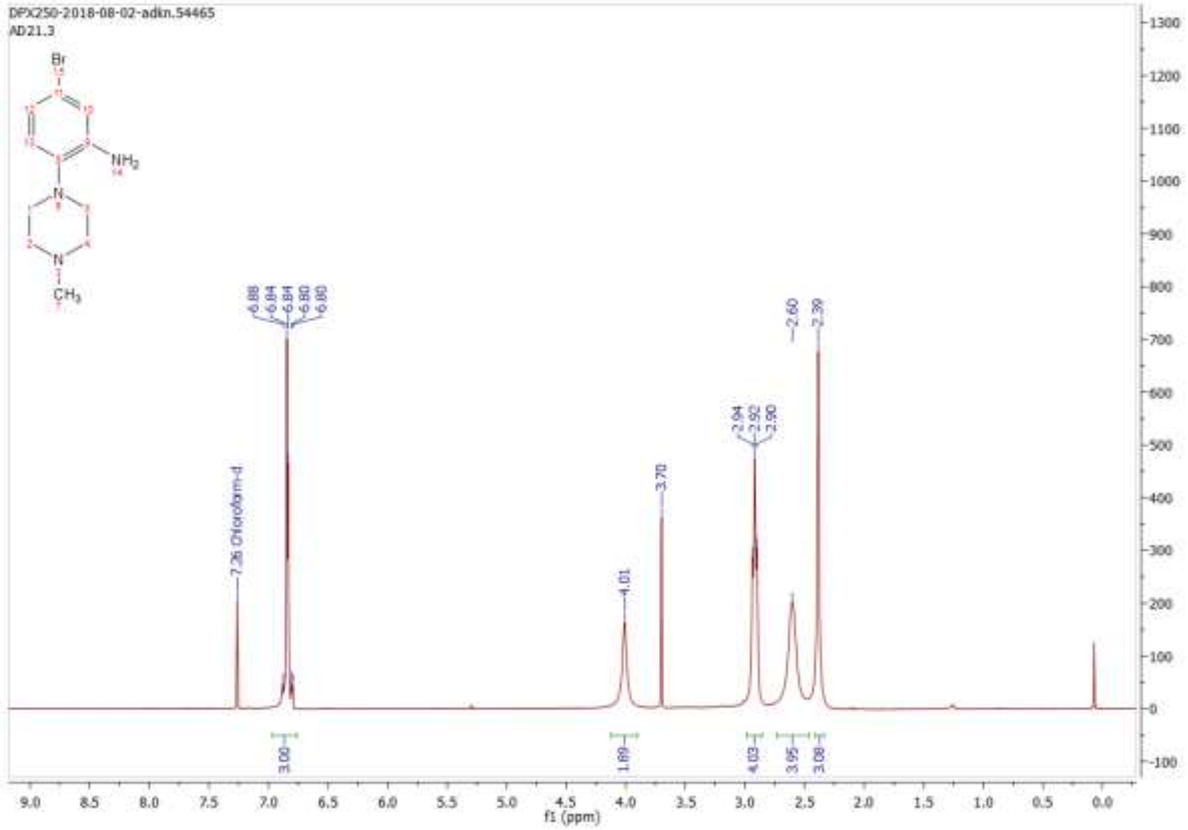

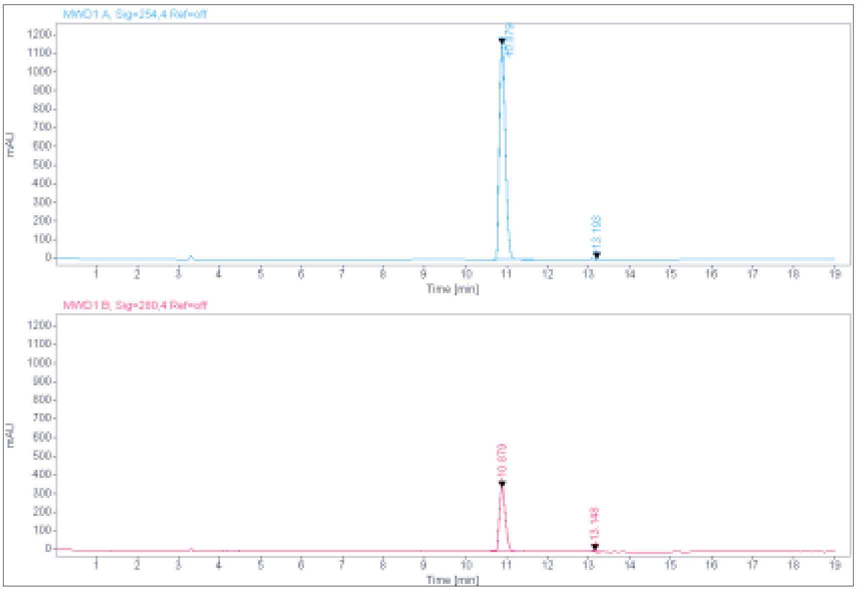

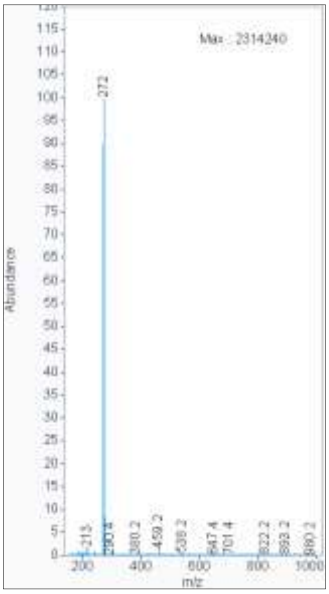

**Signal:** MWD1 A, Sig=254,4 Ref=off

| RT [min] | Type | Width [min] | Area | Height | Area% | Name |
|---|---|---|---|---|---|---|
| 10.879 | BV | 0.1668 | 11862.8955 | 1155.4844 | 99.6042 | |
| 13.193 | MM | 0.0751 | 47.1392 | 10.4557 | 0.3958 | |
| | | Sum | 11910.0347 | | | |

**Signal:** MWD1 B, Sig=280,4 Ref=off

| RT [min] | Type | Width [min] | Area | Height | Area% | Name |
|---|---|---|---|---|---|---|
| 10.879 | VV | 0.1677 | 3558.1433 | 343.9120 | 98.2896 | |
| 13.148 | MM | 0.1711 | 61.9159 | 6.0318 | 1.7104 | |
| | | Sum | 3620.0593 | | | |

**ESI, HPLC, $^{1}$H-NMR and $^{13}$C-NMR of tert-butyl 3'-amino-4'-(4-methylpiperazin-1-yl)-[1,1'-biphenyl]-4-carboxylate 5**

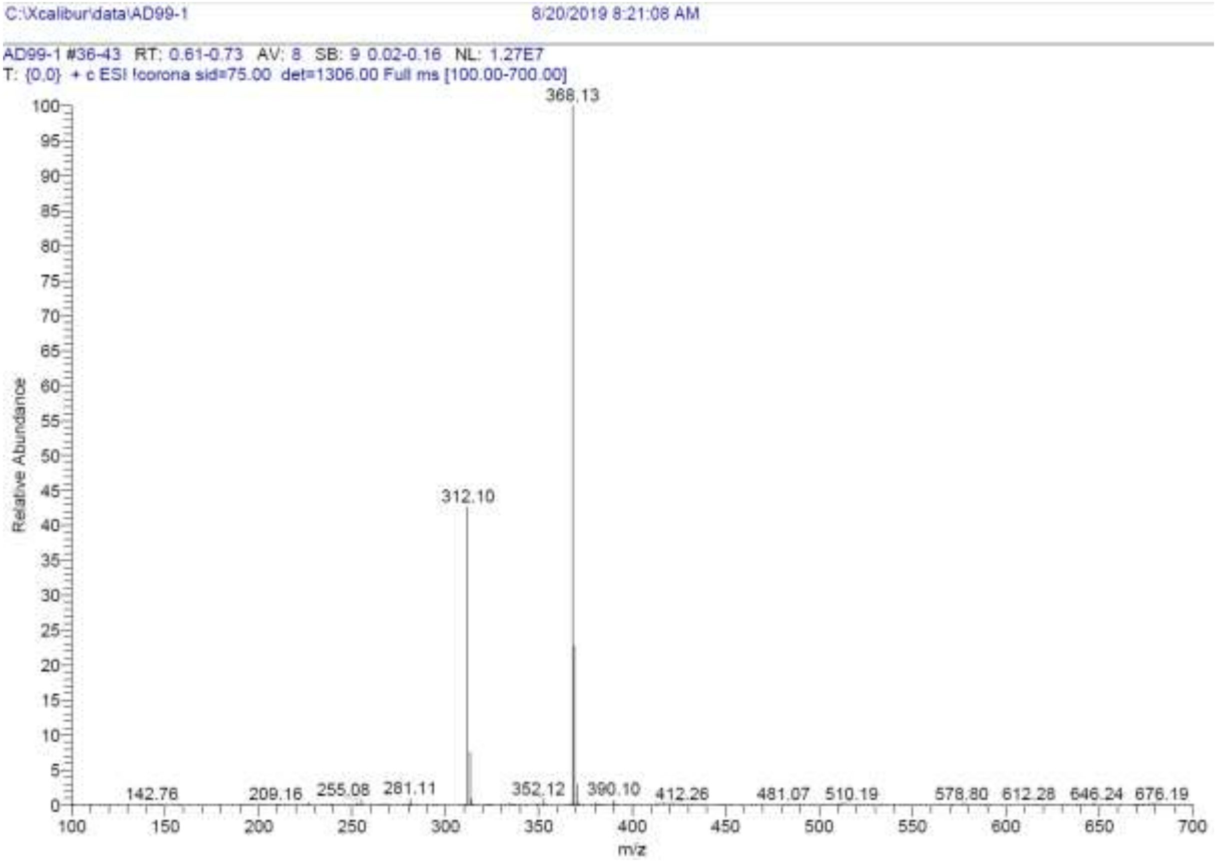

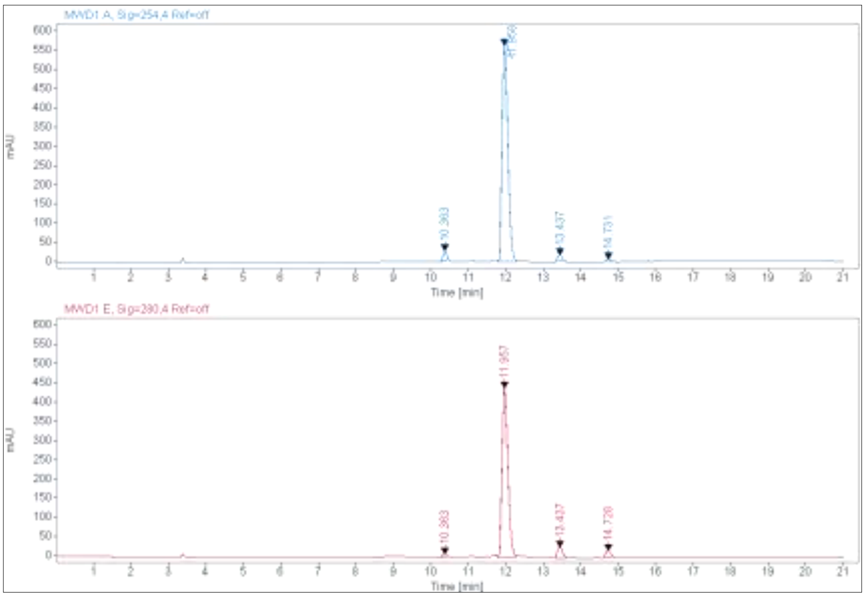

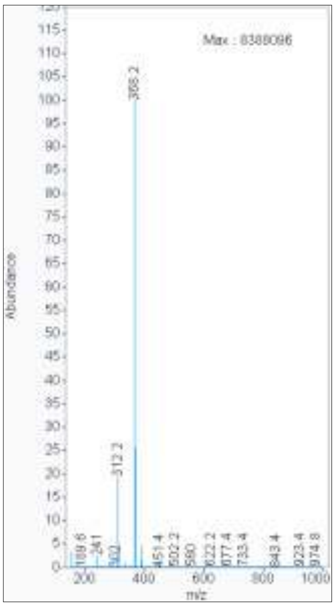

**Signal:** MWD1 A, Sig=254,4 Ref=off

| RT [min] | Type | Width [min] | Area | Height | Area% | Name |
|---|---|---|---|---|---|---|
| 10.363 | MM | 0.1143 | 179.2153 | 26.1271 | 2.7597 | |
| 11.958 | VV | 0.1658 | 6113.2773 | 562.4454 | 94.1364 | |
| 13.437 | MM | 0.1422 | 136.3742 | 15.9810 | 2.1000 | |
| 14.731 | MM | 0.1515 | 65.1996 | 7.1723 | 1.0040 | |
| | | Sum | 6494.0666 | | | |

**Signal:** MWD1 E, Sig=280,4 Ref=off

| RT [min] | Type | Width [min] | Area | Height | Area% | Name |
|---|---|---|---|---|---|---|
| 10.363 | MM | 0.1024 | 45.8921 | 7.4716 | 0.8672 | |
| 11.957 | VV | 0.1663 | 4796.4849 | 439.7085 | 90.6417 | |
| 13.437 | MM | 0.1683 | 301.3157 | 29.8367 | 5.6941 | |
| 14.728 | MM | 0.1462 | 148.0057 | 16.8705 | 2.7969 | |

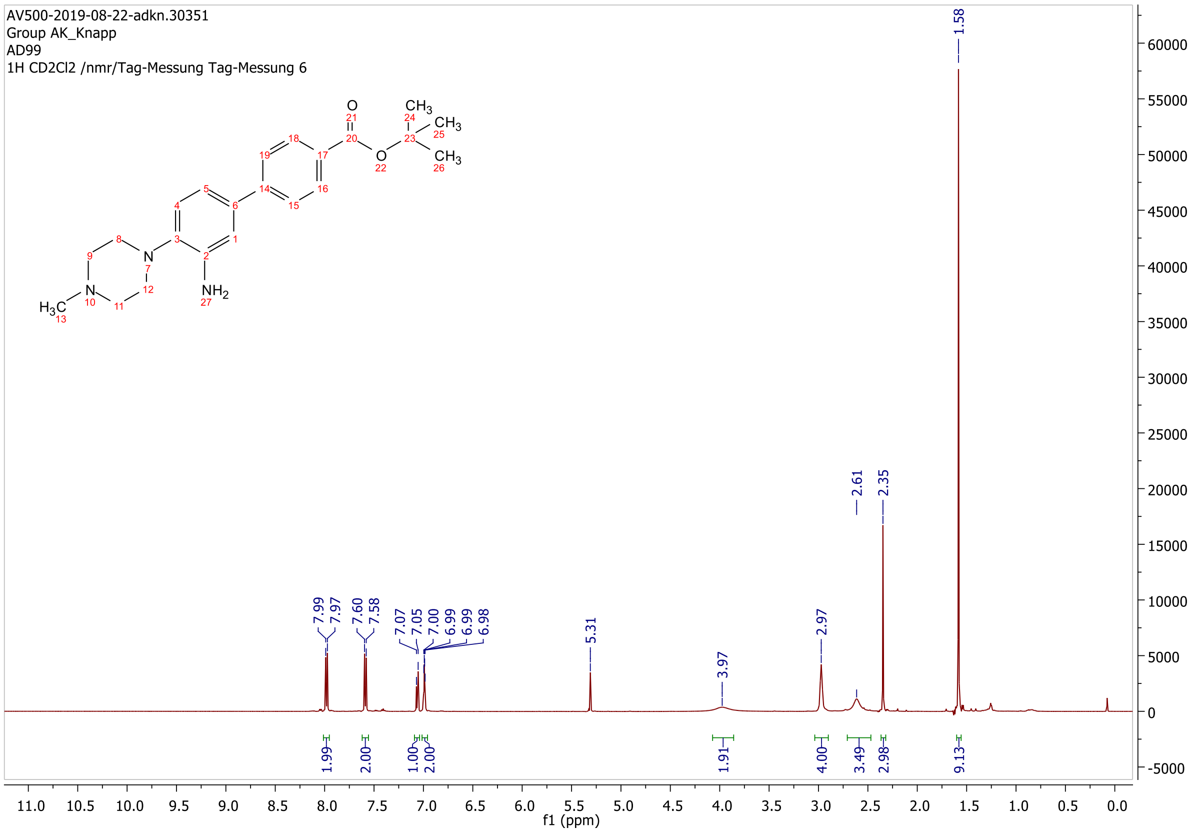

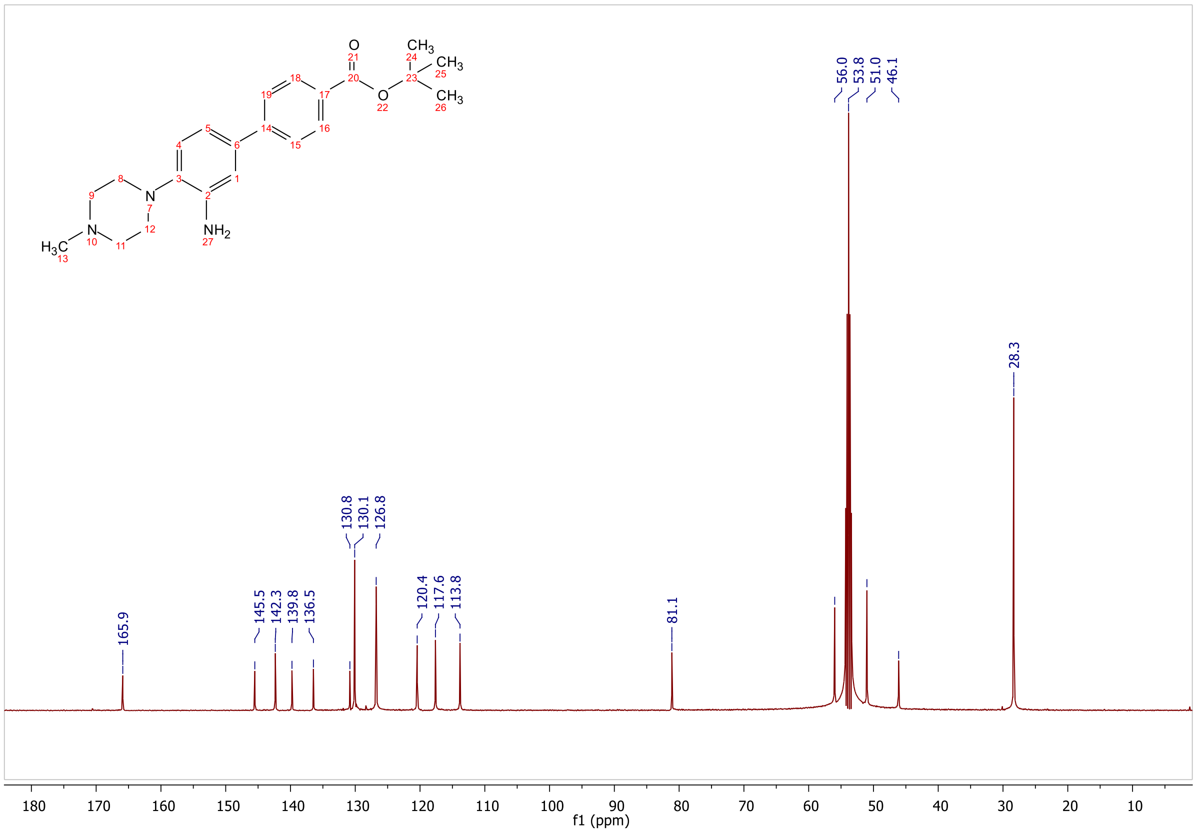
165.9
145.5
142.3
139.8
136.5
130.8
130.1
126.8
120.4
117.6
113.8
81.1
56.0
53.8
51.0
46.1
28.3
f1 (ppm)

**ESI, HPLC, $^{1}$H-NMR and $^{13}$C-NMR of tert-butyl 3'-(6-hydroxy-4-(trifluoromethyl)nicotinamido)-4'-(4-methylpiperazin-1-yl)-[1,1'-biphenyl]-4-carboxylate 6**

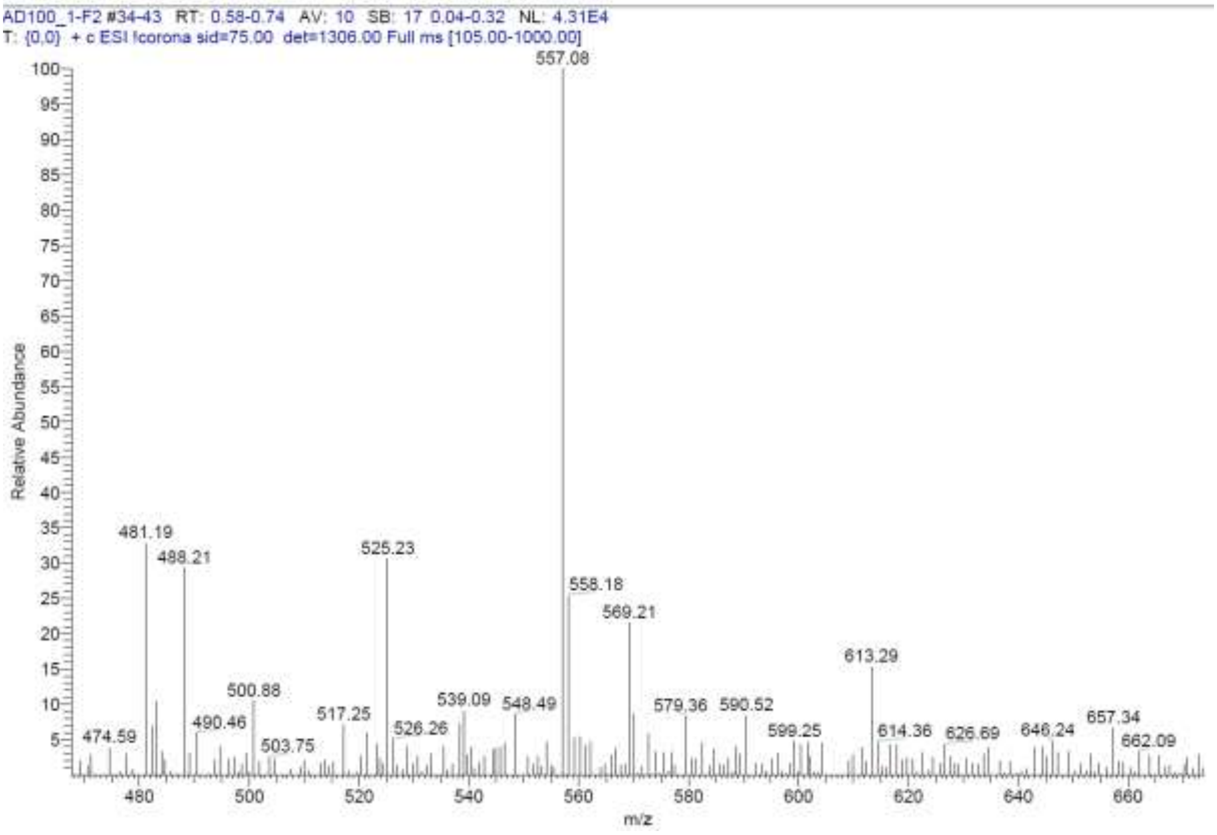

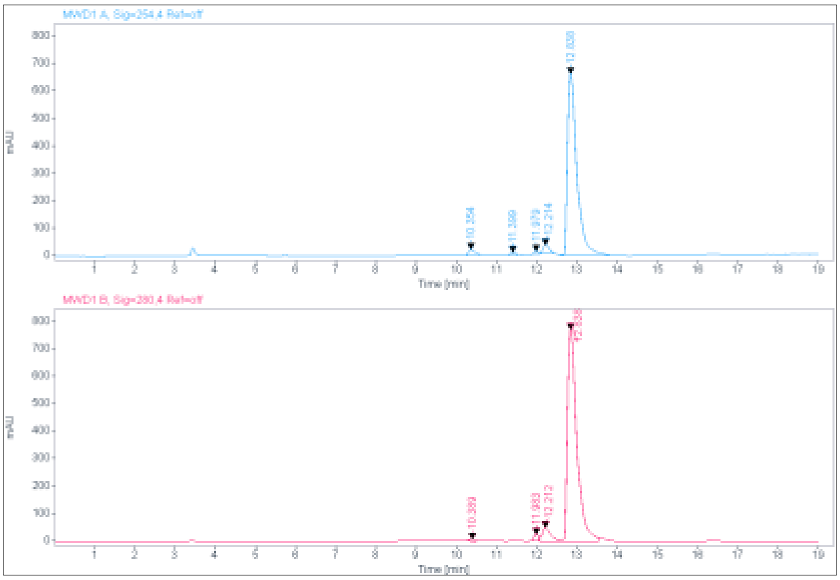

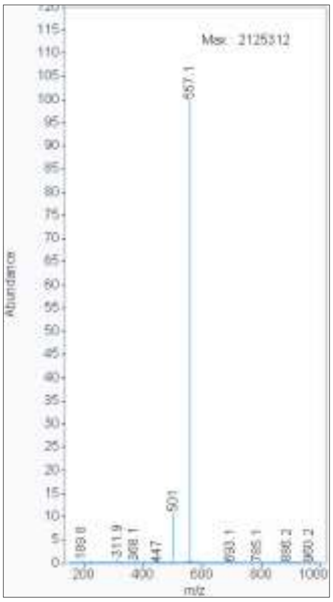

**Signal:** MWD1 A, Sig=254,4 Ref=off

| RT [min] | Type | Width [min] | Area | Height | Area% | Name |
|---|---|---|---|---|---|---|
| 10.354 | MM | 0.1955 | 264.3400 | 22.5354 | 2.2644 | |
| 11.399 | MM | 0.1602 | 109.3309 | 11.3760 | 0.9366 | |
| 11.979 | MM | 0.1438 | 97.5140 | 11.2982 | 0.8353 | |
| 12.214 | MM | 0.1826 | 324.7970 | 29.6518 | 2.7823 | |
| 12.838 | VV | 0.2499 | 10877.6895 | 661.4399 | 93.1814 | |

| | | Sum | | 11673.6714 |
|---|---|---|---|---|

**Signal:** MWD1 B, Sig=280,4 Ref=off

| RT [min] | Type | Width [min] | Area | Height | Area% | Name |
|---|---|---|---|---|---|---|
| 10.389 | MM | 0.1813 | 84.4925 | 7.7665 | 0.6195 | |
| 11.983 | MM | 0.1406 | 191.5884 | 22.7102 | 1.4046 | |
| 12.212 | VV | 0.2312 | 748.0756 | 50.1011 | 5.4845 | |
| 12.838 | VV | 0.2487 | 12615.6523 | 771.6691 | 92.4914 | |

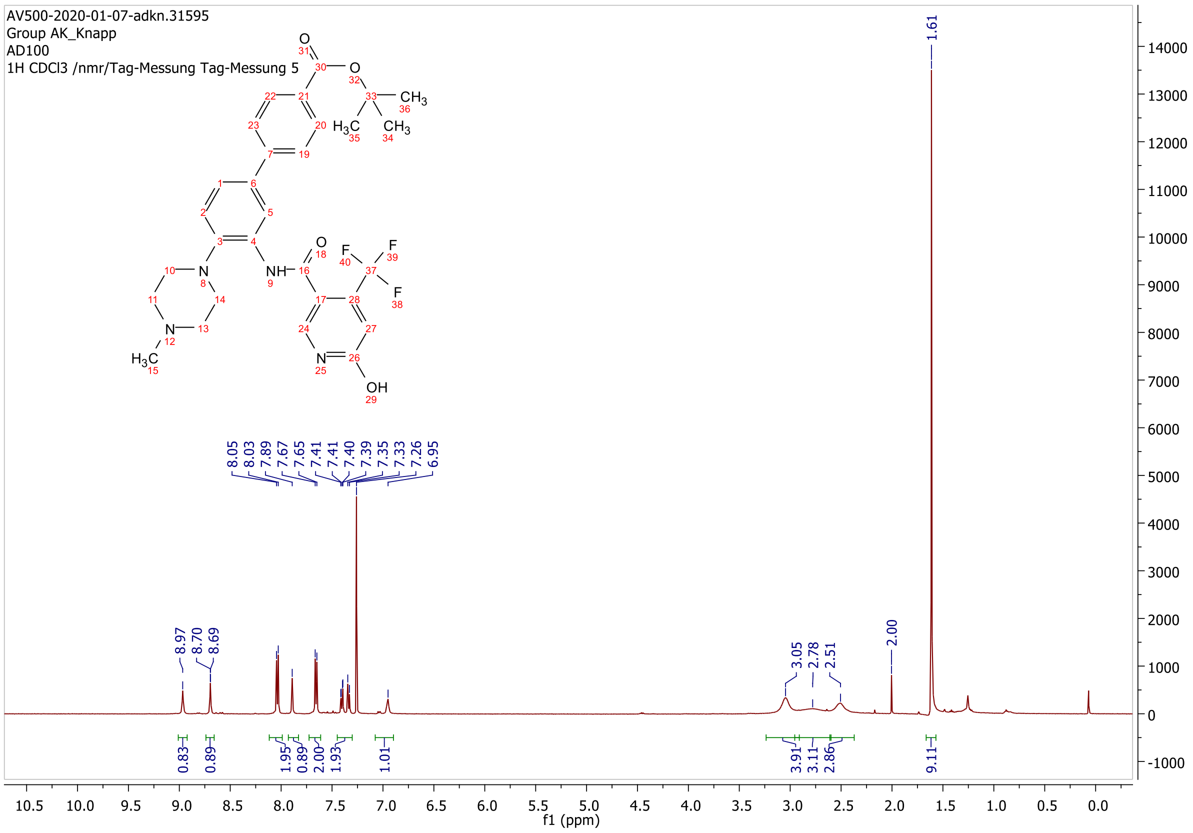

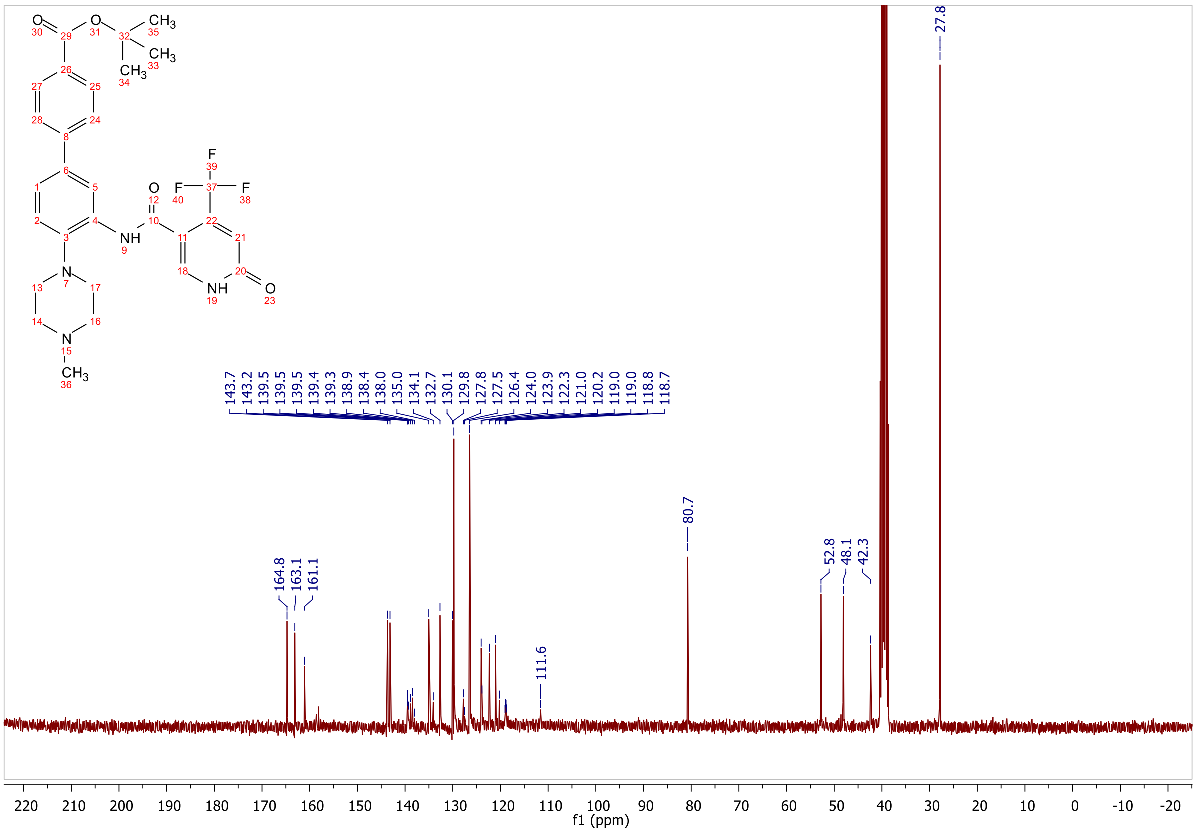
164.8
163.1
161.1
143.7
143.2
139.5
139.5
139.4
139.3
138.9
138.4
138.0
135.0
134.1
132.7
130.1
129.8
127.8
127.5
126.4
124.0
123.9
122.3
121.0
120.2
119.0
118.8
118.7
111.6
80.7
52.8
48.1
42.3
27.8
f1 (ppm)

#### 1.4 Appendix for heterobifunctional molecules 7a-e, 8a-j, 9a-c

**MALDI, HRMS, HPLC, $^{1}$H-NMR, $^{13}$C-NMR, $^{13}$C-NMR (downfield region), $^{13}$C-DEPT90, $^{1}$H-$^{13}$C-HSQC and $^{1}$H-$^{13}$C-HMBC of *N*-(4'-((2-(2-((2-(2,6-dioxopiperidin-3-yl)-1,3-dioxoisoindolin-4-yl)amino)ethoxy)ethyl)carbamoyl)-4-(4-methylpiperazin-1-yl)-[1,1'-biphenyl]-3-yl)-6-hydroxy-4-(trifluoromethyl)nicotinamide 7a**

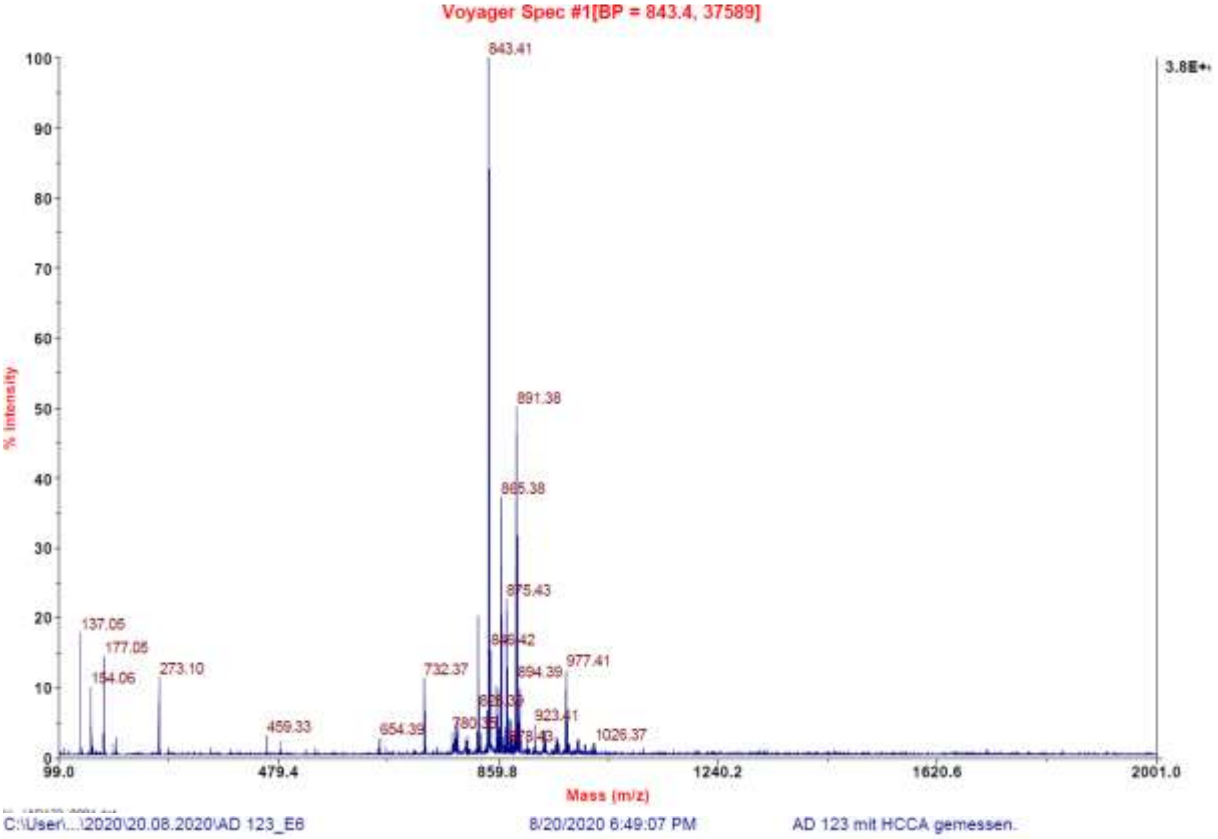

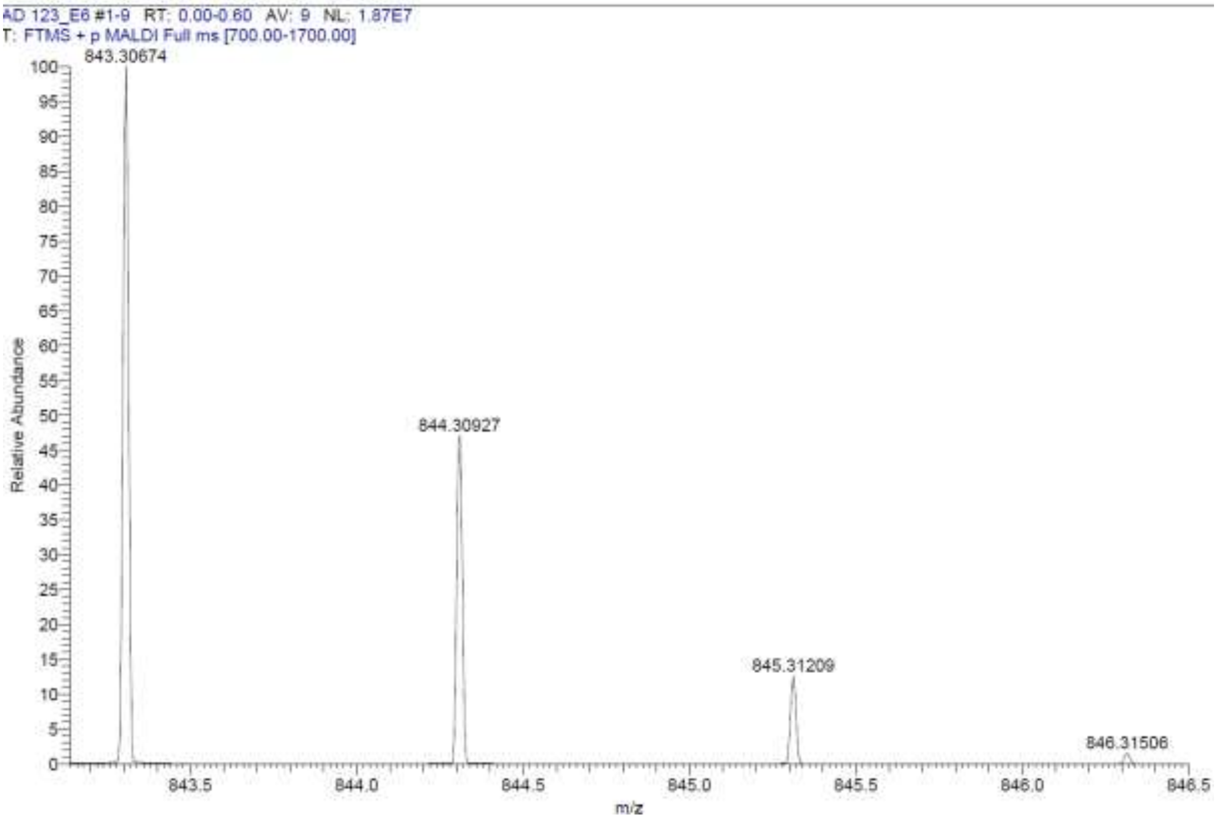

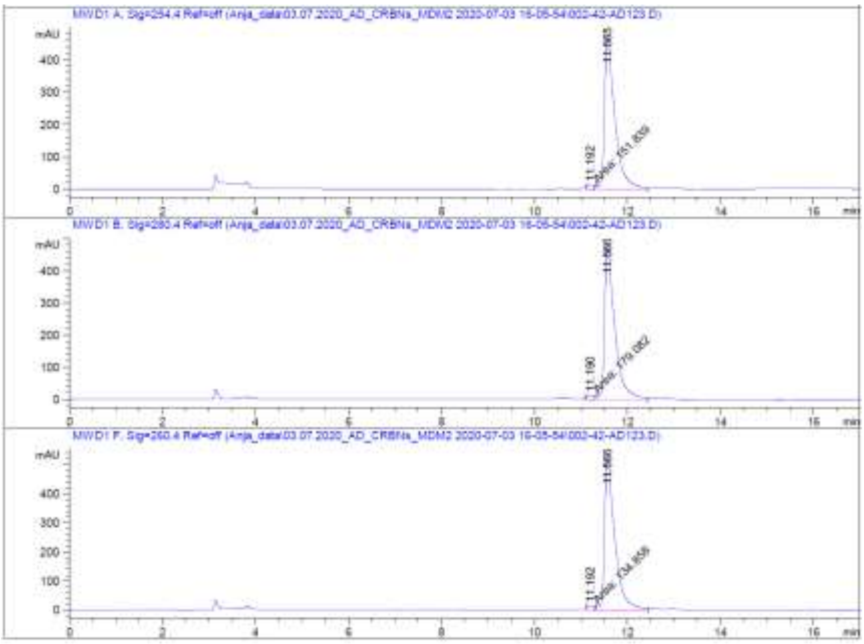

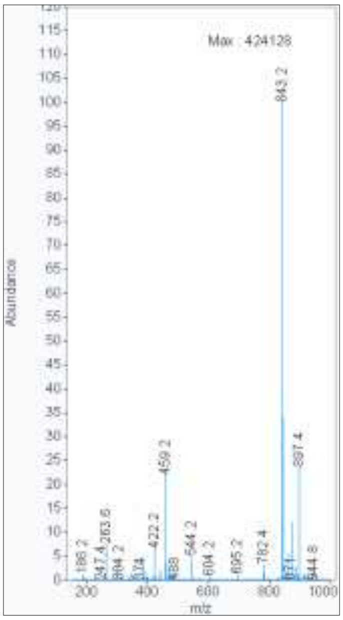

**Signal:** MWD1 A, Sig=254,4 Ref=off

| RT [min] | Type | Width [min] | Area | Height | Area% | Name |
|---|---|---|---|---|---|---|
| 11.614 | MM | 0.2199 | 202.6069 | 15.3539 | 100.0000 | |
| | | Sum | 202.6069 | | | |

**Signal:** MWD1 B, Sig=280,4 Ref=off

| RT [min] | Type | Width [min] | Area | Height | Area% | Name |
|---|---|---|---|---|---|---|
| 11.612 | MM | 0.2239 | 212.1426 | 15.7890 | 100.0000 | |
| | | Sum | 212.1426 | | | |

**Signal:** MWD1 F, Sig=260,4 Ref=off

| RT [min] | Type | Width [min] | Area | Height | Area% | Name |
|---|---|---|---|---|---|---|
| 11.612 | MM | 0.2276 | 238.7010 | 17.4816 | 100.0000 | |

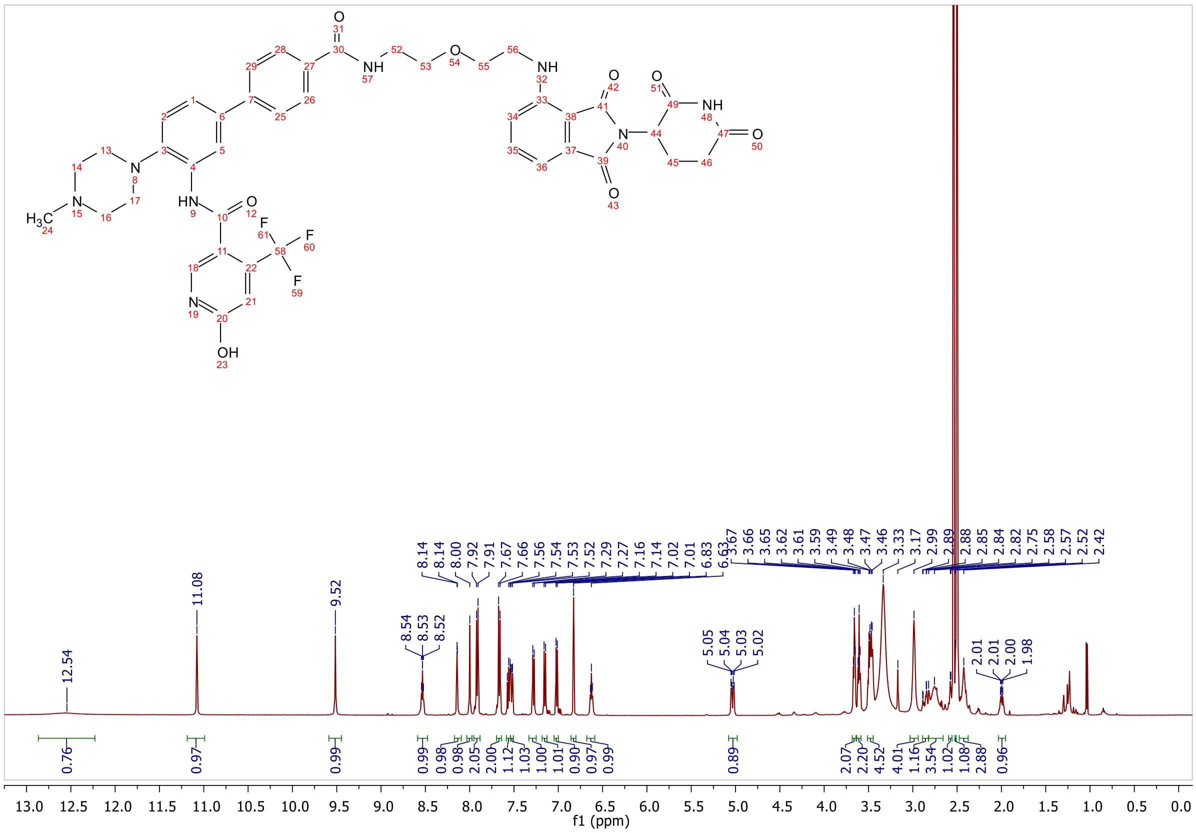

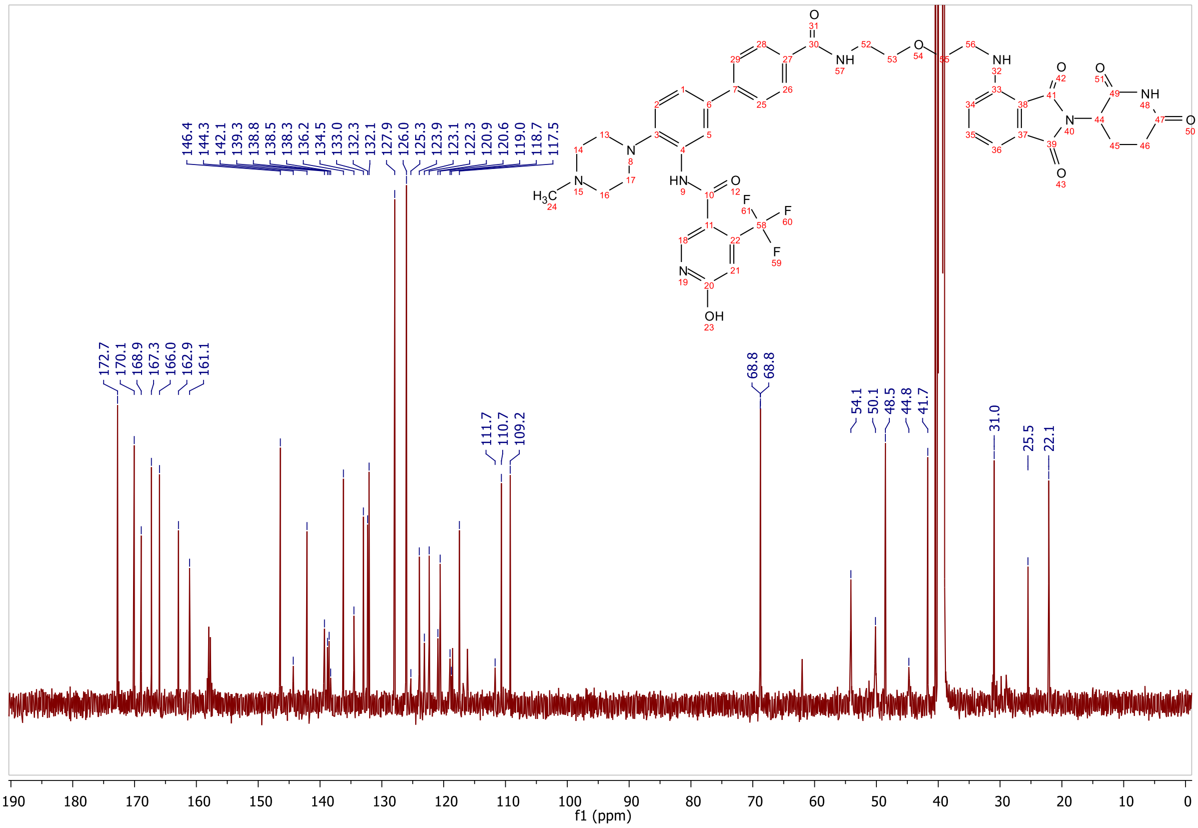

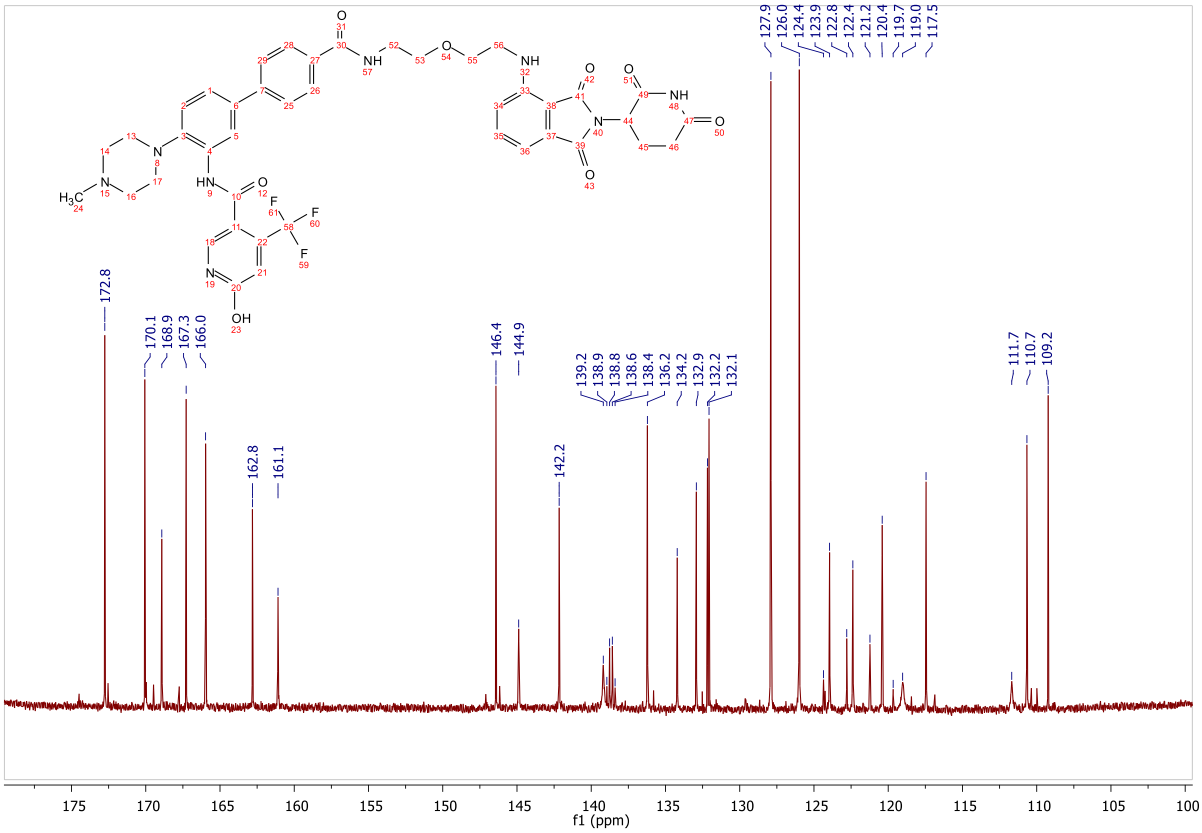

protac_700
compound #123 ~?mg in 500ul DMSO-d6 T=298K
13C DEPT90 (only CH groups detected) NS=1k Aq=1.5s D1=1.5s protac_700/63 17.3.2021
13C carrier at 140ppm, SW=80ppm, de=50us

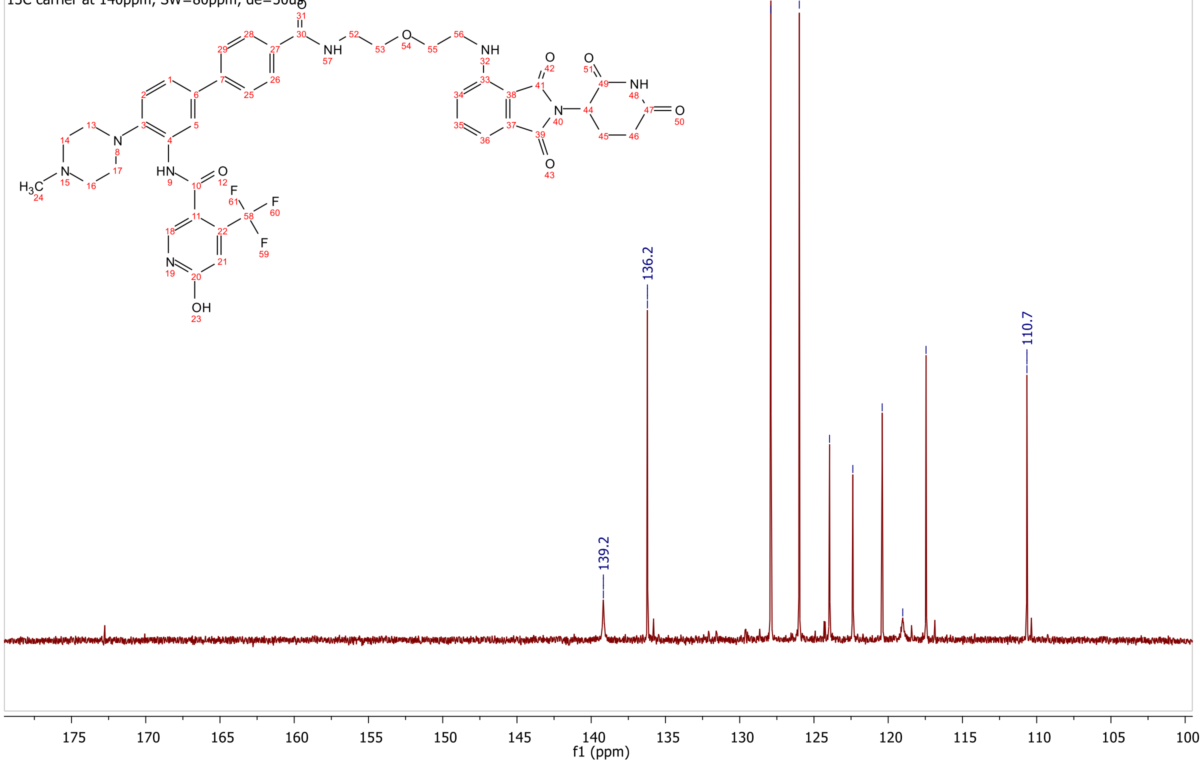

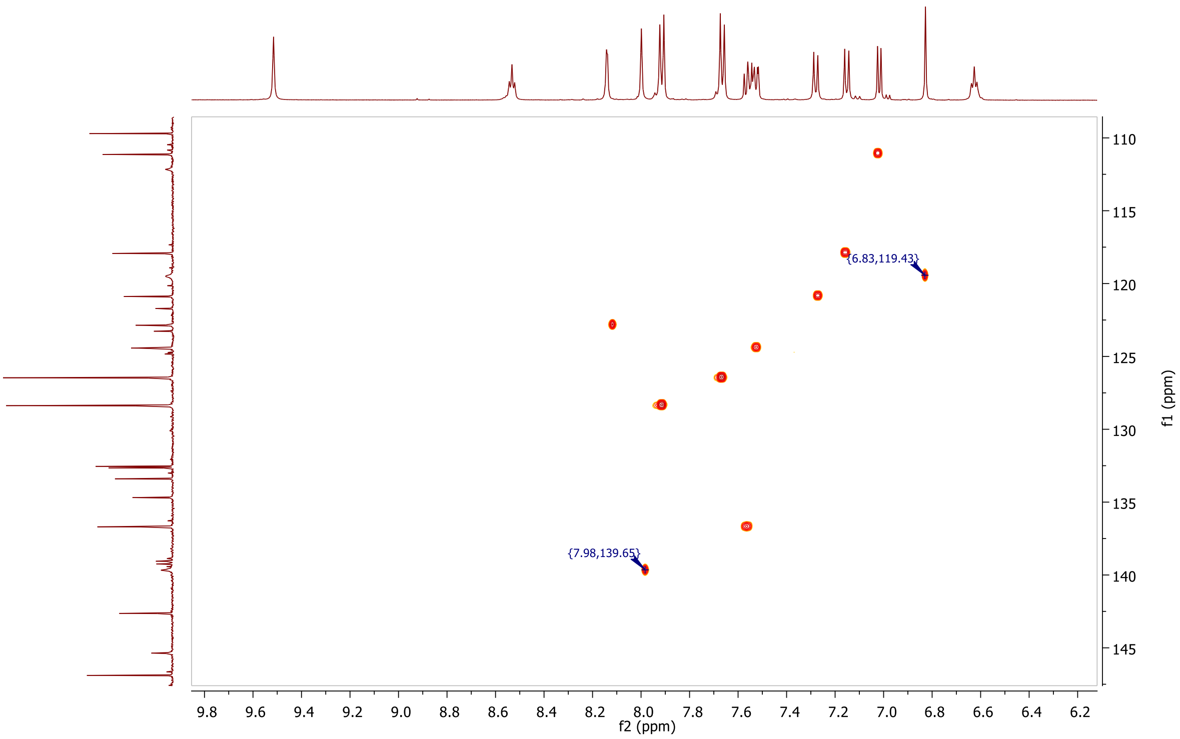
{6.83,119.43}
{7.98,139.65}
f2 (ppm)
f1 (ppm)

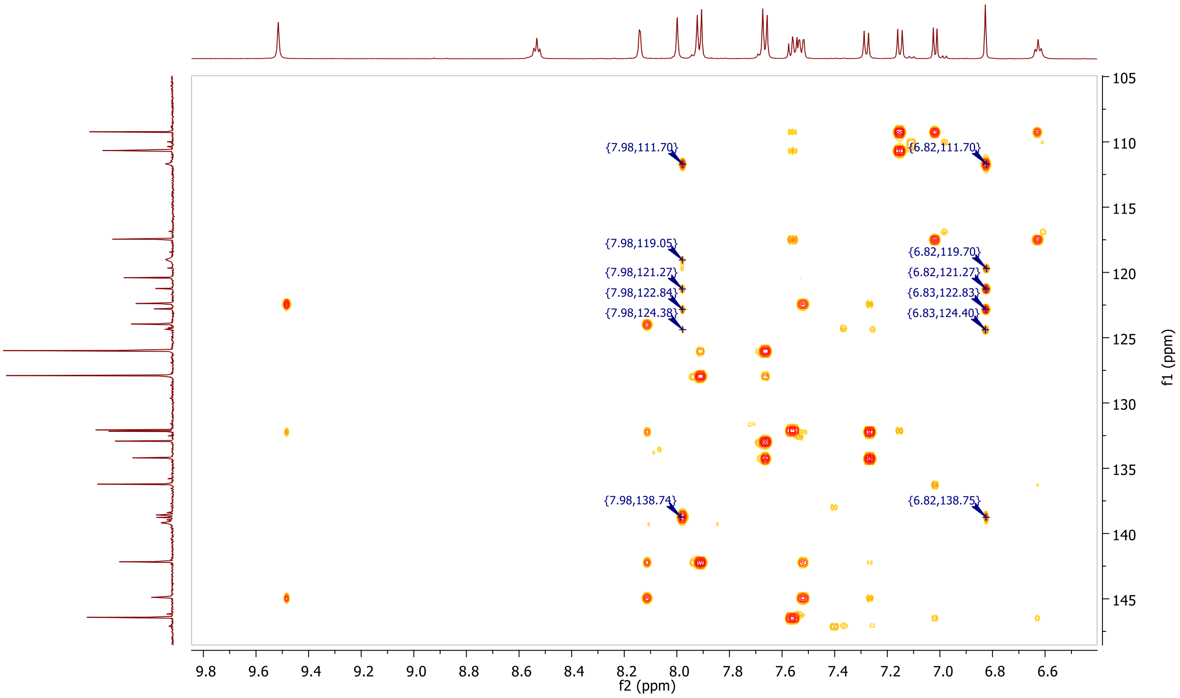
{7.98,111.70}
{6.82,111.70}
{7.98,119.05}
{6.82,119.70}
{7.98,121.27}
{6.82,121.27}
{7.98,122.84}
{6.83,122.83}
{7.98,124.38}
{6.83,124.40}
{7.98,138.74}
{6.82,138.75}
f2 (ppm)
f1 (ppm)

**HPLC, MALDI, HRMS, $^{1}$H-NMR, $^{13}$C-NMR, $^{13}$C-NMR (downfield region), $^{1}$H-$^{13}$C-HSQC and $^{1}$H-$^{13}$C-HMBC of *N*-(4'-((2-(2-(2-(2-((2-(2,6-dioxopiperidin-3-yl)-1,3-dioxoisoindolin-4-yl)amino)ethoxy)ethoxy)ethoxy)ethyl)carbamoyl)-4-(4-methylpiperazin-1-yl)-[1,1'-biphenyl]-3-yl)-6-hydroxy-4-(trifluoromethyl)nicotinamide 7b**

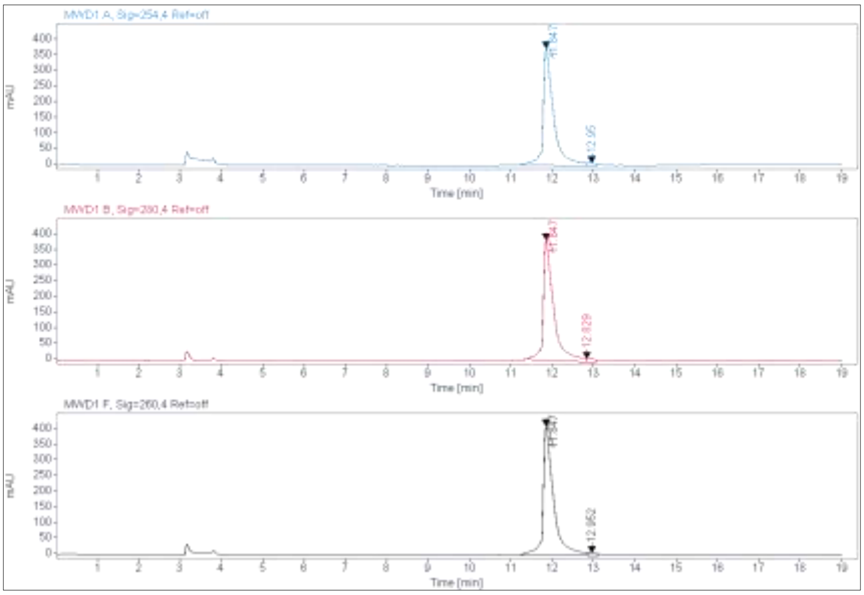

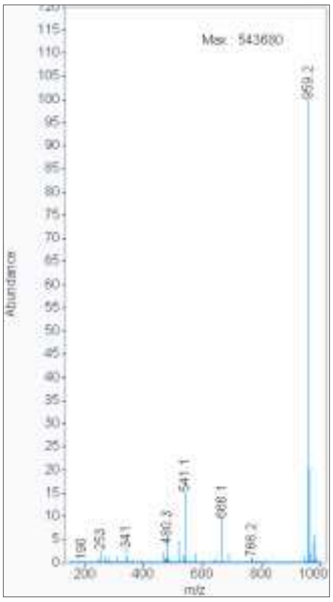

**Signal:** MWD1 A, Sig=254,4 Ref=off

| RT [min] | Type | Width [min] | Area | Height | Area% |
|---|---|---|---|---|---|
| 11.847 | MM | 0.2960 | 6658.2402 | 374.9590 | 97.7984 |
| 12.950 | MM | 0.2326 | 149.8847 | 10.7386 | 2.2016 |
| | | Sum | 6808.1249 | | |

**Signal:** MWD1 B, Sig=280,4 Ref=off

| RT [min] | Type | Width [min] | Area | Height | Area% |
|---|---|---|---|---|---|
| 11.847 | MM | 0.2924 | 6744.6553 | 384.4801 | 98.0727 |
| 12.829 | MM | 0.1510 | 132.5450 | 10.4535 | 1.9273 |
| | | Sum | 6877.2003 | | |

**Signal:** MWD1 F, Sig=260,4 Ref=off

| RT [min] | Type | Width [min] | Area | Height | Area% |
|---|---|---|---|---|---|
| 11.847 | VV | 0.2405 | 7302.5942 | 412.6076 | 98.0674 |
| 12.952 | MM | 0.2115 | 143.9127 | 11.3433 | 1.9326 |

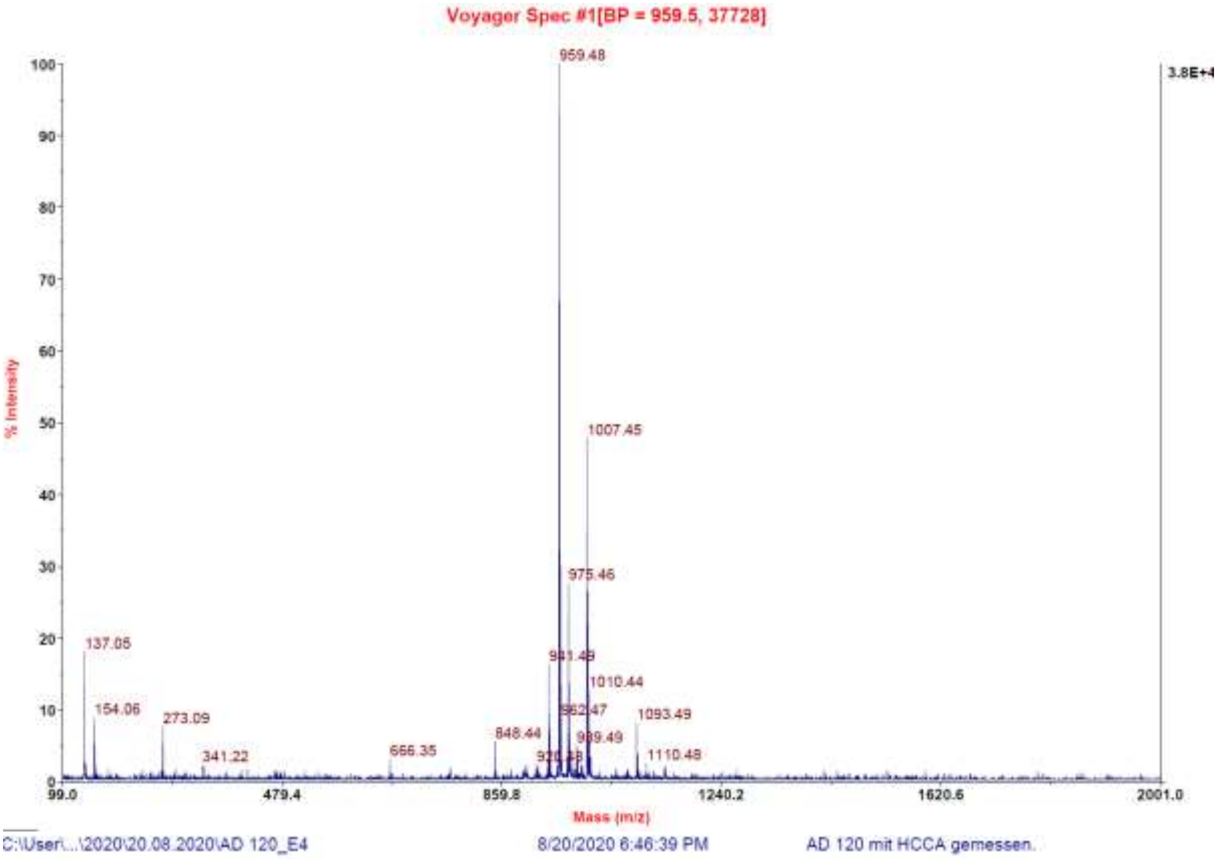
Voyager Spec #1[BP = 959.5, 37728]
% Intensity
Mass (m/z)
959.48
1007.45
975.46
941.48
1010.44
962.47
1093.49
989.49
1110.48
848.44
666.35
137.05
154.06
273.09
341.22
3.8E+4
C:\User\...\2020\20.08.2020\AD 120_E4
8/20/2020 6:46:39 PM
AD 120 mit HCCA gemessen.

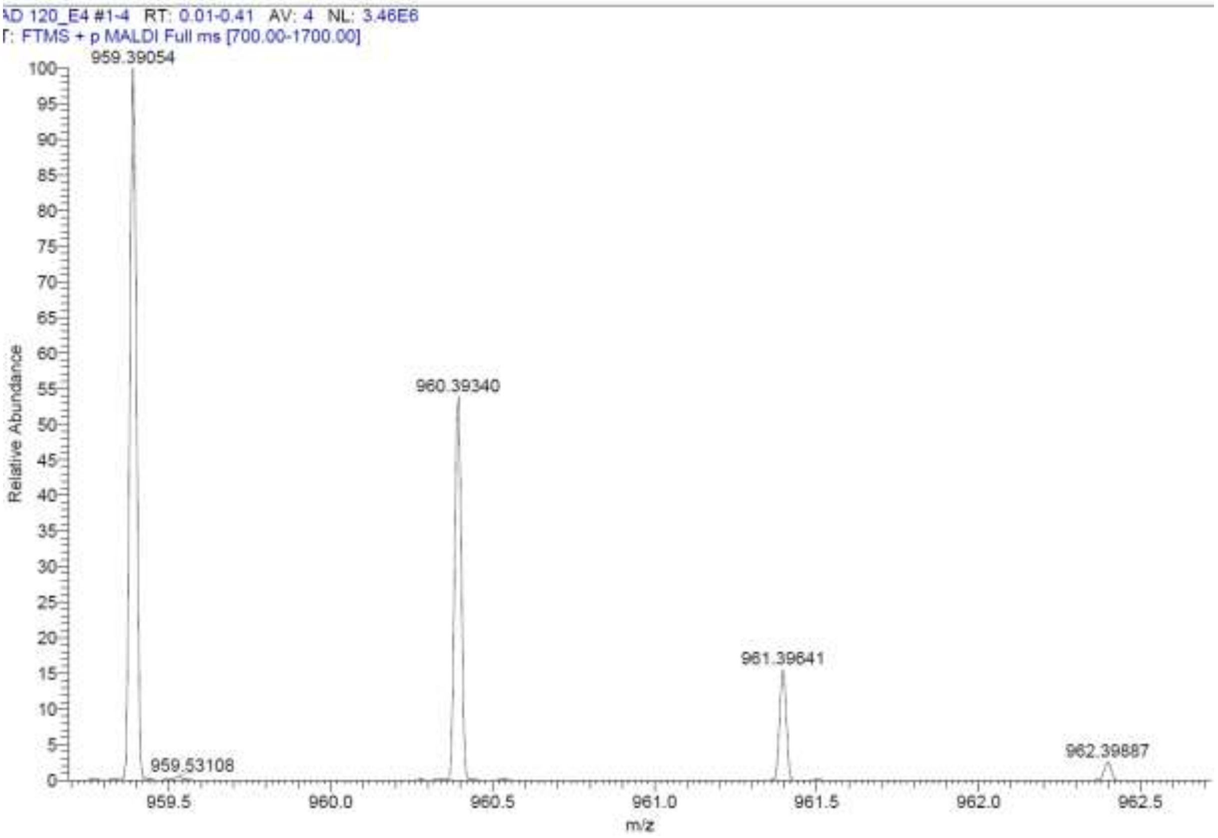
AD 120_E4 #1-4 RT: 0.01-0.41 AV: 4 NL: 3.46E6
T: FTMS + p MALDI Full ms [700.00-1700.00]
Relative Abundance
m/z
959.39054
959.53108
960.39340
961.39641
962.39887

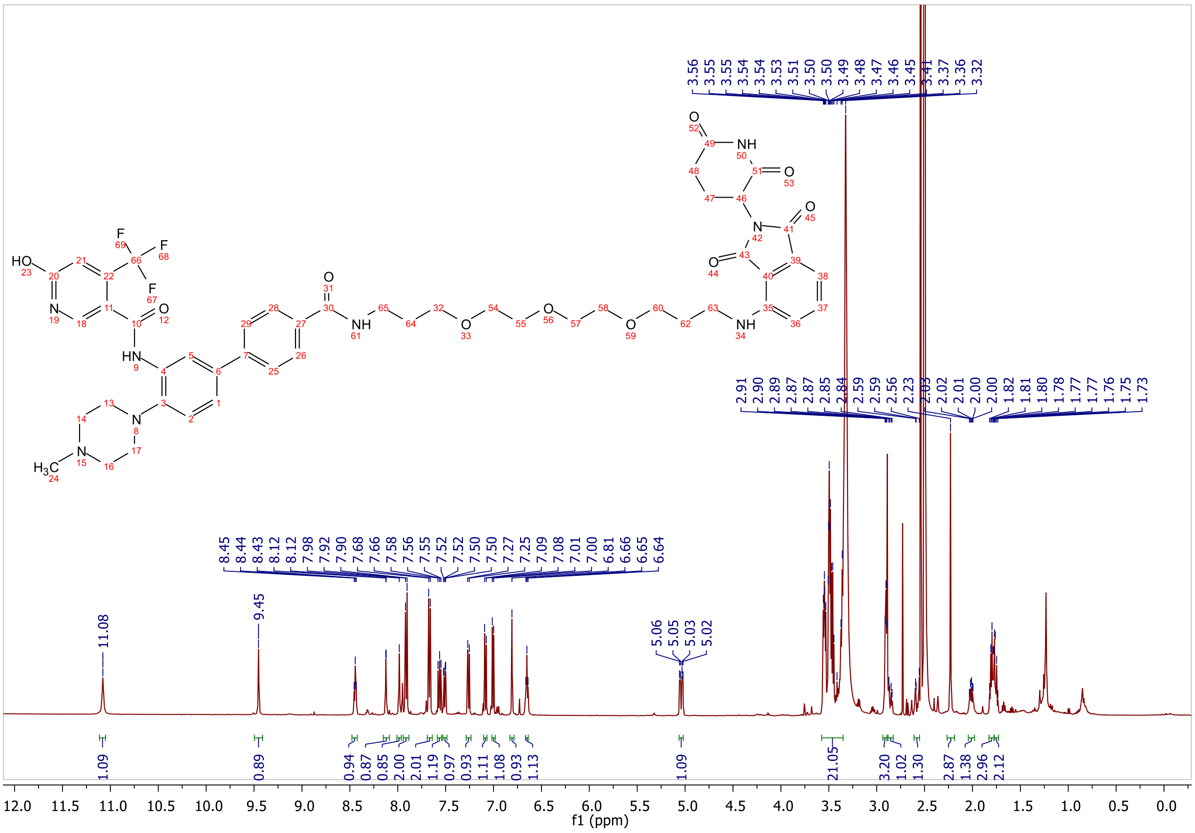
3.56 3.55 3.55 3.54 3.54 3.53 3.51 3.50 3.50 3.49 3.48 3.47 3.46 3.45 3.41 3.37 3.36 3.32
2.91 2.90 2.89 2.87 2.87 2.85 2.84 2.59 2.59 2.56 2.23 2.03 2.02 2.01 2.00 2.00 1.82 1.81 1.80 1.78 1.77 1.77 1.76 1.75 1.73
8.45 8.44 8.43 8.12 8.12 7.98 7.92 7.90 7.68 7.66 7.58 7.56 7.55 7.52 7.52 7.50 7.50 7.27 7.25 7.09 7.08 7.01 7.00 6.81 6.66 6.65 6.64
11.08
9.45
5.06 5.05 5.03 5.02
1.09 0.89 0.94 0.87 0.85 2.00 2.01 1.19 0.97 0.93 1.11 1.08 0.93 1.13 1.09 21.05 3.20 1.02 1.30 2.87 1.38 2.96 2.12
f1 (ppm)

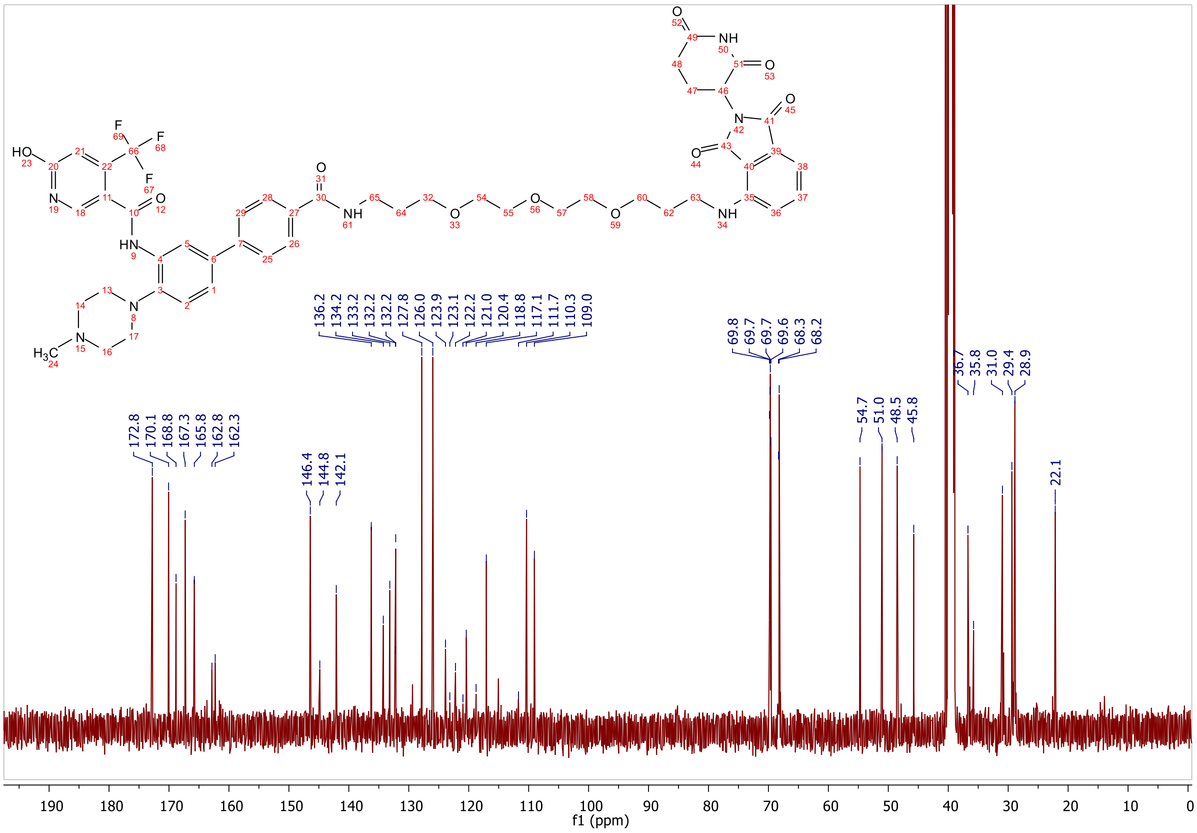
136.2 134.2 133.2 132.2 132.2 127.8 126.0 123.9 123.1 122.2 121.0 120.4 118.8 117.1 111.7 110.3 109.0
69.8 69.7 69.7 69.6 68.3 68.2
172.8 170.1 168.8 167.3 165.8 162.8 162.3
146.4 144.8 142.1
54.7 51.0 48.5 45.8
36.7 35.8 31.0 29.4 28.9
22.1
f1 (ppm)

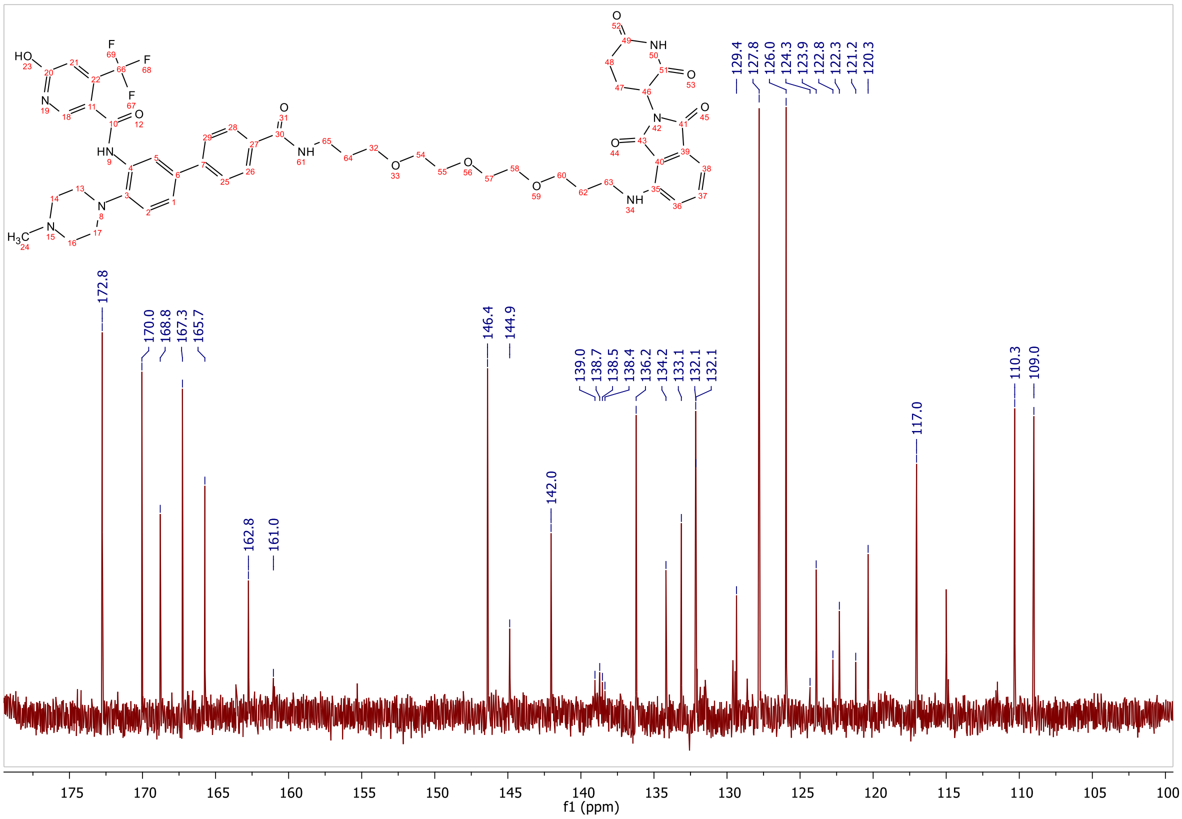
172.8
170.0
168.8
167.3
165.7
162.8
161.0
146.4
144.9
142.0
139.0
138.7
138.5
138.4
136.2
134.2
133.1
132.1
132.1
129.4
127.8
126.0
124.3
123.9
122.8
122.3
121.2
120.3
117.0
110.3
109.0
f1 (ppm)

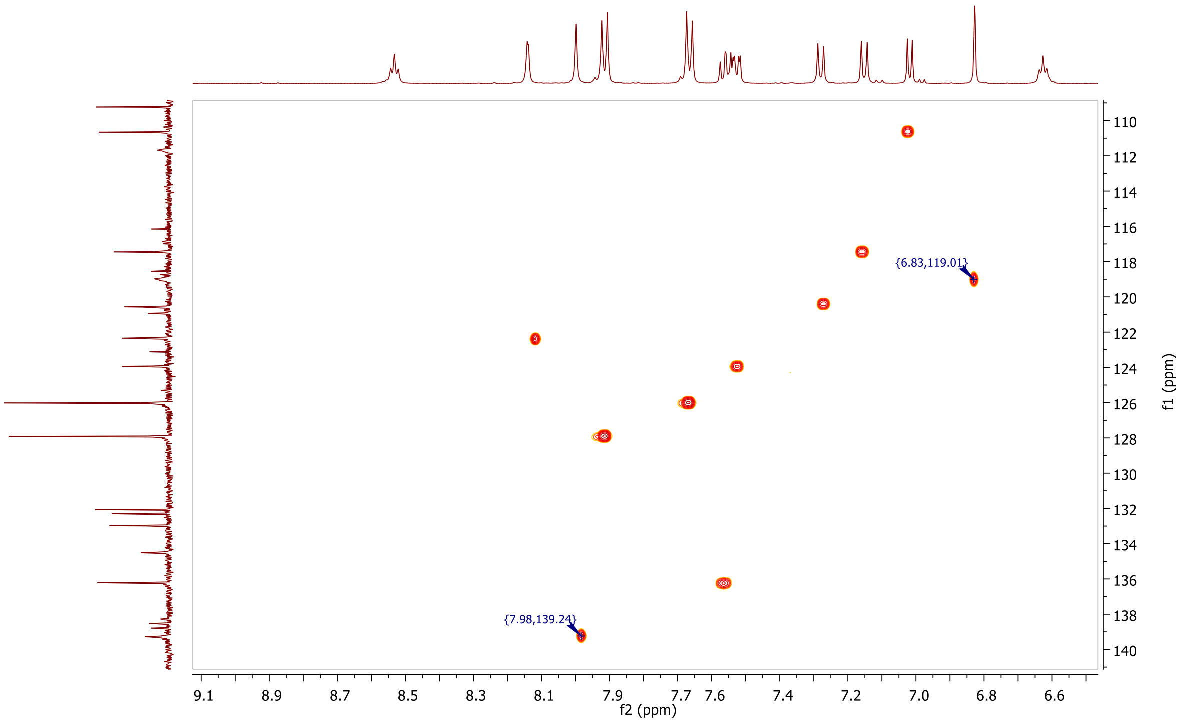
{6.83,119.01}
{7.98,139.24}
f2 (ppm)
f1 (ppm)

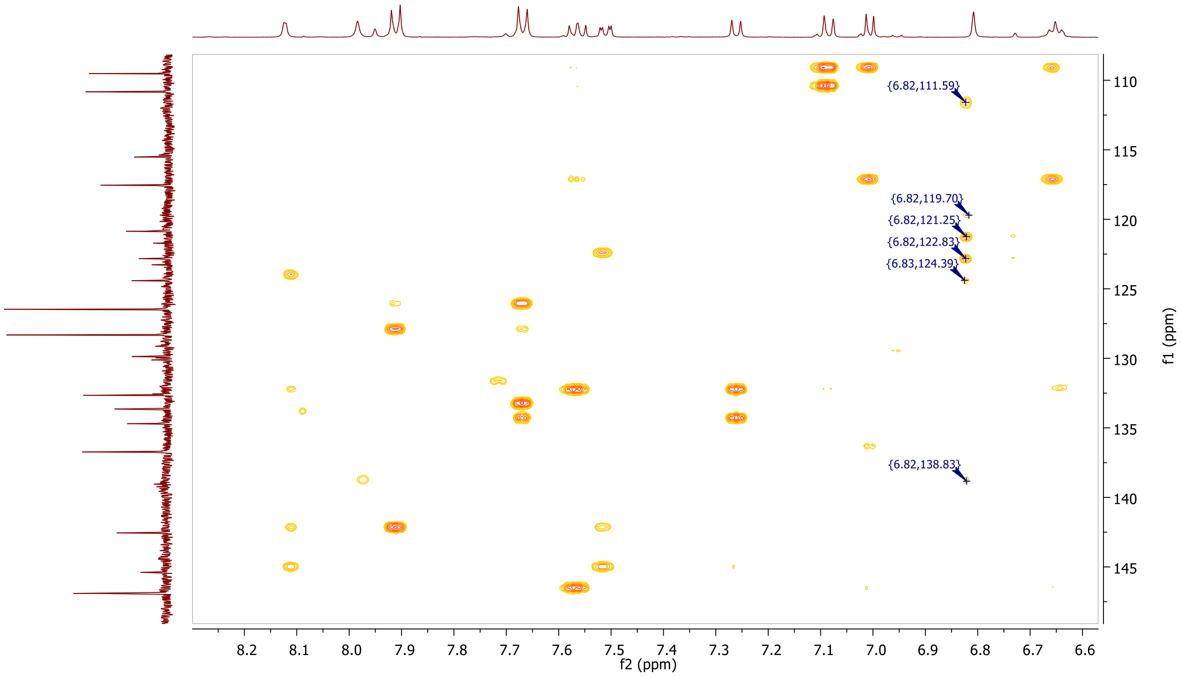
{6.82,111.59}
{6.82,119.70}
{6.82,121.25}
{6.82,122.83}
{6.83,124.39}
{6.82,138.83}
f2 (ppm)
f1 (ppm)

**MALDI, HRMS, HPLC, $^{1}$H-NMR, $^{13}$C-NMR, $^{1}$H-$^{13}$C-HSQC and $^{1}$H-$^{13}$C-HMBC of *N*-(4'-((17-((2-(2,6-dioxopiperidin-3-yl)-1,3-dioxoisoindolin-4-yl)amino)-3,6,9,12,15-pentaoxaheptadecyl)carbamoyl)-4-(4-methylpiperazin-1-yl)-[1,1'-biphenyl]-3-yl)-6-hydroxy-4-(trifluoromethyl)nicotinamide 7c**

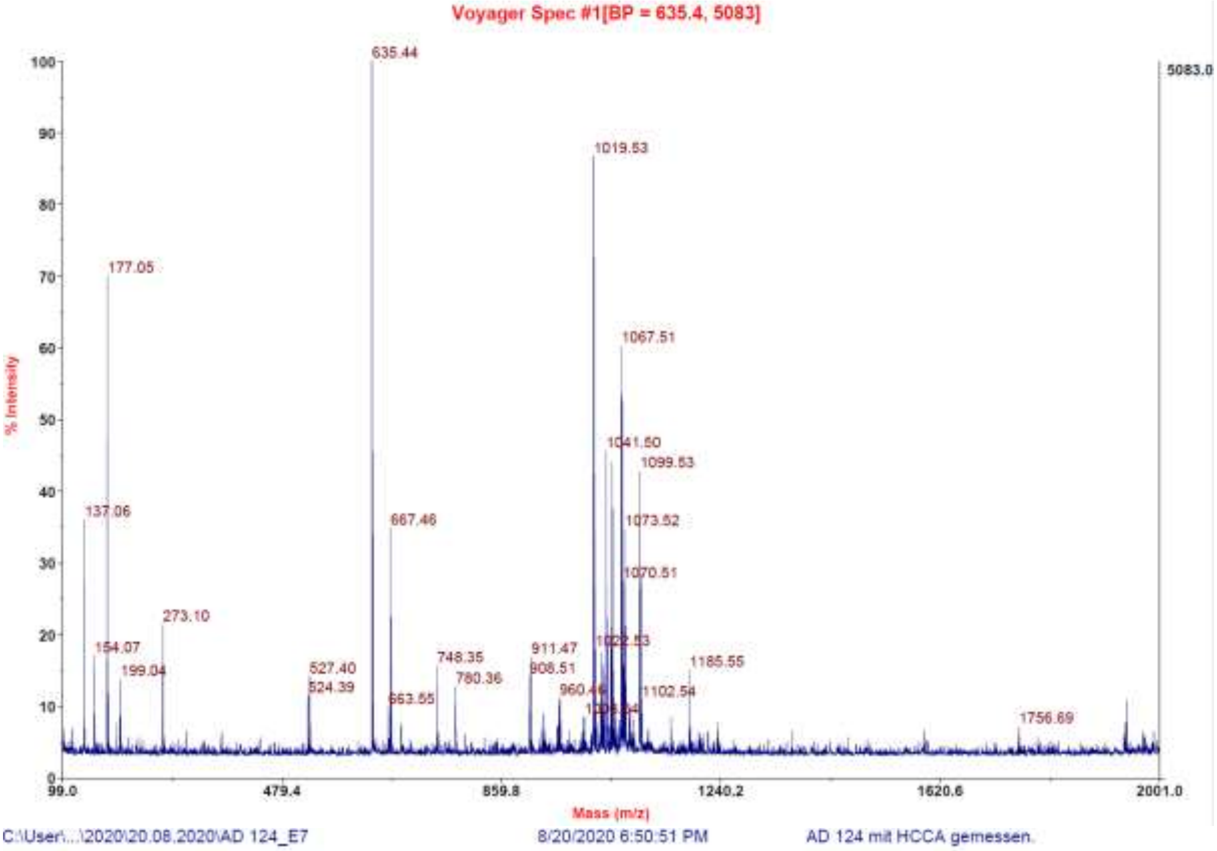

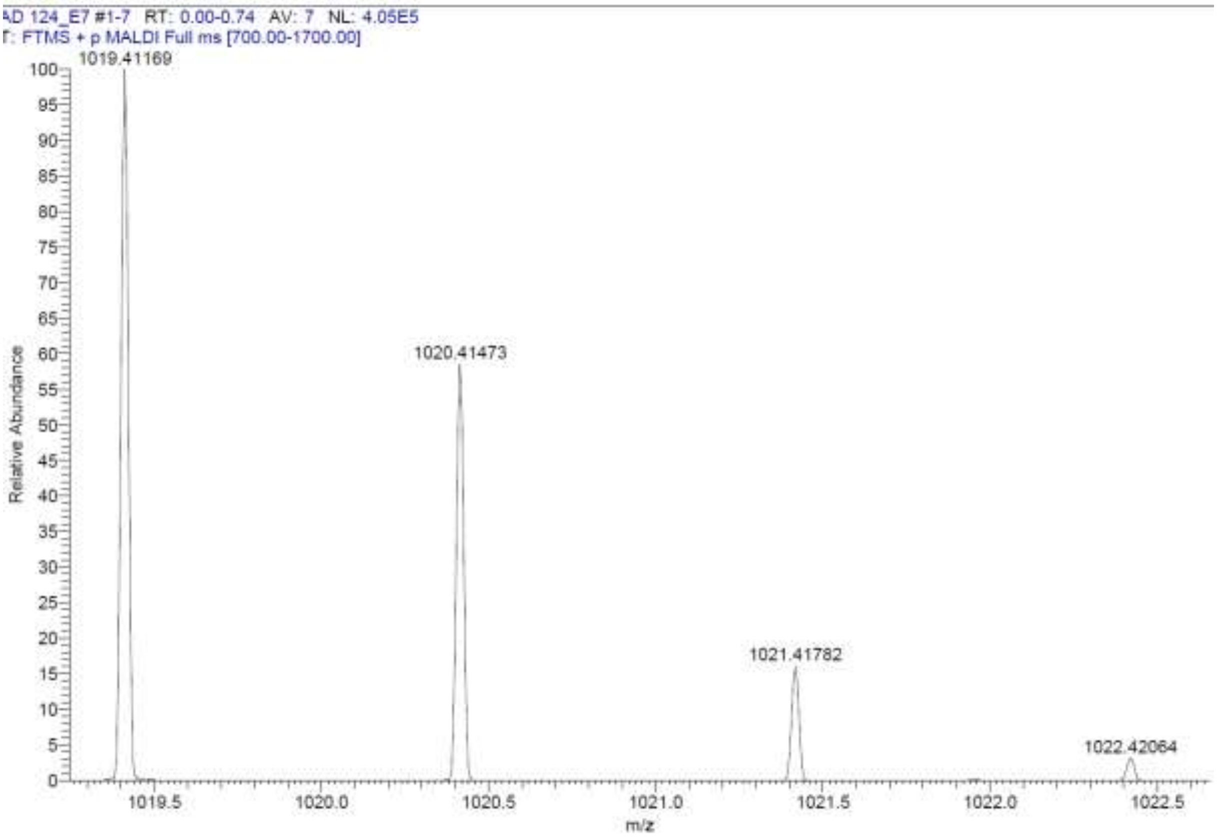

**Signal:** MWD1 A, Sig=254,4 Ref=off

| RT [min] | Type | Width [min] | Area | Height | Area% | Name |
|---|---|---|---|---|---|---|
| 11.593 | VV | 0.2390 | 18262.3613 | 1133.0564 | 100.0000 | |
| | | Sum | 18262.3613 | | | |

**Signal:** MWD1 B, Sig=280,4 Ref=off

| RT [min] | Type | Width [min] | Area | Height | Area% | Name |
|---|---|---|---|---|---|---|
| 11.593 | VV | 0.2389 | 22847.2031 | 1418.2609 | 100.0000 | |
| | | Sum | 22847.2031 | | | |

**Signal:** MWD1 F, Sig=260,4 Ref=off

| RT [min] | Type | Width [min] | Area | Height | Area% | Name |
|---|---|---|---|---|---|---|
| 11.593 | VV | 0.2401 | 19822.1172 | 1229.6807 | 100.0000 | |

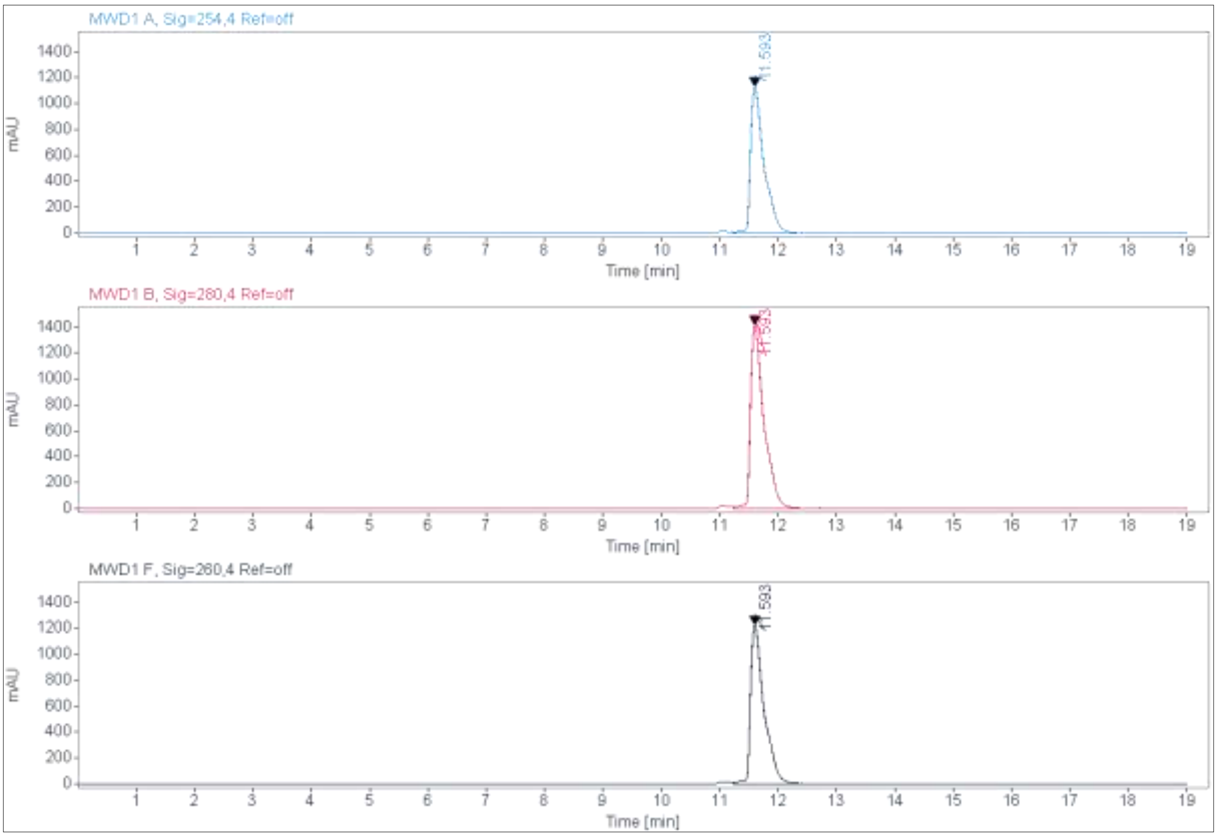

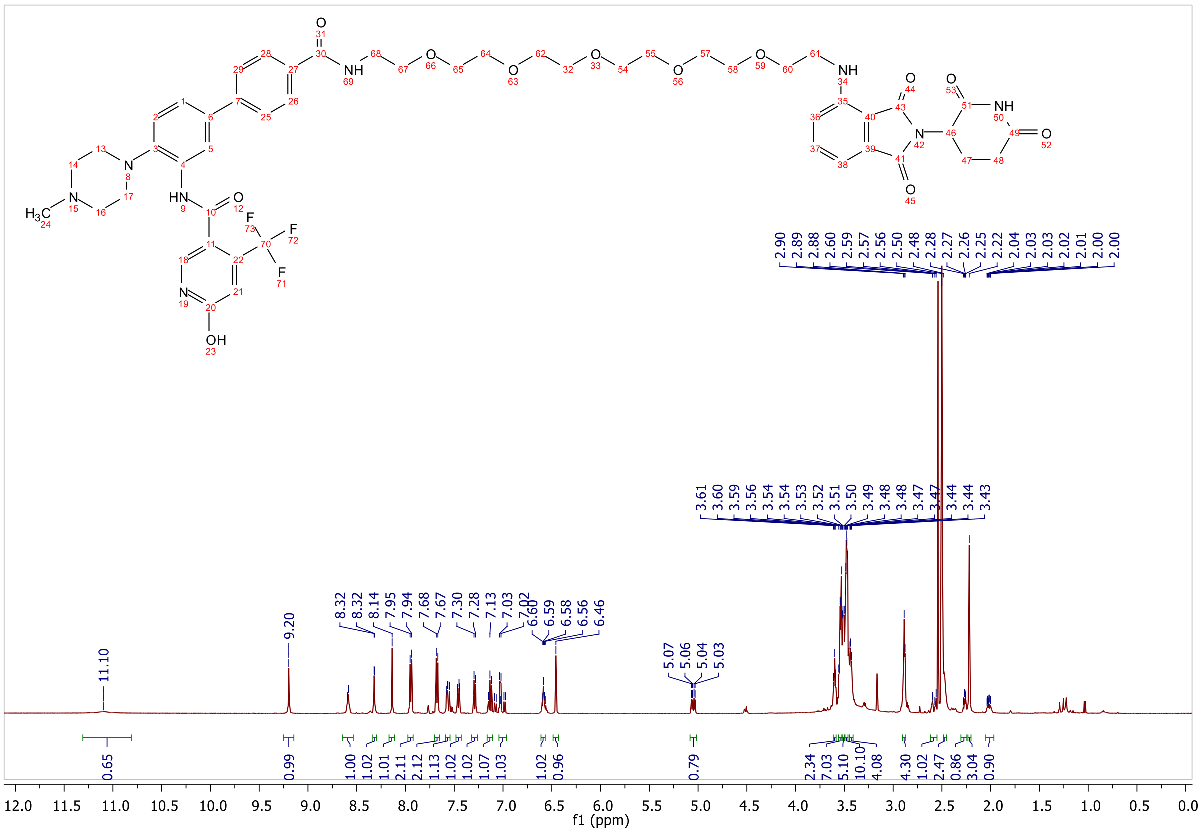

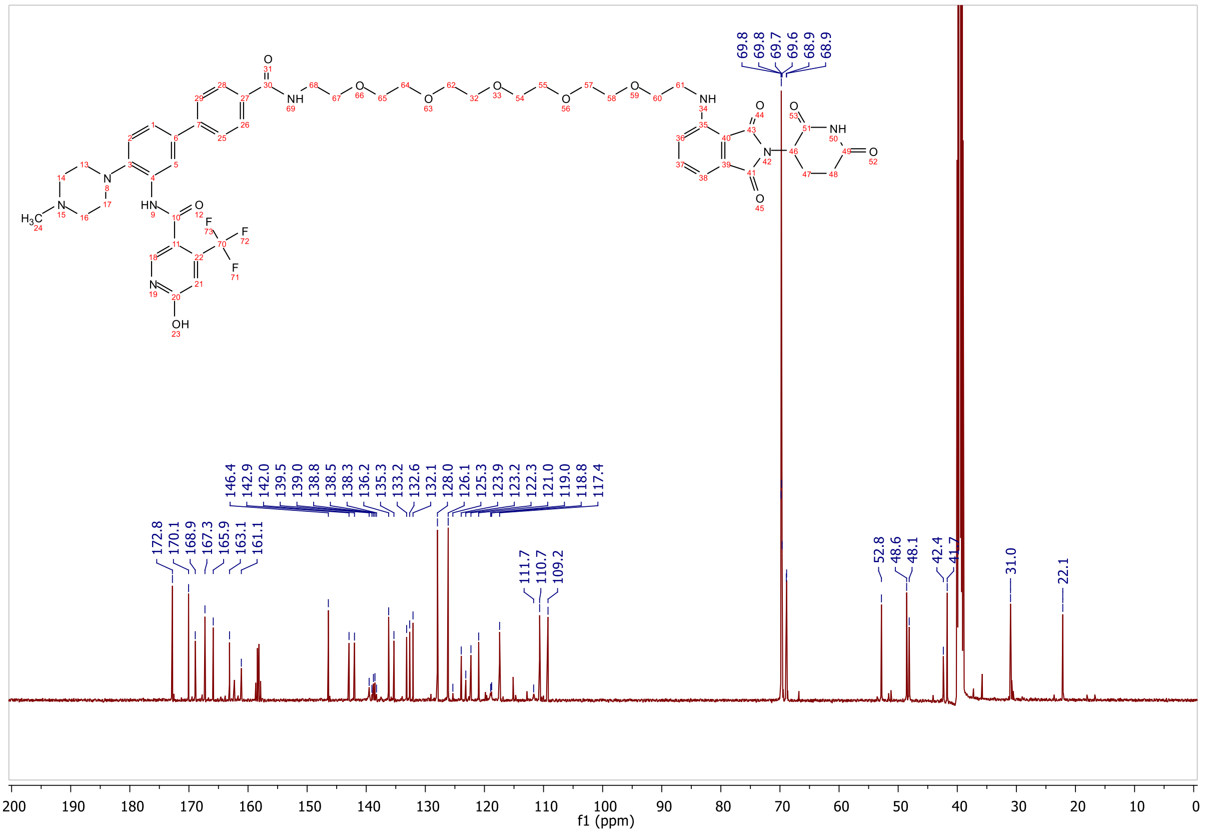

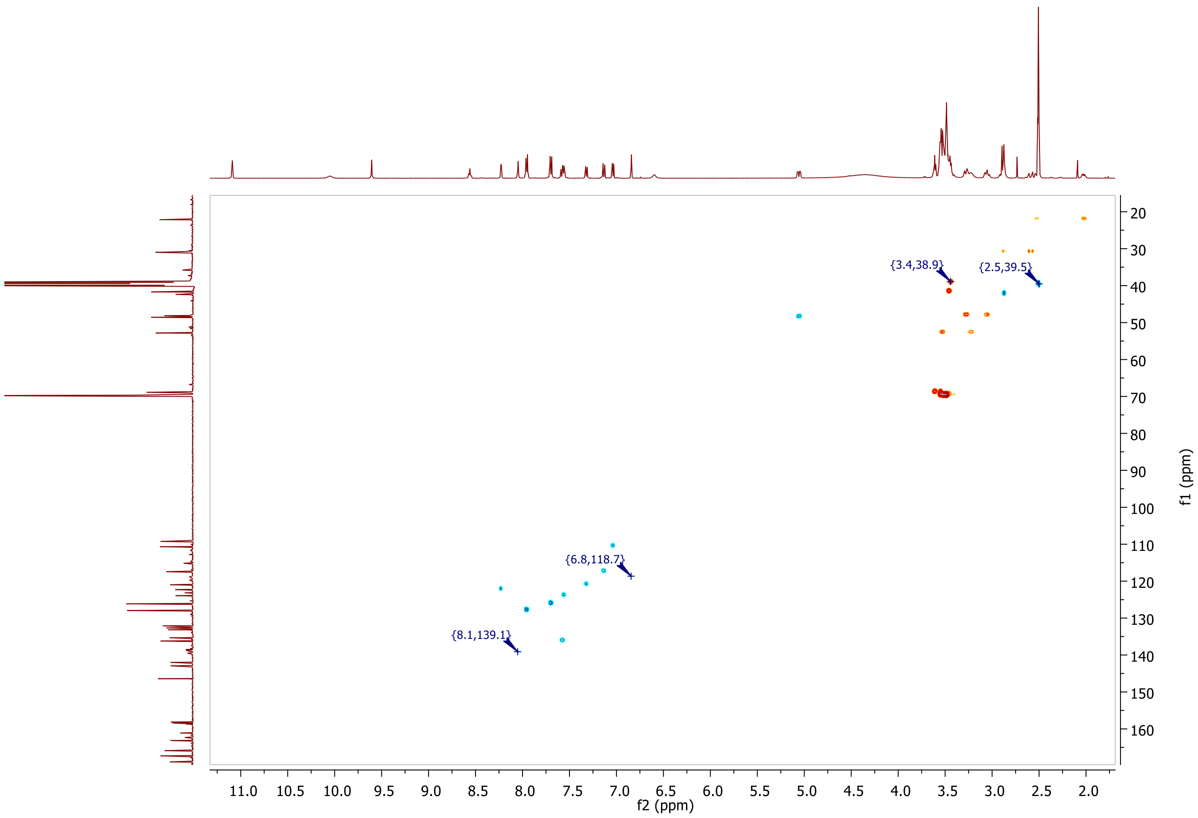
{3.4,38.9}
{2.5,39.5}
{6.8,118.7}
{8.1,139.1}
f2 (ppm)
f1 (ppm)

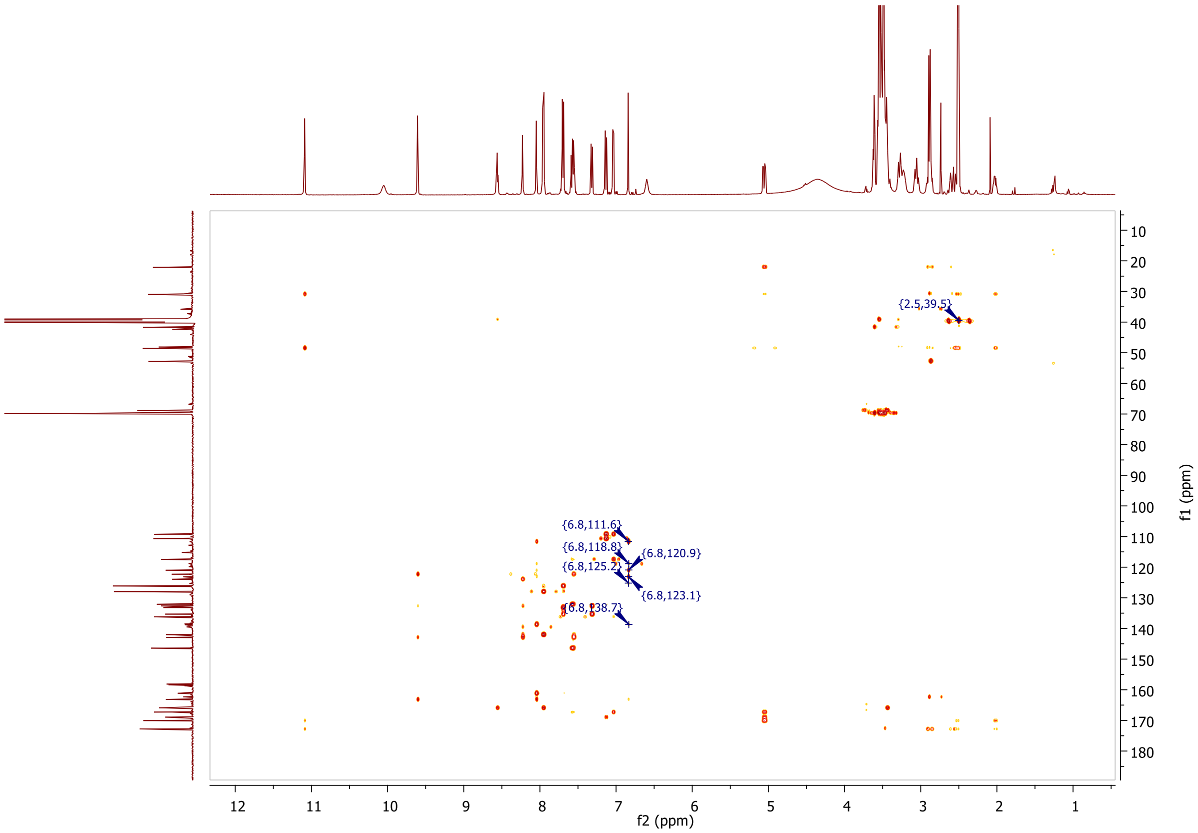
{2.5,39.5}
{6.8,111.6}
{6.8,118.8}
{6.8,120.9}
{6.8,125.2}
{6.8,123.1}
f2 (ppm)
f1 (ppm)

**HRMS, MALDI, HPLC, $^1$H-NMR, $^{13}$C-NMR, $^{13}$C-NMR (downfield region), $^1$H-$^{13}$C-HSQC and $^1$H-$^{13}$C-HMBC of *N*-(4'-((23-((2-(2,6-dioxopiperidin-3-yl)-1,3-dioxoisoindolin-4-yl)amino)-3,6,9,12,15,18,21-heptaoxatricosyl)carbamoyl)-4-(4-methylpiperazin-1-yl)-[1,1'-biphenyl]-3-yl)-6-hydroxy-4-(trifluoromethyl)nicotinamide 7d**

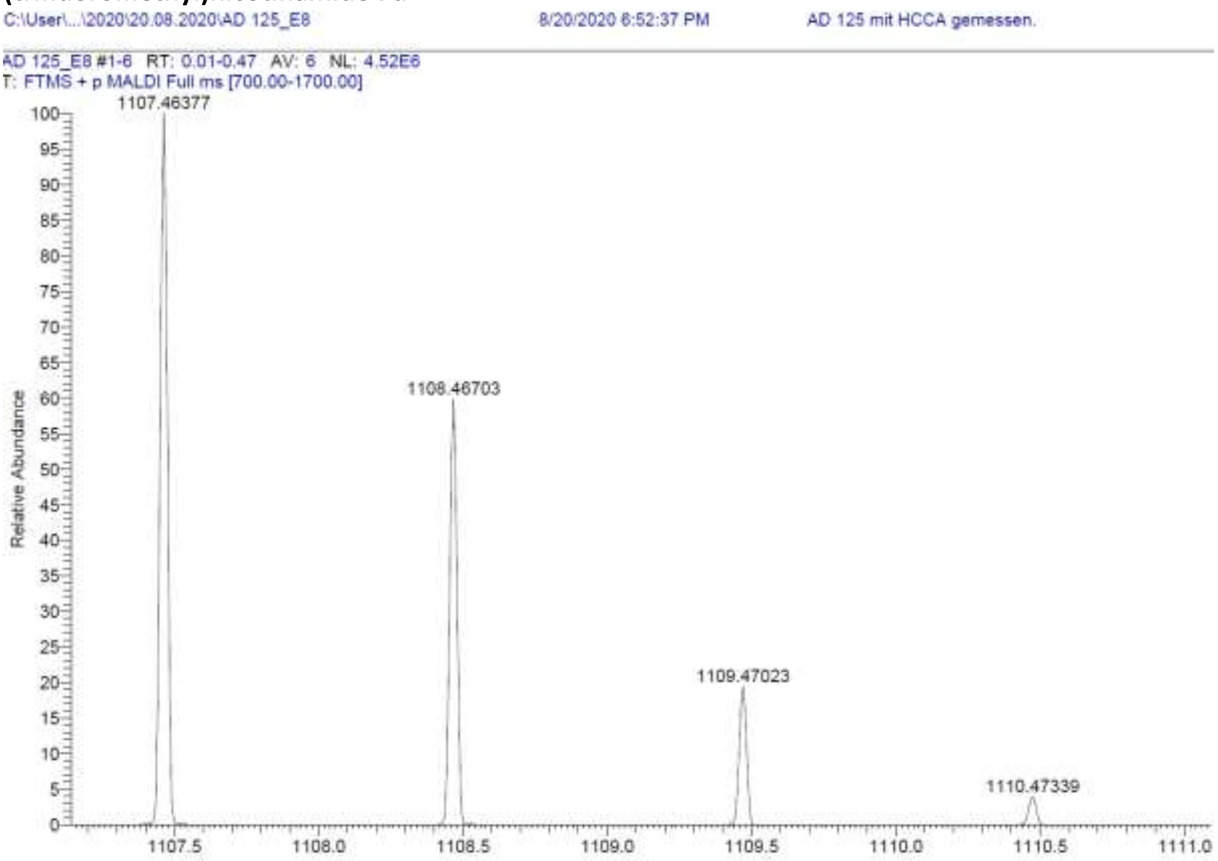

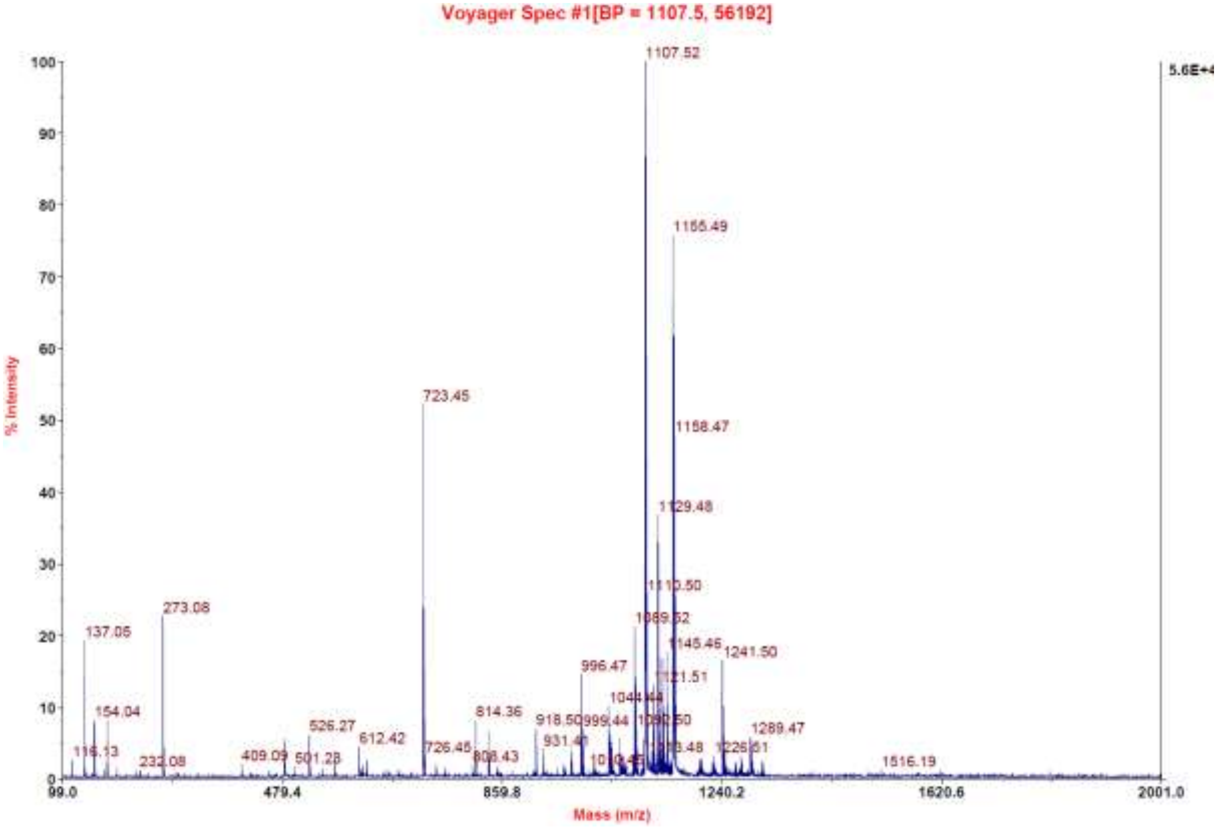

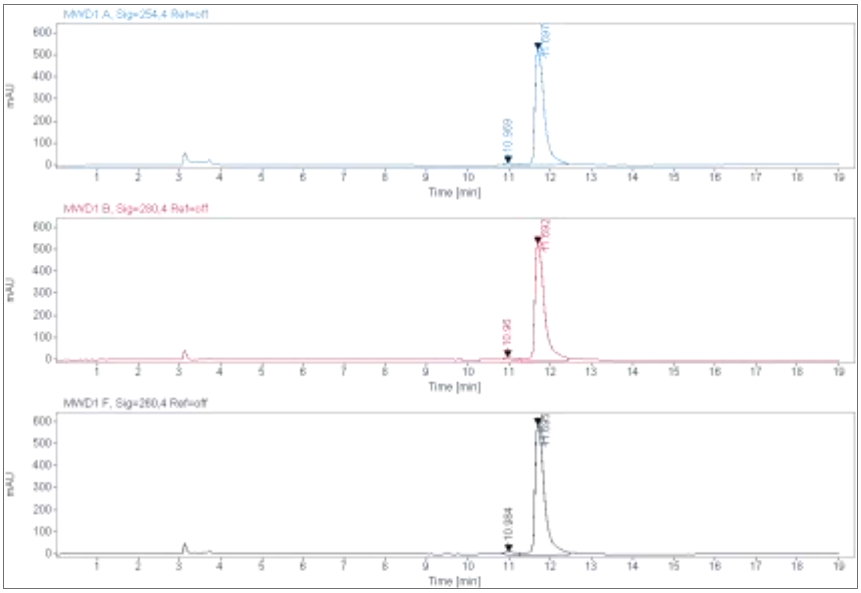

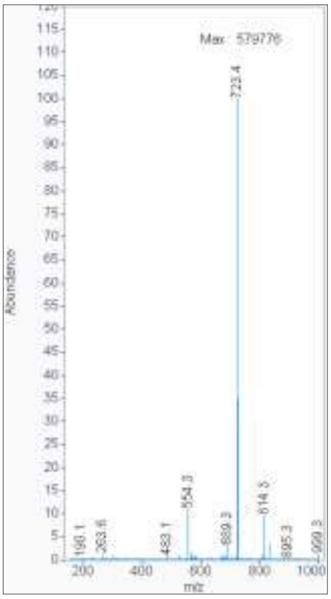

**Signal:** MWD1 A, Sig=254,4 Ref=off

| RT [min] | Type | Width [min] | Area | Height | Area% |
|---|---|---|---|---|---|
| 10.969 | MM | 0.3015 | 220.6292 | 12.1971 | 2.5413 |
| 11.697 | MM | 0.2683 | 8461.0498 | 525.6016 | 97.4587 |
| | | Sum | 8681.6790 | | |

**Signal:** MWD1 B, Sig=280,4 Ref=off

| RT [min] | Type | Width [min] | Area | Height | Area% |
|---|---|---|---|---|---|
| 10.960 | MM | 0.2779 | 234.3564 | 14.0530 | 2.5827 |
| 11.692 | MM | 0.2764 | 8839.7959 | 533.0018 | 97.4173 |
| | | Sum | 9074.1523 | | |

**Signal:** MWD1 F, Sig=260,4 Ref=off

| RT [min] | Type | Width [min] | Area | Height | Area% |
|---|---|---|---|---|---|
| 10.984 | MM | 0.3388 | 313.1158 | 15.4024 | 3.0819 |
| 11.695 | MM | 0.2776 | 9846.8340 | 591.2772 | 96.9181 |

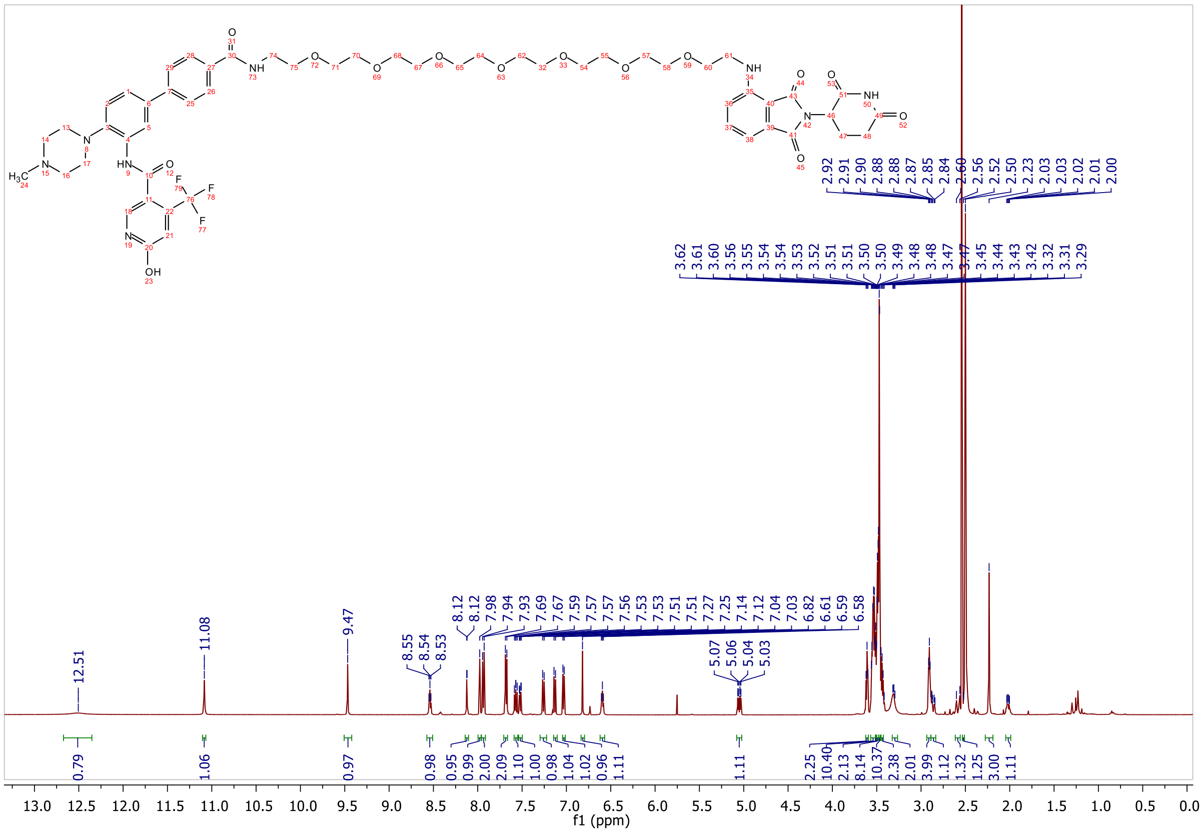

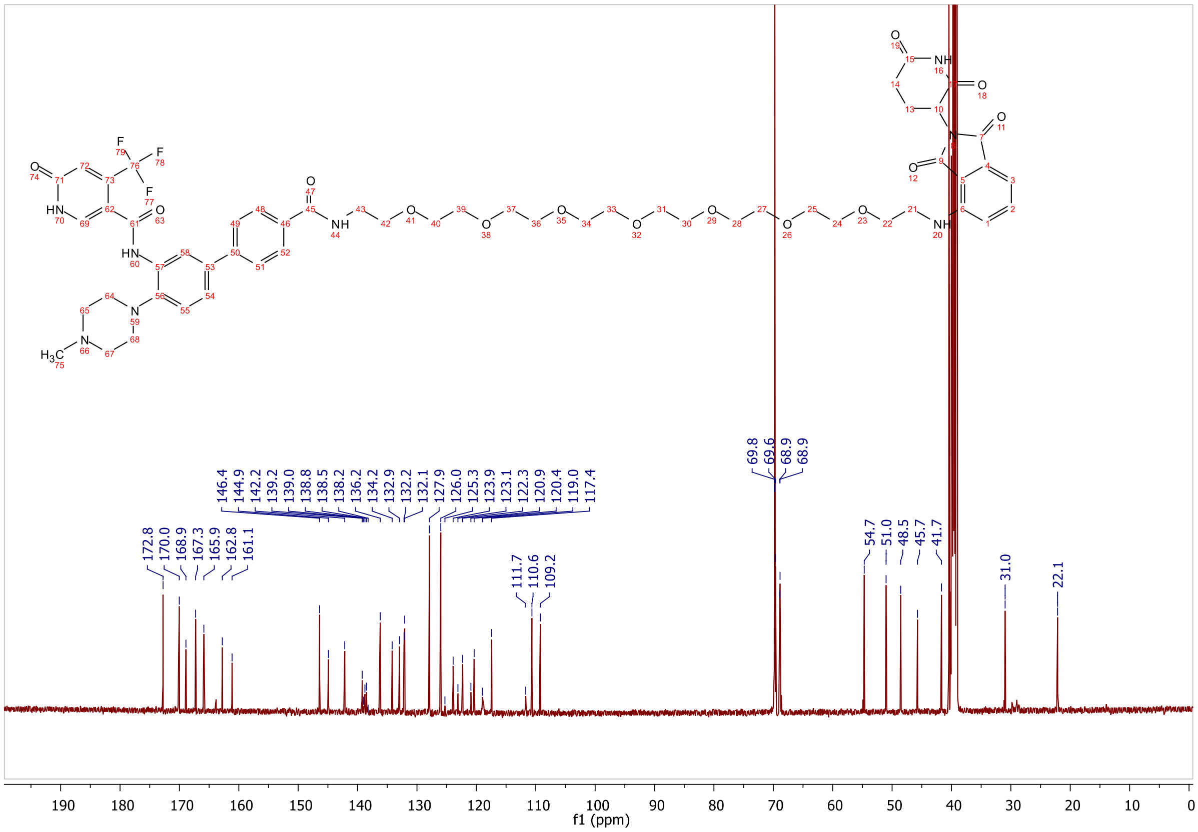

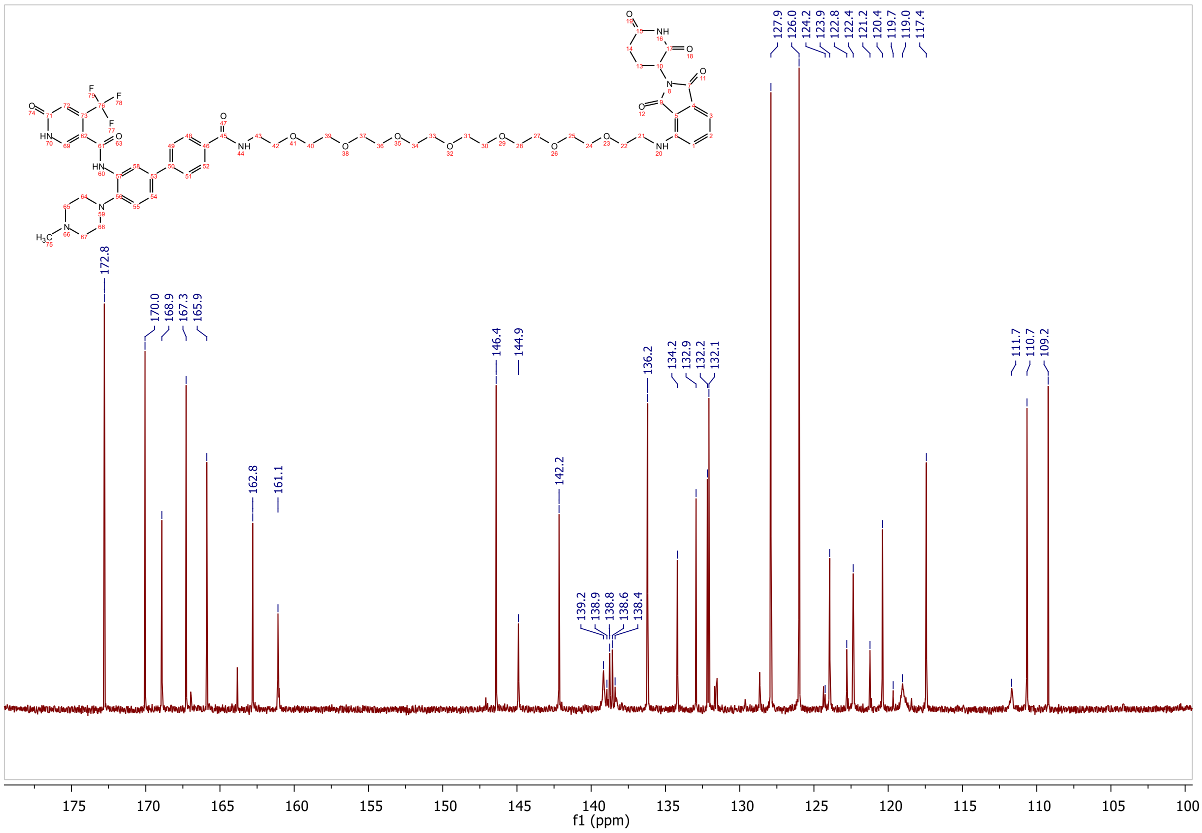
172.8
170.0
168.9
167.3
165.9
162.8
161.1
146.4
144.9
142.2
139.2
138.9
138.8
138.6
138.4
136.2
134.2
132.9
132.2
132.1
127.9
126.0
124.2
123.9
122.8
122.4
121.2
120.4
119.7
119.0
117.4
111.7
110.7
109.2
f1 (ppm)

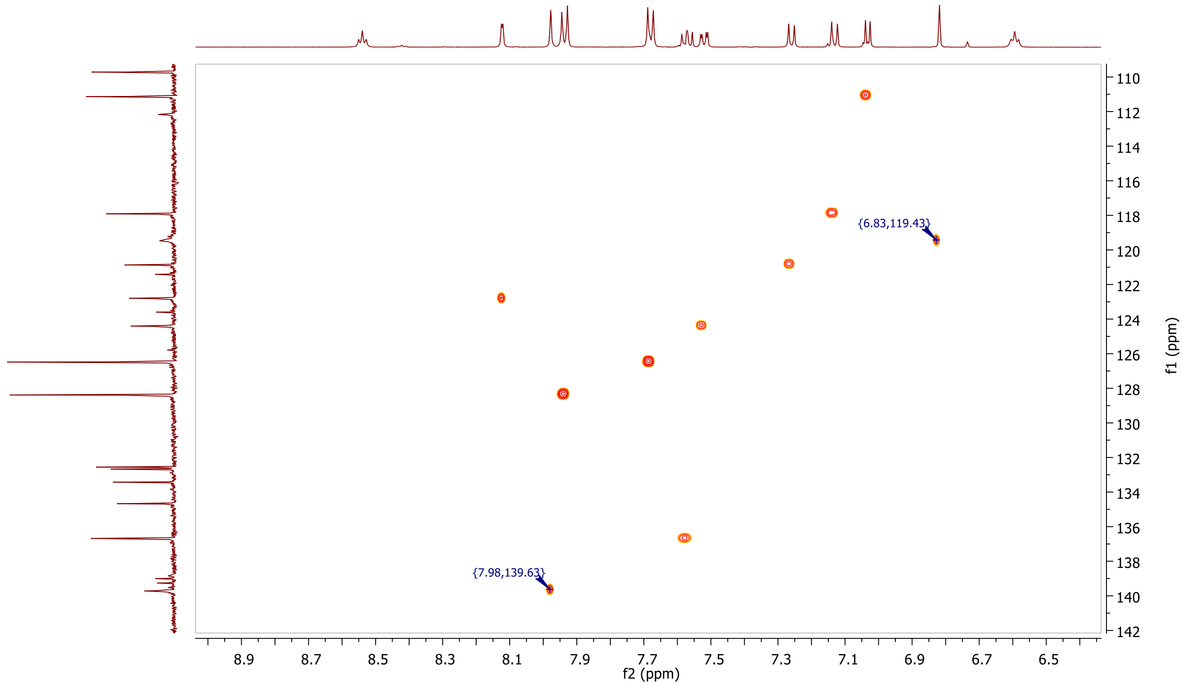
{6.83,119.43}
{7.98,139.63}
f2 (ppm)
f1 (ppm)

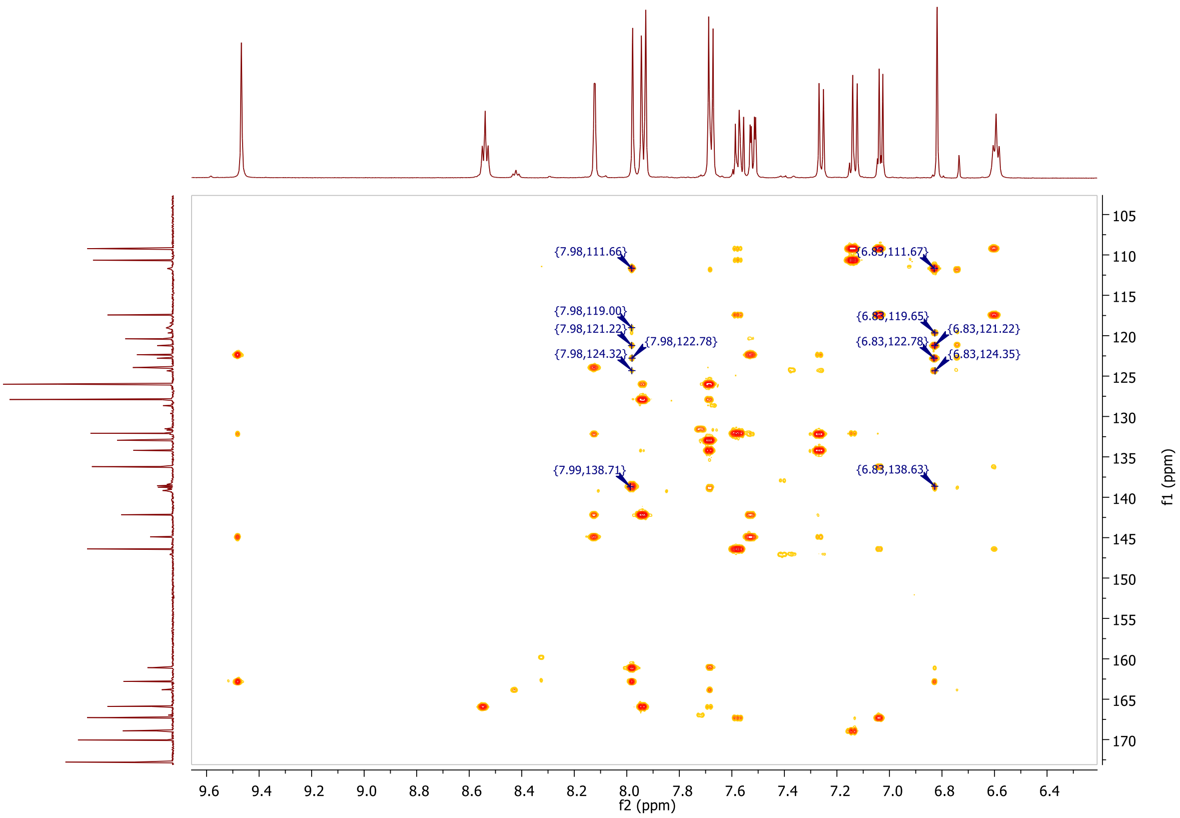
{7.98,111.66}
{6.83,111.67}
{7.98,119.00}
{7.98,121.22}
{7.98,122.78}
{7.98,124.32}
{6.83,119.65}
{6.83,121.22}
{6.83,122.78}
{6.83,124.35}
{7.99,138.71}
{6.83,138.63}
f2 (ppm)
f1 (ppm)

**MALDI, HRMS, $^{1}$H-NMR, $^{13}$C-NMR, $^{1}$H-$^{13}$C-HSQC, $^{1}$H-$^{13}$C-HMBC and HPLC of N-(4'-((4-(((2-(2,6-dioxopiperidin-3-yl)-1,3-dioxoisoindolin-4-yl)amino)methyl)benzyl)carbamoyl)-4-(4-methylpiperazin-1-yl)-[1,1'-biphenyl]-3-yl)-6-hydroxy-4-(trifluoromethyl)nicotinamide 7e**

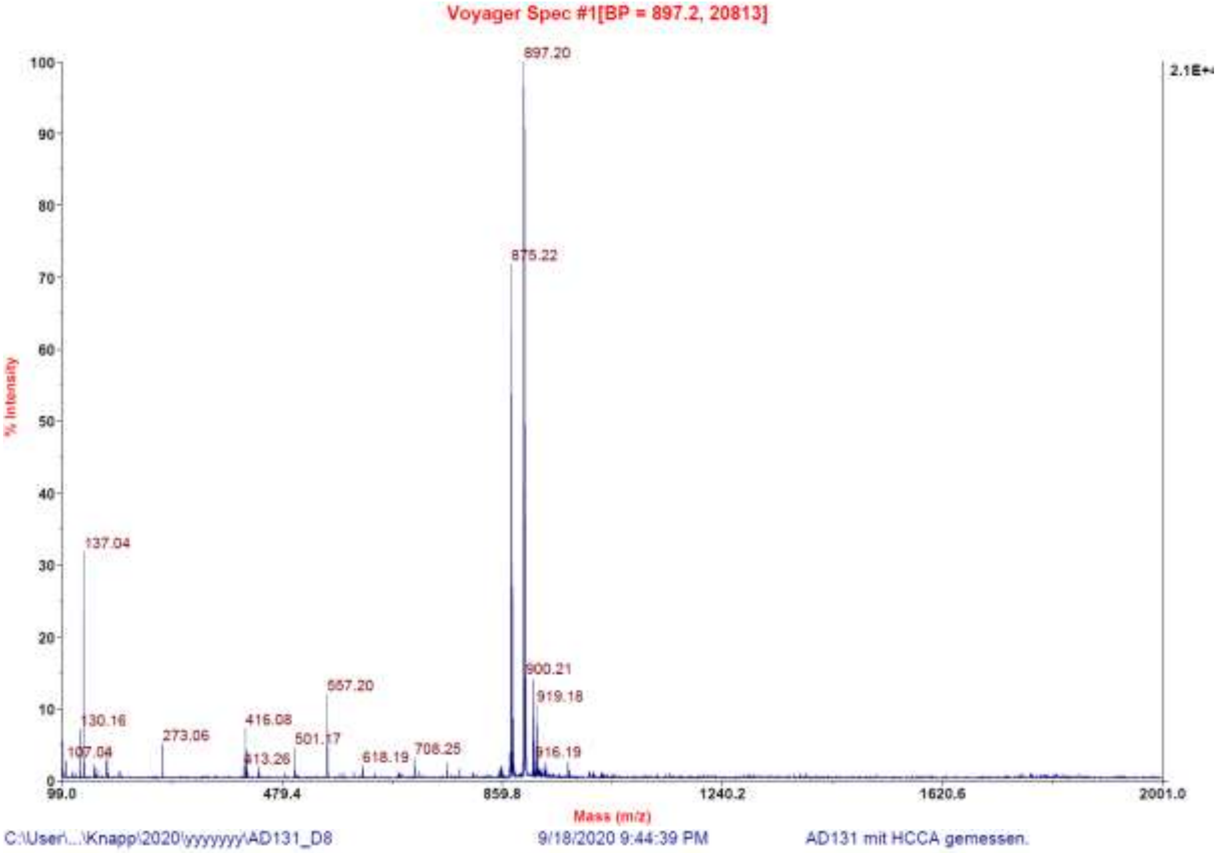

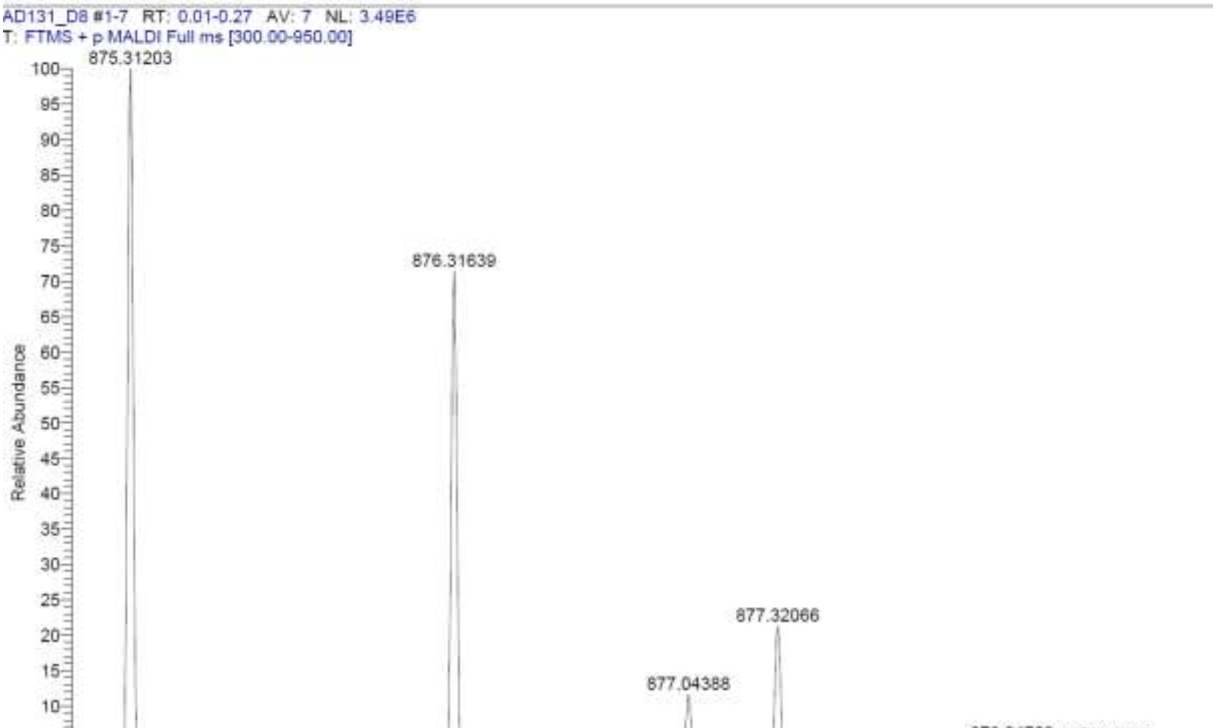

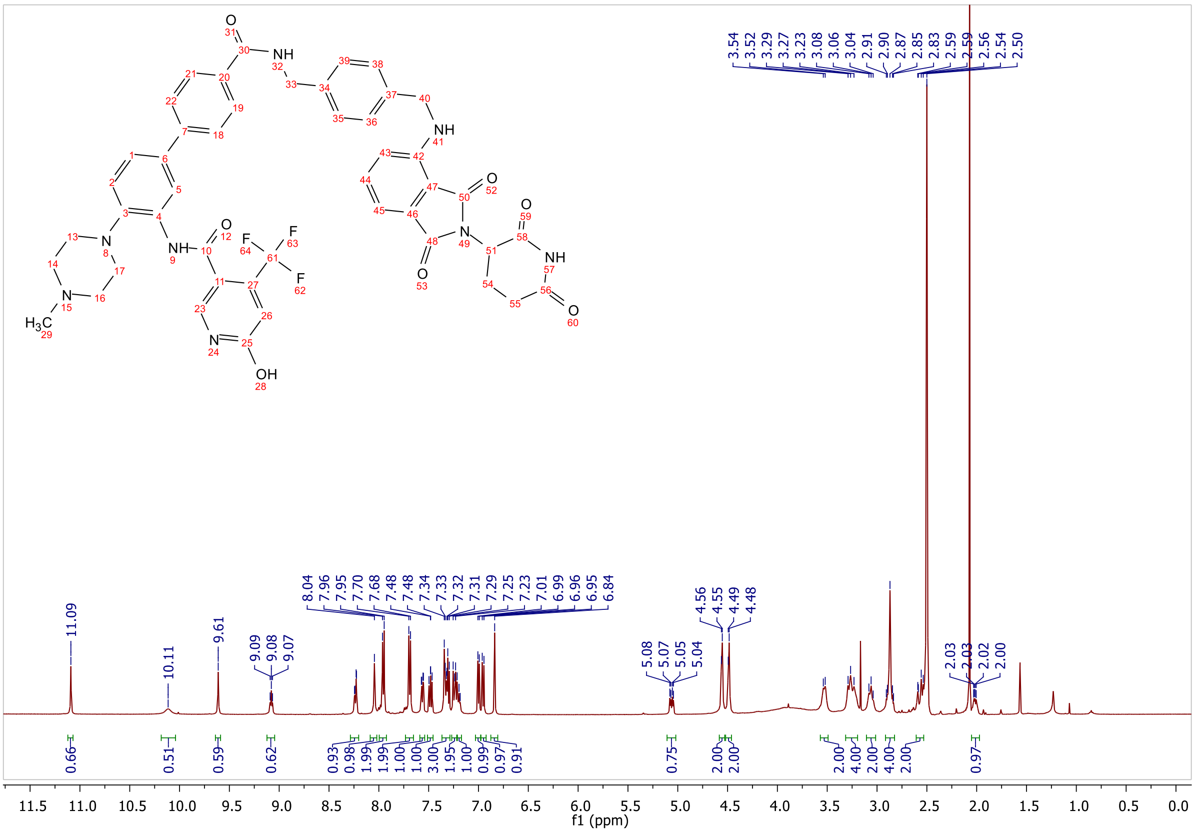

3.54 3.52 3.29 3.27 3.23 3.08 3.06 3.04 2.91 2.90 2.87 2.85 2.83 2.59 2.56 2.54 2.50
11.09
10.11
9.61
9.09 9.08 9.07
8.04 7.96 7.95 7.70 7.68 7.48 7.34 7.33 7.32 7.31 7.29 7.25 7.23 7.01 6.99 6.96 6.95 6.84
5.08 5.07 5.05 5.04
4.56 4.55 4.49 4.48
2.03 2.03 2.02 2.00
0.66 0.51 0.59 0.62 0.93 0.98 1.99 1.99 1.00 1.00 3.00 1.95 1.00 0.99 0.97 0.91 0.75 2.00 2.00 2.00 4.00 2.00 4.00 2.00 0.97
f1 (ppm)

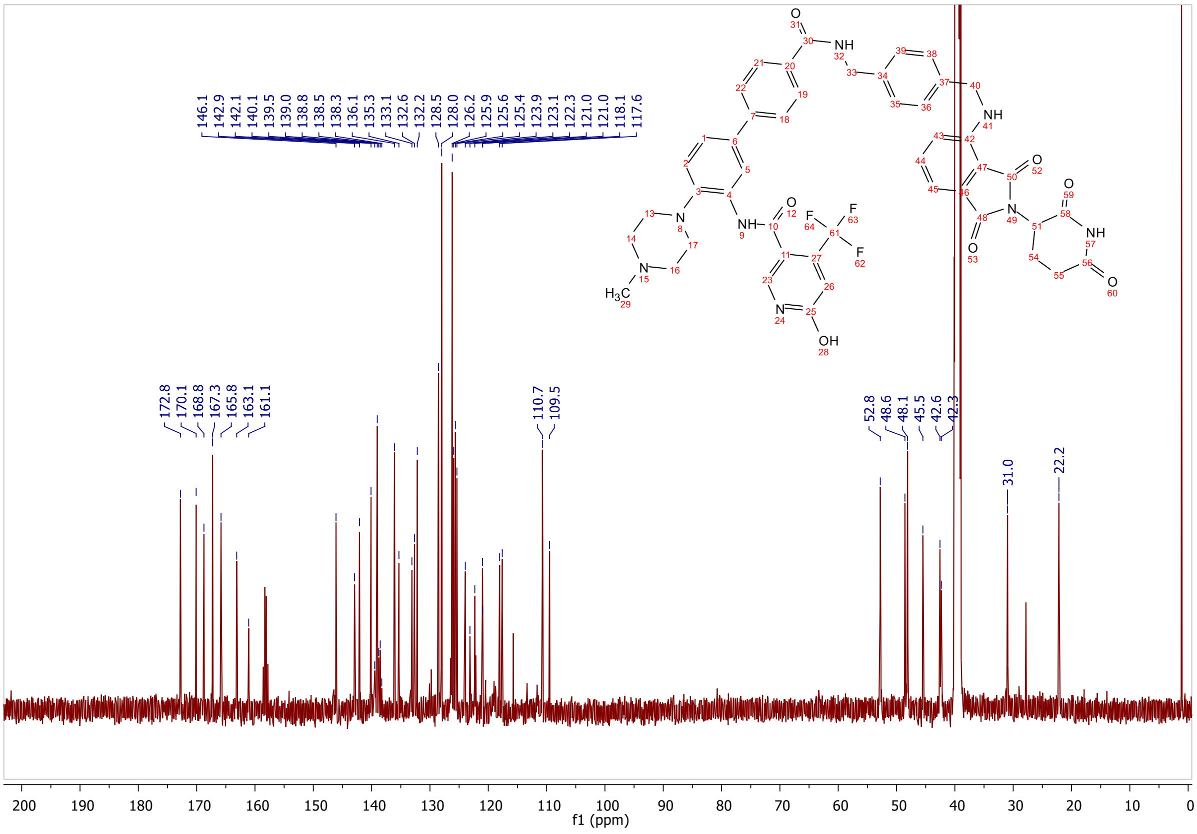

146.1 142.9 142.1 140.1 139.5 139.0 138.8 138.5 138.3 136.1 135.3 133.1 132.6 132.2 128.5 128.0 126.2 125.9 125.6 125.4 123.9 123.1 122.3 121.0 121.0 118.1 117.6
172.8 170.1 168.8 167.3 165.8 163.1 161.1
110.7 109.5
52.8 48.6 48.1 45.5 42.6 42.3
31.0
22.2
f1 (ppm)

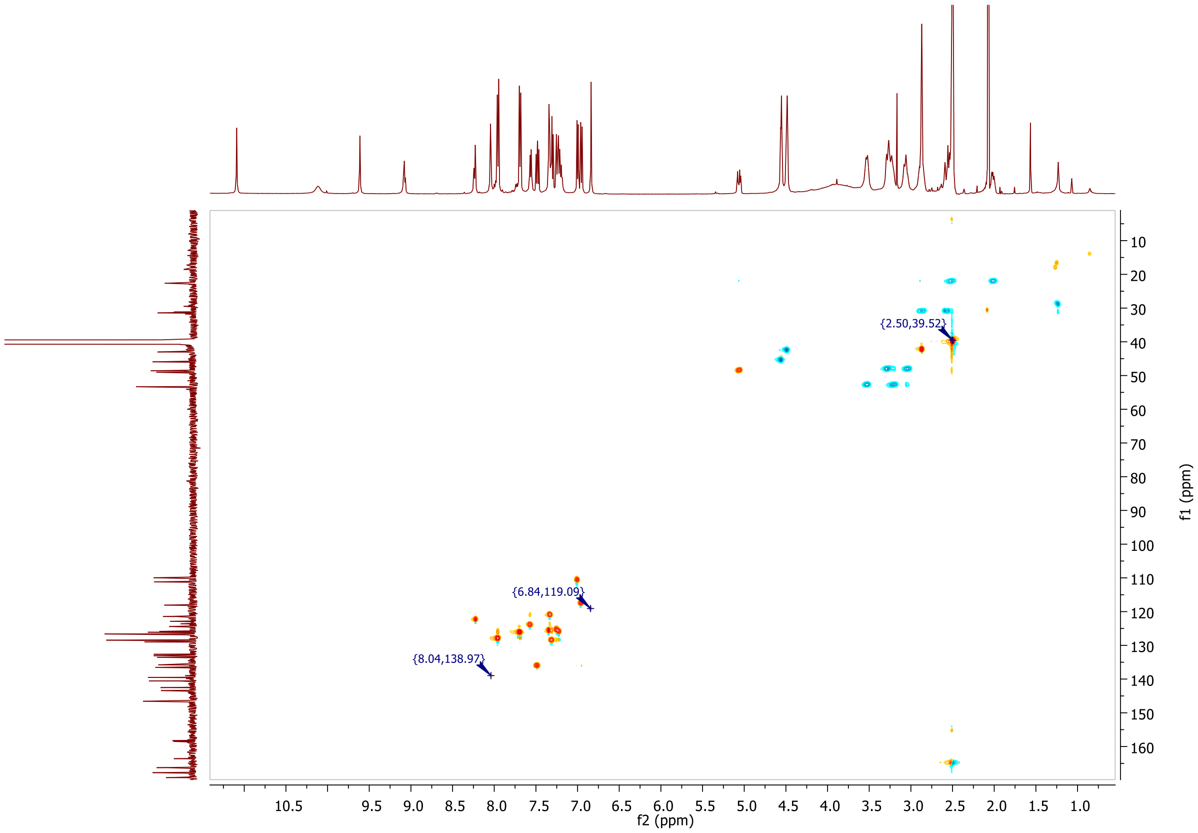

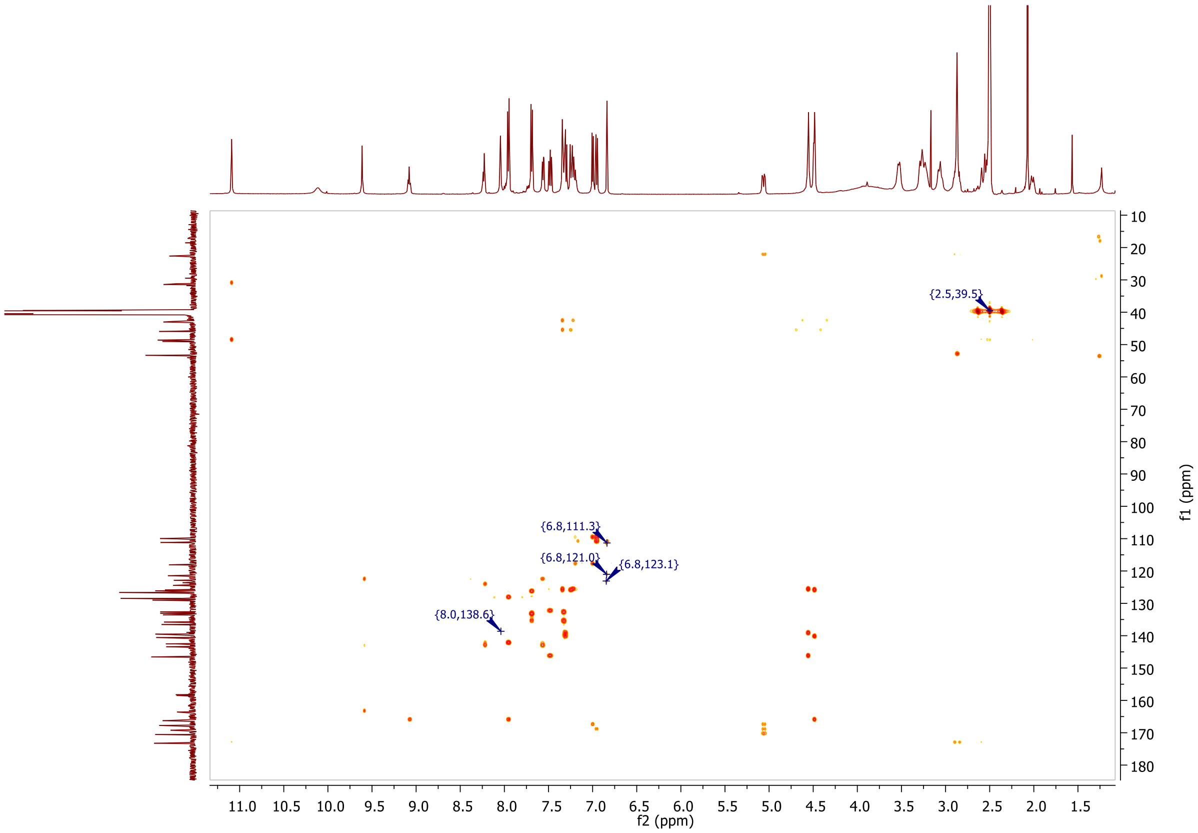

**Signal:** MWD1 A, Sig=254,4 Ref=off

| RT [min] | Type | Width [min] | Area | Height | Area% |
|---|---|---|---|---|---|
| 11.654 | MM | 0.1599 | 142.2890 | 14.8330 | 1.5723 |

| | | | | | |
|---|---|---|---|---|---|
| 12.065 | MM | 0.2775 | 8600.4414 | 516.4586 | 95.0367 |
| 12.892 | MM | 0.2353 | 306.8708 | 21.7380 | 3.3910 |
| | | Sum | 9049.6012 | | |

**Signal:** MWD1 B, Sig=280,4 Ref=off

| RT [min] | Type | Width [min] | Area | Height | Area% |
|---|---|---|---|---|---|
| 11.659 | MM | 0.1161 | 118.6666 | 12.6600 | 1.3609 |
| 12.065 | MM | 0.2644 | 8392.1240 | 529.0167 | 96.2442 |
| 12.895 | MM | 0.1790 | 208.8267 | 19.4480 | 2.3949 |
| | | Sum | 8719.6173 | | |

**Signal:** MWD1 F, Sig=260,4 Ref=off

| RT [min] | Type | Width [min] | Area | Height | Area% |
|---|---|---|---|---|---|
| 11.661 | MM | 0.1766 | 173.2003 | 16.3418 | 1.7365 |
| 12.065 | MM | 0.2720 | 9509.6045 | 582.6867 | 95.3410 |
| 12.889 | MM | 0.1962 | 291.4992 | 24.7618 | 2.9225 |

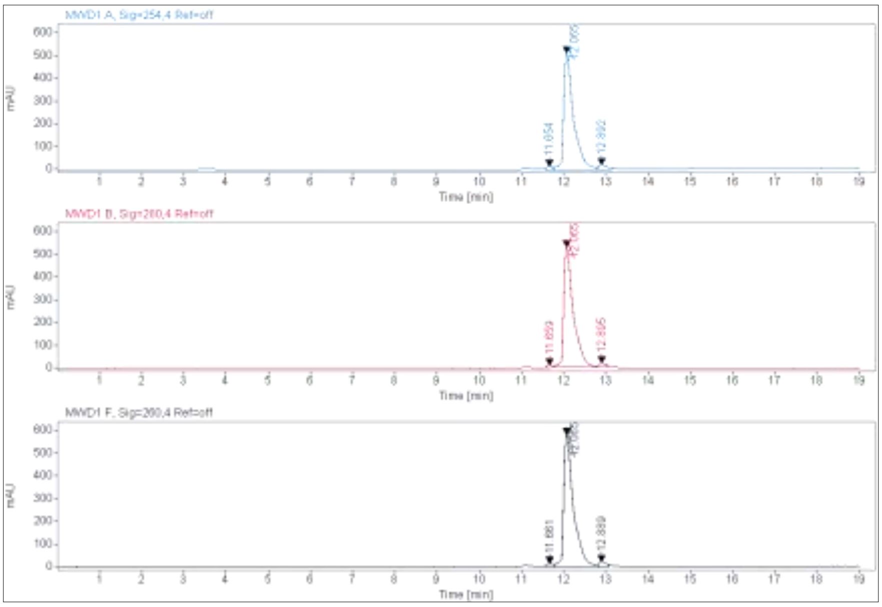

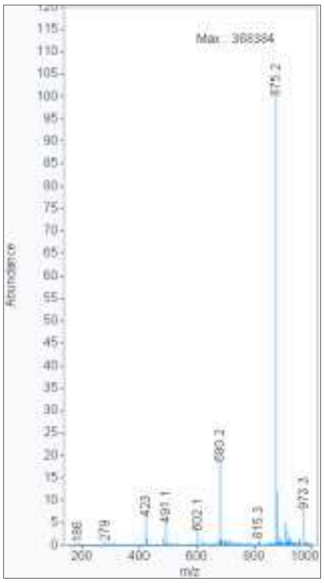

**MALDI, HRMS, HPLC, $^{1}$H-NMR, $^{13}$C-NMR, $^{1}$H-$^{13}$C-HSQC and $^{1}$H-$^{13}$C-HMBC of 6-hydroxy-N-(4'-((2-(3-(((S)-1-((2S,4R)-4-hydroxy-2-((4-(4-methylthiazol-5-yl)benzyl)carbamoyl)pyrrolidin-1-yl)-3,3-dimethyl-1-oxobutan-2-yl)amino)-3-oxopropoxy)ethyl)carbamoyl)-4-(4-methylpiperazin-1-yl)-[1,1'-biphenyl]-3-yl)-4-(trifluoromethyl)nicotinamide 8a**

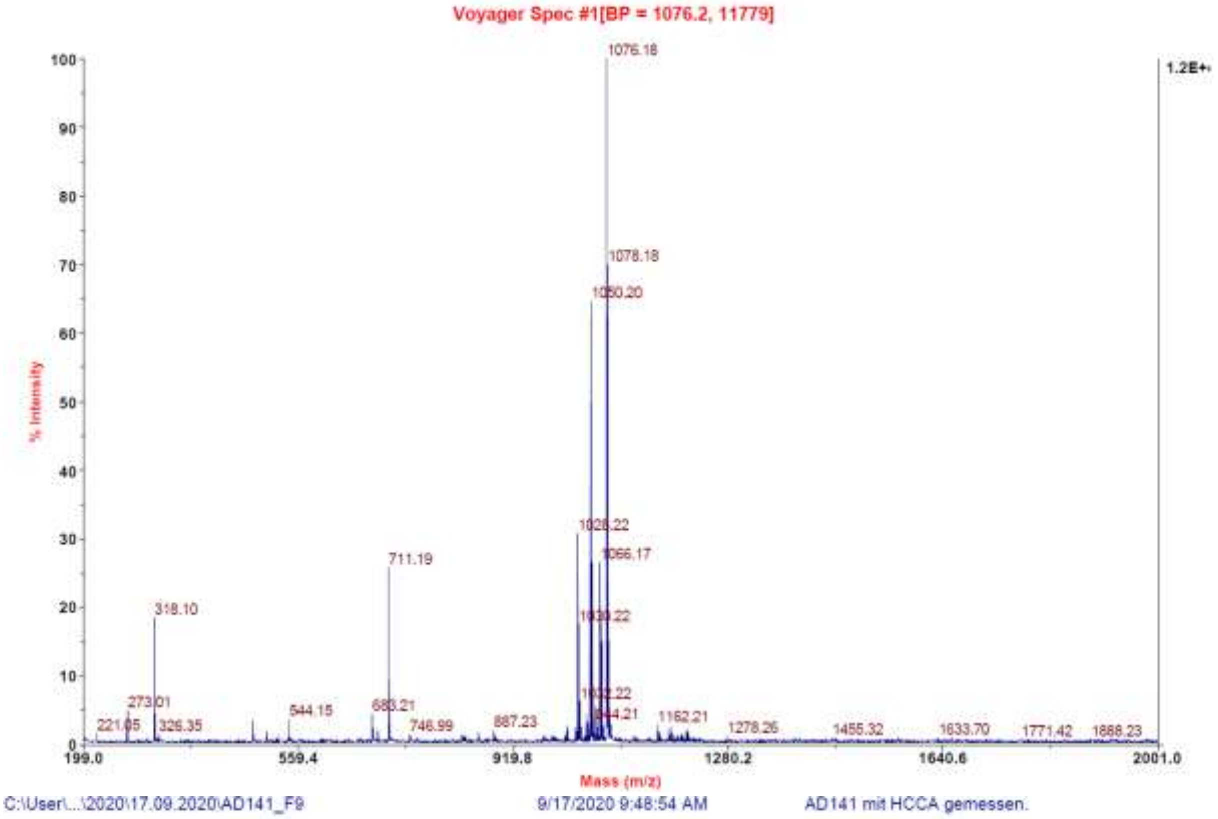

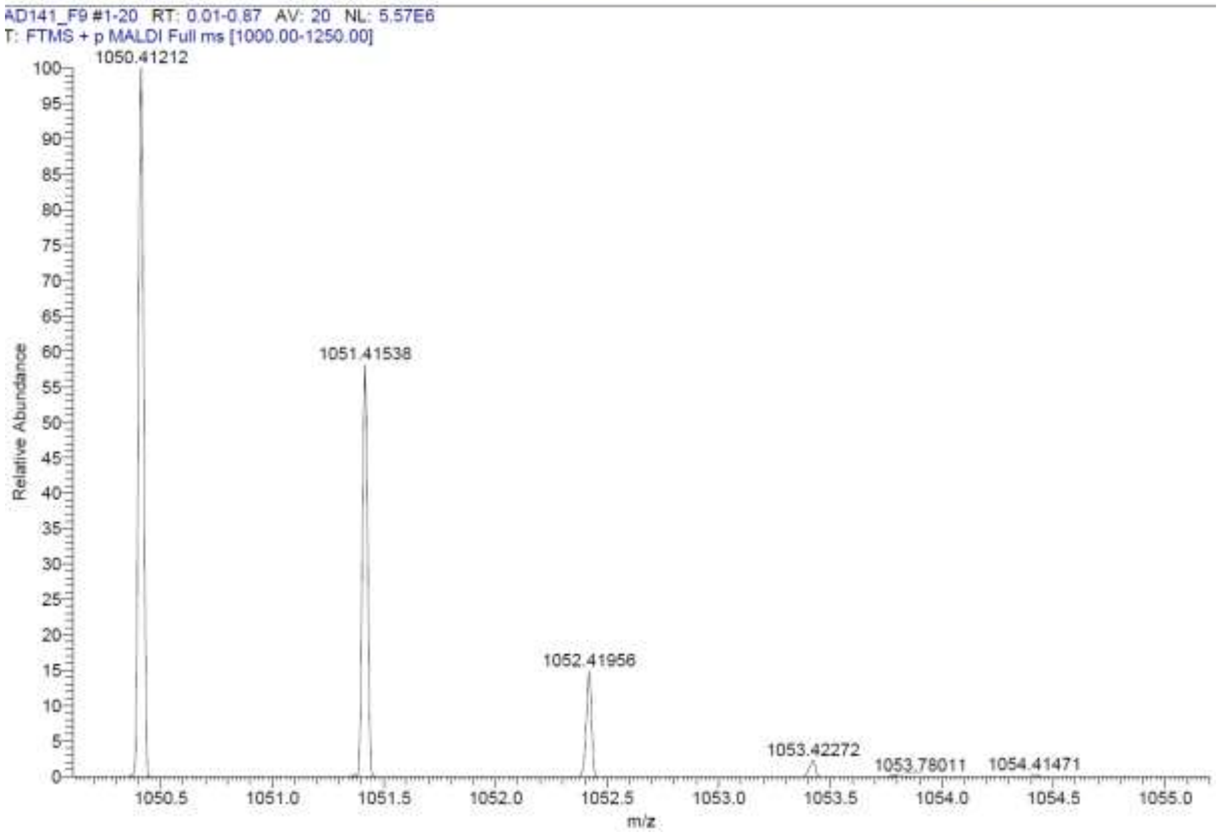

**Signal:** MWD1 A, Sig=254,4 Ref=off

| RT [min] | Type | Width [min] | Area | Height | Area% | Name |
|---|---|---|---|---|---|---|
| 11.278 | VV | 0.2050 | 7216.8145 | 486.5691 | 97.2523 | |
| 11.786 | MM | 0.1465 | 203.8960 | 23.2009 | 2.7477 | |
| | | Sum | 7420.7104 | | | |

**Signal:** MWD1 B, Sig=280,4 Ref=off

| RT [min] | Type | Width [min] | Area | Height | Area% | Name |
|---|---|---|---|---|---|---|
| 11.278 | VV | 0.2150 | 9186.0752 | 582.5713 | 100.0000 | |
| | | Sum | 9186.0752 | | | |

**Signal:** MWD1 F, Sig=260,4 Ref=off

| RT [min] | Type | Width [min] | Area | Height | Area% | Name |
|---|---|---|---|---|---|---|
| 11.278 | VV | 0.2159 | 8699.8975 | 548.8284 | 100.0000 | |

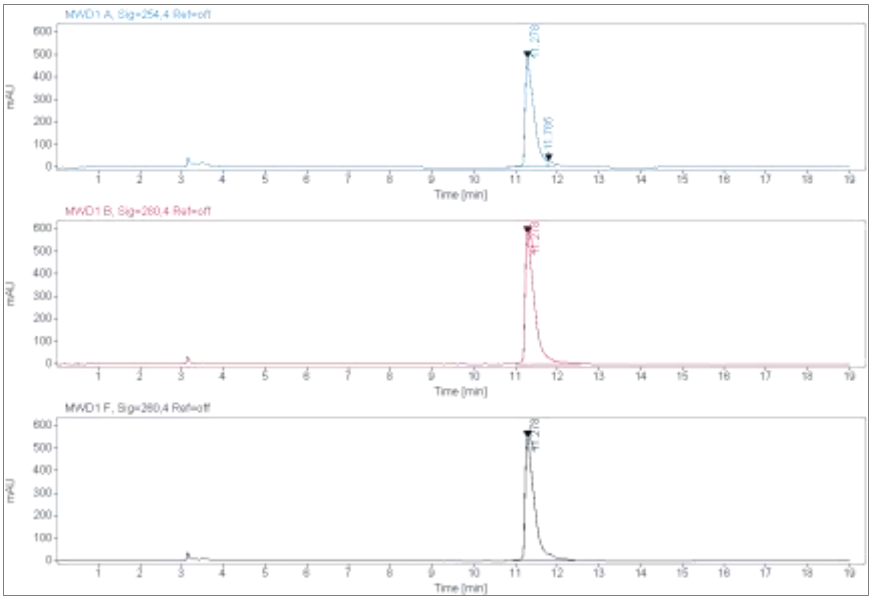

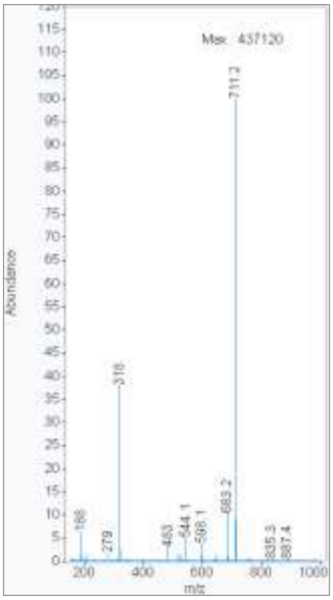

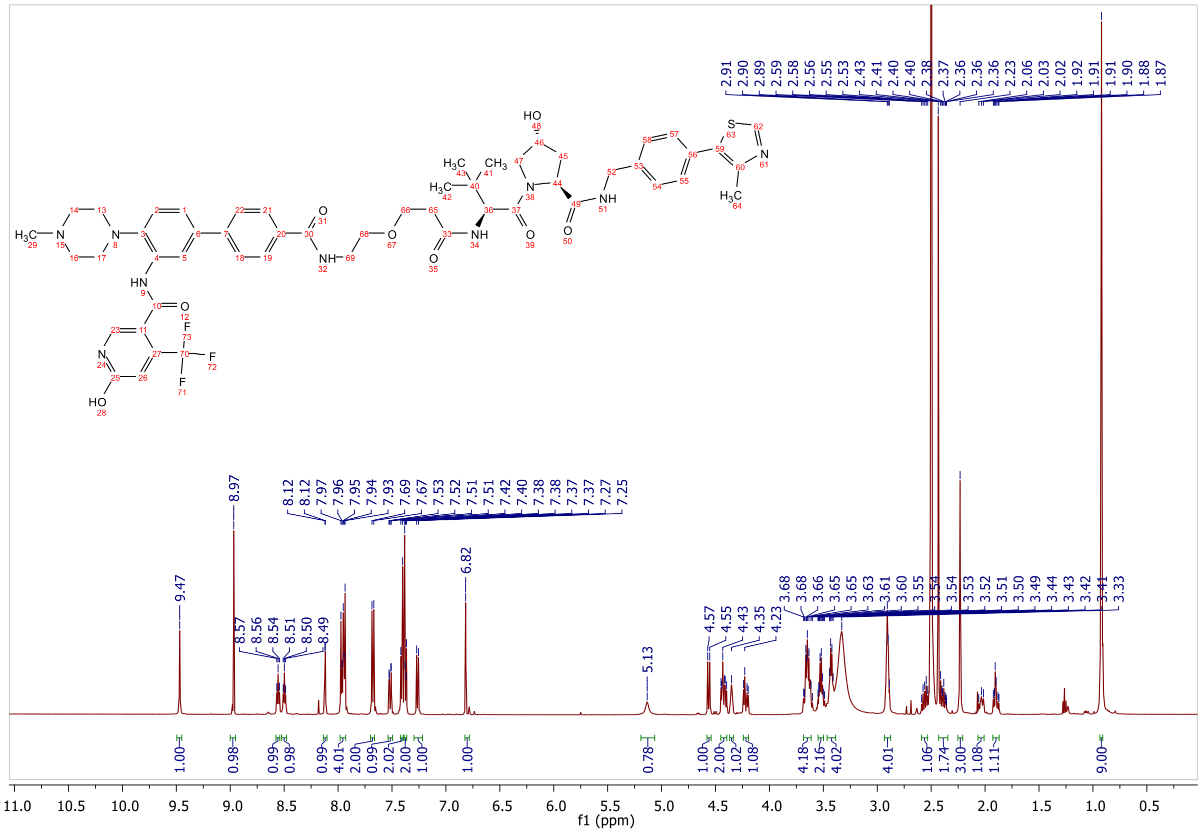

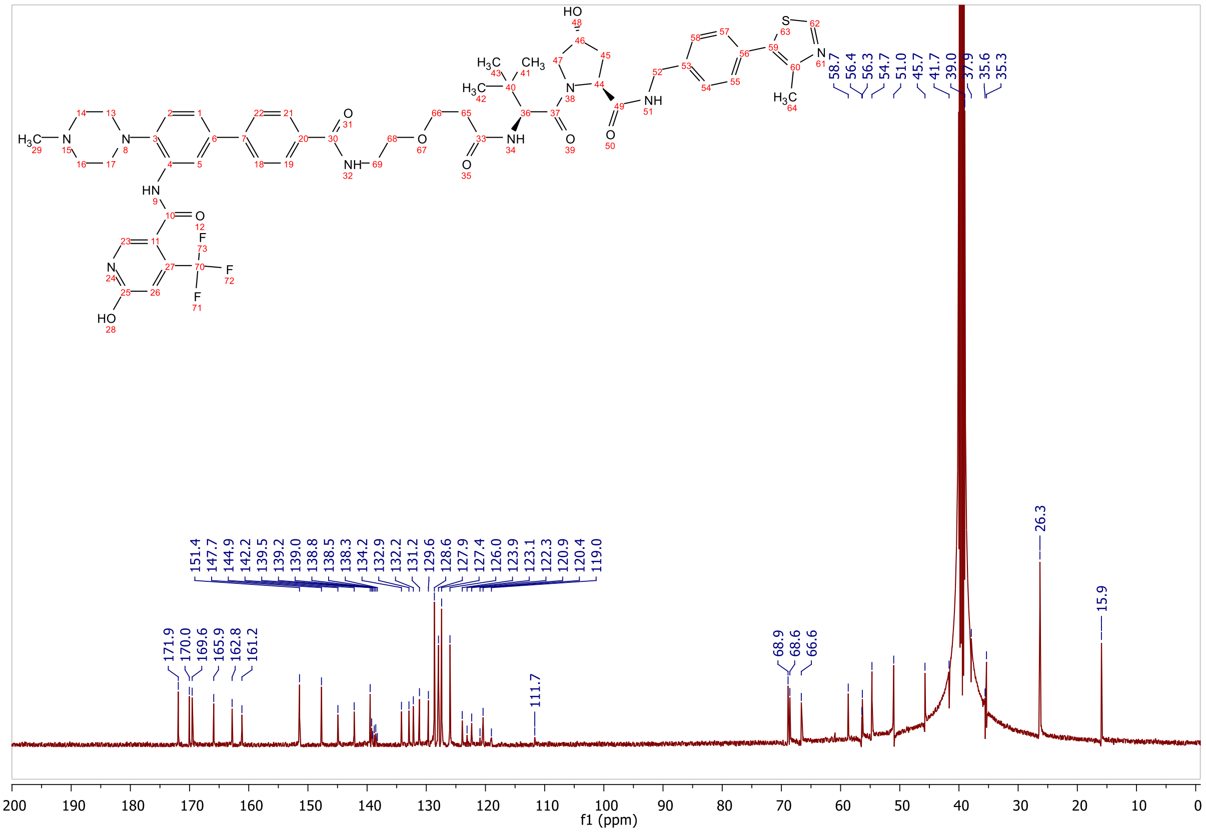

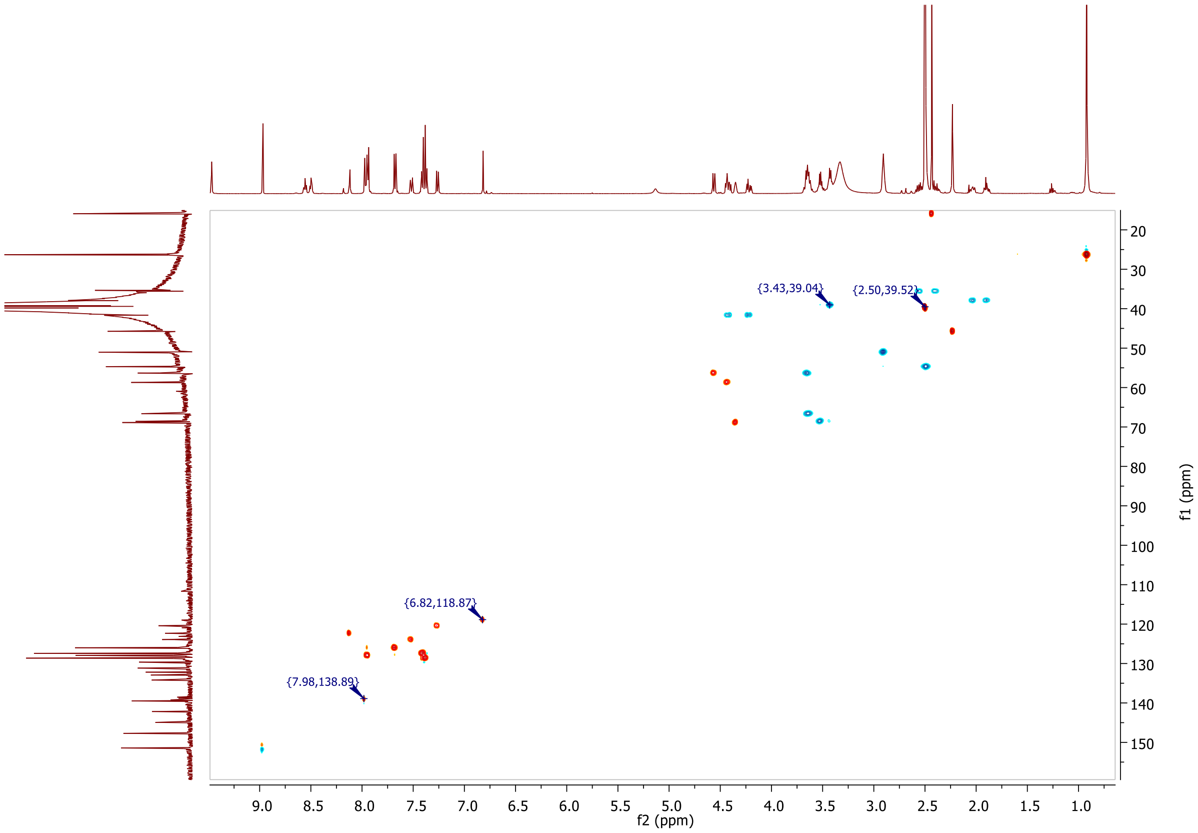
{3.43,39.04}
{2.50,39.52}
{6.82,118.87}
{7.98,138.89}
f2 (ppm)
f1 (ppm)

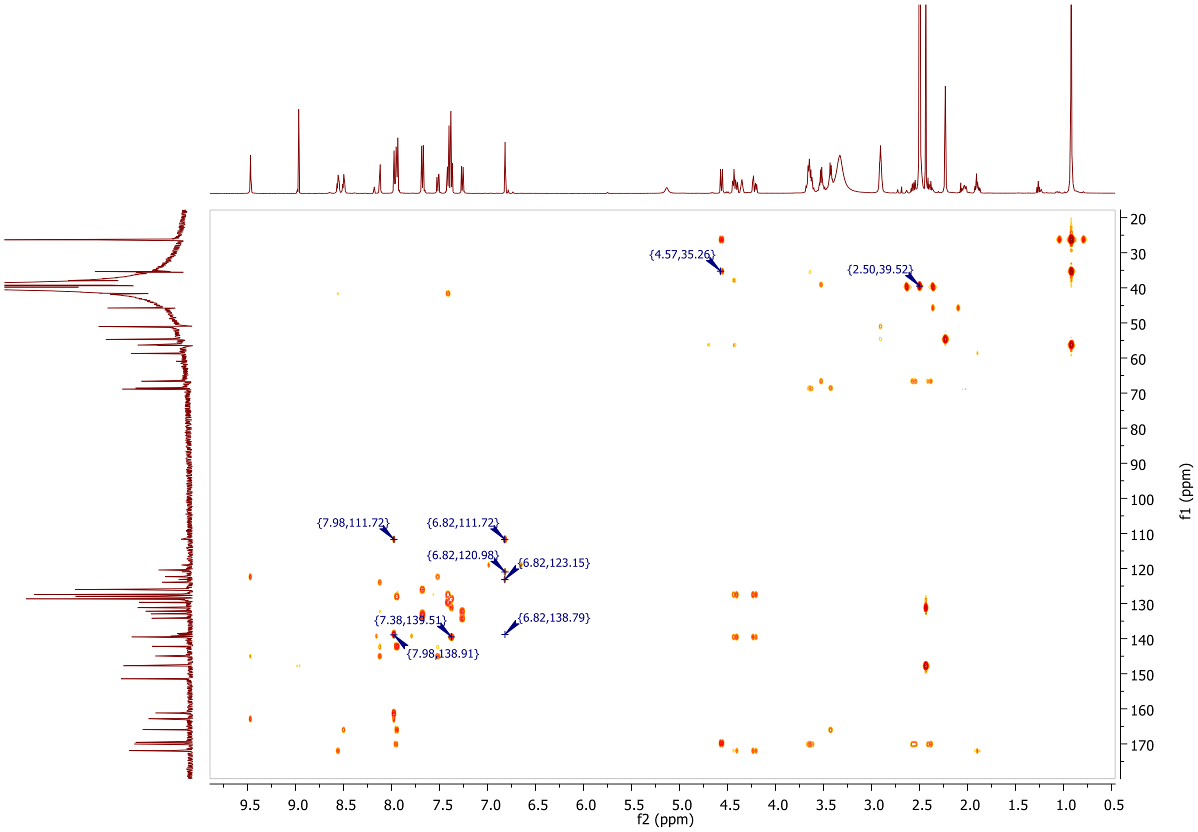
{4.57,35.26}
{2.50,39.52}
{7.98,111.72}
{6.82,111.72}
{6.82,120.98}
{6.82,123.15}
{7.38,139.51}
{6.82,138.79}
{7.98,138.91}
f2 (ppm)
f1 (ppm)

**MALDI, HRMS, HPLC, $^{1}$H-NMR and $^{13}$C-NMR of 6-hydroxy-*N*-(4'-((2-(2-(3-(((*S*)-1-((2*S*,4*R*)-4-hydroxy-2-((4-(4-methylthiazol-5-yl)benzyl)carbamoyl)pyrrolidin-1-yl)-3,3-dimethyl-1-oxobutan-2-yl)amino)-3-oxopropoxy)ethoxy)ethyl)carbamoyl)-4-(4-methylpiperazin-1-yl)-[1,1'-biphenyl]-3-yl)-4-(trifluoromethyl)nicotinamide 8b**

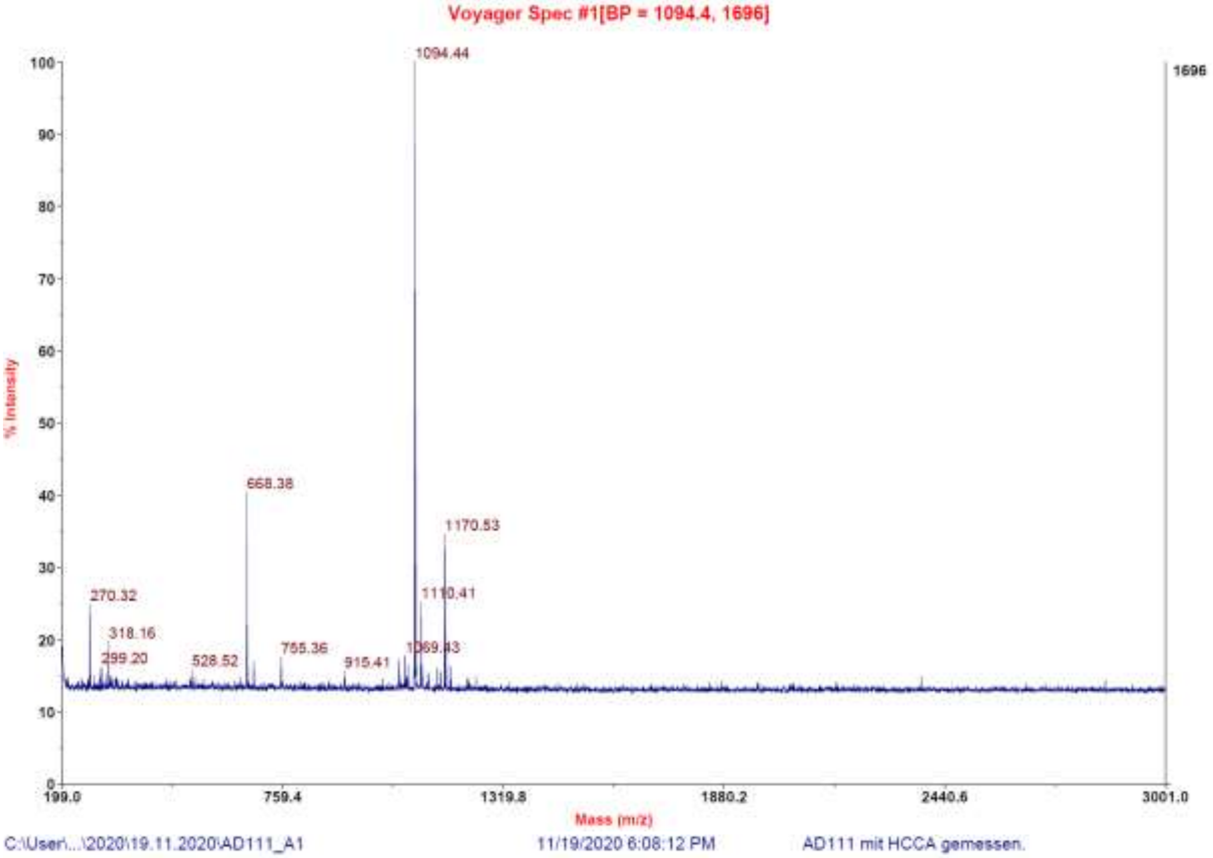

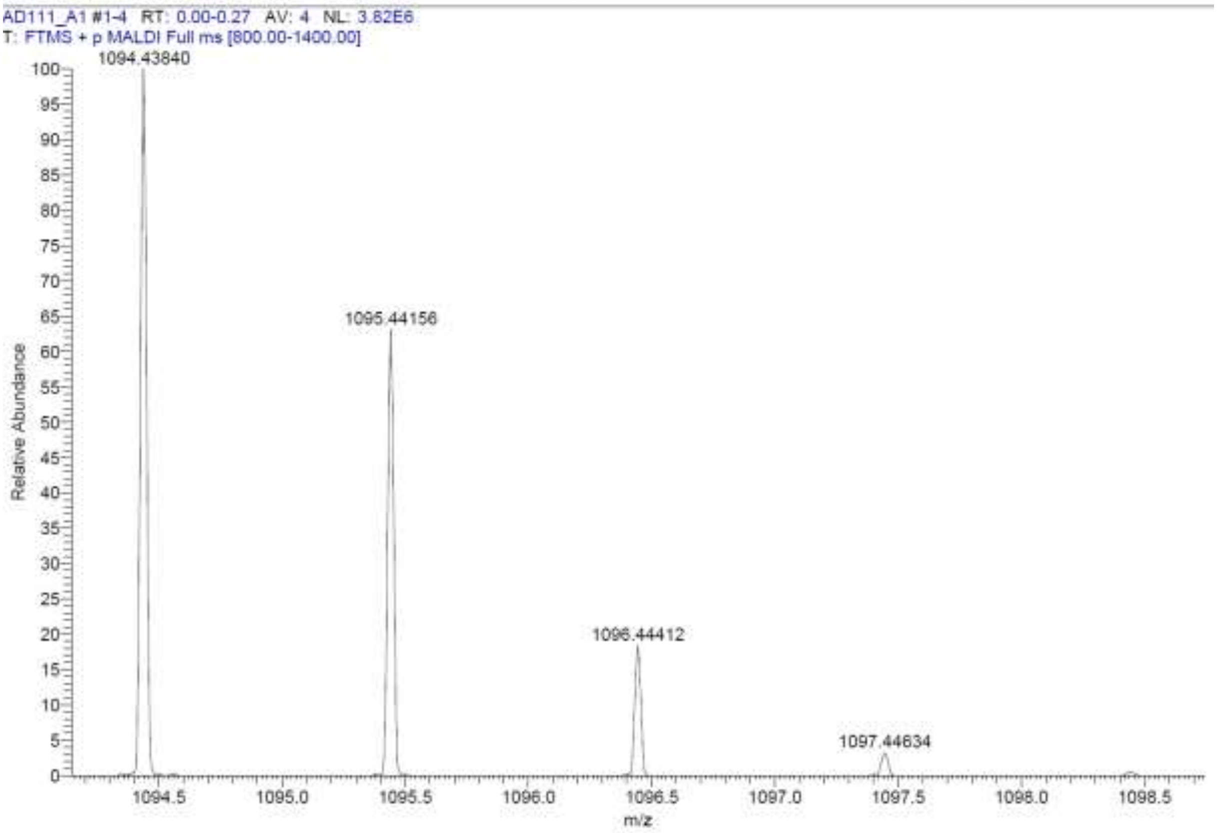

**Signal:** MWD1 A, Sig=254,4 Ref=off

| RT [min] | Type | Width [min] | Area | Height | Area% | Name |
|---|---|---|---|---|---|---|
| 11.319 | VV | 0.2071 | 54018.9102 | 3087.1851 | 100.0000 | |
| | | Sum | 54018.9102 | | | |

**Signal:** MWD1 B, Sig=280,4 Ref=off

| RT [min] | Type | Width [min] | Area | Height | Area% | Name |
|---|---|---|---|---|---|---|
| 11.316 | VV | 0.2211 | 60787.4297 | 3251.7109 | 100.0000 | |
| | | Sum | 60787.4297 | | | |

**Signal:** MWD1 F, Sig=260,4 Ref=off

| RT [min] | Type | Width [min] | Area | Height | Area% | Name |
|---|---|---|---|---|---|---|
| 11.318 | VV | 0.2136 | 59376.8555 | 3289.8892 | 100.0000 | |

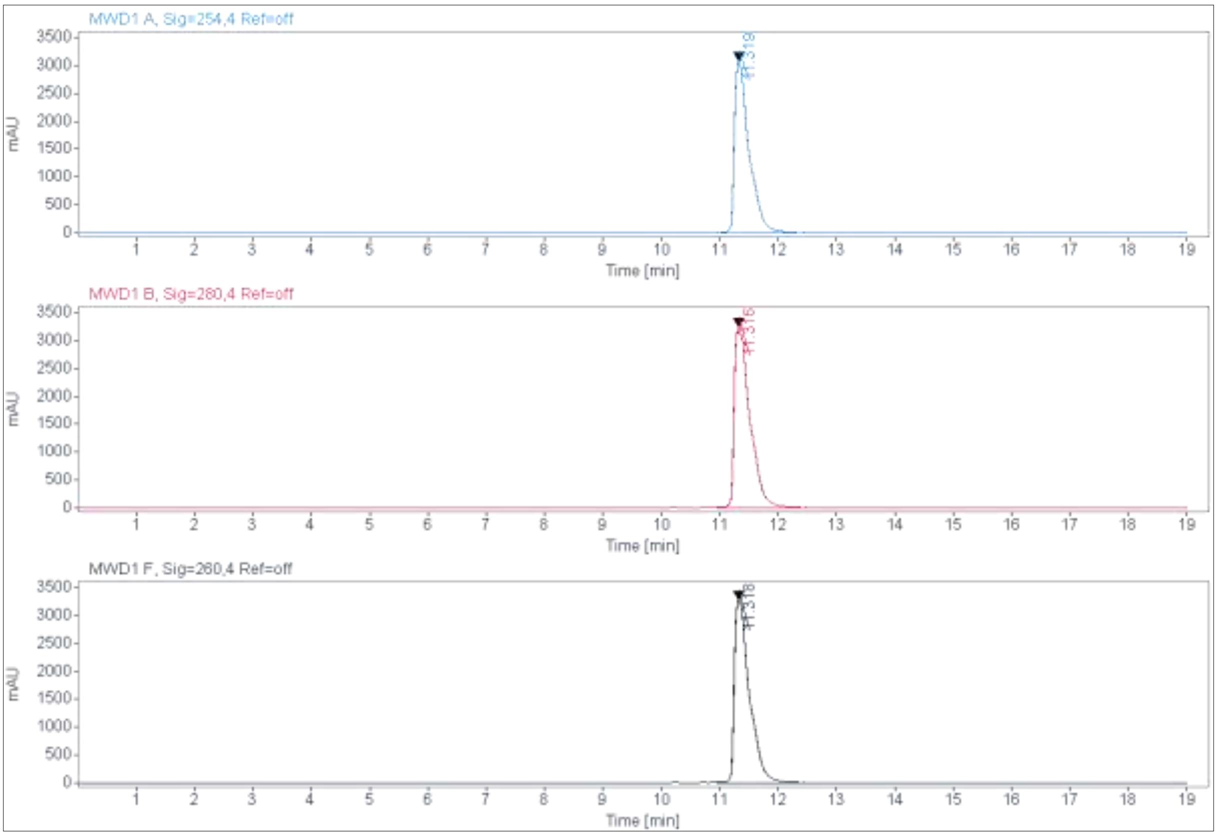

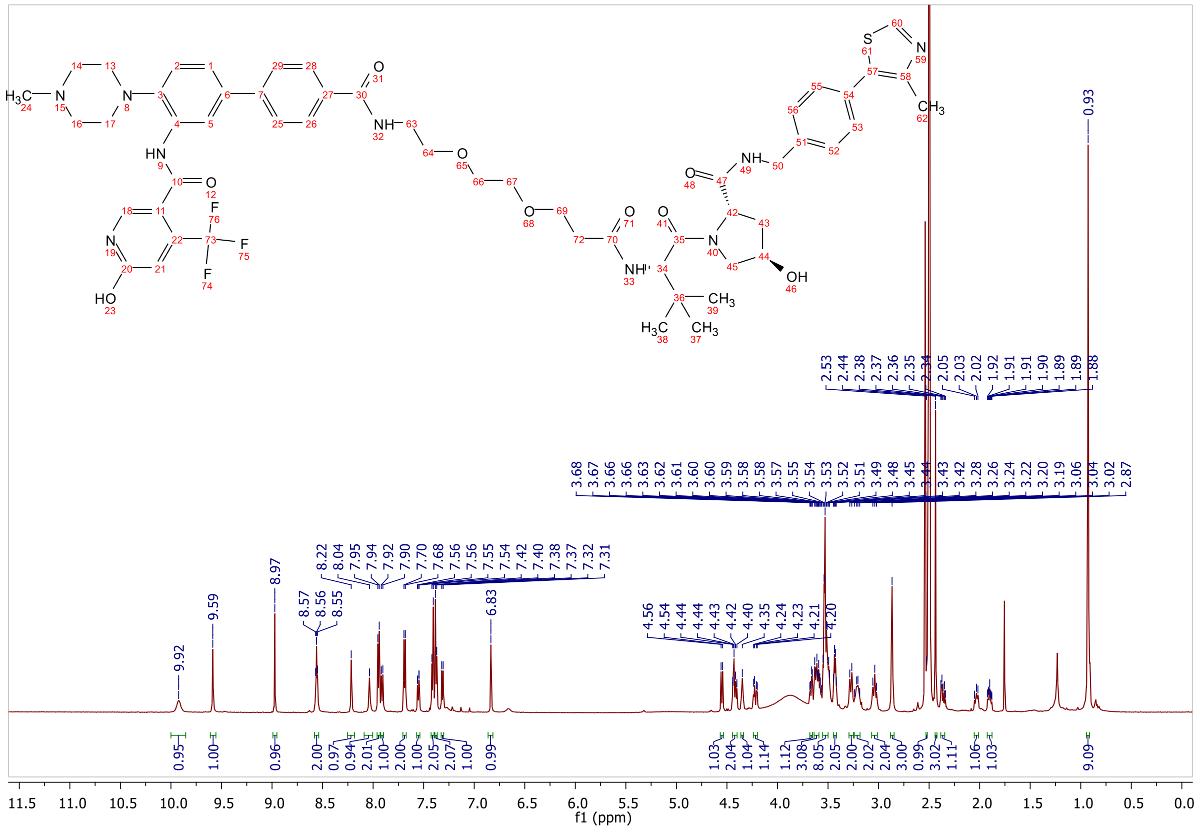
f1 (ppm)

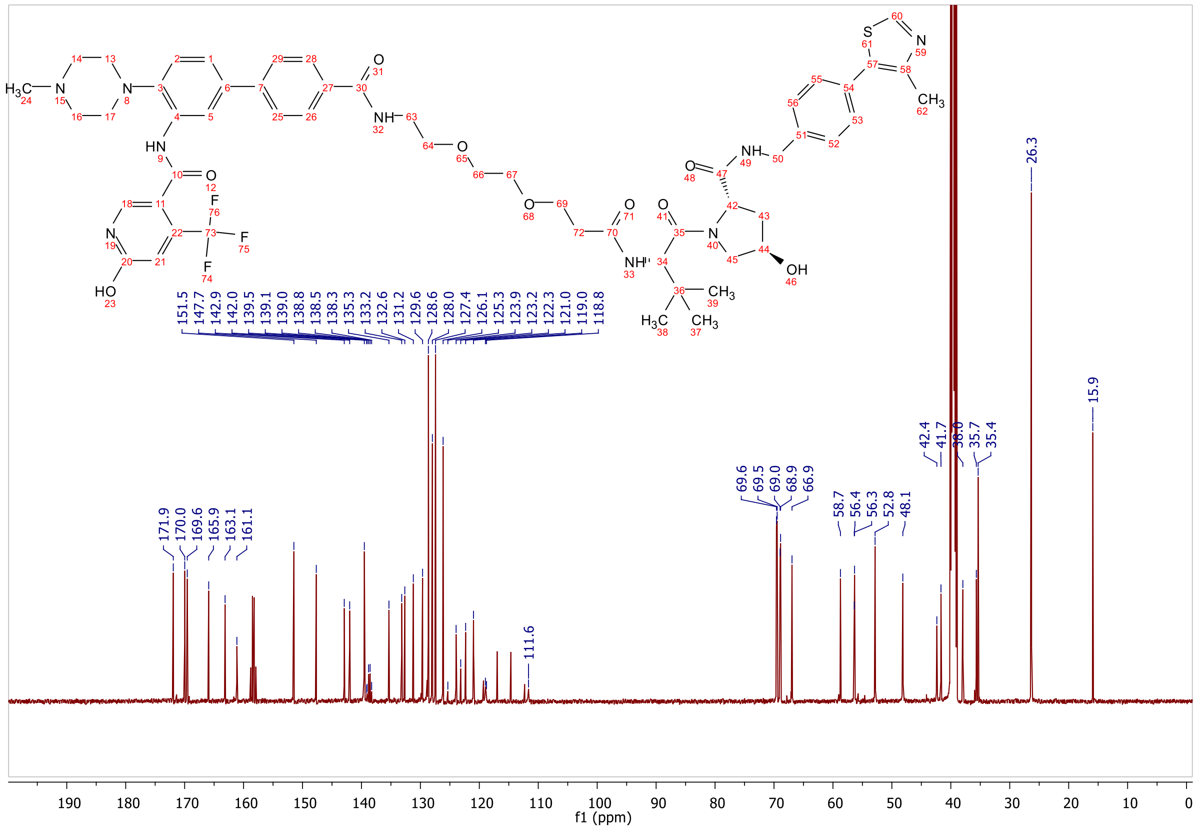
f1 (ppm)

**HRMS, MALDI, HPLC, $^{1}$H-NMR and $^{13}$C-NMR of 6-hydroxy-*N*-(4'-(((*S*)-17-((2*S*,4*R*)-4-hydroxy-2-((4-(4-methylthiazol-5-yl)benzyl)carbamoyl)pyrrolidine-1-carbonyl)-18,18-dimethyl-15-oxo-3,6,9,12-tetraoxa-16-azanonadecyl)carbamoyl)-4-(4-methylpiperazin-1-yl)-[1,1'-biphenyl]-3-yl)-4-(trifluoromethyl)nicotinamide 8c**

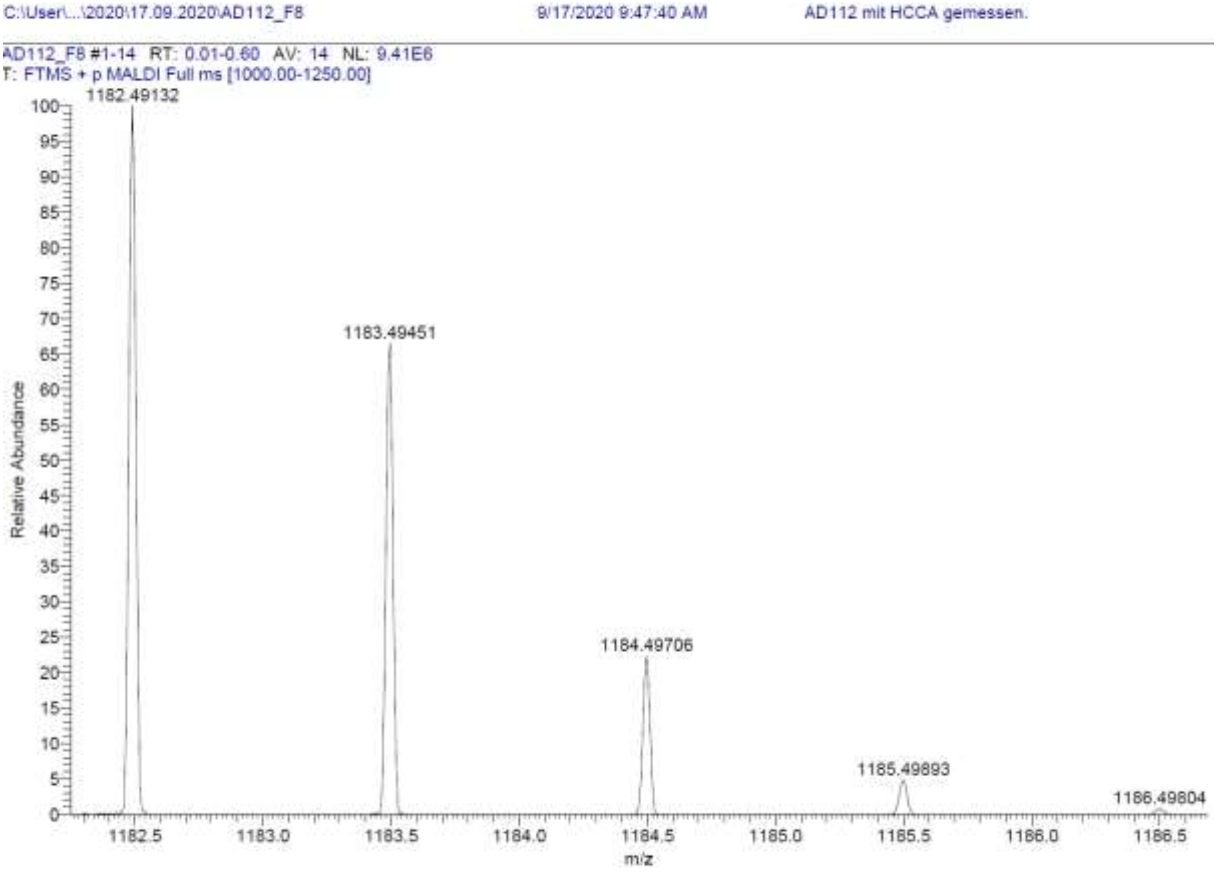

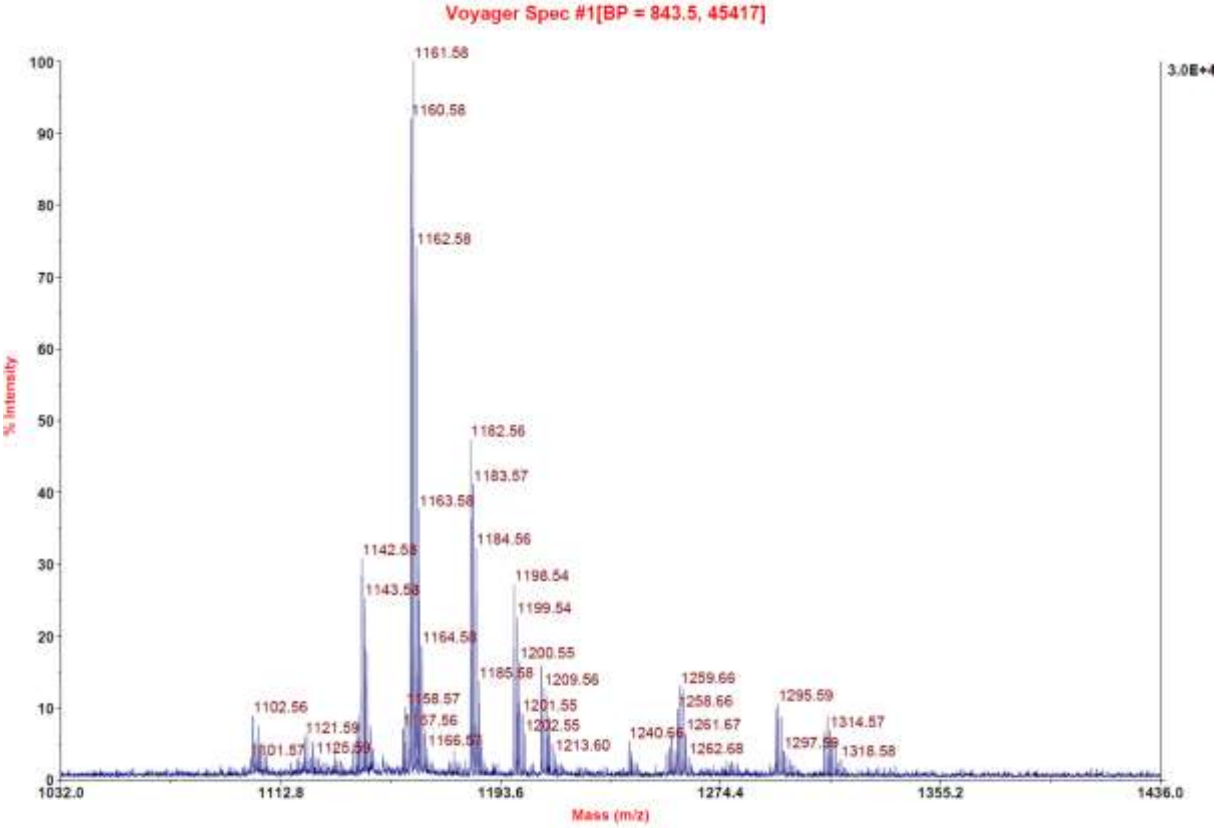

**Signal:** MWD1 A, Sig=254,4 Ref=off

| RT [min] | Type | Width [min] | Area | Height | Area% | Name |
|---|---|---|---|---|---|---|
| 11.445 | VV | 0.2008 | 1150.3380 | 79.0589 | 98.6354 | |
| 12.949 | MM | 0.1184 | 15.9151 | 2.2397 | 1.3646 | |
| | | Sum | 1166.2531 | | | |

**Signal:** MWD1 B, Sig=280,4 Ref=off

| RT [min] | Type | Width [min] | Area | Height | Area% | Name |
|---|---|---|---|---|---|---|
| 11.445 | VV | 0.2055 | 1419.3114 | 94.8723 | 100.0000 | |
| | | Sum | 1419.3114 | | | |

**Signal:** MWD1 F, Sig=260,4 Ref=off

| RT [min] | Type | Width [min] | Area | Height | Area% | Name |
|---|---|---|---|---|---|---|
| 11.445 | VV | 0.2004 | 1305.3274 | 89.3934 | 98.6329 | |
| 12.952 | MM | 0.1277 | 18.0924 | 2.3606 | 1.3671 | |

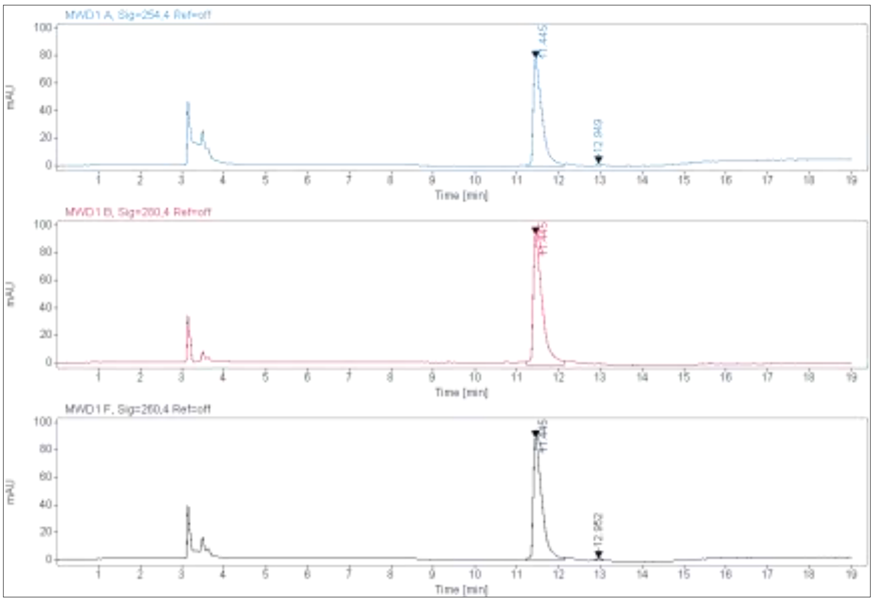

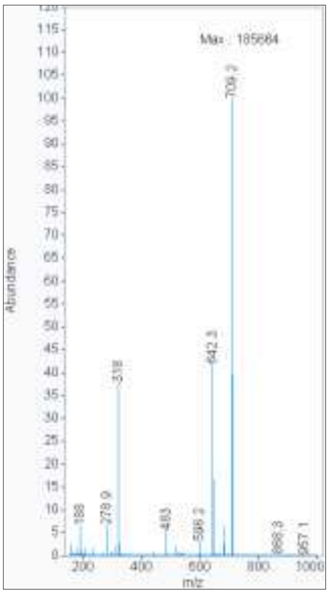

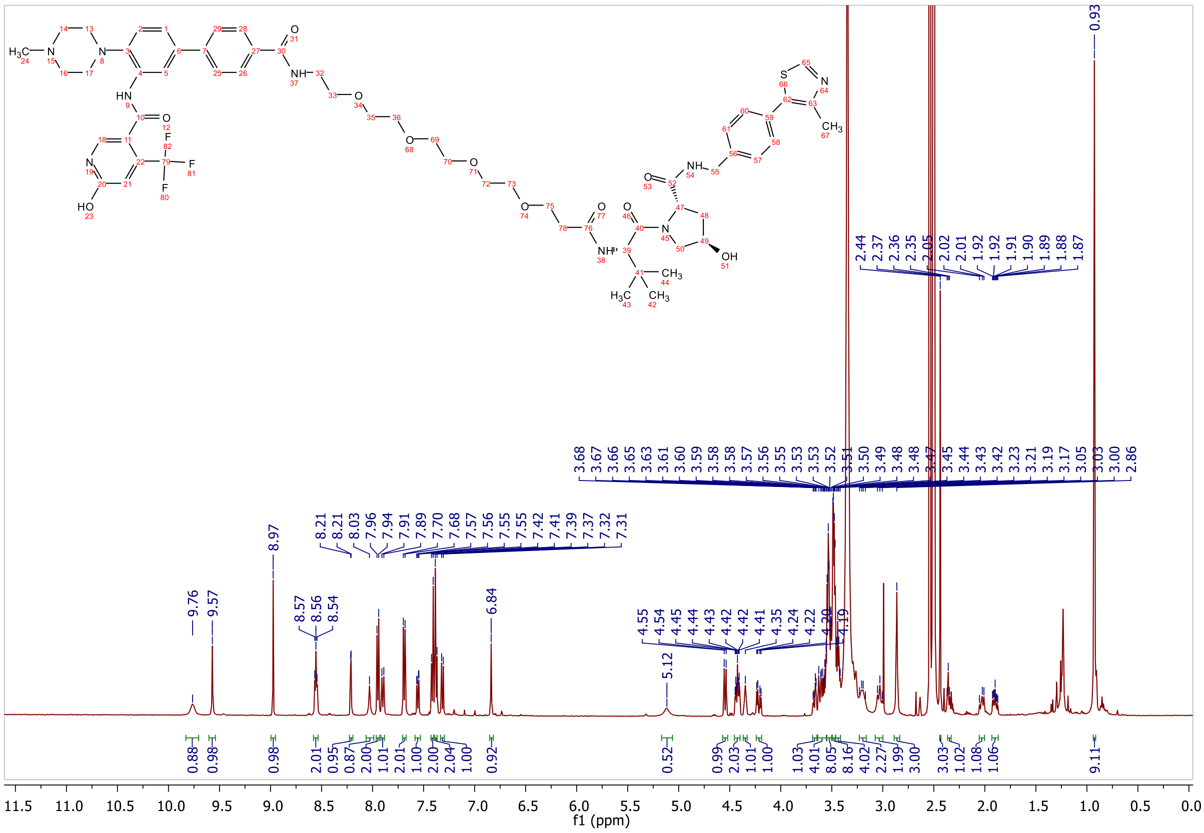

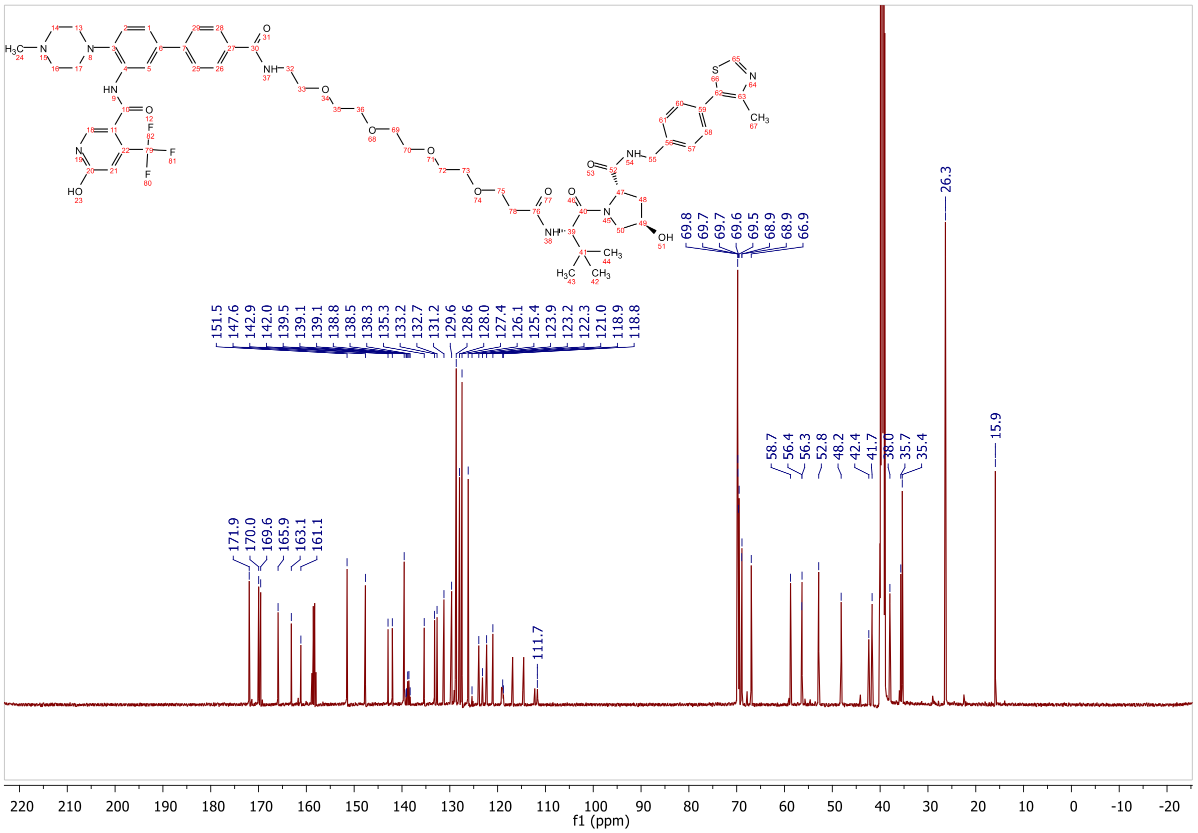

**ESI, MALDI, HRMS, HPLC, $^{1}$H-NMR, $^{13}$C-NMR, $^{13}$C-NMR (downfield region) and $^{1}$H-$^{13}$C-HMBC of 6-hydroxy-N-(4'-(((S)-1-((2S,4R)-4-hydroxy-2-((4-(4-methylthiazol-5-yl)benzyl)carbamoyl)pyrrolidin-1-yl)-3,3-dimethyl-1-oxobutan-2-yl)carbamoyl)-4-(4-methylpiperazin-1-yl)-[1,1'-biphenyl]-3-yl)-4-(trifluoromethyl)nicotinamide 8d**

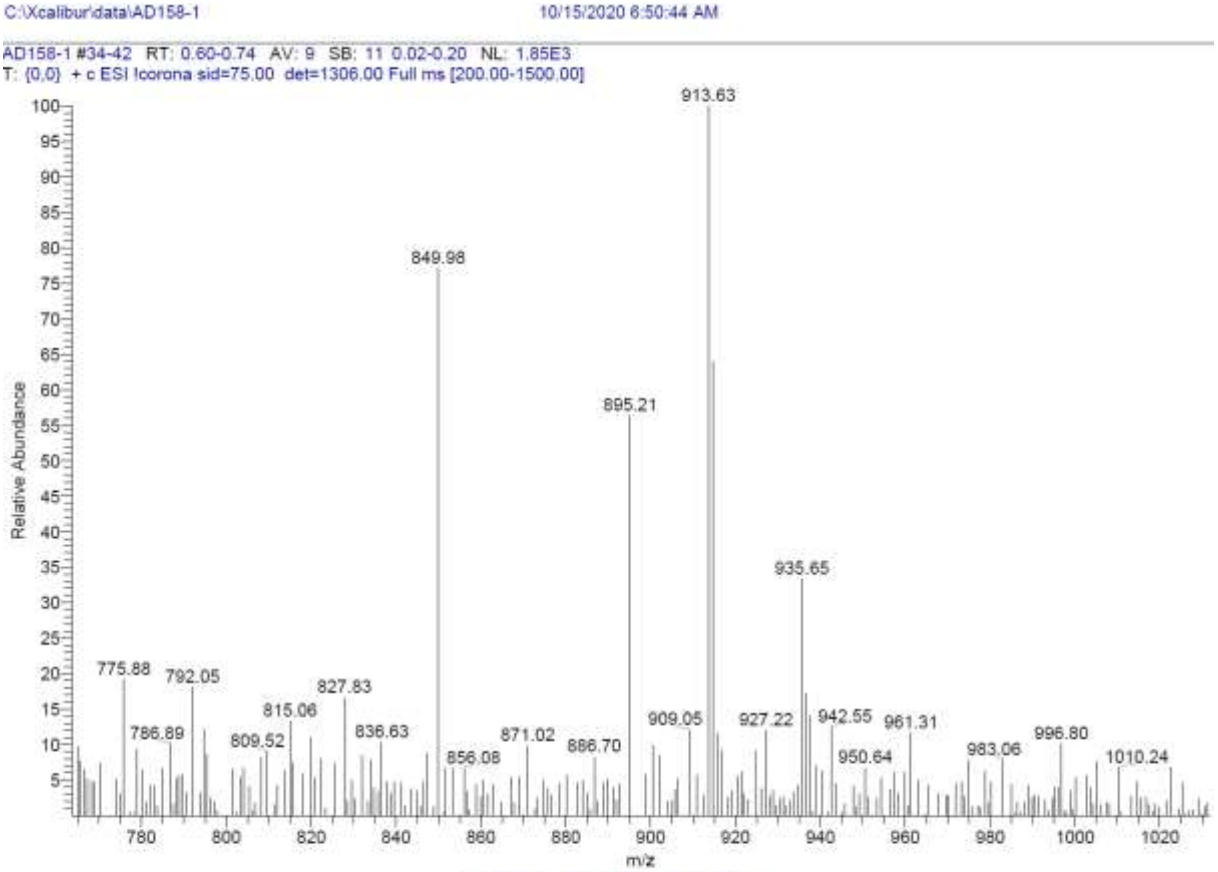

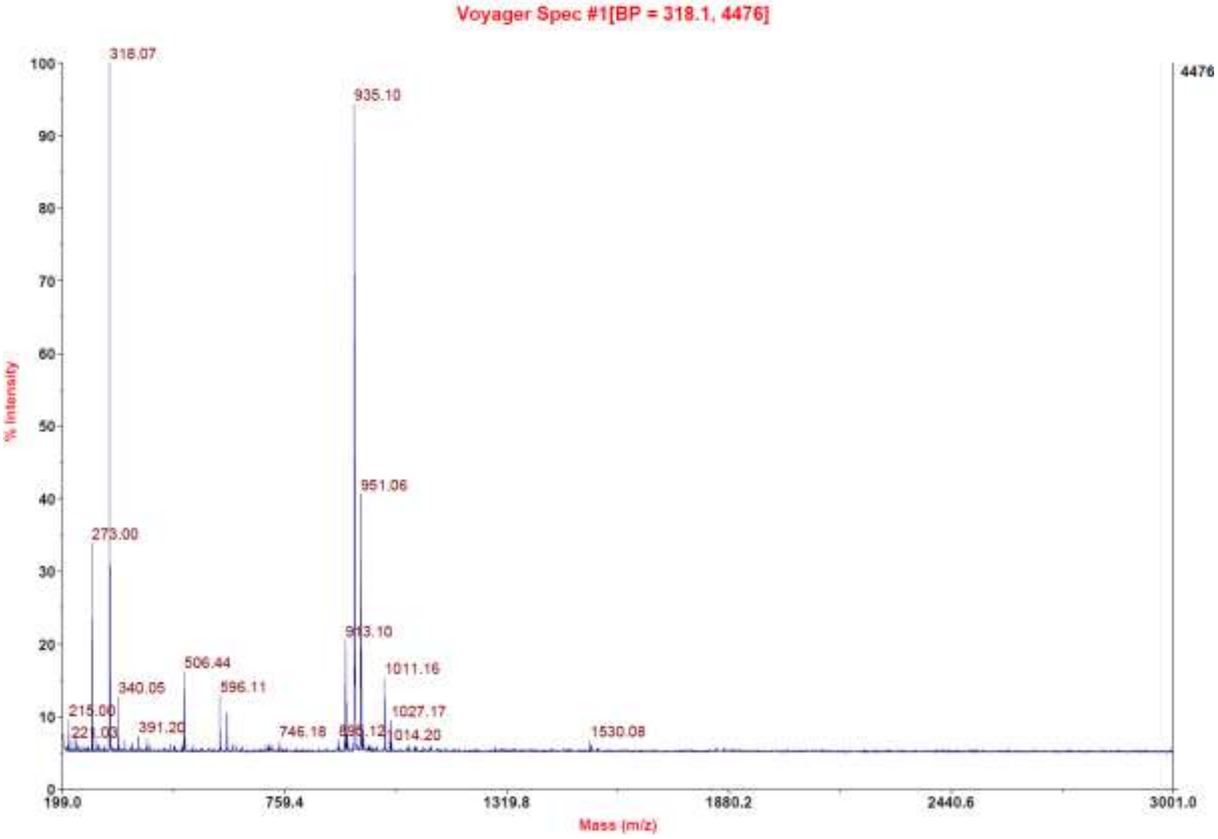

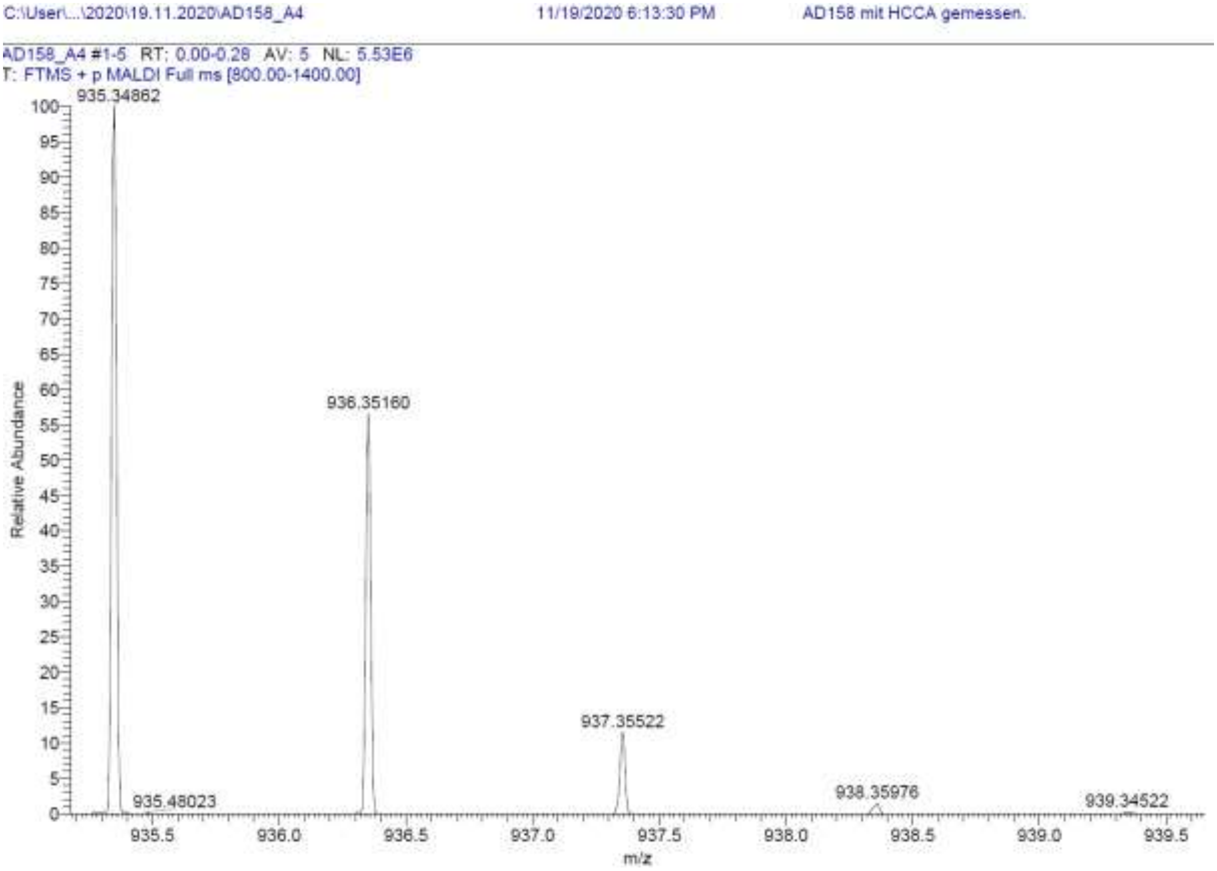

| RT [min] | Type | Width [min] | Area | Height | Area% |
|---|---|---|---|---|---|
| 10.184 | VV | 0.2023 | 725.7850 | 53.8351 | 1.6325 |
| 11.567 | VV | 0.2195 | 42875.0313 | 2667.2349 | 96.4396 |
| 12.127 | VV | 0.1624 | 857.0851 | 76.6709 | 1.9279 |
| | | Sum | 44457.9013 | | |

**Signal:** MWD1 B, Sig=280,4 Ref=off

| RT [min] | Type | Width [min] | Area | Height | Area% |
|---|---|---|---|---|---|
| 10.181 | VV | 0.2005 | 1076.0703 | 79.6896 | 2.0573 |
| 11.558 | VV | 0.1974 | 50273.9727 | 3016.5632 | 96.1188 |
| 12.123 | VV | 0.1574 | 953.9785 | 88.0350 | 1.8239 |
| | | Sum | 52304.0214 | | |

**Signal:** MWD1 F, Sig=260,4 Ref=off

| RT [min] | Type | Width [min] | Area | Height | Area% |
|---|---|---|---|---|---|
| 10.184 | VV | 0.2011 | 783.1274 | 58.1524 | 1.5765 |
| 11.563 | VV | 0.2178 | 47934.4141 | 2930.3010 | 96.4971 |
| 12.127 | VV | 0.1643 | 956.9185 | 85.6613 | 1.9264 |

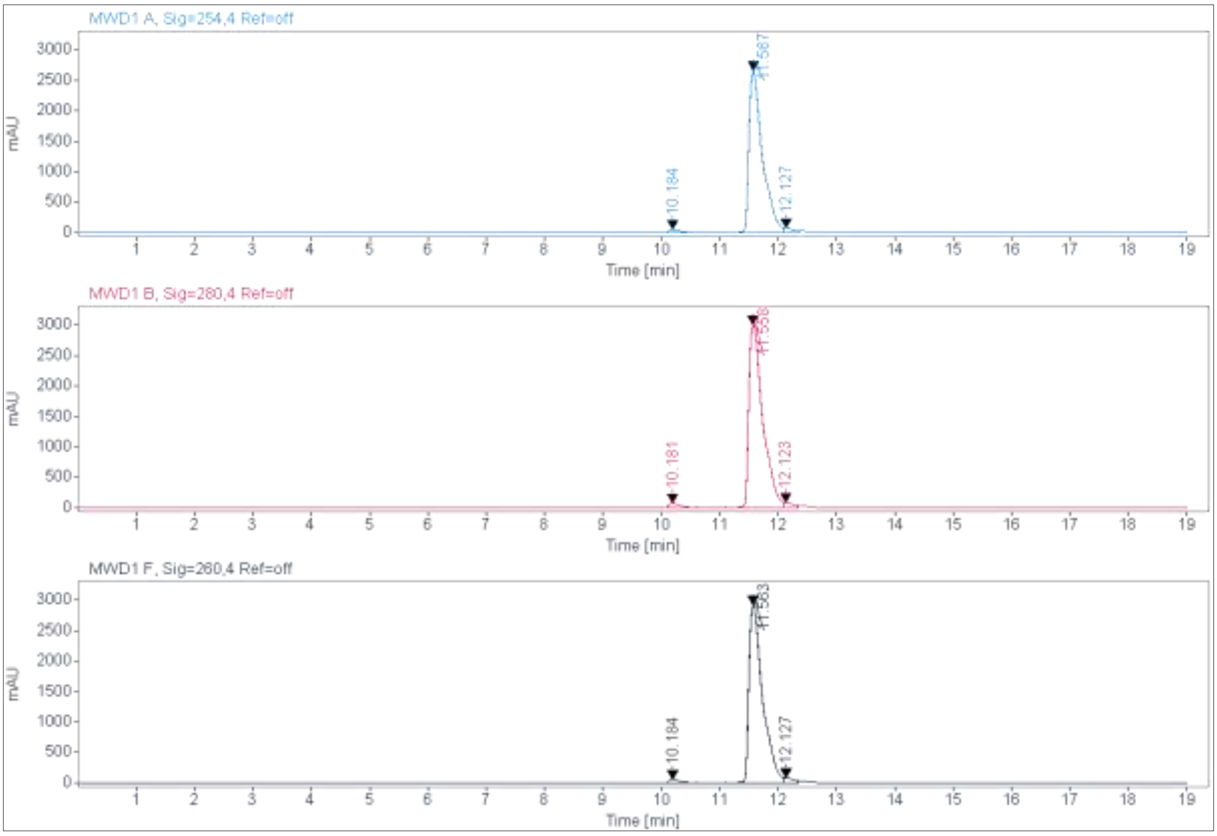
MWD1 A, Sig=254,4 Ref=off
mAU
10.184
11.587
12.127
Time [min]
MWD1 B, Sig=280,4 Ref=off
mAU
10.181
11.558
12.123
Time [min]
MWD1 F, Sig=260,4 Ref=off
mAU
10.184
11.563
12.127
Time [min]

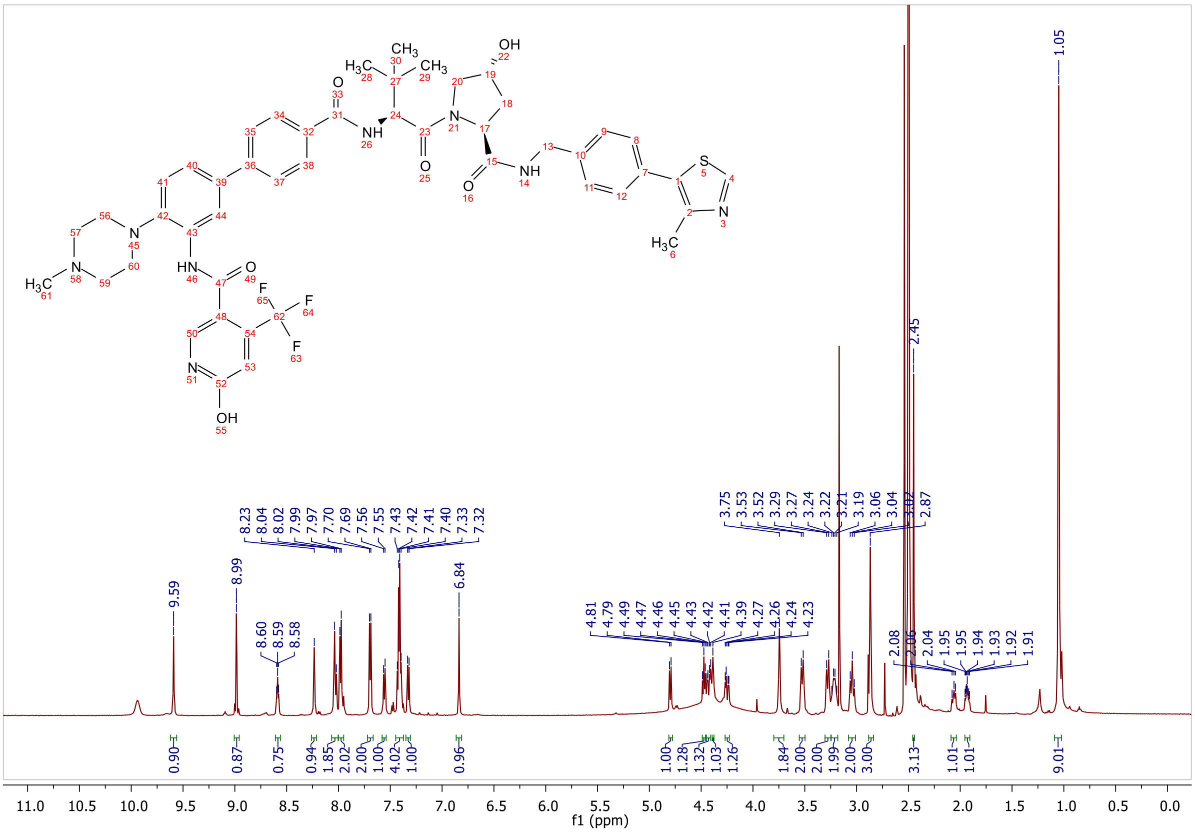
9.59
8.99
8.60 8.59 8.58
8.23 8.04 8.02 7.99 7.97 7.70 7.69 7.56 7.55 7.43 7.42 7.41 7.40 7.33 7.32
6.84
4.81 4.79 4.49 4.47 4.46 4.45 4.43 4.42 4.41 4.39 4.27 4.26 4.24 4.23
3.75 3.53 3.52 3.29 3.27 3.24 3.22 3.21 3.19 3.06 3.04 3.02 2.87
2.45
2.08 2.06 2.04 1.95 1.95 1.94 1.93 1.92 1.91
1.05
0.90 0.87 0.75 0.94 1.85 2.02 2.00 1.00 1.00 4.02 1.00 0.96
1.00 1.28 1.32 1.03 1.26 1.84 2.00 2.00 1.99 2.00 3.00 3.13 1.01 1.01 9.01
f1 (ppm)

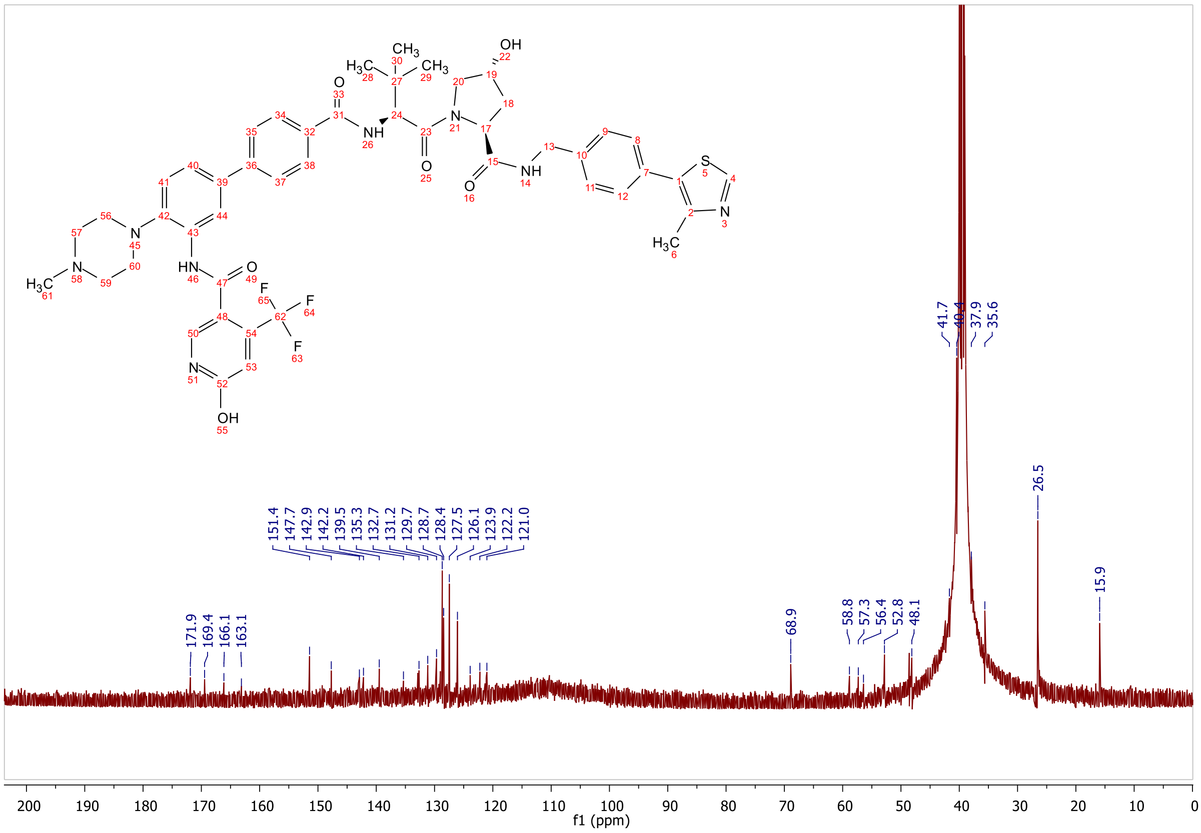
171.9
169.4
166.1
163.1
151.4
147.7
142.9
142.2
139.5
135.3
132.7
131.2
129.7
128.7
128.4
127.5
126.1
123.9
122.2
121.0
68.9
58.8
57.3
56.4
52.8
48.1
41.7
40.4
37.9
35.6
26.5
15.9
f1 (ppm)

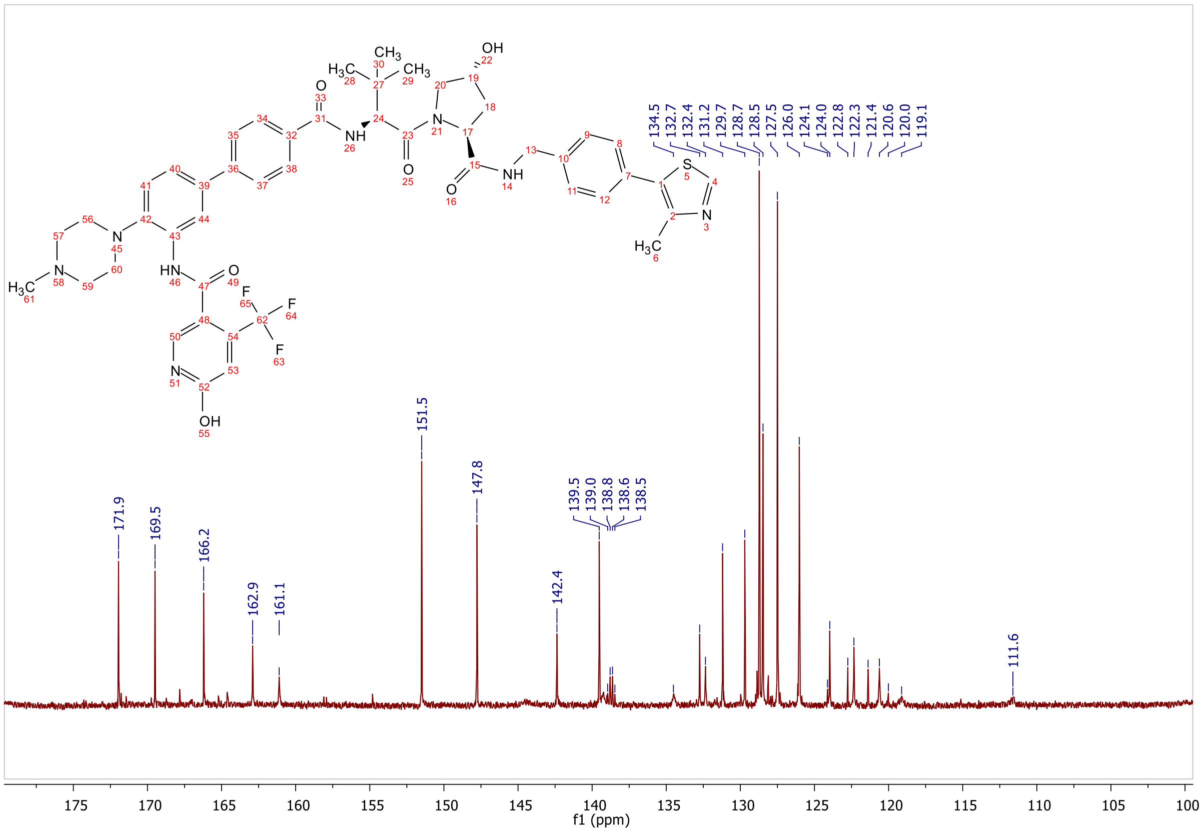
171.9
169.5
166.2
162.9
161.1
151.5
147.8
142.4
139.5
139.0
138.8
138.6
138.5
134.5
132.7
132.4
131.2
129.7
128.7
128.5
127.5
126.0
124.1
124.0
122.8
122.3
121.4
120.6
120.0
119.1
111.6
f1 (ppm)

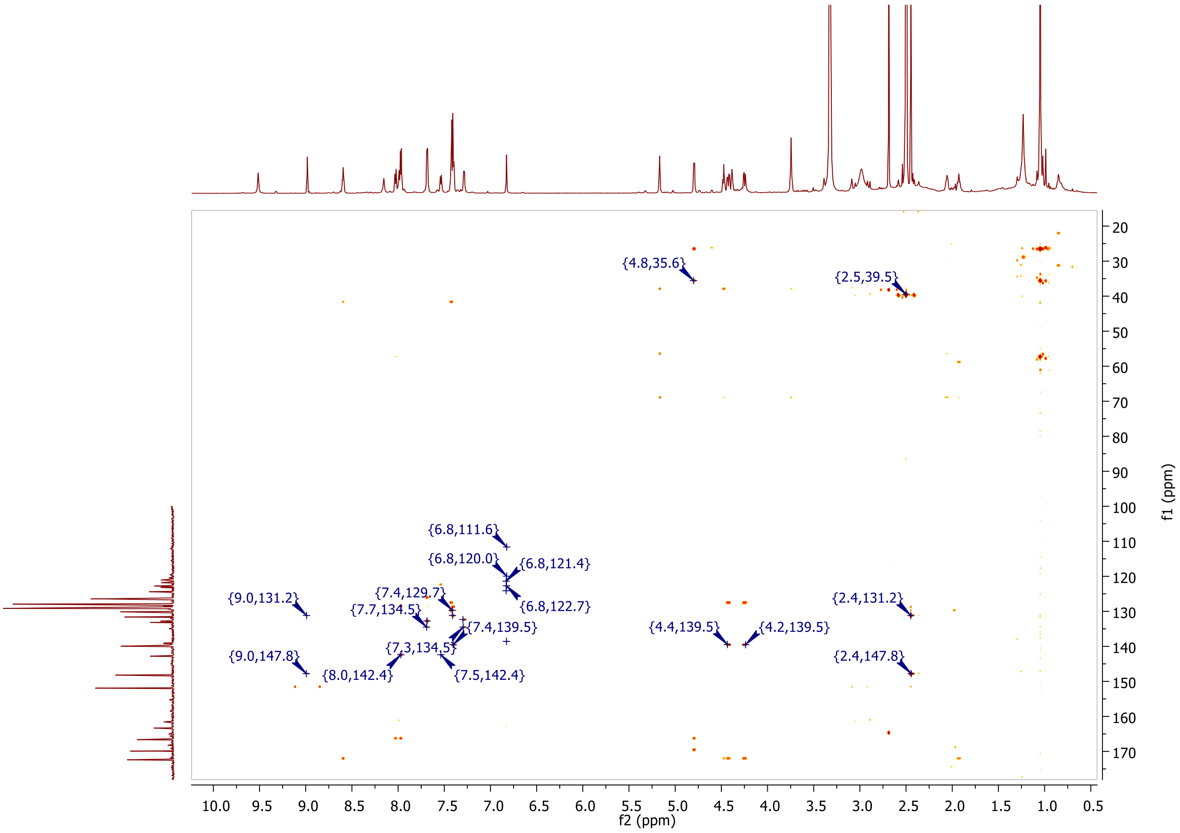
{4.8,35.6}
{2.5,39.5}
{6.8,111.6}
{6.8,120.0}
{6.8,121.4}
{9.0,131.2}
{7.4,129.7}
{7.7,134.5}
{6.8,122.7}
{7.4,139.5}
{7.3,134.5}
{9.0,147.8}
{8.0,142.4}
{7.5,142.4}
{4.4,139.5}
{4.2,139.5}
{2.4,131.2}
{2.4,147.8}
f2 (ppm)
f1 (ppm)

**MALDI, HRMS, HPLC, $^{1}$H-NMR and $^{13}$C-NMR of 6-hydroxy-N-(4'-((3-(((S)-1-((2S,4R)-4-hydroxy-2-((4-(4-methylthiazol-5-yl)benzyl)carbamoyl)pyrrolidin-1-yl)-3,3-dimethyl-1-oxobutan-2-yl)amino)-3-oxopropyl)carbamoyl)-4-(4-methylpiperazin-1-yl)-[1,1'-biphenyl]-3-yl)-4-(trifluoromethyl)nicotinamide 8e**

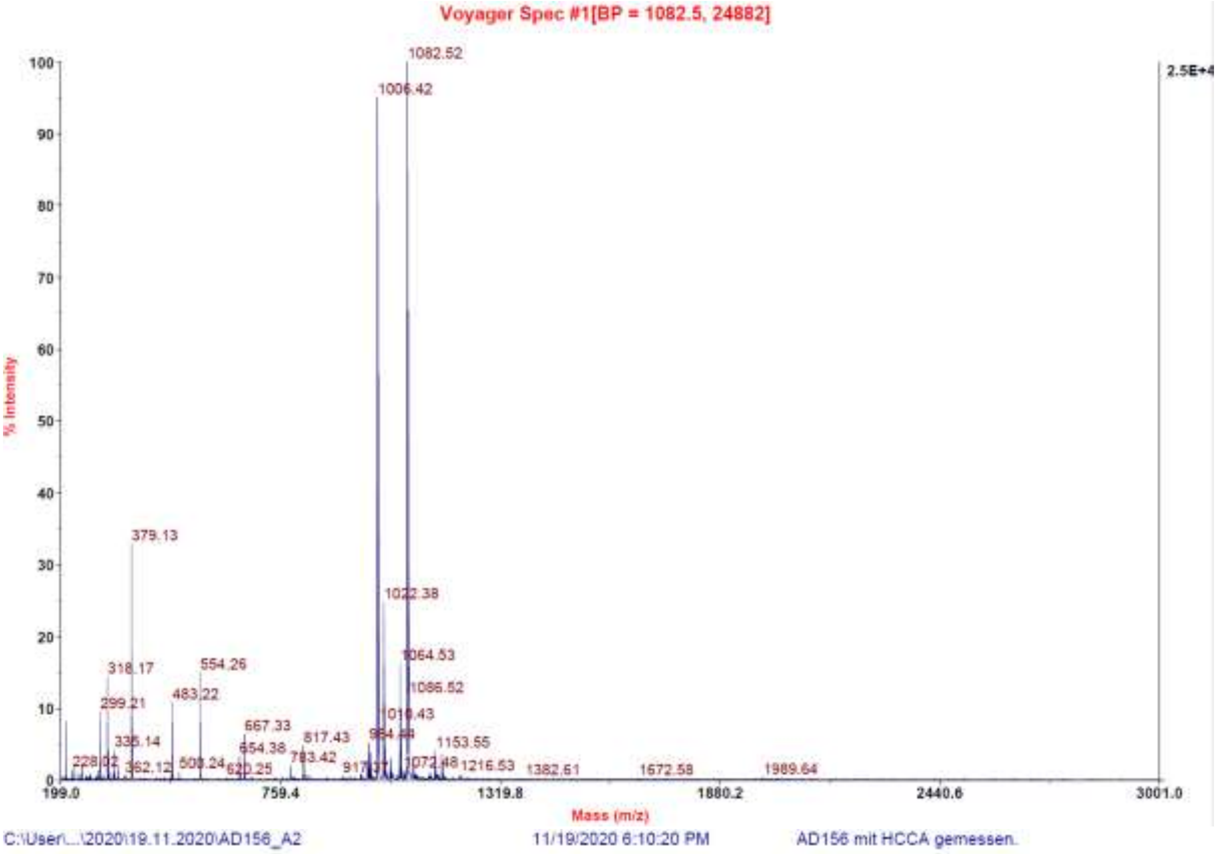

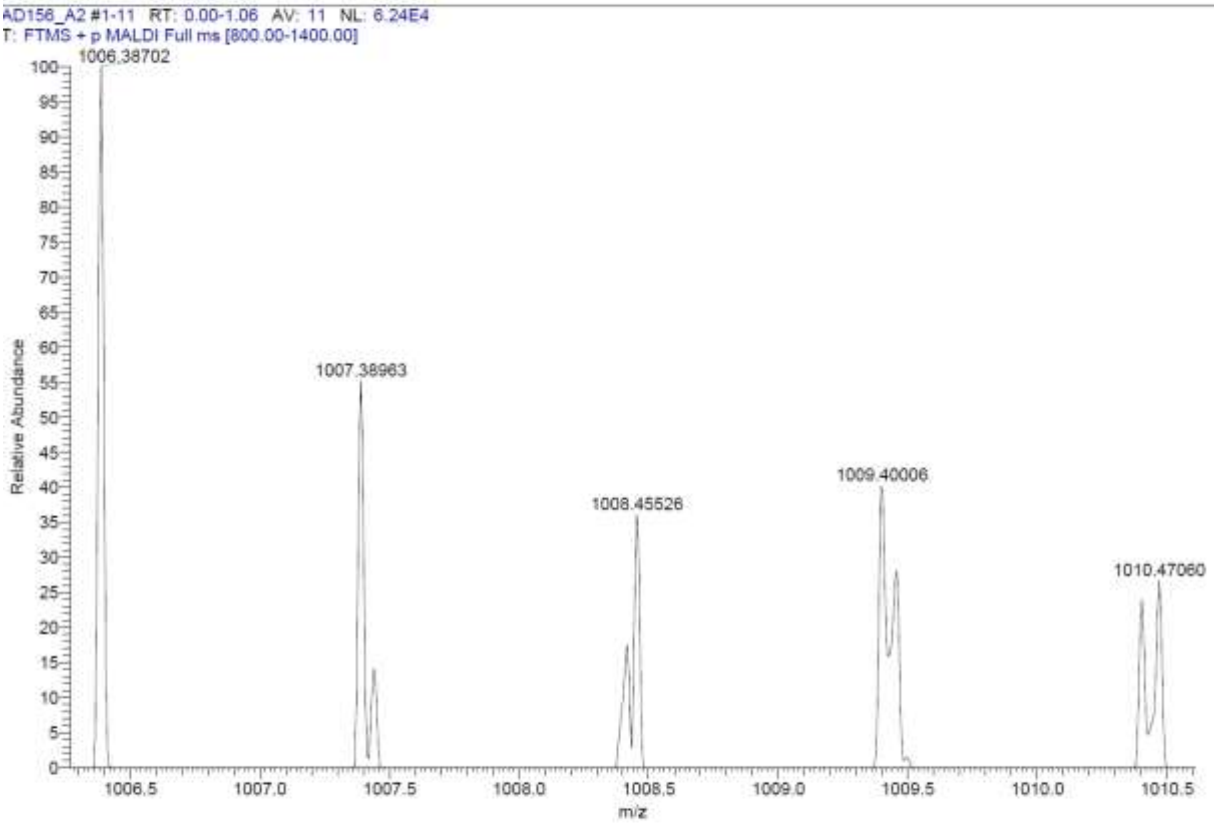

**Signal:** MWD1 A, Sig=254,4 Ref=off

| RT [min] | Type | Width [min] | Area | Height | Area% | Name |
|---|---|---|---|---|---|---|
| 11.239 | VV | 0.2244 | 10229.9883 | 672.8884 | 100.0000 | |
| | | Sum | 10229.9883 | | | |

**Signal:** MWD1 B, Sig=280,4 Ref=off

| RT [min] | Type | Width [min] | Area | Height | Area% | Name |
|---|---|---|---|---|---|---|
| 11.239 | VV | 0.2255 | 12384.9932 | 809.5344 | 100.0000 | |
| | | Sum | 12384.9932 | | | |

**Signal:** MWD1 F, Sig=260,4 Ref=off

| RT [min] | Type | Width [min] | Area | Height | Area% | Name |
|---|---|---|---|---|---|---|
| 11.239 | VV | 0.2257 | 11558.8154 | 758.7136 | 100.0000 | |

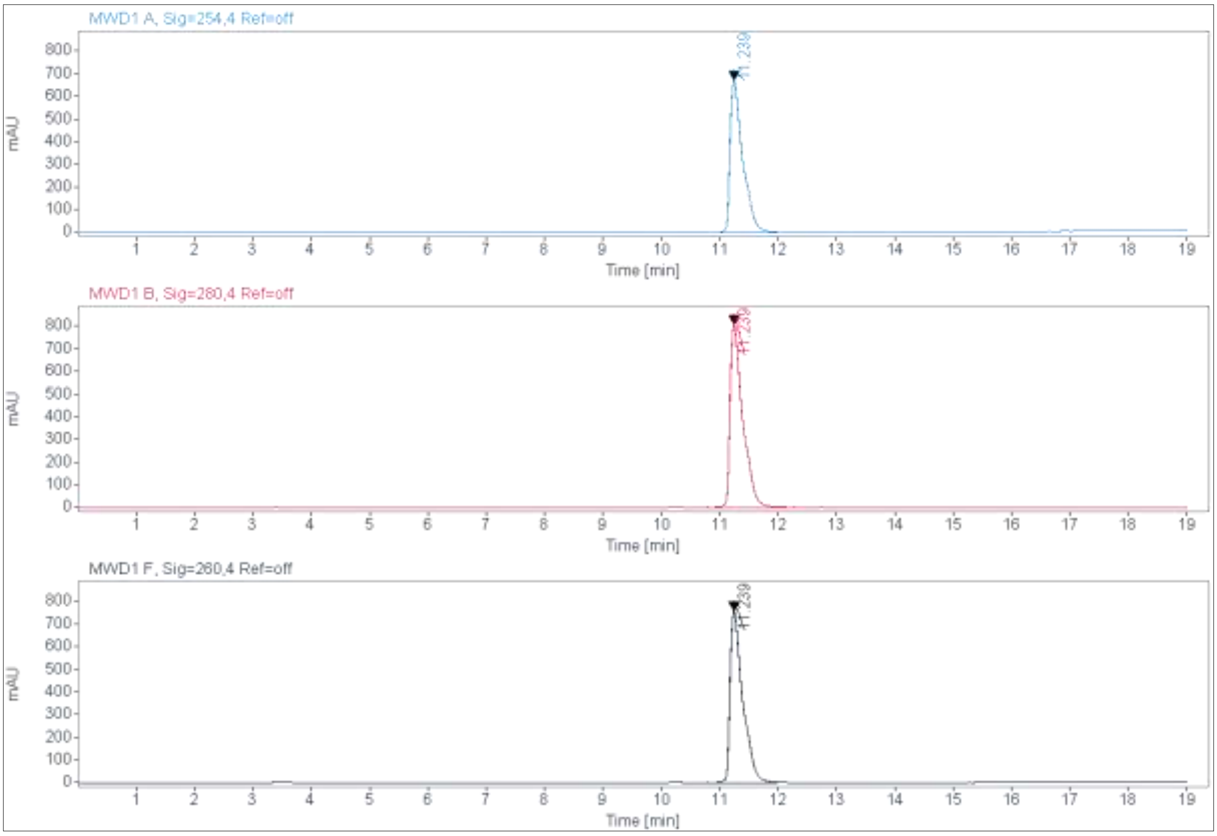

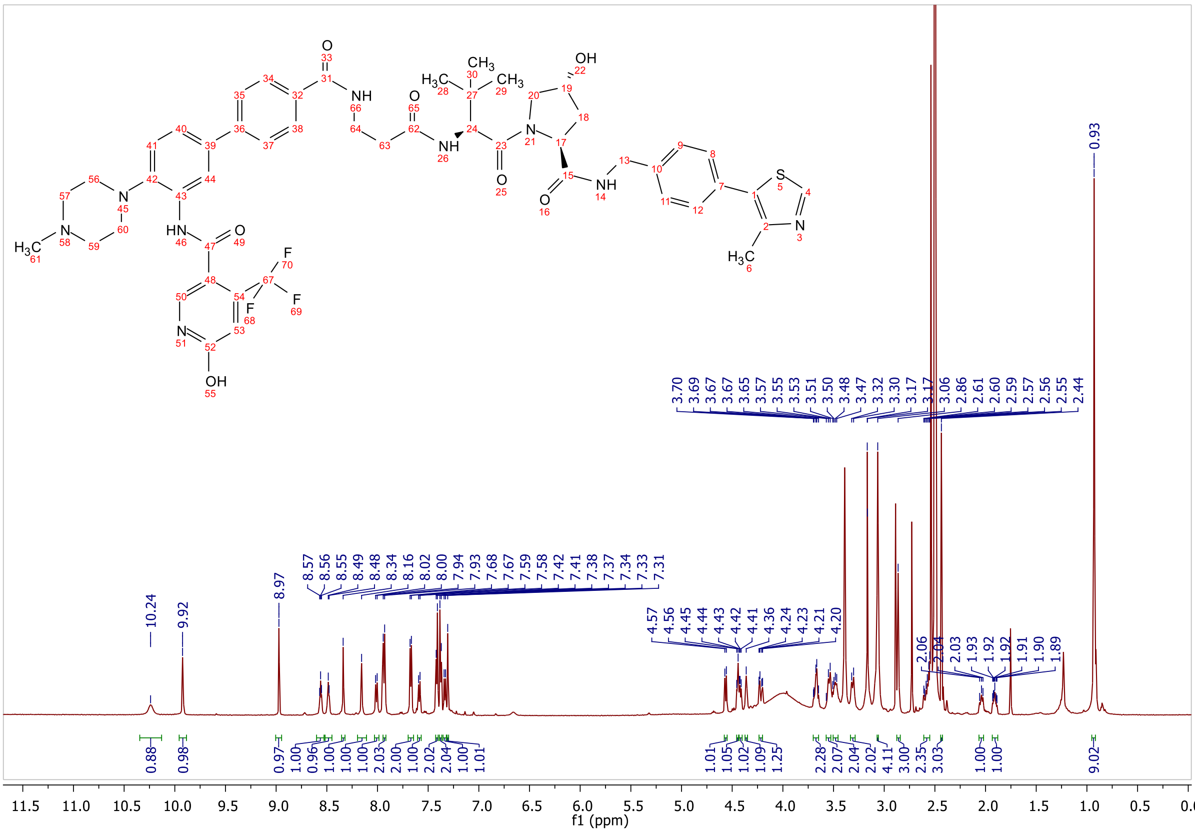
3.70 3.69 3.67 3.67 3.65 3.57 3.55 3.53 3.51 3.50 3.48 3.47 3.32 3.30 3.17 3.17 3.06 2.86 2.61 2.60 2.59 2.57 2.56 2.55 2.44
0.93
10.24 9.92 8.97 8.57 8.56 8.55 8.49 8.48 8.34 8.16 8.02 8.00 7.94 7.93 7.68 7.67 7.59 7.58 7.42 7.41 7.38 7.37 7.34 7.33 7.31
4.57 4.56 4.45 4.44 4.43 4.42 4.41 4.36 4.24 4.23 4.21 4.20
2.06 2.04 2.03 1.93 1.92 1.92 1.91 1.90 1.89
0.88 0.98 0.97 1.00 0.96 1.00 1.00 1.00 2.03 2.00 1.00 2.02 2.04 1.00 1.01
1.01 1.05 1.02 1.09 1.25 2.28 2.07 2.04 2.02 4.11 3.00 2.35 3.03 1.00 1.00 9.02
f1 (ppm)

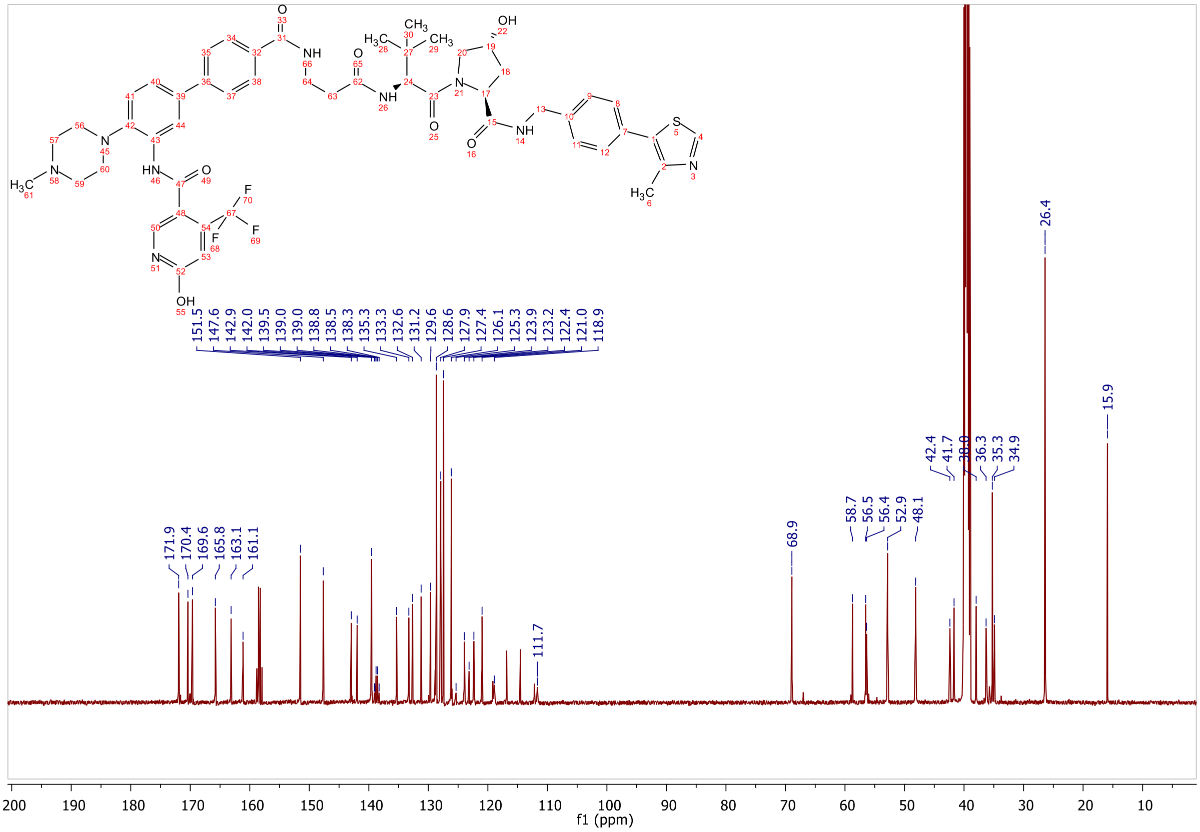
151.5 147.6 142.9 142.0 139.5 139.0 138.8 138.5 138.3 135.3 133.3 132.6 131.2 129.6 128.6 127.9 127.4 126.1 125.3 123.9 123.2 122.4 121.0 118.9
171.9 170.4 169.6 165.8 163.1 161.1
111.7
68.9
58.7 56.5 56.4 52.9 48.1
42.4 41.7 38.0 36.3 35.3 34.9
26.4
15.9
f1 (ppm)

**ESI, MALDI, HRMS, HPLC, $^{1}$H-NMR, $^{13}$C-NMR, $^{13}$C-NMR (downfield region), $^{1}$H-$^{13}$C-HSQC (downfield), $^{1}$H-$^{13}$C-HSQC (upfield), $^{1}$H-$^{13}$C-HMBC (downfield) and $^{1}$H-$^{13}$C-HMBC (upfield) of 6-hydroxy-N-(4'-((4-(((S)-1-((2S,4R)-4-hydroxy-2-((4-(4-methylthiazol-5-yl)benzyl)carbamoyl)pyrrolidin-1-yl)-3,3-dimethyl-1-oxobutan-2-yl)amino)-4-oxobutyl)carbamoyl)-4-(4-methylpiperazin-1-yl)-[1,1'-biphenyl]-3-yl)-4-(trifluoromethyl)nicotinamide 8f**

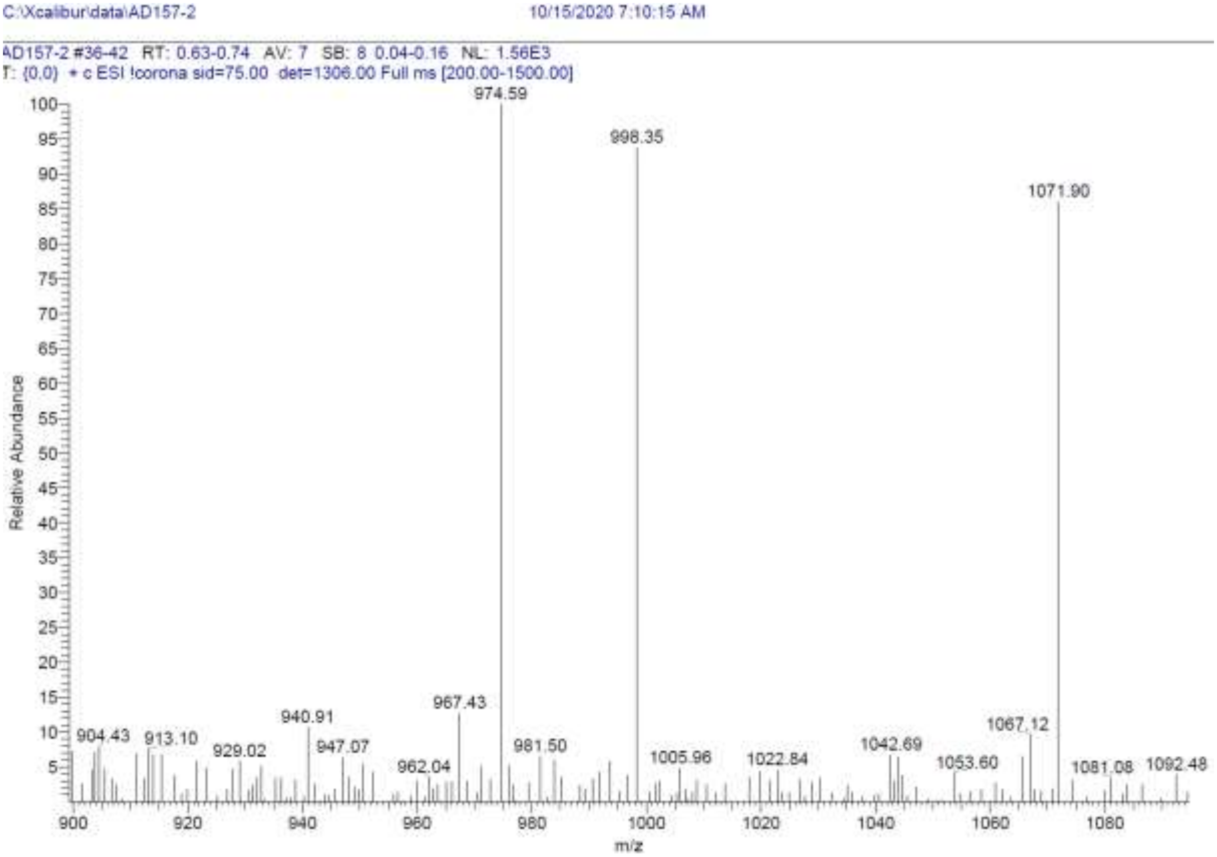

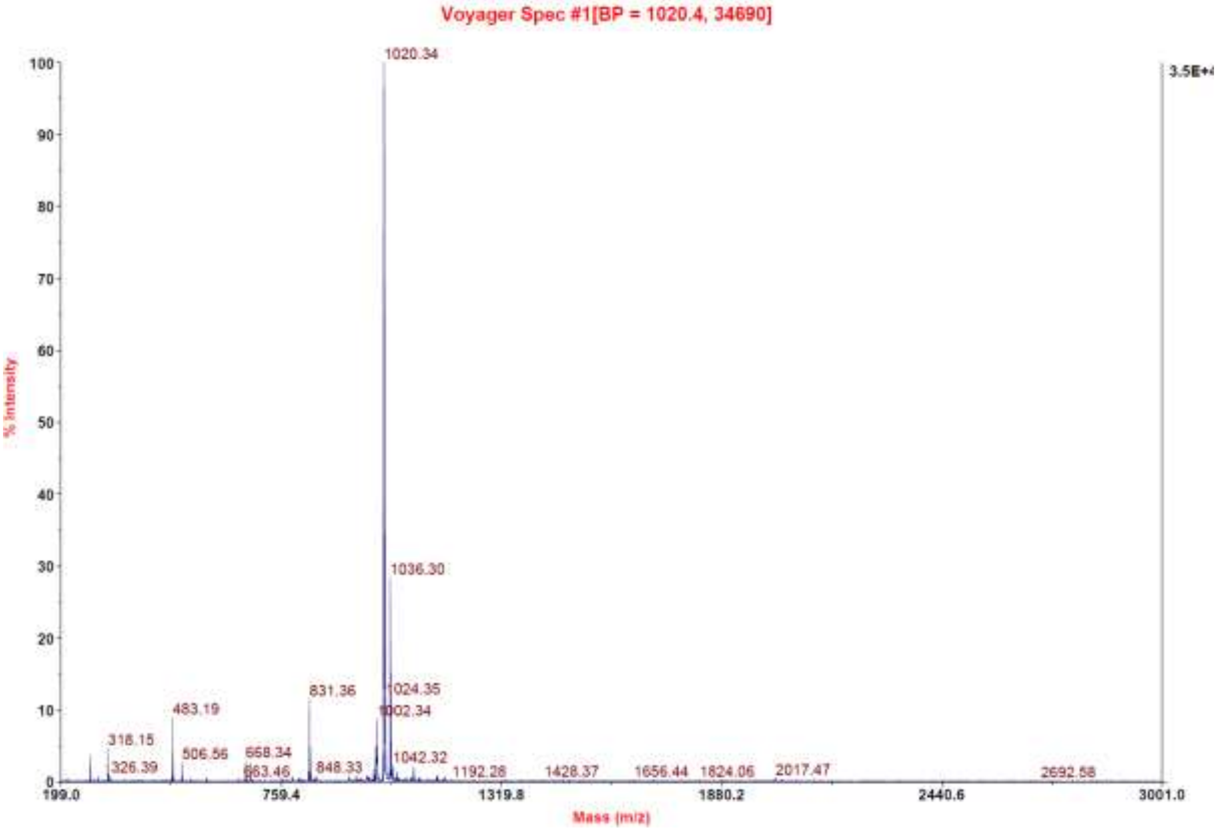

C:\User\...\2020\19.11.2020\AD157_A3 11/19/2020 6:12:15 PM AD157 mit HCCA gemessen.

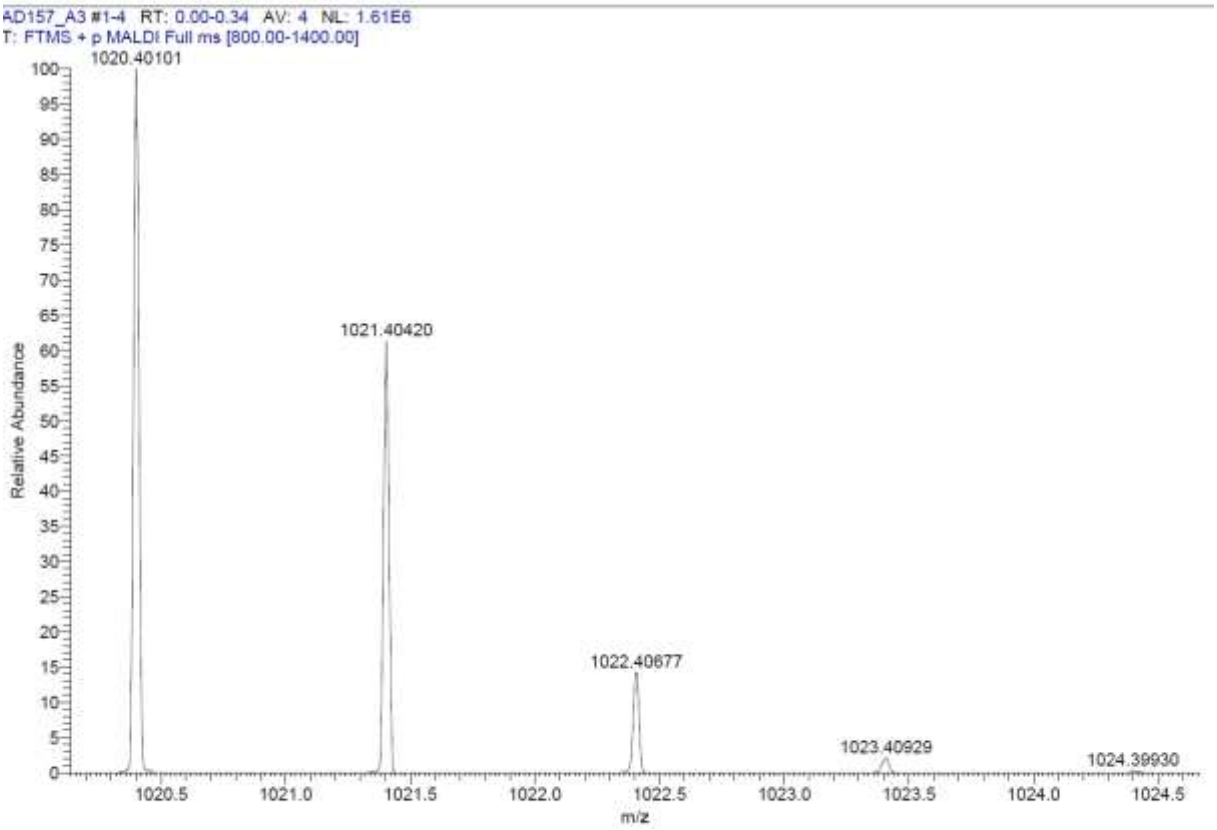

**Signal:** MWD1 A, Sig=254,4 Ref=off

| RT [min] | Type | Width [min] | Area | Height | Area% | Name |
|---|---|---|---|---|---|---|
| 11.279 | VV | 0.2370 | 9289.1738 | 585.9404 | 100.0000 | |
| | | Sum | 9289.1738 | | | |

**Signal:** MWD1 B, Sig=280,4 Ref=off

| RT [min] | Type | Width [min] | Area | Height | Area% | Name |
|---|---|---|---|---|---|---|
| 11.279 | VV | 0.2382 | 10607.9912 | 664.5951 | 100.0000 | |
| | | Sum | 10607.9912 | | | |

**Signal:** MWD1 F, Sig=260,4 Ref=off

| RT [min] | Type | Width [min] | Area | Height | Area% | Name |
|---|---|---|---|---|---|---|
| 11.279 | VV | 0.2376 | 10198.9717 | 641.1873 | 100.0000 | |

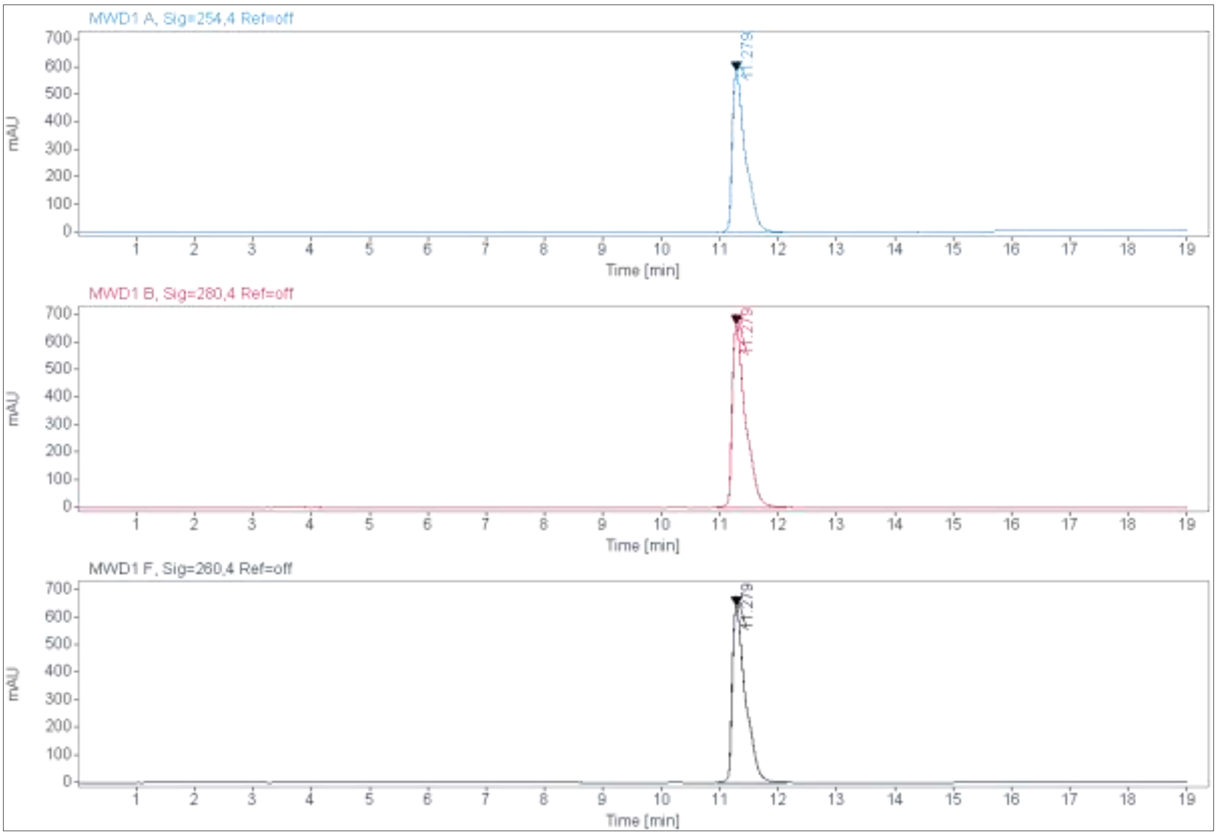
MWD1 A, Sig=254,4 Ref=off
mAU
Time [min]
11.279
MWD1 B, Sig=280,4 Ref=off
mAU
Time [min]
11.279
MWD1 F, Sig=260,4 Ref=off
mAU
Time [min]
11.279

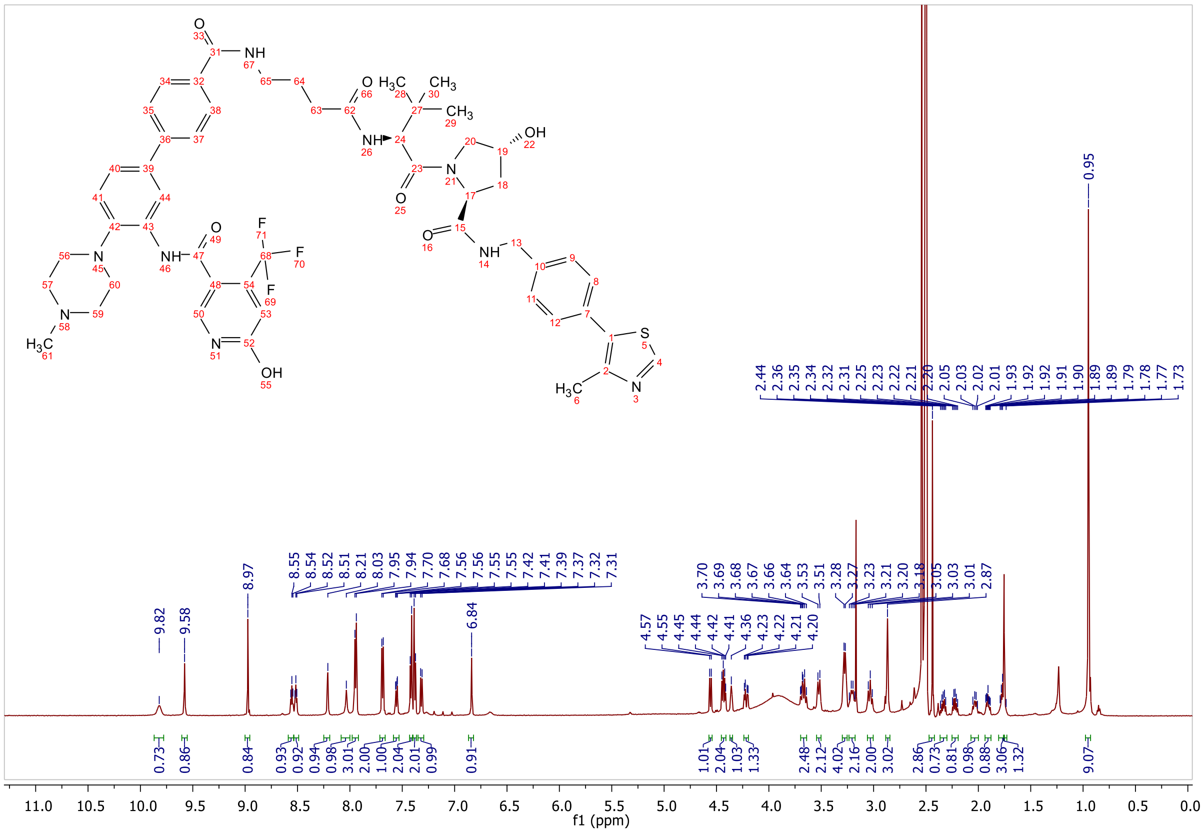
f1 (ppm)

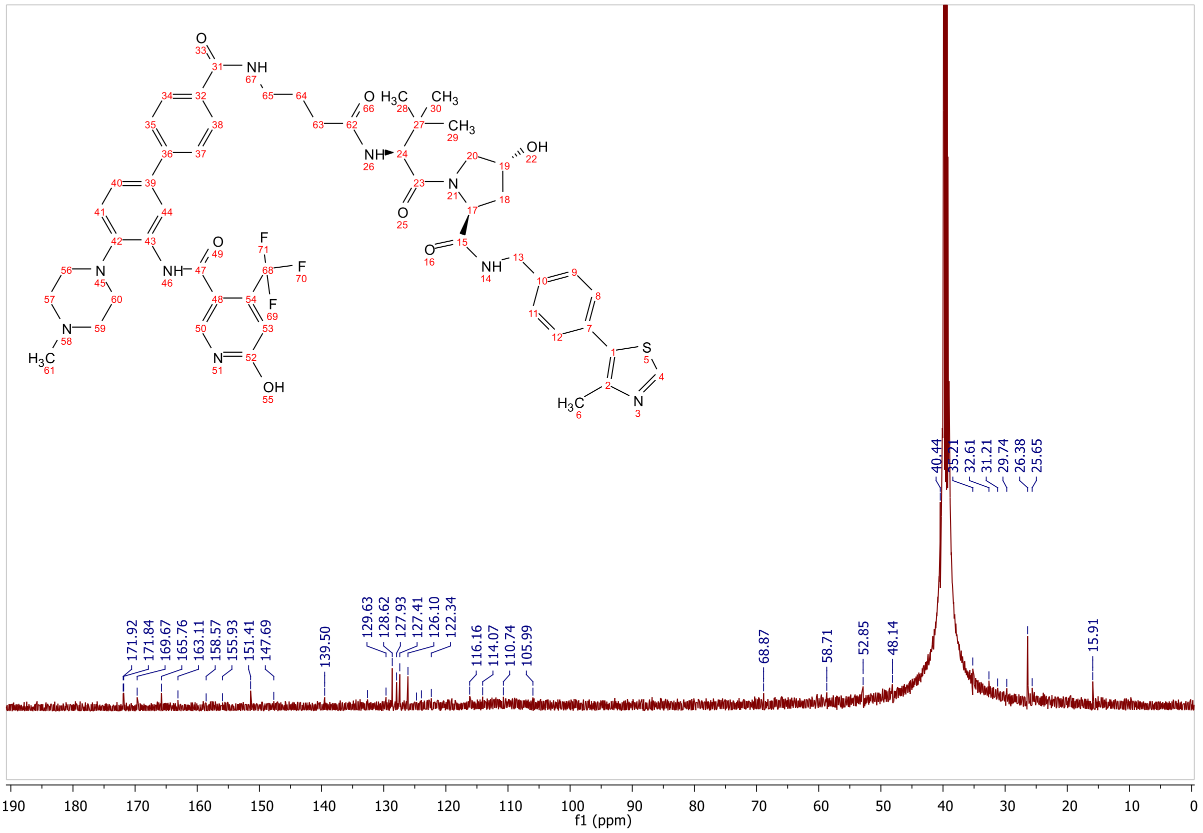
171.92
171.84
169.67
165.76
163.11
158.57
155.93
151.41
147.69
139.50
129.63
128.62
127.93
127.41
126.10
122.34
116.16
114.07
110.74
105.99
68.87
58.71
52.85
48.14
40.44
35.21
32.61
31.21
29.74
26.38
25.65
15.91
f1 (ppm)

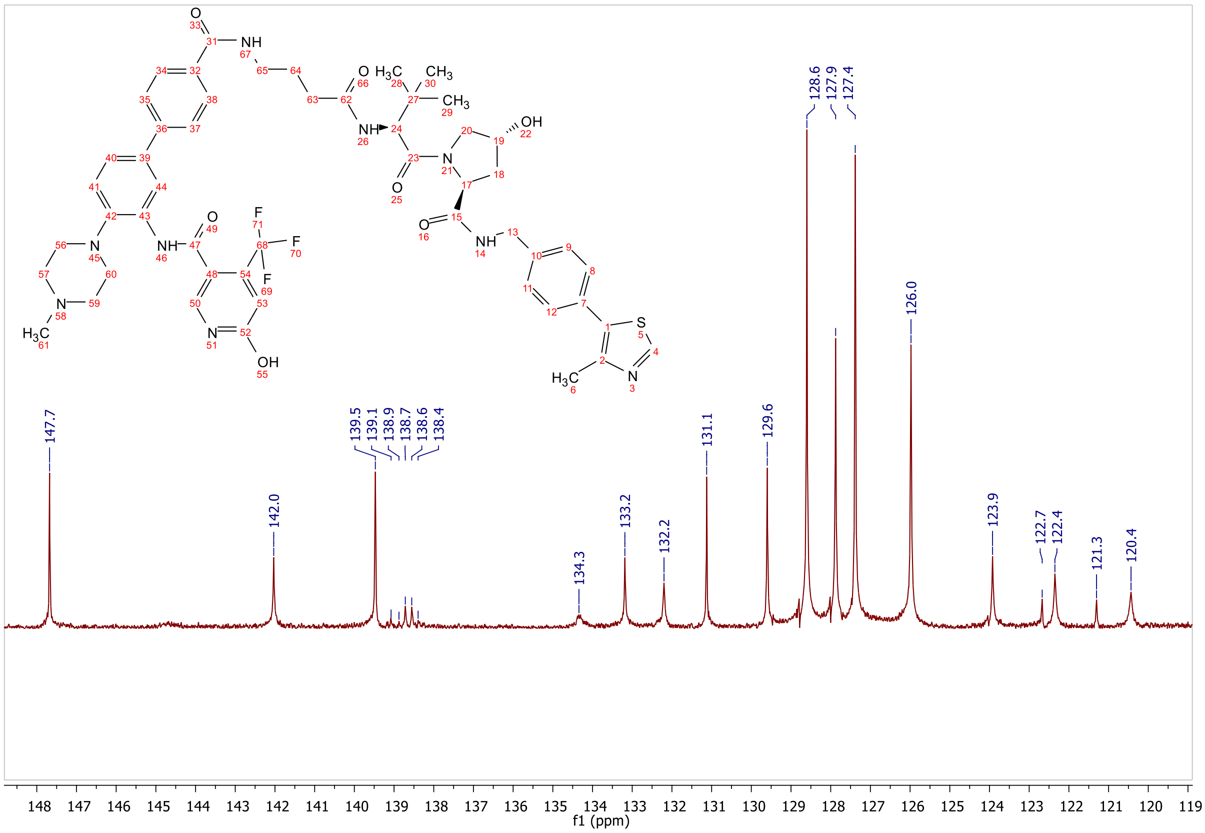
147.7
142.0
139.5
139.1
138.9
138.7
138.6
138.4
134.3
133.2
132.2
131.1
129.6
128.6
127.9
127.4
126.0
123.9
122.7
122.4
121.3
120.4
f1 (ppm)

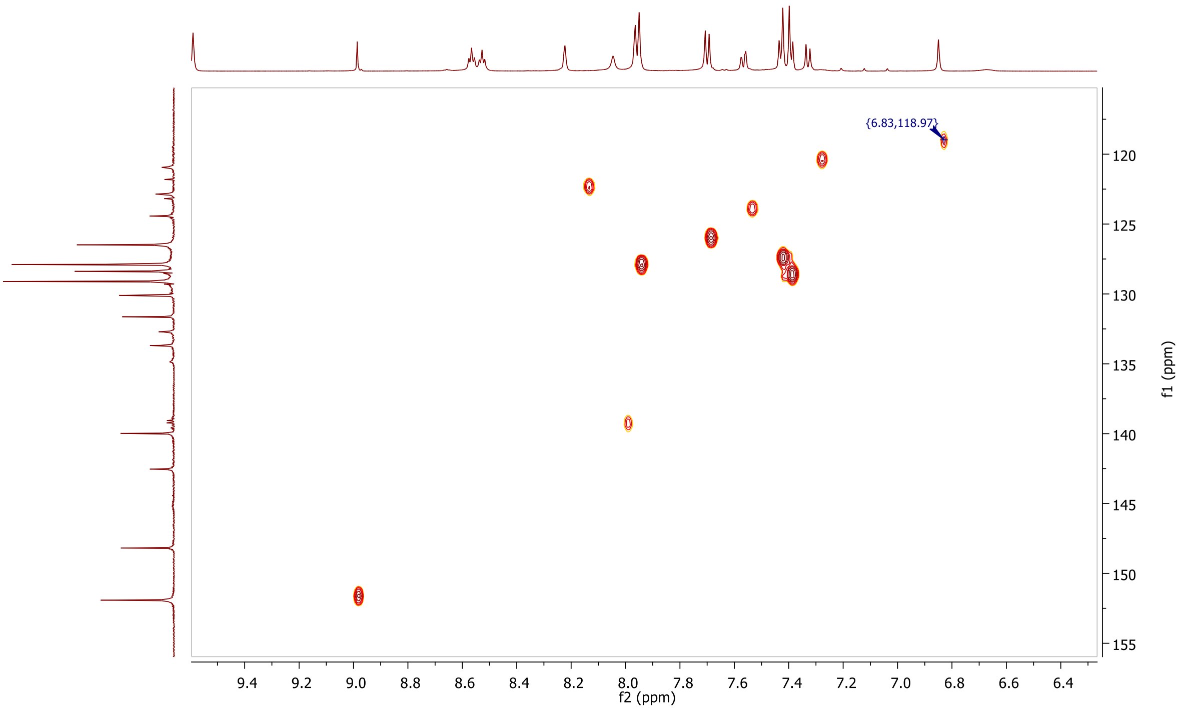
{6.83,118.97}
f2 (ppm)
f1 (ppm)

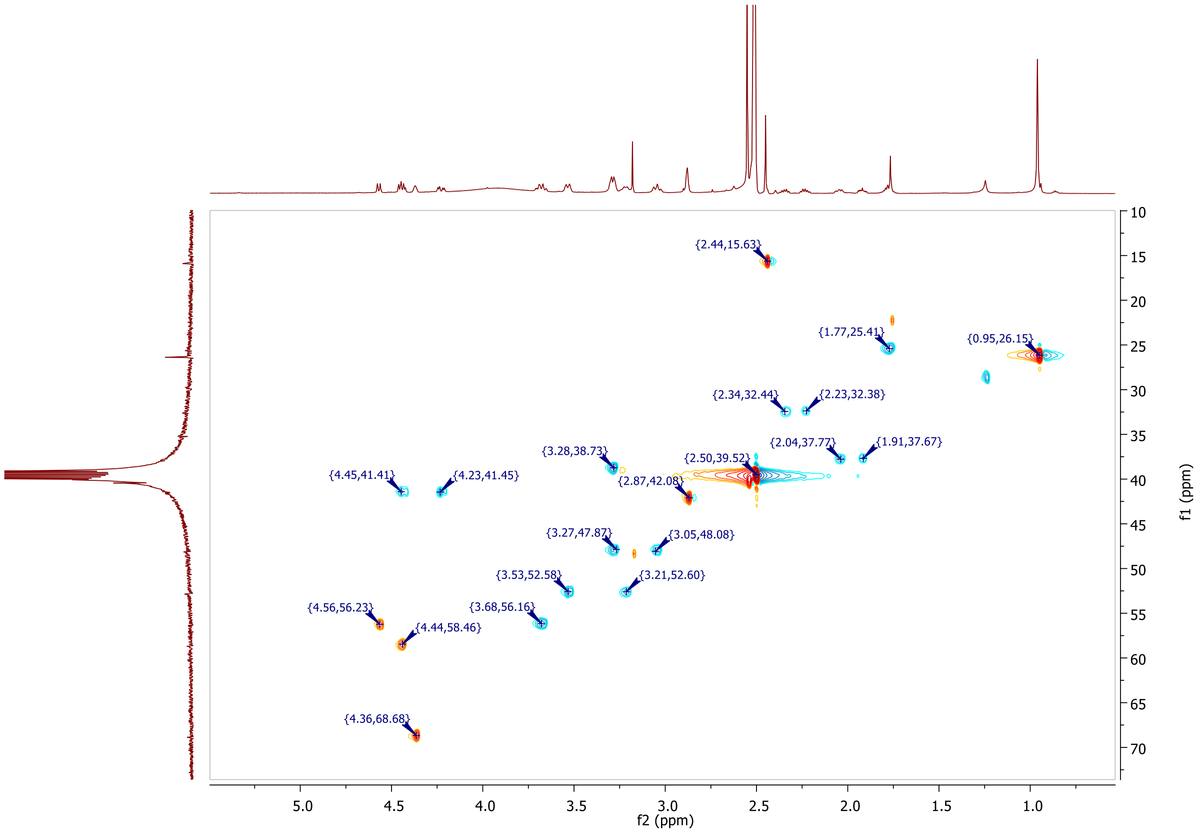
{2.44,15.63}
{1.77,25.41}
{0.95,26.15}
{2.34,32.44}
{2.23,32.38}
{2.04,37.77}
{1.91,37.67}
{3.28,38.73}
{2.50,39.52}
{4.45,41.41}
{4.23,41.45}
{2.87,42.08}
{3.27,47.87}
{3.05,48.08}
{3.53,52.58}
{3.21,52.60}
{4.56,56.23}
{3.68,56.16}
{4.44,58.46}
{4.36,68.68}
f2 (ppm)
f1 (ppm)

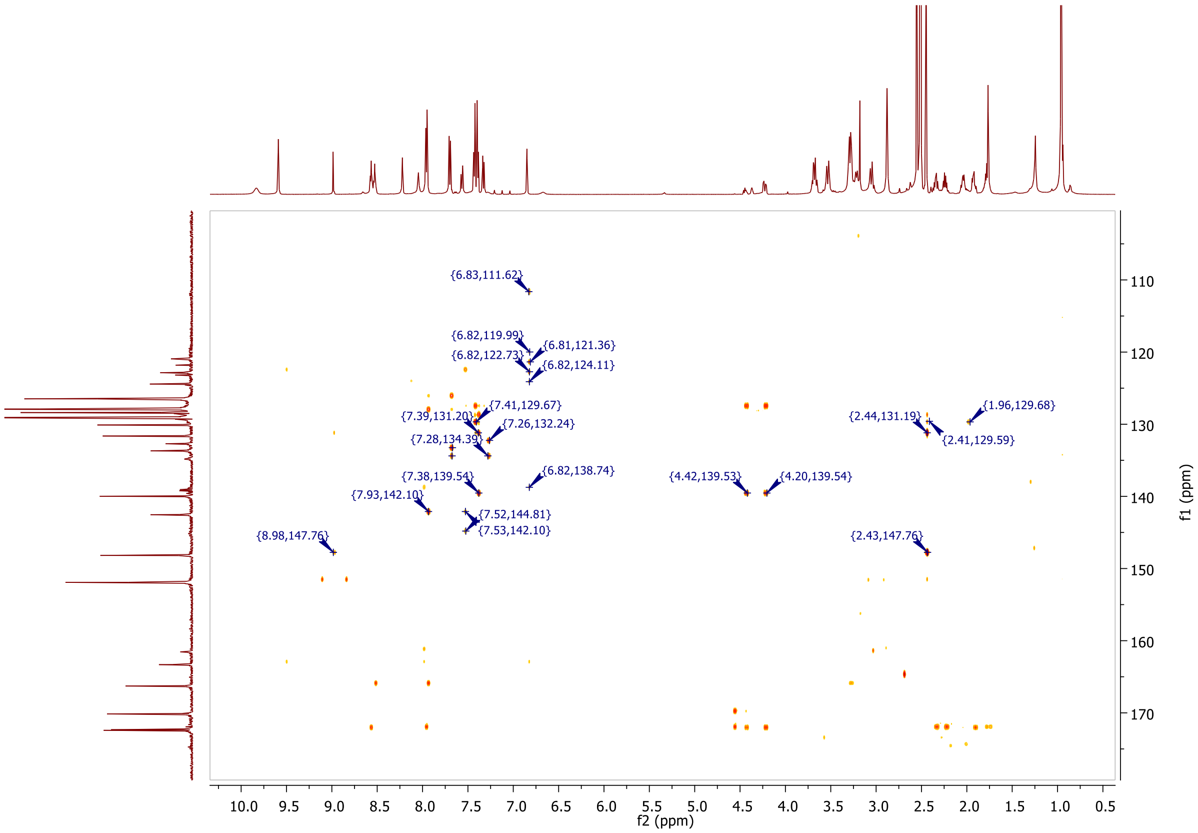
{6.83,111.62}
{6.82,119.99}
{6.81,121.36}
{6.82,122.73}
{6.82,124.11}
{7.41,129.67}
{7.39,131.20}
{7.26,132.24}
{7.28,134.39}
{6.82,138.74}
{7.38,139.54}
{4.42,139.53}
{4.20,139.54}
{7.93,142.10}
{7.52,144.81}
{7.53,142.10}
{8.98,147.76}
{1.96,129.68}
{2.44,131.19}
{2.41,129.59}
{2.43,147.76}
f2 (ppm)
f1 (ppm)

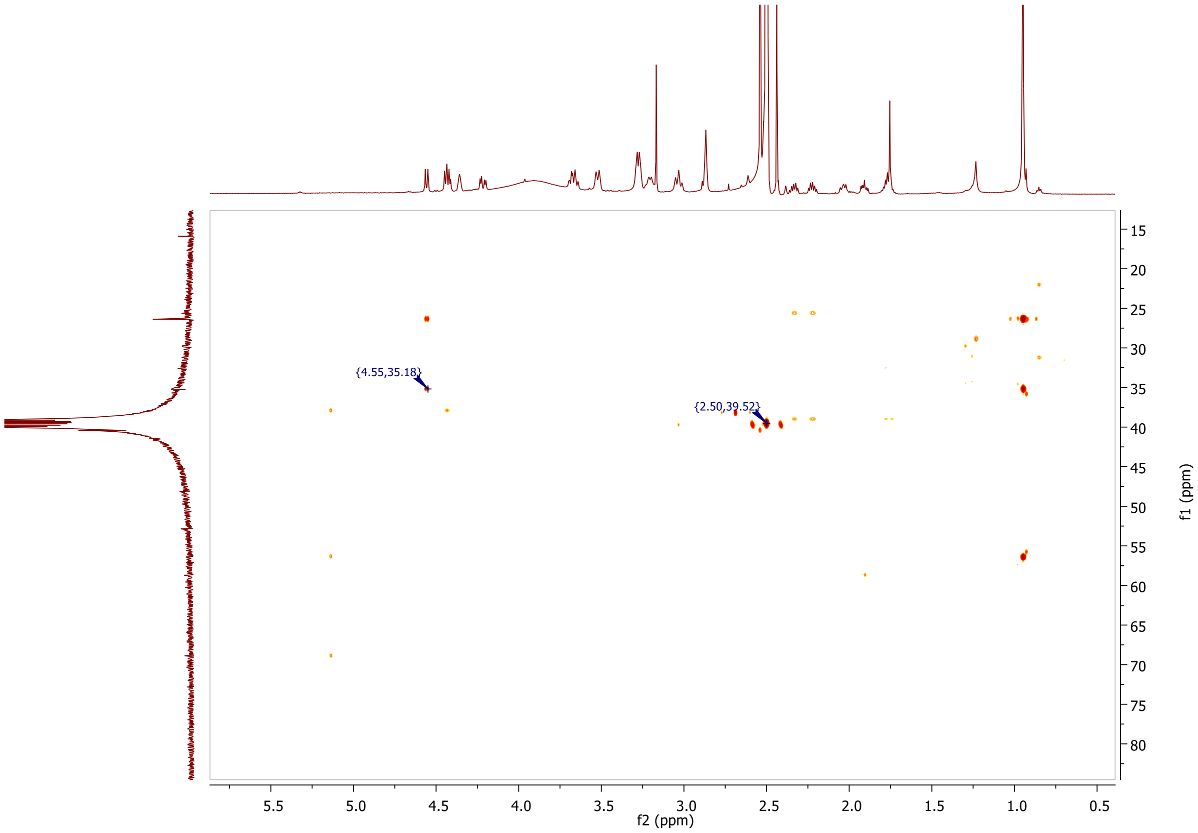

{4.55,35.18}
{2.50,39.52}
5.5
5.0
4.5
4.0
3.5
3.0
2.5
2.0
1.5
1.0
0.5
f2 (ppm)
15
20
25
30
35
40
45
50
55
60
65
70
75
80
f1 (ppm)

**ESI, HRMS, HPLC, $^{1}$H-NMR, $^{13}$C-NMR, $^{13}$C-NMR (downfield region), $^{1}$H-$^{13}$C-HSQC, $^{1}$H-$^{13}$C-HMBC (downfield) and $^{1}$H-$^{13}$C-HMBC (downfield) of 6-hydroxy-*N*-(4'-((5-(((*S*)-1-((2*S*,4*R*)-4-hydroxy-2-((4-(4-methylthiazol-5-yl)benzyl)carbamoyl)pyrrolidin-1-yl)-3,3-dimethyl-1-oxobutan-2-yl)amino)-5-oxopentyl)carbamoyl)-4-(4-methylpiperazin-1-yl)-[1,1'-biphenyl]-3-yl)-4-(trifluoromethyl)nicotinamide 8g**

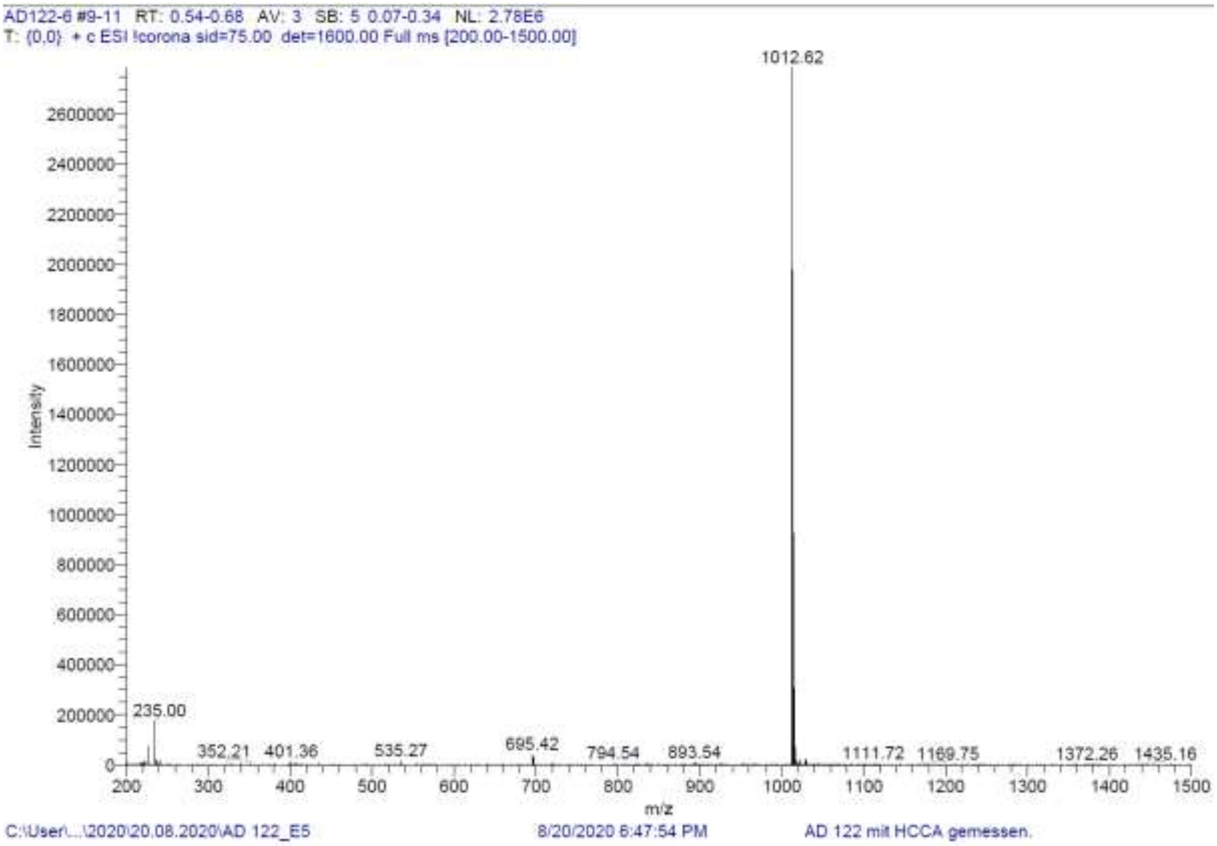

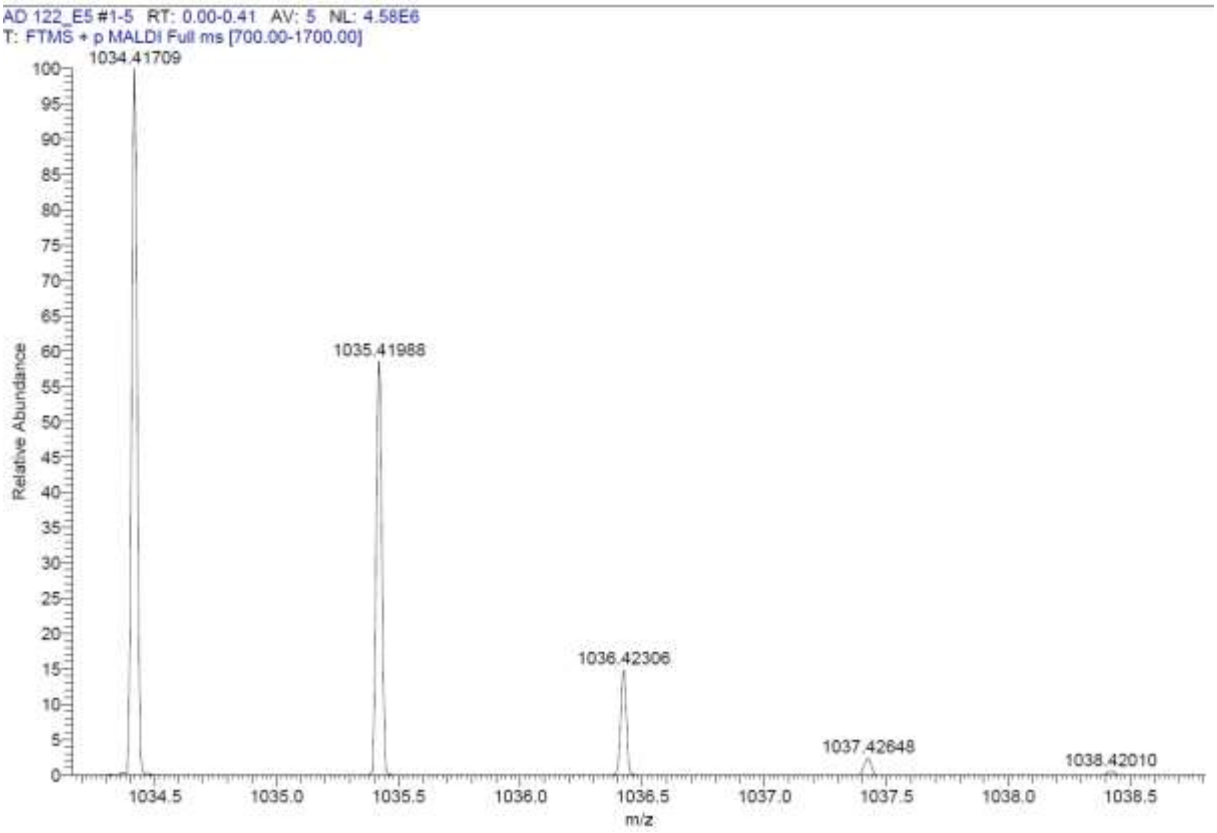

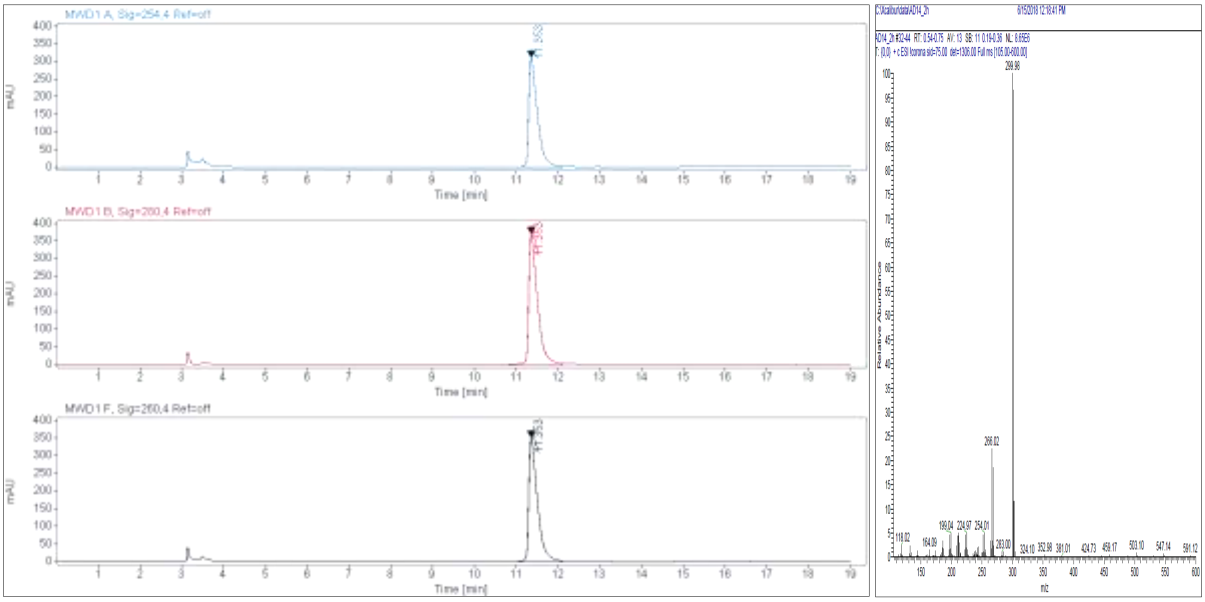

**Signal:** MWD1 A, Sig=254,4 Ref=off

| RT [min] | Type | Width [min] | Area | Height | Area% | Name |
|---|---|---|---|---|---|---|
| 11.353 | VV | 0.2008 | 4513.2095 | 311.9199 | 100.0000 | |
| | | Sum | 4513.2095 | | | |

**Signal:** MWD1 B, Sig=280,4 Ref=off

| RT [min] | Type | Width [min] | Area | Height | Area% | Name |
|---|---|---|---|---|---|---|
| 11.353 | VV | 0.2029 | 5449.8301 | 372.0602 | 100.0000 | |
| | | Sum | 5449.8301 | | | |

**Signal:** MWD1 F, Sig=260,4 Ref=off

| RT [min] | Type | Width [min] | Area | Height | Area% | Name |
|---|---|---|---|---|---|---|
| 11.353 | VV | 0.2021 | 5101.9272 | 352.0205 | 100.0000 | |

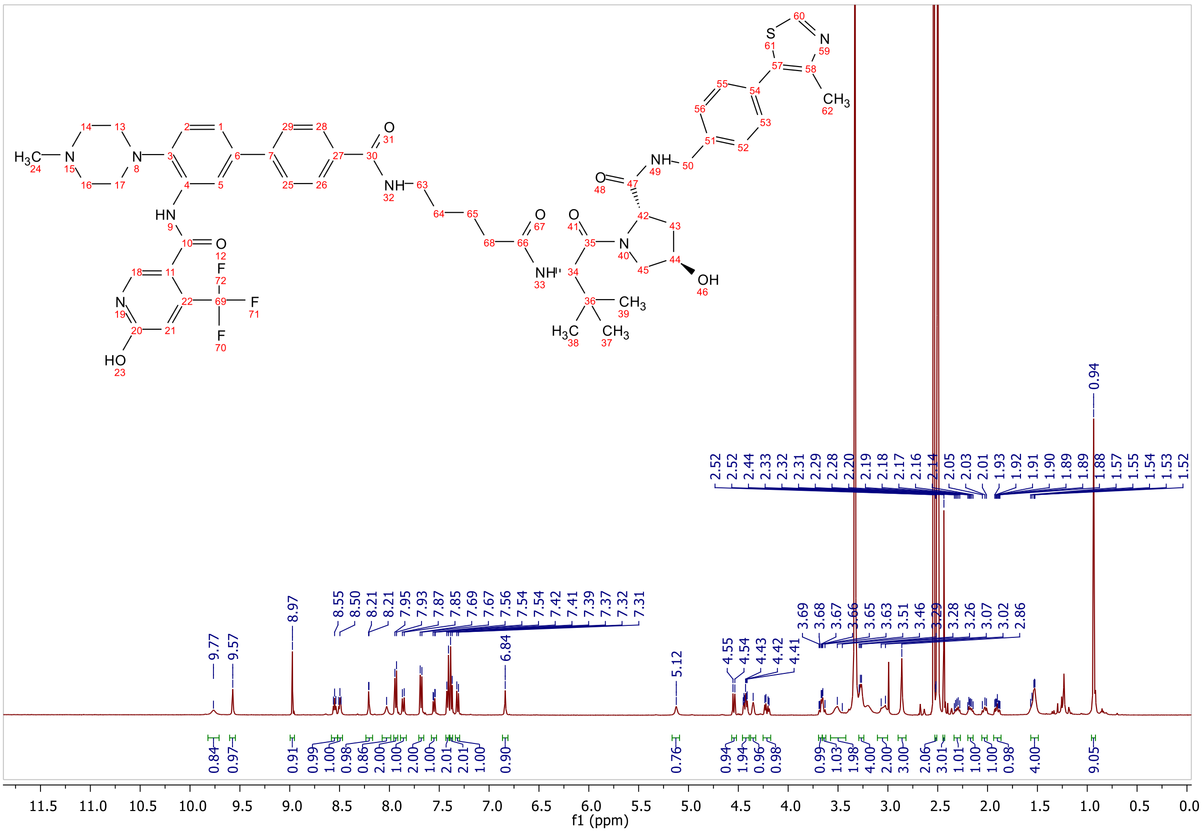
H3C N N HN O N HO F F F NH O O NH O N OH O NH S N CH3 H3C CH3 CH3
9.77 9.57 8.97 8.55 8.50 8.21 8.21 7.95 7.93 7.87 7.85 7.69 7.67 7.56 7.54 7.54 7.42 7.41 7.39 7.37 7.32 7.31 6.84 5.12 4.55 4.54 4.43 4.42 4.41 3.69 3.68 3.67 3.66 3.65 3.63 3.51 3.46 3.29 3.28 3.26 3.07 3.02 2.86
2.52 2.52 2.44 2.33 2.32 2.31 2.29 2.28 2.20 2.19 2.18 2.17 2.16 2.14 2.05 2.03 2.01 1.93 1.92 1.91 1.90 1.89 1.89 1.88 1.57 1.55 1.54 1.53 1.52 0.94
0.84 0.97 0.91 0.99 1.00 0.98 0.86 2.00 1.00 2.00 1.00 2.01 2.01 1.00 0.90 0.76 0.94 1.94 0.96 0.98 0.99 1.03 1.98 4.00 2.00 3.00 2.06 3.01 1.01 1.00 1.00 0.98 4.00 9.05
f1 (ppm)

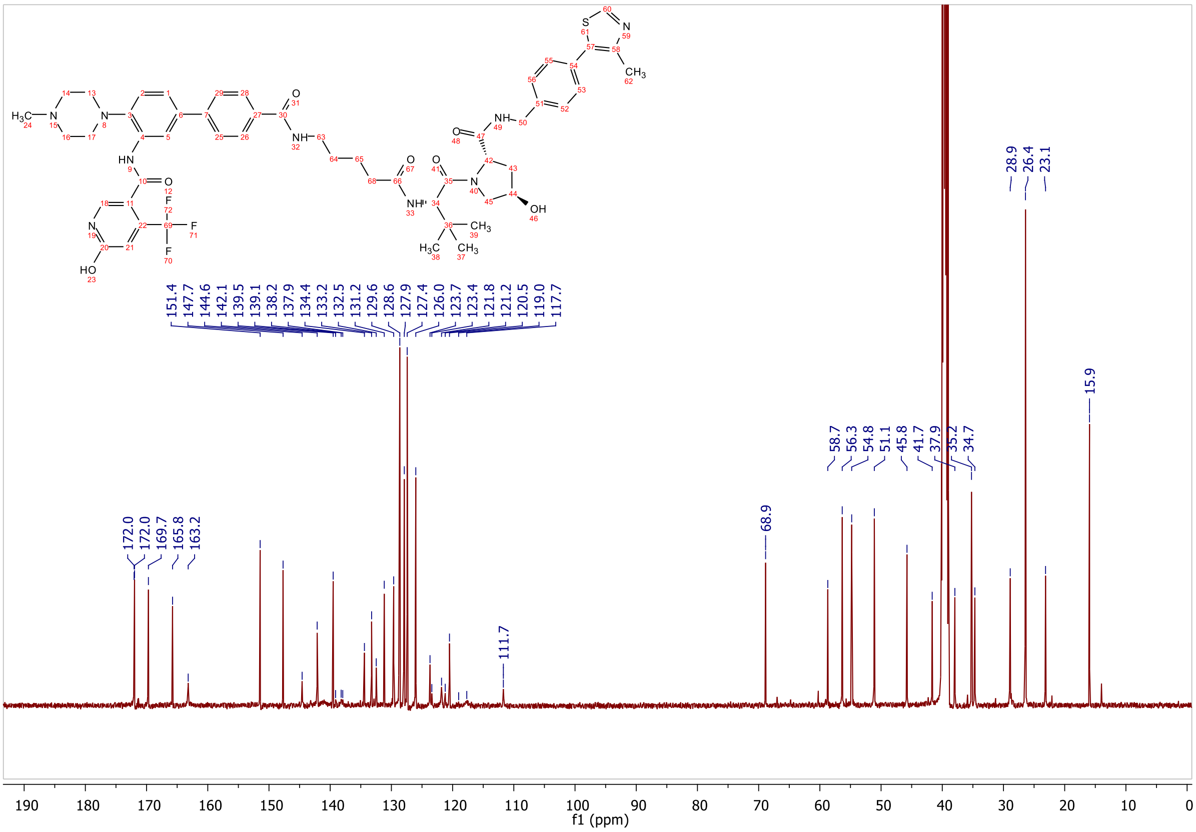
172.0 172.0 169.7 165.8 163.2
151.4 147.7 144.6 142.1 139.5 139.1 138.2 137.9 134.4 133.2 132.5 131.2 129.6 128.6 127.9 127.4 126.0 123.7 123.4 121.8 121.2 120.5 119.0 117.7
111.7 68.9 58.7 56.3 54.8 51.1 45.8 41.7 37.9 35.2 34.7 28.9 26.4 23.1 15.9
f1 (ppm)

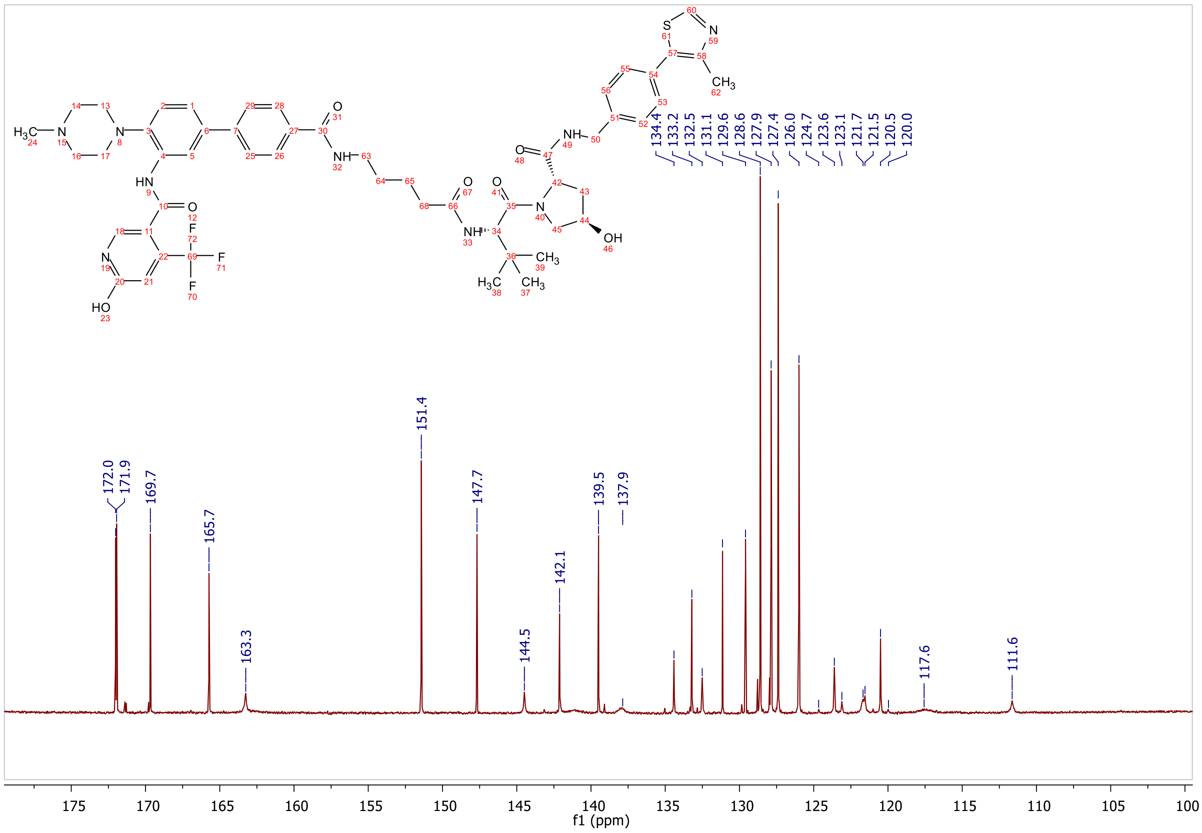
172.0
171.9
169.7
165.7
163.3
151.4
147.7
144.5
142.1
139.5
137.9
134.4
133.2
132.5
131.1
129.6
128.6
127.9
127.4
126.0
124.7
123.6
123.1
121.7
121.5
120.5
120.0
117.6
111.6
f1 (ppm)

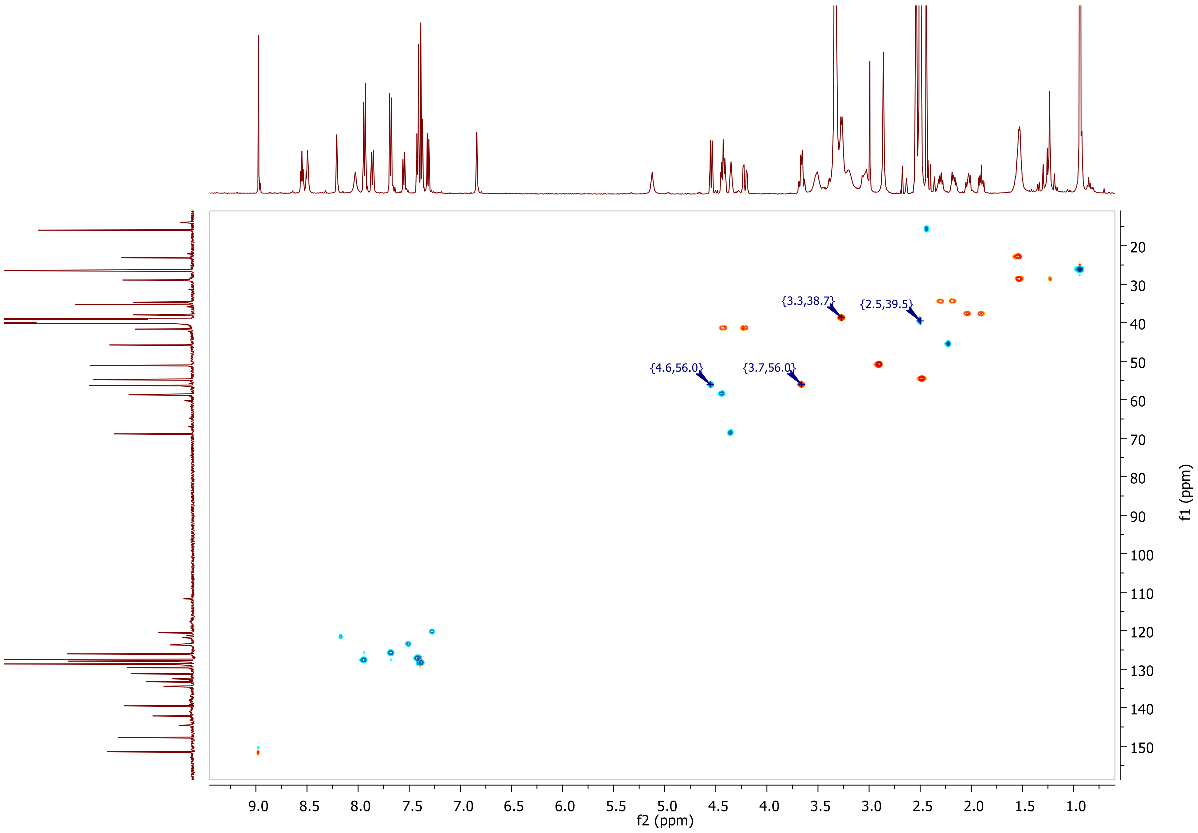
{3.3,38.7}
{2.5,39.5}
{4.6,56.0}
{3.7,56.0}
f2 (ppm)
f1 (ppm)

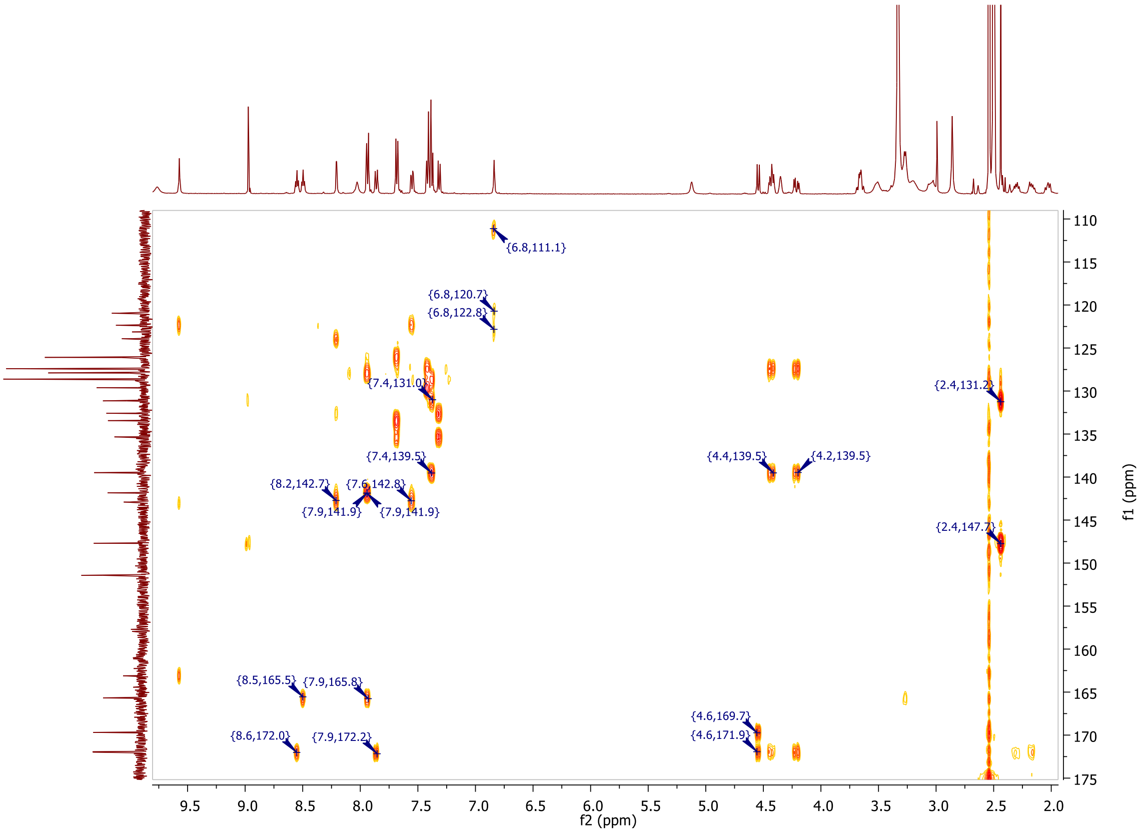
{6.8,111.1}
{6.8,120.7}
{6.8,122.8}
{7.4,131.0}
{2.4,131.2}
{7.4,139.5}
{4.4,139.5}
{4.2,139.5}
{8.2,142.7}
{7.6,142.8}
{7.9,141.9}
{7.9,141.9}
{2.4,147.7}
{8.5,165.5}
{7.9,165.8}
{4.6,169.7}
{4.6,171.9}
{8.6,172.0}
{7.9,172.2}
f2 (ppm)
f1 (ppm)

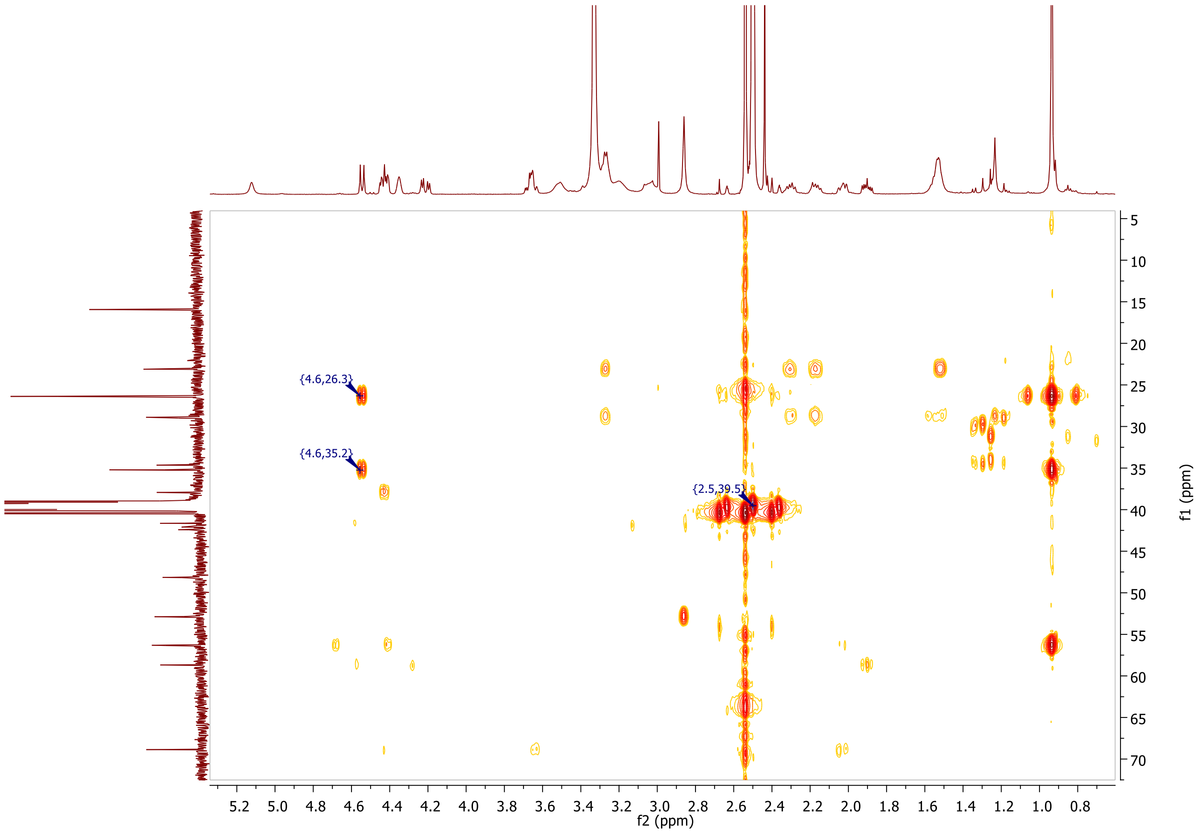
{4.6,26.3}
{4.6,35.2}
{2.5,39.5}
f2 (ppm)
f1 (ppm)

**MALDI, HRMS, HPLC,[1]H-NMR and [13]C-NMR of 6-hydroxy-*N*-(4'-((6-(((*S*)-1-((2*S*,4*R*)-4-hydroxy-2-((4-(4-methylthiazol-5-yl)benzyl)carbamoyl)pyrrolidin-1-yl)-3,3-dimethyl-1-oxobutan-2-yl)amino)-6-oxohexyl)carbamoyl)-4-(4-methylpiperazin-1-yl)-[1,1'-biphenyl]-3-yl)-4-(trifluoromethyl)nicotinamide 8h**

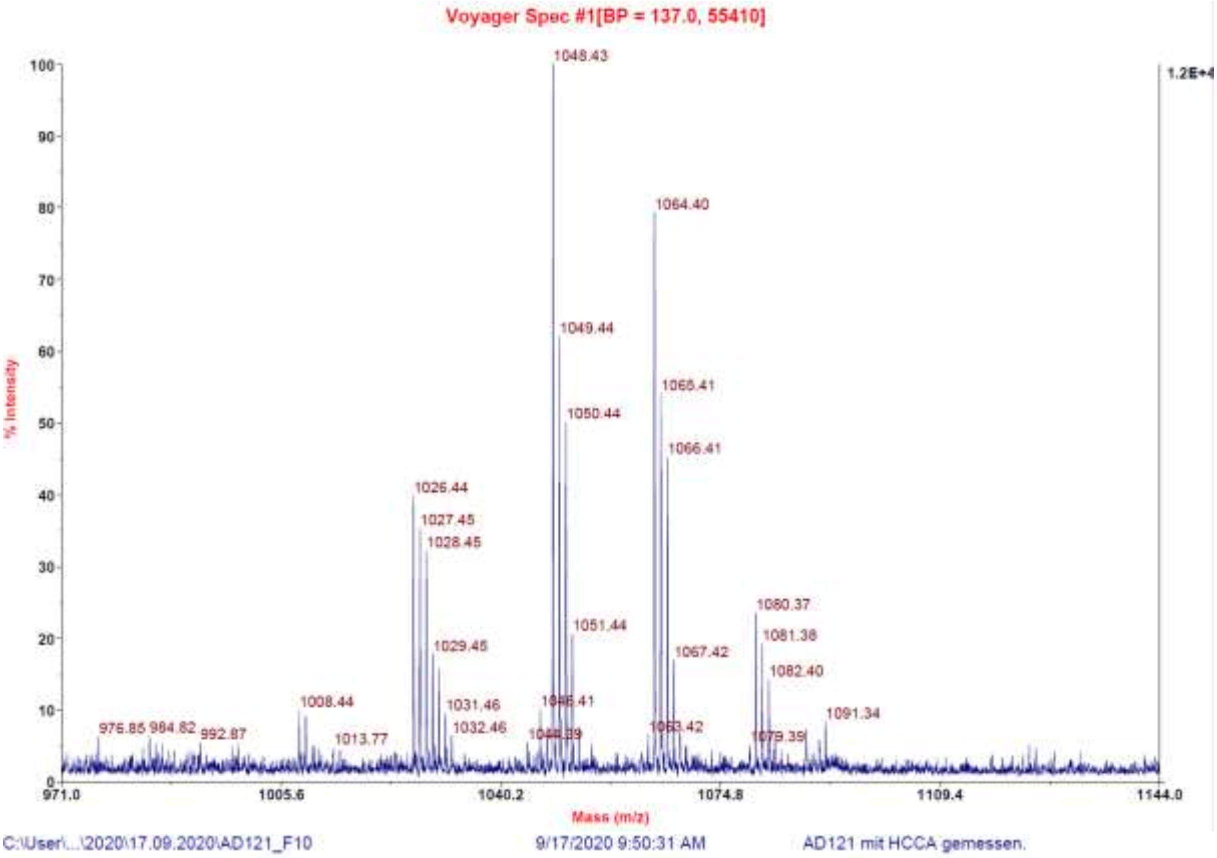

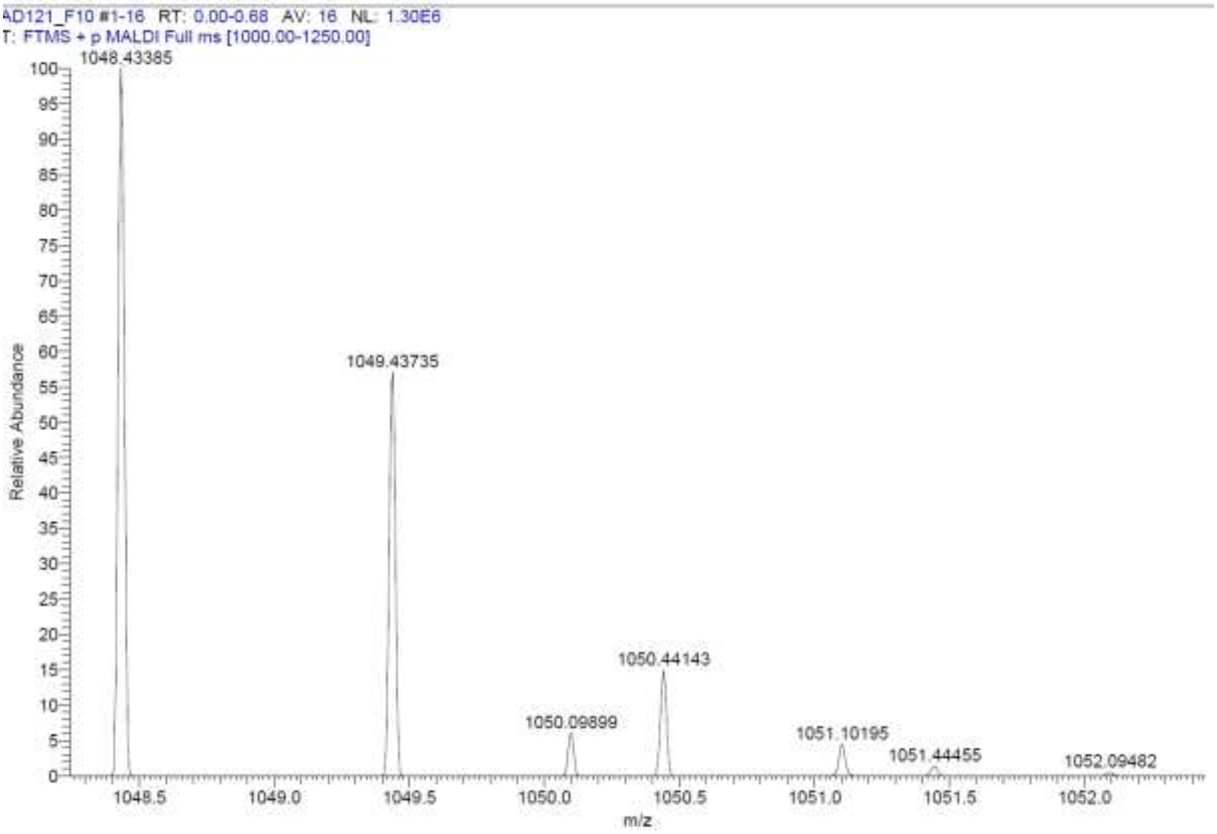

**Signal:** MWD1 A, Sig=254,4 Ref=off

| RT [min] | Type | Width [min] | Area | Height | Area% | Name |
|---|---|---|---|---|---|---|
| 11.385 | VV | 0.2094 | 4595.4873 | 303.9242 | 100.0000 | |
| | | Sum | 4595.4873 | | | |

**Signal:** MWD1 B, Sig=280,4 Ref=off

| RT [min] | Type | Width [min] | Area | Height | Area% | Name |
|---|---|---|---|---|---|---|
| 11.385 | VV | 0.2101 | 5472.0347 | 360.5278 | 100.0000 | |
| | | Sum | 5472.0347 | | | |

**Signal:** MWD1 F, Sig=260,4 Ref=off

| RT [min] | Type | Width [min] | Area | Height | Area% | Name |
|---|---|---|---|---|---|---|
| 11.385 | VV | 0.2096 | 5170.4536 | 341.5610 | 100.0000 | |

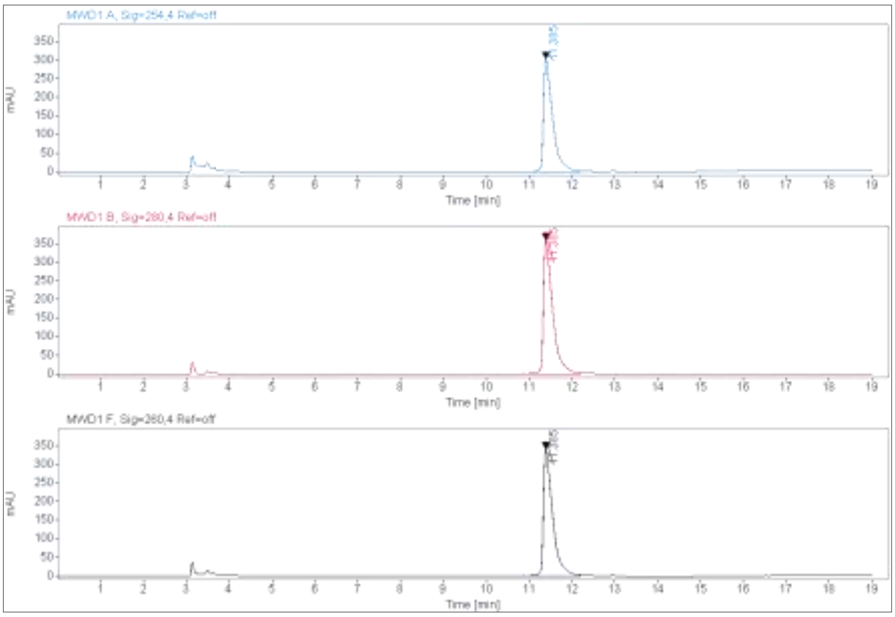

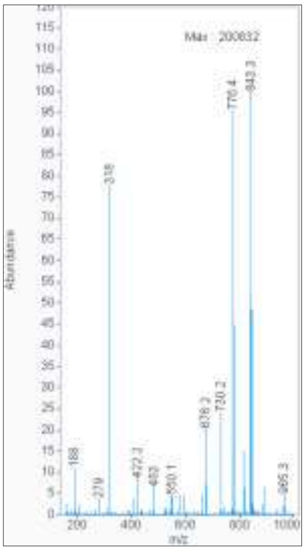

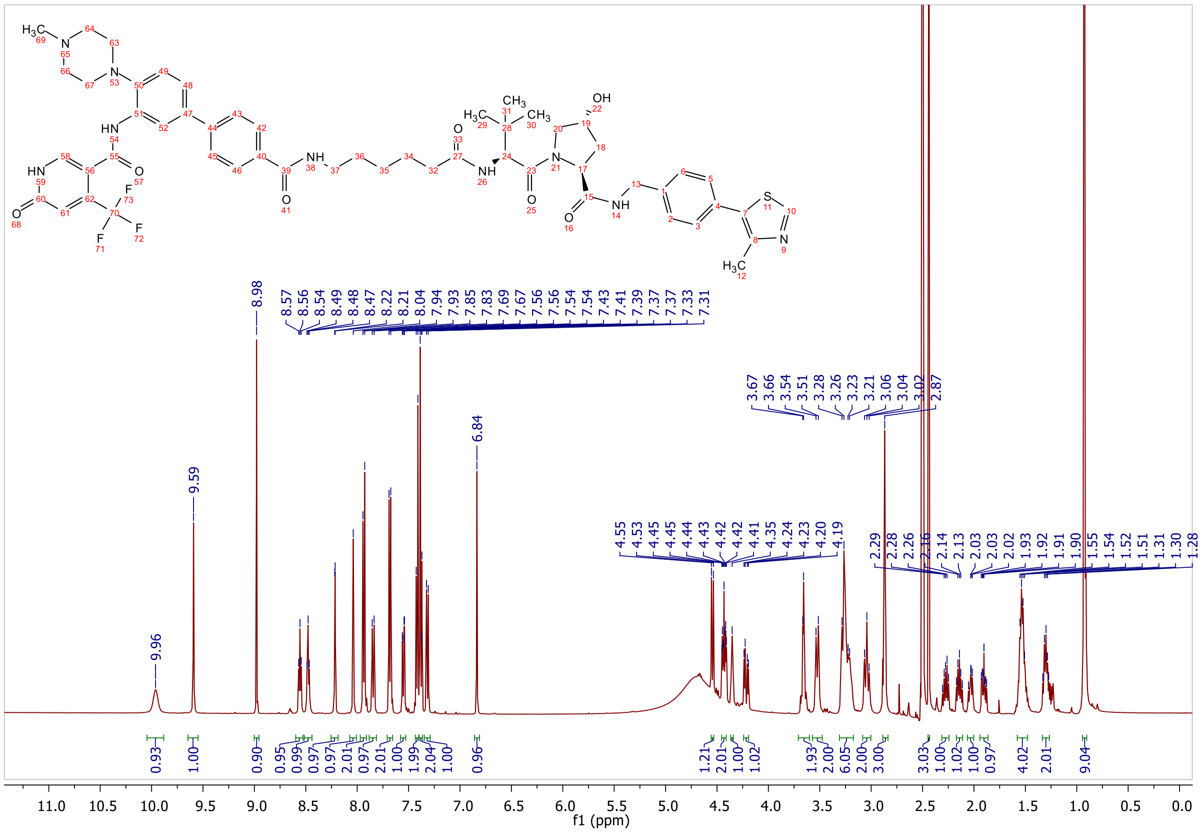
9.96
9.59
8.98
6.84
0.93
1.00
0.90
9.04
f1 (ppm)

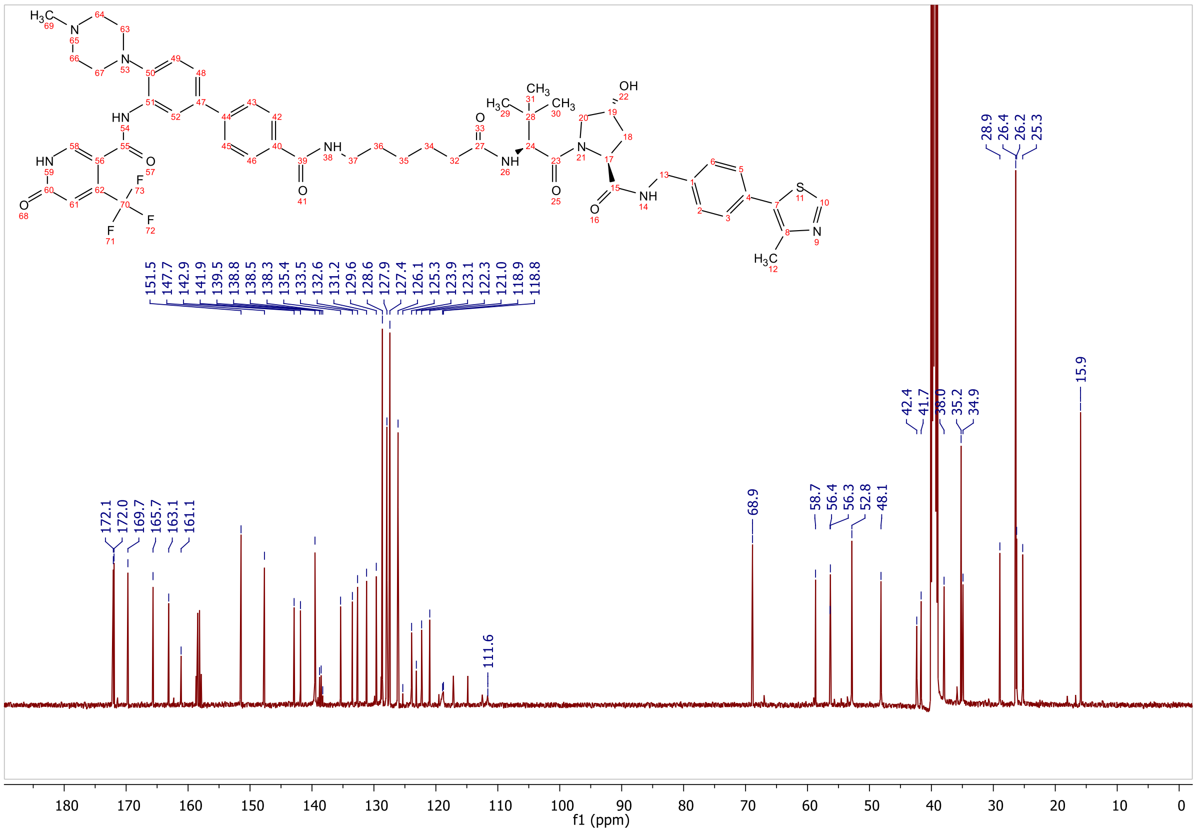
172.1
172.0
169.7
165.7
163.1
161.1
151.5
147.7
142.9
141.9
139.5
138.8
138.5
138.3
135.4
133.5
132.6
131.2
129.6
128.6
127.9
127.4
126.1
125.3
123.9
123.1
122.3
121.0
118.9
118.8
111.6
68.9
58.7
56.4
56.3
52.8
48.1
42.4
41.7
38.0
35.2
34.9
28.9
26.4
26.2
25.3
15.9
f1 (ppm)

**HPLC, MALDI, HRMS, $^{1}H$-NMR, $^{13}C$-NMR, $^{1}H$-$^{13}C$-HSQC and $^{1}H$-$^{13}C$-HMBC of 6-hydroxy-*N*-(4'-((7-(((*S*)-1-((2*S*,4*R*)-4-hydroxy-2-((4-(4-methylthiazol-5-yl)benzyl)carbamoyl)pyrrolidin-1-yl)-3,3-dimethyl-1-oxobutan-2-yl)amino)-7-oxoheptyl)carbamoyl)-4-(4-methylpiperazin-1-yl)-[1,1'-biphenyl]-3-yl)-4-(trifluoromethyl)nicotinamide 8i**

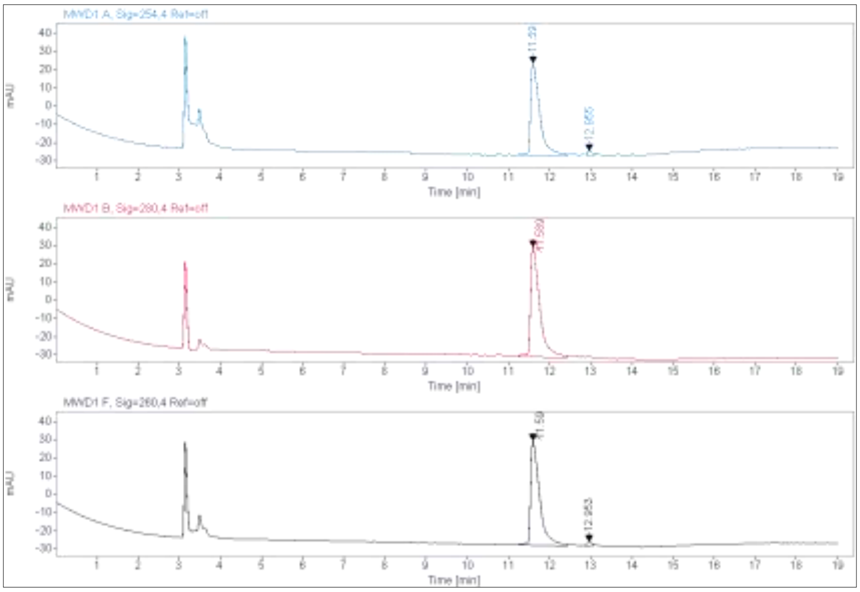

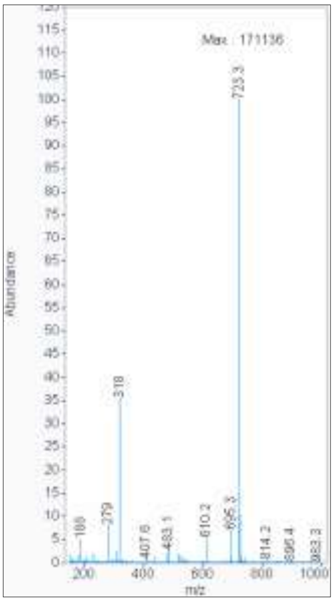

**Signal:** MWD1 A, Sig=254,4 Ref=off

| RT [min] | Type | Width [min] | Area | Height | Area% | Name |
|---|---|---|---|---|---|---|
| 11.590 | VV | 0.2077 | 772.4310 | 50.7114 | 97.5053 | |
| 12.955 | MM | 0.1361 | 19.7630 | 2.4203 | 2.4947 | |
| | | Sum | 792.1941 | | | |

**Signal:** MWD1 B, Sig=280,4 Ref=off

| RT [min] | Type | Width [min] | Area | Height | Area% | Name |
|---|---|---|---|---|---|---|
| 11.589 | VV | 0.2115 | 947.8234 | 60.6039 | 100.0000 | |
| | | Sum | 947.8234 | | | |

**Signal:** MWD1 F, Sig=260,4 Ref=off

| RT [min] | Type | Width [min] | Area | Height | Area% | Name |
|---|---|---|---|---|---|---|
| 11.590 | VV | 0.2078 | 879.9773 | 57.4239 | 97.6449 | |
| 12.953 | MM | 0.1394 | 21.2239 | 2.5383 | 2.3551 | |

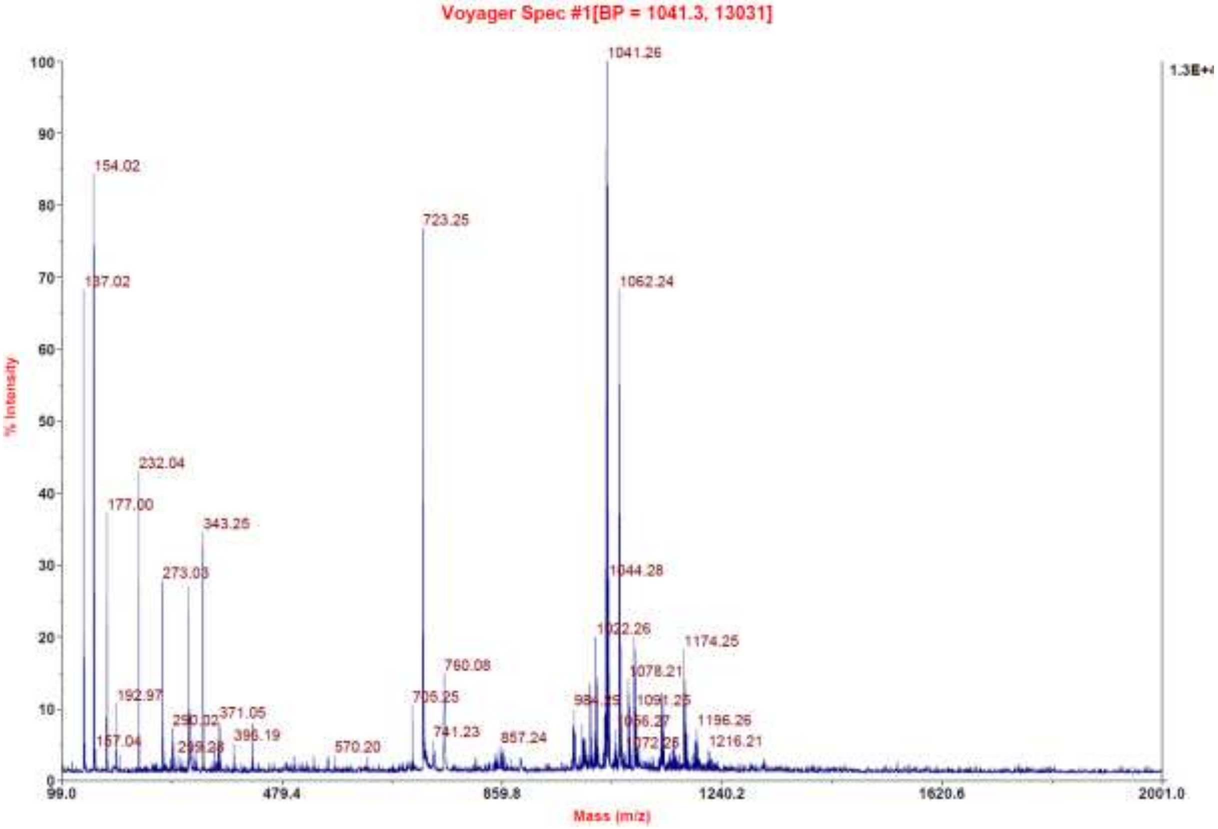

C:\User\...\2020\20.08.2020\AD 110_E3 8/20/2020 6:45:29 PM AD 110 mit HCCA gemessen.

AD 110_E3 #1-4 RT: 0.01-0.32 AV: 4 NL: 2.67E6
T: FTMS + p MALDI Full ms [700.00-1700.00]

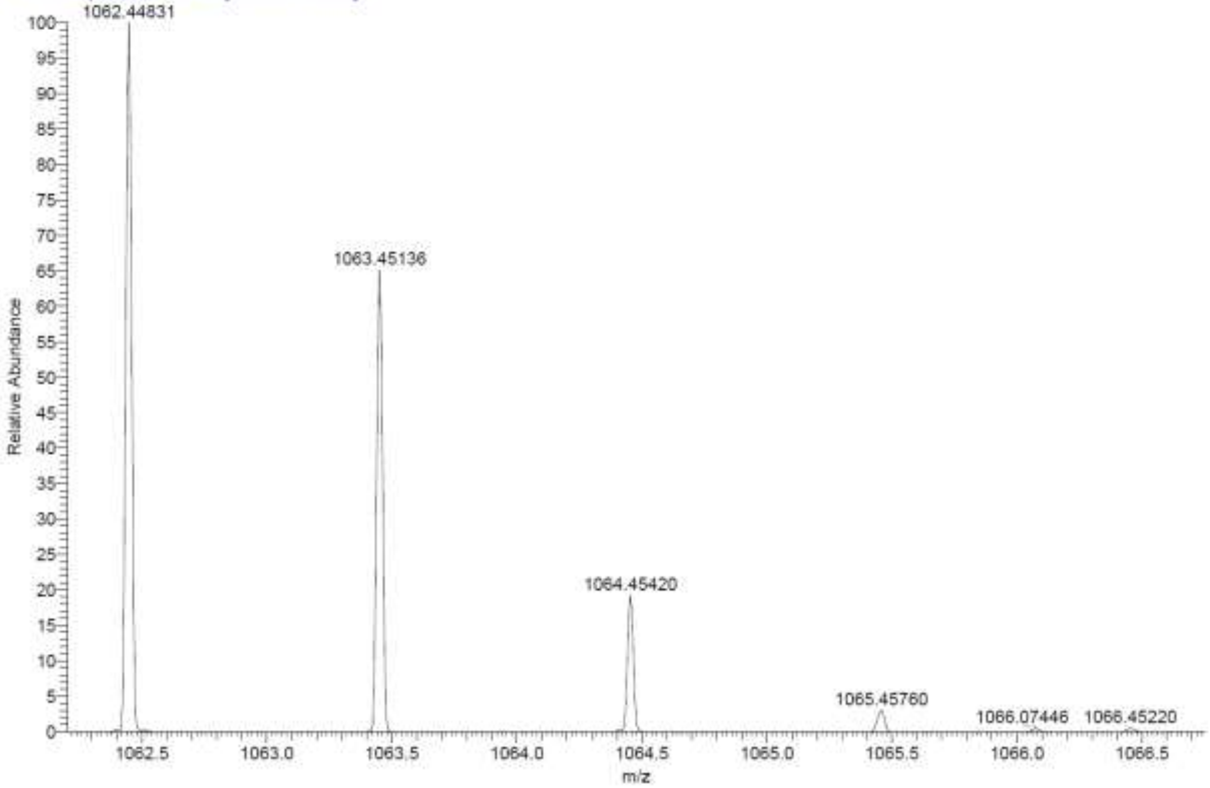

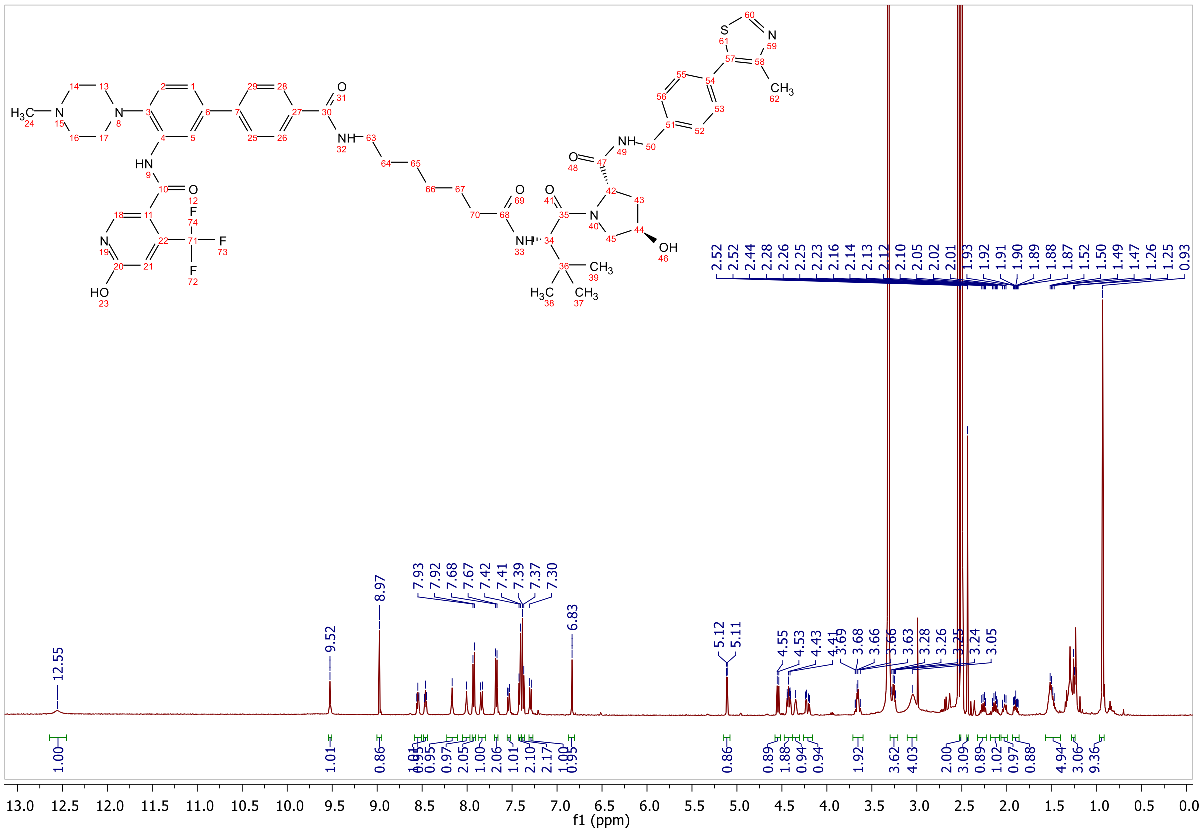

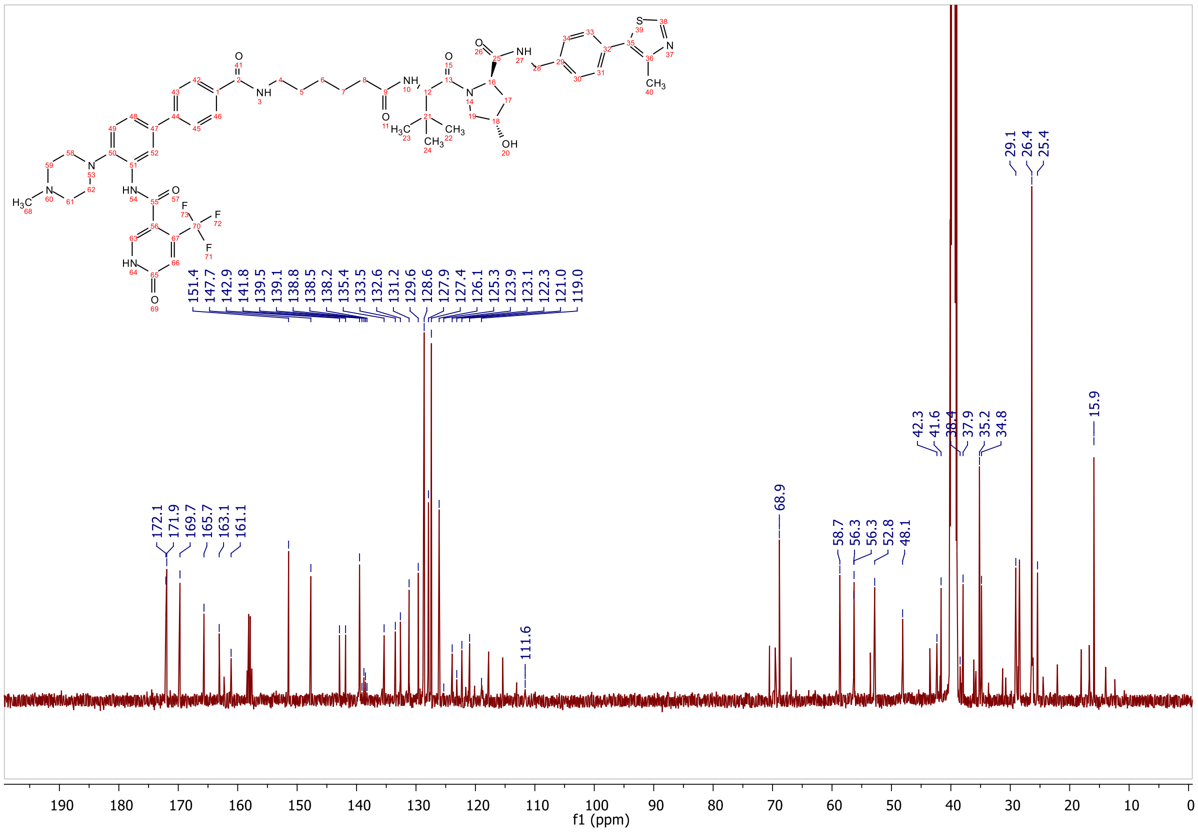

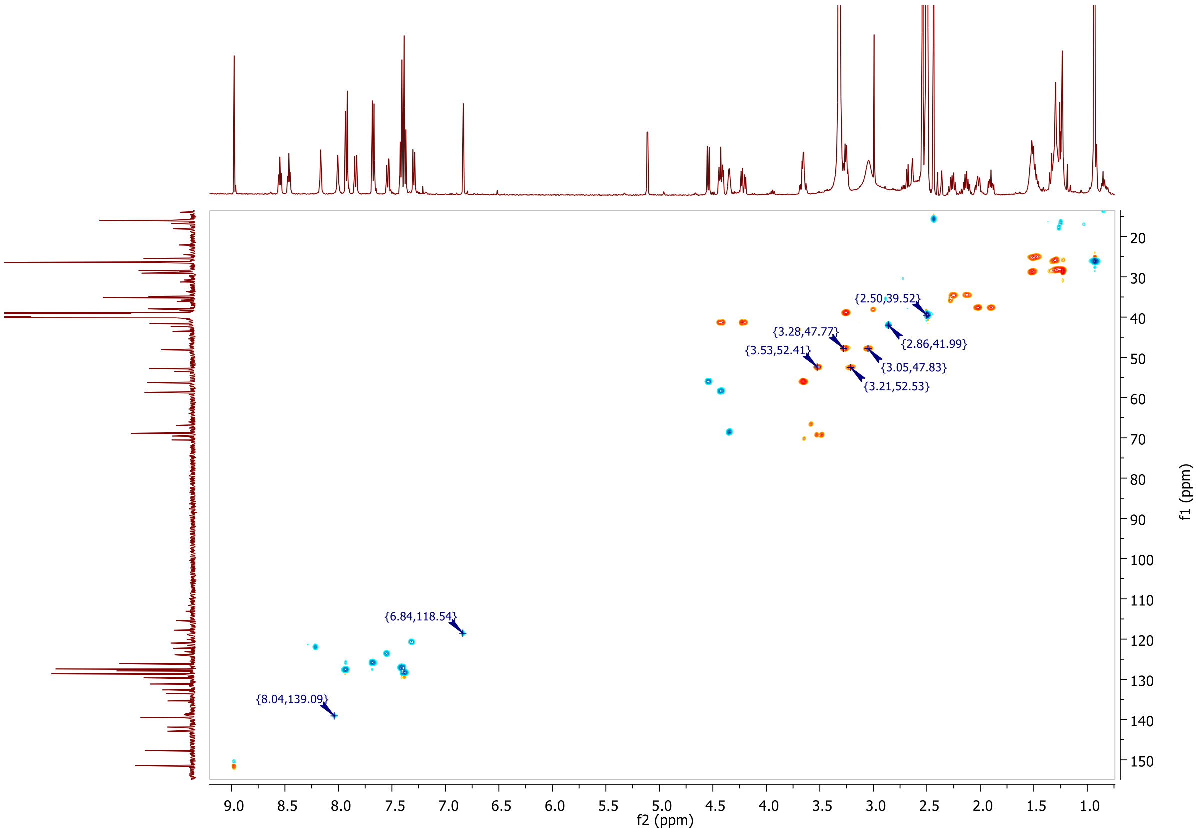
{2.50,39.52}
{3.28,47.77}
{3.53,52.41}
{2.86,41.99}
{3.05,47.83}
{3.21,52.53}
{6.84,118.54}
{8.04,139.09}
f2 (ppm)
f1 (ppm)

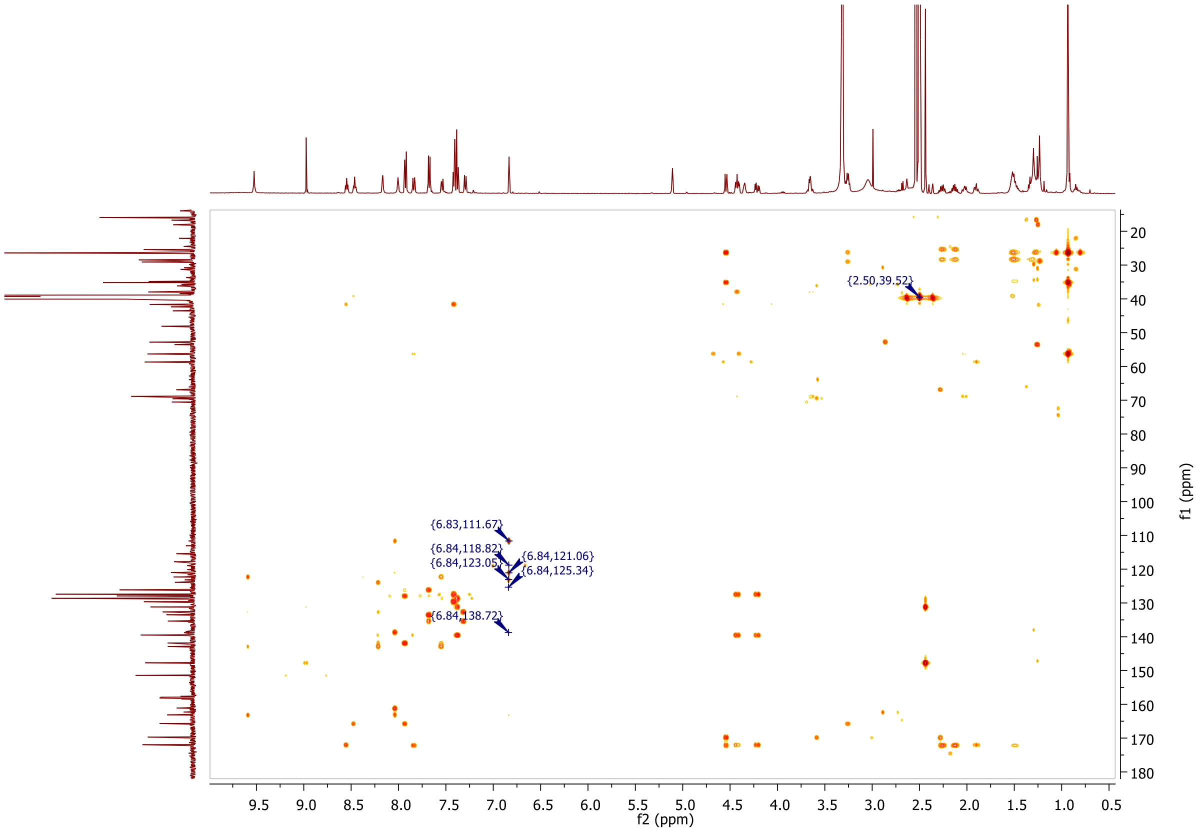
{2.50,39.52}
{6.83,111.67}
{6.84,118.82}
{6.84,123.05}
{6.84,121.06}
{6.84,125.34}
{6.84,138.72}
f2 (ppm)
f1 (ppm)

**MALDI, HRMS, HPLC, $^{1}$H-NMR, $^{13}$C-NMR, $^{1}$H-$^{13}$C-HSQC and $^{1}$H-$^{13}$C-HMBC of 6-hydroxy-N-(4'-((4-(2-(((S)-1-((2S,4R)-4-hydroxy-2-((4-(4-methylthiazol-5-yl)benzyl)carbamoyl)pyrrolidin-1-yl)-3,3-dimethyl-1-oxobutan-2-yl)amino)-2-oxoethyl)benzyl)carbamoyl)-4-(4-methylpiperazin-1-yl)-[1,1'-biphenyl]-3-yl)-4-(trifluoromethyl)nicotinamide 8j**

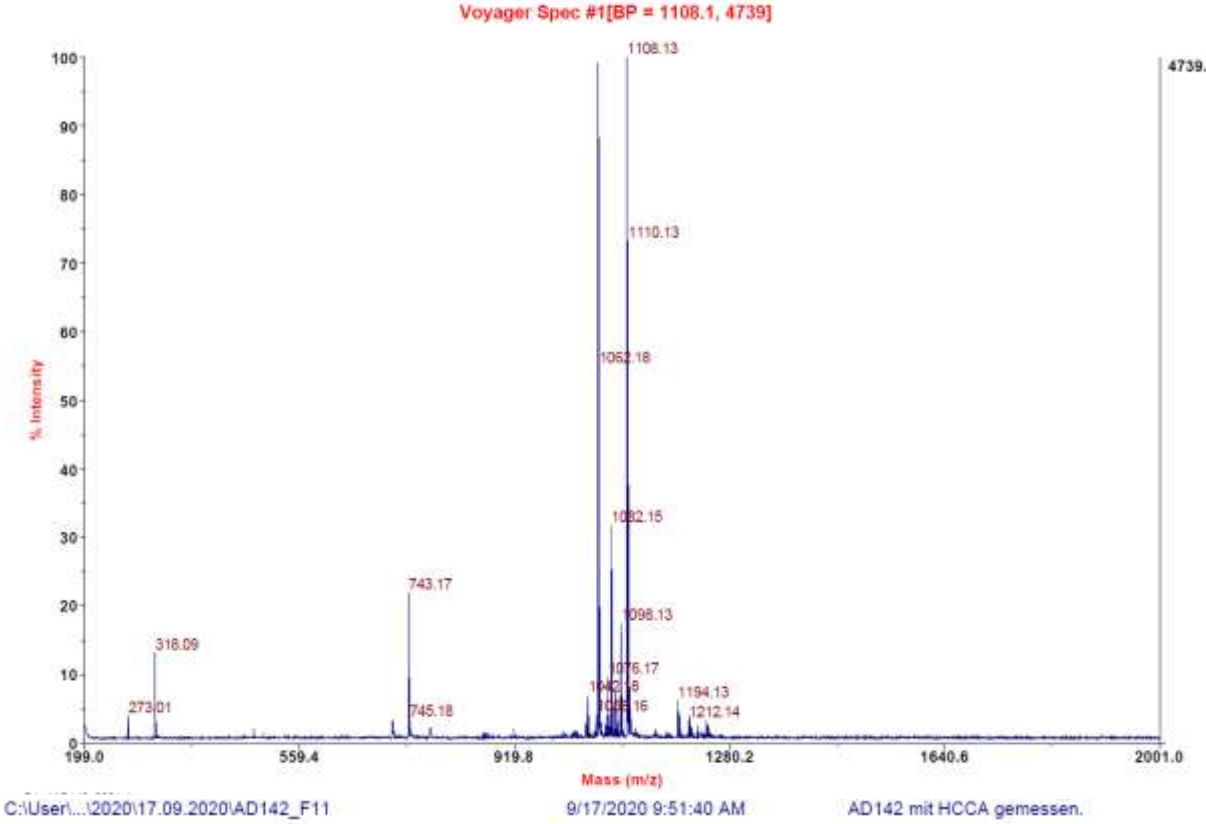

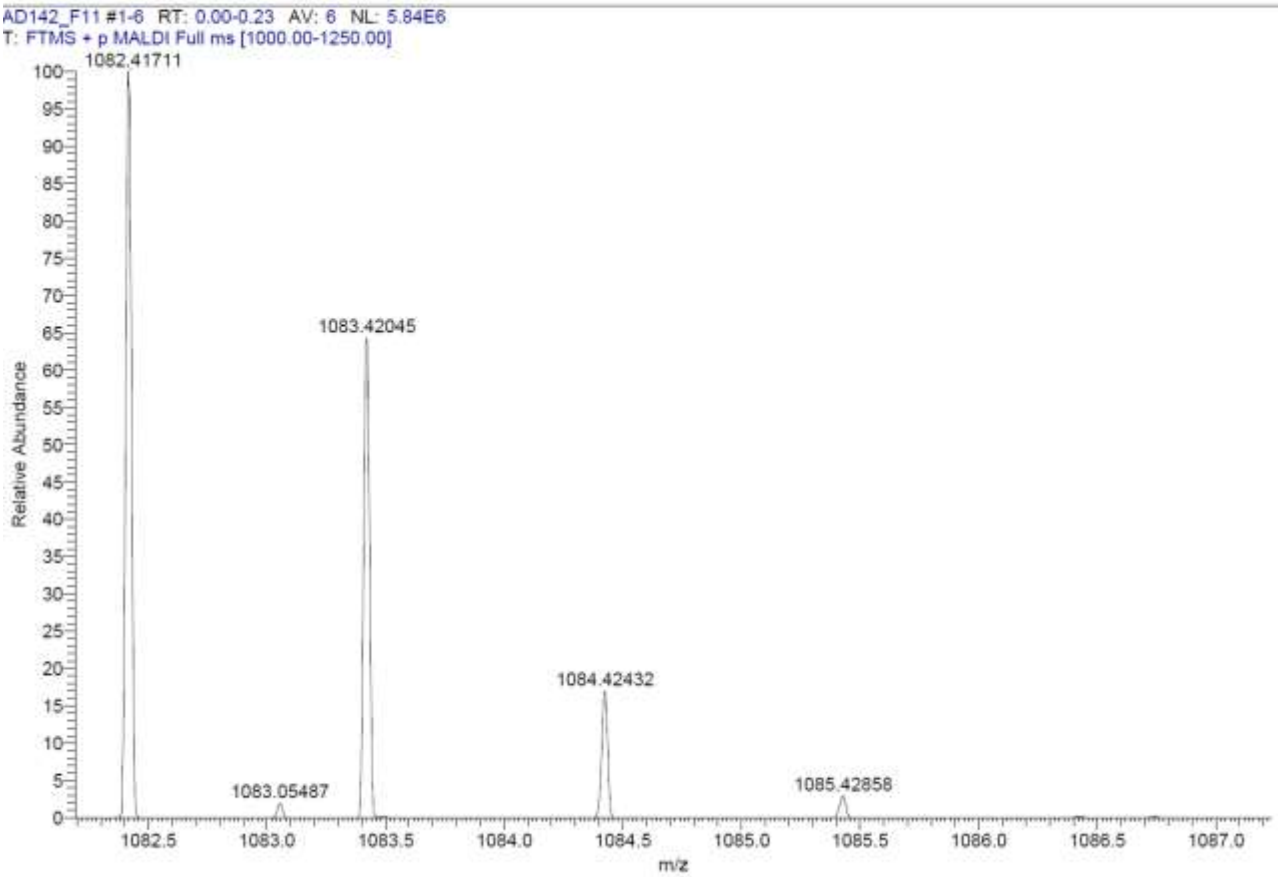

**Signal:** MWD1 A, Sig=254,4 Ref=off

| RT [min] | Type | Width [min] | Area | Height | Area% | Name |
|---|---|---|---|---|---|---|
| 11.560 | VV | 0.2134 | 4870.5161 | 308.0758 | 100.0000 | |
| | | Sum | 4870.5161 | | | |

**Signal:** MWD1 B, Sig=280,4 Ref=off

| RT [min] | Type | Width [min] | Area | Height | Area% | Name |
|---|---|---|---|---|---|---|
| 11.560 | VV | 0.2151 | 5846.5635 | 366.4924 | 100.0000 | |
| | | Sum | 5846.5635 | | | |

**Signal:** MWD1 F, Sig=260,4 Ref=off

| RT [min] | Type | Width [min] | Area | Height | Area% | Name |
|---|---|---|---|---|---|---|
| 11.560 | VV | 0.2134 | 5481.1699 | 346.8157 | 100.0000 | |

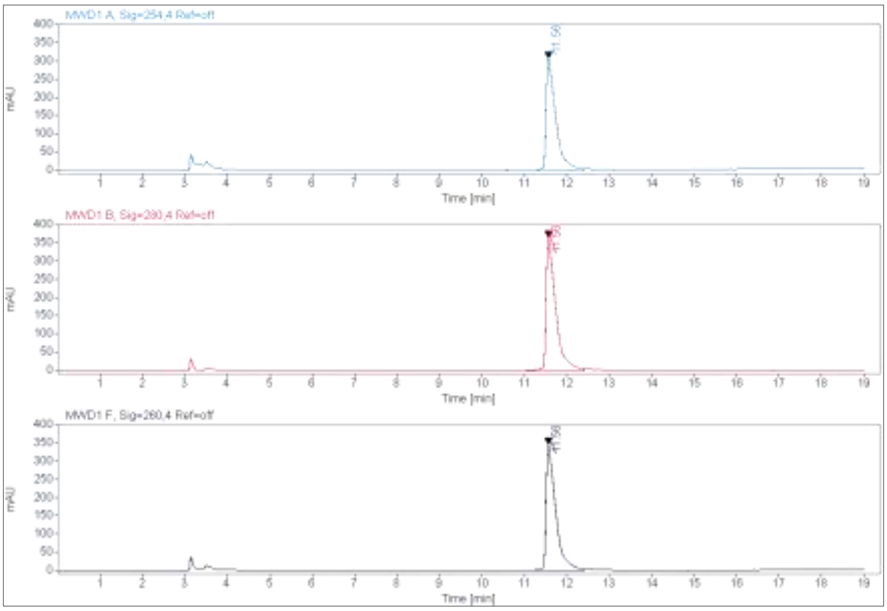

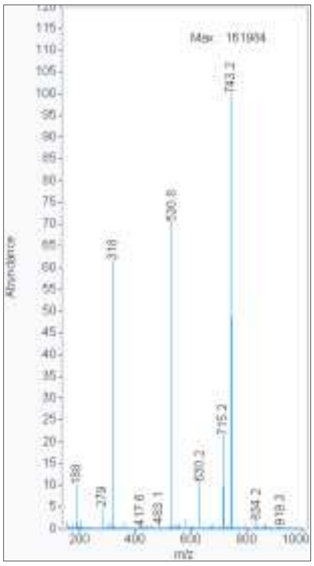

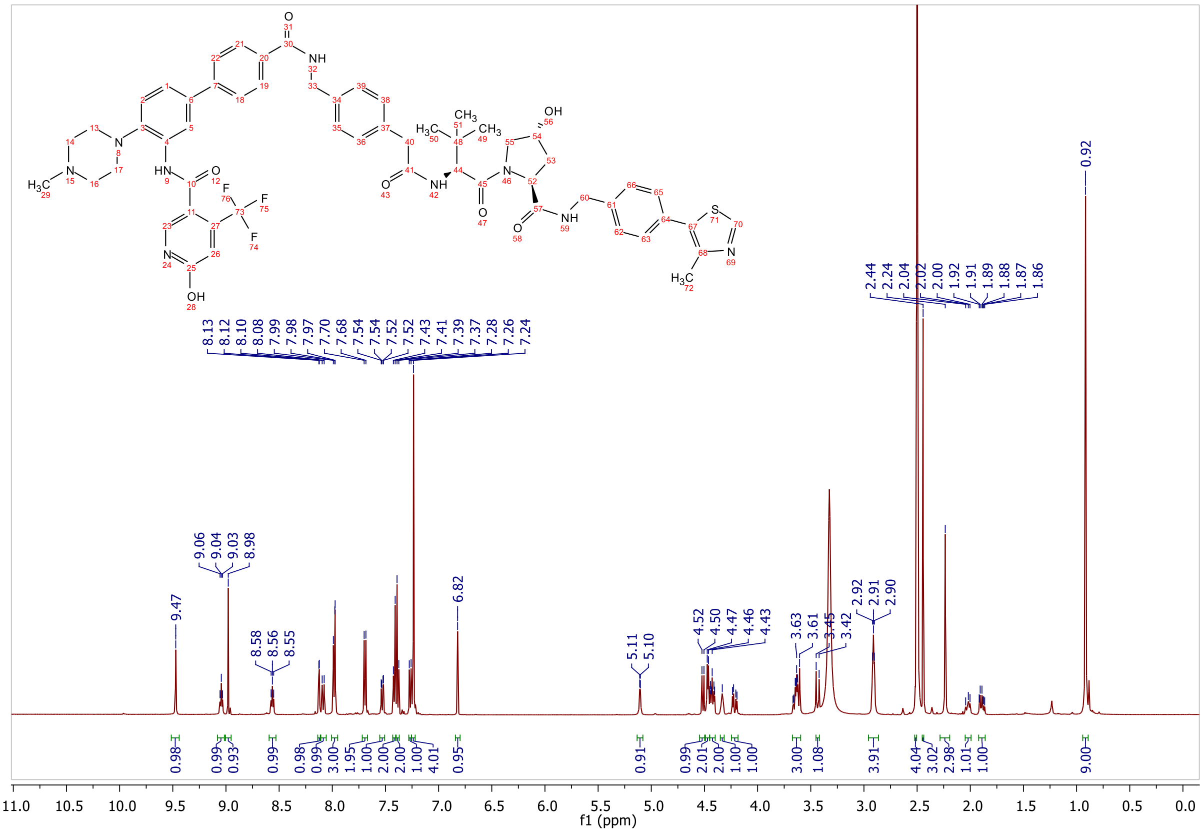

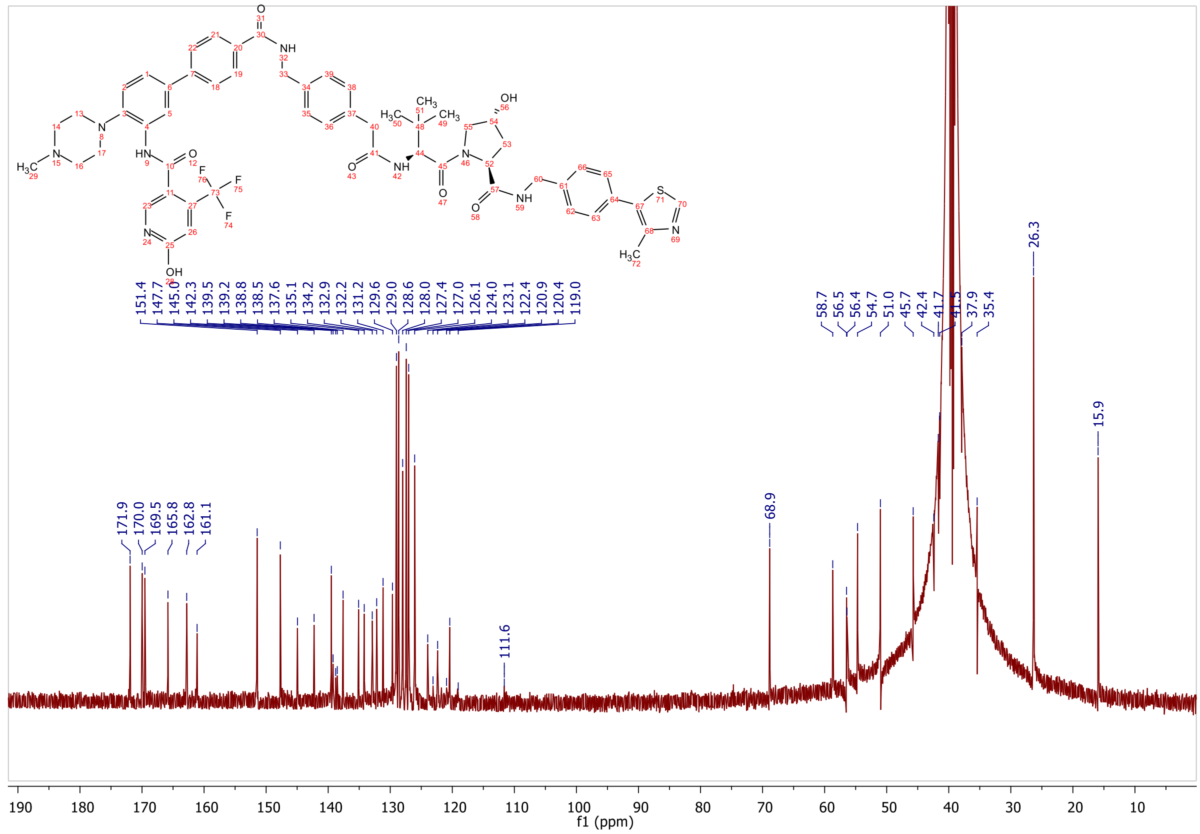

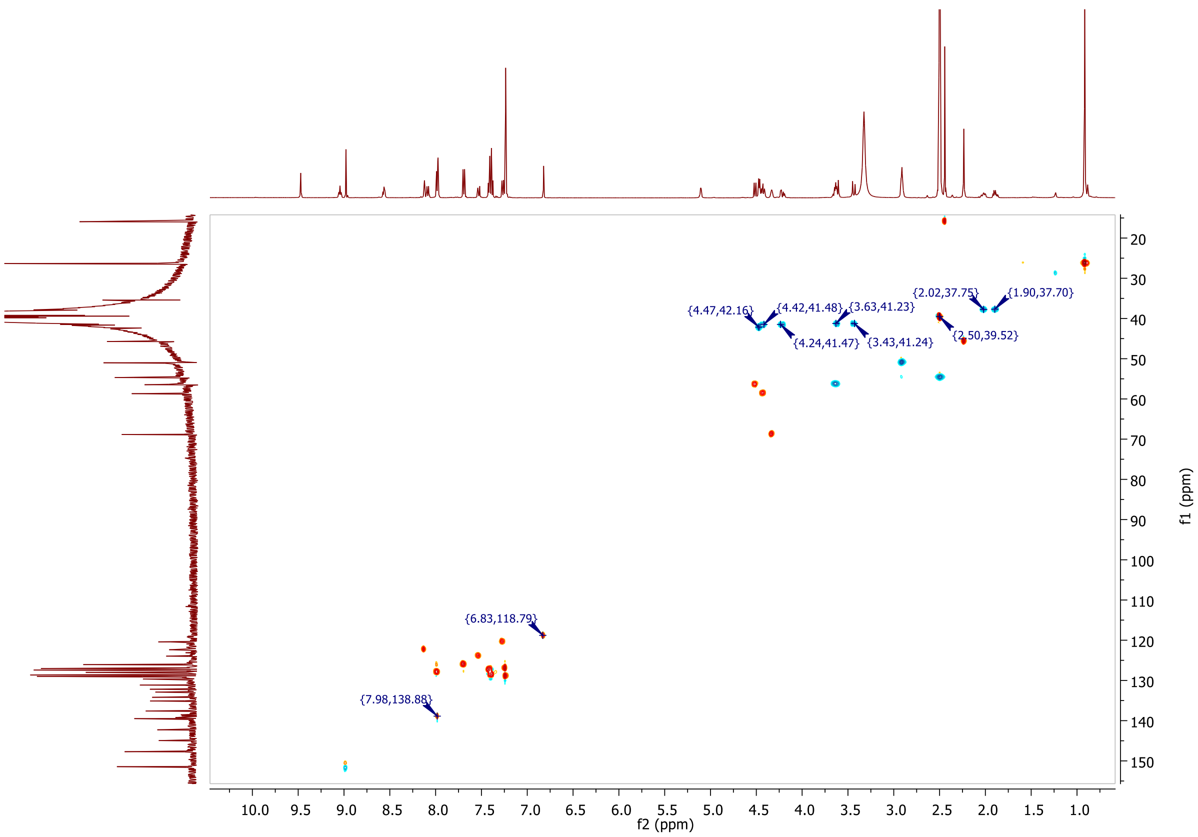
{4.47,42.16}
{4.42,41.48}
{3.63,41.23}
{2.02,37.75}
{1.90,37.70}
{4.24,41.47}
{3.43,41.24}
{2.50,39.52}
{6.83,118.79}
{7.98,138.88}
f2 (ppm)
f1 (ppm)

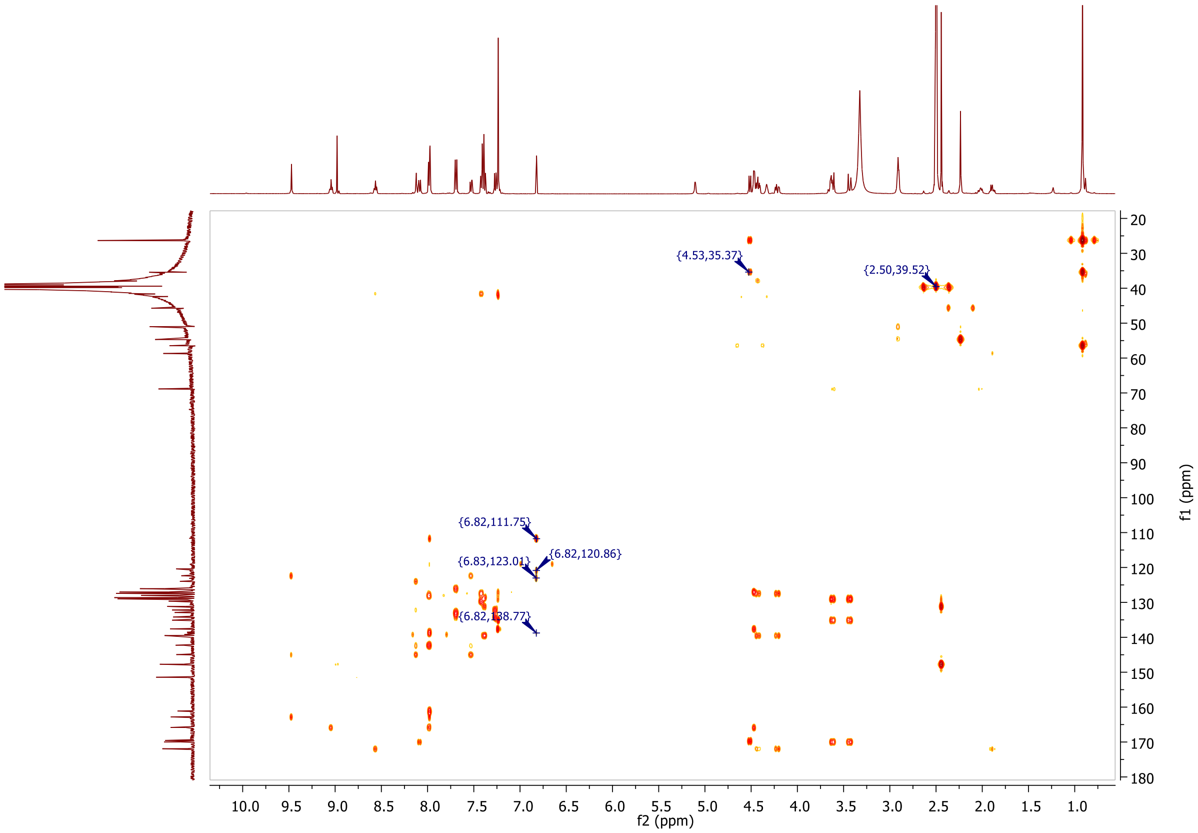
{4.53,35.37}
{2.50,39.52}
{6.82,111.75}
{6.82,120.86}
{6.83,123.01}
{6.82,138.77}
f2 (ppm)
f1 (ppm)

**MALDI, HRMS, HPLC, $^{1}$H-NMR of N-(4'-((2-(2-(4-((2R,3S,4R,5S)-3-(3-chloro-2-fluorophenyl)-4-(4-chloro-2-fluorophenyl)-4-cyano-5-neopentylpyrrolidine-2-carboxamido)-3-methoxybenzamido)ethoxy)ethyl)carbamoyl)-4-(4-methylpiperazin-1-yl)-[1,1'-biphenyl]-3-yl)-6-hydroxy-4-(trifluoromethyl)nicotinamide 9a**

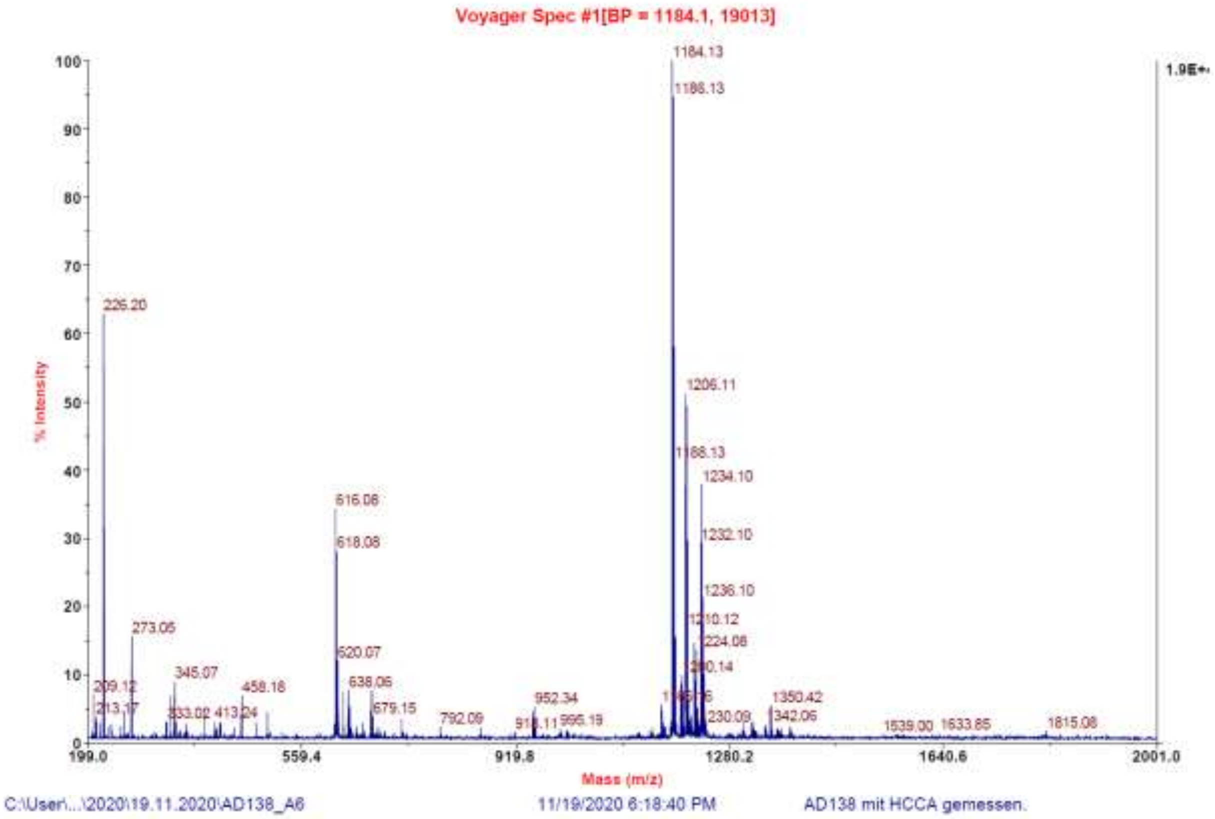

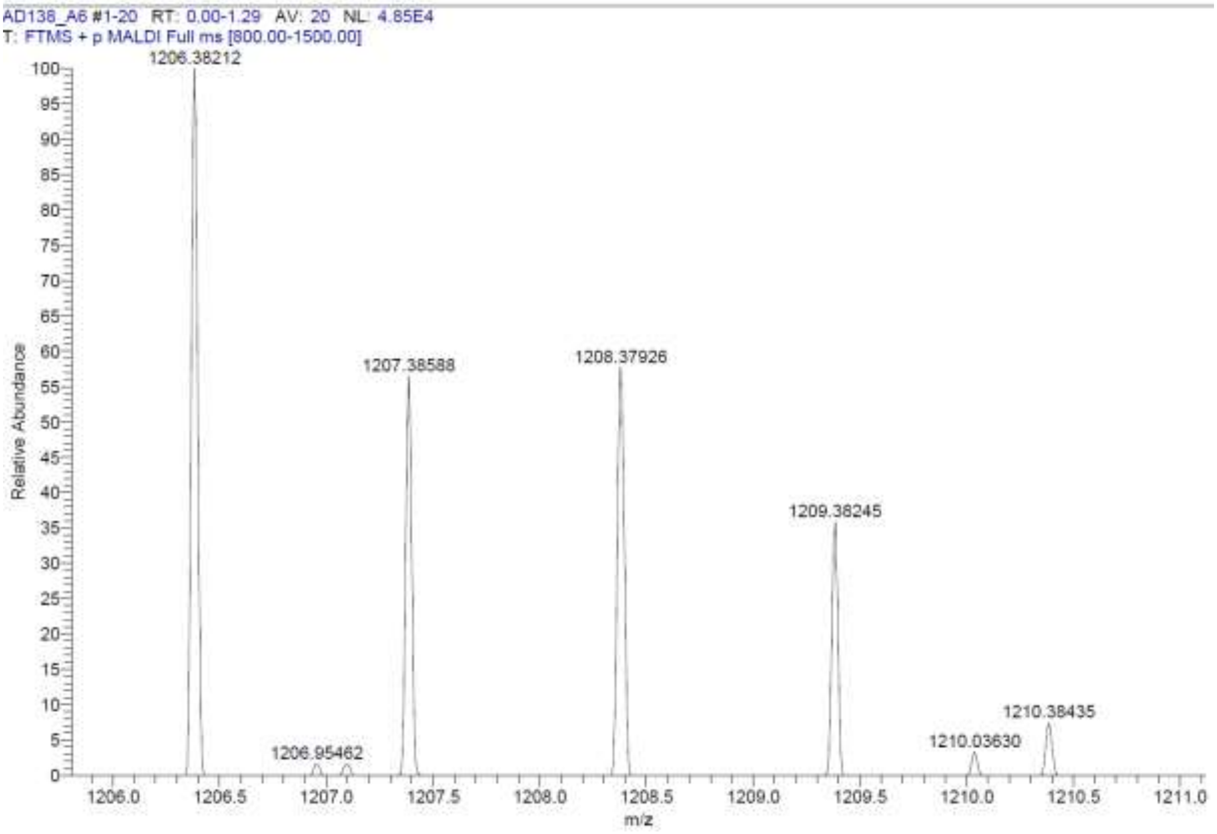

**Signal:** MWD1 A, Sig=254,4 Ref=off

| RT [min] | Type | Width [min] | Area | Height | Area% |
|---|---|---|---|---|---|
| 12.966 | MM | 0.1809 | 45.8666 | 4.2269 | 1.5833 |
| 14.363 | VV | 0.2753 | 2851.0457 | 147.6335 | 98.4167 |
| | | Sum | 2896.9123 | | |

**Signal:** MWD1 B, Sig=280,4 Ref=off

| RT [min] | Type | Width [min] | Area | Height | Area% |
|---|---|---|---|---|---|
| 12.960 | MM | 0.2518 | 15.9346 | 1.0546 | 0.5114 |
| 14.363 | VV | 0.2623 | 3099.7249 | 162.6616 | 99.4886 |
| | | Sum | 3115.6594 | | |

**Signal:** MWD1 F, Sig=260,4 Ref=off

| RT [min] | Type | Width [min] | Area | Height | Area% |
|---|---|---|---|---|---|
| 12.962 | MM | 0.2060 | 54.8831 | 4.4413 | 1.5802 |
| 14.363 | VV | 0.2919 | 3418.3169 | 178.0980 | 98.4198 |

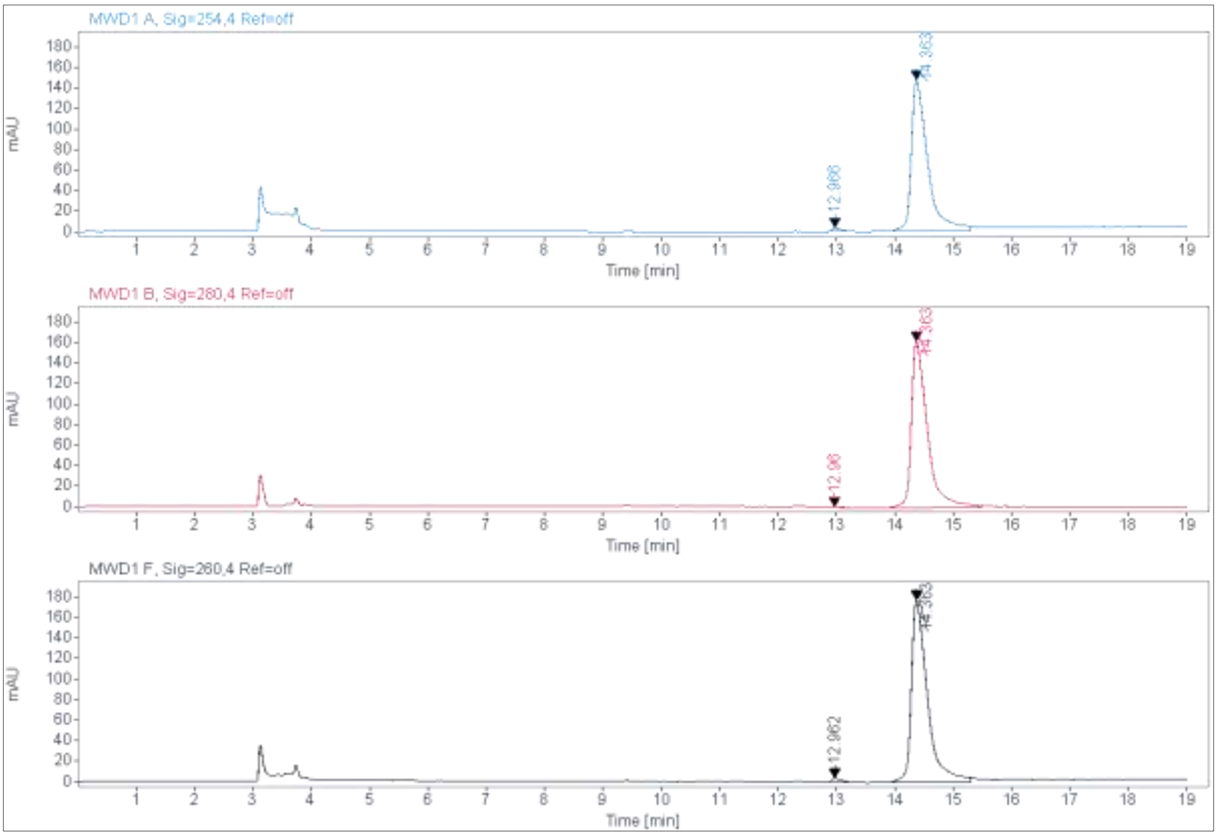

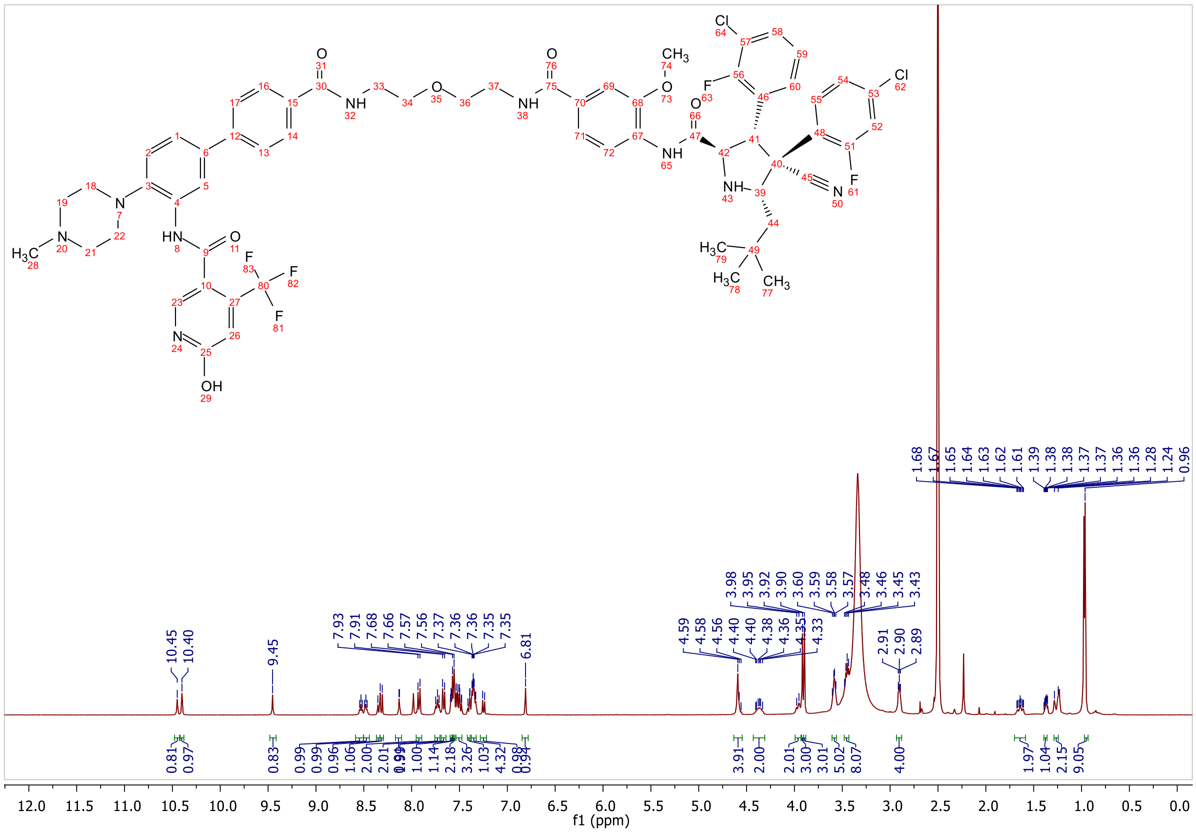

**MALDI, HRMS, HMRS (zoom), HPLC, $^{1}$H-NMR of N-(4'-((1-(4-((2R,3S,4R,5S)-3-(3-chloro-2-fluorophenyl)-4-(4-chloro-2-fluorophenyl)-4-cyano-5-neopentylpyrrolidine-2-carboxamido)-3-methoxyphenyl)-1-oxo-5,8,11,14,17,20,23-heptaoxa-2-azapentacosan-25-yl)carbamoyl)-4-(4-methylpiperazin-1-yl)-[1,1'-biphenyl]-3-yl)-6-hydroxy-4-(trifluoromethyl)nicotinamide 9b**

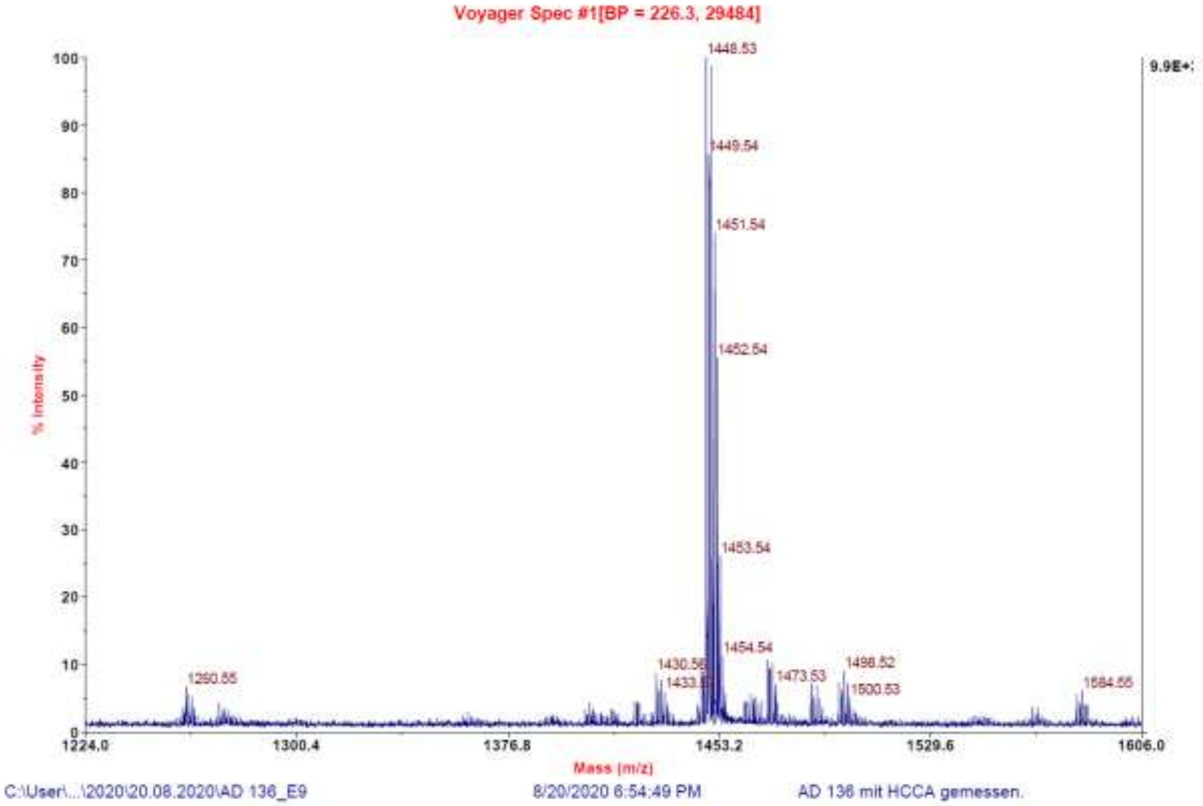

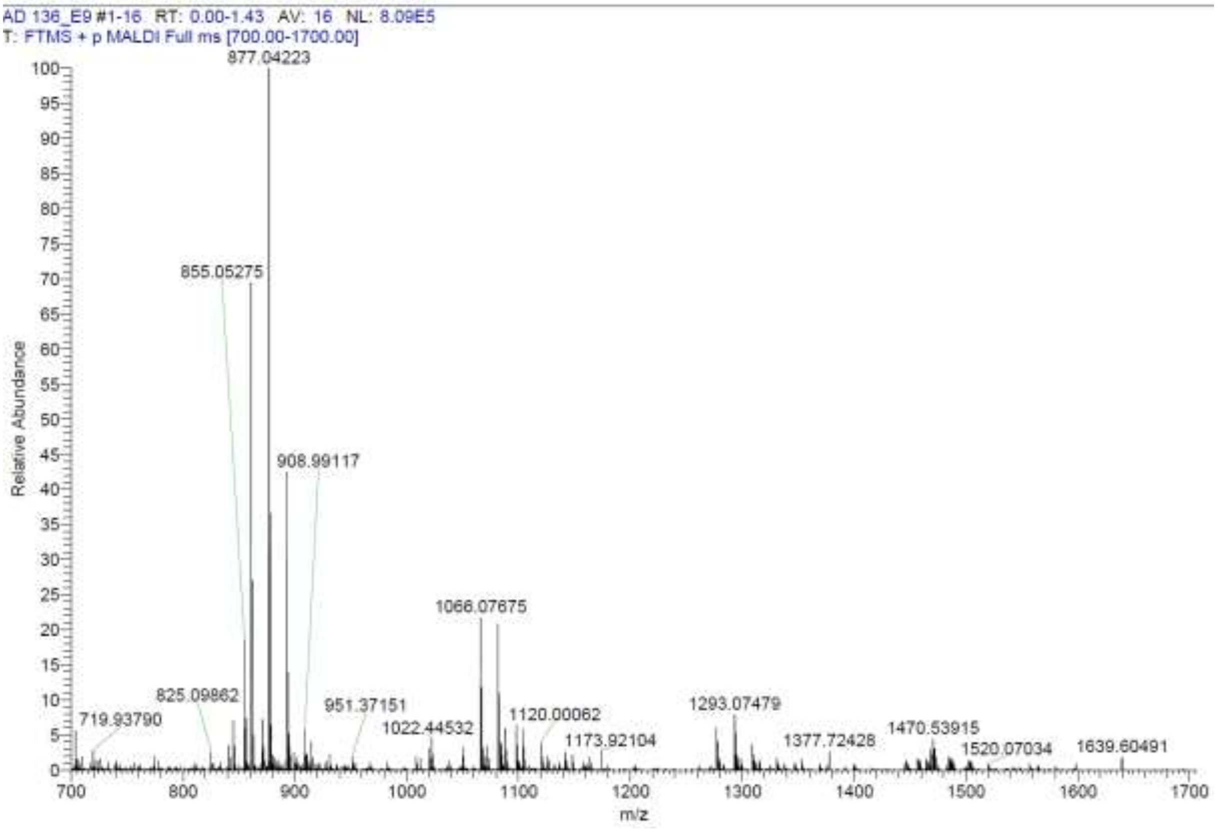

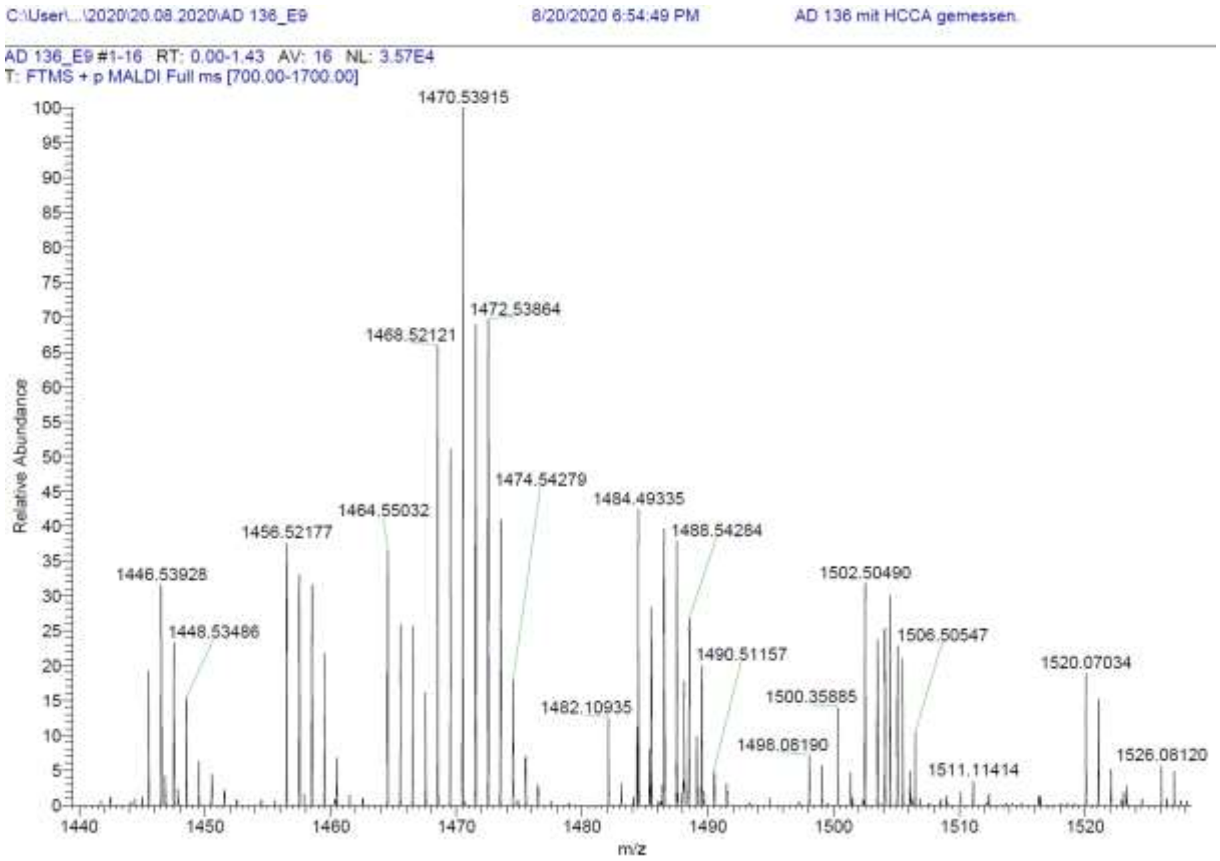

| RT [min] | Type | Width [min] | Area | Height | Area% |
|---|---|---|---|---|---|
| 13.388 | MM | 0.2391 | 223.4376 | 15.5738 | 3.8297 |
| 14.631 | VV | 0.4209 | 5610.9448 | 180.3722 | 96.1703 |
| | | Sum | 5834.3824 | | |

**Signal:** MWD1 B, Sig=280,4 Ref=off

| RT [min] | Type | Width [min] | Area | Height | Area% |
|---|---|---|---|---|---|
| 13.385 | MM | 0.1997 | 178.9707 | 14.9336 | 3.1186 |
| 14.630 | VV | 0.3882 | 5559.8906 | 195.2569 | 96.8814 |
| | | Sum | 5738.8614 | | |

**Signal:** MWD1 F, Sig=260,4 Ref=off

| RT [min] | Type | Width [min] | Area | Height | Area% |
|---|---|---|---|---|---|
| 13.386 | MM | 0.2384 | 306.7542 | 21.4423 | 4.6729 |
| 14.631 | VV | 0.4019 | 6257.7744 | 214.1512 | 95.3271 |

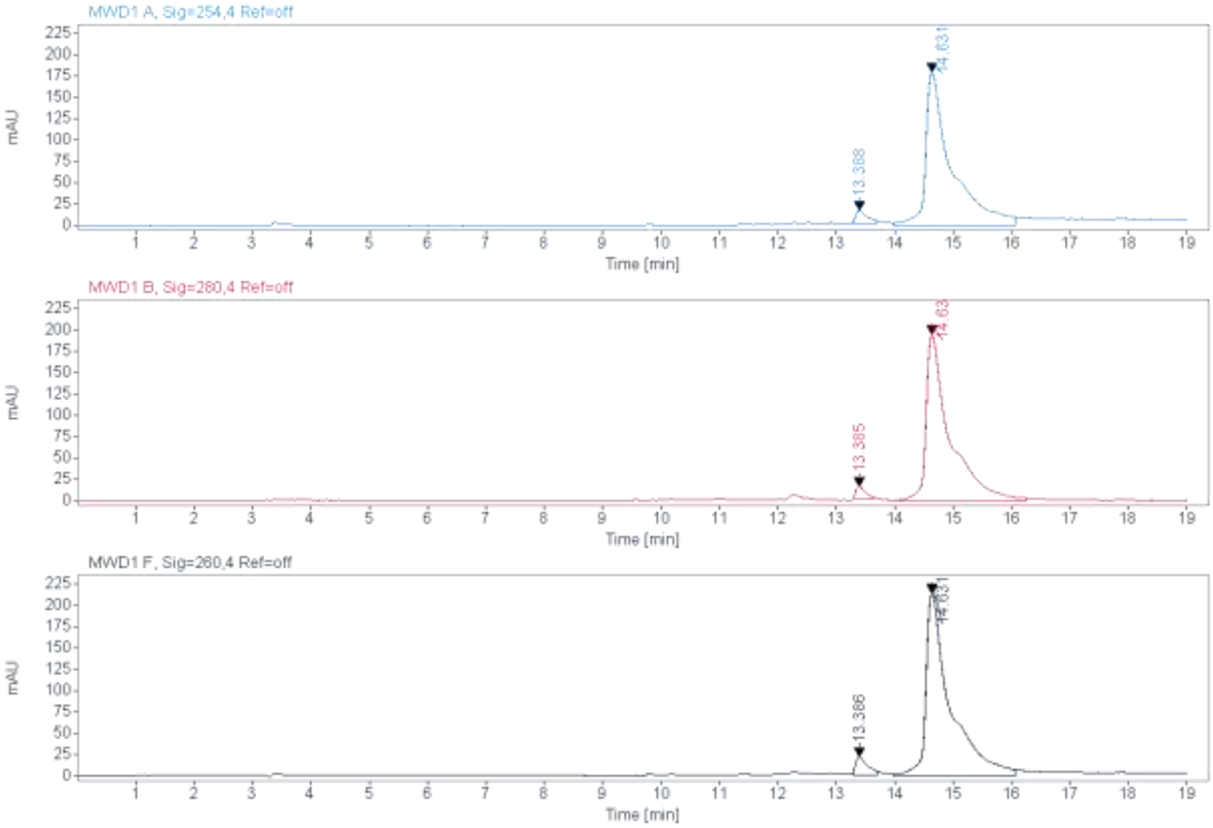

MWD1 A, Sig=254,4 Ref=off
13.388
14.631
MWD1 B, Sig=280,4 Ref=off
13.385
14.63
MWD1 F, Sig=260,4 Ref=off
13.386
14.631
mAU
Time [min]

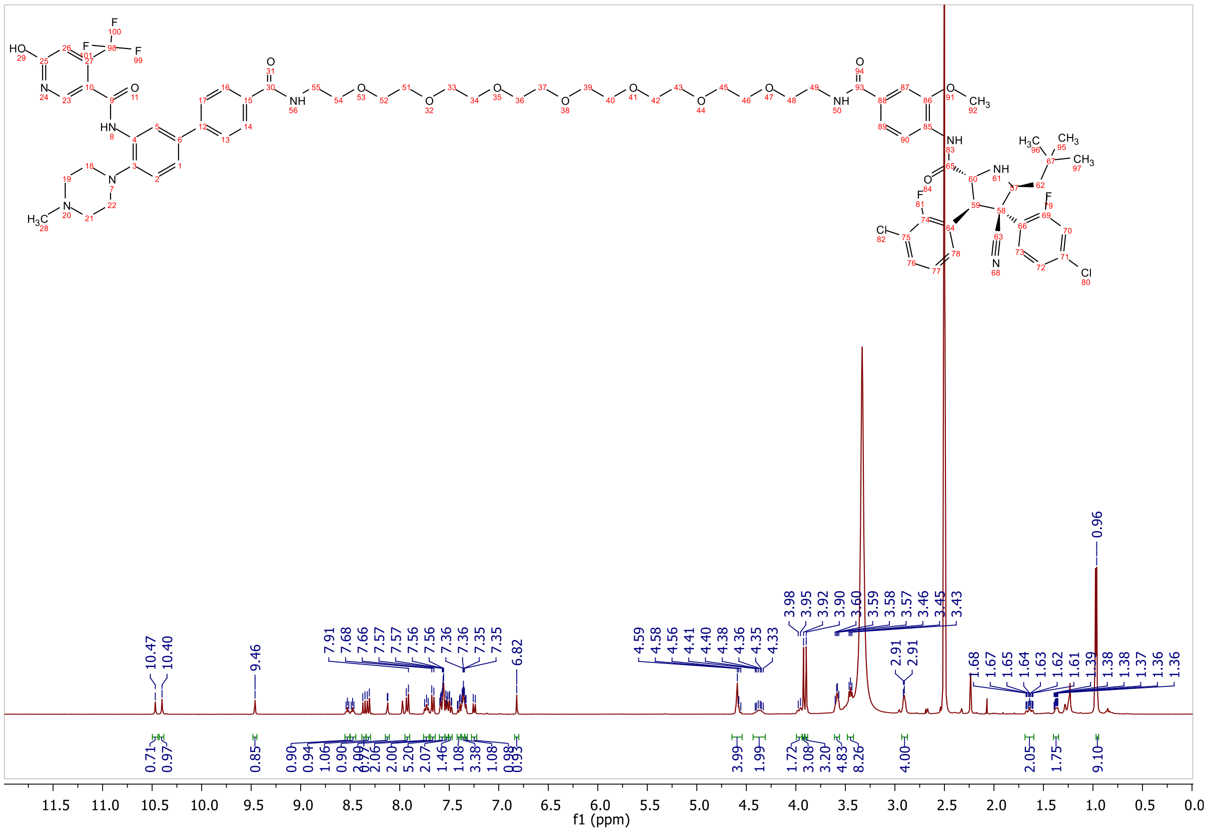

f1 (ppm)

**MALDI, HRMS, HPLC, $^{1}$H-NMR of N-(4'-((4-((4-((2R,3S,4R,5S)-3-(3-chloro-2-fluorophenyl)-4-(4-chloro-2-fluorophenyl)-4-cyano-5-neopentylpyrrolidine-2-carboxamido)-3-methoxybenzamido)methyl)benzyl)carbamoyl)-4-(4-methylpiperazin-1-yl)-[1,1'-biphenyl]-3-yl)-6-hydroxy-4-(trifluoromethyl)nicotinamide 9c**

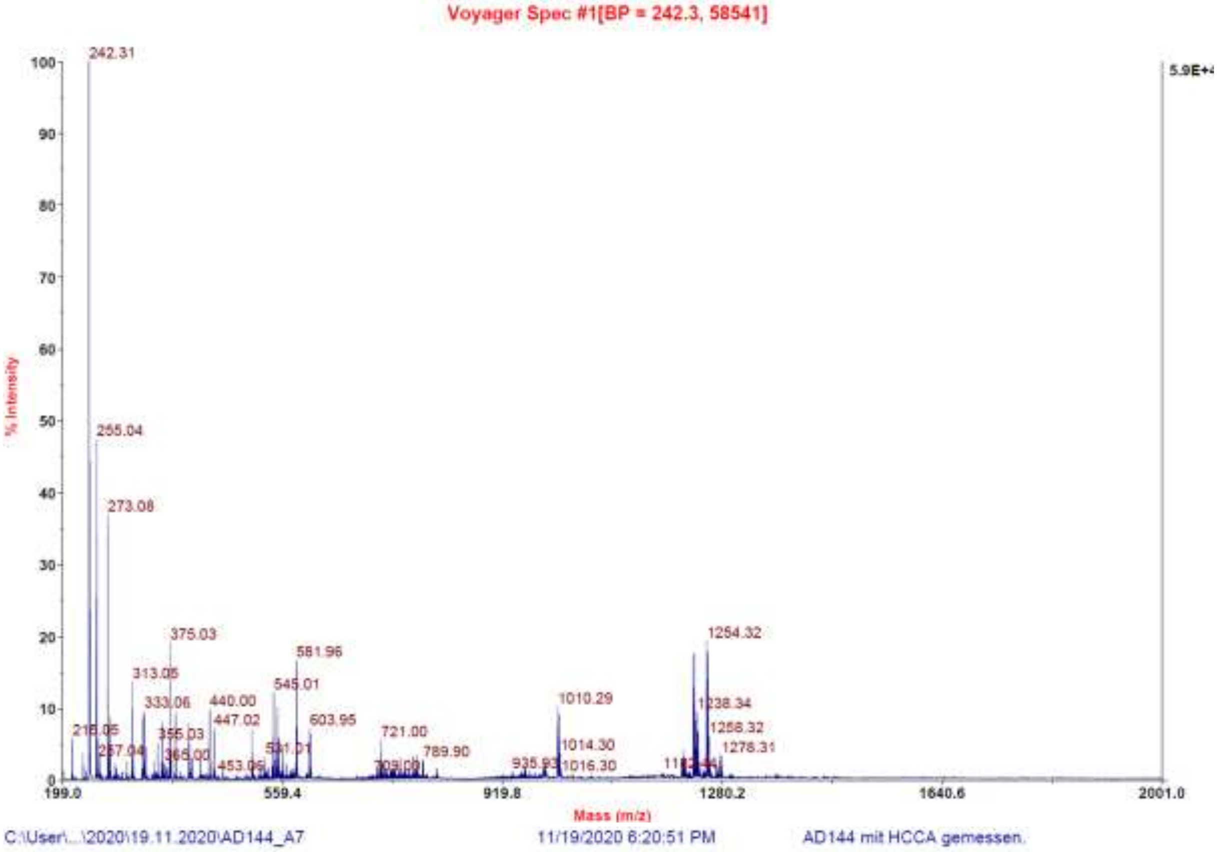

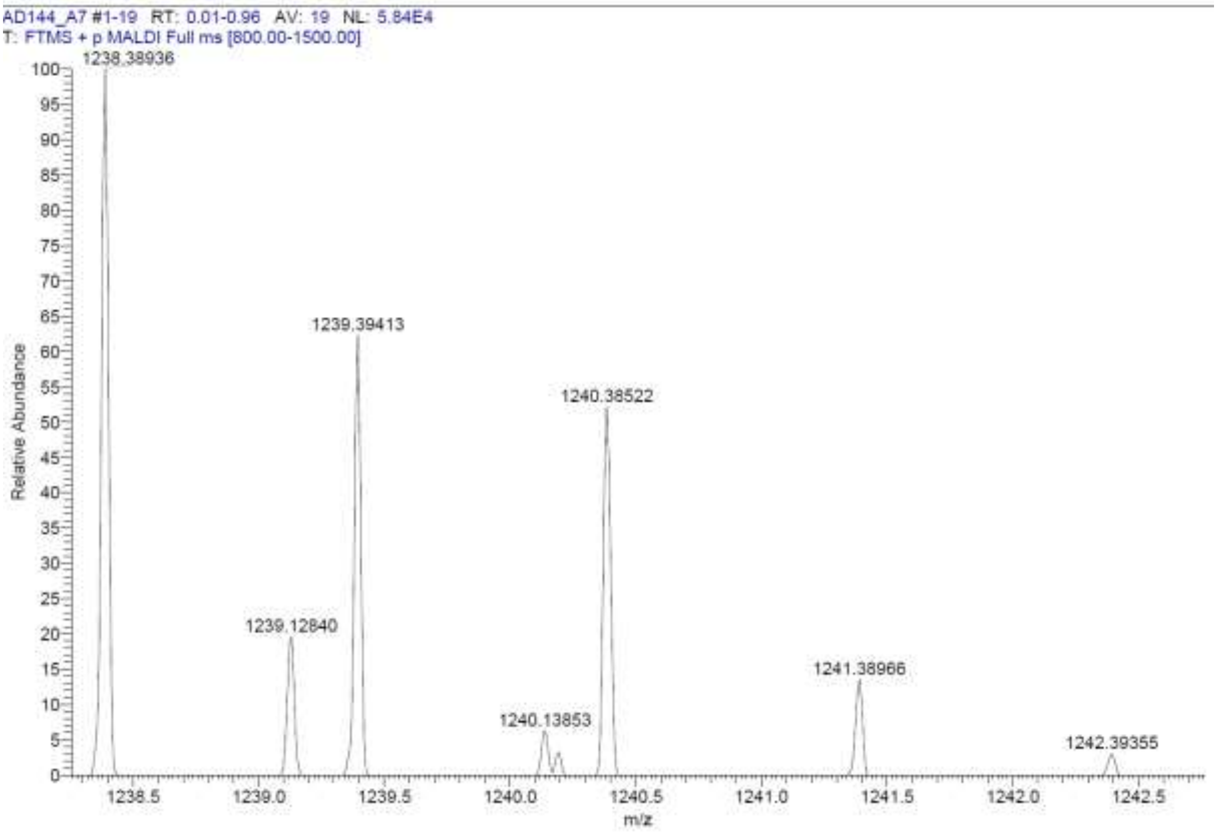

**Signal:** MWD1 A, Sig=254,4 Ref=off

| RT [min] | Type | Width [min] | Area | Height | Area% | Name |
|---|---|---|---|---|---|---|
| 12.962 | MM | 0.1568 | 35.1322 | 3.7335 | 1.8300 | |
| 14.724 | VV | 0.3252 | 1884.6840 | 75.5157 | 98.1700 | |
| | | Sum | 1919.8162 | | | |

**Signal:** MWD1 B, Sig=280,4 Ref=off

| RT [min] | Type | Width [min] | Area | Height | Area% | Name |
|---|---|---|---|---|---|---|
| 14.722 | VV | 0.3226 | 2160.1218 | 87.6254 | 100.0000 | |
| | | Sum | 2160.1218 | | | |

**Signal:** MWD1 F, Sig=260,4 Ref=off

| RT [min] | Type | Width [min] | Area | Height | Area% | Name |
|---|---|---|---|---|---|---|
| 12.967 | MM | 0.1599 | 33.1664 | 3.4567 | 1.4688 | |
| 14.724 | VV | 0.3246 | 2224.9216 | 89.0201 | 98.5312 | |

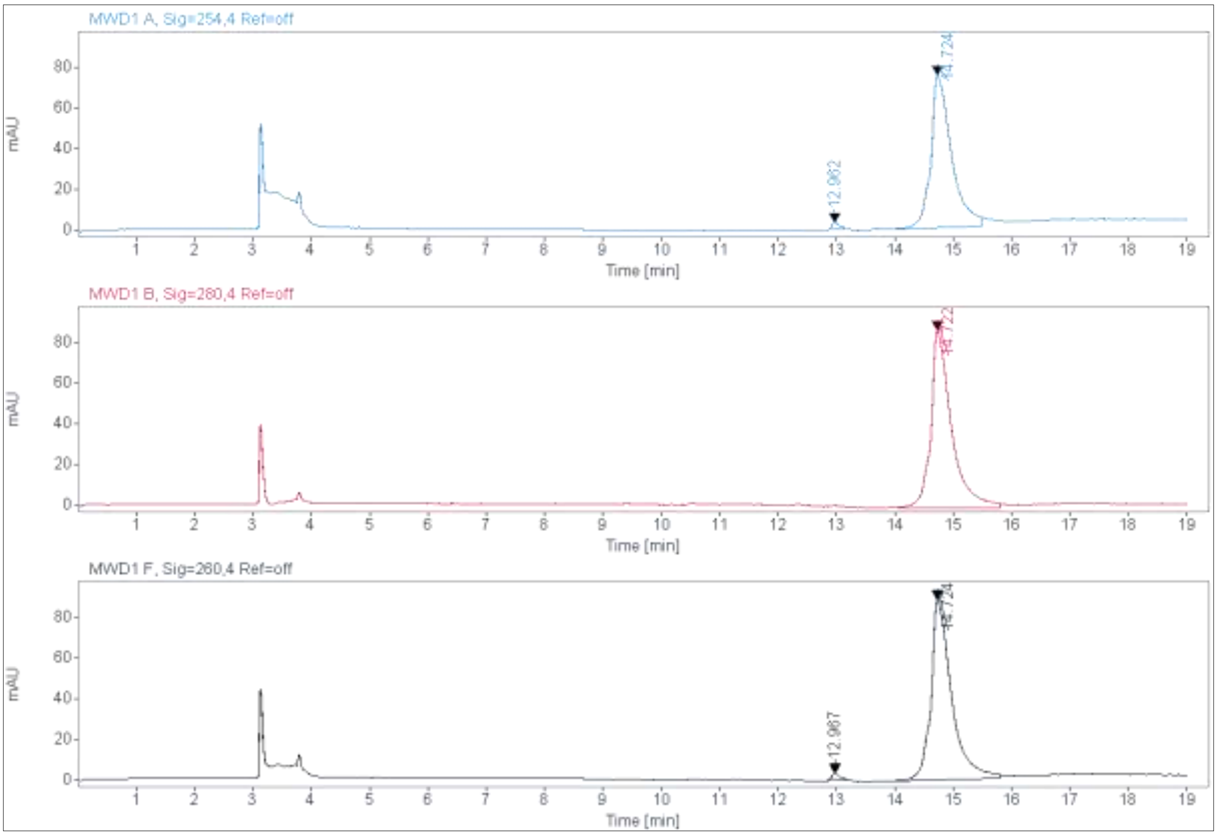

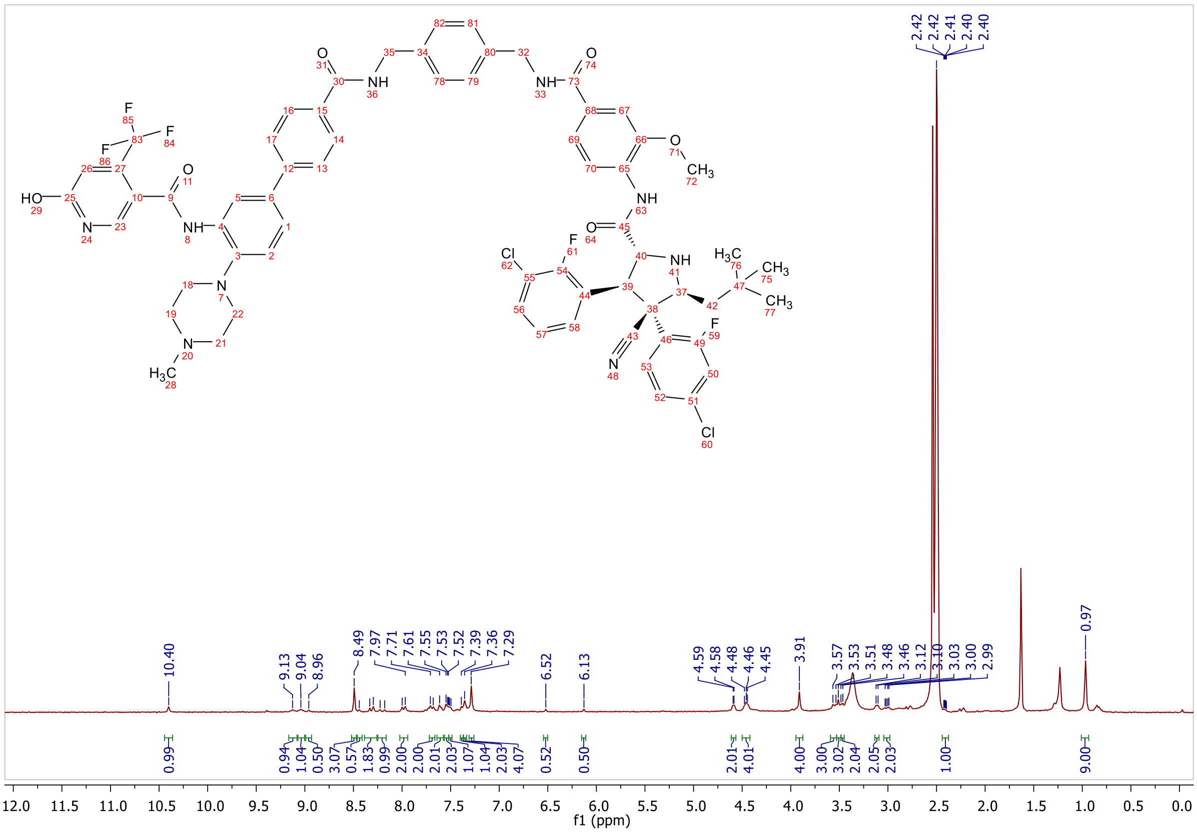

10.40
9.13
9.04
8.96
8.49
7.97
7.71
7.61
7.55
7.53
7.52
7.39
7.36
7.29
6.52
6.13
4.59
4.58
4.48
4.46
4.45
3.91
3.57
3.53
3.51
3.48
3.46
3.12
3.10
3.03
3.00
2.99
2.42
2.42
2.41
2.40
2.40
0.97
0.99
0.94
1.04
0.50
3.07
0.57
1.83
0.99
2.00
2.00
2.01
2.03
1.07
1.04
2.03
4.07
0.52
0.50
2.01
4.01
4.00
3.00
3.02
2.04
2.05
2.03
1.00
9.00
12.0 11.5 11.0 10.5 10.0 9.5 9.0 8.5 8.0 7.5 7.0 6.5 6.0 5.5 5.0 4.5 4.0 3.5 3.0 2.5 2.0 1.5 1.0 0.5 0.0
f1 (ppm)

### 1.5 Appendix for intermediates 11-16a-g

#### $^1$H-NMR, $^{13}$C-NMR and ESI of 2-Bromo-6,7-dihydro-5H-pyrrolo[1,2-a]imidazole 11

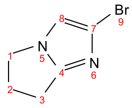

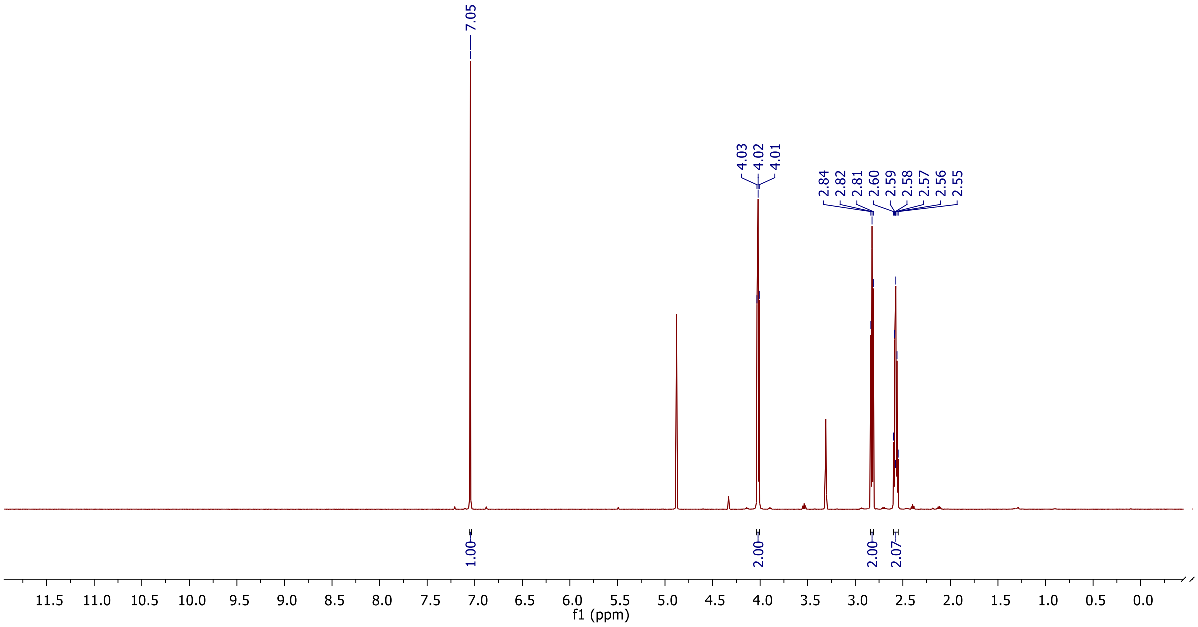

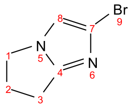

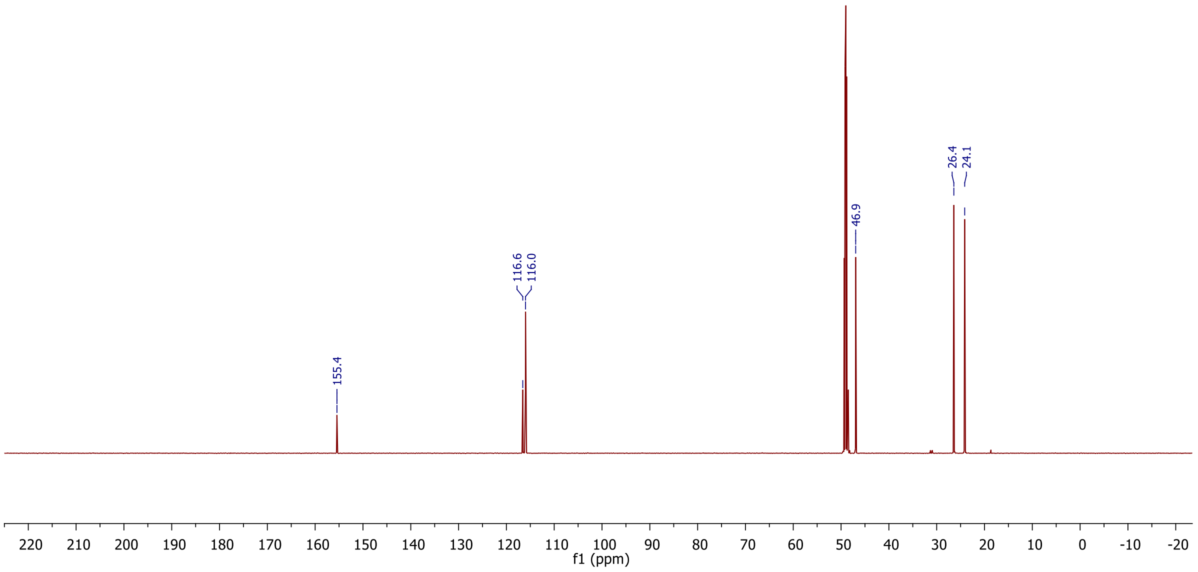

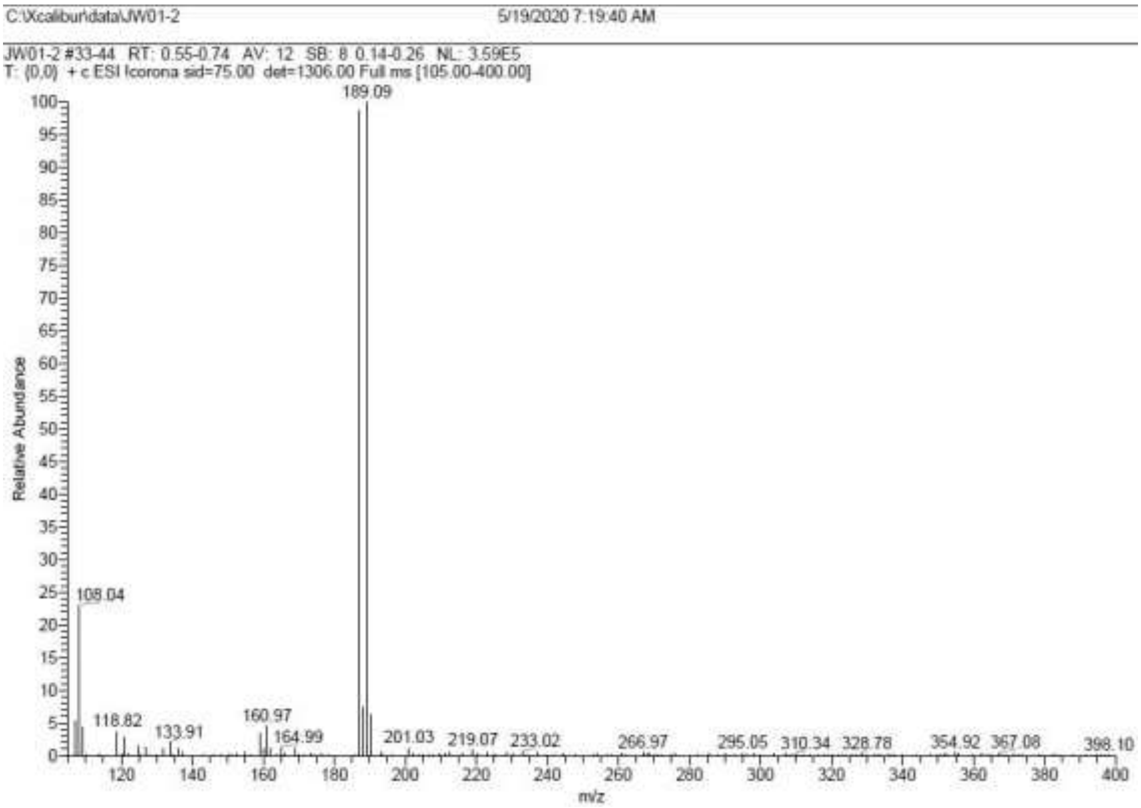
C:\Xcalibur\data\JW01-2
5/19/2020 7:19:40 AM
JW01-2 #33-44 RT: 0.55-0.74 AV: 12 SB: 8 0.14-0.26 NL: 3.59E5
T: {0,0} + c ESI Icorona sid=75.00 det=1306.00 Full ms [105.00-400.00]
Relative Abundance
m/z
189.09
108.04
118.82
133.91
160.97
164.99
201.03
219.07
233.02
266.97
295.05
310.34
328.78
354.92
367.08
398.10

**ESI, $^{1}$H-NMR and $^{13}$C-NMR of 5-(6,7-Dihydro-5H-pyrrolo[1,2-a]imidazol-2-yl)-2-methoxybenzonitrile 12**

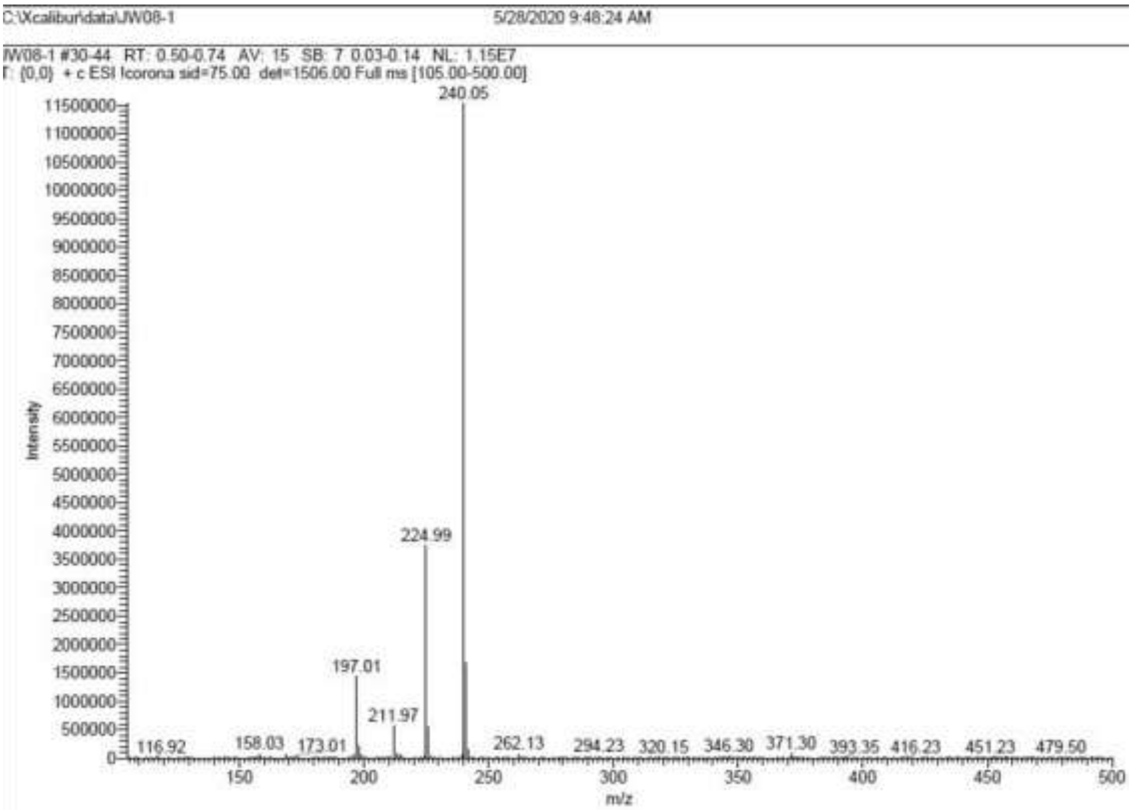

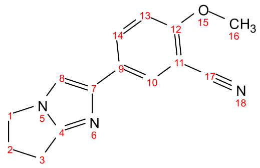

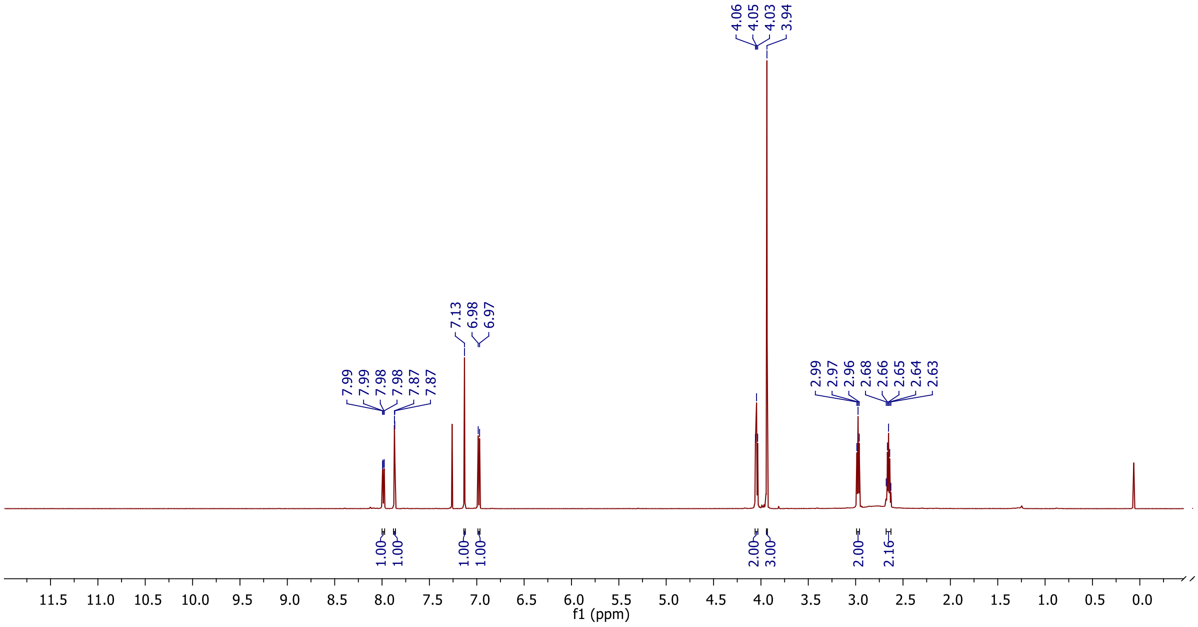

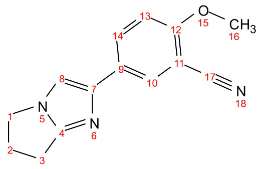

O
CH3
N
N
N

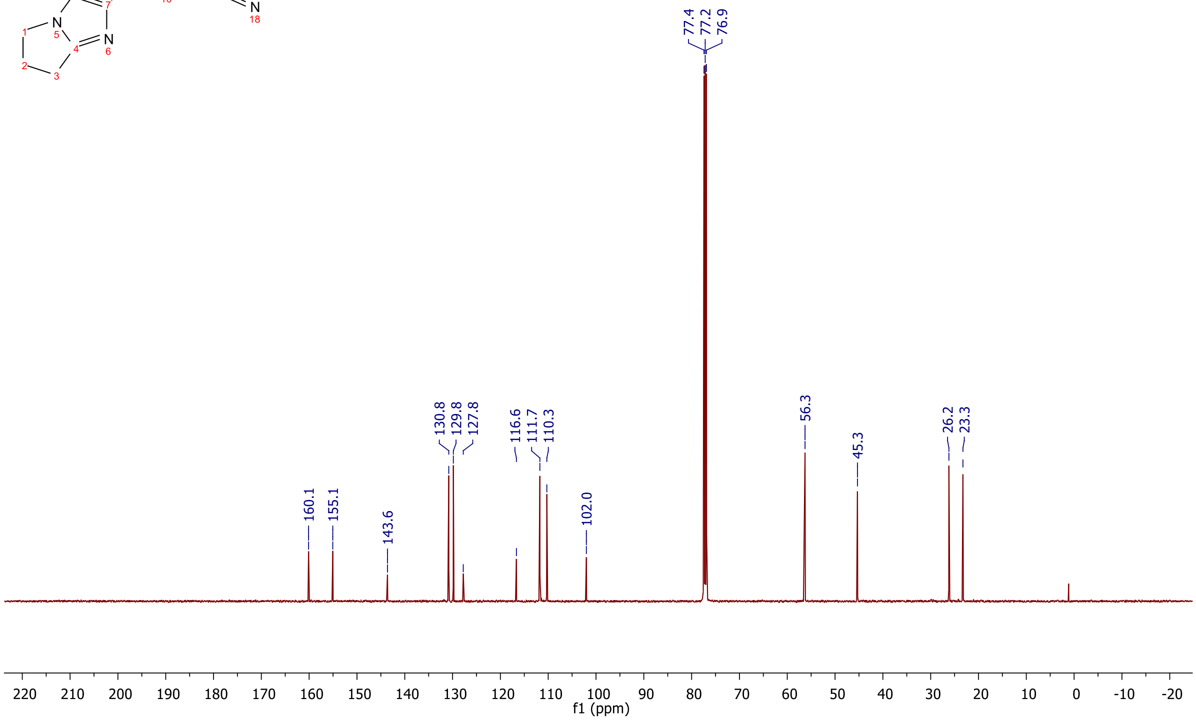

77.4
77.2
76.9
160.1
155.1
143.6
130.8
129.8
127.8
116.6
111.7
110.3
102.0
56.3
45.3
26.2
23.3
220
210
200
190
180
170
160
150
140
130
120
110
100
90
80
70
60
50
40
30
20
10
0
-10
-20
f1 (ppm)

**ESI, $^{1}$H-NMR and $^{13}$C-NMR of (5-(6,7-dihydro-5H-pyrrolo[1,2-a]imidazol-2-yl)-2-methoxyphenyl)methanamine 13**

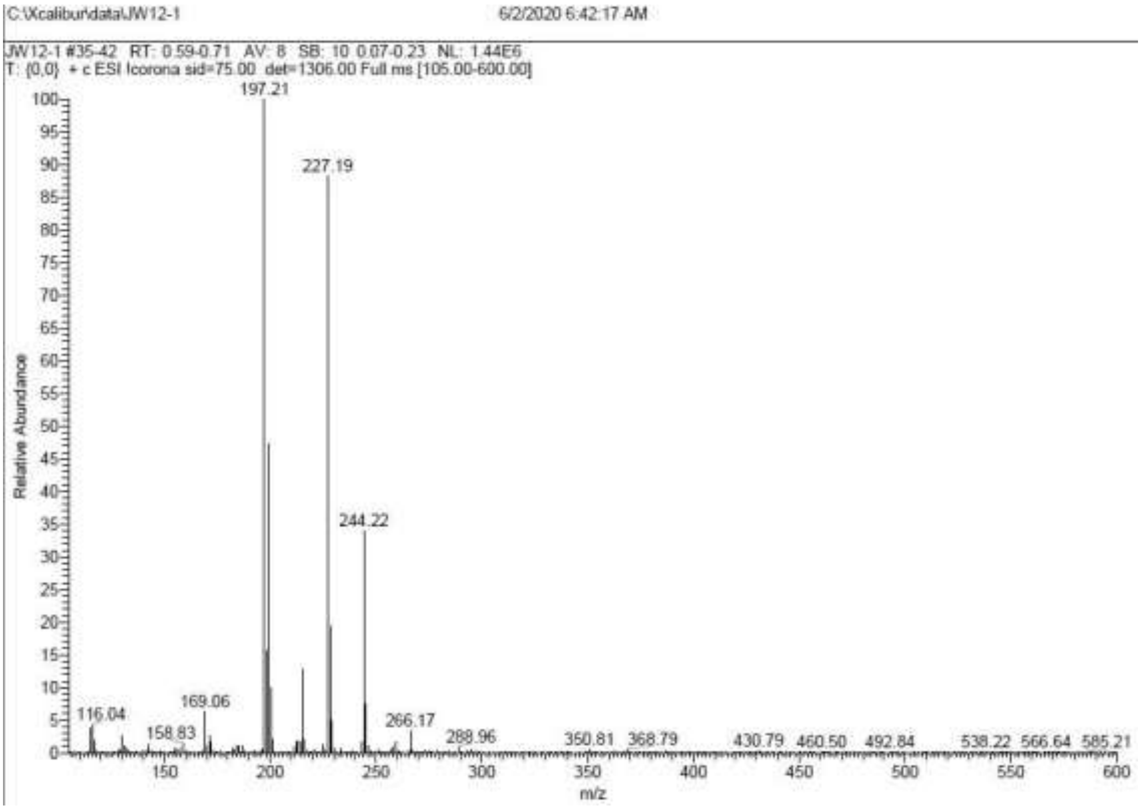

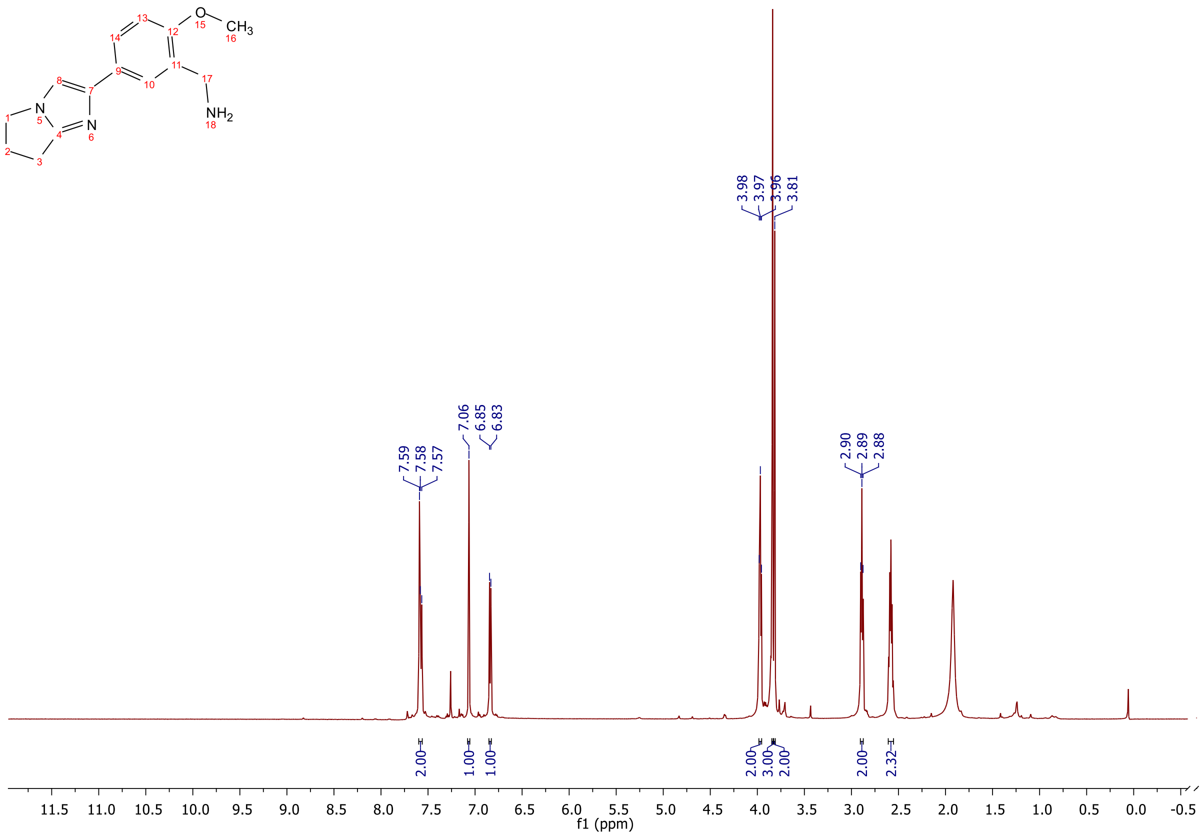

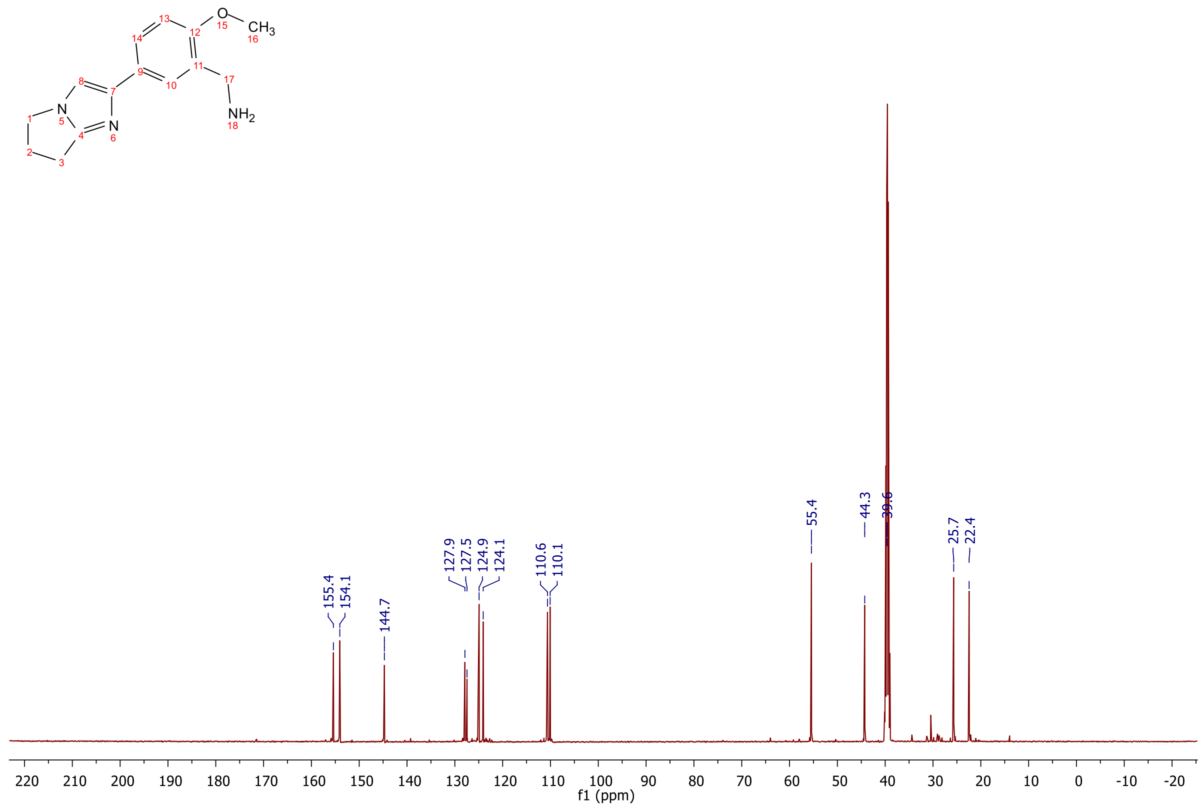
O
15
CH3
16
13
12
14
11
17
8
9
10
7
NH2
18
N
1
5
N
6
4
2
3
155.4
154.1
144.7
127.9
127.5
124.9
124.1
110.6
110.1
55.4
44.3
39.6
25.7
22.4
220
210
200
190
180
170
160
150
140
130
120
110
100
90
80
70
60
50
40
30
20
10
0
-10
-20
f1 (ppm)

**[1]H-NMR, [13]C-NMR, ESI, HRMS and HPLC of 2-(3,4-Dichlorophenyl)-N-(5-(6,7-dihydro-5H-pyrrolo[1,2-a]imidazol-2-yl)-2-methoxybenzyl)acetamide 14**

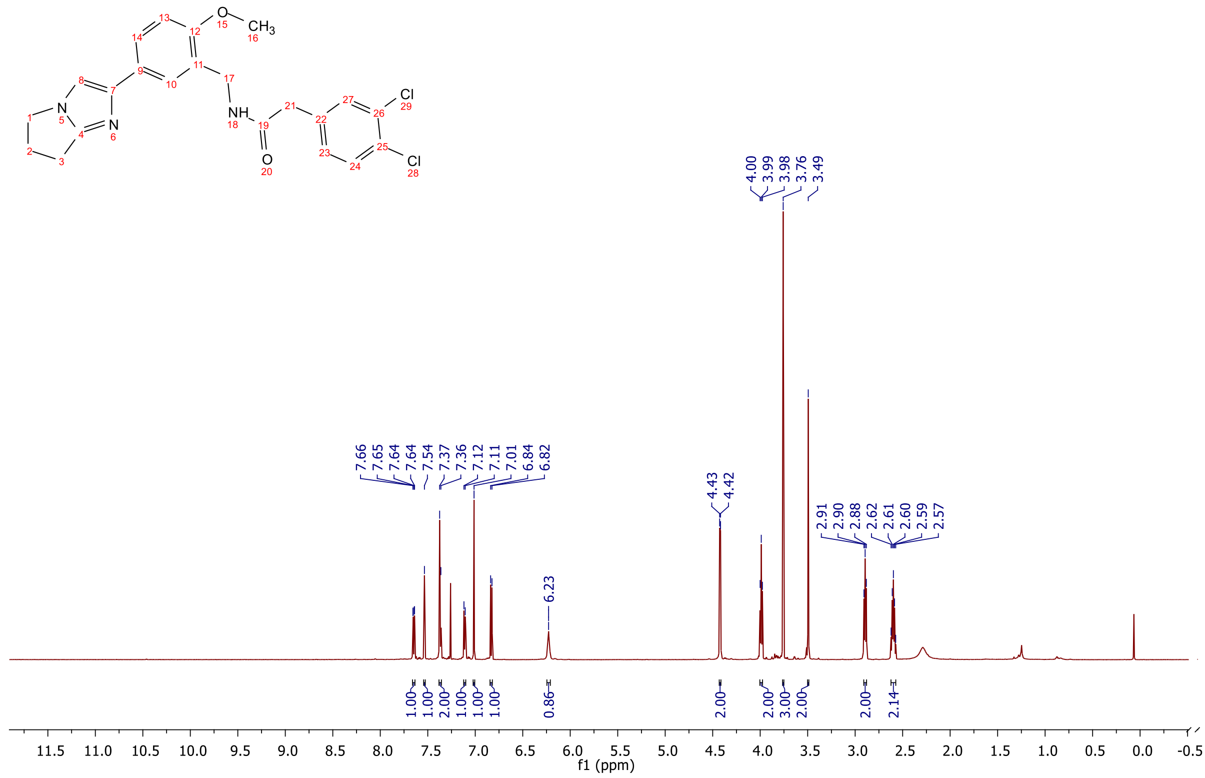

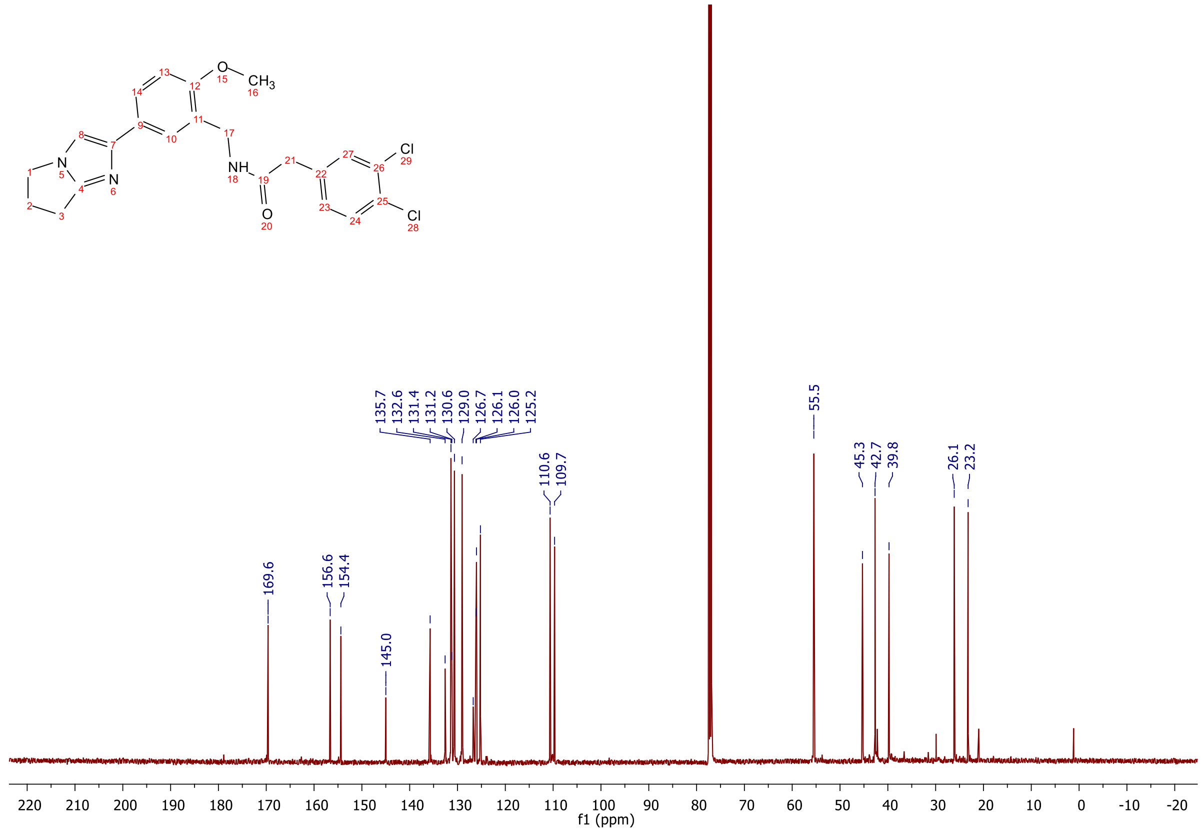

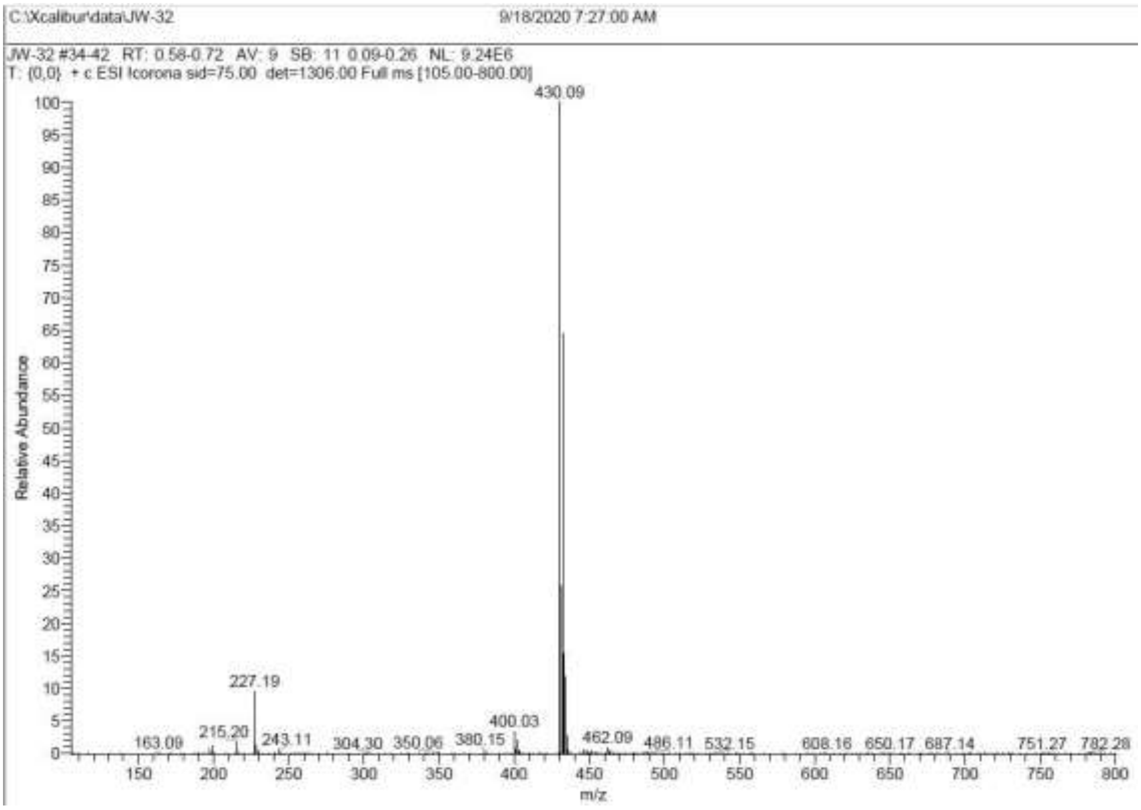
C:\Xcalibur\data\JW-32
9/18/2020 7:27:00 AM
JW-32 #34-42 RT: 0.58-0.72 AV: 9 SB: 11 0.09-0.26 NL: 9.24E6
T: {0,0} + c ESI !corona sid=75.00 det=1306.00 Full ms [105.00-800.00]
430.09
227.19
400.03
163.09
215.20
243.11
304.30
350.06
380.15
462.09
486.11
532.15
608.16
650.17
687.14
751.27
782.28
Relative Abundance
m/z

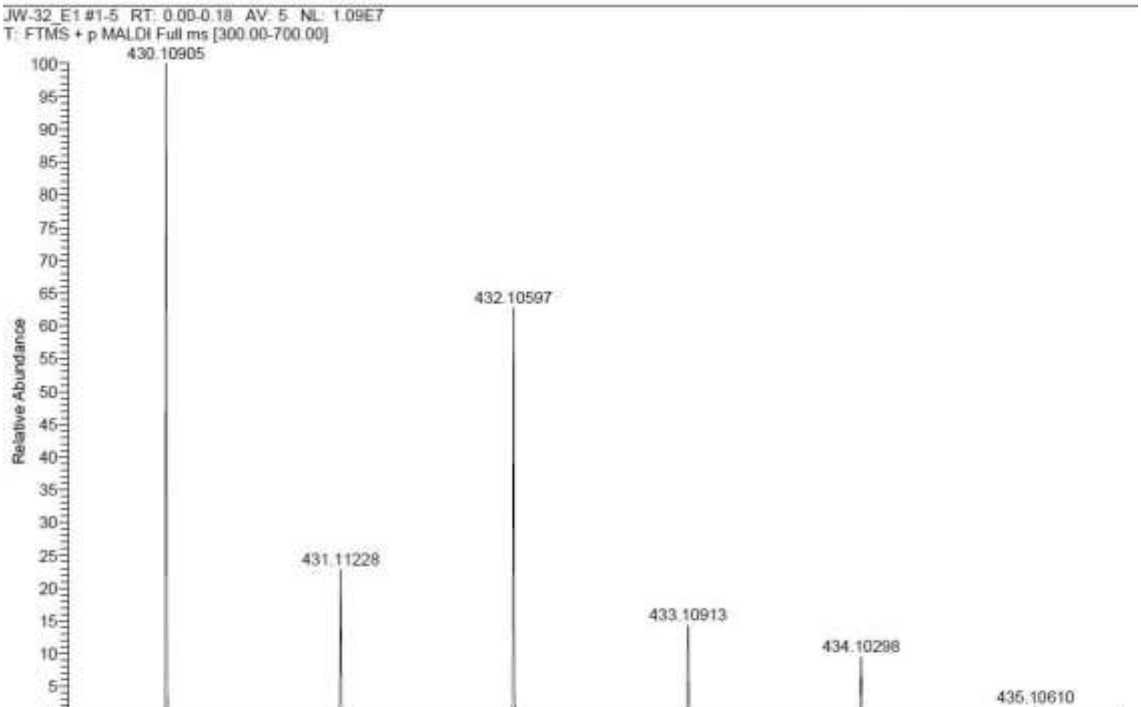
C:\User\...\Knapp\2020\xxxxxxx\JW-32_E1
9/18/2020 9:42:30 PM
JW-32 mit HCCA gemessen.
JW-32_E1 #1-5 RT: 0.00-0.18 AV: 5 NL: 1.09E7
T: FTMS + p MALDI Full ms [300.00-700.00]
430.10905
432.10597
431.11228
433.10913
434.10298
435.10610
Relative Abundance
m/z

**Signal:** MWD1 A, Sig=254,4 Ref=off

| RT [min] | Type | Width [min] | Area | Height | Area% |
|---|---|---|---|---|---|
| 12.360 | VV | 0.2502 | 4107.7275 | 238.1471 | 100.0000 |
| | | Sum | 4107.7275 | | |

**Signal:** MWD1 B, Sig=280,4 Ref=off

| RT [min] | Type | Width [min] | Area | Height | Area% |
|---|---|---|---|---|---|
| 12.360 | VV | 0.2477 | 4168.8242 | 244.8035 | 100.0000 |
| | | Sum | 4168.8242 | | |

**Signal:** MWD1 F, Sig=260,4 Ref=off

| RT [min] | Type | Width [min] | Area | Height | Area% |
|---|---|---|---|---|---|
| 12.360 | VV | 0.2483 | 4973.1641 | 291.0685 | 100.0000 |
| | | Sum | 4973.1641 | | |

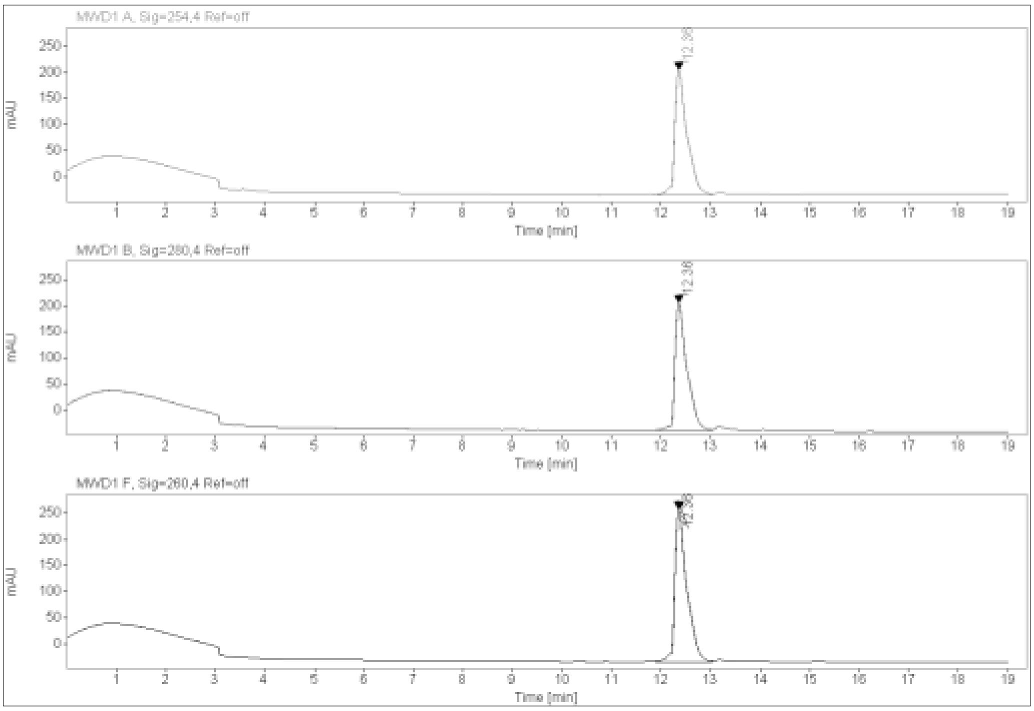

**ESI, $^{1}$H-NMR and $^{13}$C-NMR of 2-(3,4-dichlorophenyl)-N-(5-(6,7-dihydro-5H-pyrrolo[1,2-a]imidazol-2-yl)-2-hydroxybenzyl) acetamide 15**

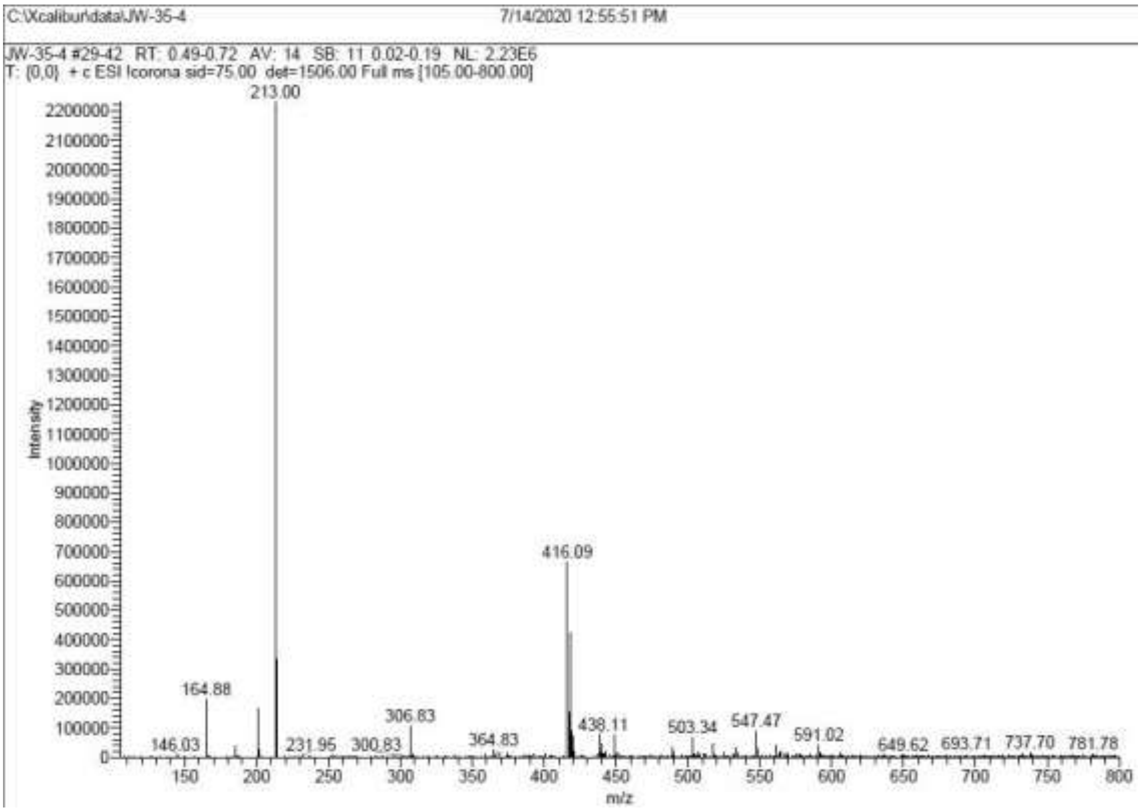

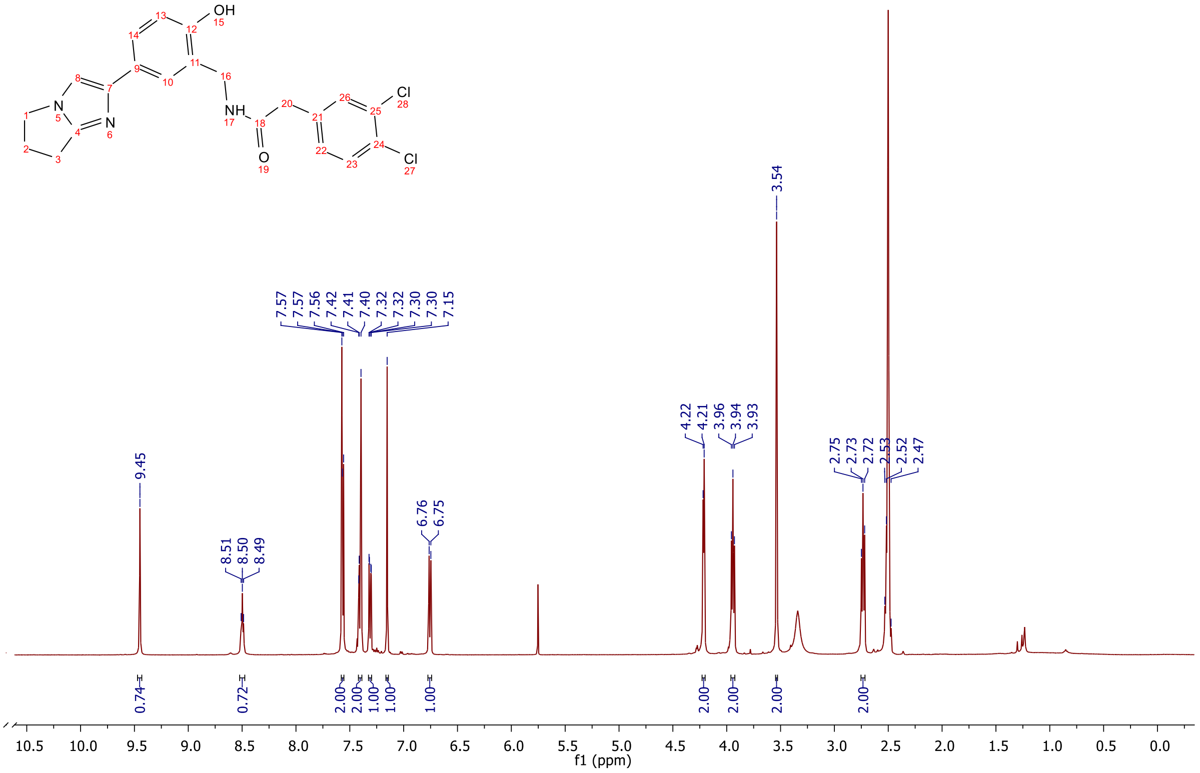

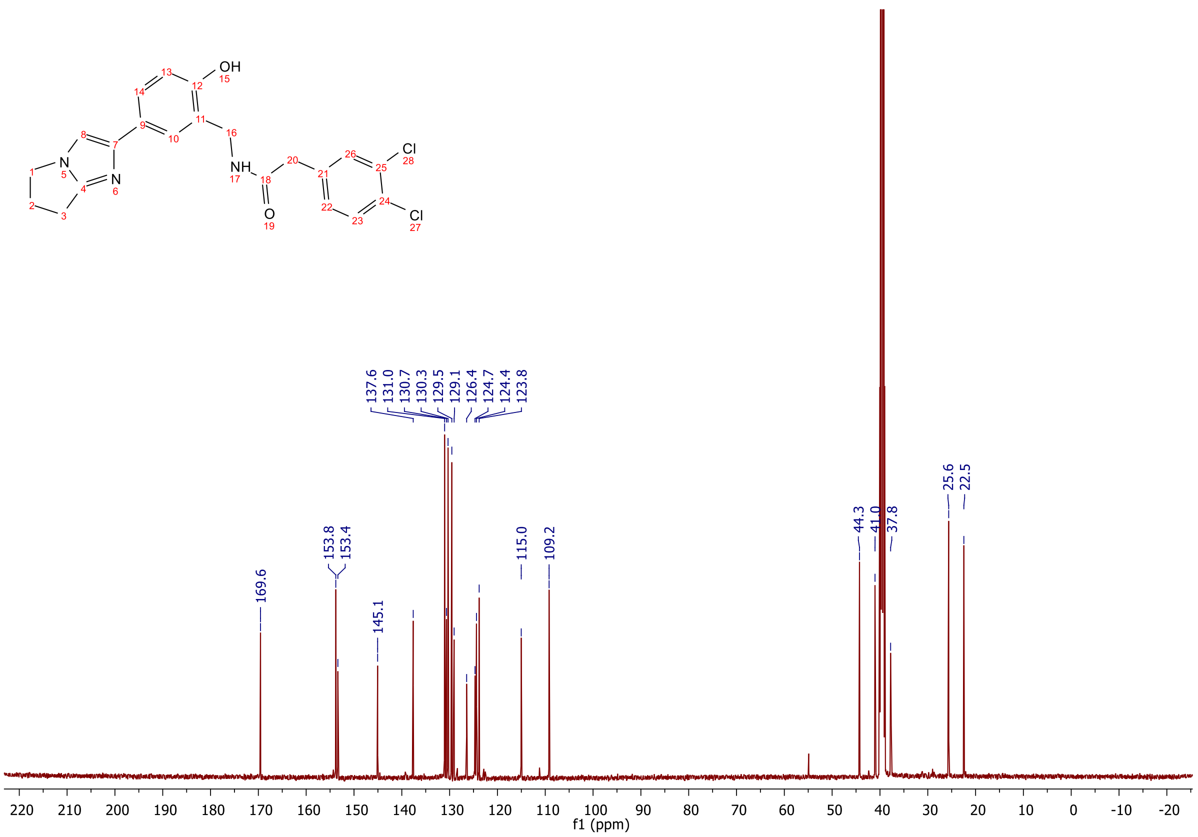
OH
NH
N
N
O
Cl
Cl
137.6
131.0
130.7
130.3
129.5
129.1
126.4
124.7
124.4
123.8
169.6
153.8
153.4
145.1
115.0
109.2
44.3
41.0
37.8
25.6
22.5
220 210 200 190 180 170 160 150 140 130 120 110 100 90 80 70 60 50 40 30 20 10 0 -10 -20
f1 (ppm)

**[1]H-NMR, [13]C-NMR and MALDI of tert-butyl 3-(2-(2-((2-(3,4-dichlorophenyl)acetamido)methyl)-4-(6,7-dihydro-5H-pyrrolo[1,2-a]imidazol-2-yl)phenoxy)ethoxy)propanoate 16a**

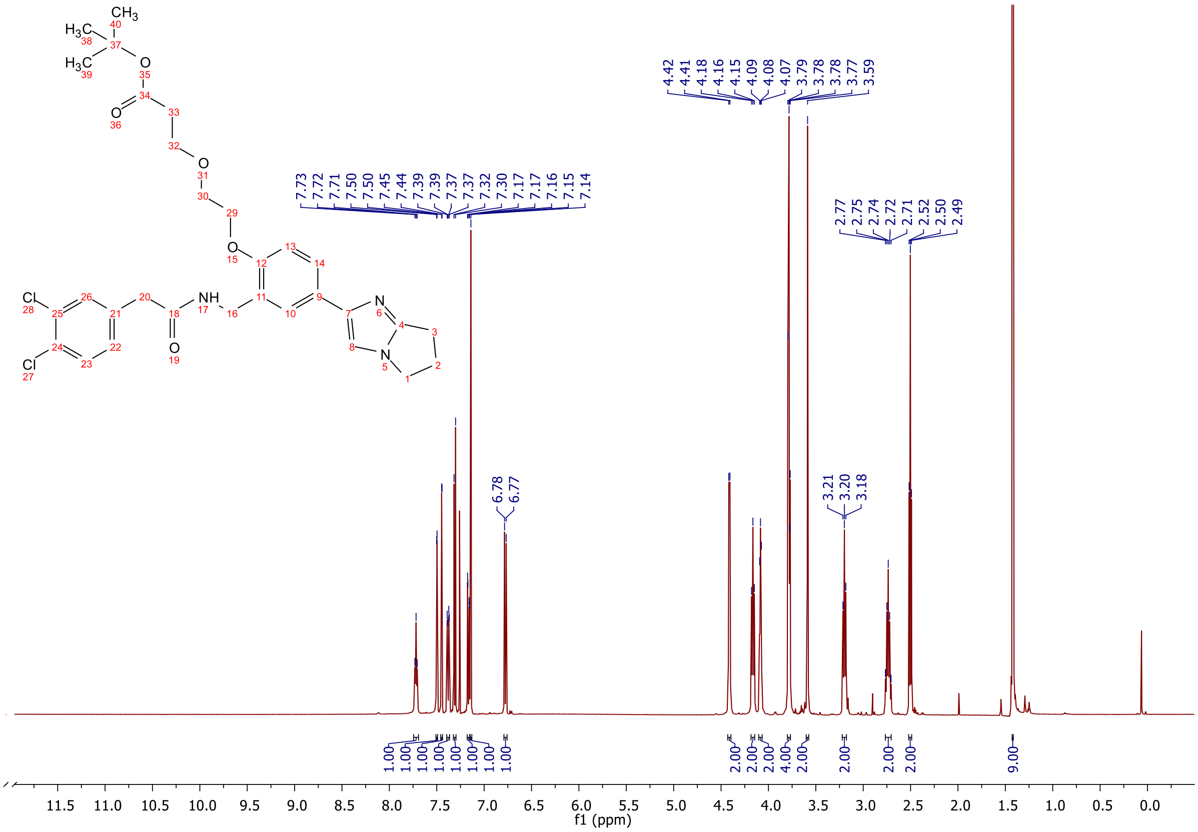

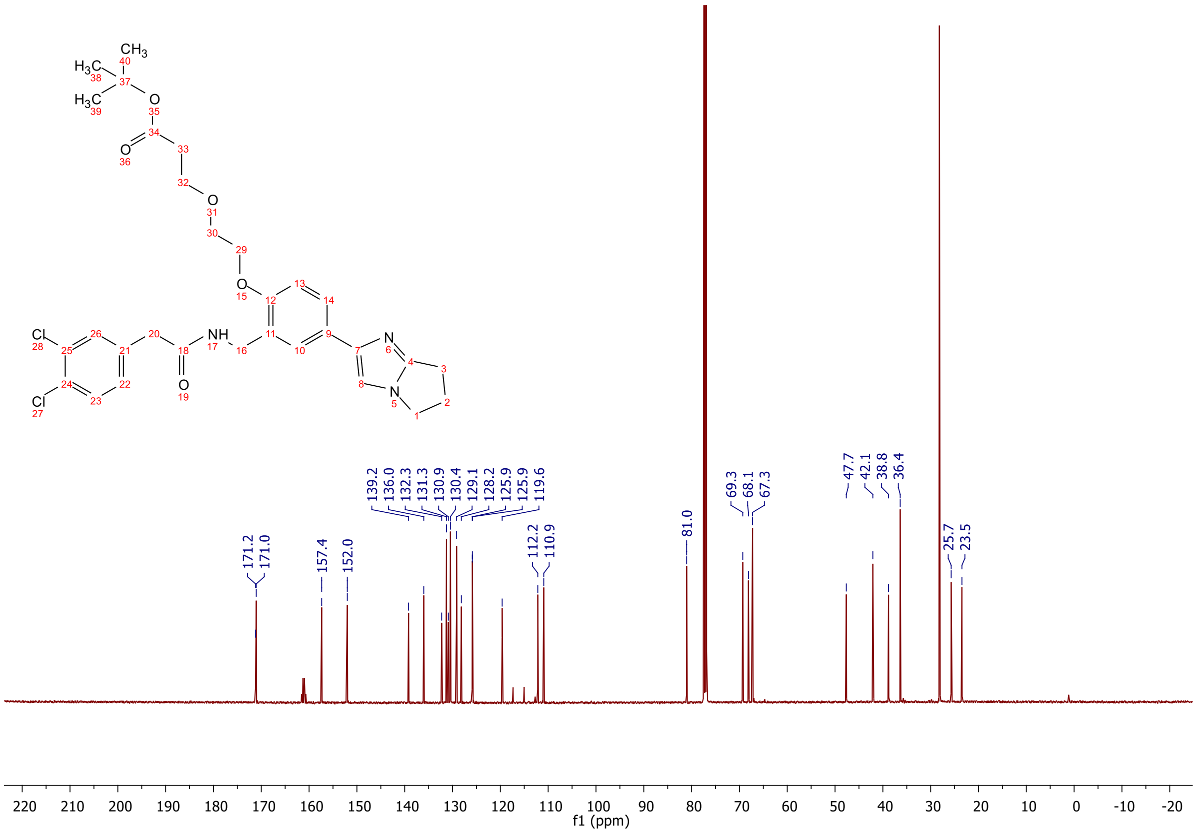

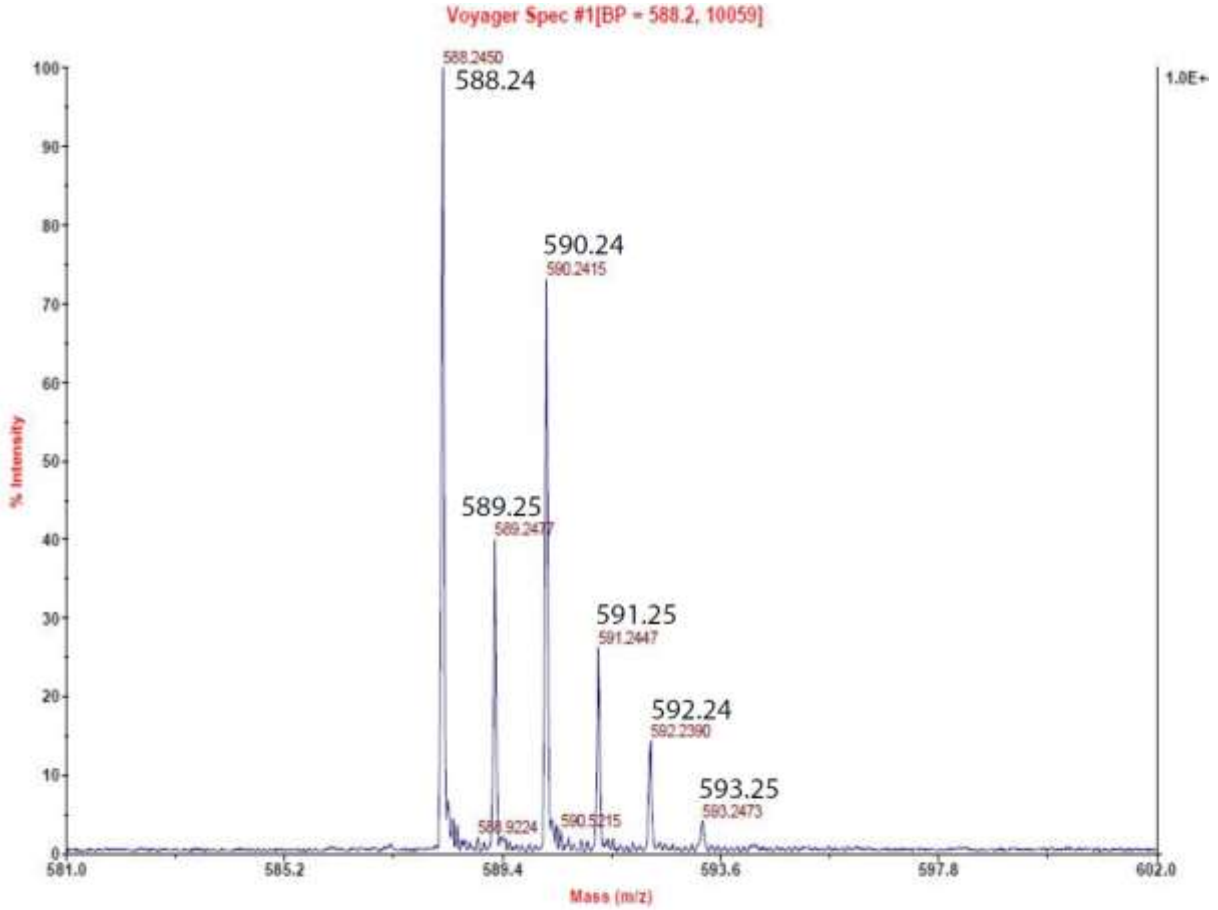
Voyager Spec #1[BP = 588.2, 10059]
588.2450
588.24
590.24
590.2415
589.25
589.2477
591.25
591.2447
592.24
592.2390
593.25
593.2473
588.9224
590.5215
% Intensity
Mass (m/z)
1.0E+

**$^{1}$H-NMR, $^{13}$C-NMR and MALDI of tert-butyl 3-(2-(2-(2-((2-(3,4-dichlorophenyl)acetamido)methyl)-4-(6,7-dihydro-5H-pyrrolo[1,2-a]imidazol-2-yl)phenoxy)ethoxy)ethoxy)propanoate 16b**

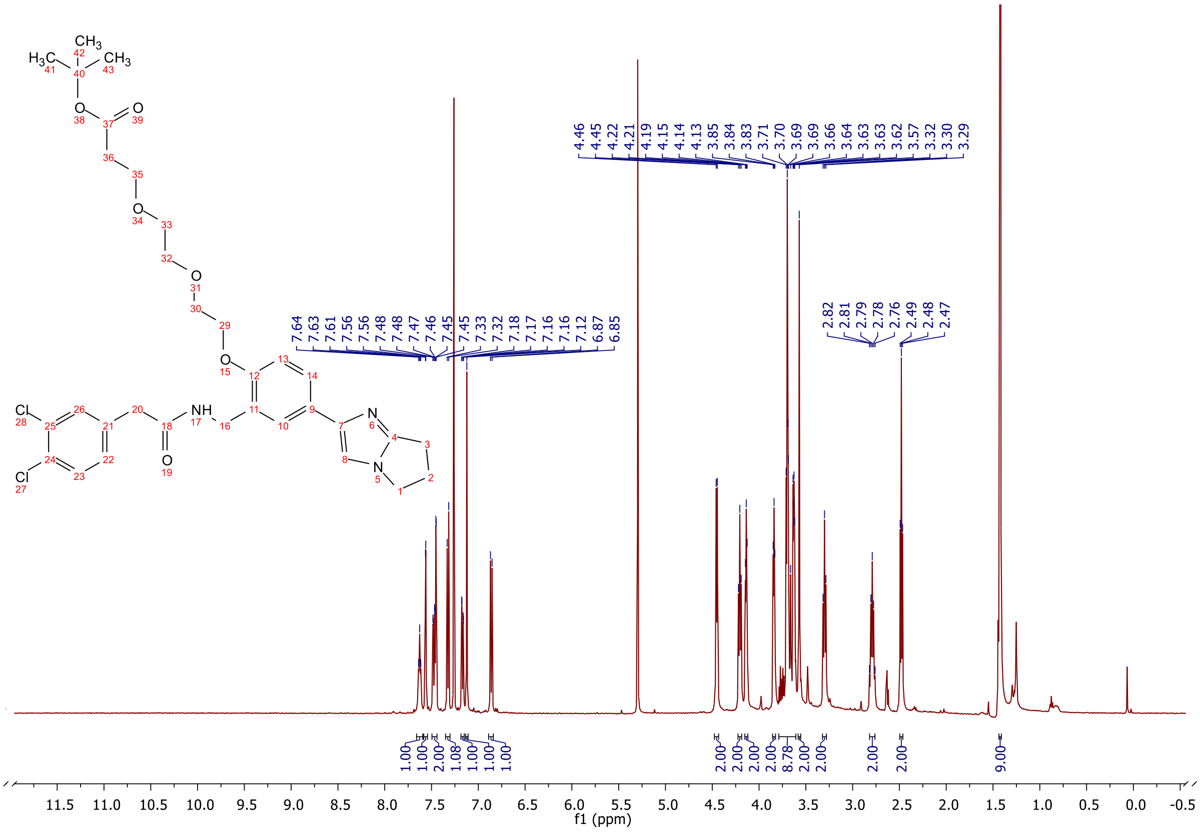

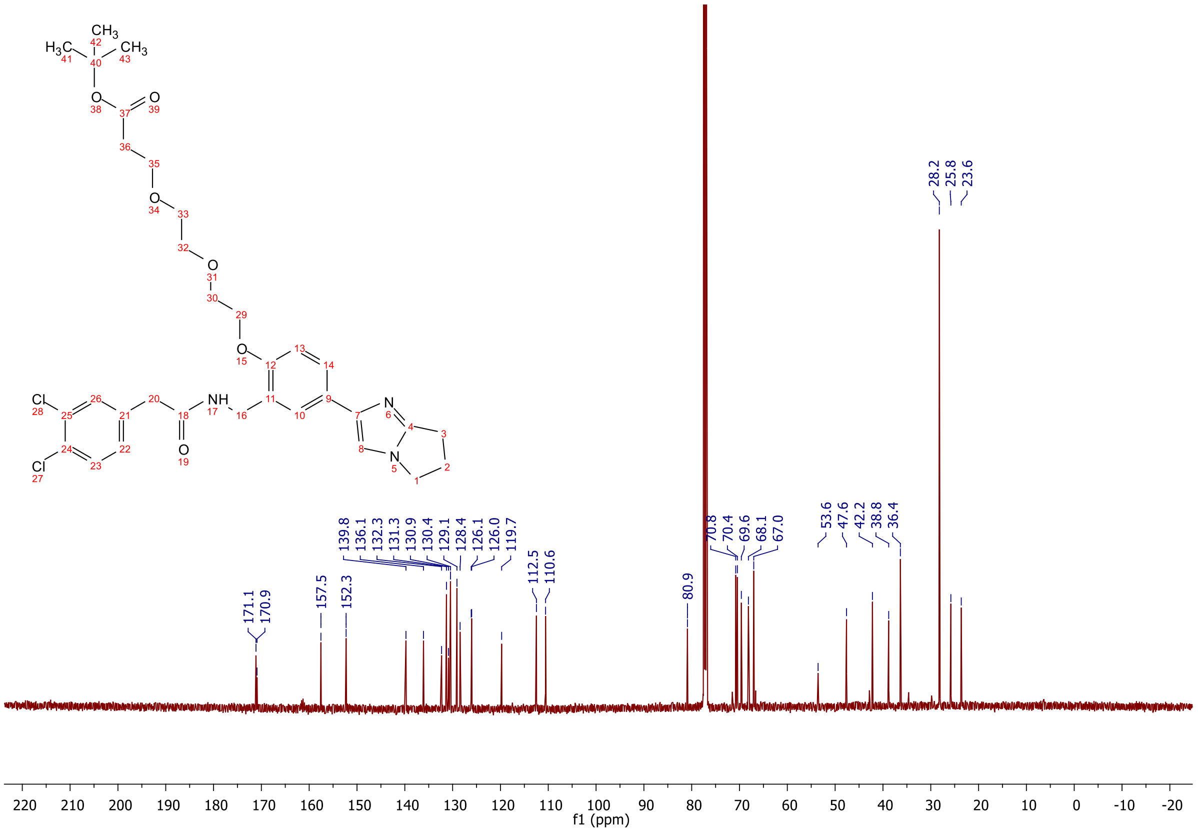

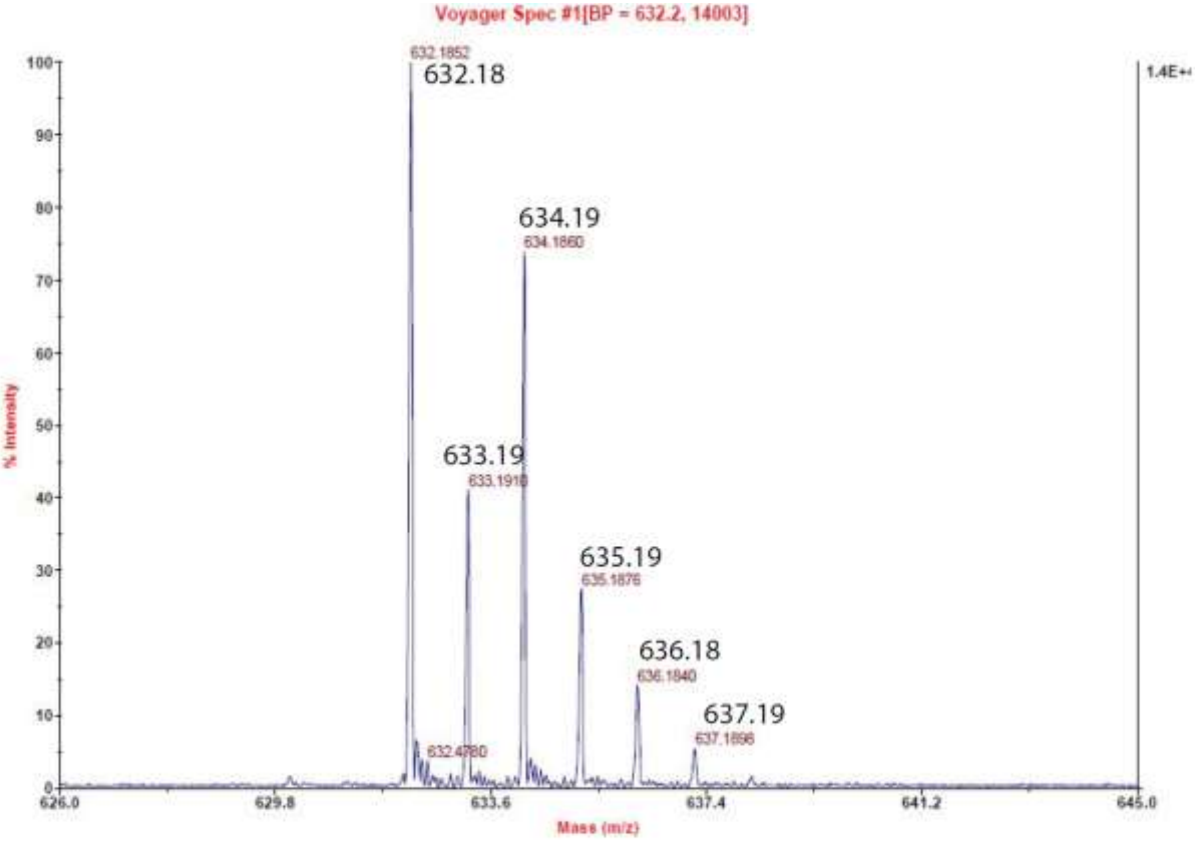
Voyager Spec #1[BP = 632.2, 14003]
632.1852
632.18
634.19
634.1860
633.19
633.1910
635.19
635.1876
636.18
636.1840
637.19
637.1898
632.4780
1.4E+4
% Intensity
Mass (m/z)
626.0
629.8
633.6
637.4
641.2
645.0

**$^{1}$H-NMR, $^{13}$C-NMR and MALDI of tert-butyl 3-(2-(2-(2-(2-((2-(3,4-dichlorophenyl)acetamido)methyl)-4-(6,7-dihydro-5H-pyrrolo[1,2-a]imidazol-2-yl)phenoxy)ethoxy)ethoxy)ethoxy)propanoate 16c**

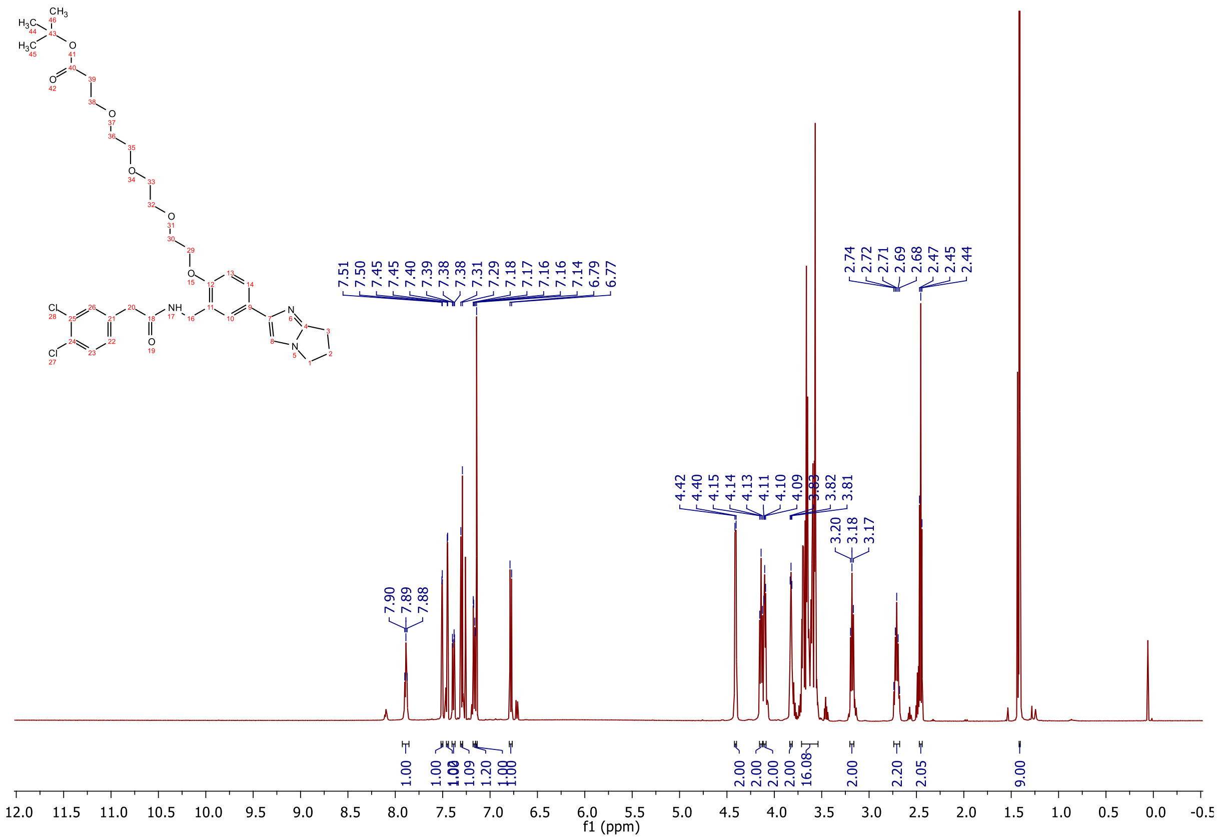

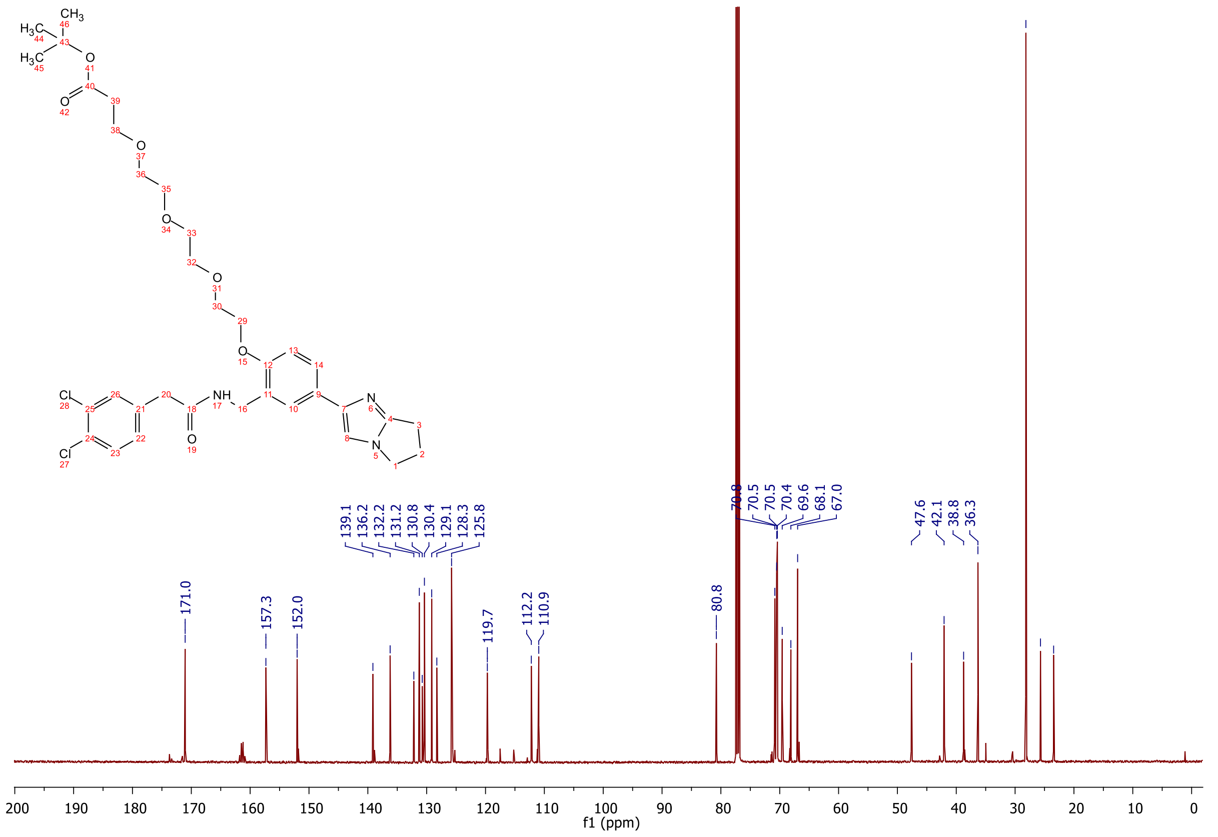

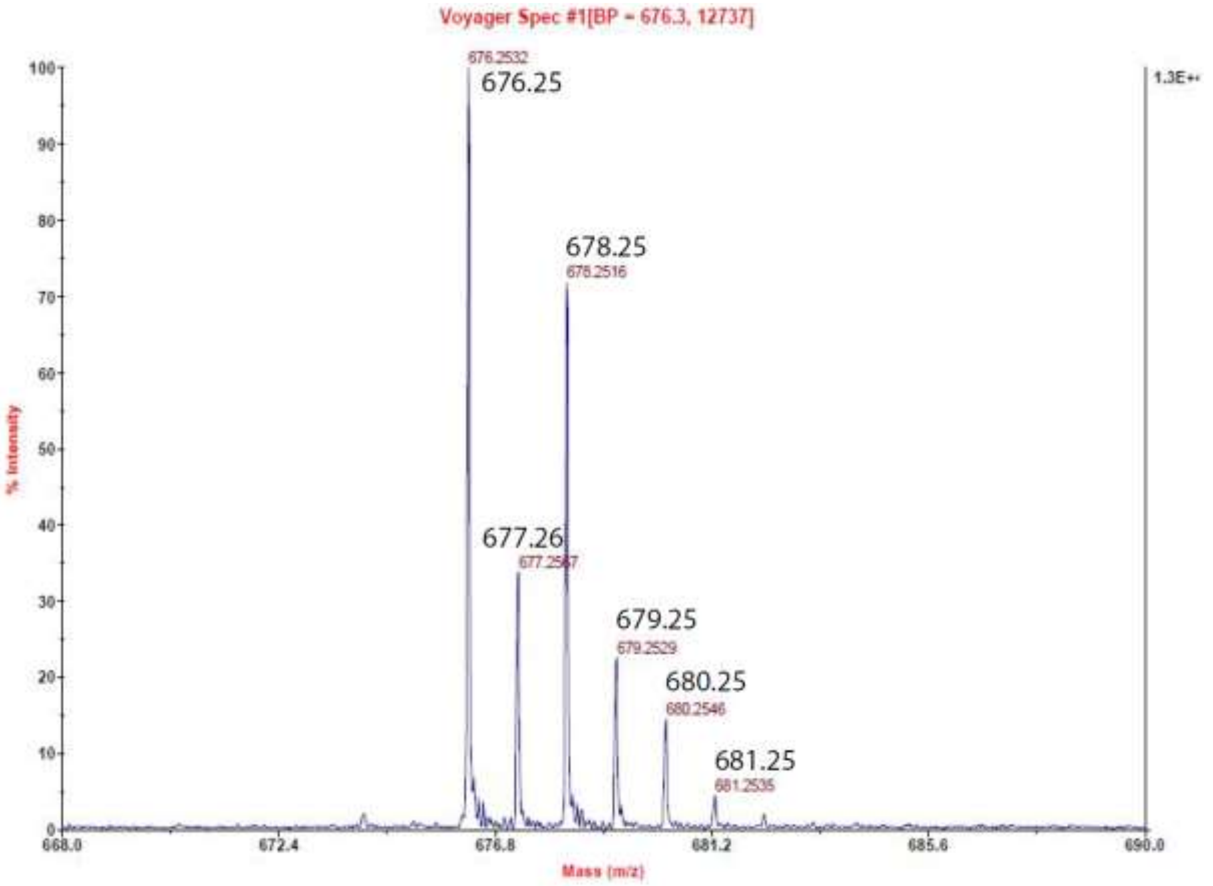

Voyager Spec #1[BP = 676.3, 12737]
676.2532
676.25
678.25
678.2516
677.26
677.2567
679.25
679.2529
680.25
680.2546
681.25
681.2535
1.3E+4
% Intensity
Mass (m/z)

**$^{1}$H-NMR, $^{13}$C-NMR and MALDI of tert-butyl 1-(2-((2-(3,4-dichlorophenyl)acetamido)methyl)-4-(6,7-dihydro-5H-pyrrolo[1,2-a]imidazol-2-yl)phenoxy)-3,6,9,12-tetraoxapentadecan-15-oate 16d**

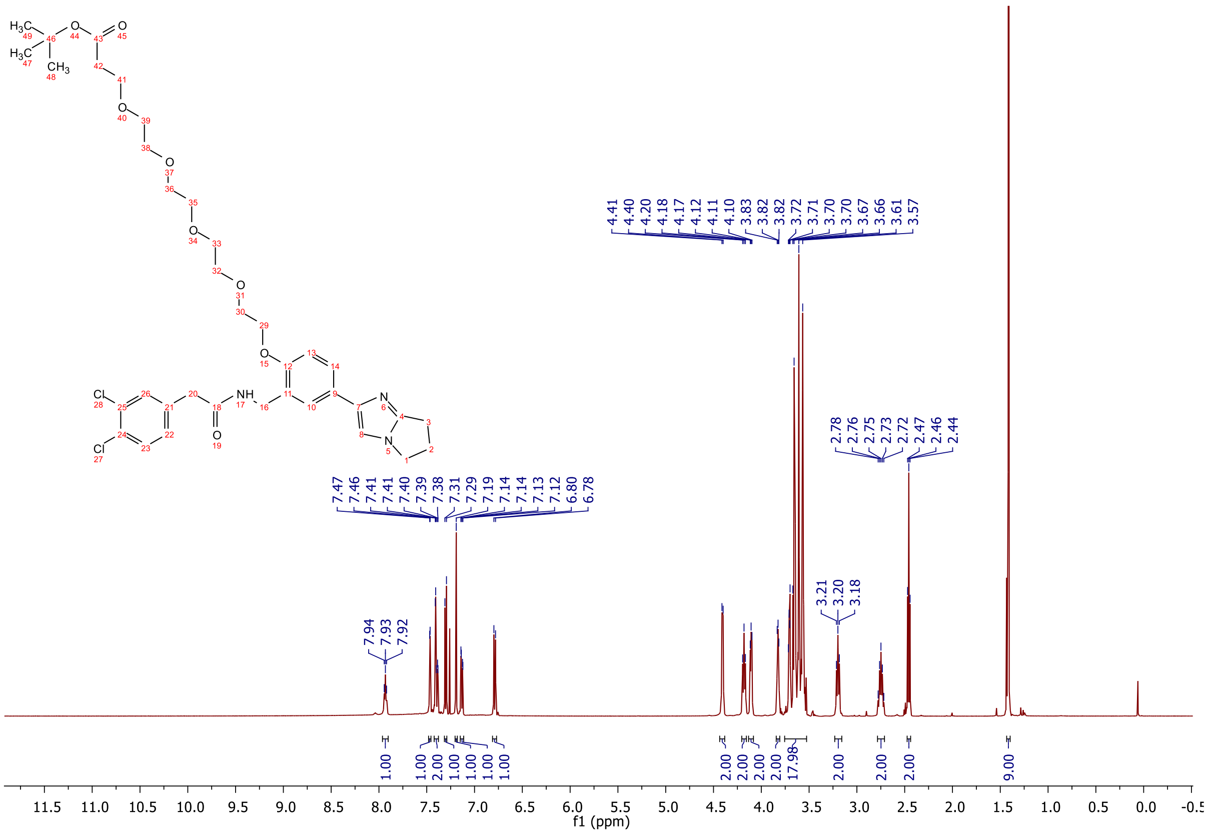

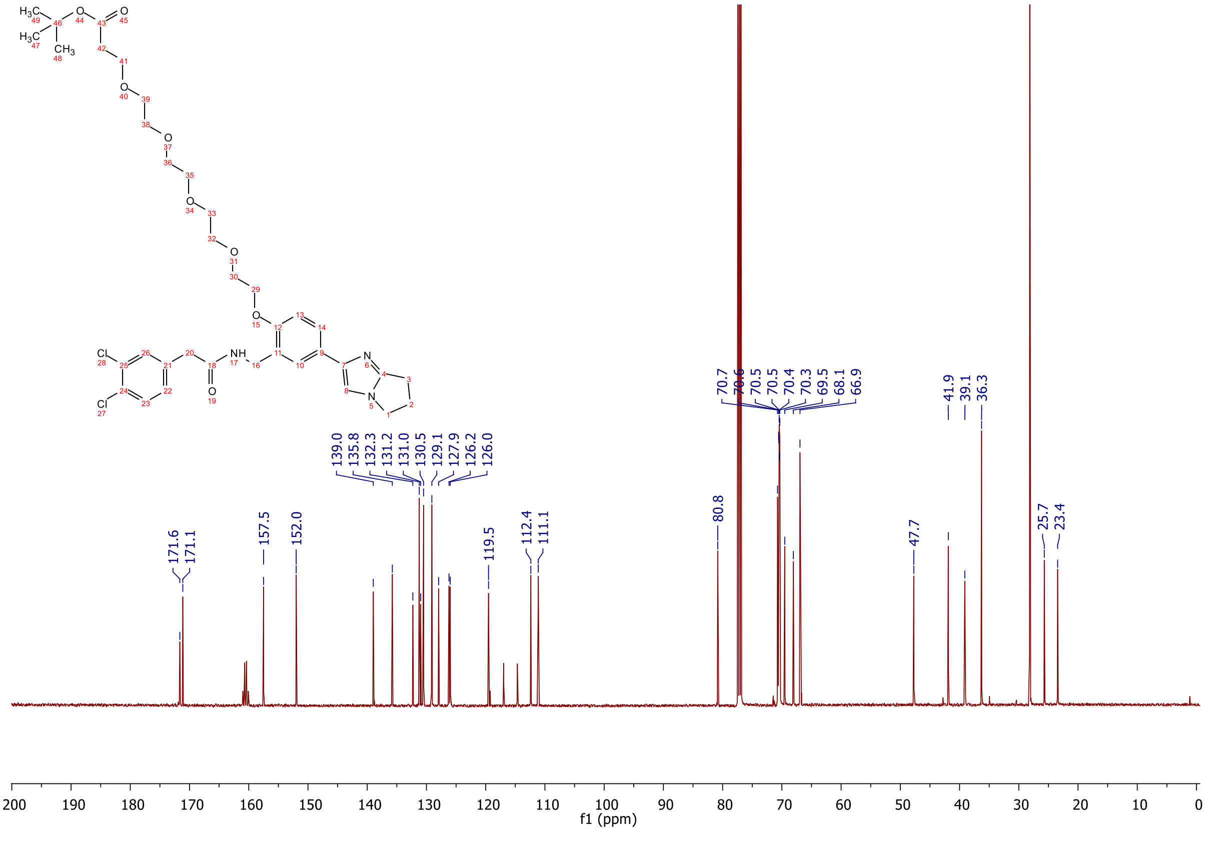

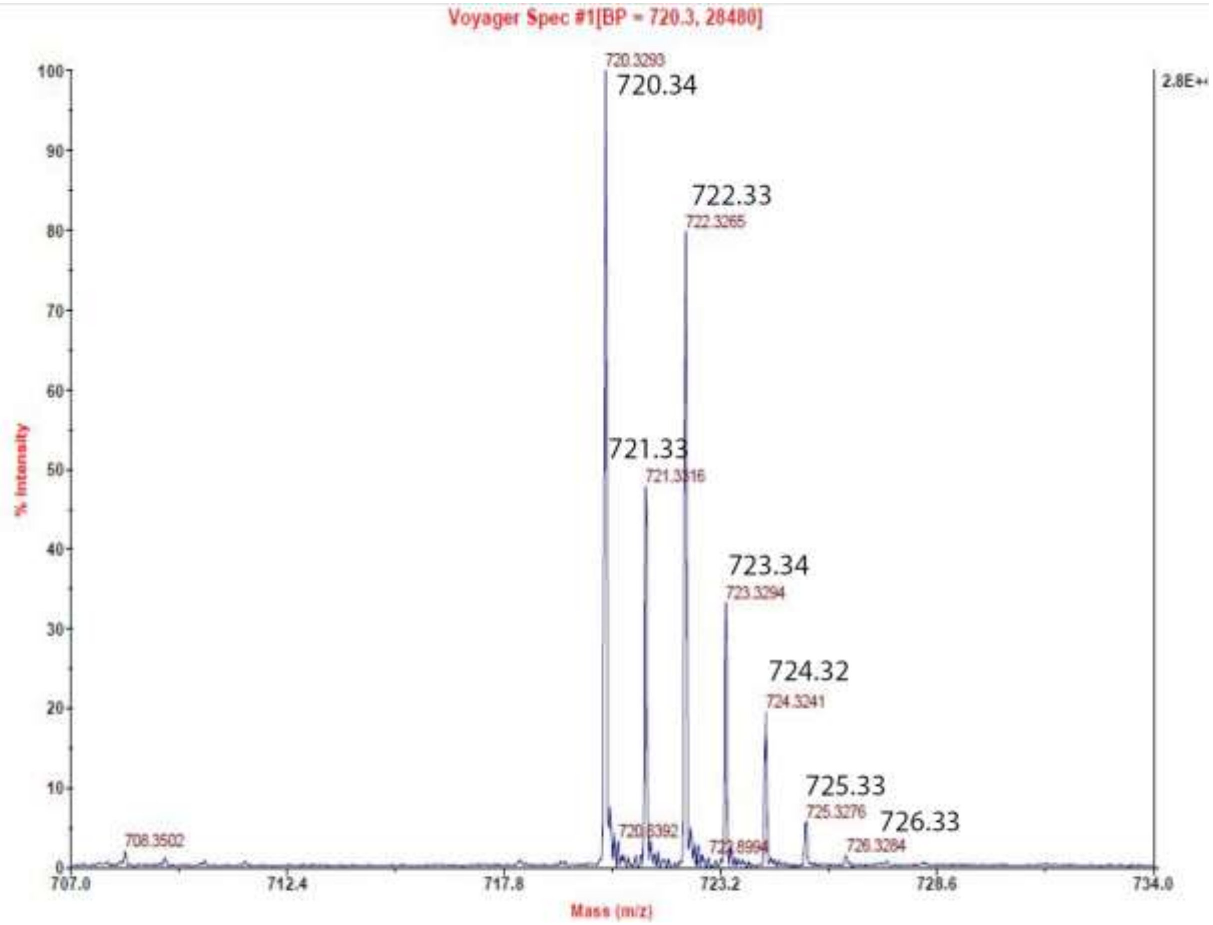

Voyager Spec #1[BP = 720.3, 28480]
720.3293
720.34
722.33
722.3265
721.33
721.3316
723.34
723.3294
724.32
724.3241
725.33
725.3276
726.33
726.3284
708.3502
% Intensity
Mass (m/z)
2.8E+

**[1]H-NMR, [13]C-NMR and MALDI of tert-butyl 1-(2-((2-(3,4-dichlorophenyl)acetamido)methyl)-4-(6,7-dihydro-5H-pyrrolo[1,2-a]imidazol-2-yl)phenoxy)-3,6,9,12,15-pentaoxaoctadecan-18-oate 16e**

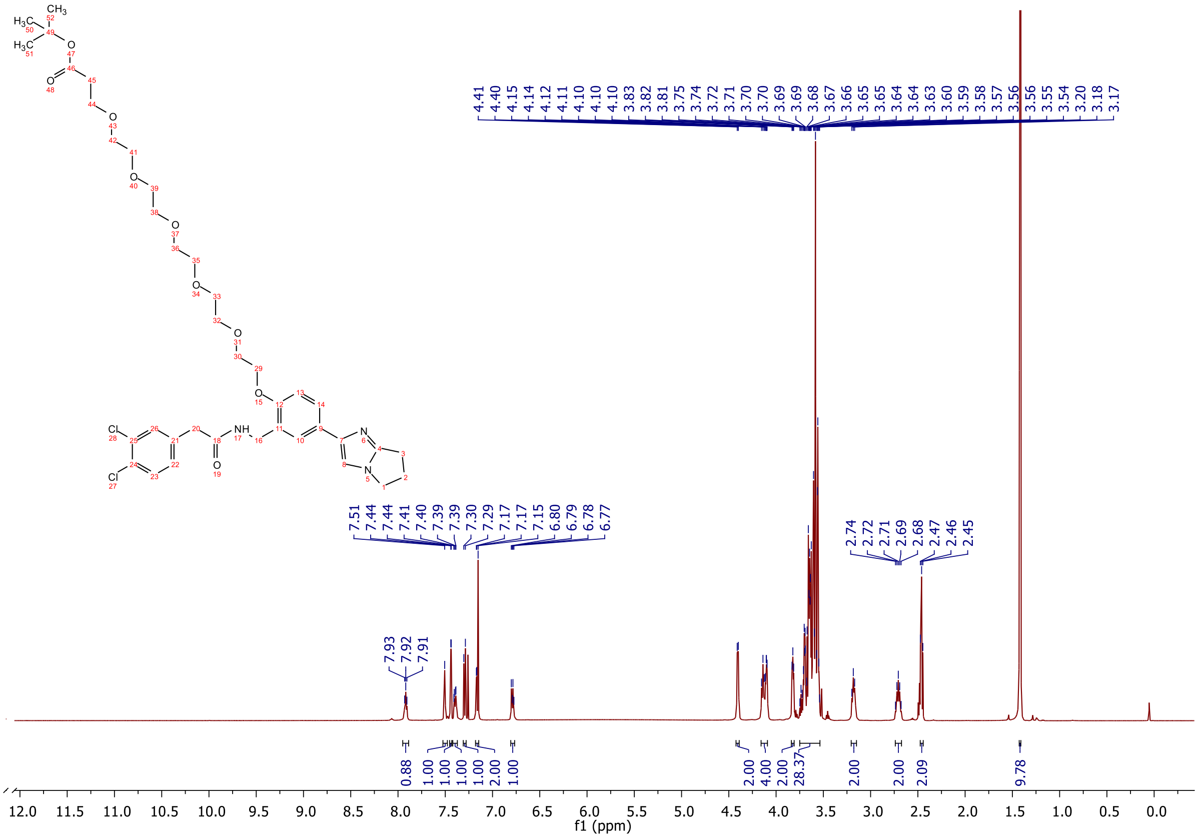

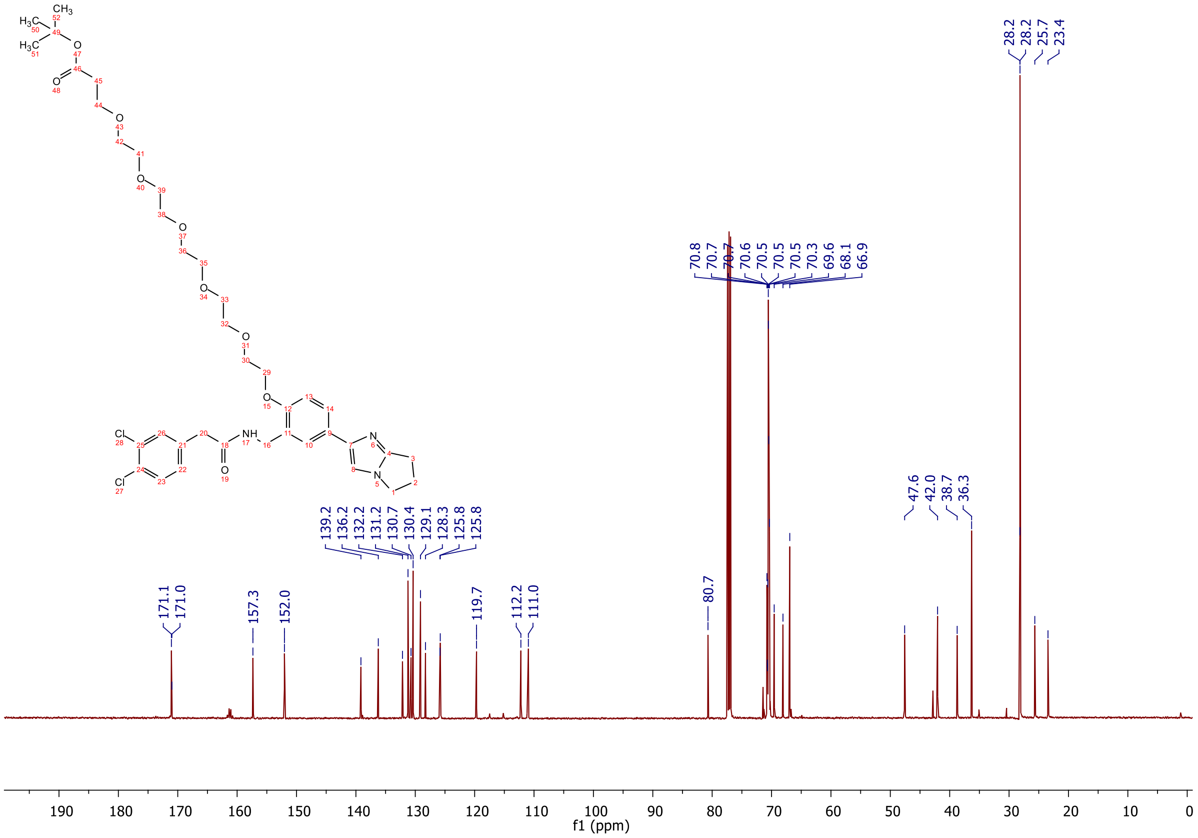

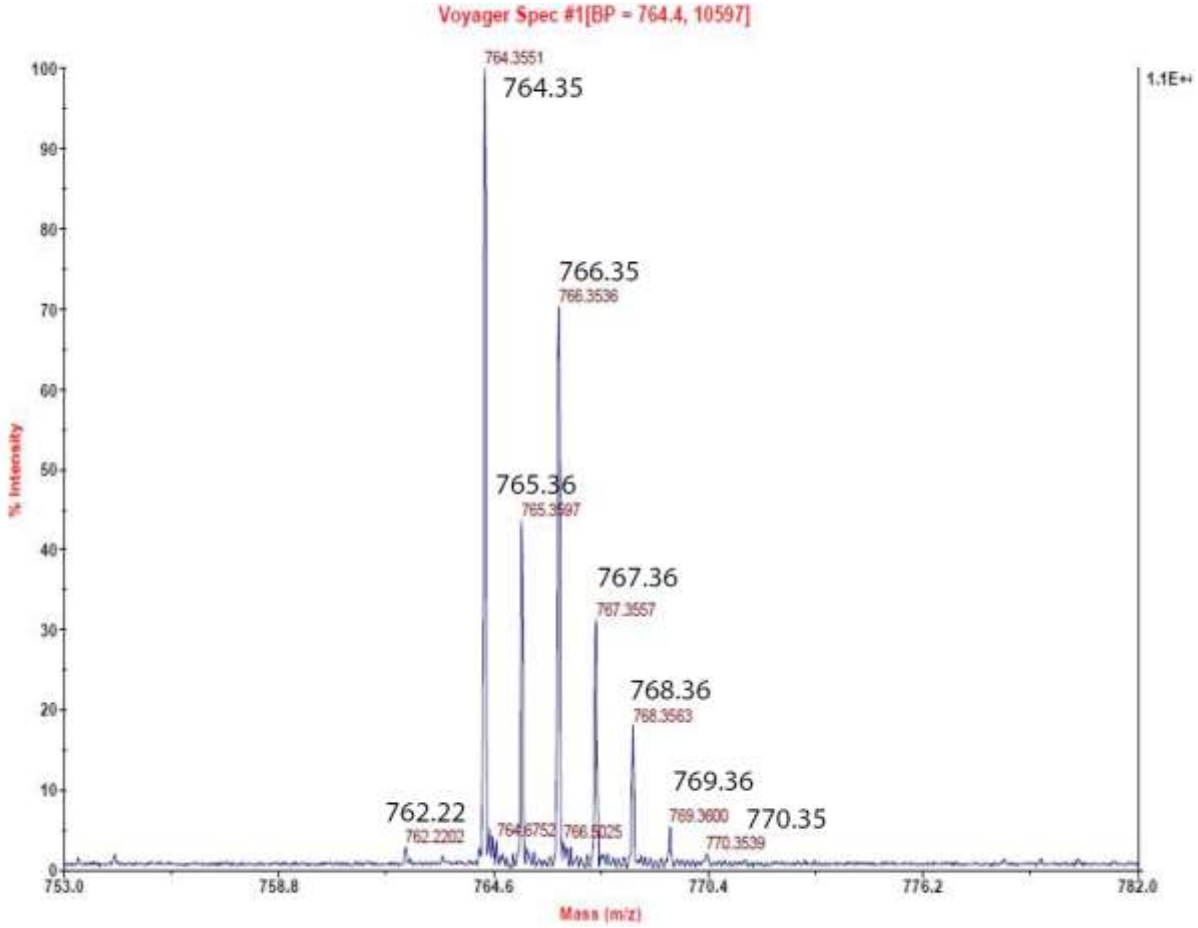
Voyager Spec #1[BP = 764.4, 10597]
% Intensity
Mass (m/z)
764.3551
764.35
766.35
766.3536
765.36
765.3597
767.36
767.3557
768.36
768.3563
769.36
769.3600
770.35
770.3539
762.22
762.2202
764.6752
766.5025
1.1E+4
753.0
758.8
764.6
770.4
776.2
782.0

**[1]H-NMR, [13]C-NMR and MALDI of tert-butyl 1-(2-((2-(3,4-dichlorophenyl)acetamido)methyl)-4-(6,7-dihydro-5H-pyrrolo[1,2-a]imidazol-2-yl)phenoxy)-3,6,9,12,15,18-hexaoxahenicosan-21-oate 16f**

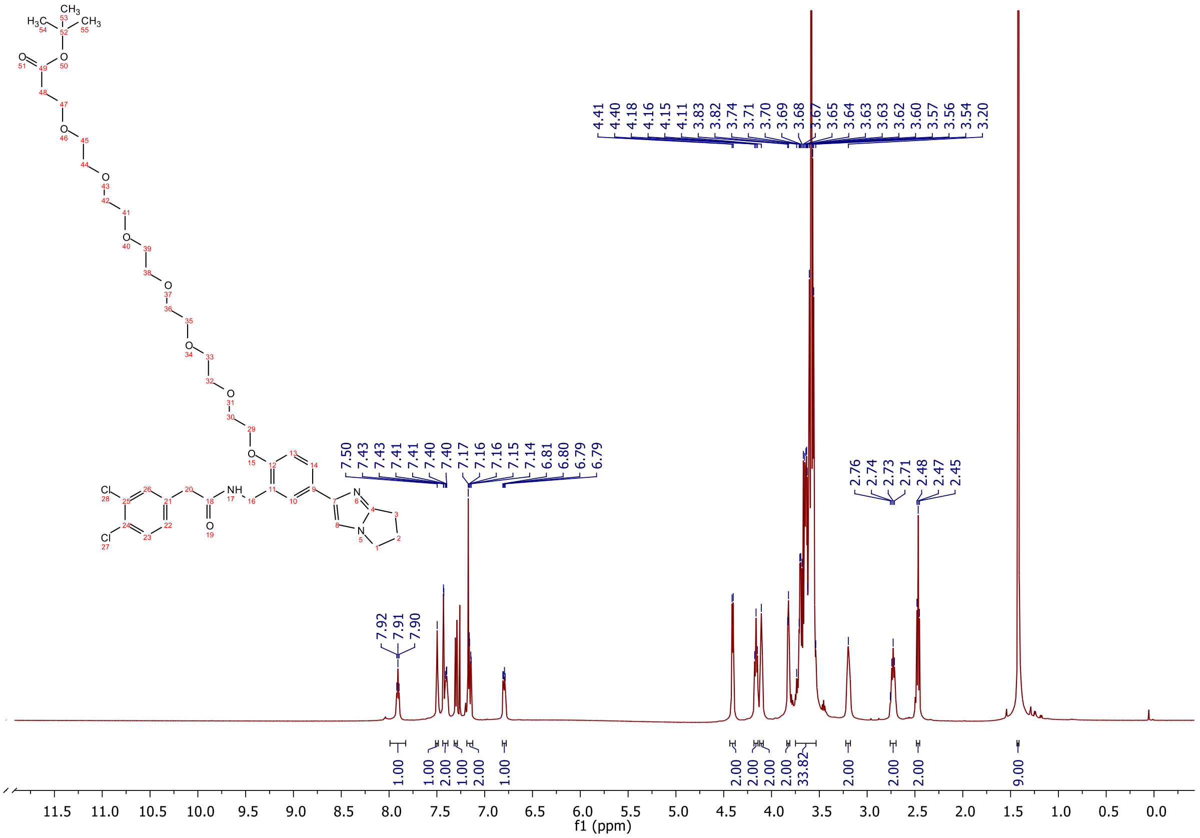

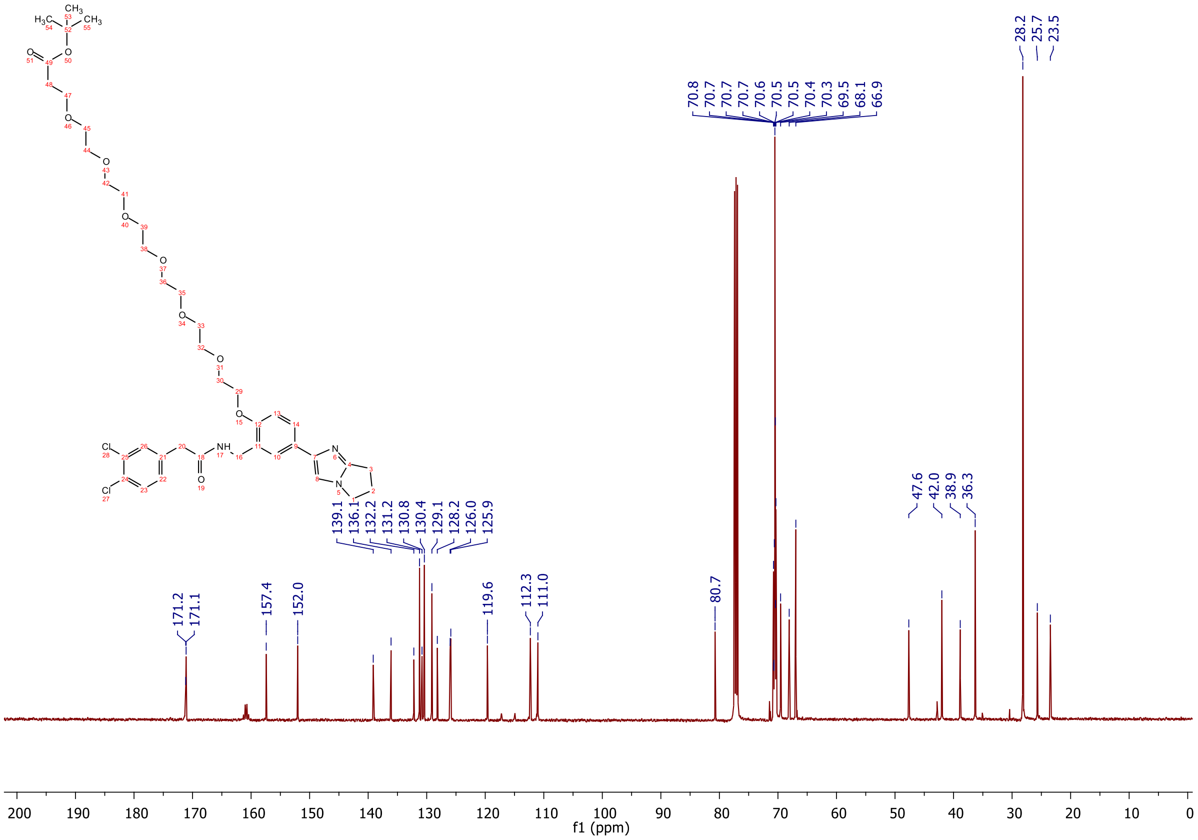

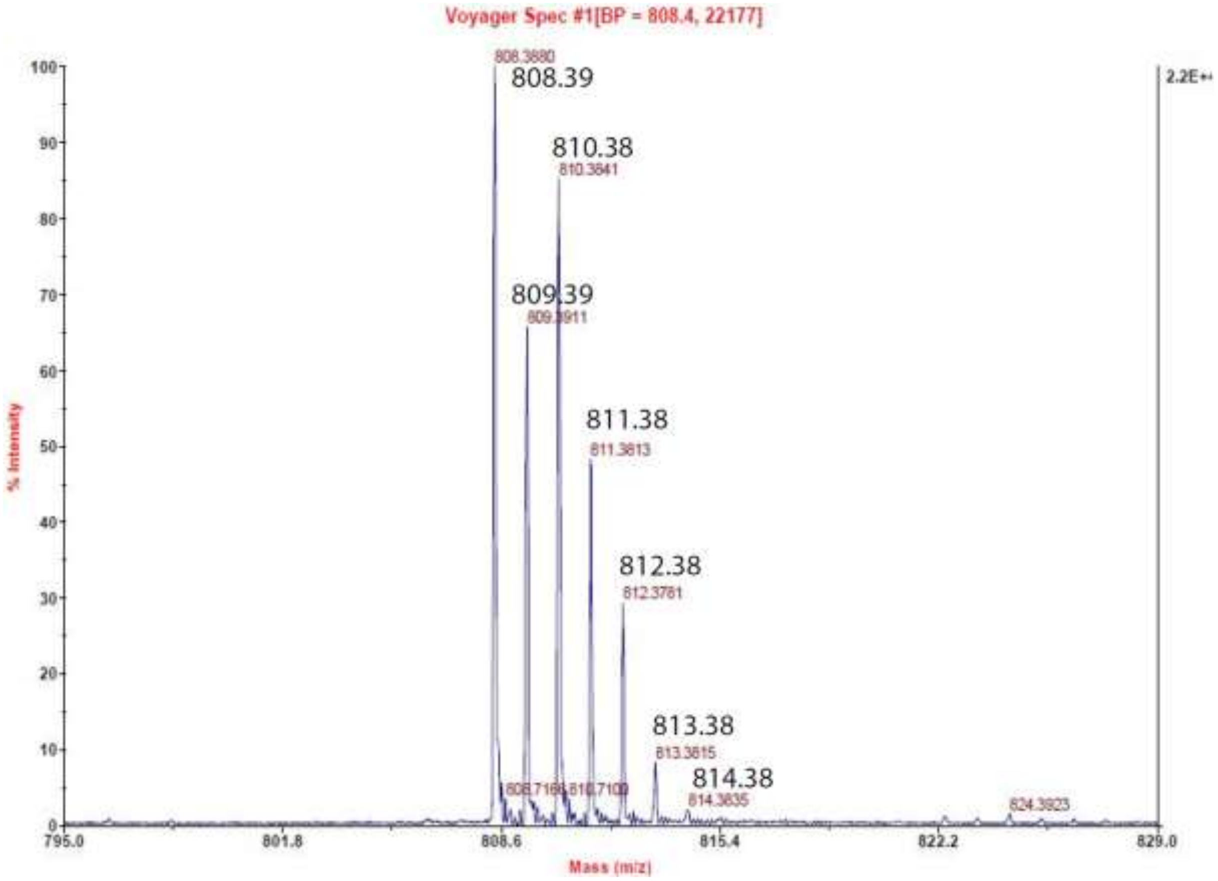

Voyager Spec #1[BP = 808.4, 22177]
808.3880
808.39
810.38
810.3841
809.39
809.3911
811.38
811.3813
812.38
812.3781
813.38
813.3815
814.38
814.3835
824.3923
2.2E+4
% Intensity
Mass (m/z)
795.0
801.8
808.6
815.4
822.2
829.0

**[1]H-NMR, [13]C-NMR and MALDI of tert-butyl 1-(2-((2-(3,4-dichlorophenyl)acetamido)methyl)-4-(6,7-dihydro-5H-pyrrolo[1,2-a]imidazol-2-yl)phenoxy)-3,6,9,12,15,18,21-heptaoxatetracosan-24-oate 16g**

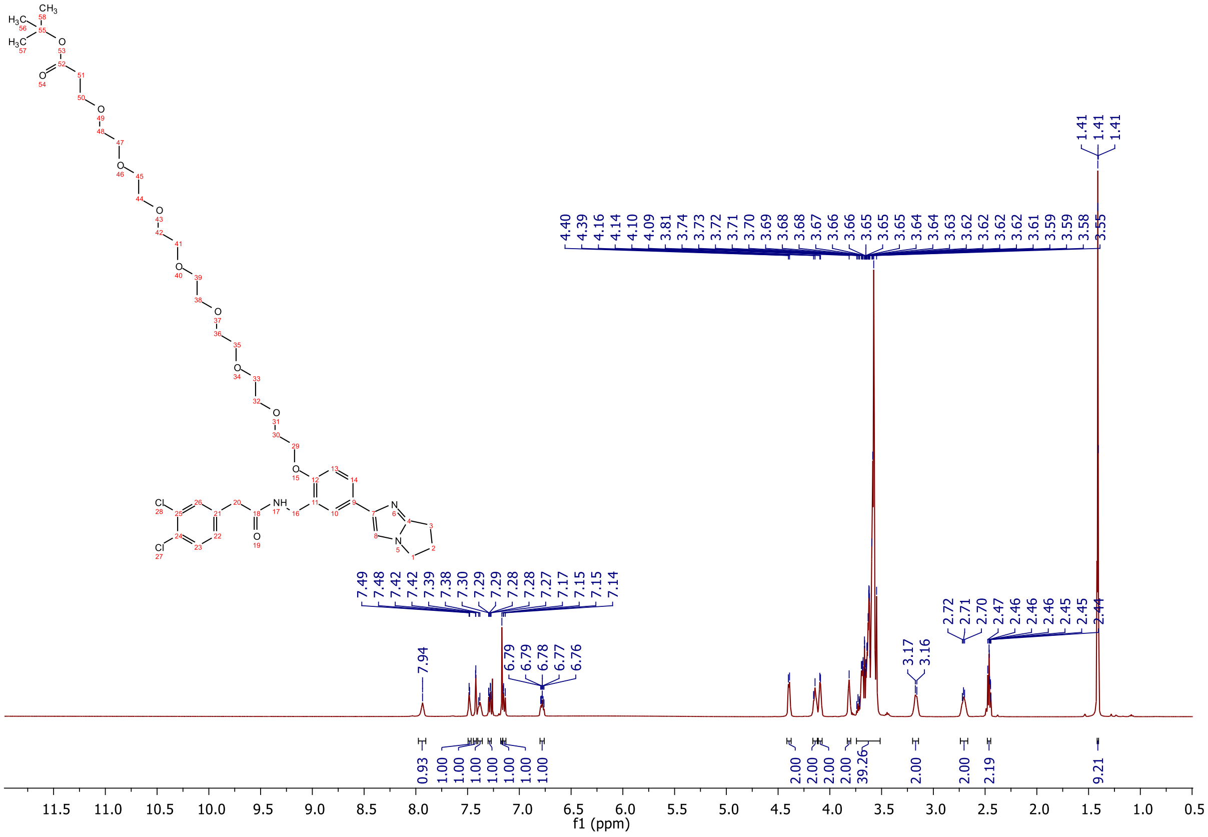

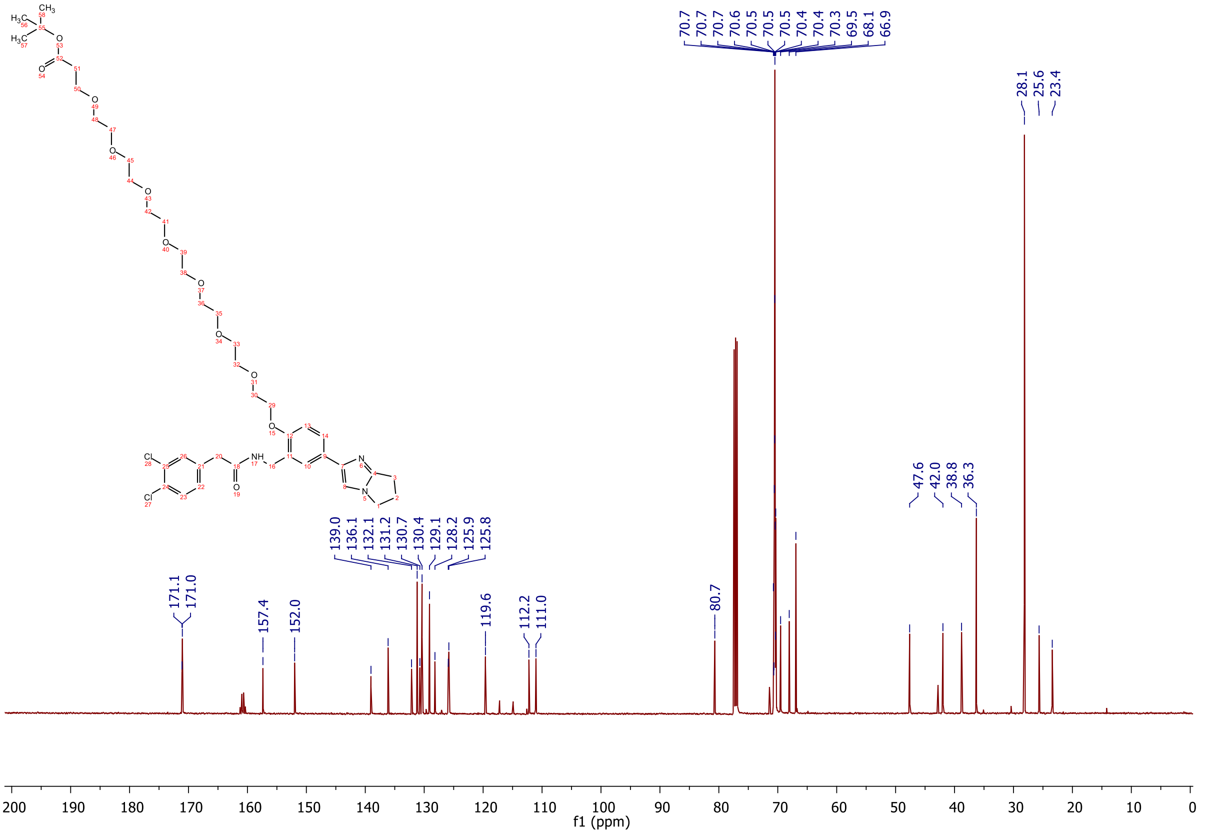

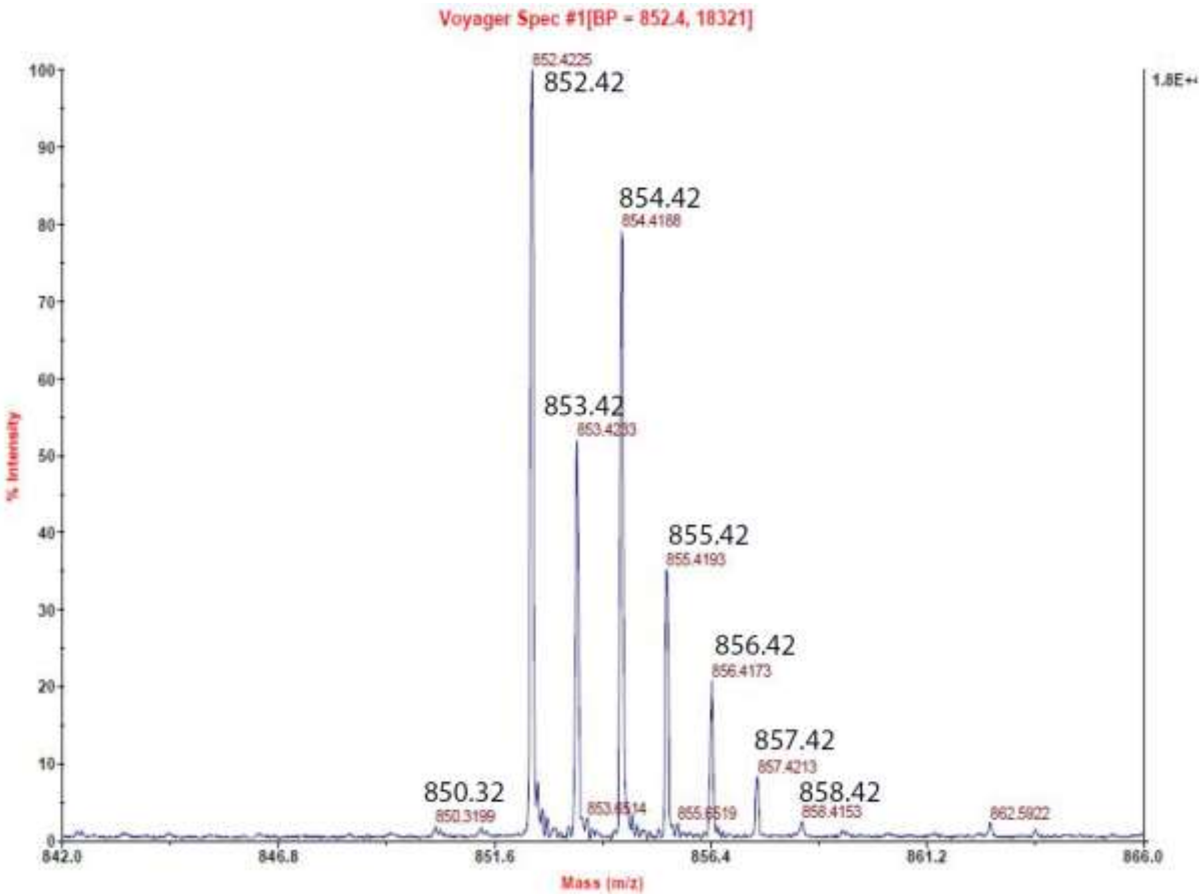
Voyager Spec #1[BP = 852.4, 18321]
852.4225
852.42
854.42
854.4188
853.42
855.42
855.4193
856.42
856.4173
857.42
857.4213
850.32
850.3199
858.42
858.4153
862.5922
% Intensity
Mass (m/z)
842.0
846.8
851.6
856.4
861.2
866.0

#### 1.6 Appendix for heterobifunctional molecules 17a-g

**$^{1}$H-NMR, $^{13}$C-NMR, MALDI, HRMS and HPLC of (2S,4R)-1-((S)-2-(3-(2-(2-((2-(3,4-dichlorophenyl)acetamido)methyl)-4-(6,7-dihydro-5H-pyrrolo[1,2-a]imidazol-2-yl)phenoxy)ethoxy)propanamido)-3,3-dimethylbutanoyl)-4-hydroxy-N-(4-(4-methylthiazol-5-yl)benzyl)pyrrolidine-2-carboxamide 17a**

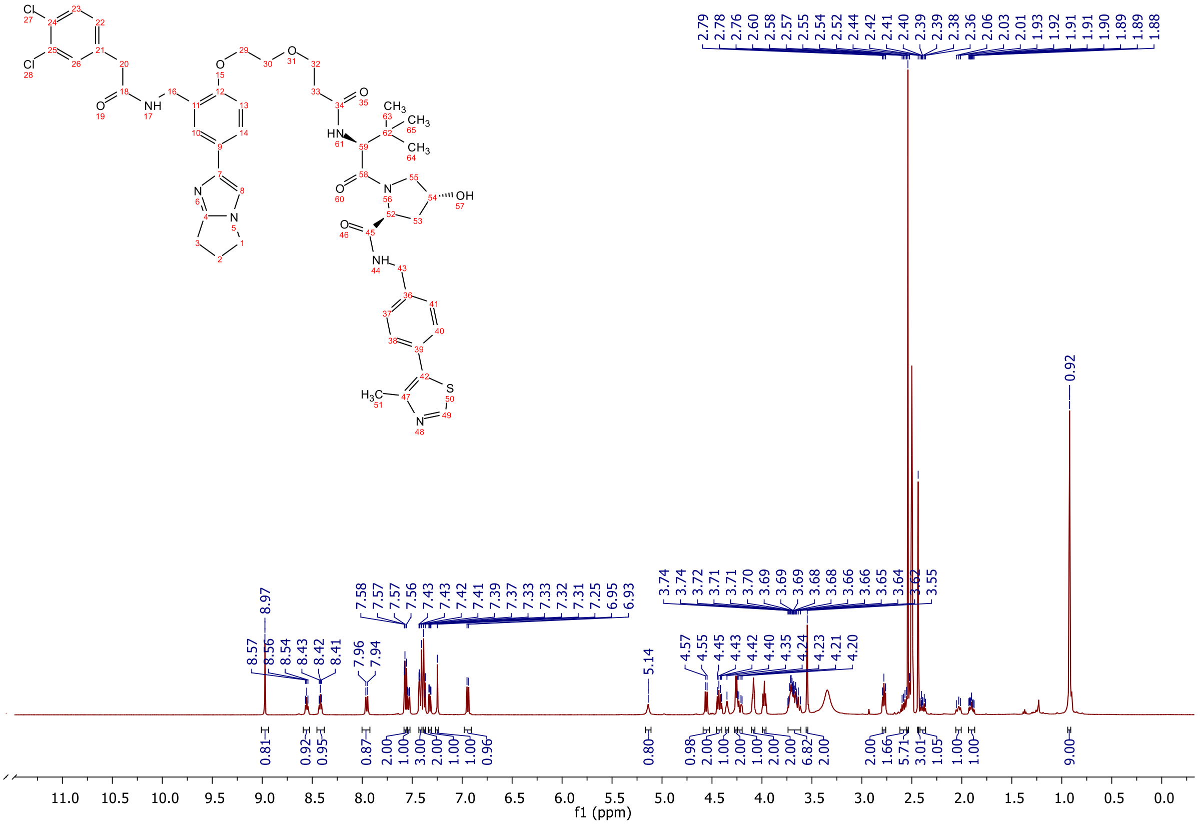

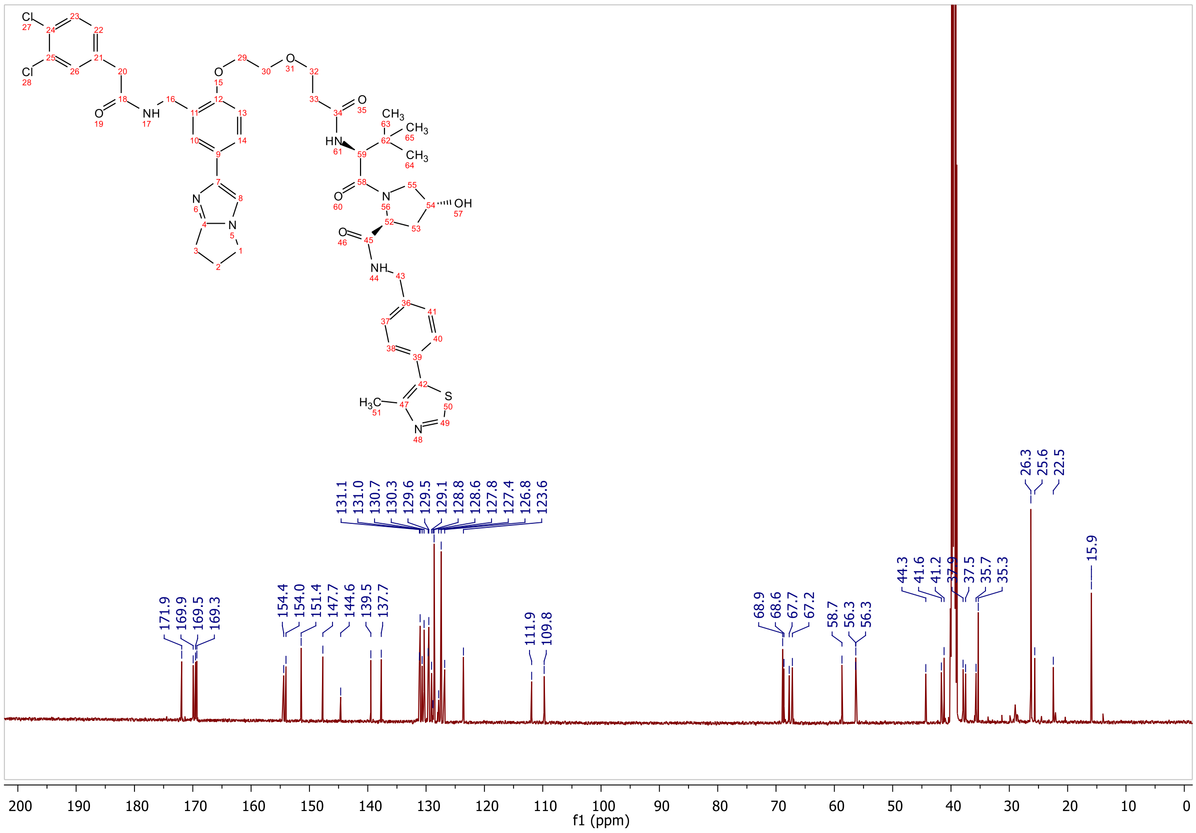

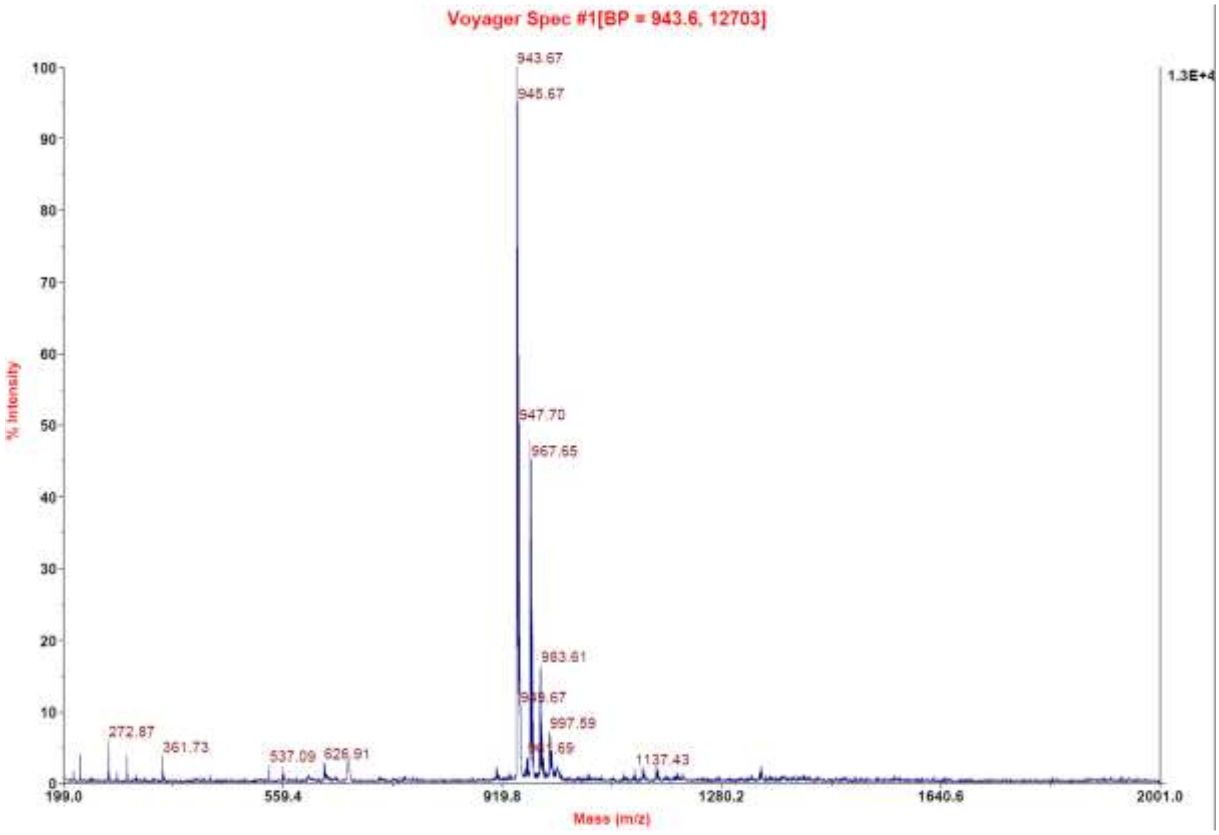
Voyager Spec #1[BP = 943.6, 12703]
% Intensity
Mass (m/z)

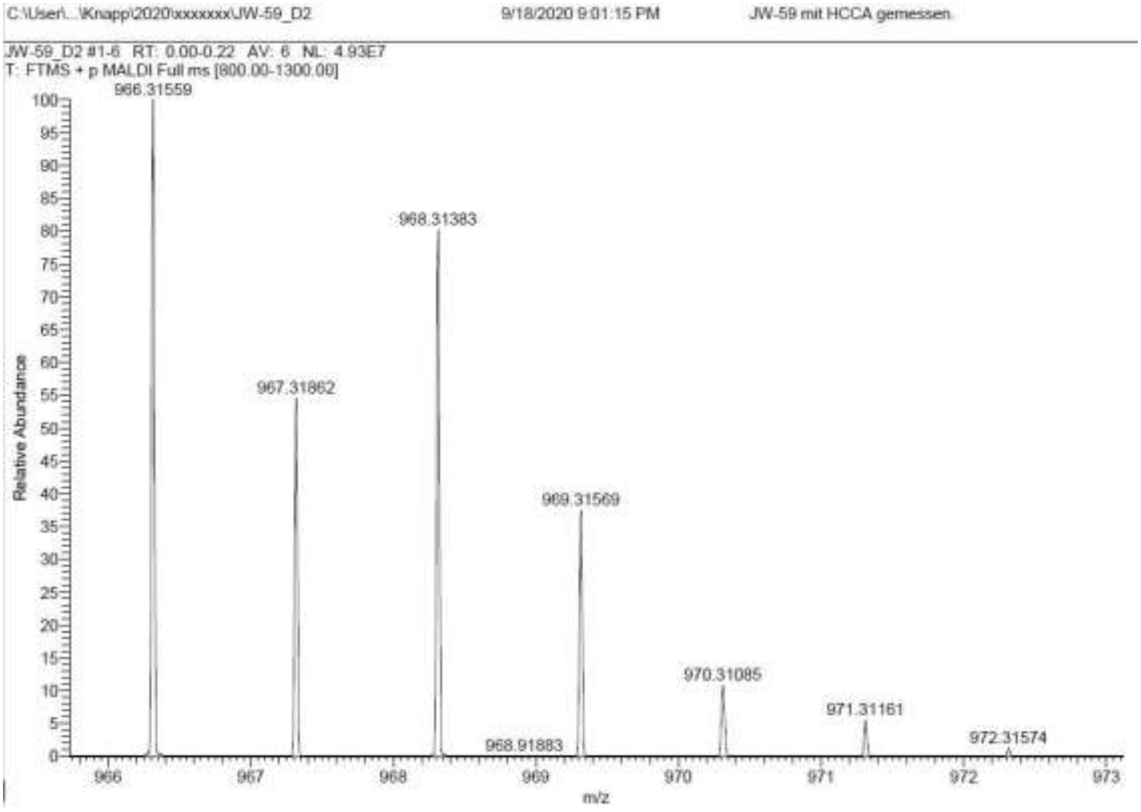
C:\User\...\Knapp\2020\xxxxxxx\JW-59_D2
9/18/2020 9:01:15 PM
JW-59 mit HCCA gemessen
JW-59_D2 #1-6 RT: 0.00-0.22 AV: 6 NL: 4.93E7
T: FTMS + p MALDI Full ms [800.00-1300.00]
966.31559
968.31383
967.31862
969.31569
970.31085
971.31161
968.91883
972.31574
Relative Abundance
m/z

**Signal:** MWD1 A, Sig=254,4 Ref=off

| RT [min] | Type | Width [min] | Area | Height | Area% |
|---|---|---|---|---|---|
| 12.256 | VV | 0.2609 | 6275.1787 | 353.9105 | 100.0000 |
| | | Sum | 6275.1787 | | |

**Signal:** MWD1 B, Sig=280,4 Ref=off

| RT [min] | Type | Width [min] | Area | Height | Area% |
|---|---|---|---|---|---|
| 12.257 | VV | 0.2518 | 7036.0225 | 412.9974 | 100.0000 |
| | | Sum | 7036.0225 | | |

**Signal:** MWD1 F, Sig=260,4 Ref=off

| RT [min] | Type | Width [min] | Area | Height | Area% |
|---|---|---|---|---|---|
| 12.256 | VV | 0.2605 | 7479.5820 | 422.5608 | 100.0000 |
| | | Sum | 7479.5820 | | |

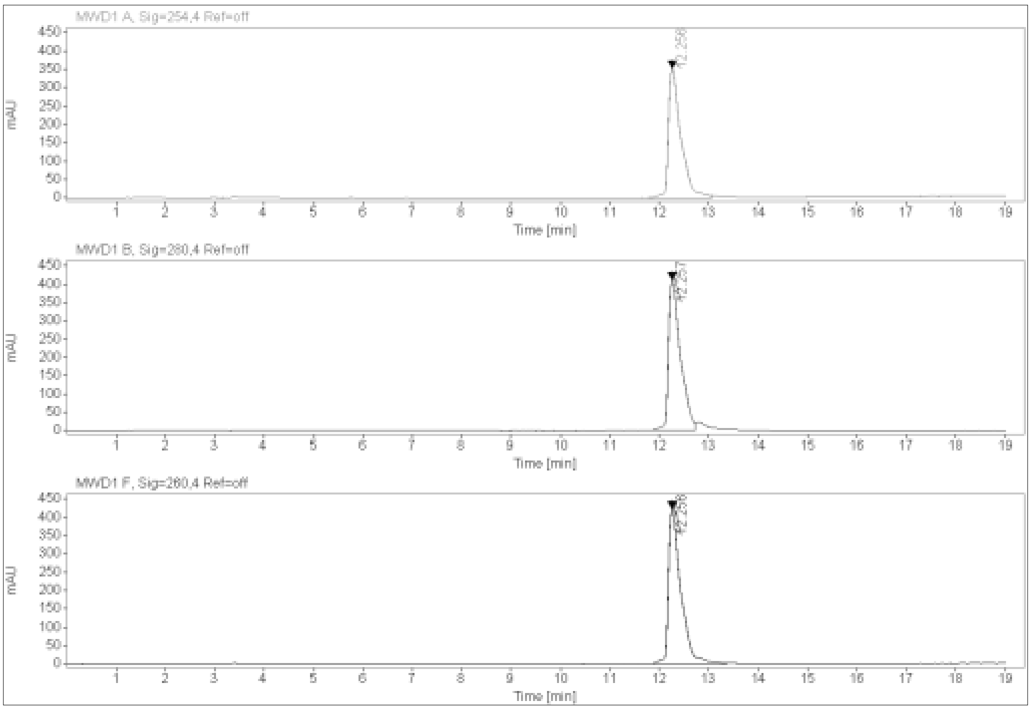

**$^{1}$H-NMR, $^{13}$C-NMR, MALDI, HRMS and HPLC of (2S,4R)-1-((S)-2-(3-(2-(2-(2-((2-(3,4-dichlorophenyl)acetamido)methyl)-4-(6,7-dihydro-5H-pyrrolo[1,2-a]imidazol-2-yl)phenoxy)ethoxy)ethoxy)propanamido)-3,3-dimethylbutanoyl)-4-hydroxy-N-(4-(4-methylthiazol-5-yl)benzyl)pyrrolidine-2-carboxamide 17b**

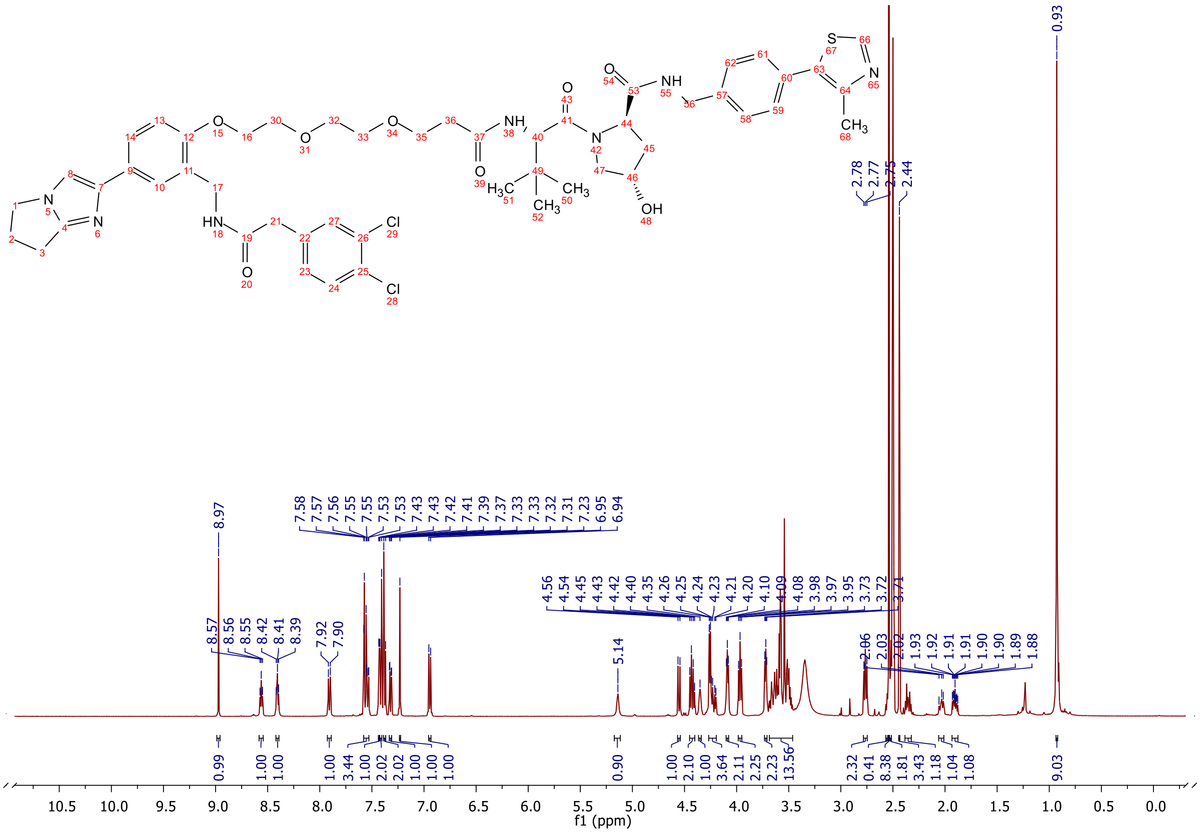

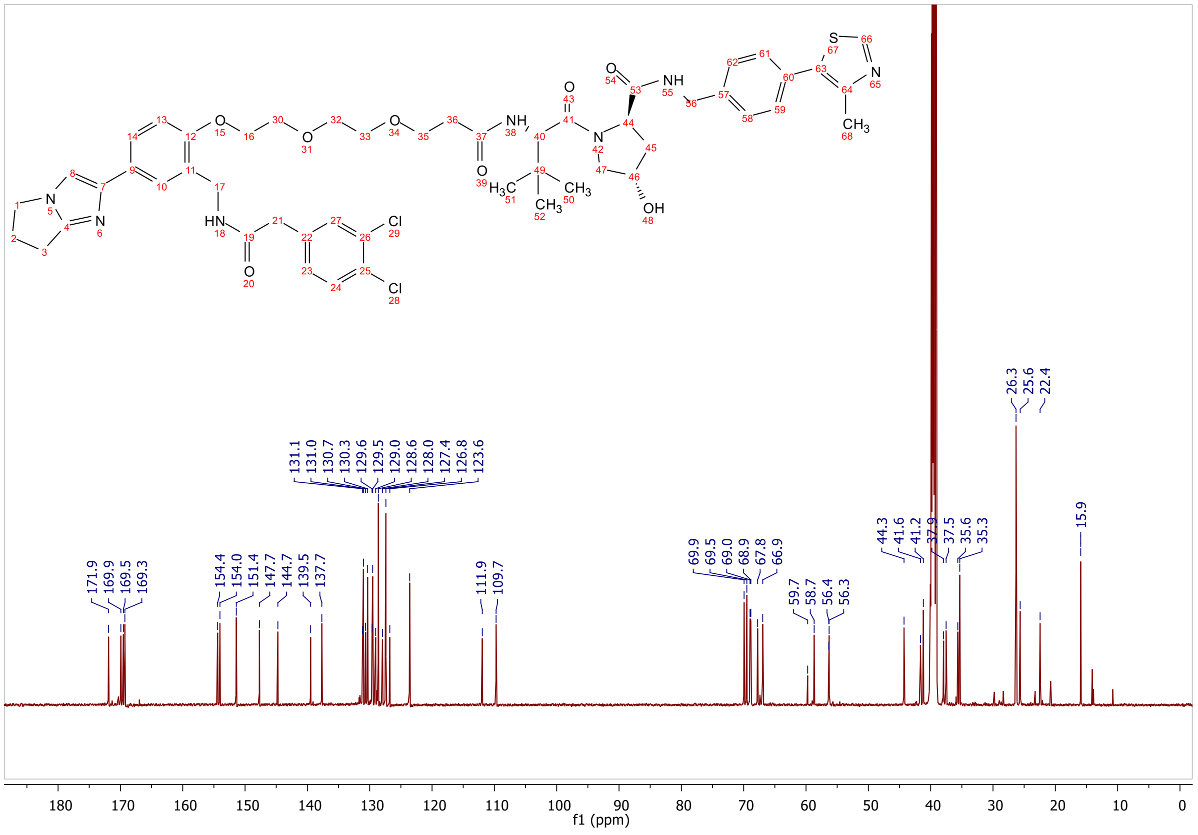

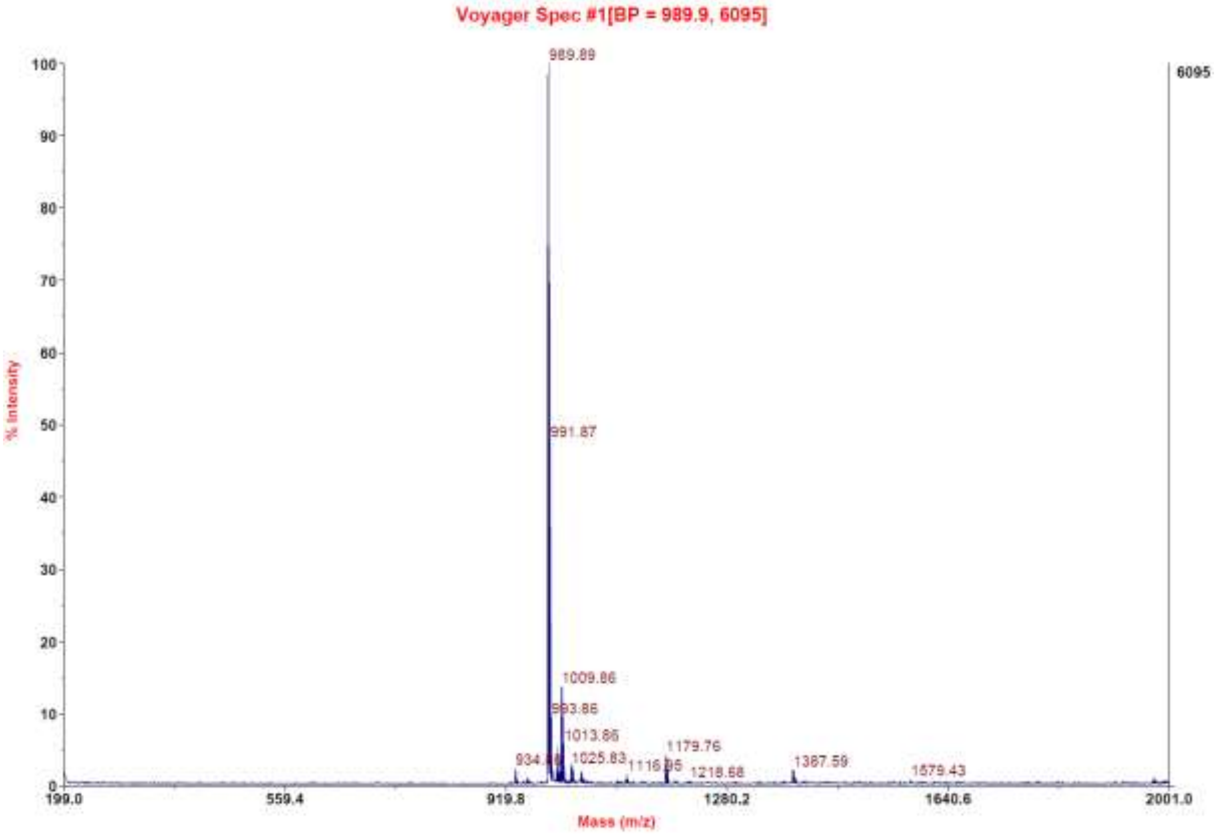
Voyager Spec #1[BP = 989.9, 6095]
989.89
991.87
1009.86
993.86
1013.86
934.
1025.83
1116.
1179.76
1218.68
1387.59
1579.43
6095
% Intensity
Mass (m/z)
199.0
559.4
919.8
1280.2
1640.6
2001.0

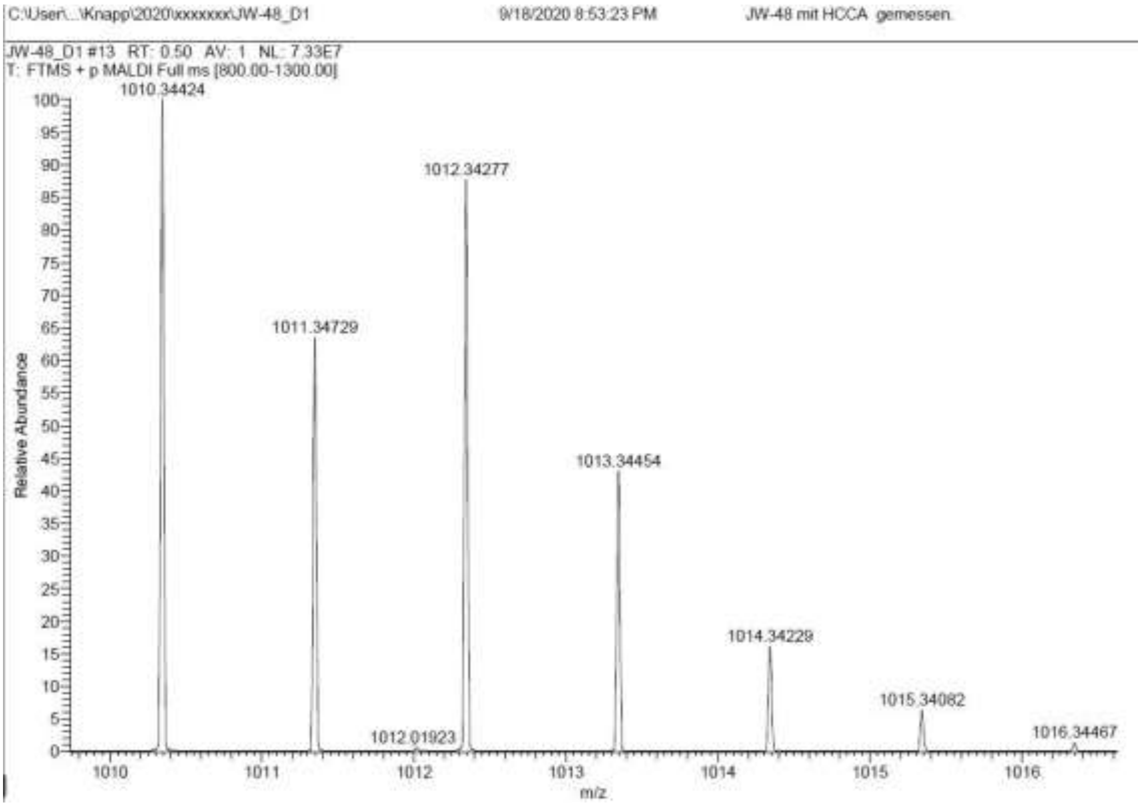
C:\User\...\Knapp\2020\xxxxxxx\JW-48_D1
9/18/2020 8:53:23 PM
JW-48 mit HCCA gemessen.
JW-48_D1 #13 RT: 0.50 AV: 1 NL: 7.33E7
T: FTMS + p MALDI Full ms [800.00-1300.00]
1010.34424
1012.34277
1011.34729
1013.34454
1014.34229
1015.34082
1012.01923
1016.34467
Relative Abundance
m/z

**Signal:** MWD1 A, Sig=254,4 Ref=off

| RT [min] | Type | Width [min] | Area | Height | Area% |
|---|---|---|---|---|---|
| 12.321 | VV | 0.2622 | 6443.7129 | 352.6285 | 100.0000 |
| | | Sum | 6443.7129 | | |

**Signal:** MWD1 B, Sig=280,4 Ref=off

| RT [min] | Type | Width [min] | Area | Height | Area% |
|---|---|---|---|---|---|
| 12.322 | VV | 0.2599 | 7143.2705 | 410.7104 | 100.0000 |
| | | Sum | 7143.2705 | | |

**Signal:** MWD1 F, Sig=260,4 Ref=off

| RT [min] | Type | Width [min] | Area | Height | Area% |
|---|---|---|---|---|---|
| 12.321 | VV | 0.2600 | 7610.5439 | 420.8916 | 100.0000 |
| | | Sum | 7610.5439 | | |

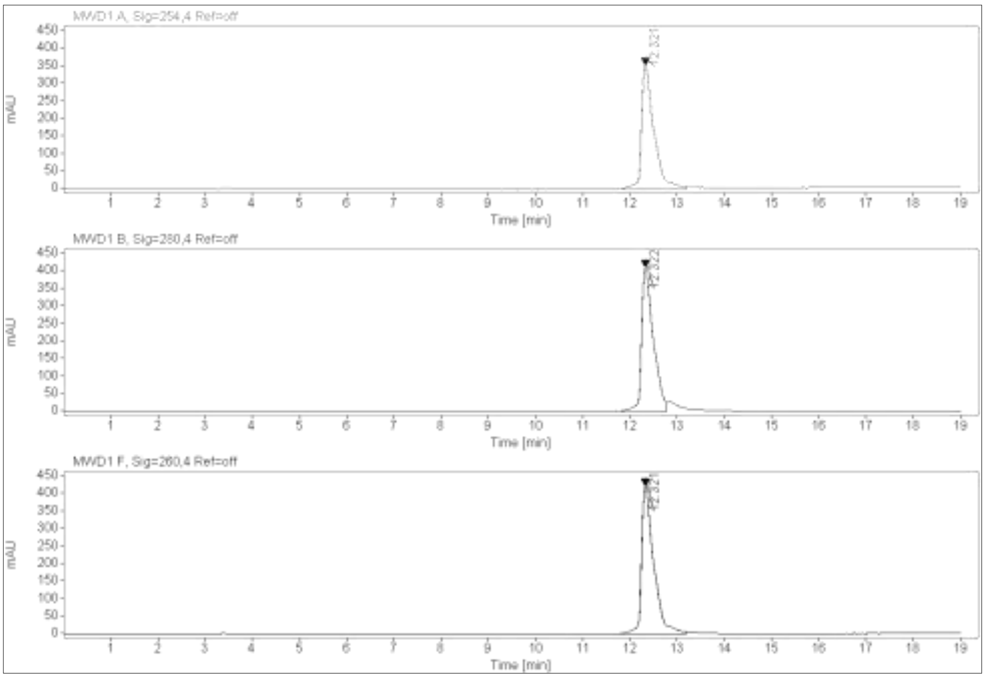

**$^{1}$H-NMR, $^{13}$C-NMR, MALDI, HRMS and HPLC of (2S,4R)-1-((S)-14-(tert-butyl)-1-(2-((2-(3,4-dichlorophenyl)acetamido)methyl)-4-(6,7-dihydro-5H-pyrrolo[1,2-a]imidazol-2-yl)phenoxy)-12-oxo-3,6,9-trioxa-13-azapentadecan-15-oyl)-4-hydroxy-N-(4-(4-methylthiazol-5-yl)benzyl)pyrrolidine-2-carboxamide 17c**

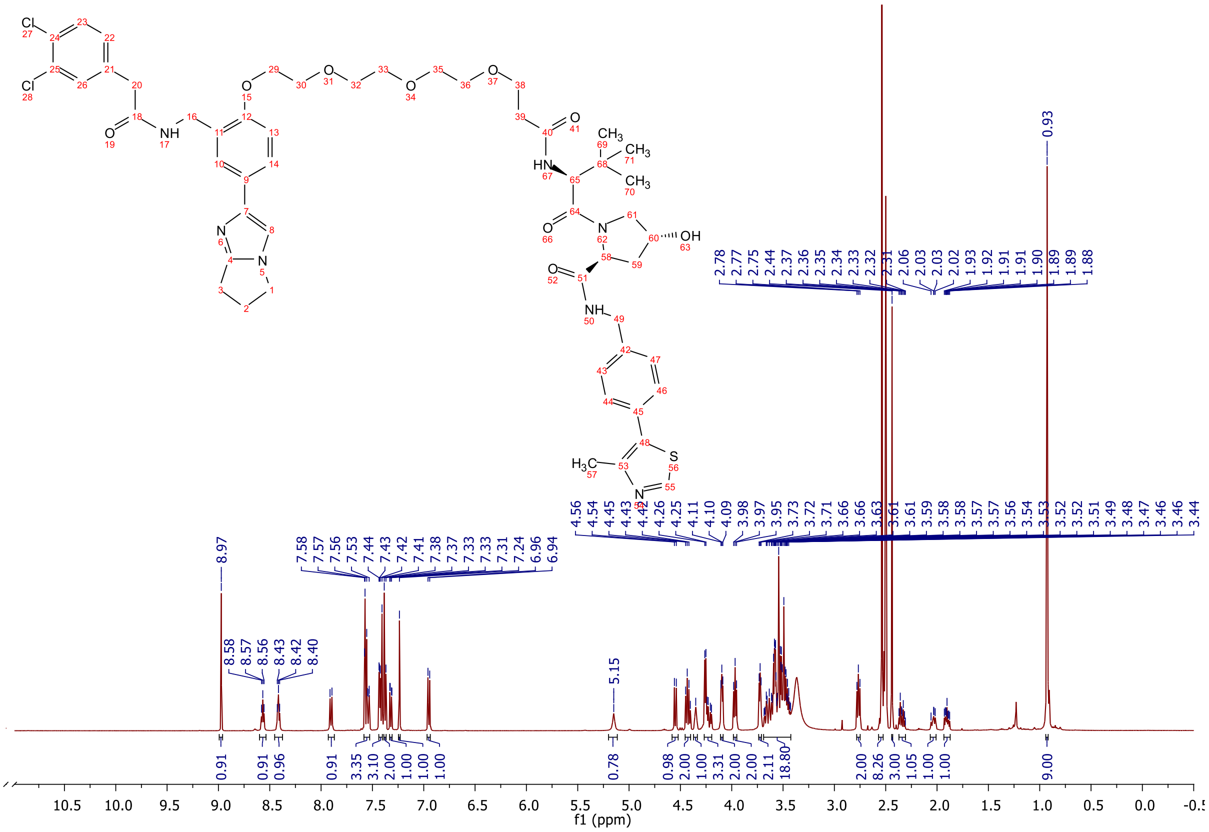

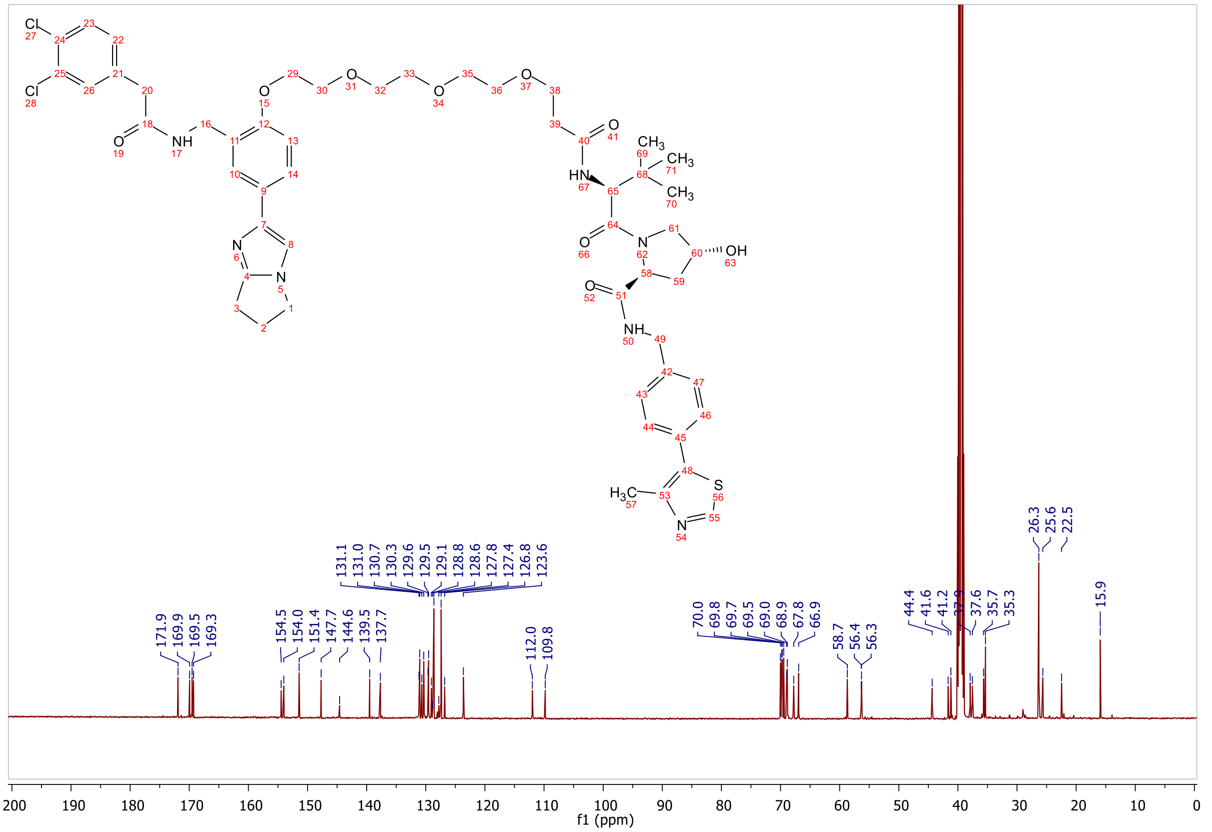

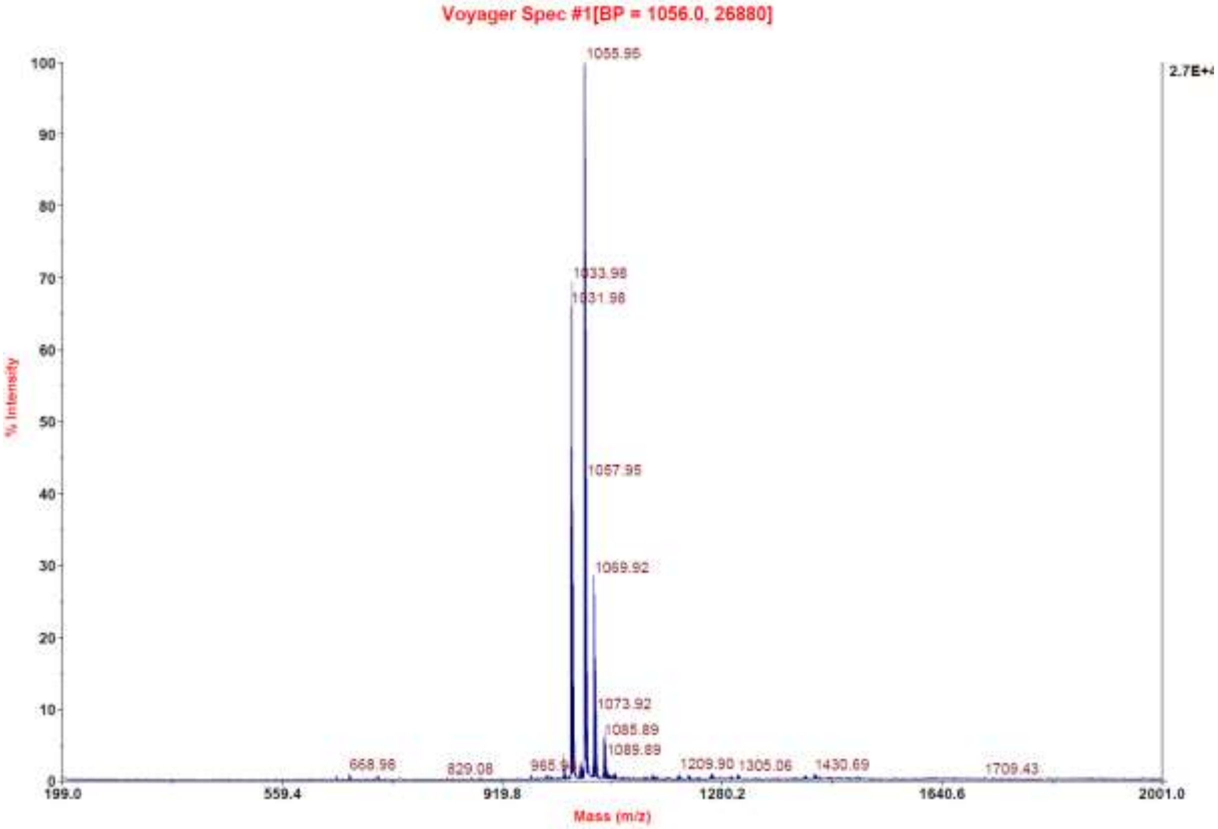

C:\User\...\Knapp\2020\xxxxxxx\JW-60_D3 9/18/2020 9:01:49 PM JW-60 mit HCCA gemessen.

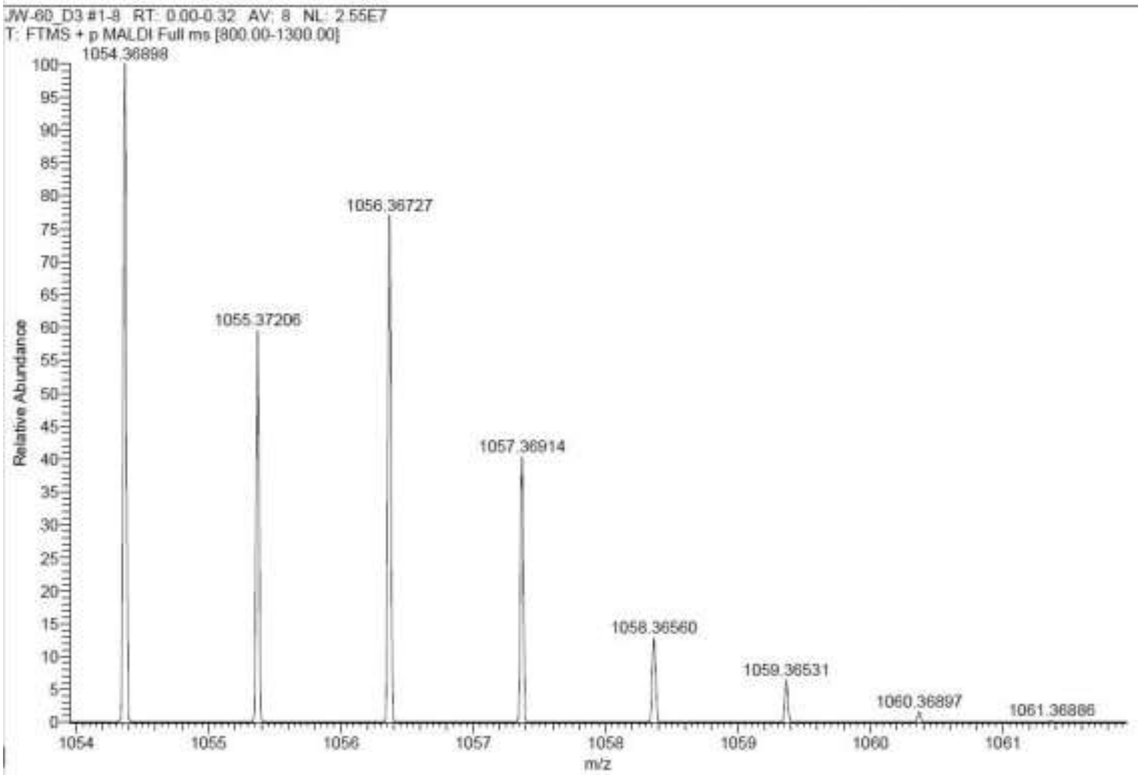

**Signal:** MWD1 A, Sig=254,4 Ref=off

| RT [min] | Type | Width [min] | Area | Height | Area% |
|---|---|---|---|---|---|
| 12.349 | VV | 0.2700 | 5961.0488 | 322.0216 | 100.0000 |
| | | Sum | 5961.0488 | | |

**Signal:** MWD1 B, Sig=280,4 Ref=off

| RT [min] | Type | Width [min] | Area | Height | Area% |
|---|---|---|---|---|---|
| 12.349 | VV | 0.2727 | 7138.7886 | 375.7160 | 100.0000 |
| | | Sum | 7138.7886 | | |

**Signal:** MWD1 F, Sig=260,4 Ref=off

| RT [min] | Type | Width [min] | Area | Height | Area% |
|---|---|---|---|---|---|
| 12.349 | VV | 0.2660 | 7048.1709 | 384.1659 | 100.0000 |
| | | Sum | 7048.1709 | | |

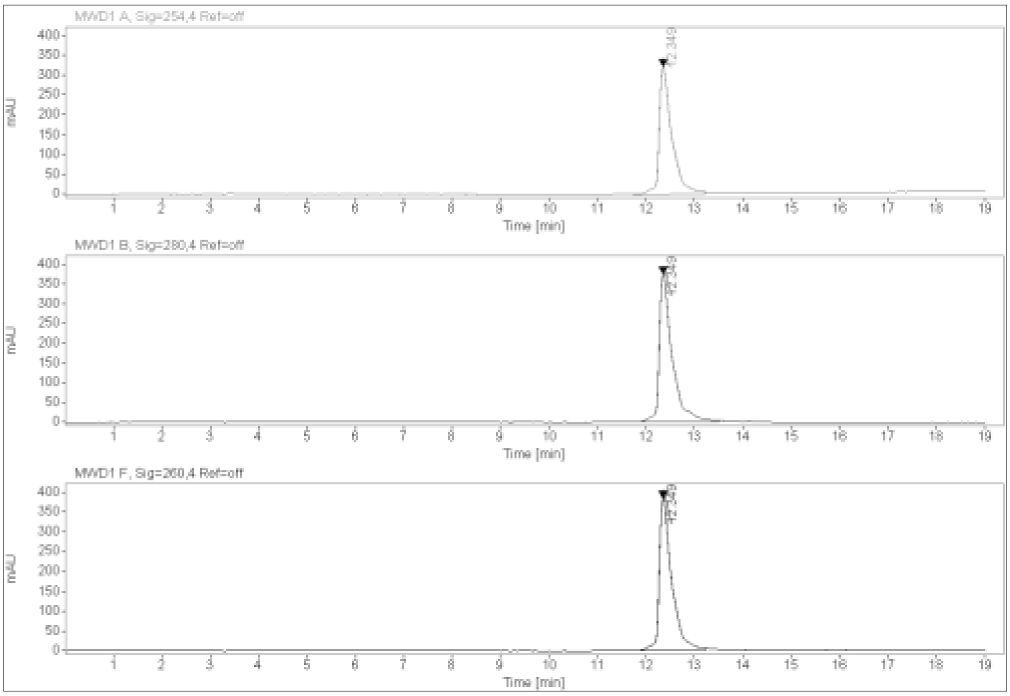

**$^{1}$H-NMR, $^{13}$C-NMR, MALDI, HRMS and HPLC of (2S,4R)-1-((S)-17-(tert-butyl)-1-(2-((2-(3,4-dichlorophenyl)acetamido)methyl)-4-(6,7-dihydro-5H-pyrrolo[1,2-a]imidazol-2-yl)phenoxy)-15-oxo-3,6,9,12-tetraoxa-16-azaoctadecan-18-oyl)-4-hydroxy-N-(4-(4-methylthiazol-5-yl)benzyl)pyrrolidine-2-carboxamide 17d**

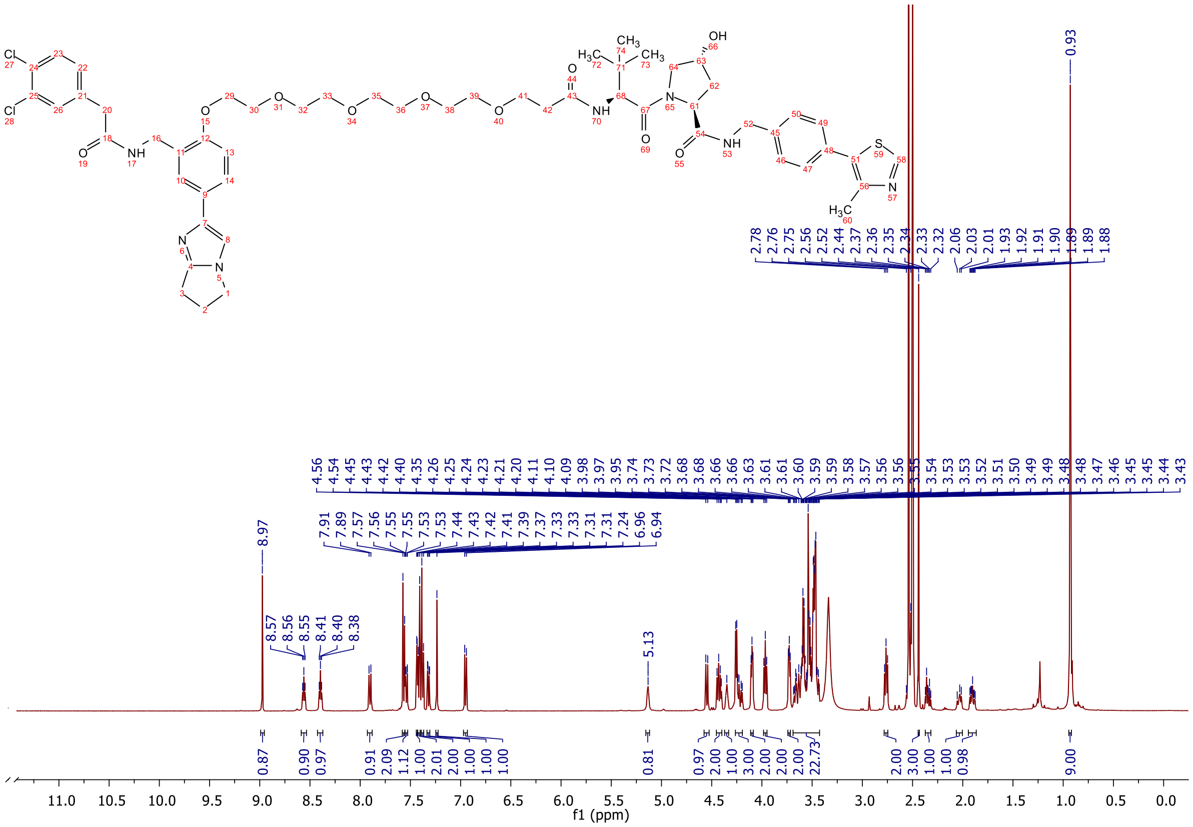

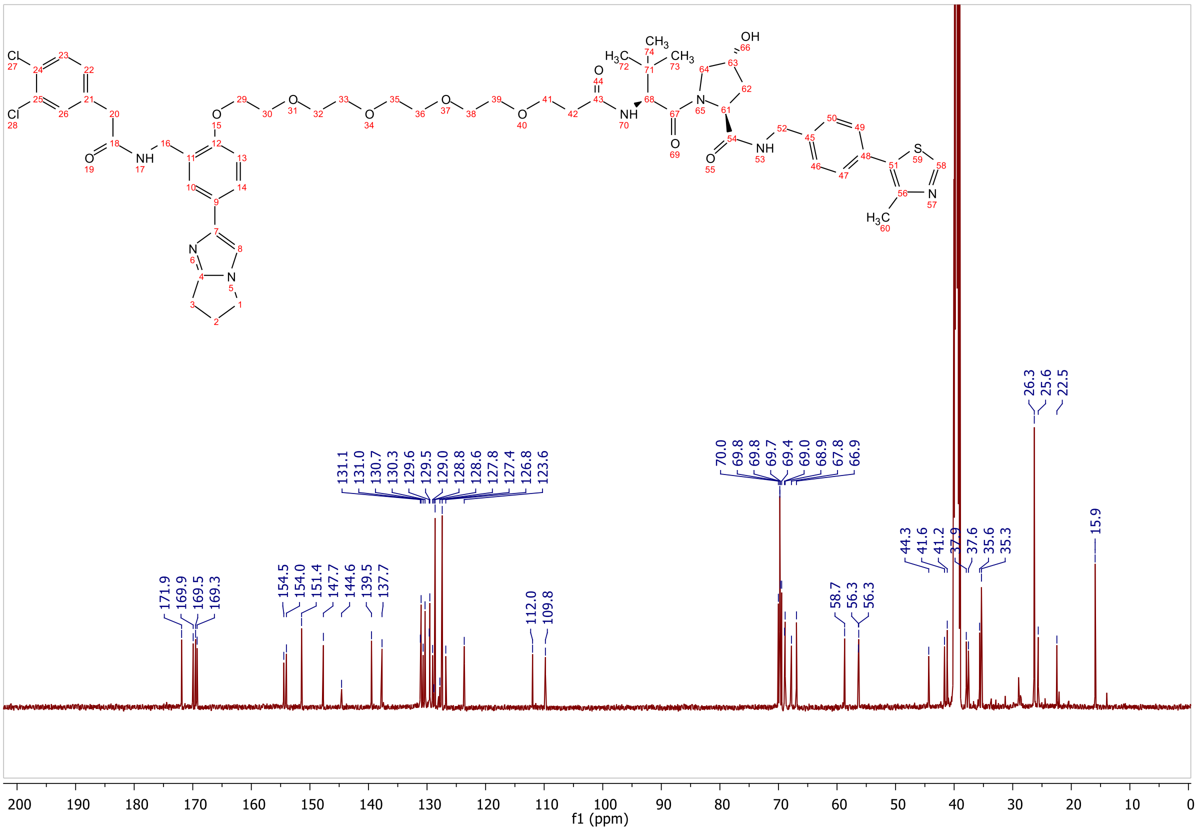

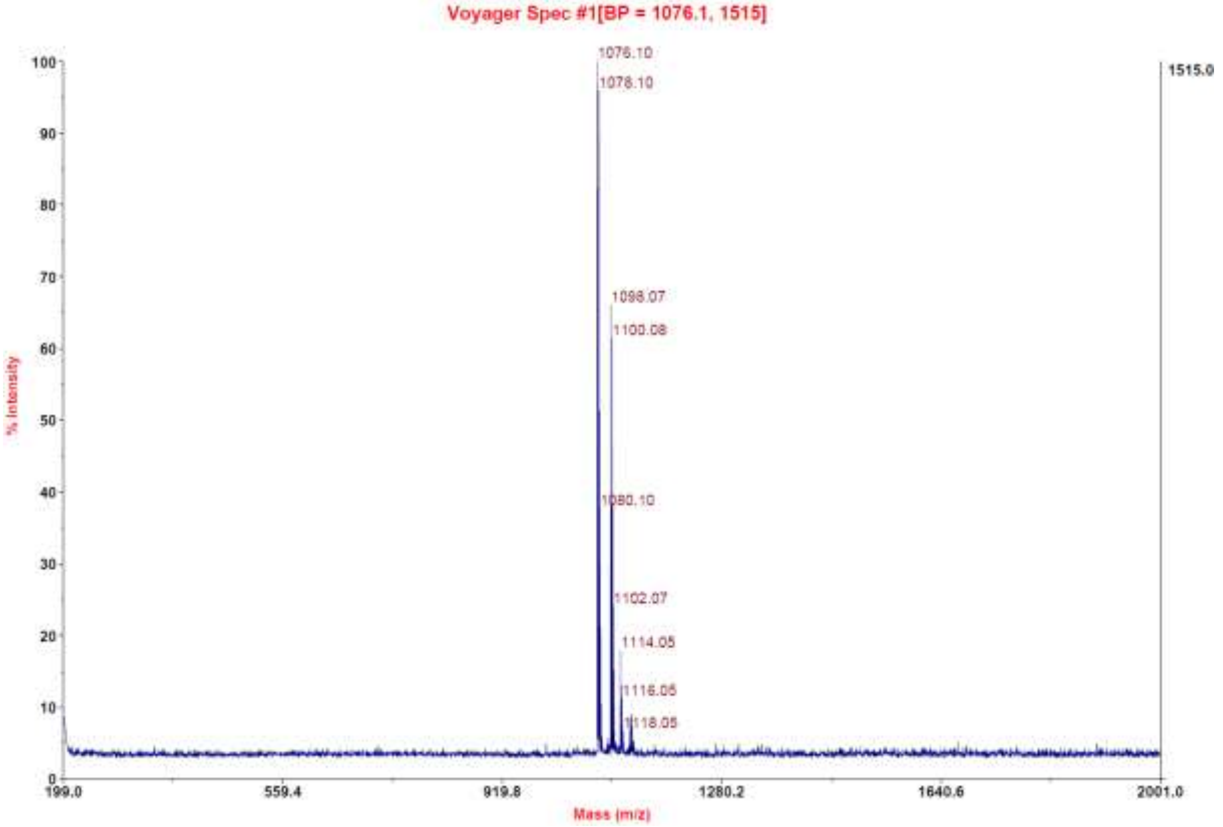
Voyager Spec #1[BP = 1076.1, 1515]
1076.10
1078.10
1098.07
1100.08
1080.10
1102.07
1114.05
1116.05
1118.05
1515.0
% Intensity
Mass (m/z)
199.0
559.4
919.8
1280.2
1640.6
2001.0

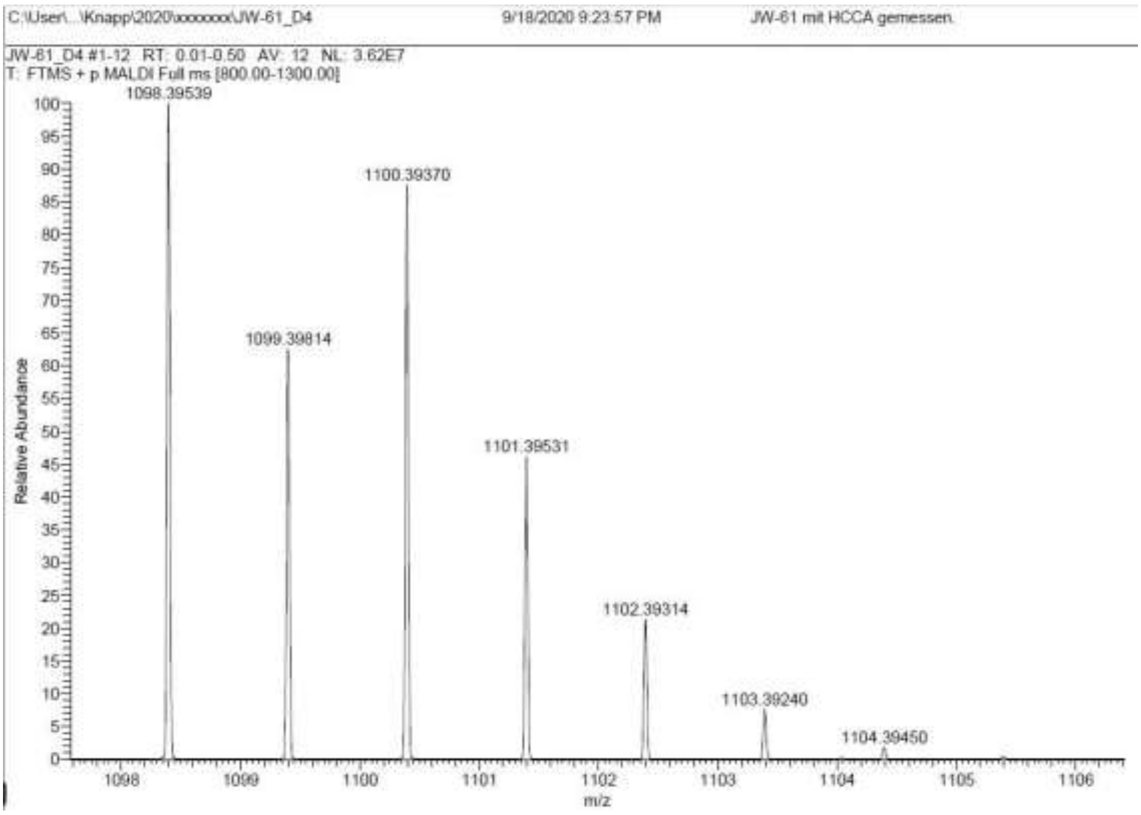
C:\User\...\Knapp\2020\xxxxxxx\JW-61_D4
9/18/2020 9:23:57 PM
JW-61 mit HCCA gemessen
JW-61_D4 #1-12 RT: 0.01-0.50 AV: 12 NL: 3.62E7
T: FTMS + p MALDI Full ms [800.00-1300.00]
1098.39539
1100.39370
1099.39814
1101.39531
1102.39314
1103.39240
1104.39450
Relative Abundance
m/z

**Signal:** MWD1 A, Sig=254,4 Ref=off

| RT [min] | Type | Width [min] | Area | Height | Area% |
|---|---|---|---|---|---|
| 12.372 | VV | 0.2762 | 6240.8027 | 332.1096 | 100.0000 |
| | | Sum | 6240.8027 | | |

**Signal:** MWD1 B, Sig=280,4 Ref=off

| RT [min] | Type | Width [min] | Area | Height | Area% |
|---|---|---|---|---|---|
| 12.373 | VV | 0.2786 | 7359.5869 | 387.4797 | 100.0000 |
| | | Sum | 7359.5869 | | |

**Signal:** MWD1 F, Sig=260,4 Ref=off

| RT [min] | Type | Width [min] | Area | Height | Area% |
|---|---|---|---|---|---|
| 12.373 | VV | 0.2674 | 7285.1592 | 394.6602 | 100.0000 |
| | | Sum | 7285.1592 | | |

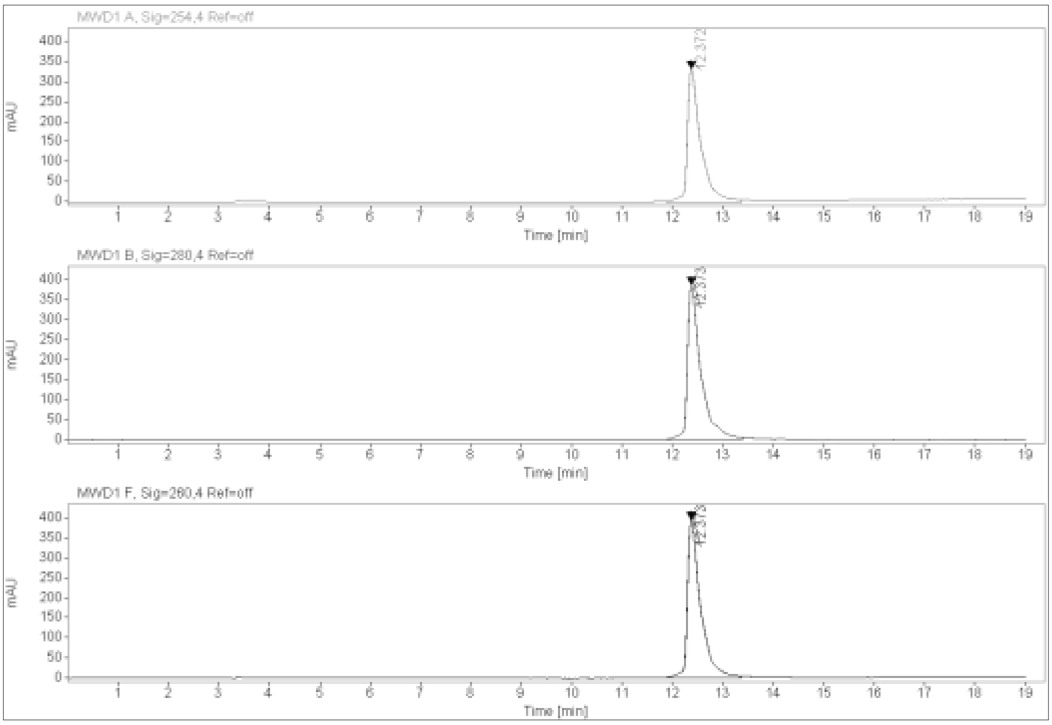

**$^{1}$H-NMR, $^{13}$C-NMR, MALDI, HRMS and HPLC of (2S,4R)-1-((S)-20-(tert-butyl)-1-(2-((2-(3,4-dichlorophenyl)acetamido)methyl)-4-(6,7-dihydro-5H-pyrrolo[1,2-a]imidazol-2-yl)phenoxy)-18-oxo-3,6,9,12,15-pentaoxa-19-azahenicosan-21-oyl)-4-hydroxy-N-(4-(4-methylthiazol-5-yl)benzyl)pyrrolidine-2-carboxamide 17e**

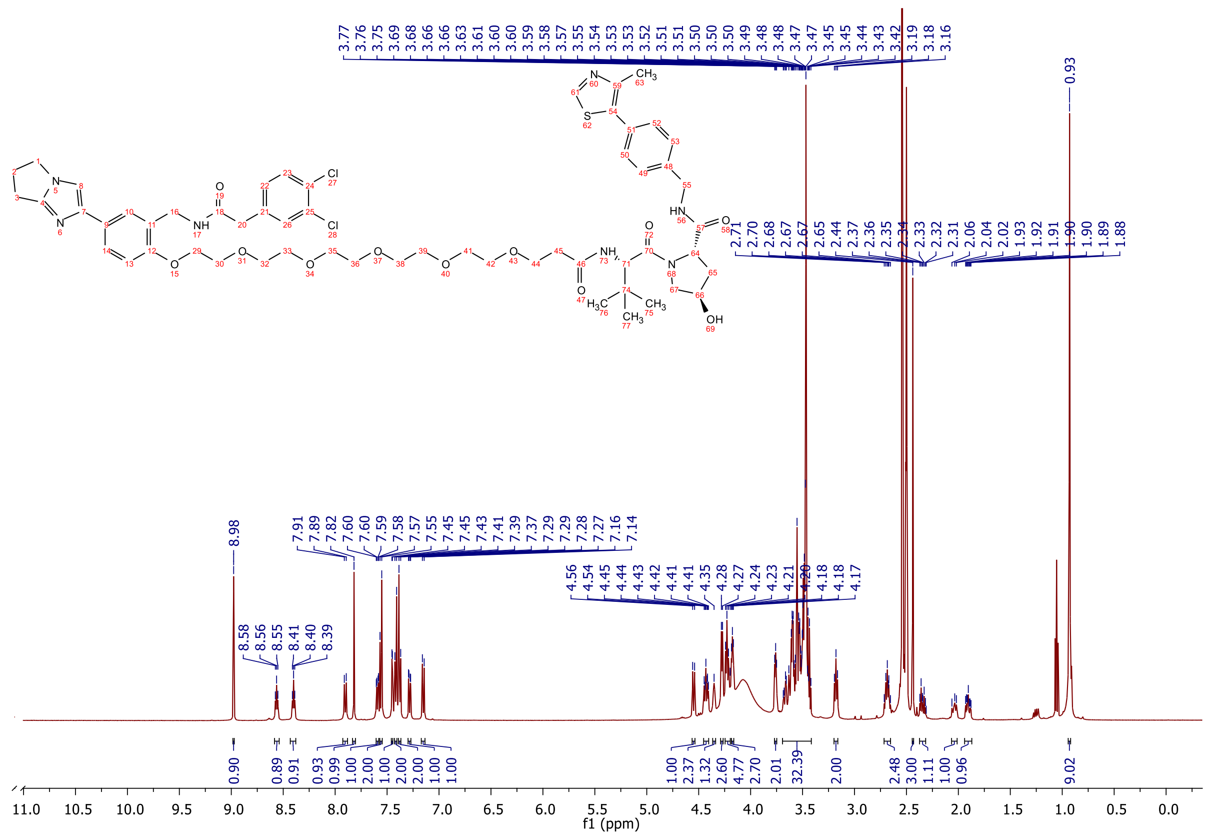

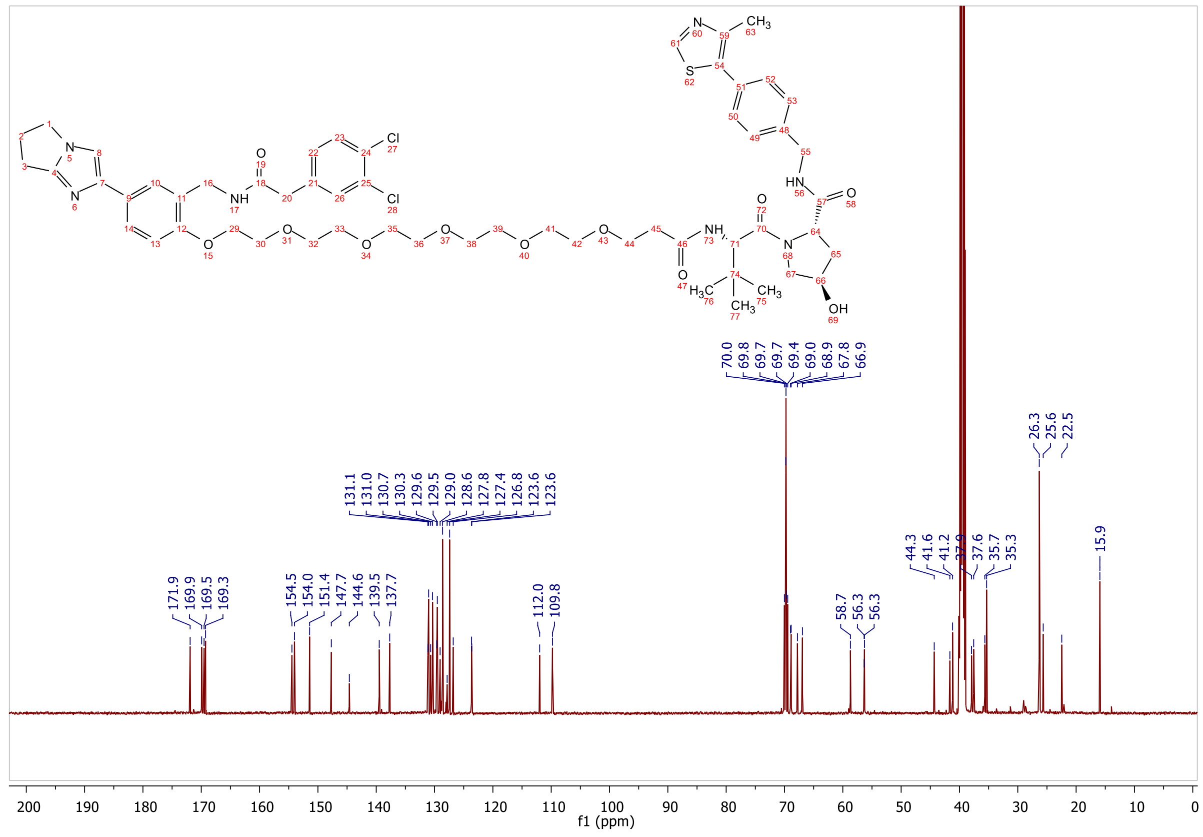

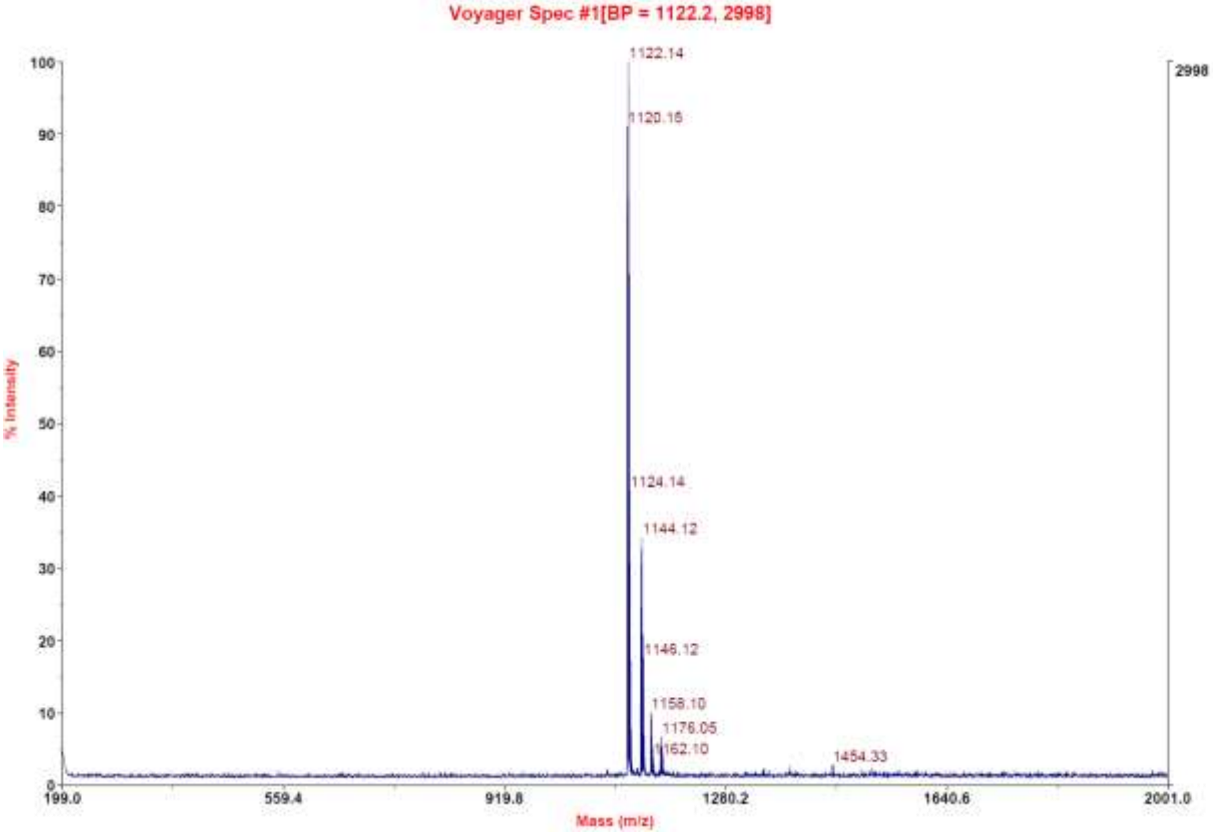
Voyager Spec #1[BP = 1122.2, 2998]
1122.14
1120.16
1124.14
1144.12
1146.12
1158.10
1176.05
1162.10
1454.33
2998
% Intensity
Mass (m/z)
199.0
559.4
919.8
1280.2
1640.6
2001.0

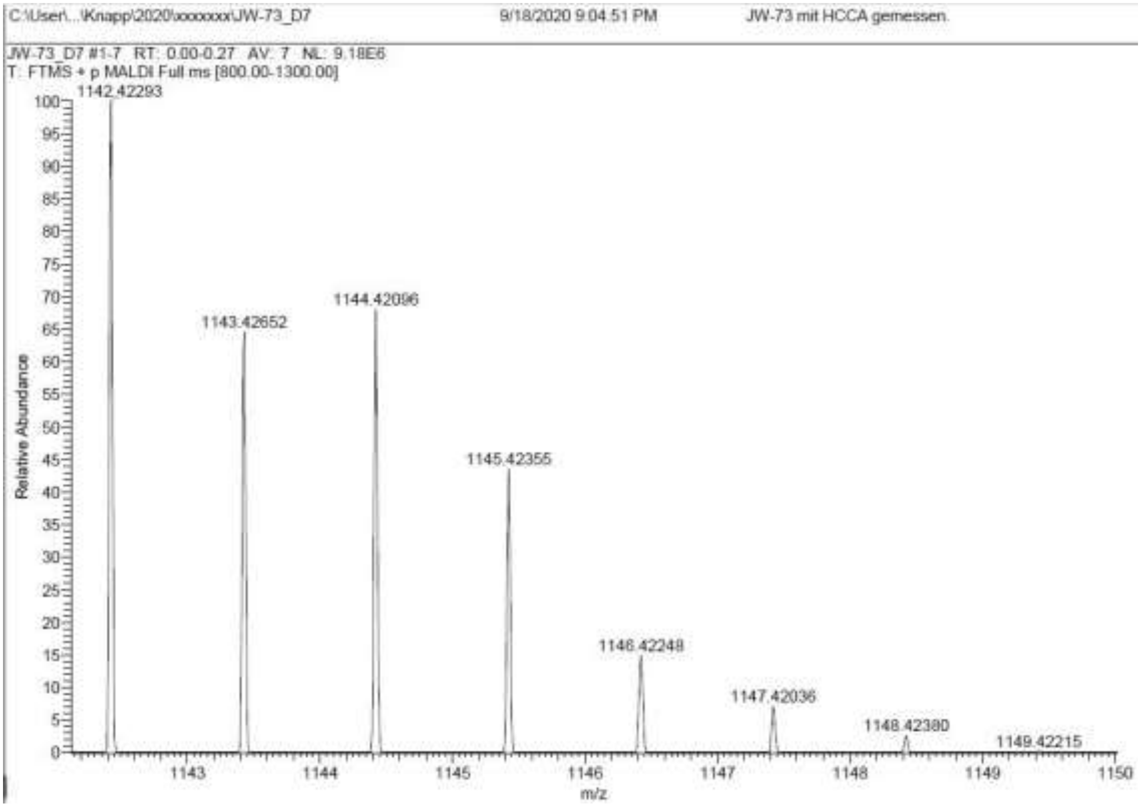
C:\User\...\Knapp\2020\xxxxxxx\JW-73_D7
9/18/2020 9:04:51 PM
JW-73 mit HCCA gemessen.
JW-73_D7 #1-7 RT: 0.00-0.27 AV: 7 NL: 9.18E6
T: FTMS + p MALDI Full ms [800.00-1300.00]
1142.42293
1143.42652
1144.42096
1145.42355
1146.42248
1147.42036
1148.42380
1149.42215
Relative Abundance
m/z

**Signal:** MWD1 A, Sig=254,4 Ref=off

| RT [min] | Type | Width [min] | Area | Height | Area% |
|---|---|---|---|---|---|
| 12.375 | VV | 0.2566 | 9890.8145 | 569.6321 | 100.0000 |
| | | Sum | 9890.8145 | | |

**Signal:** MWD1 B, Sig=280,4 Ref=off

| RT [min] | Type | Width [min] | Area | Height | Area% |
|---|---|---|---|---|---|
| 12.375 | VV | 0.2575 | 11565.6602 | 666.2625 | 100.0000 |
| | | Sum | 11565.6602 | | |

**Signal:** MWD1 F, Sig=260,4 Ref=off

| RT [min] | Type | Width [min] | Area | Height | Area% |
|---|---|---|---|---|---|
| 12.375 | VV | 0.2558 | 11773.2158 | 680.7462 | 100.0000 |
| | | Sum | 11773.2158 | | |

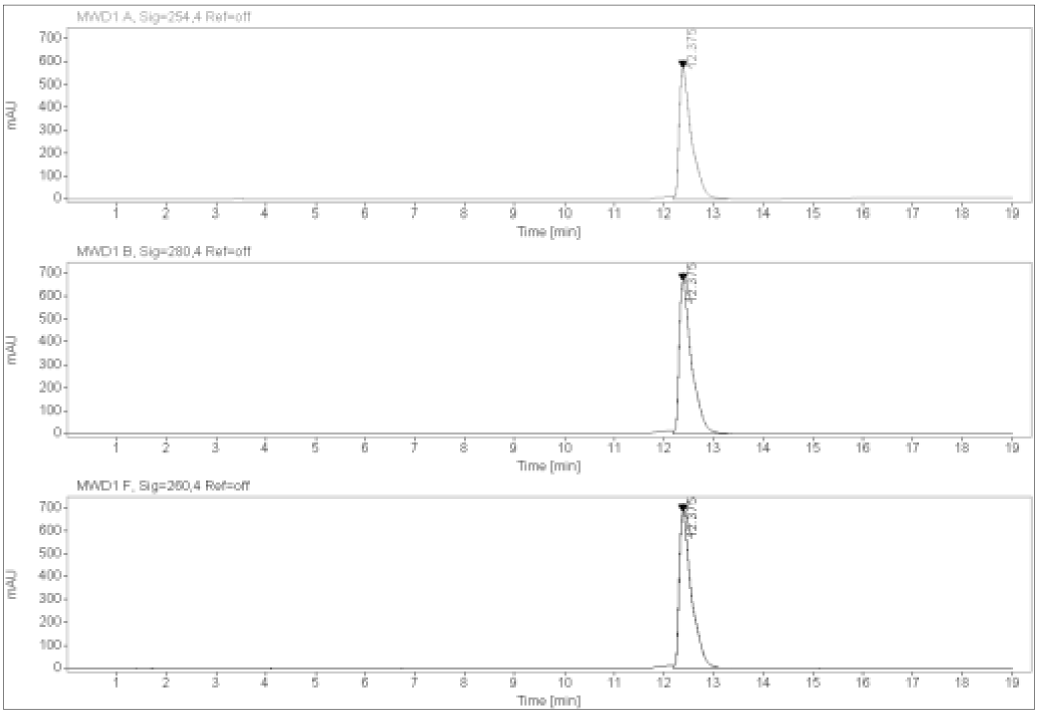

**$^{1}$H-NMR, $^{13}$C-NMR, MALDI, HRMS and HPLC of (2S,4R)-1-((S)-23-(tert-butyl)-1-(2-((2-(3,4-dichlorophenyl)acetamido)methyl)-4-(6,7-dihydro-5H-pyrrolo[1,2-a]imidazol-2-yl)phenoxy)-21-oxo-3,6,9,12,15,18-hexaoxa-22-azatetracosan-24-oyl)-4-hydroxy-N-(4-(4-methylthiazol-5-yl)benzyl)pyrrolidine-2-carboxamide 17f**

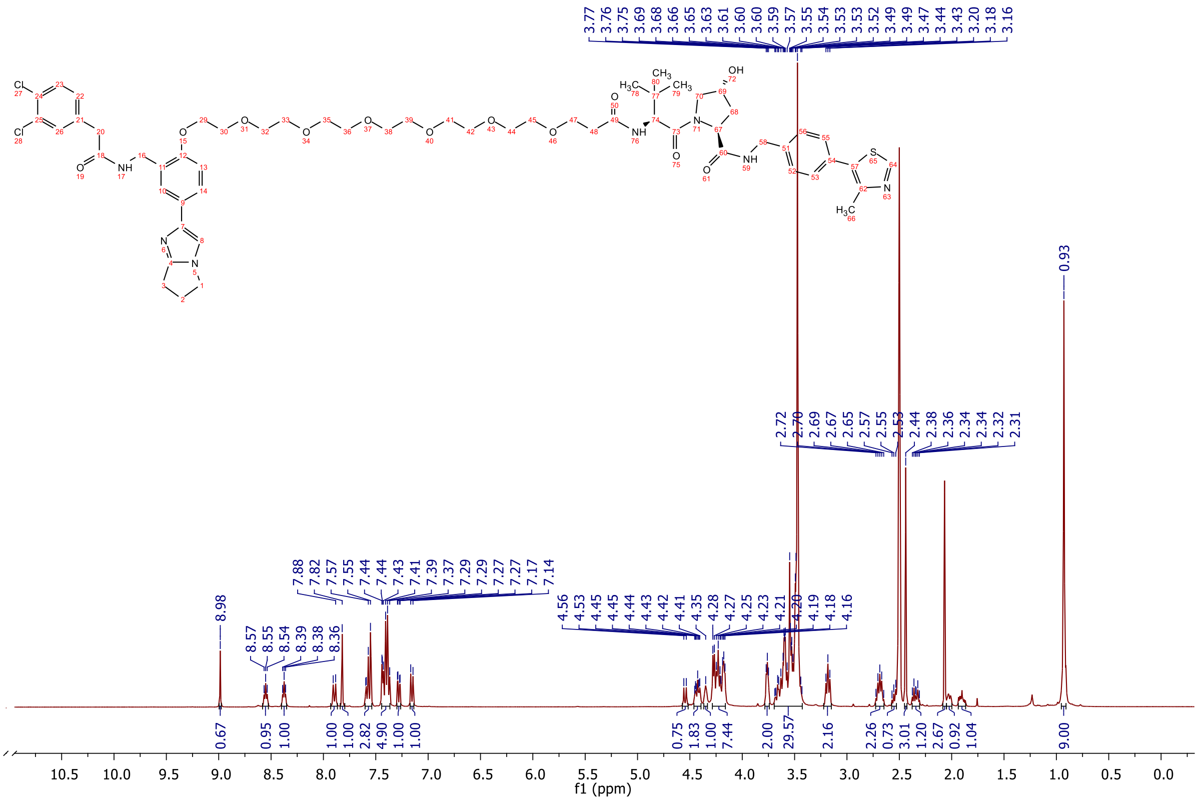

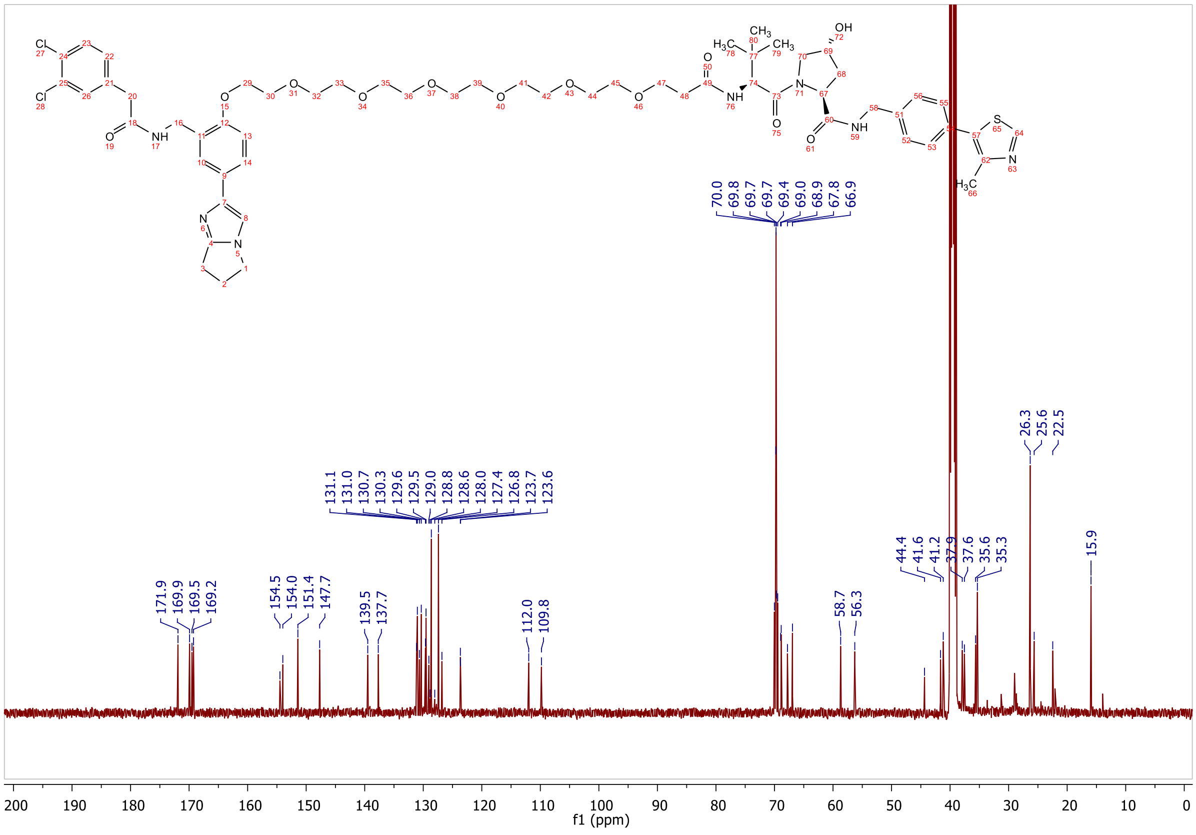

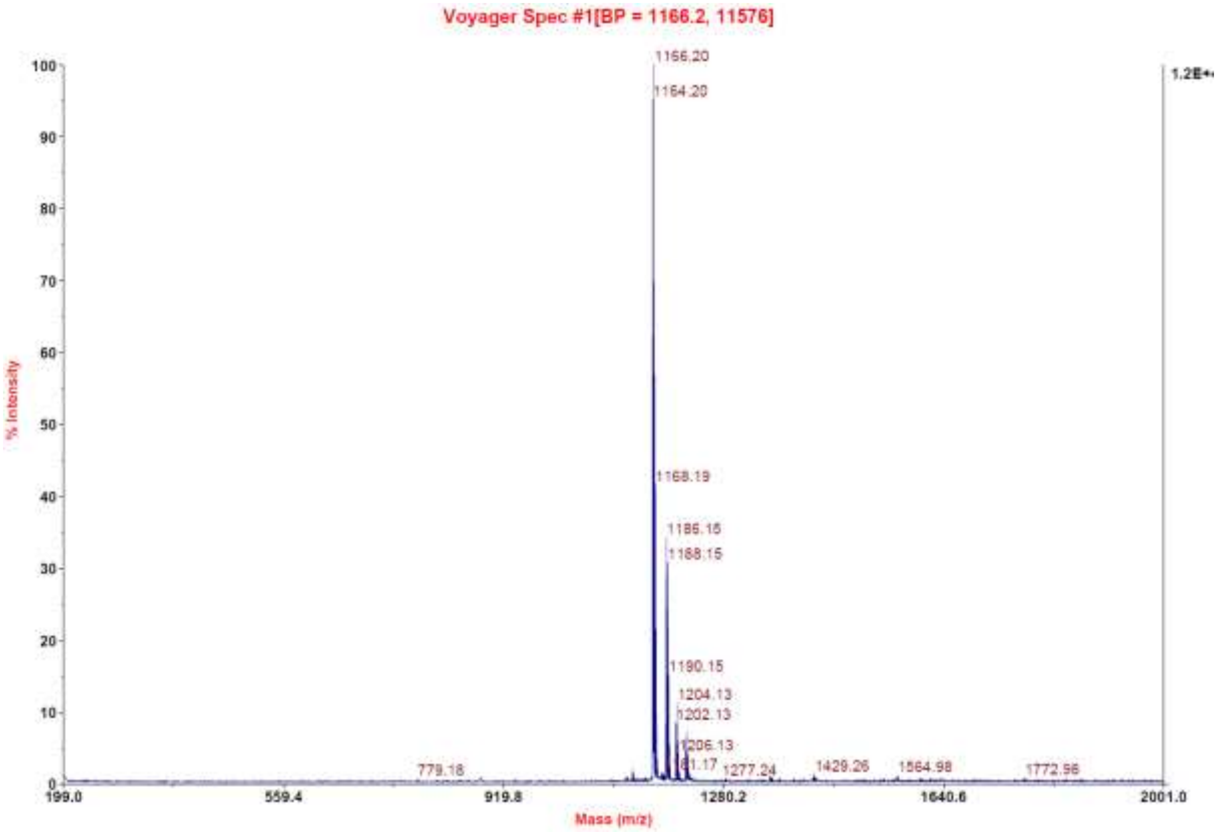
Voyager Spec #1[BP = 1166.2, 11576]
% Intensity
Mass (m/z)
1166.20
1164.20
1168.19
1186.15
1188.15
1190.15
1204.13
1202.13
1206.13
1277.24
1429.26
1564.98
1772.96
779.18
1.2E+

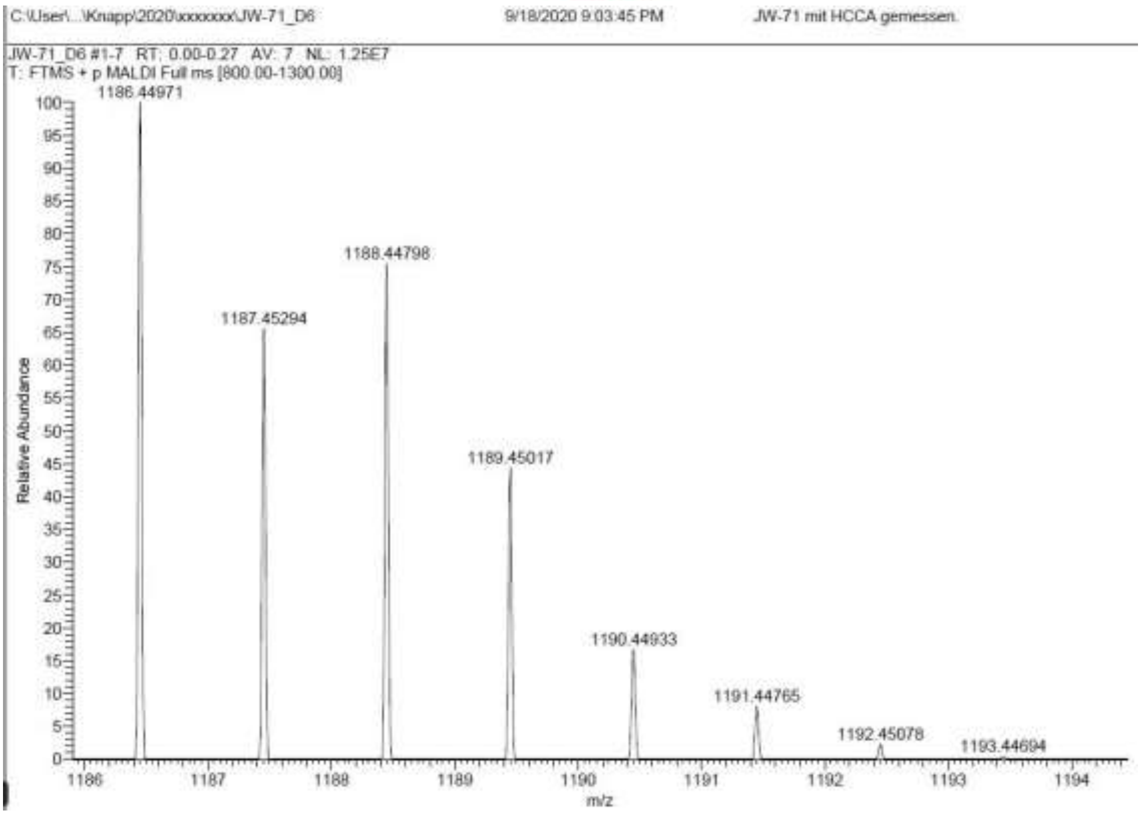
C:\User\...\Knapp\2020\xxxxxxx\JW-71_D6
9/18/2020 9:03:45 PM
JW-71 mit HCCA gemessen.
JW-71_D6 #1-7 RT: 0.00-0.27 AV: 7 NL: 1.25E7
T: FTMS + p MALDI Full ms [800.00-1300.00]
Relative Abundance
m/z
1186.44971
1187.45294
1188.44798
1189.45017
1190.44933
1191.44765
1192.45078
1193.44694

**Signal:** MWD1 A, Sig=254,4 Ref=off

| RT [min] | Type | Width [min] | Area | Height | Area% |
|---|---|---|---|---|---|
| 12.394 | VV | 0.2659 | 7949.5796 | 448.2770 | 100.0000 |
| | | Sum | 7949.5796 | | |

**Signal:** MWD1 B, Sig=280,4 Ref=off

| RT [min] | Type | Width [min] | Area | Height | Area% |
|---|---|---|---|---|---|
| 12.394 | BV | 0.2616 | 9266.4102 | 520.8672 | 100.0000 |
| | | Sum | 9266.4102 | | |

**Signal:** MWD1 F, Sig=260,4 Ref=off

| RT [min] | Type | Width [min] | Area | Height | Area% |
|---|---|---|---|---|---|
| 12.394 | VV | 0.2651 | 9394.3320 | 534.4274 | 100.0000 |
| | | Sum | 9394.3320 | | |

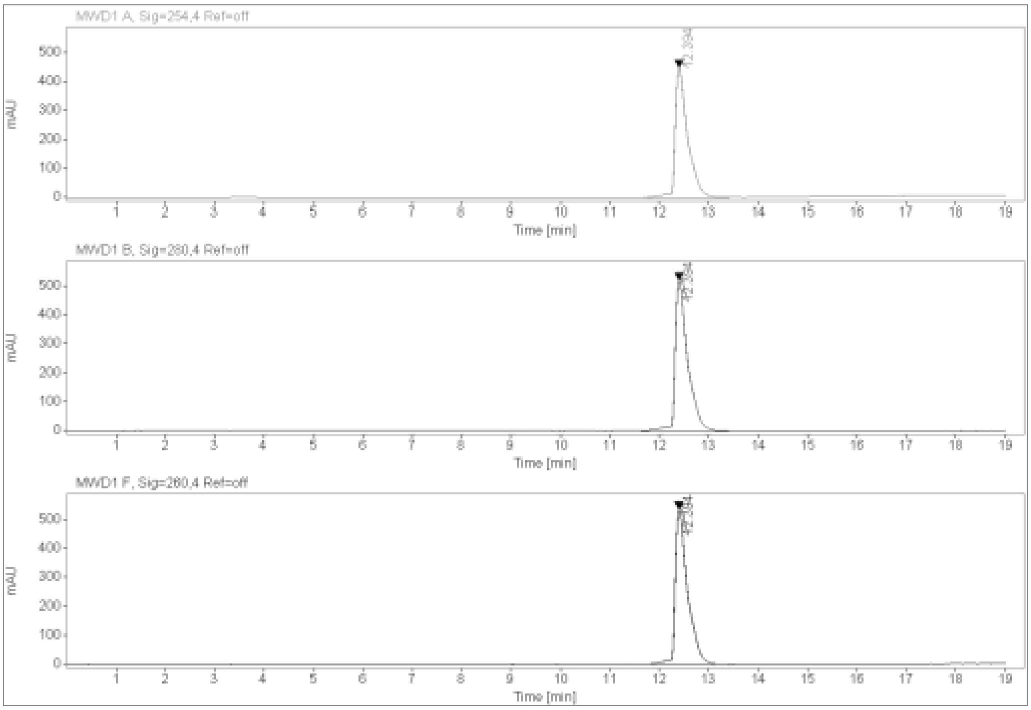
MWD1 A, Sig=254,4 Ref=off
12.394
mAU
Time [min]
MWD1 B, Sig=280,4 Ref=off
mAU
Time [min]
MWD1 F, Sig=260,4 Ref=off
mAU
Time [min]

**$^{1}$H-NMR, $^{13}$C-NMR, MALDI, HRMS and HPLC of (2S,4R)-1-((S)-26-(tert-butyl)-1-(2-((2-(3,4-dichlorophenyl)acetamido)methyl)-4-(6,7-dihydro-5H-pyrrolo[1,2-a]imidazol-2-yl)phenoxy)-24-oxo-3,6,9,12,15,18,21-heptaoxa-25-azaheptacosan-27-oyl)-4-hydroxy-N-(4-(4-methylthiazol-5-yl)benzyl)pyrrolidine-2-carboxamide 17g**

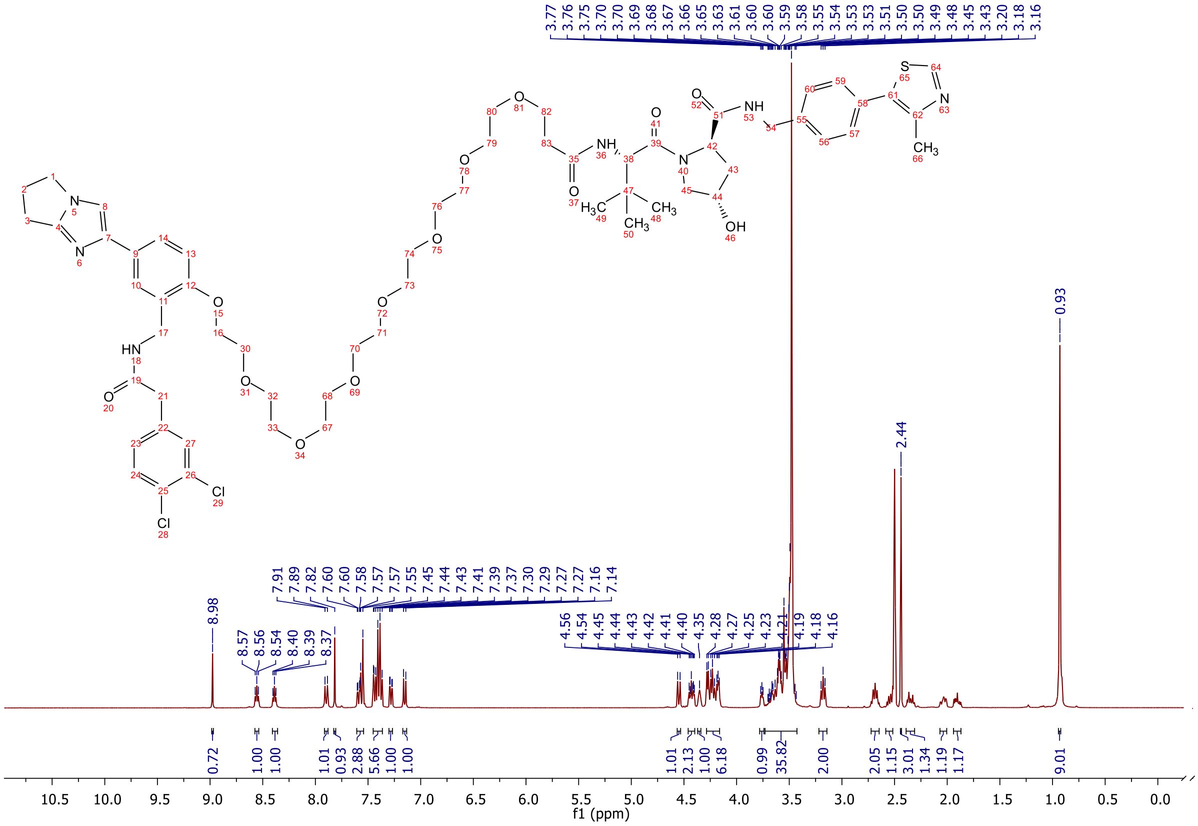

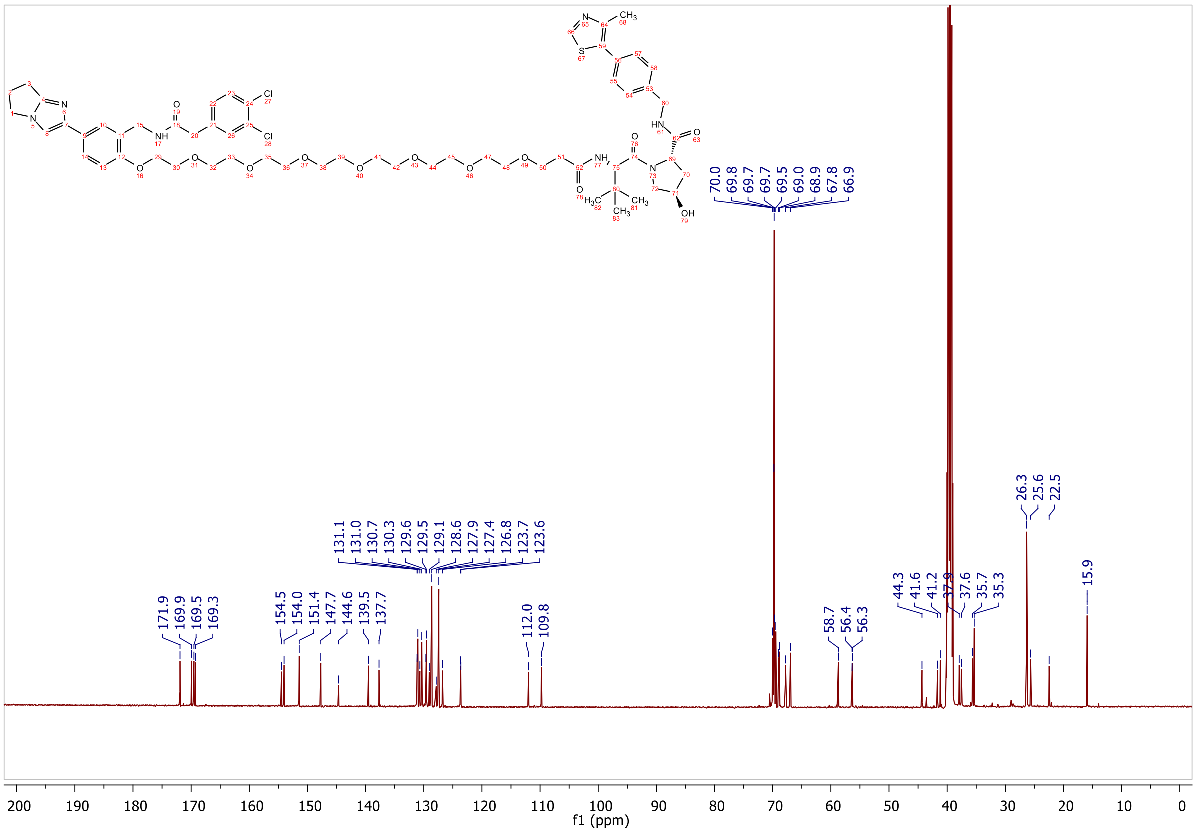

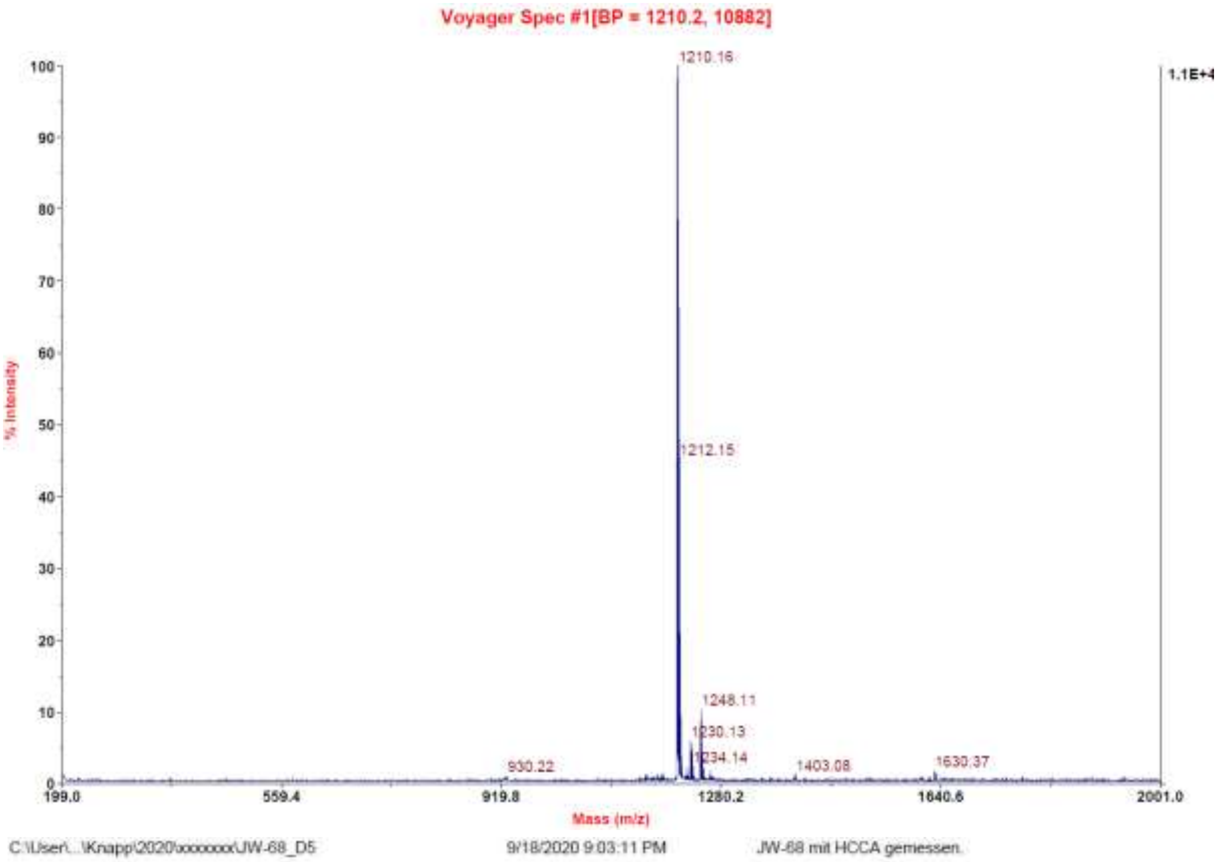
Voyager Spec #1[BP = 1210.2, 10882]
1210.16
1212.15
1248.11
1230.13
1234.14
930.22
1403.08
1630.37
1.1E+4
% Intensity
Mass (m/z)
C:\User\...\Knapp\2020\xxxxxxx\JW-68_D5
9/18/2020 9:03:11 PM
JW-68 mit HCCA gemessen.

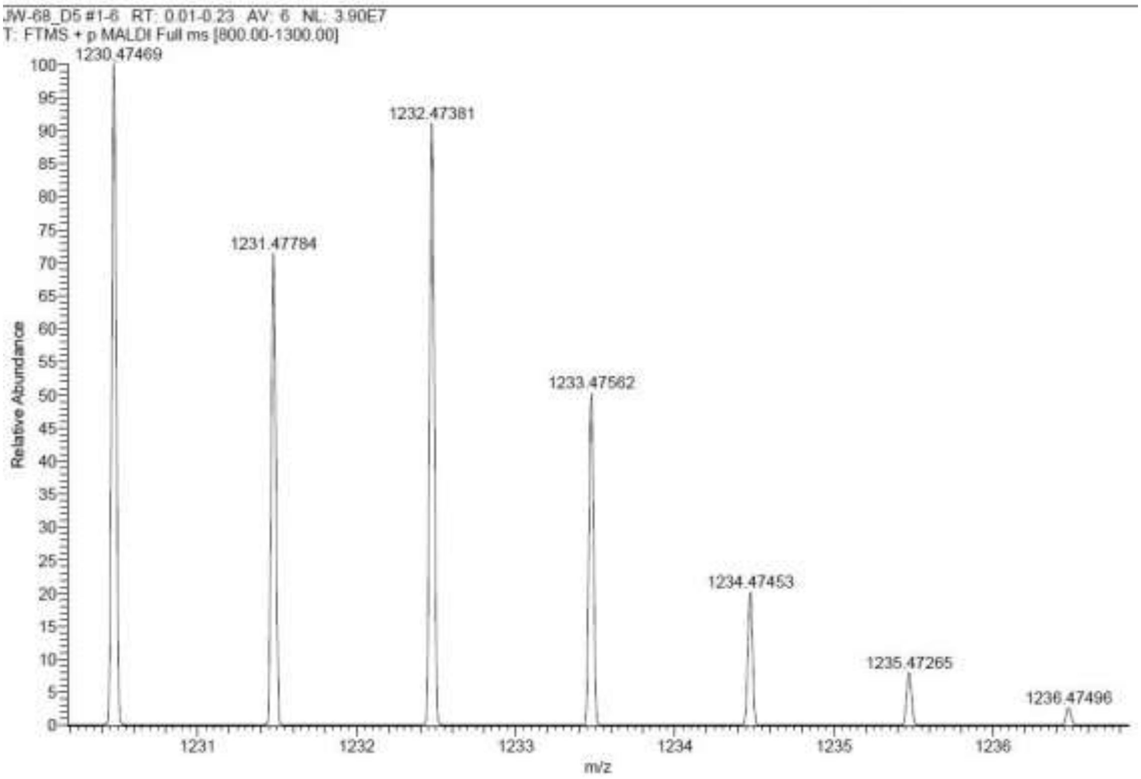
JW-68_D5 #1-6 RT: 0.01-0.23 AV: 6 NL: 3.90E7
T: FTMS + p MALDI Full ms [800.00-1300.00]
1230.47469
1231.47784
1232.47381
1233.47562
1234.47453
1235.47265
1236.47496
Relative Abundance
m/z

**Signal:** MWD1 A, Sig=254,4 Ref=off

| RT [min] | Type | Width [min] | Area | Height | Area% |
|---|---|---|---|---|---|
| 12.408 | VV | 0.2650 | 7540.5884 | 423.0270 | 100.0000 |
| | | Sum | 7540.5884 | | |

**Signal:** MWD1 B, Sig=280,4 Ref=off

| RT [min] | Type | Width [min] | Area | Height | Area% |
|---|---|---|---|---|---|
| 12.408 | VV | 0.2642 | 8727.5557 | 489.1573 | 100.0000 |
| | | Sum | 8727.5557 | | |

**Signal:** MWD1 F, Sig=260,4 Ref=off

| RT [min] | Type | Width [min] | Area | Height | Area% |
|---|---|---|---|---|---|
| 12.408 | VV | 0.2616 | 8891.1162 | 502.1700 | 100.0000 |
| | | Sum | 8891.1162 | | |

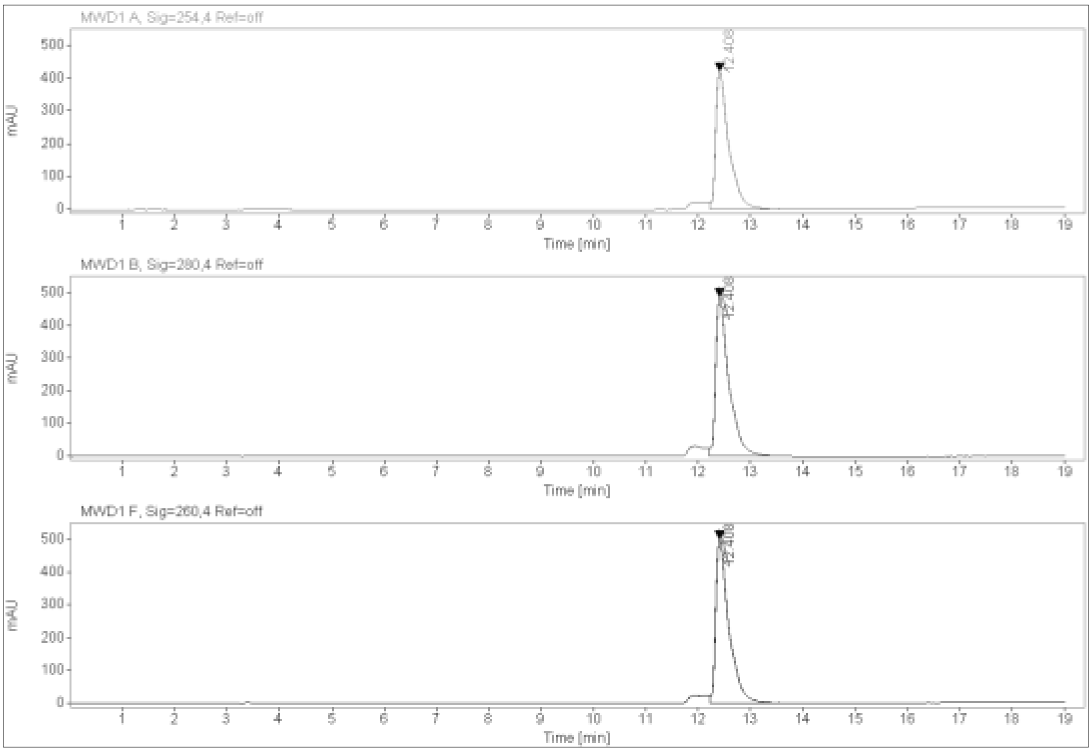

1.7 Appendix for negative controls 20 and 21

**MALDI, HRMS, HPLC and $^{1}$H-NMR of 6-hydroxy-N-(4'-((5-(((S)-1-((2S,4S)-4-hydroxy-2-((4-(4-methylthiazol-5-yl)benzyl)carbamoyl)pyrrolidin-1-yl)-3,3-dimethyl-1-oxobutan-2-yl)amino)-5-oxopentyl)carbamoyl)-4-(4-methylpiperazin-1-yl)-[1,1'-biphenyl]-3-yl)-4-(trifluoromethyl)nicotinamide 20**

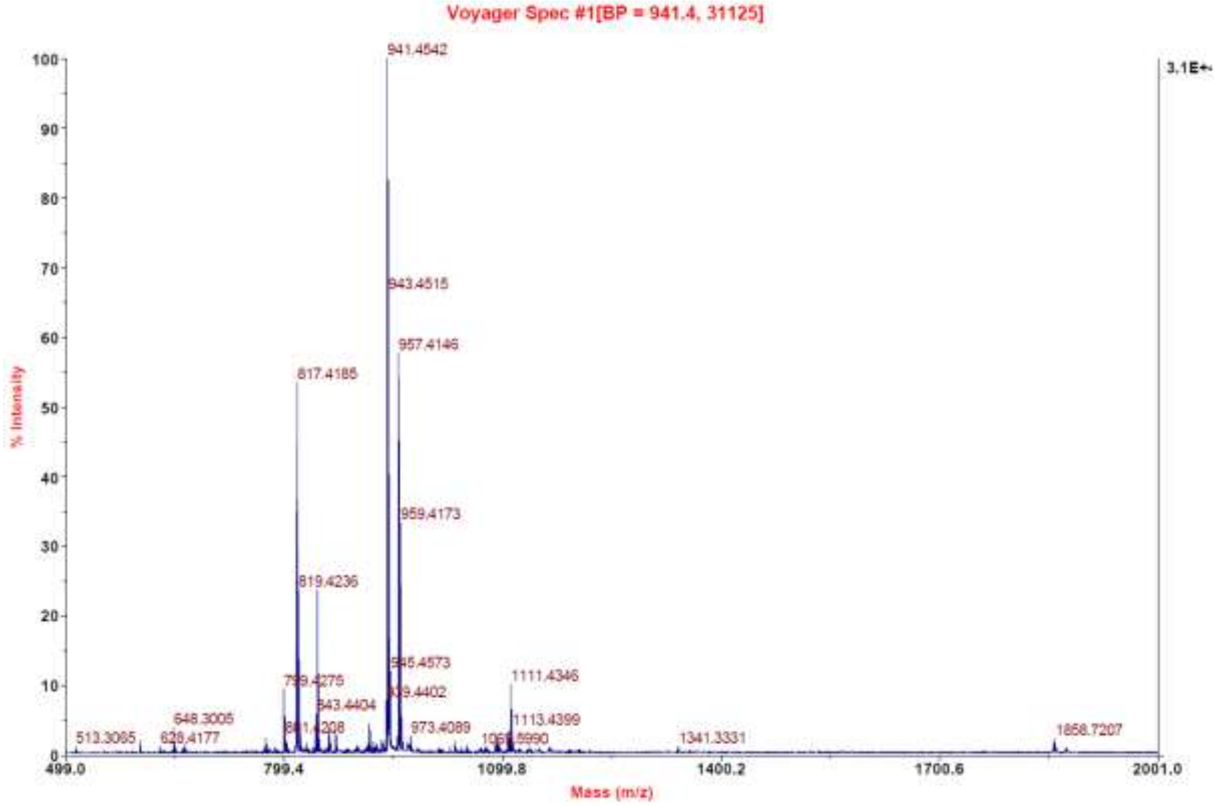

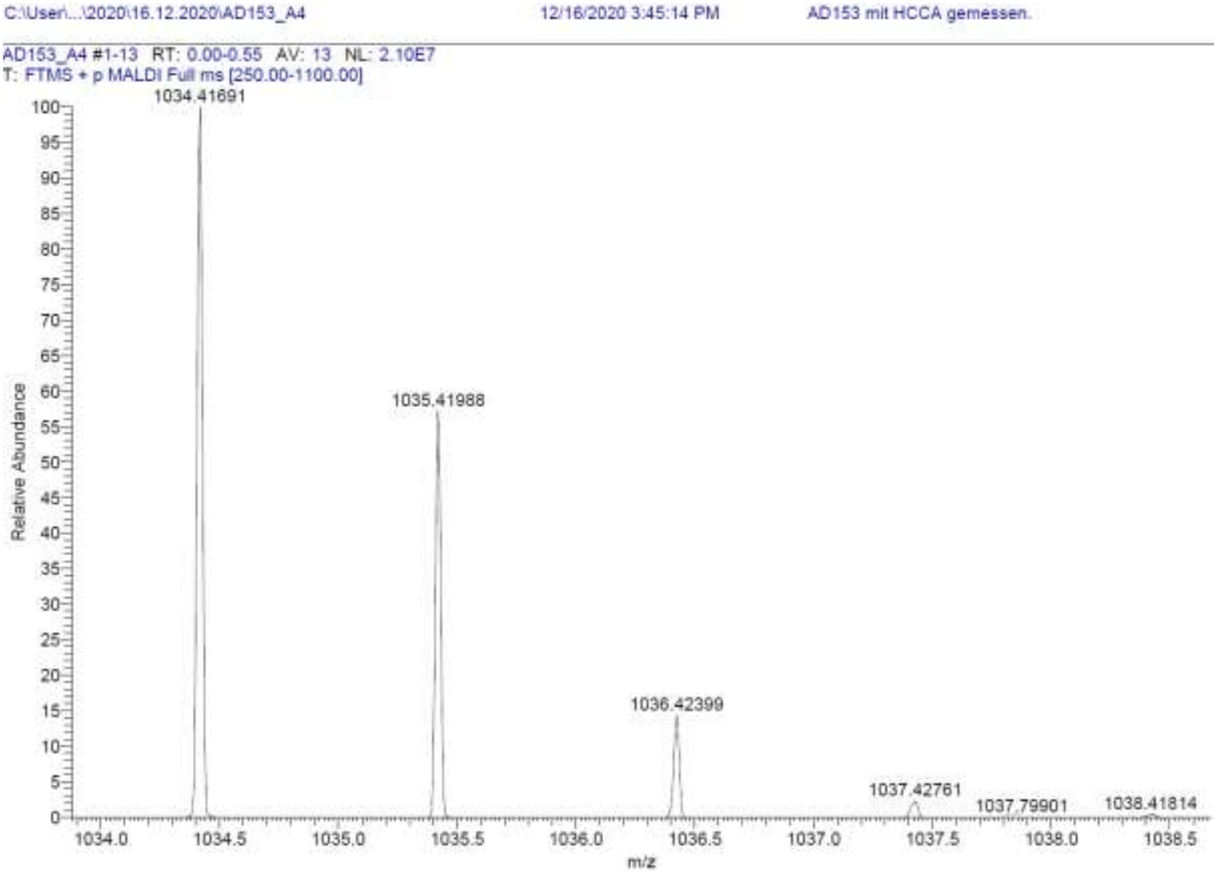

**Signal:** MSD1 TIC, MS File

| RT [min] | Type | Width [min] | Area | Height | Area% | Name |
|---|---|---|---|---|---|---|
| 11.580 | BB | 0.3359 | 26619110.0000 | 1236768.2500 | 58.3972 | |
| 18.536 | BBA | 0.3762 | 18963760.0000 | 840934.5625 | 41.6028 | |
| | | Sum | 45582870.0000 | | | |

**Signal:** MWD1 A, Sig=254,4 Ref=off

| RT [min] | Type | Width [min] | Area | Height | Area% | Name |
|---|---|---|---|---|---|---|
| 11.353 | VV | 0.2008 | 4513.2095 | 311.9199 | 100.0000 | |
| | | Sum | 4513.2095 | | | |

**Signal:** MWD1 B, Sig=280,4 Ref=off

| RT [min] | Type | Width [min] | Area | Height | Area% | Name |
|---|---|---|---|---|---|---|
| 11.353 | VV | 0.2029 | 5449.8301 | 372.0602 | 100.0000 | |
| | | Sum | 5449.8301 | | | |

**Signal:** MWD1 F, Sig=260,4 Ref=off

| RT [min] | Type | Width [min] | Area | Height | Area% | Name |
|---|---|---|---|---|---|---|
| 11.353 | VV | 0.2021 | 5101.9272 | 352.0205 | 100.0000 | |

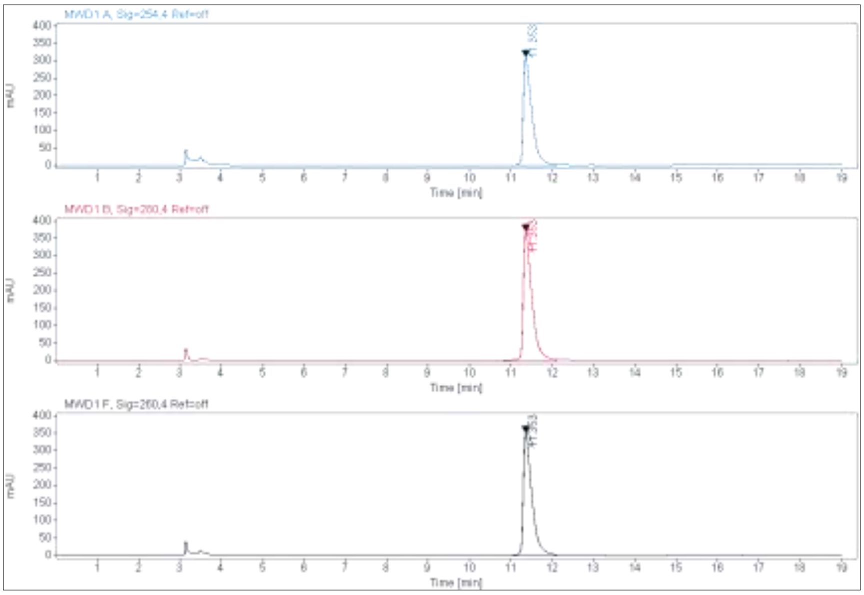
MWD1 A, Sig=254.4 Ref=off
MWD1 B, Sig=280.4 Ref=off
MWD1 F, Sig=260.4 Ref=off
mAU
Time [min]

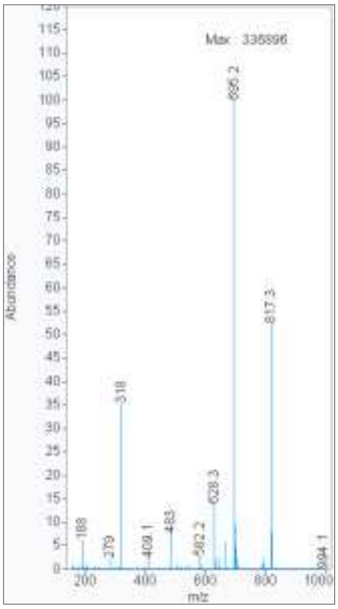
Max: 335896
Abundance
m/z
695.2
817.3
318
628.3
483
188
279
409.1
582.2
994.1

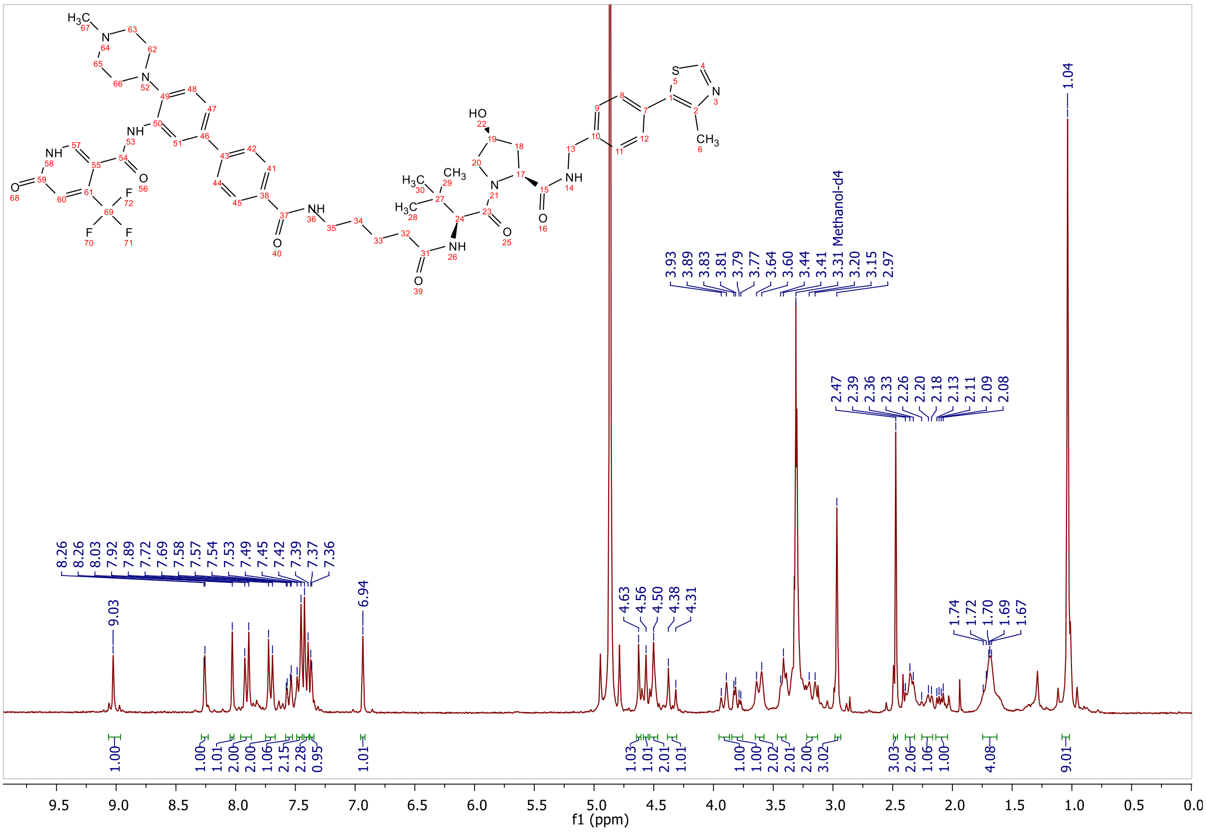
Methanol-d4
f1 (ppm)

**MALDI, HRMS, HPLC, $^{1}$H-NMR and $^{13}$C-NMR of (2S,4S)-1-((S)-2-(3-(2-(2-(2-((2-(3,4-dichlorophenyl)acetamido)methyl)-4-(6,7-dihydro-5H-pyrrolo[1,2-a]imidazol-2-yl)phenoxy)ethoxy)ethoxy)propanamido)-3,3-dimethylbutanoyl)-4-hydroxy-N-(4-(4-methylthiazol-5-yl)benzyl)pyrrolidine-2-carboxamide 21**

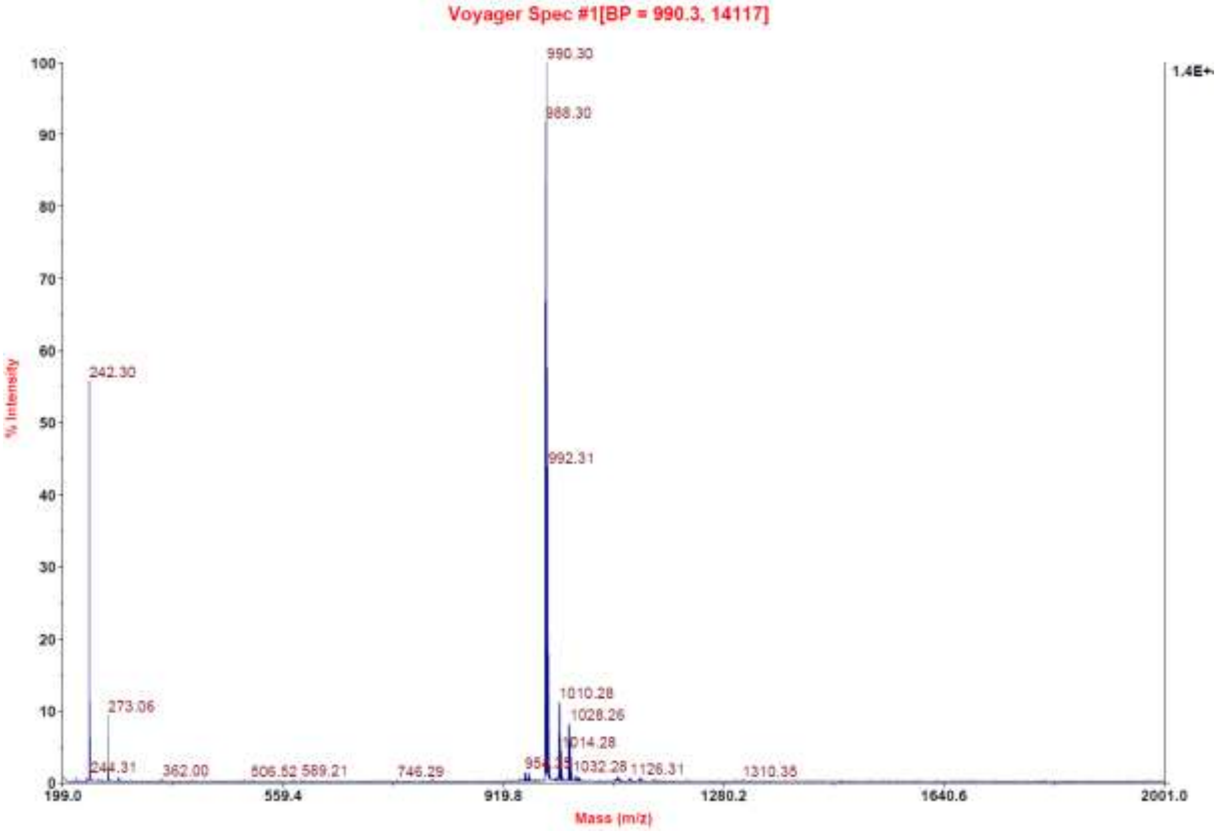

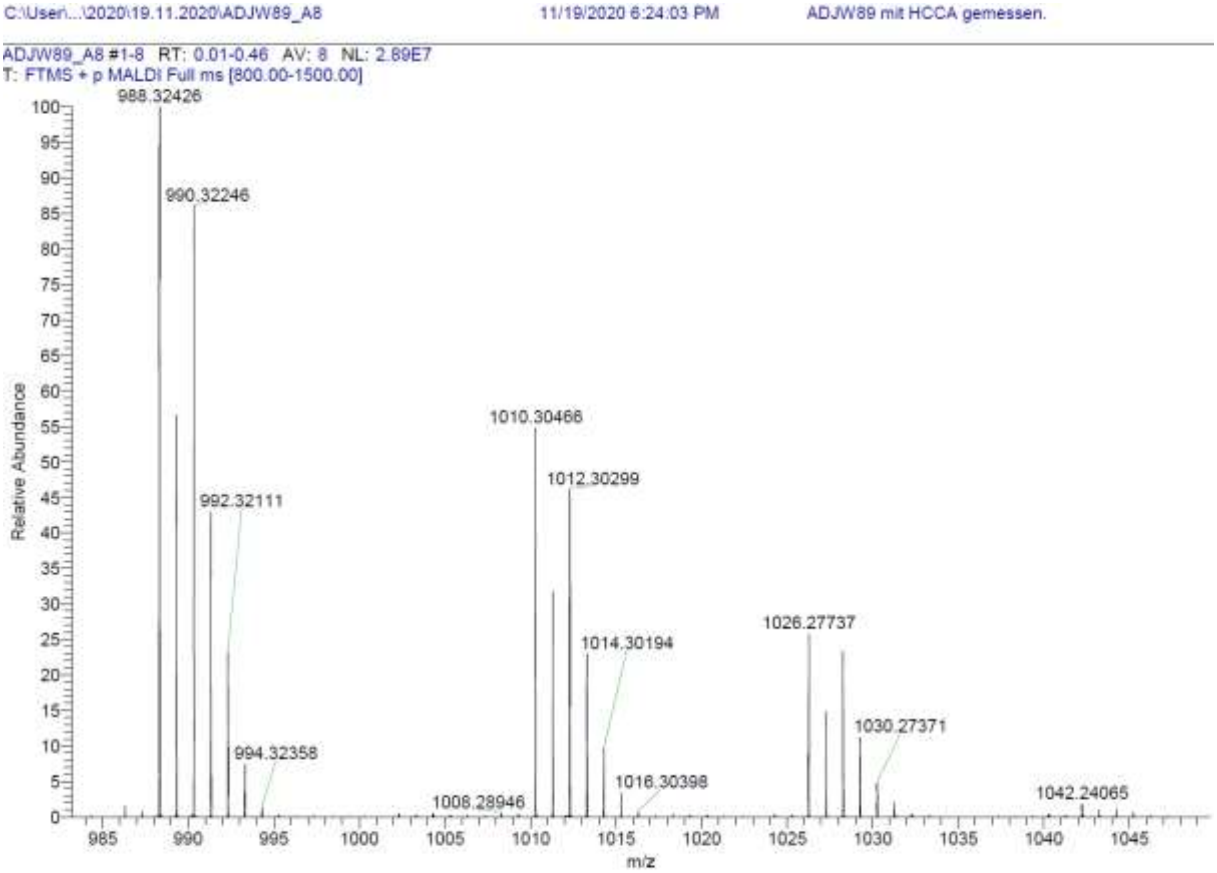

| RT [min] | Type | Width [min] | Area | Height | Area% | Name |
|---|---|---|---|---|---|---|
| 11.802 | VV | 0.2292 | 9360.8057 | 577.3460 | 100.0000 | |
| | | Sum | 9360.8057 | | | |

**Signal:** MWD1 B, Sig=280,4 Ref=off

| RT [min] | Type | Width [min] | Area | Height | Area% | Name |
|---|---|---|---|---|---|---|
| 11.802 | VV | 0.2278 | 10925.8682 | 671.5524 | 100.0000 | |
| | | Sum | 10925.8682 | | | |

**Signal:** MWD1 F, Sig=260,4 Ref=off

| RT [min] | Type | Width [min] | Area | Height | Area% | Name |
|---|---|---|---|---|---|---|
| 11.802 | VV | 0.2281 | 11161.8379 | 688.7037 | 100.0000 | |

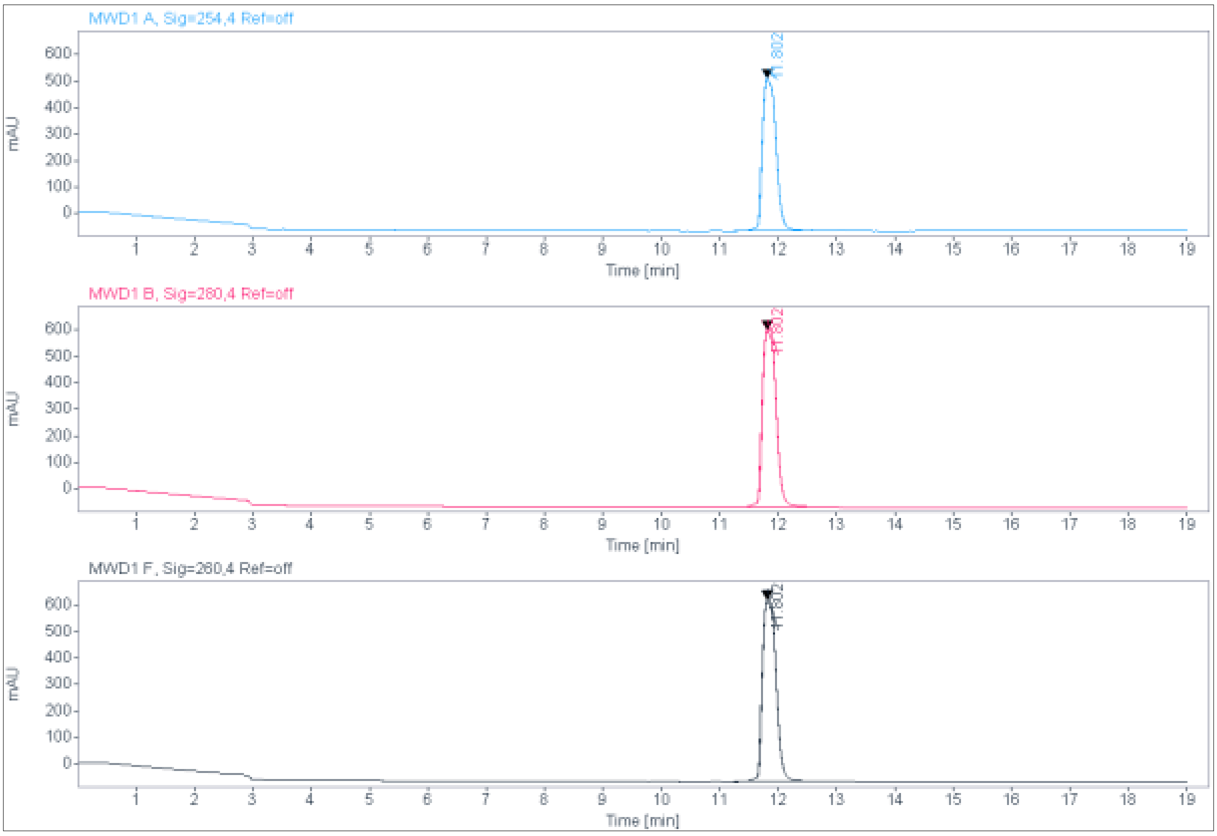

MWD1 A, Sig=254,4 Ref=off
11.802
MWD1 B, Sig=280,4 Ref=off
11.802
MWD1 F, Sig=280,4 Ref=off
11.802
mAU
Time [min]

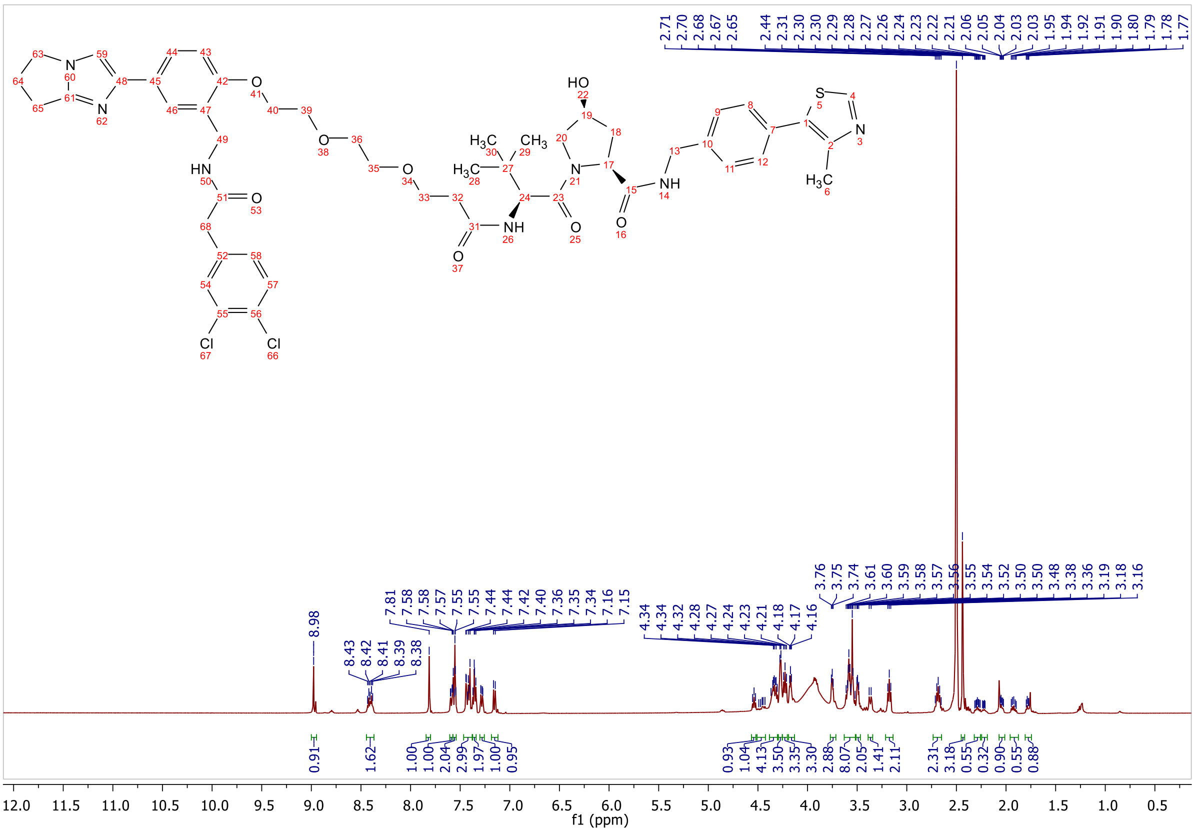

f1 (ppm)

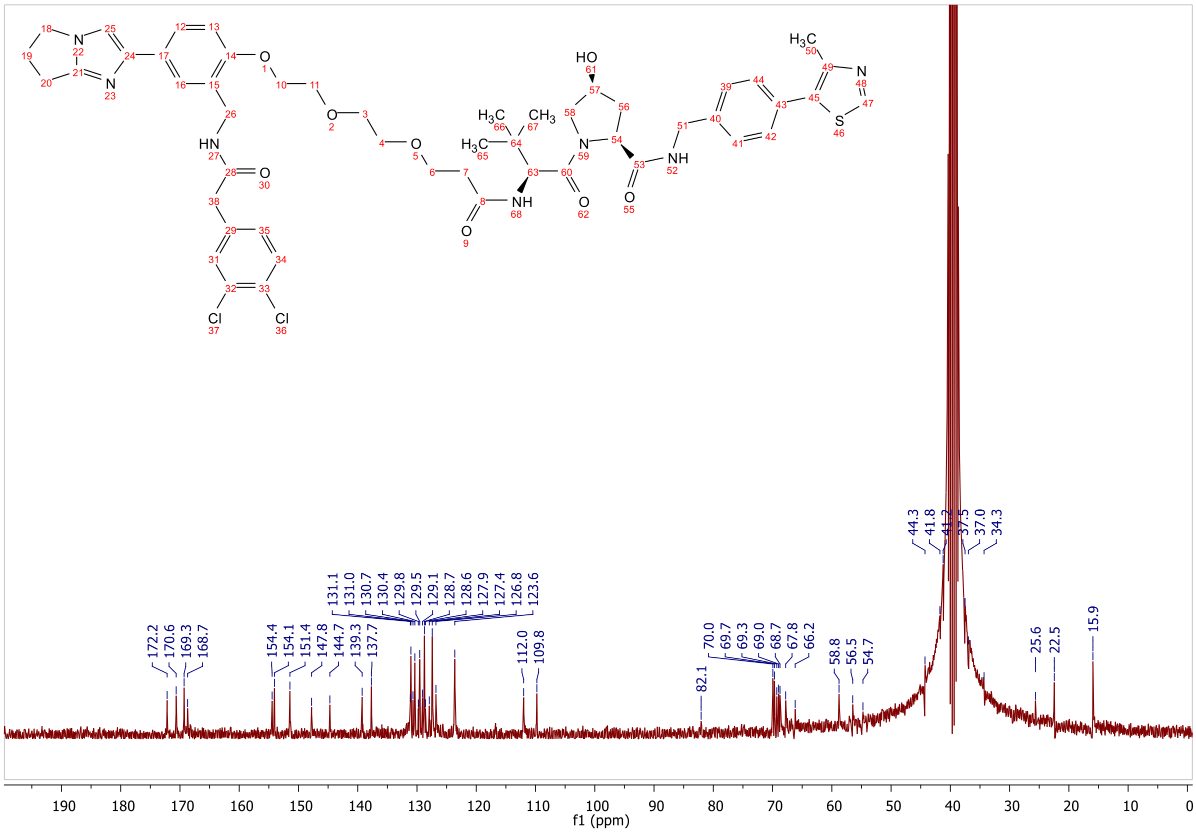
